# The clinical, genomic, and transcriptomic landscape of BRAF mutant cancers

**DOI:** 10.1101/2023.09.15.557847

**Authors:** Suzanne Kazandjian, Emmanuelle Rousselle, Matthew Dankner, David W. Cescon, Anna Spreafico, Kim Ma, Petr Kavan, Gerald Batist, April A. N. Rose

## Abstract

**Background:** BRAF mutations are classified into 4 molecularly distinct groups, and Class 1 (V600) mutant tumors are treated with targeted therapies. Effective treatment has not been established for Class 2/3 or BRAF Fusions. We investigated whether BRAF mutation class differed according to clinical, genomic, and transcriptomic variables in cancer patients.

**Methods:** Using the AACR GENIE (v.12) cancer database, the distribution of BRAF mutation class in adult cancer patients was analyzed according to sex, age, primary race, and tumor type. Genomic alteration data and transcriptomic analysis was performed using The Cancer Genome Atlas.

**Results:** BRAF mutations were identified in 9515 (6.2%) samples among 153,834, with melanoma (31%), CRC (20.7%), and NSCLC (13.9%) being the most frequent cancer types. Class 1 harbored co-mutations outside of the MAPK pathway (TERT, RFN43) vs Class 2/3 mutations (RAS, NF1). Across all tumour types, Class 2/3 were enriched for alterations in genes involved in UV response and WNT/β-catenin. Pathway analysis revealed enrichment of WNT/β-catenin and Hedgehog signaling in non-V600 mutated CRC. Males had a higher proportion of Class 3 mutations vs. females (17.4% vs 12.3% q = 0.003). Non-V600 mutations were generally more common in older patients (aged 60+) vs younger (38% vs 15% p<0.0001), except in CRC (15% vs 30% q = 0.0001). Black race was associated with non-V600 BRAF alterations (OR: 1.58; p<0.0001).

**Conclusions:** Class 2/3 BRAF are more present in Black, male patients with co-mutations outside of the MAPK pathway, likely requiring additional oncogenic input for tumorigenesis. Improving access to NGS and trial enrollment will help development of targeted therapies for non-V600 BRAF mutations.

**Statement of Translational Relevance:** BRAF mutations are classified in 4 categories based on molecular characteristics, but only Class 1 BRAF V600 have effective targeted treatment strategies. With increasing access to next-generation sequencing, oncologists are more frequently uncovering non-V600 BRAF mutations, where there remains a scarcity of effective therapies. Responsiveness to MAPK pathway inhibitors differs according to BRAF mutation class and primary tumor type. For this reason, we sought to determine whether key demographic, genomic, and transcriptomic differences existed between classes. This cross-sectional study analyzes the largest dataset of BRAF-mutated cancers to date. Our findings propose insights to optimize clinical trial design and patient selection in the pursuit of developing effective treatment strategies for patients whose tumors harbor non-V600 BRAF mutations. This study also offers insights into the potential of targeting alternative pathways in addition to the MAPK pathway as part of combinatorial treatment strategies.

## Introduction

BRAF is a serine/threonine kinase and a key signalling molecule within the mitogen-activated protein kinase (MAPK) pathway. The MAPK pathway serves to transmit extracellular mitogenic signals to the nucleus of receptive cells, promoting cellular survival and proliferation [1]. BRAF genomic alterations, such as mutations, fusions, and structural variants, are common in many cancer types and have proven to be potent oncogenic drivers [2].

Yao et. al., developed a classification system for BRAF mutations. Oncogenic BRAF alterations are categorized based on kinase activity, RAS-dependency, and dimerization requirements [3, 4]. The four distinct categories of BRAF alterations are: Class 1, Class 2, Class 3 and BRAF Fusions [1]. Class 1 BRAF mutants occur at the V600 codon. These V600 mutants signal as monomers, independent of upstream RAS activation and exhibit substantially increased kinase activity [5]. Class 2 BRAF mutations are non-V600 mutants that signal as dimers with intermediate to high kinase activity in a RAS-independent fashion [6]. Class 3 mutations are kinase impaired or kinase dead; however, Class 3 mutants have augmented ability to form dimers with wild-type BRAF or CRAF, exhibit enhanced binding to upstream RAS, and still result in MAPK pathway hyper-activation, albeit in a RAS-dependent manner [3]. BRAF Fusions function as RAS-independent obligate dimers that signal similarly to Class 2 BRAF mutations.

Class 1 BRAF mutant cancers are targetable with combinations of BRAF, MEK, and EGFR targeted therapies as part of the treatment armamentarium for advanced stage BRAF V600-mutant melanoma, non-small cell lung cancer (NSCLC), and colorectal cancer (CRC) [7–11]. Recently, the FDA granted tumor-type agnostic approval of dabrafenib and trametinib, BRAF and MEK inhibitors, respectively, for the treatment of any metastatic BRAF V600E mutant cancer [12]. Conversely, there have been no targeted therapy treatments that have been approved for cancers with Class 2 & 3 non-V600 BRAF mutations or BRAF fusions – which collectively comprise approximately 35% of all oncogenic BRAF alterations in adult solid tumors [1]. Retrospective clinical data suggests that some patients with Class 2 and 3 BRAF mutations benefit from MAPK pathway inhibitors, but the response rates are lower than in patients with Class 1 BRAF mutations [13].

There are multiple mechanisms of acquired resistance to BRAF +/-MEK inhibitors, including via oncogenic co-mutations in genes such as PTEN, NRAS, NF1, and AKT [14]. Tumors with Class 2 or 3 BRAF mutations are more likely to have co-occurring RAS mutations than tumors with Class 1 BRAF mutations, which may be associated with resistance to MAPK pathway inhibitors [3, 13, 15, 16]. Alternatively, some additional co-occurring mutations, in genes such as RNF43 in Class 1 BRAF mutant CRC are associated with increased responsiveness to BRAF targeted therapies [17]. This highlights the need to characterize the broader genomic profiles of BRAF mutant tumours as it could lead to the identification of potential mechanisms of therapeutic response and resistance that vary according to BRAF mutation classes.

Several clinical variables have also been associated with differences in response to MAPK pathway inhibitors in Class 1 BRAF mutant tumors. For example, the treatment response to BRAF and MEK inhibitors varies according to patient gender, in that women derive more benefit from targeted therapies than men [10]. Intriguingly, it has also been established that the incidence and genotype of RAS mutations differs according to primary tumour site, gender and race [18] [19]. However, the interactions between gender, age, race and BRAF mutation class across multiple tumor types are not well described.

Given the lack of established, effective therapies for Class 2 and 3 BRAF mutations, we sought to identify clinical, genomic, and transcriptomic variables that could underlie some of the differences in responsiveness to MAPK inhibitors between BRAF mutation classes. To do so, we interrogated the AACR Project GENIE and The Cancer Genome Atlas (TCGA) datasets. The AACR GENIE database is a comprehensive, cancer genomics database with clinical and next-generation sequencing (NGS) data available from more than 150,000 tumor samples [20] [21]. Our analysis provides insight into the molecular mechanisms underlying the tumorigenesis of non-V600 BRAF mutant tumors and identifies subgroups of patient most likely to benefit from novel therapeutic approaches.

## Materials & Methods

### Search Strategy

Using the American Association for Cancer Research Project Genomics Evidence Neoplasia Information Exchange GENIE (v12) cancer database, we analyzed the incidence and distribution of BRAF mutation class in cancer patients according to: sex, age, primary race, sample type, tumor type, and co-occurring mutations [21] [20]. Key inclusion criteria were: age above 20 years old and presence of a BRAF mutation. Key exclusion criteria were: patients with missing information on sex and age, unclassifiable BRAF mutations, and VUS mutations (variant of uncertain significance). Unclassifiable mutations were BRAF mutations with mechanisms of actions that did not follow the kinase activity, RAS-dependency, and dimerization requirements of the BRAF classification system. Variants of uncertain significance were BRAF mutations whose function and association to disease are unknown. The remaining samples were then classified into four categories according to previously published criteria. The incidence of specific BRAF alterations is included in **Supplementary Table S1** [1] [22].

### Statistical Analysis

The chi-square test was used with contingency tables to test whether differences exist in gender, race, and age across different classes of BRAF mutation. These values were then corrected using the Benjamini–Hochberg method to determine false discovery rate– corrected q values, which were considered significant when q<0.05. Odds ratio and 95% confidence intervals (CI) were calculated using a probit logistic regression model. The relationship between age according to BRAF mutation class and type of cancer was calculated using the two-way ANOVA test.

### Genomic Analysis

Genomic alteration data was obtained using the GENIEv12 database. We compared tumors with Class 1, 2 or 3 BRAF mutations. BRAF fusions were not included in the genomic analysis due to small sample size. To identify the top 30 most frequently co-occurring mutated genes that were differentially altered according to BRAF mutation class, we limited our analysis to: 1) genes that were altered in a minimum of 15 samples across classes, and 2) genes that were significantly (q<0.05) altered across the 3 classes. Additional inclusion criteria for NSCLC, CRC, and melanoma cancer-type specific analyses required genes to have been tested in at least 50% of the samples evaluated. Oncoprint figures were generated using the 4 BRAF Classes. We excluded samples for which sequencing data was not available for all queried genes. Pathway analyses on the list of genes that were significantly (q<0.05) differentially altered between Class 1 vs Class 2 or Class 1 vs Class 3 were performed using MSigDB Hallmark analysis and Enrichr [22, 23, 24].

Of the 153 834 samples in the AACR GENIE (v12) database, 9515 had identifiable BRAF mutations. 2022 of these samples harbored unclassifiable mutations or variants of unknown significance (VUS), and were excluded from further analyses. 1274 pediatric samples were removed, 1079 samples were excluded due to incomplete information on age and gender, and 20 duplicate samples were removed. A total of 5120 samples with BRAF Class 1, 2, 3 mutations and BRAF Fusions were included in the study (**Supplemental Figure S1**). 1264 samples were used to generate oncoprints across all BRAF mutant cancers, 221 samples in NSCLC, 462 samples in CRC, and 579 samples in melanoma.

### Transcriptomic Analysis

RNA sequencing data for BRAF mutant Melanoma (Skin Cutaneous Melanoma), NSCLC, and CRC cancers was obtained from the TCGA database [23]. Data pre-processing of these samples was performed in MATLAB verifying Spearman correlation, total counts, and IQR values of the samples. Outliers were removed and data was normalized in R using the DESeq2 normalization algorithm. The TCGA Melanoma RNAseq data included 195 BRAF mutant samples (167 Class 1, 11 Class 2, 14 Class 3, and 3 Fusions). The NSCLC RNAseq data is composed of 32 BRAF mutant samples (9 Class 1, 9 Class 2, and 14 Class 3). The CRC RNAseq data included 53 BRAF mutant samples (45 Class 1, 2 Class 2, 5 Class 3, and 1 Fusion). Heatmaps were generated in R using the ComplexHeatmap package and the following cutoffs: absolute LogFold ≥ 2 (BRAF Class 1 vs non-V600 BRAF), baseMean ≥ 50, and padj ≤ 0.01 (Supplemental Tables S19-S21). The normalized gene expression was then used for Gene Set Enrichment Analysis (GSEA) to compare the BRAF Class 1 gene expression signature to that of the BRAF non-V600 samples. The analysis was performed using the MSigDB Hallmark gene sets, 1000 permutations by phenotype, and using the Ratio of classes metric [24–26]. Fusion samples were excluded from the GSEA analyses.

## Results

### Characteristics of BRAF mutant Patient Cohort

We interrogated v12 of the AACR GENIE data set to identify all tumors with BRAF mutations or fusions. BRAF alterations were identified in 6.2% of samples **(Supplemental Table S1).** Among the classifiable mutations, 3358 (65.6%) were Class 1 mutants, 782 (15.3%) were Class 2 mutants, 759 (14.9%) were Class 3 mutations, and 221 (4.3%) were BRAF Fusions. The median age for all patients with BRAF mutant cancers was 62 years old. Median age was 62 years old, 65 years old, 64 years old and 59 years old for Class 1, 2, 3 and Fusion respectively (**Table 1**). The most frequent BRAF altered cancer types were melanoma (n=1591), CRC (n=1061) and NSCLC (n=714). This cohort of patients had sequenced samples from both primary (47.2%) and metastatic (41.2%) tumors (**Table 1**).

**Table 1:**
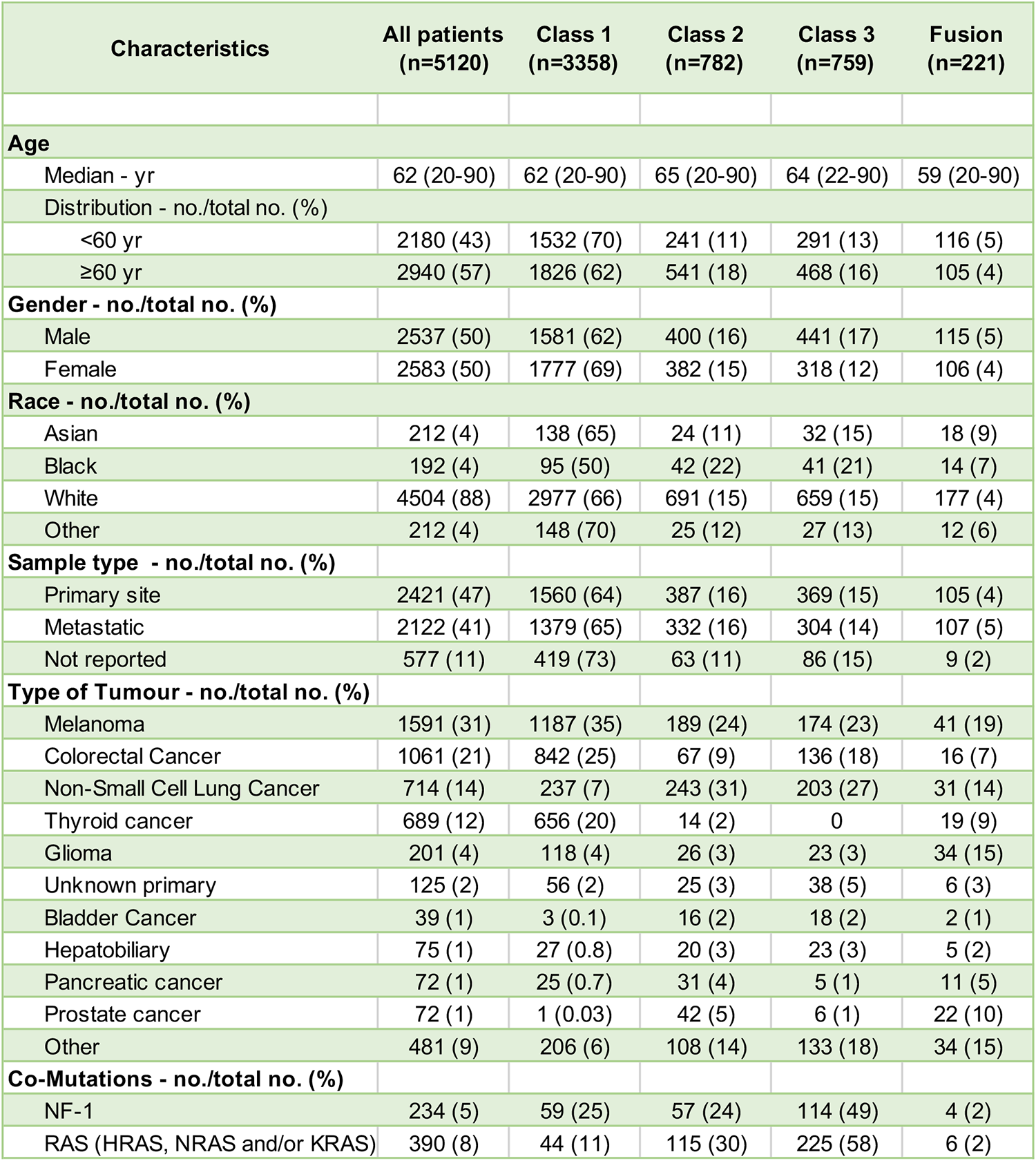
Baseline Demographic and Clinical Characteristics of Patients.

### Relationship between BRAF mutation class and co-occurring genomic alterations

We analyzed the incidence of co-occurring genomic alterations in BRAF altered tumors. Across all BRAF mutant cancers, 228 genes were significantly differentially altered in BRAF Class 1/2/3 mutant tumors (**Supplemental Table S2**). The most frequently altered genes that were differentially altered according to BRAF mutation Class are indicated in **Figure 1A and B (Supplemental Tables S3-S6)**. We validated previously published observations indicating that KRAS, NRAS and NF1 mutations were more common in tumors with Class 2 and 3 mutations compared to tumors with Class 1 mutations [6, 10, 27]. However, we also report several novel gene alterations that differ according to BRAF Class. These include TERT, TP53, APC, and PIK3CA. Most of these gene alterations were more common in Class 2 and 3 BRAF mutant tumors, but some gene alterations (TERT, RNF43) were more common in Class 1 BRAF mutant tumors. BRAF Fusions were associated with the lowest co-mutation burden overall but had a high prevalence of co-occurring EGFR alterations. Some of these associations may reflect higher incidence of these co-occurring alterations within different cancer types, as the prevalence of BRAF mutation Class varies by cancer type (**Table 1**). Therefore, we also examined the relationship between BRAF mutation Class and co-occurring genomic alterations within specific cancer types. In melanoma, NF1, KMT2A, ARID2, ATM, APC, ARID1A, and NOTCH1 mutations commonly co-occur in BRAF Class 2 and 3 mutant cancers (**Supplemental Figure S2, Supplemental Tables S7-S10**). Of note, most co-occurring mutations in melanoma BRAF Class 2 and 3 are of unknown significance, whereas those in Class 1 are amplifications and putative drivers such as NOTCH2 and MET amplifications. Co-occurring mutations were uncommon in melanomas with BRAF Fusions; indeed, the genomic landscape of melanomas with BRAF fusions was similar to that of BRAF Class 1 melanomas. Among BRAF mutant CRC, Class 1 samples had a higher co-mutation burden compared to other BRAF classes (**Supplemental Figure S3, Supplemental Tables S11-S14**). RNF43 mutations occurred in 60% of BRAF Class 1 CRC but less than 20% of BRAF Class 2 and 3 CRC samples harbored these truncating mutations. Almost 100% of CRCs with BRAF Fusions had RNF43 mutations, and colorectal tumors with BRAF Fusions again had a genomic landscape that was similar to BRAF Class 1 mutant CRCs. Conversely, RAS isomers (KRAS, and NRAS) were more likely to be co-mutated in BRAF Class 2 and 3 CRCs. BRAF Class 1 NSCLC predominantly have TP53 and SETD2 co-mutations (**Supplemental Figure S4, Supplemental Tables S15-18**). Class 2 and 3 BRAF mutant NSCLC were more likely to have co-occurring mutations in STK11, KEAP1 and KRAS. Again, in NSCLC, the mutational landscape of BRAF Fusions was most similar to tumors with Class 1 BRAF mutations.

**Figure 1:**
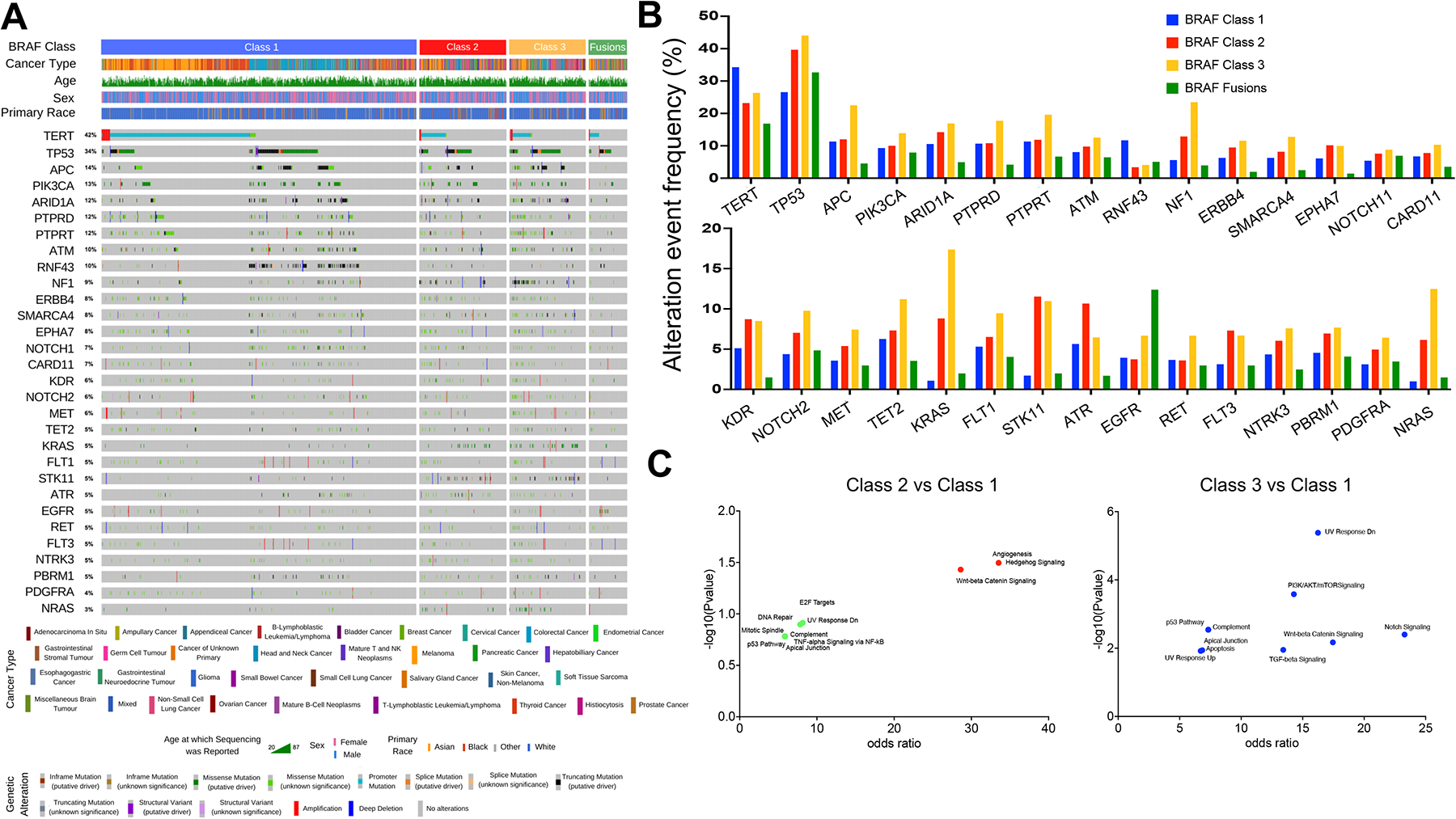
The genomic landscape of BRAF mutant tumors. **A)** Oncoprint highlighting the top 30 most frequent genes that are differentially altered between tumors with Class 1/2/3 BRAF mutations. **B)** Histogram highlighting the incidence of gene alterations within each BRAF class. **C)** The filtered list of genes that were significantly differentially altered according to BRAF Class 1/2 and 1/3 status across all cancers (n=18, and n=59 respectively, Q<0.05) was subjected to pathway analysis using the using MSigDB Hallmark algorithm. Pathways that were over-represented in this list of genes are indicated in blue, red, and green (P<0.05 & Q<0.05, P<0.05, Q<0.2, and P<0.2 & Q<0.2, respectively).

Next, we performed pathway enrichment analysis to identify the pathways that are over-represented amongst significantly differentially altered genes between BRAF Class 1 V600 mutant tumors vs. tumors with Class 2 or 3 non-V600 BRAF mutations. Across all cancers, the comparison of differentially altered genes in Class 2 vs. Class 1 did not yield any significantly enriched pathways. However, Notch Signaling, UV Response, TGF-Beta Signaling, Wnt-beta Catenin Signaling, and PI3K/AKT/mTOR Signaling pathways were differentially altered between Class 3 and Class 1 BRAF mutant tumors (**Figure 1C**). Within melanoma, Apical Junction, Wnt-beta Catenin Signaling, and UV Response pathways were significantly altered between either Class 2 or Class 3 BRAF mutant tumors. Notably in the Class 3 vs. Class 1 comparison, there was an enrichment for several additional pathways, such as E2F targets, G2M Checkpoint, KRAS Signaling, and Apoptosis (**Figure S2C**). There was overlap in the pathways enriched between Class 3 vs. Class 1 BRAF mutant CRCs and Class 3 vs. Class 1 BRAF mutant melanomas (**Supplemental Figure 3C**). Pathway analysis of altered genes in NSCLC did not yield any significant findings (**Supplemental Figure 4C**). Overall, across multiple tumor types, Class 2 BRAF mutant tumors were enriched for alterations in genes involved in ultraviolet (UV) response and Wnt-beta Catenin signaling. Class 3 BRAF mutant tumors were enriched for alterations in genes involved in UV Response and Wnt-beta Catenin Signaling, Notch Signaling, E2F targets, G2M Checkpoint, and Hedgehog Signaling.

### Relationship between BRAF mutation Class and gene expression

To validate the pathway enrichment findings from our genomic analysis, we performed Gene Set Enrichment Analysis on RNA sequencing data from BRAF mutant melanoma, NSCLC, and CRCs. The top 10 gene sets enriched amongst the genes differentially expressed between non-V600 vs. V600 BRAF mutant cancers for each cancer type are indicated in **Figure 2**. In melanoma, PI3K/AKT Signaling, Estrogen Response (Early&Late), Mitotic Spindle, G2M Checkpoint, and UV Response Dn (genes downregulated in response to UV radiation) pathways were at the level of Genomic alteration (**Supplemental Figure 2**) and RNA expression (**Figure 2A & B**). Pathway analysis of differentially expressed genes between non-V600 and V600 BRAF mutant CRCs revealed once again the enrichment of Wnt-Beta Catenin and Hedgehog Signaling pathways in the non-V600 BRAF mutants. Conversely, the only pathway enriched at the genomic and transcriptomic levels in NSCLC analyses was the enrichment of the Apical Junction gene set in the non-V600 BRAF mutants.

**Figure 2:**
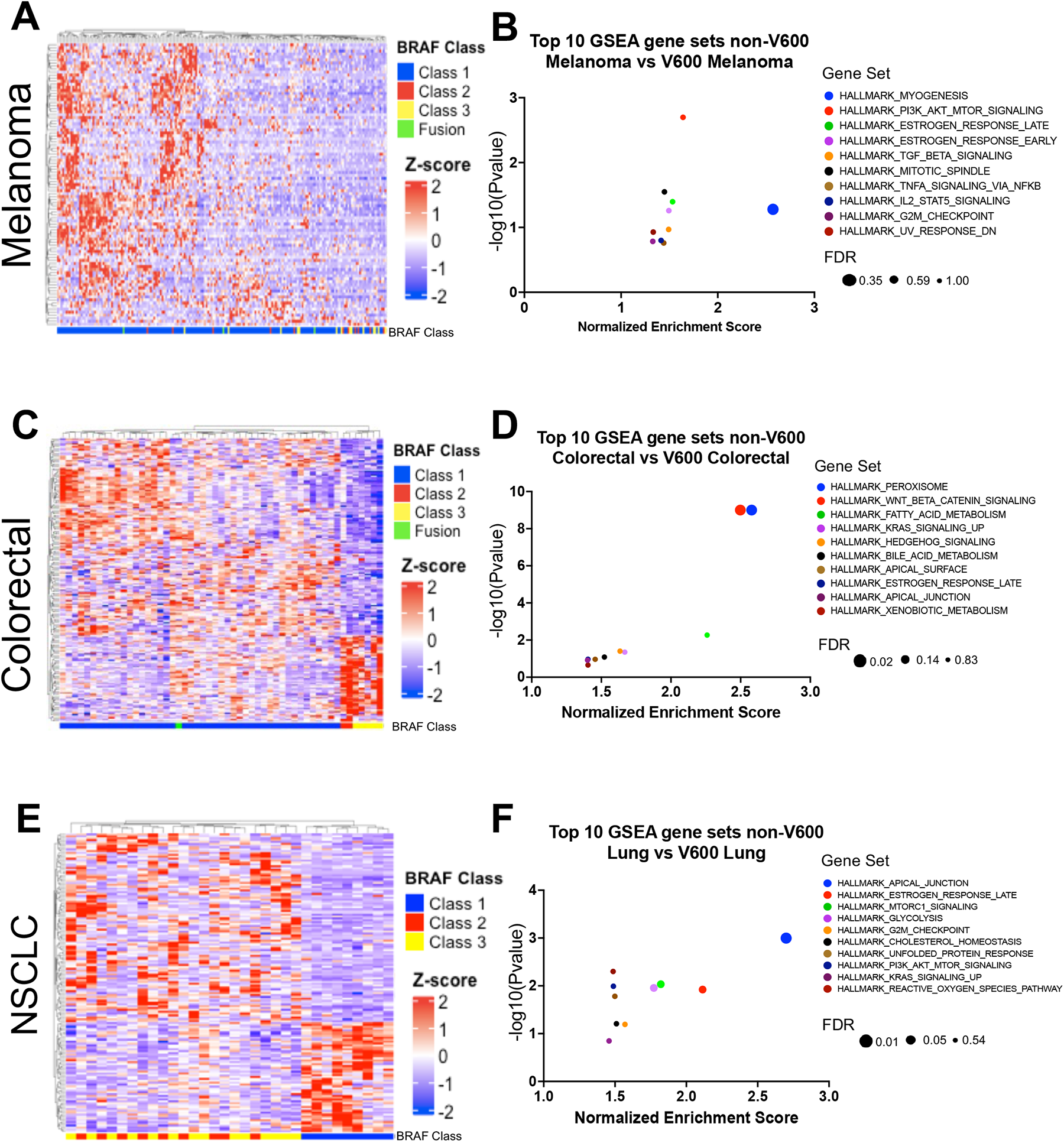
Distribution of BRAF mutation Class according to sex, age and race across cancer type. The frequency of each BRAF Class is shown in subgroups defined by sex (A), age (B), and by primary race (C) among all patients, melanoma, CRC, NSCLC and all other cancers (all patients excluding melanoma, CRC & NSCLC). Values shown within each category represent the proportion of patients expressing each BRAF Class within cancer types according to sex (A), age (B), and primary race (C). P-value was calculated through the chi-square test for each contingency table, and was then corrected using the Benjamini–Hochberg method to determine false discovery rate–corrected q value, which was considered significant when q was less than 0.05.

### Relationship between patients’ sex and BRAF mutation Class

The incidence of specific oncogenic genomic alterations varies significantly according to patient sex [19, 28]. Therefore, we inquired whether BRAF mutation Class varies with patient sex. In the entire data set, 50.4% patients were female and 49.6% were male (**Table 1**). The distribution of BRAF mutation Class differed significantly according to patient sex. Across all cancer types, males had a higher proportion of Class 3 mutations vs. females (17.4% vs 12.3% q = 0.003) (**Figure 3A**). This trend for an increased proportion of Class 3 BRAF mutations in males was also observed within specific cancer types, including: melanoma, CRCs, and NSCLC (**Figure 3A**). Conversely, females had a higher proportion of Class 1 BRAF mutations compared to males across all cancer types (69% vs. 62%) and within specific cancer types, including melanoma, CRC, and NSCLC. The relationship between patient BRAF mutation class and patient sex was independent of other key clinical and genomic variables in multivariable analysis (OR 1.55 p<0.0001) (**Table 2**).

**Table 2:** Multivariable Analyses of factors associated to non-V600 BRAF mutations (Class2/3/Fusion) vs. V600 BRAF mutations (Class 1) OR = Odd’s ratio; CI = Confidence Interval.

|  | Univariable |  |  | Multivariable |  |  |
| --- | --- | --- | --- | --- | --- | --- |
|  | OR | 95% CI | P value | OR | 95% CI | P value |
| Gender |  |  |  |  |  |  |
| Male vs Female | 1.33 | 1.19 to 1.50 | <0.0001 | 1.55 | 1.35 to 1.76 | <0.0001 |
| Age |  |  |  |  |  |  |
| ≥60 vs <60 | 1.44 | 1.28 to 1.62 | <0.0001 | 1.28 | 1.12 to 1.46 | <0.0001 |
| Primary Race |  |  |  |  |  |  |
| Black vs other | 2.00 | 1.50 to 2.67 | <0.0001 | 1.58 | 1.13 to 2.20 | 0.007 |
| Primary tumour type |  |  |  |  |  |  |
| Melanoma | 0.54 | 0.48 to 0.62 | <0.0001 | 0.50 | 0.43 to 0.59 | <0.0001 |
| Colorectal | 0.42 | 0.36 to 0.50 | <0.0001 | 0.38 | 0.31 to 0.46 | <0.0001 |
| NSCLC | 4.89 | 4.13 to 5.79 | <0.0001 | 3.08 | 2.53 to 3.75 | <0.0001 |
| Genomic co-mutations |  |  |  |  |  |  |
| RAS mt vs wild-type | 18.4 | 13.37 to 25.34 | <0.0001 | 19.18 | 13.80 to 26.64 | <0.0001 |

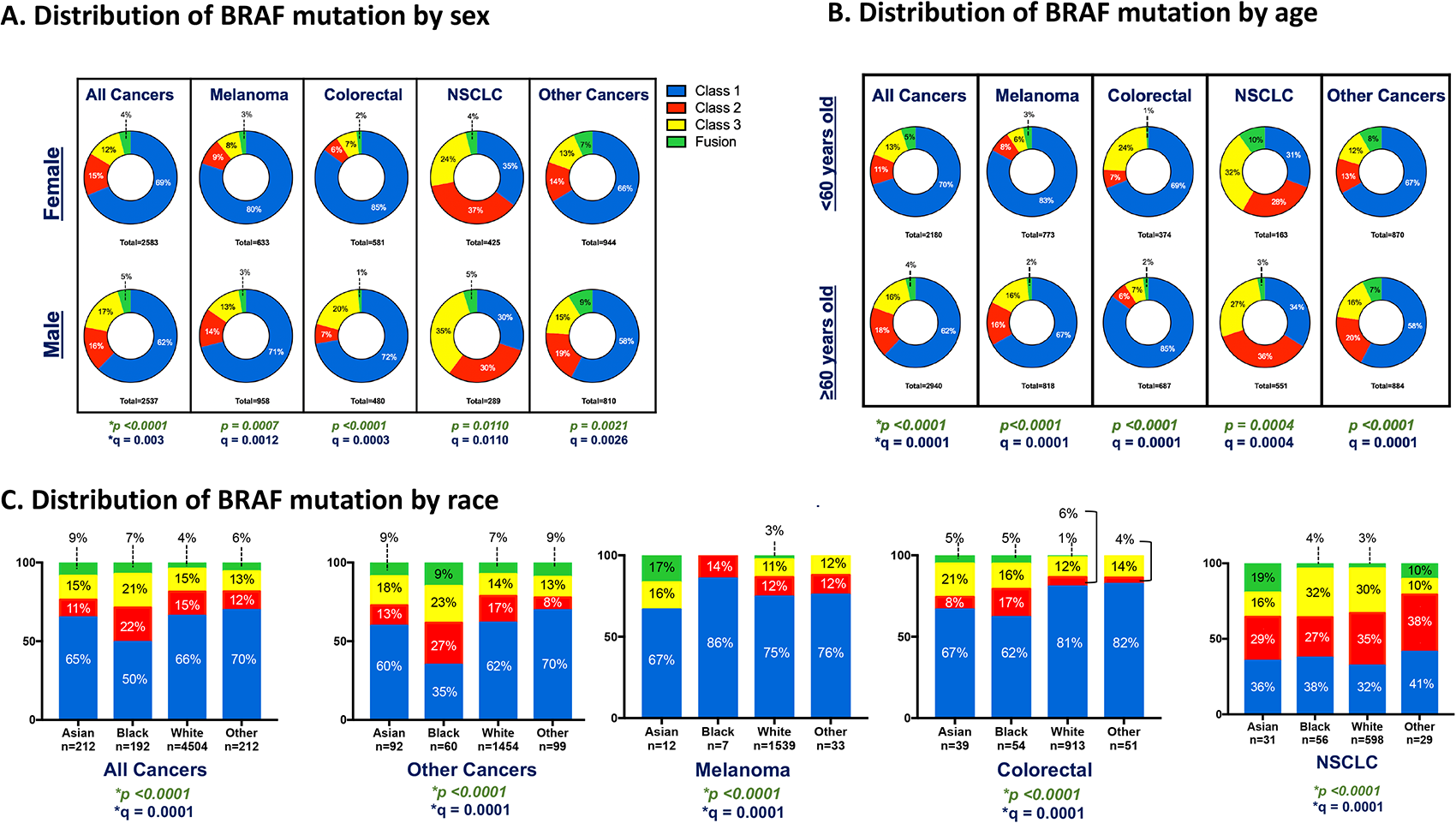

### Relationship between age and BRAF mutation Class

The age distribution of patients also varied according to BRAF Class (**Figure 3B**). The majority (57.5%) of patients included in this analysis were 60 years or older (**Table 1**). Across all cancer types, the relative proportion of non-V600 BRAF mutations (Class 2, 3, and Fusions) was higher in older patients, (age 60+) compared to younger patients (age < 60) (p<0.0001) (**Figure 3B, Supplemental Figure 5**). This relationship between BRAF class and age was independent of other clinical or genomic variables – such as RAS mutation status – in multivariable analyses (OR: 1.28; 95% CI 1.12 to 1.46; p<0.0001) (**Table 2**). This relationship with age was observed in melanoma and other cancer types. However, in NSCLC and in CRC, non-V600 BRAF mutations were more common in younger patients (**Figure 3B, Supplemental Figure 5**). In NSCLC, non-V600 mutations made up 69.3% of all BRAF mutations in patients aged 60 years old and less, compared to 66% of all BRAF mutations in older patients (q = 0.0004 – (**Figure 3B**)). In CRC, this incidence was even more pronounced with non-V600 BRAF mutations representing 31% of all BRAF mutant cancers for patients younger than 60, compared to only 15% of all BRAF mutations in CRC patients who were older than 60 (q = 0.0001 – **Figure 3B**). More specifically, in younger CRC patients 23.5% of all BRAF mutations were Class 3 mutations, whereas only 7% of BRAF mutations in older patients were Class 3 mutations (q = 0.0001 – **Figure 3B**). Early onset CRC is increasing in incidence and is defined as colorectal cancer diagnosed before age 50 [29]. Therefore, we asked if non-V600 BRAF mutations were over-represented in patients younger than 50. Indeed, CRC with non-V600 BRAF mutations were more frequent in patients who were younger than 50 (34.1% of all BRAF mutations) compared to only 17.9% in patients older than 50 years old (p<0.0001) **(Supplemental Figure 5).**

### Relationship between primary race and BRAF mutation class

The distribution of BRAF mutation class also varied according to the patients’ primary race (**Figure 2C**). Across all cancers, and specifically within CRCs and “other” cancer types, Black patients had a lower proportion of Class 1 BRAF mutations compared to patients of other races. Across all cancers, Class 1, 2, 3 and Fusion BRAF mutations made up 50%, 22%, 21% and 15% in Black patients, respectively compared to 65%, 11%, 15%, 9% in Asian patients, and 66%, 15%, 15%, and 4% in White patients (q = 0.0001) (**Figure 2C**). For patients with cancer types other than melanoma, CRC, or NSCLC, Class 1, 2, 3 and Fusion BRAF mutants made up 35%, 27%, 23% and 15% of BRAF mutations in Black patients compared to 62%, 17%, 14% and 7%, respectively in white patients (q = 0.0001) (**Figure 2C**). A similar trend was observed in CRC. In NSCLC, Asian patients were less likely to have Class 3 BRAF mutations (16%) and more likely to have BRAF fusions (19%) compared to Black patients (32% Class 3 and 4% Fusions) or white patients (30% Class 3 and 3% Fusions). In multivariable analysis, that adjusted for primary tumor type, Black race was independently associated with increased odds of having a non-V600 BRAF alteration (OR: 1.58; 95% CI 1.13 to 2.20; p<0.0001) (**Table 2**).

## Discussion

As the clinical implementation of next-generation sequencing (NGS) is becoming increasingly accessible for many tumor types, so too is the identification of non-V600 BRAF mutations in patients’ tumors in a clinical setting. While treatment strategies for Class 1 BRAF mutant tumours with MAPK targeted therapies are well described, effective targeted therapies have yet to be established for non-V600 BRAF mutant cancers [7–10]. A wide array of clinical and genomic factors – beyond the molecular characteristics of the BRAF mutation itself – affect targeted therapy efficacy in BRAF mutant tumors [30]. This cross-sectional analysis performed on a large sample of BRAF mutant cancers in adult patients reveals key demographics and genomic differences among patients with these mutations. Our findings highlight a number of potential explanations for the lackluster efficacy of standard MAPK inhibitor approaches in tumors with oncogenic, non-V600 BRAF mutations compared to those with Class 1 V600 BRAF mutations. Our results also identify additional potential therapeutic targets in non-V600 BRAF mutant tumors and identify patient populations most in need of novel therapeutic approaches.

The genomic analyses presented herein highlight the fact that Class 1 BRAF mutant tumors tend to harbor co-mutations outside of the MAPK pathway such as TERT or RNF43. Whereas Class 2, and to an even greater extent, Class 3 BRAF mutations co-exist with other members of the MAPK signaling cascade. Most prominently, these genes include RAS (KRAS, HRAS, NRAS), NF1 and genes encoding receptor tyrosine kinases, such as EGFR, ERBB2, MET and RET. This finding aligns well with the fact that non-V600 Class 2 & 3 BRAF mutations are less effective at promoting downstream MAPK pathway activation compared to Class 1 BRAF mutations [4]. As such, additional oncogenic inputs are required to cooperate with Class 2 & 3 BRAF mutations in order to elicit sufficient MAPK signalling output to promote tumorigenesis [3, 4].

We observe significant differences in the genomic landscape of BRAF mutant tumors across three common cancer types with a relatively high proportion of BRAF mutations: melanoma, NSCLC, and CRC. In NSCLC, non-V600 BRAF mutant tumors co-exist with mutations of the MAPK pathway signalling proteins, such as KRAS, but also frequently with loss of function mutations in tumor suppressor genes that are involved in other cellular processes such as metabolism (STK11) and oxidative stress response (KEAP1). STK11 (also known as liver kinase B1 – LKB1) is a kinase that acts as a metabolic sensor. In low energy/nutrient deprivation conditions, LKB1 phosphorylates and activates AMPK. Phosphorylated AMPK inhibits mTOR, thus wild-type LKB1 limits anabolic processes such as protein synthesis and promotes a switch to catabolic processes in low-energy conditions. In STK11/LKB1 mutant tumors there is increased mTOR activity and cell proliferation even in low nutrient conditions [31]. Nearly 30% of NSCLC with Class 2 or 3 BRAF mutations had co-occurring loss of function STK11 mutations. These findings suggest that Class 2 and 3 mutations – which have lower transformation capacity than Class 1 BRAF mutations [2] – may cooperate with other signalling pathways to drive tumorigenesis. These findings also suggest that therapeutic strategies targeting mTOR may cooperate with MAPK inhibitors in NSCLC. KEAP1 (Kelch-like ECH associated protein 1) is a ubiquitin ligase and a component of the NRF2-KEAP1 complex which regulates intracellular homeostasis to reactive oxygen species. KEAP1 regulates ROS accumulation by promoting the transcription of oxidative stress proteins and detoxifying enzymes [31]. Mutant KEAP1 protein has reduced affinity to the NRF2 transcription factor, leading to its constitutive activation. NRF2’s role as a tumor suppressor or oncogene is still unclear as it promotes the transcription of anti-inflammatory genes in healthy cells while its involvement in the metabolic activation of lung carcinogens may overshadow the protective effects of NRF2 and promote tumor growth[32]. Co-alterations of these two proteins on the short arm of chromosome 19 (KEAP1 and STK11/LKB1) have been shown to be associated with lung cancers [31]. The presence of KEAP1 and STK11 mutations is associated with resistance to chemotherapy and immunotherapy in NSCLC and is a negative prognostic factor [33]. Combination or novel agents are being studied in KEAP1/STK11 mutant NSCLC. Some promising results were shown in a phase II trial of KEAP1 mutant squamous lung cancers treated with single agent novel mammalian target of rapamycin complex 1 and 2 inhibitors (mTORC 1/2 inhibitor – Sapanisertib) [34]. Glutaminase inhibitor, telaglenastat, has also been proposed as a targeted treatment pathway for KEAP1/STK11 mutant tumours and is currently being evaluated in combination with Sapanisertib (NCT03872427; NCT04250545). Together, this data suggests that there may be a role for investigating combinations of these agents with MAPK inhibitors in NSCLCs with co-occurring Class 2 or 3 BRAF mutations and STK11 and/or KEAP1 mutations in preclinical models and potentially in clinical trials.

In melanoma, we found that BRAF Class 1 mutations occur in younger patients compared to Class 2/3 non-V600 BRAF mutations. BRAF mutant melanomas are strongly associated with UV exposure [35]. BRAF mutant melanoma cancers carry a high co-mutation burden mostly due to varying exposure to UV radiation and many of these mutations and genomic alterations are of unknown significance. Notably, when Class 2 and 3 alterations were compared to those of BRAF Class 1, UV Response pathways were significantly enriched in the non-V600 melanomas. Alterations in Wnt-beta Catenin signaling was significantly enriched in both Class 2 and 3 melanomas compared to Class 1. It has been reported that melanoma cells utilize Wnt signaling for proliferation and transformation, but the relative requirement of this pathway in different genomic contexts for melanoma progression remains unclear [36]. Alterations in G2M Checkpoint and E2F targets were also enriched in BRAF Class 3 melanoma. The genes within these gene-sets are critical for cell cycle progression, which could indicate some dysregulation of the cell cycle in Class 3 BRAF mutant tumors. Cyclins are critical regulators of the activity of cyclin-dependent kinases (ie. CDK2, CDK4, CDK6) that are responsible cell cycle progression from G1 to S-phase. Cyclin D is regulated at the transcriptional level by the MAPK pathway [37]. Class 1 BRAF mutant mutations induce expression of cyclin D. In BRAF mutant melanoma treatment with BRAF/MEK inhibitors dramatically down-regulates Cyclin D1 expression, thereby preventing cell cycle progression [38]. Interestingly, amplification of CCND1, the gene coding for Cyclin D1, promotes resistance to BRAF inhibitors in melanoma [39]. However, we did not observe a significant increase in Cyclin D or Cyclin E amplifications in the Class 2 & 3 BRAF mutant melanomas. Alternatively, Cyclin D can also be induced at the transcriptional level by non-MAPK pathways– including the Wnt-beta catenin signalling pathway [40]. Importantly, we did observe an enrichment for Wnt-beta catenin signalling pathway alterations in melanoma, NSCLC and CRCs with Class 2 or 3 BRAF mutations. Previously, we found that MAPK pathway inhibitors were less effective at inhibiting Cyclin D expression in Class 2 vs. Class 1 BRAF mutant tumors [16]. Taken together, these findings suggest that multiple non-MAPK signalling pathways may contribute to the induction of genes responsible for cell cycle progression in Class 2 & 3 BRAF mutant tumors. This would indicate less of reliance on MAPK pathway, which could explain decreased efficacy of MAPK pathway inhibitors in tumors with Class 2 & 3 BRAF mutations.

In our analysis, alterations in genes regulating the SWI/SNF chromatin remodeling complex were most frequent in Class 3 melanomas: these include ARID2, ARID1A and SMARCA4. The mutations in ARID1A and ARID2 were primarily loss of function mutations. ARID1A and ARID2 are components of the SWI/SWF chromatin remodeling complex (also known as BAF, Brg1-Associated Factors) and loss of these genes disrupts SWI/SNF activity and nucleosome remodeling, giving rise to abnormal gene regulation [41]. Loss-of-function ARID1A mutations independent of other mutations are not sufficient for tumorigenesis but may accelerate tumor development driven by co-occurring oncogenes [42]. With ARID2 depletion, BAF is redistributed where there is unfolded chromatin leading to transcription changes in genes regulating melanoma metastasis [43]. Thus, the relatively high incidence of co-occurring alterations of SWI/SNF complex related genes (ARID1A, ARID2, and SMARCA4), may represent a novel therapeutic opportunity in Class 3 BRAF mutant tumors. Currently, there is preclinical data to support the use of multiple classes of targeted therapies, including PARP inhibitors, Aurora kinase inhibitors, and SMARCA2 degraders in tumors ARID1A, ARID2 and SMARCA4 loss of function mutations [44, 45]. These inhibitors may also warrant further investigation in subsets of Class 3 BRAF mutant melanoma.

In CRC, we observe similar frequencies of co-existing RAS mutations in Class 2 and Class 3 mutant tumors. It has previously been reported that Class 1 BRAF mutations commonly occur in cancers that arise in the right colon whereas Class 2 & 3 non-V600 BRAF mutations more commonly arise in the left colon [46]. Importantly, there are different embryological origins of the cells that give rise to tumors in the left and right colon (hind-gut vs. mid-gut derivatives) [47]. Right-sided CRC frequently harbors Class 1 BRAF mutations, a finding that is compatible with the fact that tumors deriving from this tissue are largely driven by EGFR-independent oncogenic inputs [13]. Meanwhile, left-sided disease relies upon EGFR as a critical driver of cell proliferation through the MAPK pathway, and therefore this EGFR signalling is amplified by additional downstream MAPK driver mutations, such as Class 2 & 3 BRAF mutations. From retrospective data, it appears there is a better response rate to EGFR inhibitors in mCRC with Class 3 vs. Class 2 BRAF mutations [49]. However, it is not yet known whether the targeted therapy combination of BRAF + EGFR inhibitors that are effective for Class 1 BRAF mutant mCRC are also effective in Class 2 and 3 BRAF mutant mCRC – but ongoing clinical trials are actively investigating this question [9, 50].

We identified a strong relationship between Black race and increased prevalence of non-V600 BRAF mutations. This relationship is most pronounced in CRC, the one tumor type where non-V600 BRAF mutations tended to occur in younger patients. Indeed, the median age (at the time of sequencing) for patients with Class 2 and 3 BRAF mutant CRC was 62 and 55 years, respectively, compared to 66 years for patients with Class 1 BRAF mutant CRC. Many of these patients would have been even younger at the time of their cancer diagnosis. The incidence of early-onset CRC has been increasing, and early-onset CRC is even more predominant in the Black community [51]. Despite a lower incidence of BRAF V600 mutated CRC in Black patients, the mortality for this population remains high [28]. Early-onset CRC is typically left–sided cancer and has a unique genomic composition compared to cancers diagnosed after age 50 [29]. APC mutations occur more commonly in early-onset CRC and are associated with a poor prognosis [52]. We observed a high incidence of co-occurring APC mutations in Class 2/3 BRAF mutant CRC (29.35%, 45.45%, 68.75% for Class 1, 2, 3 respectively) compared to Class 1 CRC. Loss of function APC mutations result in increased Wnt-beta Catenin pro-oncogenic signalling [53]. Indeed, we found that alterations in the Wnt-beta catenin signalling pathway were enriched in Class 3 vs. Class 1 BRAF mutant tumors. Black patients with early-onset CRC are more likely to present with metastatic disease and experience shorter overall survival compared to patients of other races [51]. Moreover, Black and other non-white patients are typically under-represented in cancer sequencing datasets and in clinical trials in oncology [54]. Clinicians should have a high index of suspicion for the presence of non-V600 BRAF mutations in younger, Black patients with metastatic CRC. These are potentially actionable mutations for which treatment strategies have not yet been defined. However, there are several on-going clinical trials investigating novel MAPK pathway inhibitors that are enrolling patients with metastatic non-V600 BRAF mutant cancers (NCT03839342; NCT05503797; NCT04913285; NCT04249843).

Finally, we found that BRAF Class 3 mutations were more common in men than in women. In many cancer types, including melanoma, men have worse outcomes than women [55]. Moreover, men with Class 1 BRAF mutant melanoma are less likely to benefit from BRAF + MEK inhibitor therapy [56]. This finding suggests that there may be a hormonal explanation for these differences. Indeed, supplemental testosterone mitigates the effects of BRAF + MEK inhibition, and blockade of the androgen receptor promotes the anti-tumor activity of BRAF + MEK inhibitors in mouse models of Class 1 BRAF mutant melanoma [56]. Non-genomic signalling from the androgen receptor results in upstream activation of the MAPK pathway, which is essential for the oncogenic function of Class 3 BRAF mutant tumors [57]. As such, the increased incidence of Class 3 BRAF mutations in men, and the limited efficacy of MAPK inhibitors in these tumors suggests that there may be a role for investigating inhibitors of androgen receptor activity in men with Class 3 BRAF mutant tumors [13].

Together, this study describes analysis of the largest dataset of BRAF mutant tumors published to date. However, this analysis also comes with several limitations. The age at diagnosis is not reported for these patients. The age reported is the age at the time of tumor sequencing which may have occurred many years after of diagnosis. Primary race is self-reported and there is insufficient data for patients with mixed race backgrounds. Frequently, biological and health outcomes associated with gender and/or race are a consequence of differences in environmental exposures, diet, health care access, systemic and structural racism – rather than genetic differences between individuals of different races [58]. This dataset lacks information on social health determinants and lifestyle factors known to increase tumorigenesis, which limits our ability to interpret our results.

A key limitation of our genomic analysis is that the sequencing assay varied substantially between samples and many genes were not sequenced in all samples. Thus, we limited our analysis to genes with minimum 15 samples mutated across the BRAF Classes. Therefore, we are missing information on many genes that were sequenced in a minority of samples. We performed pathway enrichment analysis on significantly differentially altered genes. However, the functional significance of these alterations was not always known. Thus, our pathway enrichment analysis should be considered hypothesis-generating.

We believe this analysis of a large cohort of BRAF mutant cancers will help identify subsets of populations that would benefit from novel targeted therapies. It is likely that non-V600 BRAF mutations will become more frequently identified in the future, as these mutations are increasingly being identified as drivers of resistance to new targeted agents [59]. This work emphasizes the importance of continuing cancer sequencing projects, and the insight that can be gained from these large-scale sequencing efforts – particularly for rare driver mutations.

## Acknowledgments

ER acknowledges a Marathon of Hope Cancer Centers Network Health Informatics and Data Science Award. AANR acknowledges salary support from a Fonds de recherche du Québec – Santé (FRQS) Chercheuses-Boursières Cliniciennes Award. This research was supported by funding from the TransMedTech Institute and Apogee Canada Research Excellence Fund, a Canadian Cancer Society Challenge Grant (grant #707457) to AANR and AS, and a Conquer Cancer Foundation of ASCO Career Development Award to AANR.

## Conflict of interest

The following authors declare the listed conflicts of interest:

Anna Spreafico

Honoraria: Bristol Myers Squibb, Medison & Immunocore

Consulting or Advisory Role: Novartis, Merck, Bristol Myers Squibb, Oncorus, Medison & Immunocore

Research Funding: Bristol Myers Squibb, Novartis, Merck, Symphogen, AstraZeneca/MedImmune, Bayer, Surface Oncology, Janssen Oncology, Northern Biologics, Replimune, Roche, Alkermes, Array BioPharma, GlaxoSmithKline, Treadwell Therapeutics (Inst), Amgen (Inst)

Travel, Accommodations, Expenses: Merck, Bristol Myers Squibb, Idera, Bayer, Janssen Oncology, Roche

Gerald Batist

Partner in a Genome Canada proteomics grant with AstraZeneca.

April Rose

Employment: Merck, Recipient: an immediate family member

Stock and Other Ownership interests: Merck by an immediate family member

Consulting or Advisory Role: EMD Serono, Advanced Accelerator Applications/Novartis Research Funding: Canadian Institutes of Health Research (CIHR), Canadian Cancer Society, Conquer Cancer Foundation, Jewish General Hospital Foundation, TransMedTech Institute, Canada Foundation for Innovation, AstraZeneca Canada, Merck, Pfizer, Seattle Genetics,

The remaining authors have no conflicts of interests to declare.

