## Supplemental Table for "The clinical, genomic, and transcriptomic landscape of BRAF mutant cancers"

### Supplemental Figures & Tables

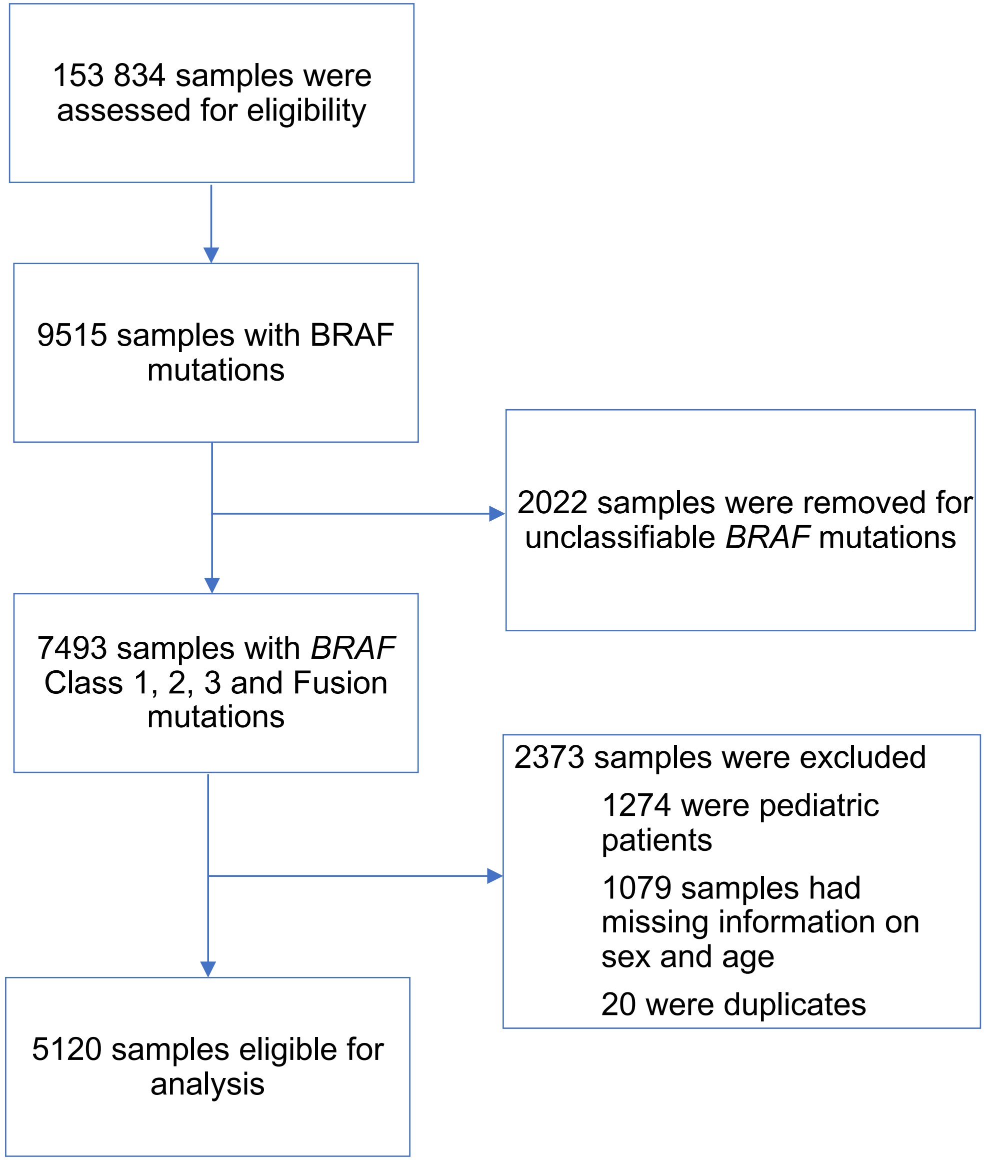
153 834 samples were assessed for eligibility
9515 samples with BRAF mutations
2022 samples were removed for unclassifiable *BRAF* mutations
7493 samples with *BRAF* Class 1, 2, 3 and Fusion mutations
2373 samples were excluded
1274 were pediatric patients
1079 samples had missing information on sex and age
20 were duplicates
5120 samples eligible for analysis

Figure S1

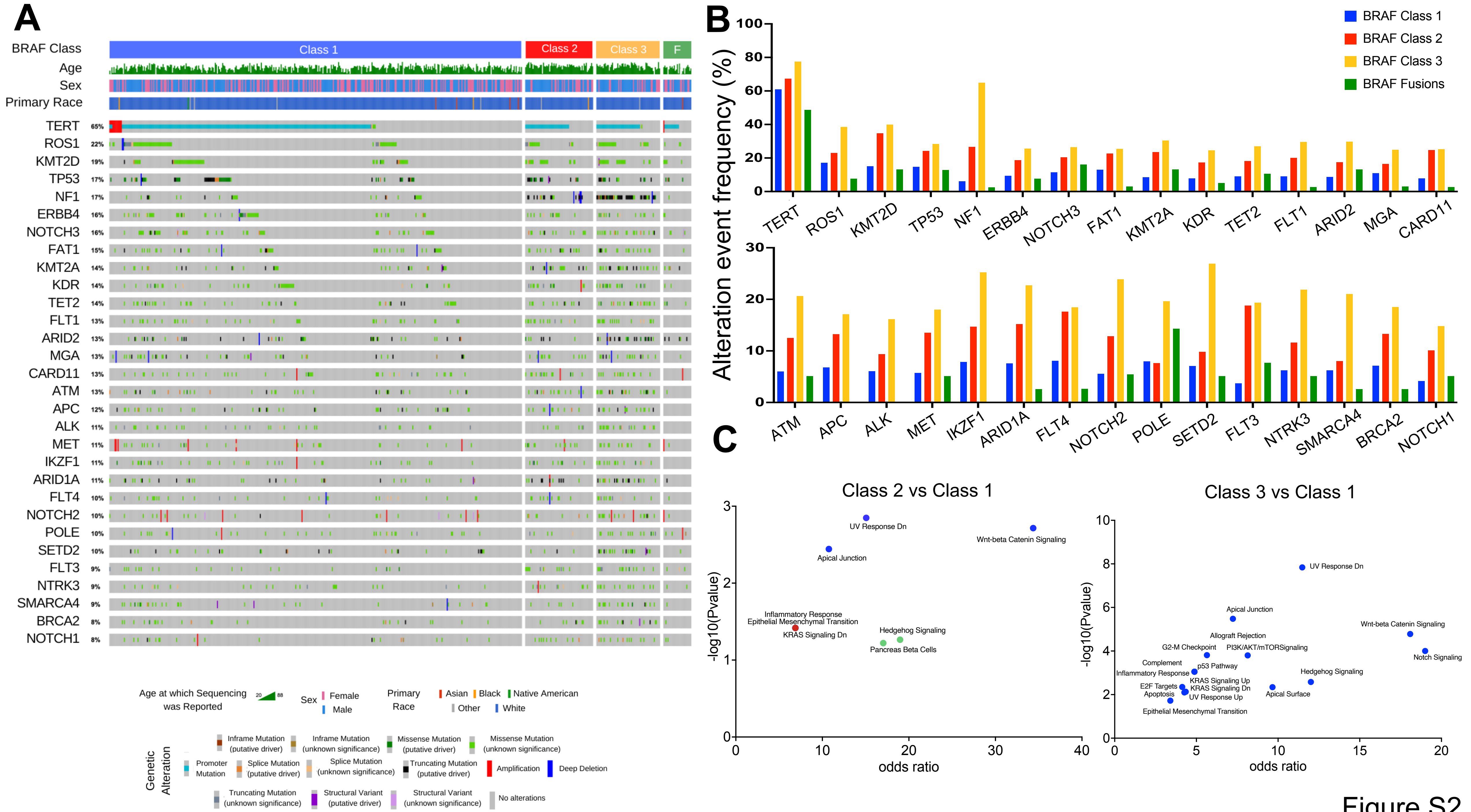
A
BRAF Class
Age
Sex
Primary Race
Class 1
Class 2
Class 3
F
TERT 65%
ROS1 22%
KMT2D 19%
TP53 17%
NF1 17%
ERBB4 16%
NOTCH3 16%
FAT1 15%
KMT2A 14%
KDR 14%
TET2 14%
FLT1 13%
ARID2 13%
MGA 13%
CARD11 13%
ATM 13%
APC 12%
ALK 11%
MET 11%
IKZF1 11%
ARID1A 11%
FLT4 10%
NOTCH2 10%
POLE 10%
SETD2 10%
FLT3 9%
NTRK3 9%
SMARCA4 9%
BRCA2 8%
NOTCH1 8%
Age at which Sequencing was Reported
20
88
Sex
Female
Male
Primary Race
Asian
Black
Native American
Other
White
Genetic Alteration
Inframe Mutation (putative driver)
Inframe Mutation (unknown significance)
Missense Mutation (putative driver)
Missense Mutation (unknown significance)
Promoter Mutation
Splice Mutation (putative driver)
Splice Mutation (unknown significance)
Truncating Mutation (putative driver)
Amplification
Deep Deletion
Truncating Mutation (unknown significance)
Structural Variant (putative driver)
Structural Variant (unknown significance)
No alterations
B
Alteration event frequency (%)
BRAF Class 1
BRAF Class 2
BRAF Class 3
BRAF Fusions
100
80
60
40
20
0
TERT
ROS1
KMT2D
TP53
NF1
ERBB4
NOTCH3
FAT1
KMT2A
KDR
TET2
FLT1
ARID2
MGA
CARD11
30
20
10
0
ATM
APC
ALK
MET
IKZF1
ARID1A
FLT4
NOTCH2
POLE
SETD2
FLT3
NTRK3
SMARCA4
BRCA2
NOTCH1
C
Class 2 vs Class 1
-log10(Pvalue)
odds ratio
3
2
1
0
0
10
20
30
40
UV Response Dn
Wnt-beta Catenin Signaling
Apical Junction
Inflammatory Response
Epithelial Mesenchymal Transition
KRAS Signaling Dn
Hedgehog Signaling
Pancreas Beta Cells
Class 3 vs Class 1
-log10(Pvalue)
odds ratio
10
8
6
4
2
0
0
5
10
15
20
UV Response Dn
Apical Junction
Allograft Rejection
Wnt-beta Catenin Signaling
G2-M Checkpoint
PI3K/AKT/mTORSignaling
Notch Signaling
Complement
p53 Pathway
Inflammatory Response
Hedgehog Signaling
KRAS Signaling Up
E2F Targets
KRAS Signaling Dn
Apoptosis
UV Response Up
Apical Surface
Epithelial Mesenchymal Transition

Figure S2

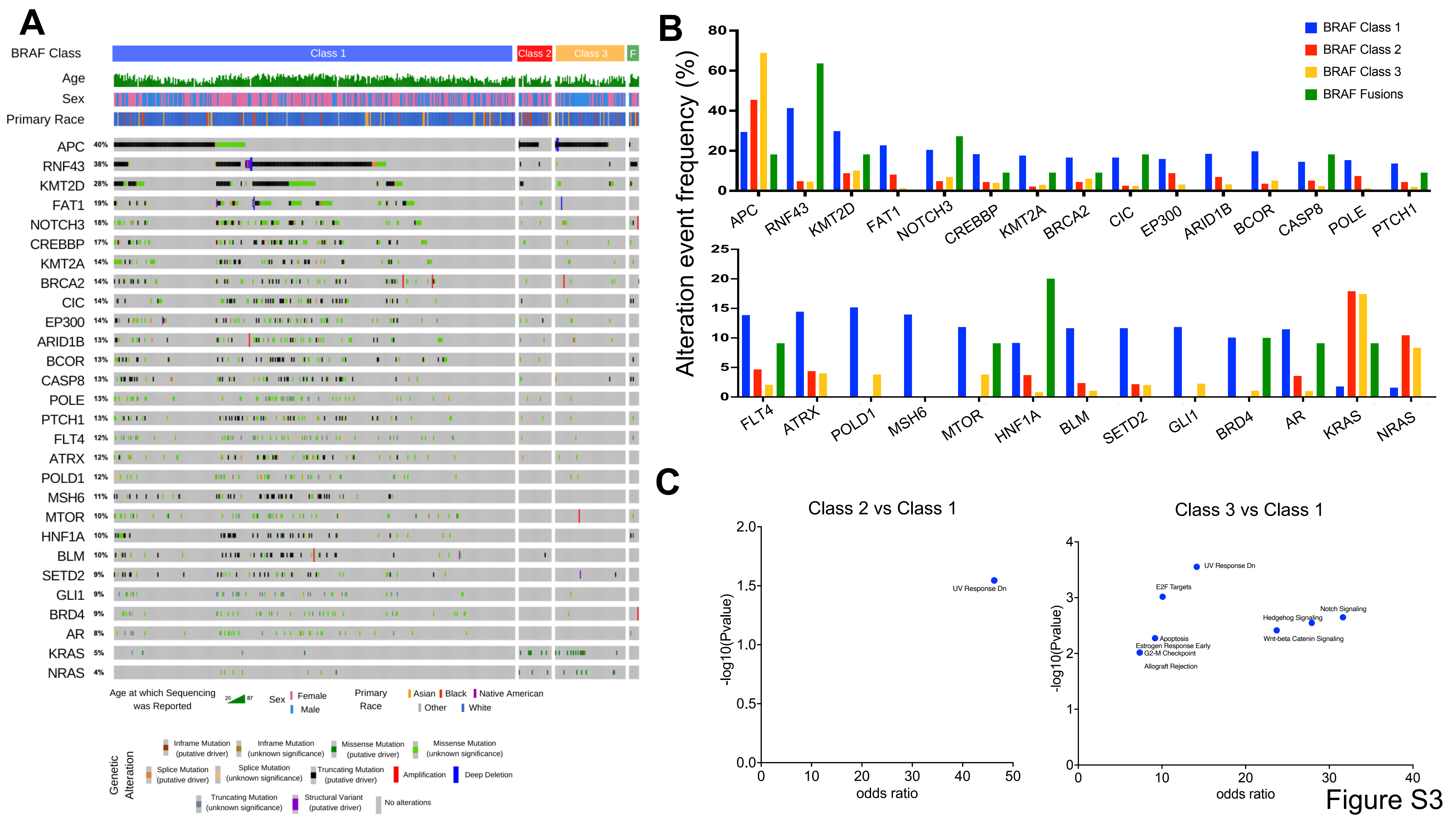
A
BRAF Class
Class 1
Class 2
Class 3
F
Age
Sex
Primary Race
APC 40%
RNF43 38%
KMT2D 28%
FAT1 19%
NOTCH3 18%
CREBBP 17%
KMT2A 14%
BRCA2 14%
CIC 14%
EP300 14%
ARID1B 13%
BCOR 13%
CASP8 13%
POLE 13%
PTCH1 13%
FLT4 12%
ATRX 12%
POLD1 12%
MSH6 11%
MTOR 10%
HNF1A 10%
BLM 10%
SETD2 9%
GLI1 9%
BRD4 9%
AR 8%
KRAS 5%
NRAS 4%
Age at which Sequencing was Reported
20 87
Sex
Female
Male
Primary Race
Asian
Black
Native American
Other
White
Genetic Alteration
Inframe Mutation (putative driver)
Inframe Mutation (unknown significance)
Missense Mutation (putative driver)
Missense Mutation (unknown significance)
Splice Mutation (putative driver)
Splice Mutation (unknown significance)
Truncating Mutation (putative driver)
Amplification
Deep Deletion
Truncating Mutation (unknown significance)
Structural Variant (putative driver)
No alterations
B
Alteration event frequency (%)
BRAF Class 1
BRAF Class 2
BRAF Class 3
BRAF Fusions
APC
RNF43
KMT2D
FAT1
NOTCH3
CREBBP
KMT2A
BRCA2
CIC
EP300
ARID1B
BCOR
CASP8
POLE
PTCH1
FLT4
ATRX
POLD1
MSH6
MTOR
HNF1A
BLM
SETD2
GLI1
BRD4
AR
KRAS
NRAS
C
Class 2 vs Class 1
-log10(Pvalue)
odds ratio
UV Response Dn
Class 3 vs Class 1
UV Response Dn
E2F Targets
Notch Signaling
Hedgehog Signaling
Wnt-beta Catenin Signaling
Apoptosis
Estrogen Response Early
G2-M Checkpoint
Allograft Rejection

Figure S3

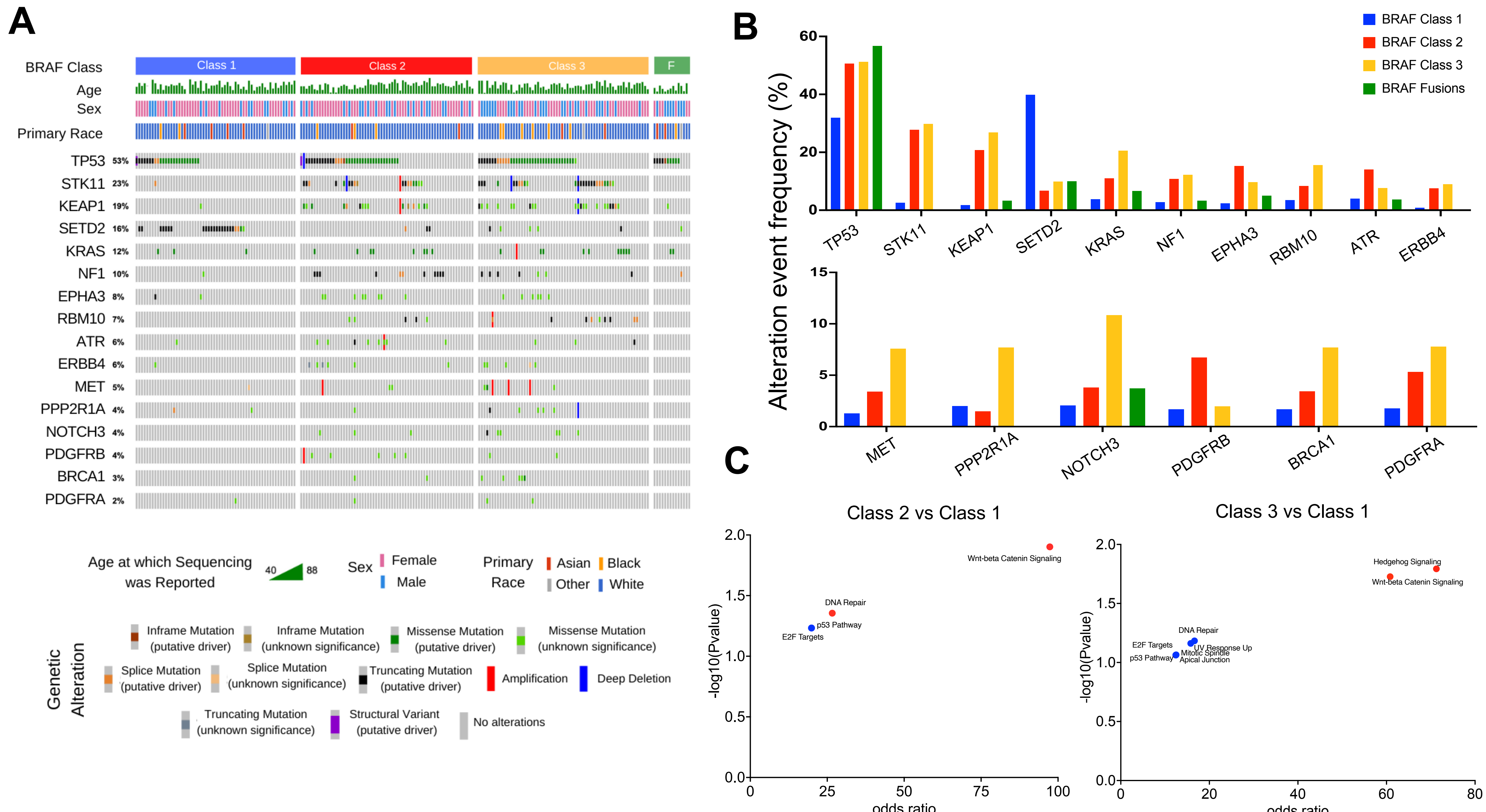
A
BRAF Class
Class 1
Class 2
Class 3
F
Age
Sex
Primary Race
TP53 53%
STK11 23%
KEAP1 19%
SETD2 16%
KRAS 12%
NF1 10%
EPHA3 8%
RBM10 7%
ATR 6%
ERBB4 6%
MET 5%
PPP2R1A 4%
NOTCH3 4%
PDGFRB 4%
BRCA1 3%
PDGFRA 2%
Age at which Sequencing was Reported
40
88
Sex
Female
Male
Primary Race
Asian
Black
Other
White
Genetic Alteration
Inframe Mutation (putative driver)
Inframe Mutation (unknown significance)
Missense Mutation (putative driver)
Missense Mutation (unknown significance)
Splice Mutation (putative driver)
Splice Mutation (unknown significance)
Truncating Mutation (putative driver)
Amplification
Deep Deletion
Truncating Mutation (unknown significance)
Structural Variant (putative driver)
No alterations
B
BRAF Class 1
BRAF Class 2
BRAF Class 3
BRAF Fusions
Alteration event frequency (%)
TP53
STK11
KEAP1
SETD2
KRAS
NF1
EPHA3
RBM10
ATR
ERBB4
MET
PPP2R1A
NOTCH3
PDGFRB
BRCA1
PDGFRA
C
Class 2 vs Class 1
Wnt-beta Catenin Signaling
DNA Repair
p53 Pathway
E2F Targets
-log10(Pvalue)
odds ratio
Class 3 vs Class 1
Hedgehog Signaling
Wnt-beta Catenin Signaling
DNA Repair
E2F Targets
UV Response Up
Mitotic Spindle
p53 Pathway
Apical Junction

Figure S4

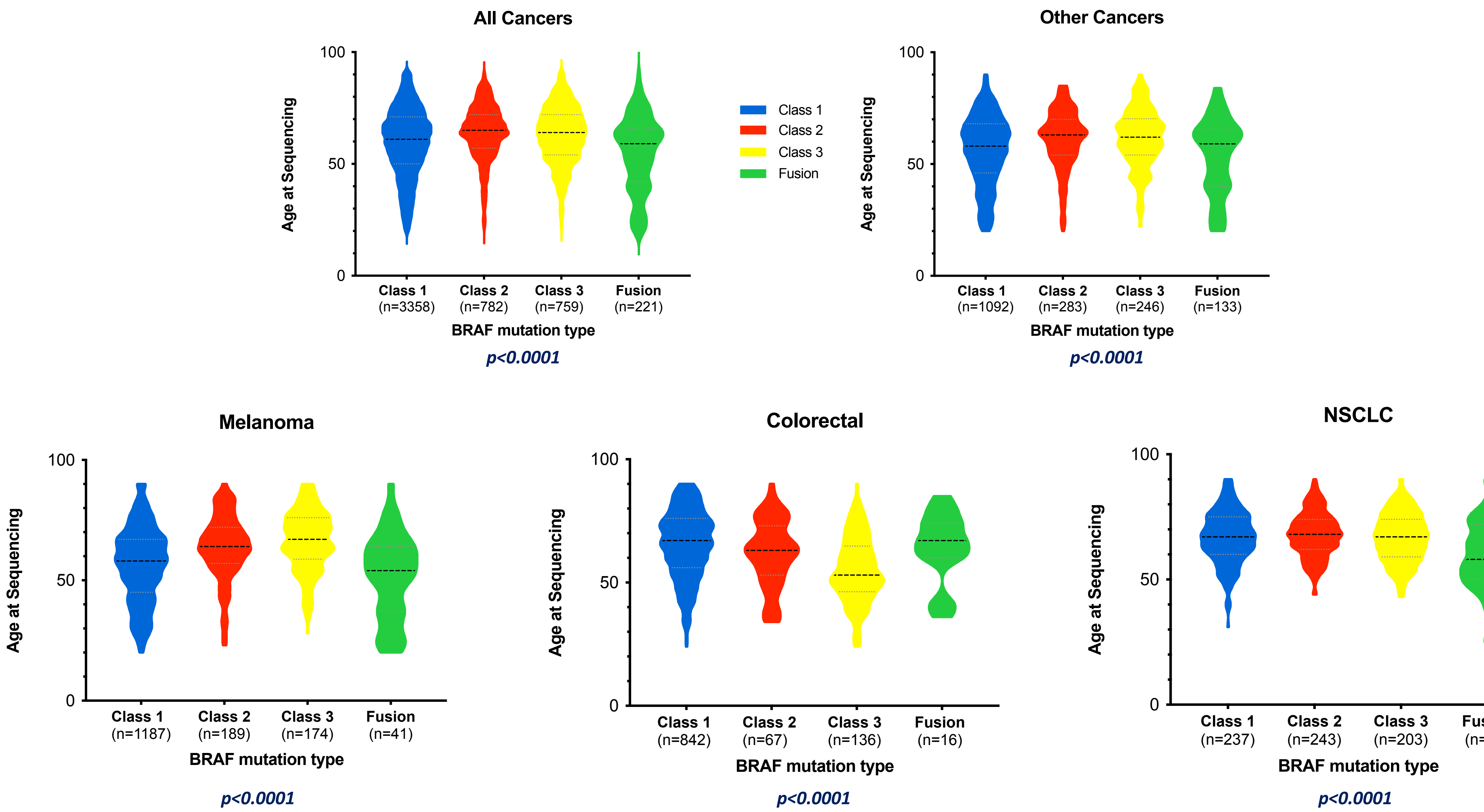
All Cancers
Age at Sequencing
100
50
0
Class 1
(n=3358)
Class 2
(n=782)
Class 3
(n=759)
Fusion
(n=221)
BRAF mutation type
p<0.0001
Class 1
Class 2
Class 3
Fusion
Other Cancers
Age at Sequencing
100
50
0
Class 1
(n=1092)
Class 2
(n=283)
Class 3
(n=246)
Fusion
(n=133)
BRAF mutation type
p<0.0001
Melanoma
Age at Sequencing
100
50
0
Class 1
(n=1187)
Class 2
(n=189)
Class 3
(n=174)
Fusion
(n=41)
BRAF mutation type
p<0.0001
Colorectal
Age at Sequencing
100
50
0
Class 1
(n=842)
Class 2
(n=67)
Class 3
(n=136)
Fusion
(n=16)
BRAF mutation type
p<0.0001
NSCLC
Age at Sequencing
100
50
0
Class 1
(n=237)
Class 2
(n=243)
Class 3
(n=203)
Fusion
(n=31)
BRAF mutation type
p<0.0001

**Figure S5**

**Supplementary Table 1**

| BRAF Class | Protein Change | Occurrence in dataset | BRAF Class | Protein Change | Occurrence in dataset | BRAF Class | Protein Change | Occurrence in dataset | BRAF Class | Protein Change | Occurrence in dataset |
|---|---|---|---|---|---|---|---|---|---|---|---|
| 1 | V600_K601delinsEI | 1 | 2 | K601E/I/N/Q | 181 | 2 | V504_R506dup | 1 | Fusion | AGK-BRAF fusion | 11 |
| 1 | V600_K601delinsEN | 1 | 2 | L485_A489delinsFS | 1 | 2 | V600_K601delinsE | 14 | Fusion | BRAF-AGK | 3 |
| 1 | V600_S602delinsRKT | 1 | 2 | L485_P490delinsF | 1 | 2 | V600_R603del | 2 | Fusion | BRAF-CDK5RAP2 fusion | 3 |
| 1 | V600D/E/G/K/L/R | 3355 | 2 | L485F/S/W | 16 | 2 | V600_S602del | 1 | Fusion | BRAF-INTERGENIC | 16 |
| 2 | A598_T599insI | 1 | 2 | L505H | 2 | 2 | V600_W604delinsDG/G/R | 5 | Fusion | BRAF-KIAA1549 | 10 |
| 2 | A598D | 1 | 2 | L597P/Q/R/S/V | 70 | 2 | X506_X506insRYSG | 2 | Fusion | BRAF-MKRN1 | 16 |
| 2 | A598dup | 1 | 2 | M484_L485delinsIM | 1 | 3 | D594A/E/G/H/N/V/Y | 303 | Fusion | BRAF-NDUFB2 | 3 |
| 2 | D594_T599dup | 23 | 2 | N486_A489delinsK | 1 | 3 | E501K/Q/V | 15 | Fusion | BRAF-SND1 | 18 |
| 2 | E586K | 13 | 2 | N486_A489delinsR | 1 | 3 | F247L | 6 | Fusion | BRAF-TMPRSS2 fusion | 3 |
| 2 | F468S | 8 | 2 | N486_P490del | 30 | 3 | G466A/E/L/R/V | 174 | Fusion | BRAF-TRIM24 fusion | 3 |
| 2 | F595I/L/S | 12 | 2 | P367L/S | 12 | 3 | G469E | 31 | Fusion | KIAA1549-BRAF fusion | 6 |

| | | | | | | | | | | | |
|---|---|---|---|---|---|---|---|---|---|---|---|
| 2 | G464A/E/I/R/V | 48 | 2 | S602_R603dup | 1 | 3 | G596C/D/R/S/V | 43 | Fusion | Other Fusions | 126 |
| 2 | G469A/I/K/L/L/R/S/V | 295 | 2 | T599_V600delinsR | 1 | 3 | K483E/N/T | 13 | | | |
| 2 | I592_A598dup | 1 | 2 | T599_V600insDT/EIAT/H/Q | 5 | 3 | N581D/H/I/S/T/Y | 134 | | | |
| 2 | K499E | 1 | 2 | T599delinsIP | 3 | 3 | Q257R | 2 | | | |
| 2 | K601_S602delinsNT | 1 | 2 | T599dup | 17 | 3 | S467L | 34 | | | |
| 2 | K601_S605delinsD | 1 | 2 | T599I | 5 | 3 | V471F | 4 | | | |
| 2 | K601_S605delinsN | 2 | 2 | V487_P492delinsA | 1 | Fusion | AGAP3-BRAF fusion | 3 | | | |

**Supplementary Table 2**

| Gene | Cytoband | Class 1 | Class 1 % | Class 2 | Class 2 % | Class 3 | Class 3 % | pValue | qValue | Most Enriched In |
|---|---|---|---|---|---|---|---|---|---|---|
| KEAP1 | 19p13.2 | 53 | 2.24 | 43 | 8.07 | 62 | 10.95 | 0 | 0 | (C) BRAF Class 3 |
| KRAS | 12p12.1 | 36 | 1.08 | 66 | 8.81 | 129 | 17.36 | 0 | 0 | (C) BRAF Class 3 |
| NF1 | 17q11.2 | 143 | 5.62 | 79 | 12.87 | 138 | 23.23 | 0 | 0 | (C) BRAF Class 3 |
| NRAS | 1p13.2 | 33 | 0.99 | 46 | 6.14 | 92 | 12.38 | 0 | 0 | (C) BRAF Class 3 |
| SERPINB4 | 18q21.33 | 0 | 0 | 1 | 20 | 0 | 0 | 0 | 0 | (B) BRAF Class 2 |
| STK11 | 19p13.3 | 56 | 1.71 | 79 | 11.53 | 79 | 11 | 0 | 0 | (B) BRAF Class 2 |
| TP53 | 17p13.1 | 894 | 26.71 | 278 | 39.66 | 326 | 44.23 | 0 | 0 | (C) BRAF Class 3 |
| APC | 5q22.2 | 356 | 11.33 | 85 | 12.01 | 159 | 22.62 | 7.77E-15 | 1.14E-12 | (C) BRAF Class 3 |
| RNF43 | 17q22 | 261 | 11.73 | 17 | 3.4 | 21 | 4.04 | 2.24E-12 | 2.85E-10 | (A) BRAF Class 1 v12 |
| HRAS | 11p15.5 | 17 | 0.51 | 13 | 1.74 | 27 | 3.64 | 2.43E-12 | 2.85E-10 | (C) BRAF Class 3 |
| FLT3 | 13q12.2 | 100 | 3.1 | 49 | 7.31 | 47 | 6.63 | 1.83E-08 | 1.95E-06 | (B) BRAF Class 2 |
| TERT | 5p15.33 | 842 | 34.19 | 126 | 23.2 | 144 | 26.23 | 8.07E-08 | 7.88E-06 | (A) BRAF Class 1 v12 |
| ATXN2 | 12q24.12 | 1 | 4 | 0 | 0 | 0 | 0 | 3.25E-07 | 2.72E-05 | (A) BRAF Class 1 v12 |
| DUSP22 | 6p25.3 | 1 | 4 | 0 | 0 | 0 | 0 | 3.25E-07 | 2.72E-05 | (A) BRAF Class 1 v12 |
| POT1 | 7q31.33 | 13 | 1.55 | 16 | 8.89 | 12 | 5.97 | 3.69E-07 | 2.88E-05 | (B) BRAF Class 2 |
| SMARCA4 | 19p13.2 | 160 | 6.3 | 46 | 8.17 | 74 | 12.63 | 1.06E-06 | 7.78E-05 | (C) BRAF Class 3 |
| PPP2R1A | 19q13.41 | 34 | 1.54 | 9 | 1.81 | 27 | 5.22 | 1.29E-06 | 8.87E-05 | (C) BRAF Class 3 |
| RMRP | 9p13.3 | 0 | 0 | 0 | 0 | 1 | 100 | 1.37E-06 | 8.92E-05 | (C) BRAF Class 3 |
| ERBB4 | 2q34 | 204 | 6.29 | 69 | 9.47 | 82 | 11.47 | 1.53E-06 | 9.41E-05 | (C) BRAF Class 3 |
| NOTCH2 | 1p12 | 110 | 4.4 | 39 | 7.03 | 55 | 9.67 | 1.67E-06 | 9.77E-05 | (C) BRAF Class 3 |
| MCL1 | 1q21.2 | 25 | 1.02 | 8 | 1.45 | 22 | 3.82 | 5.22E-06 | 0.00029 | (C) BRAF Class 3 |
| ERBB2 | 17q12 | 87 | 2.6 | 34 | 4.54 | 44 | 5.97 | 5.85E-06 | 0.00031 | (C) BRAF Class 3 |
| MET | 7q31.2 | 118 | 3.56 | 40 | 5.38 | 55 | 7.49 | 6.86E-06 | 0.00035 | (C) BRAF Class 3 |
| ETAA1 | 2p14 | 1 | 3.13 | 0 | 0 | 0 | 0 | 8.27E-06 | 0.00037 | (A) BRAF Class 1 v12 |
| FOXF1 | 16q24.1 | 1 | 3.13 | 0 | 0 | 0 | 0 | 8.27E-06 | 0.00037 | (A) BRAF Class 1 v12 |
| SLFN11 | 17q12 | 1 | 3.13 | 0 | 0 | 0 | 0 | 8.27E-06 | 0.00037 | (A) BRAF Class 1 v12 |
| ARID1A | 1p36.11 | 267 | 10.51 | 80 | 14.21 | 99 | 16.89 | 2.68E-05 | 0.00115 | (C) BRAF Class 3 |
| CBLB | 3q13.11 | 16 | 1.52 | 13 | 5.86 | 14 | 5.58 | 2.78E-05 | 0.00115 | (B) BRAF Class 2 |
| GNA14 | 9q21.2 | 0 | 0 | 3 | 20 | 0 | 0 | 2.84E-05 | 0.00115 | (B) BRAF Class 2 |
| KDR | 4q12 | 162 | 5.08 | 58 | 8.71 | 59 | 8.43 | 4.16E-05 | 0.00162 | (B) BRAF Class 2 |
| PTPN11 | 12q24.13 | 36 | 1.11 | 14 | 2.06 | 24 | 3.36 | 4.62E-05 | 0.00175 | (C) BRAF Class 3 |
| MAP3K5 | 6q23.3 | 0 | 0 | 1 | 4 | 4 | 13.33 | 5.38E-05 | 0.00197 | (C) BRAF Class 3 |
| PDGFRA | 4q12 | 101 | 3.1 | 36 | 4.95 | 46 | 6.37 | 5.81E-05 | 0.00206 | (C) BRAF Class 3 |
| COL2A1 | 12q13.11 | 5 | 3.21 | 6 | 24 | 6 | 20 | 6.40E-05 | 0.00218 | (B) BRAF Class 2 |
| EPHA3 | 3p11.1 | 92 | 4.84 | 36 | 8.49 | 42 | 9.74 | 6.53E-05 | 0.00218 | (C) BRAF Class 3 |
| FGFR1 | 8p11.23 | 54 | 1.62 | 29 | 3.87 | 24 | 3.26 | 8.70E-05 | 0.00276 | (B) BRAF Class 2 |
| SYNE1 | 6q25.2 | 11 | 2.14 | 6 | 8 | 10 | 10.64 | 8.72E-05 | 0.00276 | (C) BRAF Class 3 |
| ERBB3 | 12q13.2 | 81 | 3.15 | 29 | 4.71 | 41 | 6.86 | 9.78E-05 | 0.00301 | (C) BRAF Class 3 |
| IL2RB | 22q12.3 | 0 | 0 | 2 | 16.67 | 0 | 0 | 0.00013 | 0.00376 | (B) BRAF Class 2 |
| RHPN2 | 19q13.11 | 6 | 1.7 | 4 | 5.88 | 8 | 11.59 | 0.0002 | 0.00579 | (C) BRAF Class 3 |
| TET2 | 4q24 | 155 | 6.22 | 41 | 7.32 | 65 | 11.09 | 0.00022 | 0.00615 | (C) BRAF Class 3 |
| RHOH | 4p14 | 3 | 0.44 | 0 | 0 | 6 | 3.57 | 0.00027 | 0.00757 | (C) BRAF Class 3 |
| ATR | 3q23 | 126 | 5.7 | 53 | 10.66 | 33 | 6.38 | 0.00029 | 0.00794 | (B) BRAF Class 2 |
| HLA-A | 6p22.1 | 48 | 3.94 | 3 | 1.07 | 0 | 0 | 0.0003 | 0.00801 | (A) BRAF Class 1 v12 |

| | | | | | | | | | | |
|---|---|---|---|---|---|---|---|---|---|---|
| PTPRT | 0q12-q13.1 | 159 | 11.26 | 37 | 11.86 | 60 | 19.54 | 0.00035 | 0.00918 | (C) BRAF Class 3 |
| CDKN1C | 11p15.4 | 5 | 0.51 | 7 | 3.33 | 1 | 0.44 | 0.00036 | 0.00918 | (B) BRAF Class 2 |
| NFE2L2 | 2q31.2 | 31 | 1.22 | 19 | 3.37 | 16 | 2.73 | 0.0004 | 0.00999 | (B) BRAF Class 2 |
| FOXA1 | 14q21.1 | 39 | 1.98 | 22 | 4.97 | 7 | 1.54 | 0.00042 | 0.0103 | (B) BRAF Class 2 |
| PIK3CA | 3q26.32 | 311 | 9.29 | 75 | 10.01 | 104 | 14.05 | 0.00053 | 0.0127 | (C) BRAF Class 3 |
| FGFR2 | 10q26.13 | 86 | 2.57 | 30 | 4.01 | 38 | 5.14 | 0.00059 | 0.0139 | (C) BRAF Class 3 |
| PTPRD | 9p24.1-p23 | 185 | 10.66 | 41 | 10.82 | 69 | 17.47 | 0.00062 | 0.014 | (C) BRAF Class 3 |
| SRC | 20q11.23 | 22 | 0.83 | 10 | 1.85 | 15 | 2.69 | 0.00062 | 0.014 | (C) BRAF Class 3 |
| FANCL | 2p16.1 | 16 | 1.26 | 11 | 4.33 | 2 | 0.67 | 0.00065 | 0.0143 | (B) BRAF Class 2 |
| PRKACA | 19p13.12 | 0 | 0 | 2 | 5.56 | 0 | 0 | 0.00066 | 0.0143 | (B) BRAF Class 2 |
| MSH3 | 5q14.1 | 95 | 8.89 | 8 | 3.49 | 8 | 3.36 | 0.00072 | 0.0154 | (A) BRAF Class 1 v12 |
| KDM6A | Xp11.3 | 66 | 2.68 | 30 | 5.42 | 28 | 4.85 | 0.00081 | 0.0168 | (B) BRAF Class 2 |
| RET | 10q11.21 | 122 | 3.65 | 27 | 3.6 | 49 | 6.62 | 0.00082 | 0.0168 | (C) BRAF Class 3 |
| ATM | 11q22.3 | 256 | 8.04 | 65 | 9.76 | 87 | 12.43 | 0.00083 | 0.0168 | (C) BRAF Class 3 |
| RAF1 | 3p25.2 | 37 | 1.42 | 16 | 2.55 | 22 | 3.64 | 0.00093 | 0.0185 | (C) BRAF Class 3 |
| BRIP1 | 17q23.2 | 67 | 2.72 | 23 | 4.16 | 33 | 5.72 | 0.00103 | 0.0201 | (C) BRAF Class 3 |
| FLT1 | 13q12.3 | 130 | 5.28 | 36 | 6.51 | 54 | 9.36 | 0.00107 | 0.0206 | (C) BRAF Class 3 |
| NOTCH1 | 9q34.3 | 178 | 5.42 | 52 | 7.58 | 63 | 8.73 | 0.00115 | 0.0218 | (C) BRAF Class 3 |
| CASR | q13.33-q21 | 3 | 3.26 | 3 | 20 | 3 | 30 | 0.0017 | 0.0313 | (C) BRAF Class 3 |
| IGF1 | 12q23.2 | 6 | 0.47 | 1 | 0.34 | 7 | 2.39 | 0.00171 | 0.0313 | (C) BRAF Class 3 |
| CARD11 | 7p22.2 | 157 | 6.38 | 43 | 7.78 | 61 | 10.57 | 0.00199 | 0.0359 | (C) BRAF Class 3 |
| SMAD3 | 15q22.33 | 25 | 1.69 | 7 | 2.13 | 16 | 4.86 | 0.00209 | 0.0371 | (C) BRAF Class 3 |
| EPHA7 | 6q16.1 | 108 | 6.08 | 39 | 10.16 | 38 | 9.84 | 0.00217 | 0.0379 | (B) BRAF Class 2 |
| DDX31 | 9q34.13 | 1 | 1.09 | 3 | 20 | 1 | 10 | 0.0023 | 0.0391 | (B) BRAF Class 2 |
| DGKH | 13q14.11 | 0 | 0 | 2 | 16.67 | 2 | 10 | 0.00231 | 0.0391 | (B) BRAF Class 2 |
| SMC1A | Xp11.22 | 9 | 1 | 7 | 4.67 | 2 | 1.17 | 0.00244 | 0.0409 | (B) BRAF Class 2 |
| PBRM1 | 3p21.1 | 107 | 4.51 | 37 | 6.94 | 43 | 7.6 | 0.00318 | 0.0522 | (C) BRAF Class 3 |
| EGFR | 7p11.2 | 131 | 3.92 | 28 | 3.74 | 49 | 6.62 | 0.00324 | 0.0522 | (C) BRAF Class 3 |
| H3F3B | 17q25.1 | 6 | 0.32 | 5 | 1.19 | 7 | 1.65 | 0.00332 | 0.0522 | (C) BRAF Class 3 |
| CBL | 11q23.3 | 57 | 2.22 | 17 | 2.98 | 28 | 4.71 | 0.00334 | 0.0522 | (C) BRAF Class 3 |
| TGFBR2 | 3p24.1 | 112 | 7.13 | 9 | 2.51 | 19 | 5.25 | 0.00335 | 0.0522 | (A) BRAF Class 1 v12 |
| DICER1 | 14q32.13 | 91 | 3.78 | 19 | 3.58 | 38 | 6.87 | 0.00355 | 0.0547 | (C) BRAF Class 3 |
| NTRK3 | 15q25.3 | 110 | 4.33 | 34 | 6.04 | 44 | 7.51 | 0.00377 | 0.0573 | (C) BRAF Class 3 |
| ANKRD26 | 10p12.1 | 45 | 13.51 | 1 | 2.5 | 0 | 0 | 0.0039 | 0.0579 | (A) BRAF Class 1 v12 |
| TP63 | 3q28 | 57 | 4.55 | 16 | 5.61 | 26 | 9.63 | 0.00391 | 0.0579 | (C) BRAF Class 3 |
| MKRN1 | 7q34 | 0 | 0 | 0 | 0 | 2 | 0.27 | 0.00403 | 0.0589 | (C) BRAF Class 3 |
| MAP3K13 | 3q27.2 | 20 | 1.52 | 7 | 2.22 | 14 | 4.52 | 0.00419 | 0.0606 | (C) BRAF Class 3 |
| ARID2 | 12q12 | 180 | 7.4 | 48 | 8.96 | 65 | 11.61 | 0.00429 | 0.0613 | (C) BRAF Class 3 |
| AIP | 11q13.2 | 0 | 0 | 0 | 0 | 1 | 10 | 0.00453 | 0.0634 | (C) BRAF Class 3 |
| EFNB3 | 17p13.1 | 0 | 0 | 0 | 0 | 1 | 20 | 0.00455 | 0.0634 | (C) BRAF Class 3 |
| KIT | 4q12 | 79 | 2.36 | 26 | 3.47 | 33 | 4.44 | 0.00467 | 0.0644 | (C) BRAF Class 3 |
| EZH1 | 17q21.2 | 7 | 1.05 | 1 | 0.62 | 7 | 4.4 | 0.00487 | 0.0663 | (C) BRAF Class 3 |
| ELOC | 8q21.11 | 6 | 0.33 | 3 | 0.74 | 7 | 1.7 | 0.00518 | 0.0697 | (C) BRAF Class 3 |
| EPHA2 | 1p36.13 | 0 | 0 | 0 | 0 | 2 | 4.08 | 0.00553 | 0.0736 | (C) BRAF Class 3 |
| PMS1 | 2q32.2 | 55 | 2.38 | 14 | 2.75 | 26 | 4.97 | 0.00564 | 0.0736 | (C) BRAF Class 3 |

| | | | | | | | | | | |
|---|---|---|---|---|---|---|---|---|---|---|
| KMT2D | 12q13.12 | 361 | 14.69 | 88 | 15.91 | 116 | 20.1 | 0.00566 | 0.0736 | (C) BRAF Class 3 |
| GATA6 | 18q11.2 | 28 | 1.98 | 14 | 4.91 | 13 | 4.02 | 0.00611 | 0.0786 | (B) BRAF Class 2 |
| OR5L1 | 11q12.1 | 1 | 1.16 | 2 | 16.67 | 3 | 15 | 0.00625 | 0.0786 | (B) BRAF Class 2 |
| CXCR4 | 2q22.1 | 12 | 0.63 | 9 | 2.07 | 8 | 1.81 | 0.00627 | 0.0786 | (B) BRAF Class 2 |
| PLCB4 | )p12.3-p12 | 5 | 2.31 | 4 | 10 | 5 | 11.9 | 0.00633 | 0.0786 | (C) BRAF Class 3 |
| XRCC3 | 14q32.33 | 20 | 2.92 | 0 | 0 | 0 | 0 | 0.00638 | 0.0786 | (A) BRAF Class 1 v12 |
| TRAF7 | 16p13.3 | 44 | 2.26 | 6 | 1.37 | 20 | 4.48 | 0.00666 | 0.0803 | (C) BRAF Class 3 |
| JAK3 | 19p13.11 | 102 | 3.17 | 34 | 4.74 | 38 | 5.36 | 0.00671 | 0.0803 | (C) BRAF Class 3 |
| EBF1 | 5q33.3 | 1 | 0.44 | 2 | 5.13 | 3 | 6.67 | 0.00672 | 0.0803 | (C) BRAF Class 3 |
| DROSHA | 5p13.3 | 24 | 4 | 1 | 0.66 | 11 | 7.91 | 0.0072 | 0.0852 | (C) BRAF Class 3 |
| ROS1 | 6q22.1 | 242 | 9.29 | 53 | 8.45 | 80 | 13.16 | 0.00732 | 0.0857 | (C) BRAF Class 3 |
| DDR2 | 1q23.3 | 55 | 2.06 | 22 | 3.42 | 25 | 4.03 | 0.00755 | 0.0874 | (C) BRAF Class 3 |
| PTK2B | 8p21.2 | 22 | 3.41 | 6 | 4.26 | 15 | 9.04 | 0.00766 | 0.0874 | (C) BRAF Class 3 |
| QKI | 6q26 | 48 | 3.62 | 3 | 1.13 | 3 | 0.96 | 0.00769 | 0.0874 | (A) BRAF Class 1 v12 |
| RUNX1T1 | 8q21.3 | 56 | 4.49 | 17 | 7.14 | 25 | 8.83 | 0.00801 | 0.0902 | (C) BRAF Class 3 |
| ASXL1 | 20q11.21 | 130 | 5.22 | 31 | 5.55 | 50 | 8.53 | 0.00825 | 0.092 | (C) BRAF Class 3 |
| IKX2-1-AS | 14q13.3 | 0 | 0 | 1 | 2.5 | 0 | 0 | 0.00845 | 0.0926 | (B) BRAF Class 2 |
| SPOP | 17q21.33 | 20 | 0.91 | 9 | 1.81 | 13 | 2.51 | 0.00846 | 0.0926 | (C) BRAF Class 3 |
| JAK2 | 9p24.1 | 62 | 1.95 | 20 | 2.8 | 27 | 3.82 | 0.00875 | 0.0949 | (C) BRAF Class 3 |
| LIFR | 5p13.1 | 4 | 2.15 | 3 | 9.68 | 5 | 11.9 | 0.00907 | 0.0974 | (C) BRAF Class 3 |
| RRAS2 | 11p15.2 | 3 | 0.45 | 3 | 1.85 | 5 | 3.14 | 0.00922 | 0.0974 | (C) BRAF Class 3 |
| SF3B1 | 2q33.1 | 58 | 2.21 | 24 | 4.1 | 23 | 3.79 | 0.00925 | 0.0974 | (B) BRAF Class 2 |
| FGFR4 | 5q35.2 | 71 | 2.76 | 18 | 2.9 | 31 | 5.17 | 0.00939 | 0.0974 | (C) BRAF Class 3 |
| BCOR | Xp11.4 | 156 | 6.35 | 30 | 5.42 | 18 | 3.1 | 0.00948 | 0.0974 | (A) BRAF Class 1 v12 |
| MPL | 1p34.2 | 51 | 1.63 | 22 | 3.33 | 18 | 2.58 | 0.00949 | 0.0974 | (B) BRAF Class 2 |
| SPTA1 | 1q23.1 | 95 | 15.13 | 25 | 26.04 | 27 | 22.31 | 0.00966 | 0.0982 | (B) BRAF Class 2 |
| CASP8 | 2q33.1 | 92 | 4.51 | 8 | 1.68 | 15 | 3.07 | 0.00972 | 0.0982 | (A) BRAF Class 1 v12 |
| CDKN2A | 9p21.3 | 449 | 13.7 | 121 | 17.66 | 118 | 16.43 | 0.0104 | 0.104 | (B) BRAF Class 2 |
| EZH2 | 7q36.1 | 46 | 1.52 | 18 | 2.8 | 20 | 2.97 | 0.0106 | 0.105 | (C) BRAF Class 3 |
| PTPRB | 12q15 | 8 | 5.13 | 5 | 20 | 5 | 16.67 | 0.0107 | 0.105 | (B) BRAF Class 2 |
| FBXW7 | 4q31.3 | 185 | 5.64 | 27 | 3.94 | 55 | 7.63 | 0.0113 | 0.111 | (C) BRAF Class 3 |
| EDNRB | 13q22.3 | 0 | 0 | 1 | 8.33 | 0 | 0 | 0.0116 | 0.112 | (B) BRAF Class 2 |
| PRKCH | 14q23.1 | 0 | 0 | 1 | 8.33 | 0 | 0 | 0.0116 | 0.112 | (B) BRAF Class 2 |
| HLA-B | 6p21.33 | 25 | 4.08 | 1 | 0.63 | 1 | 0.63 | 0.0118 | 0.112 | (A) BRAF Class 1 v12 |
| HIF1A | 14q23.2 | 0 | 0 | 1 | 2.33 | 2 | 3.92 | 0.0118 | 0.112 | (C) BRAF Class 3 |
| PRKCA | 17q24.2 | 0 | 0 | 1 | 2.27 | 2 | 4.17 | 0.0122 | 0.114 | (C) BRAF Class 3 |
| NCKAP5 | 2q21.2 | 5 | 5.81 | 3 | 25 | 5 | 25 | 0.0125 | 0.117 | (C) BRAF Class 3 |
| PKHD1 | p12.3-p12. | 1 | 0.85 | 2 | 10.53 | 0 | 0 | 0.0134 | 0.123 | (B) BRAF Class 2 |
| DCLRE1C | 10p13 | 7 | 1.2 | 6 | 4.58 | 1 | 0.68 | 0.0137 | 0.124 | (B) BRAF Class 2 |
| FGA | 4q31.3 | 0 | 0 | 0 | 0 | 1 | 20 | 0.0137 | 0.124 | (C) BRAF Class 3 |
| CD74 | 5q33.1 | 3 | 0.65 | 3 | 3.95 | 0 | 0 | 0.0141 | 0.127 | (B) BRAF Class 2 |
| RSPO2 | 8q23.1 | 12 | 1.35 | 9 | 4.48 | 6 | 2.55 | 0.0146 | 0.13 | (B) BRAF Class 2 |
| FGF6 | 12p13.32 | 10 | 1.41 | 6 | 4.8 | 1 | 0.65 | 0.0147 | 0.13 | (B) BRAF Class 2 |
| MAP3K1 | 5q11.2 | 98 | 3.99 | 24 | 4.34 | 39 | 6.76 | 0.0152 | 0.134 | (C) BRAF Class 3 |
| H3F3A | 1q42.12 | 13 | 0.52 | 7 | 1.28 | 0 | 0 | 0.0156 | 0.136 | (B) BRAF Class 2 |

| | | | | | | | | | | |
|---|---|---|---|---|---|---|---|---|---|---|
| SETBP1 | 18q12.3 | 103 | 7.03 | 29 | 10.32 | 36 | 11.21 | 0.0158 | 0.136 | (C) BRAF Class 3 |
| TERC | 3q26.2 | 0 | 0 | 1 | 0.74 | 2 | 1.23 | 0.0158 | 0.136 | (C) BRAF Class 3 |
| SMAD2 | 18q21.1 | 42 | 1.71 | 5 | 0.9 | 18 | 3.12 | 0.016 | 0.137 | (C) BRAF Class 3 |
| SOX17 | 8q11.23 | 55 | 4.65 | 6 | 2.21 | 19 | 7.42 | 0.0178 | 0.15 | (C) BRAF Class 3 |
| KEL | 7q34 | 30 | 5.73 | 12 | 13.33 | 11 | 10.19 | 0.0179 | 0.15 | (B) BRAF Class 2 |
| HIST1H3C | 6p22.2 | 8 | 0.46 | 3 | 0.76 | 7 | 1.79 | 0.018 | 0.15 | (C) BRAF Class 3 |
| BEND2 | Xp22.13 | 1 | 1.09 | 2 | 13.33 | 0 | 0 | 0.018 | 0.15 | (B) BRAF Class 2 |
| PRDM14 | 8q13.3 | 14 | 2.64 | 2 | 1.44 | 9 | 6.98 | 0.0182 | 0.15 | (C) BRAF Class 3 |
| ZFHX4 | 8q21.13 | 18 | 11.54 | 6 | 24 | 9 | 30 | 0.0183 | 0.15 | (C) BRAF Class 3 |
| B2M | 15q21.1 | 87 | 4.01 | 10 | 2.06 | 10 | 2.03 | 0.0188 | 0.153 | (A) BRAF Class 1 v12 |
| ESR1 | q25.1-q25. | 55 | 2.08 | 12 | 1.9 | 24 | 3.91 | 0.0192 | 0.155 | (C) BRAF Class 3 |
| ADGRL3 | 4q13.1 | 0 | 0 | 1 | 12.5 | 0 | 0 | 0.0196 | 0.157 | (B) BRAF Class 2 |
| FOXO1 | 13q14.11 | 28 | 2.06 | 4 | 1.33 | 13 | 4.48 | 0.0209 | 0.167 | (C) BRAF Class 3 |
| PRSS1 | 7q34 | 20 | 2.75 | 8 | 4.91 | 13 | 6.88 | 0.0217 | 0.172 | (C) BRAF Class 3 |
| ZRSR2 | Xp22.2 | 41 | 1.82 | 10 | 2.02 | 1 | 0.2 | 0.0221 | 0.173 | (B) BRAF Class 2 |
| ZNF521 | 18q11.2 | 1 | 0.82 | 0 | 0 | 2 | 9.09 | 0.0225 | 0.173 | (C) BRAF Class 3 |
| RBM10 | Xp11.3 | 81 | 3.92 | 19 | 4.01 | 33 | 6.71 | 0.0227 | 0.173 | (C) BRAF Class 3 |
| TCF12 | 15q21.3 | 1 | 0.5 | 2 | 5.56 | 0 | 0 | 0.0227 | 0.173 | (B) BRAF Class 2 |
| NFKBIZ | 3q12.3 | 14 | 1.53 | 2 | 1.02 | 9 | 4.19 | 0.0228 | 0.173 | (C) BRAF Class 3 |
| THBS1 | 15q14 | 1 | 0.85 | 0 | 0 | 2 | 9.52 | 0.0228 | 0.173 | (C) BRAF Class 3 |
| CCNQ | Xq28 | 6 | 1.13 | 2 | 1.44 | 6 | 4.65 | 0.0229 | 0.173 | (C) BRAF Class 3 |
| ADAMTS2( | 12q12 | 3 | 2.56 | 0 | 0 | 3 | 14.29 | 0.0234 | 0.175 | (C) BRAF Class 3 |
| CIITA | 16p13.13 | 48 | 4.83 | 20 | 9.35 | 17 | 7.33 | 0.0238 | 0.177 | (B) BRAF Class 2 |
| ABCB11 | 2q31.1 | 33 | 5.06 | 9 | 6.25 | 17 | 10.9 | 0.025 | 0.183 | (C) BRAF Class 3 |
| FZR1 | 19p13.3 | 0 | 0 | 1 | 5.26 | 0 | 0 | 0.0259 | 0.183 | (B) BRAF Class 2 |
| LTF | 3p21.31 | 0 | 0 | 1 | 5.26 | 0 | 0 | 0.0259 | 0.183 | (B) BRAF Class 2 |
| SOX11 | 2p25.2 | 0 | 0 | 1 | 5.26 | 0 | 0 | 0.0259 | 0.183 | (B) BRAF Class 2 |
| IKZF1 | 7p12.2 | 121 | 4.93 | 26 | 4.72 | 44 | 7.65 | 0.026 | 0.183 | (C) BRAF Class 3 |
| PTPRS | 19p13.3 | 110 | 9.3 | 20 | 7.35 | 11 | 4.3 | 0.026 | 0.183 | (A) BRAF Class 1 v12 |
| MECOM | 3q26.2 | 70 | 7.35 | 13 | 6.28 | 28 | 12.39 | 0.026 | 0.183 | (C) BRAF Class 3 |
| PCM1 | 8p22 | 0 | 0 | 1 | 25 | 0 | 0 | 0.027 | 0.183 | (B) BRAF Class 2 |
| POLD1 | 19q13.33 | 131 | 6.36 | 17 | 3.61 | 21 | 4.35 | 0.0271 | 0.183 | (A) BRAF Class 1 v12 |
| CAD | 2p23.3 | 0 | 0 | 1 | 7.14 | 0 | 0 | 0.0272 | 0.183 | (B) BRAF Class 2 |
| CPS1 | 2q34 | 0 | 0 | 1 | 7.14 | 0 | 0 | 0.0272 | 0.183 | (B) BRAF Class 2 |
| INPP5D | 2q37.1 | 0 | 0 | 1 | 7.14 | 0 | 0 | 0.0272 | 0.183 | (B) BRAF Class 2 |
| MEF2C | 5q14.3 | 0 | 0 | 1 | 7.14 | 0 | 0 | 0.0272 | 0.183 | (B) BRAF Class 2 |
| MKI67 | 10q26.2 | 0 | 0 | 1 | 7.14 | 0 | 0 | 0.0272 | 0.183 | (B) BRAF Class 2 |
| PTPN6 | 12p13.31 | 0 | 0 | 1 | 7.14 | 0 | 0 | 0.0272 | 0.183 | (B) BRAF Class 2 |
| TNFRSF11/ | 18q21.33 | 0 | 0 | 1 | 7.14 | 0 | 0 | 0.0272 | 0.183 | (B) BRAF Class 2 |
| WDR90 | 16p13.3 | 0 | 0 | 1 | 7.14 | 0 | 0 | 0.0272 | 0.183 | (B) BRAF Class 2 |
| HIST1H3B | 6p22.2 | 19 | 0.91 | 11 | 2.37 | 5 | 1.07 | 0.0283 | 0.189 | (B) BRAF Class 2 |
| YES1 | 18p11.32 | 21 | 1.78 | 4 | 1.47 | 11 | 4.3 | 0.0283 | 0.189 | (C) BRAF Class 3 |
| PLCG1 | 20q12 | 0 | 0 | 1 | 5 | 0 | 0 | 0.0288 | 0.191 | (B) BRAF Class 2 |
| GRIN2A | 16p13.2 | 159 | 10.37 | 40 | 11.73 | 53 | 15.36 | 0.0295 | 0.193 | (C) BRAF Class 3 |
| ICK | 6p12.1 | 0 | 0 | 1 | 5 | 0 | 0 | 0.0295 | 0.193 | (B) BRAF Class 2 |

| | | | | | | | | | | |
|---|---|---|---|---|---|---|---|---|---|---|
| IRF4 | 6p25.3 | 21 | 1.34 | 12 | 3.35 | 7 | 1.93 | 0.0305 | 0.198 | (B) BRAF Class 2 |
| FANCE | 6p21.31 | 19 | 1.2 | 2 | 0.66 | 10 | 2.82 | 0.0312 | 0.202 | (C) BRAF Class 3 |
| JARID2 | 6p22.3 | 0 | 0 | 1 | 6.67 | 0 | 0 | 0.0313 | 0.202 | (B) BRAF Class 2 |
| TRIP11 | 14q32.12 | 0 | 0 | 1 | 4.76 | 0 | 0 | 0.0318 | 0.203 | (B) BRAF Class 2 |
| BIRC2 | 11q22.2 | 0 | 0 | 0 | 0 | 1 | 3.7 | 0.032 | 0.204 | (C) BRAF Class 3 |
| RRAS | 19q13.33 | 3 | 0.45 | 1 | 0.62 | 4 | 2.52 | 0.0322 | 0.204 | (C) BRAF Class 3 |
| HTRA1 | 10q26.13 | 0 | 0 | 1 | 6.67 | 0 | 0 | 0.0324 | 0.204 | (B) BRAF Class 2 |
| RASA2 | 3q23 | 2 | 0.91 | 1 | 2.78 | 3 | 6.98 | 0.0331 | 0.207 | (C) BRAF Class 3 |
| TFEB | 6p21.1 | 1 | 0.45 | 0 | 0 | 2 | 4.55 | 0.0335 | 0.208 | (C) BRAF Class 3 |
| CCND2 | 12p13.32 | 25 | 0.99 | 5 | 0.89 | 13 | 2.23 | 0.0344 | 0.213 | (C) BRAF Class 3 |
| CSDE1 | 1p13.2 | 10 | 1.19 | 6 | 3.39 | 6 | 3.39 | 0.0355 | 0.217 | (C) BRAF Class 3 |
| IPMK | 10q21.1 | 1 | 0.64 | 2 | 8 | 1 | 3.33 | 0.0357 | 0.217 | (B) BRAF Class 2 |
| BRCA1 | 17q21.31 | 101 | 3.98 | 29 | 4.74 | 38 | 6.39 | 0.036 | 0.217 | (C) BRAF Class 3 |
| NF2 | 22q12.2 | 58 | 2.25 | 18 | 3.17 | 24 | 4.05 | 0.0366 | 0.217 | (C) BRAF Class 3 |
| MN1 | 22q12.1 | 0 | 0 | 1 | 2.94 | 0 | 0 | 0.0366 | 0.217 | (B) BRAF Class 2 |
| HIST1H1E | 6p22.2 | 0 | 0 | 1 | 4.76 | 0 | 0 | 0.0375 | 0.217 | (B) BRAF Class 2 |
| DNM2 | 19p13.2 | 0 | 0 | 1 | 5.56 | 0 | 0 | 0.0378 | 0.217 | (B) BRAF Class 2 |
| MAML2 | 11q21 | 1 | 0.39 | 0 | 0 | 2 | 3.85 | 0.038 | 0.217 | (C) BRAF Class 3 |
| BCL3 | 19q13.32 | 0 | 0 | 0 | 0 | 1 | 4.55 | 0.038 | 0.217 | (C) BRAF Class 3 |
| HOOK3 | 8p11.21 | 0 | 0 | 0 | 0 | 1 | 4.55 | 0.038 | 0.217 | (C) BRAF Class 3 |
| LPP | 3q27.3-q2ε | 0 | 0 | 0 | 0 | 1 | 4.55 | 0.038 | 0.217 | (C) BRAF Class 3 |
| NUMA1 | 11q13.4 | 0 | 0 | 0 | 0 | 1 | 4.55 | 0.038 | 0.217 | (C) BRAF Class 3 |
| RALGDS | ɥ34.13-q34 | 0 | 0 | 0 | 0 | 1 | 4.55 | 0.038 | 0.217 | (C) BRAF Class 3 |
| G6PD | Xq28 | 0 | 0 | 0 | 0 | 1 | 4.76 | 0.0384 | 0.217 | (C) BRAF Class 3 |
| MAGEA1 | Xq28 | 0 | 0 | 0 | 0 | 1 | 4.76 | 0.0384 | 0.217 | (C) BRAF Class 3 |
| MAGI1 | 3p14.1 | 0 | 0 | 0 | 0 | 1 | 4.76 | 0.0384 | 0.217 | (C) BRAF Class 3 |
| MMP2 | 16q12.2 | 0 | 0 | 0 | 0 | 1 | 4.76 | 0.0384 | 0.217 | (C) BRAF Class 3 |
| USP9X | Xp11.4 | 0 | 0 | 0 | 0 | 1 | 4.76 | 0.0384 | 0.217 | (C) BRAF Class 3 |
| RARA | 17q21.2 | 43 | 1.75 | 18 | 3.25 | 8 | 1.39 | 0.039 | 0.219 | (B) BRAF Class 2 |
| DUSP4 | 8p12 | 8 | 1.2 | 6 | 3.7 | 1 | 0.63 | 0.0401 | 0.224 | (B) BRAF Class 2 |
| KLF6 | 10p15.2 | 0 | 0 | 0 | 0 | 1 | 4.35 | 0.0402 | 0.224 | (C) BRAF Class 3 |
| MAP2K1 | 15q22.31 | 71 | 2.63 | 29 | 4.5 | 21 | 3.36 | 0.0413 | 0.228 | (B) BRAF Class 2 |
| NOTCH3 | 19p13.12 | 219 | 9.99 | 38 | 7.74 | 64 | 12.52 | 0.0413 | 0.228 | (C) BRAF Class 3 |
| SPRED1 | 15q14 | 11 | 1.66 | 8 | 4.94 | 5 | 3.14 | 0.0429 | 0.232 | (B) BRAF Class 2 |
| NUP98 | 11p15.4 | 0 | 0 | 0 | 0 | 1 | 4 | 0.0434 | 0.232 | (C) BRAF Class 3 |
| ZMYM2 | 13q12.11 | 0 | 0 | 0 | 0 | 1 | 4.55 | 0.0436 | 0.232 | (C) BRAF Class 3 |
| FGF8 | 10q24.32 | 2 | 0.6 | 2 | 5 | 1 | 2.08 | 0.0437 | 0.232 | (B) BRAF Class 2 |
| NTRK2 | 9q21.33 | 53 | 2.09 | 19 | 3.37 | 21 | 3.58 | 0.0438 | 0.232 | (C) BRAF Class 3 |
| BOD1L1 | 4p15.33 | 0 | 0 | 1 | 12.5 | 1 | 12.5 | 0.0452 | 0.232 | (C) BRAF Class 3 |
| CDKN1A | 6p21.2 | 15 | 0.63 | 7 | 1.3 | 9 | 1.6 | 0.0453 | 0.232 | (C) BRAF Class 3 |
| PAX8 | 2q14.1 | 8 | 1.39 | 1 | 1.19 | 5 | 4.9 | 0.0462 | 0.232 | (C) BRAF Class 3 |
| AURKB | 17p13.1 | 24 | 0.98 | 10 | 1.81 | 2 | 0.35 | 0.0465 | 0.232 | (B) BRAF Class 2 |
| DDX3X | Xp11.4 | 3 | 1.2 | 0 | 0 | 3 | 5.77 | 0.0465 | 0.232 | (C) BRAF Class 3 |
| PAK5 | 20p12.2 | 80 | 6.21 | 26 | 8.72 | 29 | 9.9 | 0.0468 | 0.232 | (C) BRAF Class 3 |
| LMO3 | 12p12.3 | 0 | 0 | 0 | 0 | 1 | 1.45 | 0.047 | 0.232 | (C) BRAF Class 3 |

| | | | | | | | | | | |
|---|---|---|---|---|---|---|---|---|---|---|
| GLI1 | 12q13.3 | 114 | 4.96 | 16 | 3.17 | 34 | 6.5 | 0.0473 | 0.232 | (C) BRAF Class 3 |
| BARD1 | 2q35 | 38 | 1.79 | 10 | 2.05 | 18 | 3.54 | 0.0473 | 0.232 | (C) BRAF Class 3 |
| DAXX | 6p21.32 | 41 | 1.98 | 8 | 1.68 | 18 | 3.69 | 0.0486 | 0.232 | (C) BRAF Class 3 |
| SESN2 | 1p35.3 | 5 | 0.94 | 2 | 1.44 | 5 | 3.88 | 0.0491 | 0.232 | (C) BRAF Class 3 |
| AXL | 19q13.2 | 97 | 3.82 | 28 | 4.97 | 35 | 5.97 | 0.0505 | 0.232 | (C) BRAF Class 3 |
| FANCF | 11p14.3 | 10 | 0.63 | 6 | 1.97 | 5 | 1.41 | 0.0511 | 0.232 | (B) BRAF Class 2 |
| ITK | 5q33.3 | 12 | 2.01 | 6 | 4.44 | 8 | 5.37 | 0.0519 | 0.232 | (C) BRAF Class 3 |
| PDK1 | 2q31.1 | 8 | 1.13 | 5 | 4 | 2 | 1.3 | 0.0522 | 0.232 | (B) BRAF Class 2 |
| FANCA | 16q24.3 | 90 | 3.54 | 28 | 4.97 | 32 | 5.46 | 0.0526 | 0.232 | (C) BRAF Class 3 |
| E2F3 | 6p22.3 | 10 | 0.85 | 6 | 2.21 | 6 | 2.34 | 0.0529 | 0.232 | (C) BRAF Class 3 |
| PAX5 | 9p13.2 | 53 | 2.13 | 17 | 3.04 | 22 | 3.77 | 0.0537 | 0.232 | (C) BRAF Class 3 |
| ATRX | Xq21.1 | 167 | 6.57 | 38 | 6.75 | 55 | 9.39 | 0.0541 | 0.232 | (C) BRAF Class 3 |
| USP6 | 17p13.2 | 2 | 1.18 | 2 | 6.67 | 2 | 7.69 | 0.0547 | 0.232 | (C) BRAF Class 3 |
| NRG1 | 8p12 | 52 | 5.07 | 13 | 6.74 | 20 | 9.22 | 0.055 | 0.232 | (C) BRAF Class 3 |
| ASH2L | 8p11.23 | 0 | 0 | 1 | 4 | 1 | 3.33 | 0.0553 | 0.232 | (B) BRAF Class 2 |
| AKAP9 | 7q21.2 | 0 | 0 | 1 | 4.76 | 1 | 4.55 | 0.0565 | 0.232 | (B) BRAF Class 2 |
| PBX1 | 1q23.3 | 0 | 0 | 1 | 4.76 | 1 | 4.55 | 0.0565 | 0.232 | (B) BRAF Class 2 |
| CTNNB1 | 3p22.1 | 113 | 3.38 | 32 | 4.27 | 38 | 5.14 | 0.0572 | 0.232 | (C) BRAF Class 3 |
| SETD2 | 3p21.31 | 224 | 8.82 | 40 | 7.1 | 65 | 11.09 | 0.0573 | 0.232 | (C) BRAF Class 3 |
| COL1A1 | 17q21.33 | 4 | 1.56 | 3 | 6.82 | 3 | 5.77 | 0.0592 | 0.232 | (B) BRAF Class 2 |
| ERCC4 | 16p13.12 | 52 | 2.27 | 13 | 2.48 | 4 | 0.74 | 0.0596 | 0.232 | (B) BRAF Class 2 |
| RASA1 | 5q14.3 | 39 | 2.08 | 15 | 3.54 | 16 | 3.7 | 0.0597 | 0.232 | (C) BRAF Class 3 |
| WAS | Xp11.23 | 10 | 1.45 | 2 | 1.32 | 7 | 4.12 | 0.0611 | 0.232 | (C) BRAF Class 3 |
| USP8 | 15q21.2 | 14 | 2.36 | 5 | 3.73 | 9 | 6.16 | 0.0615 | 0.232 | (C) BRAF Class 3 |
| STAG2 | Xq25 | 73 | 2.97 | 22 | 3.98 | 28 | 4.85 | 0.0617 | 0.232 | (C) BRAF Class 3 |
| NCOA2 | 8q13.3 | 6 | 2.34 | 4 | 9.09 | 3 | 5.77 | 0.0625 | 0.232 | (B) BRAF Class 2 |
| BCL2 | 18q21.33 | 13 | 0.53 | 6 | 1.09 | 8 | 1.39 | 0.0628 | 0.232 | (C) BRAF Class 3 |
| CDKN2B | 9p21.3 | 208 | 8.21 | 58 | 10.38 | 63 | 10.79 | 0.0631 | 0.232 | (C) BRAF Class 3 |
| LEF1 | 4q25 | 0 | 0 | 1 | 2.7 | 1 | 2.27 | 0.0631 | 0.232 | (B) BRAF Class 2 |
| RPS6KB2 | 11q13.2 | 21 | 1.78 | 9 | 3.31 | 10 | 3.91 | 0.0632 | 0.232 | (C) BRAF Class 3 |
| ACP2 | .1.2\|11p12 | 0 | 0 | 0 | 0 | 1 | 0.13 | 0.0635 | 0.232 | (C) BRAF Class 3 |
| ARHGEF3{ | 4q24 | 0 | 0 | 0 | 0 | 1 | 0.13 | 0.0635 | 0.232 | (C) BRAF Class 3 |
| C1ORF147 | 1q32.1 | 0 | 0 | 0 | 0 | 1 | 0.13 | 0.0635 | 0.232 | (C) BRAF Class 3 |
| CACNA1C | 12p13.33 | 0 | 0 | 0 | 0 | 1 | 0.13 | 0.0635 | 0.232 | (C) BRAF Class 3 |
| CCHCR1 | 6p21.33 | 0 | 0 | 0 | 0 | 1 | 0.13 | 0.0635 | 0.232 | (C) BRAF Class 3 |
| DAP | 5p15.2 | 0 | 0 | 0 | 0 | 1 | 0.13 | 0.0635 | 0.232 | (C) BRAF Class 3 |
| )OCK8-AS: | 9p24.3 | 0 | 0 | 0 | 0 | 1 | 0.13 | 0.0635 | 0.232 | (C) BRAF Class 3 |
| EIF2S3 | Xp22.11 | 0 | 0 | 0 | 0 | 1 | 0.13 | 0.0635 | 0.232 | (C) BRAF Class 3 |
| ENSA | 1q21.3 | 0 | 0 | 0 | 0 | 1 | 0.13 | 0.0635 | 0.232 | (C) BRAF Class 3 |
| FNDC3B | 3q26.31 | 0 | 0 | 0 | 0 | 1 | 0.13 | 0.0635 | 0.232 | (C) BRAF Class 3 |
| GON4L | 1q22 | 0 | 0 | 0 | 0 | 1 | 0.13 | 0.0635 | 0.232 | (C) BRAF Class 3 |
| GPD1 | 12q13.12 | 0 | 0 | 0 | 0 | 1 | 0.13 | 0.0635 | 0.232 | (C) BRAF Class 3 |
| HIPK2 | 7q34 | 0 | 0 | 0 | 0 | 1 | 0.13 | 0.0635 | 0.232 | (C) BRAF Class 3 |
| IGLJ5 | 22q11.22 | 0 | 0 | 0 | 0 | 1 | 0.13 | 0.0635 | 0.232 | (C) BRAF Class 3 |
| INHBE | 12q13.3 | 0 | 0 | 0 | 0 | 1 | 0.13 | 0.0635 | 0.232 | (C) BRAF Class 3 |

| | | | | | | | | | | |
|---|---|---|---|---|---|---|---|---|---|---|
| LINC0099€ | 7q36.1 | 0 | 0 | 0 | 0 | 1 | 0.13 | 0.0635 | 0.232 | (C) BRAF Class 3 |
| LINC01514 | 10q24.31 | 0 | 0 | 0 | 0 | 1 | 0.13 | 0.0635 | 0.232 | (C) BRAF Class 3 |
| MIR99AHG | 21q21.1 | 0 | 0 | 0 | 0 | 1 | 0.13 | 0.0635 | 0.232 | (C) BRAF Class 3 |
| NSRP1 | 17q11.2 | 0 | 0 | 0 | 0 | 1 | 0.13 | 0.0635 | 0.232 | (C) BRAF Class 3 |
| PAX8-AS1 | 2q14.1 | 0 | 0 | 0 | 0 | 1 | 0.13 | 0.0635 | 0.232 | (C) BRAF Class 3 |
| PCMT1 | 6q25.1 | 0 | 0 | 0 | 0 | 1 | 0.13 | 0.0635 | 0.232 | (C) BRAF Class 3 |
| PDE4D | q11.2-q12. | 0 | 0 | 0 | 0 | 1 | 0.13 | 0.0635 | 0.232 | (C) BRAF Class 3 |
| PTTG1IP | 21q22.3 | 0 | 0 | 0 | 0 | 1 | 0.13 | 0.0635 | 0.232 | (C) BRAF Class 3 |
| RAB11FIP4 | 17q11.2 | 0 | 0 | 0 | 0 | 1 | 0.13 | 0.0635 | 0.232 | (C) BRAF Class 3 |
| RLF | 1p34.2 | 0 | 0 | 0 | 0 | 1 | 0.13 | 0.0635 | 0.232 | (C) BRAF Class 3 |
| SDHAP3 | 5p15.33 | 0 | 0 | 0 | 0 | 1 | 0.13 | 0.0635 | 0.232 | (C) BRAF Class 3 |
| SEPTIN14 | 7p11.2 | 0 | 0 | 0 | 0 | 1 | 0.13 | 0.0635 | 0.232 | (C) BRAF Class 3 |
| SNTG1 | 8q11.21 | 0 | 0 | 0 | 0 | 1 | 0.13 | 0.0635 | 0.232 | (C) BRAF Class 3 |
| TANC2 | 'q23.2-q23 | 0 | 0 | 0 | 0 | 1 | 0.13 | 0.0635 | 0.232 | (C) BRAF Class 3 |
| TMIE | 3p21.31 | 0 | 0 | 0 | 0 | 1 | 0.13 | 0.0635 | 0.232 | (C) BRAF Class 3 |
| TRAJ22 | 14q11.2 | 0 | 0 | 0 | 0 | 1 | 0.13 | 0.0635 | 0.232 | (C) BRAF Class 3 |
| TTF1 | 9q34.13 | 0 | 0 | 0 | 0 | 1 | 0.13 | 0.0635 | 0.232 | (C) BRAF Class 3 |
| TTF2 | 1p13.1 | 0 | 0 | 0 | 0 | 1 | 0.13 | 0.0635 | 0.232 | (C) BRAF Class 3 |
| UAP1L1 | 9q34.3 | 0 | 0 | 0 | 0 | 1 | 0.13 | 0.0635 | 0.232 | (C) BRAF Class 3 |
| ZFP30 | 19q13.12 | 0 | 0 | 0 | 0 | 1 | 0.13 | 0.0635 | 0.232 | (C) BRAF Class 3 |
| ZNF135 | 19q13.43 | 0 | 0 | 0 | 0 | 1 | 0.13 | 0.0635 | 0.232 | (C) BRAF Class 3 |
| ZNF284 | 19q13.31 | 0 | 0 | 0 | 0 | 1 | 0.13 | 0.0635 | 0.232 | (C) BRAF Class 3 |
| MTAP | 9p21.3 | 39 | 6.27 | 12 | 7.95 | 19 | 11.66 | 0.0648 | 0.232 | (C) BRAF Class 3 |
| ADGRV1 | 5q14.3 | 0 | 0 | 1 | 0.13 | 0 | 0 | 0.0652 | 0.232 | (B) BRAF Class 2 |
| ALG10 | 12p11.1 | 0 | 0 | 1 | 0.13 | 0 | 0 | 0.0652 | 0.232 | (B) BRAF Class 2 |
| C5ORF22 | 5p13.3 | 0 | 0 | 1 | 0.13 | 0 | 0 | 0.0652 | 0.232 | (B) BRAF Class 2 |
| CALB1 | 8q21.3 | 0 | 0 | 1 | 0.13 | 0 | 0 | 0.0652 | 0.232 | (B) BRAF Class 2 |
| CAMKMT | 2p21 | 0 | 0 | 1 | 0.13 | 0 | 0 | 0.0652 | 0.232 | (B) BRAF Class 2 |
| CRLF3 | 17q11.2 | 0 | 0 | 1 | 0.13 | 0 | 0 | 0.0652 | 0.232 | (B) BRAF Class 2 |
| FAM124B | 2q36.2 | 0 | 0 | 1 | 0.13 | 0 | 0 | 0.0652 | 0.232 | (B) BRAF Class 2 |
| HEXIM2 | 17q21.31 | 0 | 0 | 1 | 0.13 | 0 | 0 | 0.0652 | 0.232 | (B) BRAF Class 2 |
| IGHD2-2 | 14q32.33 | 0 | 0 | 1 | 0.13 | 0 | 0 | 0.0652 | 0.232 | (B) BRAF Class 2 |
| IRX3 | 16q12.2 | 0 | 0 | 1 | 0.13 | 0 | 0 | 0.0652 | 0.232 | (B) BRAF Class 2 |
| KCNAB1 | 3q25.31 | 0 | 0 | 1 | 0.13 | 0 | 0 | 0.0652 | 0.232 | (B) BRAF Class 2 |
| KCNJ1 | 11q24.3 | 0 | 0 | 1 | 0.13 | 0 | 0 | 0.0652 | 0.232 | (B) BRAF Class 2 |
| KYNU | 2q22.2 | 0 | 0 | 1 | 0.13 | 0 | 0 | 0.0652 | 0.232 | (B) BRAF Class 2 |
| LAMC3 | 9q34.12 | 0 | 0 | 1 | 0.13 | 0 | 0 | 0.0652 | 0.232 | (B) BRAF Class 2 |
| LGALS17A | 19q13.2 | 0 | 0 | 1 | 0.13 | 0 | 0 | 0.0652 | 0.232 | (B) BRAF Class 2 |
| LILRB1 | 19q13.42 | 0 | 0 | 1 | 0.13 | 0 | 0 | 0.0652 | 0.232 | (B) BRAF Class 2 |
| LINC00484 | 9q22.2 | 0 | 0 | 1 | 0.13 | 0 | 0 | 0.0652 | 0.232 | (B) BRAF Class 2 |
| LINC02544 | 6q27 | 0 | 0 | 1 | 0.13 | 0 | 0 | 0.0652 | 0.232 | (B) BRAF Class 2 |
| MTMR3 | 22q12.2 | 0 | 0 | 1 | 0.13 | 0 | 0 | 0.0652 | 0.232 | (B) BRAF Class 2 |
| NBPF1 | 1p36.13 | 0 | 0 | 1 | 0.13 | 0 | 0 | 0.0652 | 0.232 | (B) BRAF Class 2 |
| NUFIP1 | 13q14.12 | 0 | 0 | 1 | 0.13 | 0 | 0 | 0.0652 | 0.232 | (B) BRAF Class 2 |
| PAXIP1-AS: | 7q36.2 | 0 | 0 | 1 | 0.13 | 0 | 0 | 0.0652 | 0.232 | (B) BRAF Class 2 |

| | | | | | | | | | | |
|---|---|---|---|---|---|---|---|---|---|---|
| PEX5 | 12p13.31 | 0 | 0 | 1 | 0.13 | 0 | 0 | 0.0652 | 0.232 | (B) BRAF Class 2 |
| PRAG1 | 8p23.1 | 0 | 0 | 1 | 0.13 | 0 | 0 | 0.0652 | 0.232 | (B) BRAF Class 2 |
| PXMP2 | 12q24.33 | 0 | 0 | 1 | 0.13 | 0 | 0 | 0.0652 | 0.232 | (B) BRAF Class 2 |
| RERE | 1p36.23 | 0 | 0 | 1 | 0.13 | 0 | 0 | 0.0652 | 0.232 | (B) BRAF Class 2 |
| ROCK1 | 18q11.1 | 0 | 0 | 1 | 0.13 | 0 | 0 | 0.0652 | 0.232 | (B) BRAF Class 2 |
| RUNX3 | 1p36.11 | 0 | 0 | 1 | 0.13 | 0 | 0 | 0.0652 | 0.232 | (B) BRAF Class 2 |
| SBF1P1 | 8q12.1 | 0 | 0 | 1 | 0.13 | 0 | 0 | 0.0652 | 0.232 | (B) BRAF Class 2 |
| EC24B-AS | 4q25 | 0 | 0 | 1 | 0.13 | 0 | 0 | 0.0652 | 0.232 | (B) BRAF Class 2 |
| SELENOI | 2p23.3 | 0 | 0 | 1 | 0.13 | 0 | 0 | 0.0652 | 0.232 | (B) BRAF Class 2 |
| ST7 | 7q31.2 | 0 | 0 | 1 | 0.13 | 0 | 0 | 0.0652 | 0.232 | (B) BRAF Class 2 |
| TECRL | 4q13.1 | 0 | 0 | 1 | 0.13 | 0 | 0 | 0.0652 | 0.232 | (B) BRAF Class 2 |
| TMEM132( | q24.32-q24 | 0 | 0 | 1 | 0.13 | 0 | 0 | 0.0652 | 0.232 | (B) BRAF Class 2 |
| TTC28 | 22q12.1 | 0 | 0 | 1 | 0.13 | 0 | 0 | 0.0652 | 0.232 | (B) BRAF Class 2 |
| VPREB1 | 22q11.22 | 0 | 0 | 1 | 0.13 | 0 | 0 | 0.0652 | 0.232 | (B) BRAF Class 2 |
| VRK2 | 2p16.1 | 0 | 0 | 1 | 0.13 | 0 | 0 | 0.0652 | 0.232 | (B) BRAF Class 2 |
| CNOT3 | 19q13.42 | 2 | 1.18 | 2 | 6.9 | 0 | 0 | 0.0653 | 0.232 | (B) BRAF Class 2 |
| IDH1 | 2q34 | 80 | 2.39 | 26 | 3.47 | 12 | 1.62 | 0.0655 | 0.232 | (B) BRAF Class 2 |
| CRKL | 22q11.21 | 18 | 0.73 | 7 | 1.27 | 10 | 1.73 | 0.0661 | 0.233 | (C) BRAF Class 3 |
| CREB1 | 2q33.3 | 0 | 0 | 1 | 2.27 | 1 | 1.92 | 0.0667 | 0.235 | (B) BRAF Class 2 |
| RSPO3 | 6q22.33 | 20 | 2.57 | 9 | 5.45 | 3 | 1.59 | 0.0672 | 0.236 | (B) BRAF Class 2 |
| EED | 11q14.2 | 14 | 0.88 | 3 | 0.83 | 8 | 2.25 | 0.0678 | 0.237 | (C) BRAF Class 3 |
| INSIG1 | 7q36.3 | 0 | 0 | 0 | 0 | 1 | 1.72 | 0.068 | 0.237 | (C) BRAF Class 3 |
| PAX7 | 1p36.13 | 15 | 2.65 | 2 | 2.44 | 7 | 7 | 0.0682 | 0.237 | (C) BRAF Class 3 |
| RB1 | 13q14.2 | 104 | 3.23 | 27 | 4.03 | 35 | 4.96 | 0.0683 | 0.237 | (C) BRAF Class 3 |
| SMARCA2 | 9p24.3 | 2 | 1.2 | 2 | 7.14 | 2 | 6.67 | 0.0688 | 0.237 | (B) BRAF Class 2 |
| CDH4 | 20q13.33 | 34 | 5.84 | 7 | 5.34 | 16 | 10.96 | 0.0689 | 0.237 | (C) BRAF Class 3 |
| NEIL3 | 4q34.3 | 23 | 3.95 | 0 | 0 | 5 | 3.42 | 0.0703 | 0.241 | (A) BRAF Class 1 v12 |
| ZNF708 | 19p12 | 3 | 0.85 | 3 | 4.41 | 2 | 2.9 | 0.0705 | 0.241 | (B) BRAF Class 2 |
| RIT1 | 1q22 | 26 | 1.33 | 3 | 0.68 | 11 | 2.44 | 0.0705 | 0.241 | (C) BRAF Class 3 |
| MAGI2 | 7q21.11 | 38 | 7.69 | 11 | 15.07 | 11 | 12.09 | 0.0707 | 0.241 | (B) BRAF Class 2 |
| BAP1 | 3p21.1 | 62 | 2.44 | 22 | 3.9 | 11 | 1.88 | 0.0717 | 0.243 | (B) BRAF Class 2 |
| PDGFRB | 5q32 | 92 | 3.62 | 32 | 5.68 | 26 | 4.44 | 0.0718 | 0.243 | (B) BRAF Class 2 |
| AKT1 | 14q32.33 | 109 | 3.26 | 17 | 2.27 | 14 | 1.89 | 0.0723 | 0.244 | (A) BRAF Class 1 v12 |
| RAD54B | 8q22.1 | 17 | 2.92 | 9 | 6.87 | 4 | 2.74 | 0.0727 | 0.245 | (B) BRAF Class 2 |
| SOX10 | 22q13.1 | 13 | 1.92 | 6 | 5.56 | 4 | 2.92 | 0.0744 | 0.249 | (B) BRAF Class 2 |
| TLR4 | 9q33.1 | 13 | 3.08 | 3 | 4 | 8 | 8.08 | 0.0747 | 0.249 | (C) BRAF Class 3 |
| LATS1 | 6q25.1 | 24 | 2.03 | 12 | 4.41 | 7 | 2.73 | 0.0748 | 0.249 | (B) BRAF Class 2 |
| IL7R | 5p13.2 | 98 | 4.67 | 22 | 4.68 | 35 | 7.13 | 0.0748 | 0.249 | (C) BRAF Class 3 |
| MSH6 | 2p16.3 | 140 | 5.44 | 22 | 3.88 | 21 | 3.55 | 0.0749 | 0.249 | (A) BRAF Class 1 v12 |
| PTP4A1 | 6q12 | 1 | 0.19 | 2 | 1.44 | 0 | 0 | 0.0752 | 0.249 | (B) BRAF Class 2 |
| STAT6 | 12q13.3 | 17 | 1.51 | 8 | 3.4 | 8 | 3.07 | 0.0773 | 0.255 | (B) BRAF Class 2 |
| ELF3 | 1q32.1 | 10 | 1.51 | 7 | 4.32 | 4 | 2.52 | 0.0789 | 0.259 | (B) BRAF Class 2 |
| H3F3C | 12p11.21 | 9 | 0.76 | 3 | 1.1 | 6 | 2.34 | 0.0791 | 0.259 | (C) BRAF Class 3 |
| EP400 | 12q24.33 | 5 | 4.17 | 0 | 0 | 3 | 14.29 | 0.0793 | 0.259 | (C) BRAF Class 3 |
| CTCF | 16q22.1 | 51 | 2.4 | 9 | 1.85 | 20 | 3.94 | 0.0798 | 0.26 | (C) BRAF Class 3 |

| | | | | | | | | | | |
|---|---|---|---|---|---|---|---|---|---|---|
| PML | 15q24.1 | 28 | 4.05 | 5 | 3.29 | 1 | 0.59 | 0.0801 | 0.261 | (A) BRAF Class 1 v12 |
| RUNX1 | 21q22.12 | 68 | 2.73 | 21 | 3.85 | 25 | 4.35 | 0.0804 | 0.261 | (C) BRAF Class 3 |
| TEX12 | 11q23.1 | 0 | 0 | 0 | 0 | 1 | 1.08 | 0.0813 | 0.263 | (C) BRAF Class 3 |
| FGF14 | 13q33.1 | 13 | 1.84 | 5 | 4 | 7 | 4.55 | 0.0817 | 0.263 | (C) BRAF Class 3 |
| PPM1D | 17q23.2 | 45 | 2.48 | 9 | 2.22 | 18 | 4.38 | 0.0818 | 0.263 | (C) BRAF Class 3 |
| AFF1 | q21.3-q22. | 2 | 1.64 | 0 | 0 | 2 | 9.09 | 0.0832 | 0.267 | (C) BRAF Class 3 |
| PTPRK | 6q22.33 | 19 | 12.18 | 7 | 28 | 3 | 10 | 0.0836 | 0.268 | (B) BRAF Class 2 |
| SLIT2 | 4p15.31 | 52 | 8.28 | 12 | 12.5 | 17 | 14.05 | 0.0838 | 0.268 | (C) BRAF Class 3 |
| TRIM28 | 19q13.43 | 0 | 0 | 0 | 0 | 1 | 5 | 0.0845 | 0.269 | (C) BRAF Class 3 |
| PIK3C2G | 12p12.3 | 81 | 6.15 | 22 | 6.98 | 30 | 9.68 | 0.0854 | 0.271 | (C) BRAF Class 3 |
| XRCC1 | 19q13.31 | 11 | 1.71 | 2 | 1.41 | 7 | 4.4 | 0.0869 | 0.274 | (C) BRAF Class 3 |
| ATP1B2 | 17p13.1 | 0 | 0 | 1 | 10 | 0 | 0 | 0.087 | 0.274 | (B) BRAF Class 2 |
| CDH12 | 5p14.3 | 0 | 0 | 1 | 10 | 0 | 0 | 0.087 | 0.274 | (B) BRAF Class 2 |
| SRY | Yp11.2 | 0 | 0 | 0 | 0 | 1 | 0.66 | 0.0872 | 0.274 | (C) BRAF Class 3 |
| SMC3 | 10q25.2 | 27 | 1.85 | 8 | 2.87 | 12 | 3.79 | 0.0881 | 0.275 | (C) BRAF Class 3 |
| XRCC4 | 5q14.2 | 10 | 1.72 | 3 | 2.29 | 7 | 4.79 | 0.0881 | 0.275 | (C) BRAF Class 3 |
| VEGFA | 6p21.1 | 21 | 1.04 | 6 | 1.3 | 11 | 2.3 | 0.0886 | 0.276 | (C) BRAF Class 3 |
| NSD3 | 8p11.23 | 30 | 2.67 | 14 | 4.88 | 13 | 4.44 | 0.0921 | 0.285 | (B) BRAF Class 2 |
| CD70 | 19p13.3 | 0 | 0 | 0 | 0 | 1 | 3.23 | 0.0921 | 0.285 | (C) BRAF Class 3 |
| NR0B1 | Xp21.2 | 11 | 1.89 | 2 | 1.53 | 7 | 4.79 | 0.0922 | 0.285 | (C) BRAF Class 3 |
| PARP3 | 3p21.2 | 0 | 0 | 1 | 2.27 | 0 | 0 | 0.0932 | 0.287 | (B) BRAF Class 2 |
| XPC | 3p25.1 | 22 | 2.15 | 5 | 2.29 | 11 | 4.6 | 0.0943 | 0.29 | (C) BRAF Class 3 |
| NPRL3 | 16p13.3 | 1 | 0.38 | 1 | 2.08 | 2 | 3.45 | 0.0961 | 0.294 | (C) BRAF Class 3 |
| ADGRB3 | 6q12-q13 | 3 | 5.56 | 0 | 0 | 2 | 25 | 0.097 | 0.296 | (C) BRAF Class 3 |
| NKX3-1 | 8p21.2 | 17 | 0.98 | 9 | 2.24 | 4 | 1 | 0.098 | 0.299 | (B) BRAF Class 2 |
| EPHA5 | q13.1-q13. | 90 | 4.83 | 30 | 7.37 | 25 | 6.04 | 0.0987 | 0.3 | (B) BRAF Class 2 |
| PRKDC | 8q11.21 | 169 | 10.73 | 41 | 13.53 | 50 | 14.2 | 0.102 | 0.308 | (C) BRAF Class 3 |
| KRIT1 | 7q21.2 | 1 | 1.09 | 0 | 0 | 1 | 10 | 0.102 | 0.308 | (C) BRAF Class 3 |
| BCL11B | 14q32.2 | 31 | 4.46 | 13 | 8.44 | 7 | 4.09 | 0.102 | 0.308 | (B) BRAF Class 2 |
| FGF19 | 11q13.3 | 24 | 1.46 | 11 | 2.99 | 5 | 1.35 | 0.103 | 0.31 | (B) BRAF Class 2 |
| CUX1 | 7q22.1 | 117 | 8.21 | 17 | 6.27 | 34 | 11.07 | 0.104 | 0.313 | (C) BRAF Class 3 |
| MAP3K4 | 6q26 | 30 | 9.01 | 1 | 2.5 | 1 | 2.08 | 0.105 | 0.315 | (A) BRAF Class 1 v12 |
| AURKA | 20q13.2 | 26 | 1.06 | 12 | 2.17 | 8 | 1.39 | 0.106 | 0.317 | (B) BRAF Class 2 |
| PIK3R1 | 5q13.1 | 67 | 2.6 | 17 | 2.99 | 25 | 4.22 | 0.107 | 0.317 | (C) BRAF Class 3 |
| NTRK1 | 1q23.1 | 100 | 3.85 | 31 | 5.36 | 32 | 5.35 | 0.107 | 0.317 | (B) BRAF Class 2 |
| TOP2A | 17q21.2 | 32 | 5.1 | 1 | 1.04 | 3 | 2.48 | 0.108 | 0.32 | (A) BRAF Class 1 v12 |
| SESN3 | 11q21 | 8 | 1.51 | 1 | 0.72 | 5 | 3.88 | 0.11 | 0.324 | (C) BRAF Class 3 |
| PIK3C2B | 1q32.1 | 94 | 6.5 | 16 | 5.5 | 31 | 9.37 | 0.11 | 0.324 | (C) BRAF Class 3 |
| ZNRF3 | 22q12.1 | 41 | 5.87 | 3 | 1.9 | 8 | 4.47 | 0.11 | 0.325 | (A) BRAF Class 1 v12 |
| CHEK2 | 22q12.1 | 60 | 2.36 | 7 | 1.24 | 18 | 3.07 | 0.111 | 0.325 | (C) BRAF Class 3 |
| MAP2K2 | 19p13.3 | 32 | 1.45 | 12 | 2.2 | 14 | 2.68 | 0.113 | 0.33 | (C) BRAF Class 3 |
| IFNGR1 | 6q23.3 | 19 | 1.61 | 3 | 1.1 | 0 | 0 | 0.113 | 0.33 | (A) BRAF Class 1 v12 |
| EXO1 | 1q43 | 14 | 2.41 | 6 | 4.58 | 8 | 5.48 | 0.113 | 0.33 | (C) BRAF Class 3 |
| KMT2C | 7q36.1 | 184 | 12.62 | 28 | 8.54 | 40 | 12.54 | 0.114 | 0.33 | (A) BRAF Class 1 v12 |
| SUZ12 | 17q11.2 | 33 | 1.47 | 10 | 2 | 14 | 2.78 | 0.117 | 0.34 | (C) BRAF Class 3 |

| | | | | | | | | | | |
|---|---|---|---|---|---|---|---|---|---|---|
| FLI1 | 11q24.3 | 7 | 1.64 | 3 | 5.26 | 3 | 4.35 | 0.118 | 0.343 | (B) BRAF Class 2 |
| MCM8 | 20p12.3 | 14 | 2.41 | 7 | 5.34 | 7 | 4.79 | 0.12 | 0.346 | (B) BRAF Class 2 |
| NEGR1 | 1p31.1 | 28 | 2.07 | 7 | 2.33 | 12 | 4.1 | 0.121 | 0.348 | (C) BRAF Class 3 |
| NT5C2 | ı24.32-q24 | 11 | 1.33 | 6 | 3.19 | 6 | 2.86 | 0.122 | 0.351 | (B) BRAF Class 2 |
| CDH1 | 16q22.1 | 72 | 2.3 | 11 | 1.67 | 23 | 3.31 | 0.124 | 0.355 | (C) BRAF Class 3 |
| HABP2 | 10q25.3 | 16 | 2.75 | 1 | 0.76 | 7 | 4.79 | 0.126 | 0.359 | (C) BRAF Class 3 |
| SOX2 | 3q26.33 | 33 | 1.34 | 3 | 0.54 | 11 | 1.91 | 0.126 | 0.359 | (C) BRAF Class 3 |
| SYK | 9q22.2 | 44 | 2.32 | 14 | 3.3 | 17 | 3.94 | 0.126 | 0.359 | (C) BRAF Class 3 |
| CDH23 | 10q22.1 | 3 | 6.25 | 0 | 0 | 2 | 25 | 0.127 | 0.361 | (C) BRAF Class 3 |
| CD79B | 17q23.3 | 15 | 0.61 | 8 | 1.45 | 5 | 0.87 | 0.128 | 0.362 | (B) BRAF Class 2 |
| FANCD2 | 3p25.3 | 59 | 3.85 | 14 | 4.78 | 21 | 6.31 | 0.129 | 0.363 | (C) BRAF Class 3 |
| RICTOR | 5p13.1 | 74 | 3.36 | 24 | 4.83 | 25 | 4.84 | 0.129 | 0.363 | (C) BRAF Class 3 |
| LYN | 8q12.1 | 27 | 2.62 | 6 | 2.64 | 12 | 5.06 | 0.13 | 0.366 | (C) BRAF Class 3 |
| ETS1 | 11q24.3 | 7 | 1.24 | 2 | 2.47 | 4 | 4 | 0.133 | 0.372 | (C) BRAF Class 3 |
| FRK | 6q22.1 | 1 | 1.09 | 1 | 6.67 | 1 | 10 | 0.134 | 0.372 | (C) BRAF Class 3 |
| EGLN1 | 1q42.2 | 5 | 0.86 | 3 | 2.29 | 0 | 0 | 0.134 | 0.372 | (B) BRAF Class 2 |
| LRP1B | q22.1-q22. | 146 | 21.35 | 25 | 23.15 | 40 | 29.2 | 0.134 | 0.372 | (C) BRAF Class 3 |
| MBD4 | 3q21.3 | 15 | 2.58 | 0 | 0 | 2 | 1.37 | 0.135 | 0.376 | (A) BRAF Class 1 v12 |
| NR4A3 | 9q31.1 | 2 | 1.25 | 2 | 7.14 | 1 | 4.17 | 0.137 | 0.379 | (B) BRAF Class 2 |
| MST1 | 3p21.31 | 9 | 0.81 | 0 | 0 | 0 | 0 | 0.137 | 0.379 | (A) BRAF Class 1 v12 |
| HFE | 6p22.2 | 6 | 1.03 | 4 | 3.05 | 1 | 0.68 | 0.139 | 0.382 | (B) BRAF Class 2 |
| PRKCZ | 1p36.33 | 5 | 1.42 | 3 | 4.41 | 3 | 4.35 | 0.139 | 0.382 | (B) BRAF Class 2 |
| MYCL | 1p34.2 | 33 | 1.3 | 10 | 1.77 | 14 | 2.39 | 0.14 | 0.384 | (C) BRAF Class 3 |
| KLHL6 | 3q27.1 | 21 | 2.93 | 0 | 0 | 5 | 3.21 | 0.141 | 0.385 | (C) BRAF Class 3 |
| TFRC | 3q29 | 7 | 2.07 | 3 | 7.14 | 1 | 2.04 | 0.142 | 0.387 | (B) BRAF Class 2 |
| PRAME | 22q11.22 | 4 | 1.13 | 3 | 4.41 | 2 | 2.9 | 0.142 | 0.387 | (B) BRAF Class 2 |
| JAK1 | 1p31.3 | 79 | 3.52 | 19 | 3.45 | 28 | 5.29 | 0.143 | 0.387 | (C) BRAF Class 3 |
| MLH1 | 3p22.2 | 52 | 1.62 | 17 | 2.54 | 17 | 2.42 | 0.143 | 0.387 | (B) BRAF Class 2 |
| RAD50 | 5q31.1 | 94 | 4.31 | 14 | 2.92 | 14 | 2.8 | 0.144 | 0.39 | (A) BRAF Class 1 v12 |
| WWTR1 | 3q25.1 | 3 | 0.5 | 2 | 1.3 | 3 | 2.14 | 0.146 | 0.394 | (C) BRAF Class 3 |
| PTPN14 | 1q32.3-q41 | 17 | 2.92 | 2 | 1.53 | 8 | 5.48 | 0.147 | 0.395 | (C) BRAF Class 3 |
| TNFRSF14 | 1p36.32 | 17 | 1.08 | 8 | 2.25 | 3 | 0.83 | 0.15 | 0.402 | (B) BRAF Class 2 |
| TSC2 | 16p13.3 | 133 | 5.23 | 28 | 4.96 | 42 | 7.16 | 0.152 | 0.407 | (C) BRAF Class 3 |
| KAT6A | 8p11.21 | 68 | 5.43 | 18 | 7.53 | 23 | 8.07 | 0.153 | 0.409 | (C) BRAF Class 3 |
| NIPBL | 5p13.2 | 6 | 3.64 | 3 | 11.11 | 3 | 9.38 | 0.154 | 0.409 | (B) BRAF Class 2 |
| SLC25A13 | 7q21.3 | 9 | 1.55 | 5 | 3.82 | 5 | 3.42 | 0.154 | 0.409 | (B) BRAF Class 2 |
| HGF | 7q21.11 | 61 | 3.9 | 20 | 5.59 | 21 | 5.8 | 0.154 | 0.409 | (C) BRAF Class 3 |
| TOP1 | 20q12 | 27 | 1.68 | 5 | 1.43 | 11 | 3.13 | 0.155 | 0.41 | (C) BRAF Class 3 |
| GALNT12 | 9q22.33 | 12 | 1.63 | 6 | 3.64 | 2 | 1.08 | 0.156 | 0.41 | (B) BRAF Class 2 |
| DHX15 | 4p15.2 | 14 | 4.2 | 0 | 0 | 4 | 8.33 | 0.156 | 0.41 | (C) BRAF Class 3 |
| CDK12 | 17q12 | 71 | 3.21 | 24 | 4.83 | 22 | 4.26 | 0.156 | 0.41 | (B) BRAF Class 2 |
| ANKRD11 | 16q24.3 | 88 | 7.9 | 12 | 4.72 | 14 | 5.96 | 0.156 | 0.41 | (A) BRAF Class 1 v12 |
| MAPKAP1 | 9q33.3 | 5 | 0.94 | 0 | 0 | 3 | 2.33 | 0.157 | 0.412 | (C) BRAF Class 3 |
| ROBO1 | 3p12.3 | 7 | 4.4 | 2 | 7.69 | 4 | 13.33 | 0.158 | 0.413 | (C) BRAF Class 3 |
| FGF12 | 3q28-q29 | 0 | 0 | 0 | 0 | 1 | 1.85 | 0.159 | 0.413 | (C) BRAF Class 3 |

| | | | | | | | | | | |
|---|---|---|---|---|---|---|---|---|---|---|
| TIPARP | 3q25.31 | 0 | 0 | 0 | 0 | 1 | 1.85 | 0.159 | 0.413 | (C) BRAF Class 3 |
| CEBPA | 19q13.11 | 27 | 1.13 | 8 | 1.49 | 12 | 2.15 | 0.159 | 0.413 | (C) BRAF Class 3 |
| APOBEC3G | 22q13.1 | 1 | 1.16 | 1 | 8.33 | 0 | 0 | 0.16 | 0.414 | (B) BRAF Class 2 |
| INPP4B | 4q31.21 | 32 | 2.08 | 11 | 3.12 | 13 | 3.66 | 0.16 | 0.414 | (C) BRAF Class 3 |
| BCL6 | 3q27.3 | 57 | 2.32 | 10 | 1.81 | 20 | 3.47 | 0.16 | 0.414 | (C) BRAF Class 3 |
| TRAF3 | 14q32.32 | 13 | 1.61 | 6 | 3.47 | 2 | 1.01 | 0.163 | 0.419 | (B) BRAF Class 2 |
| KLLN | 10q23.31 | 14 | 2.41 | 0 | 0 | 2 | 1.37 | 0.164 | 0.421 | (A) BRAF Class 1 v12 |
| TBX3 | 12q24.21 | 69 | 4.51 | 23 | 6.57 | 22 | 6.23 | 0.165 | 0.424 | (B) BRAF Class 2 |
| TENT5C | 1p12 | 37 | 1.51 | 8 | 1.45 | 15 | 2.6 | 0.166 | 0.424 | (C) BRAF Class 3 |
| PPARG | 3p25.2 | 39 | 2.2 | 15 | 3.86 | 11 | 2.66 | 0.166 | 0.424 | (B) BRAF Class 2 |
| ALK | p23.2-p23. | 181 | 5.42 | 38 | 5.07 | 52 | 7.06 | 0.17 | 0.432 | (C) BRAF Class 3 |
| IKZF2 | 2q34 | 4 | 1.82 | 1 | 2.7 | 3 | 6.82 | 0.17 | 0.432 | (C) BRAF Class 3 |
| HIST3H3 | 1q42.13 | 8 | 0.72 | 5 | 1.97 | 2 | 0.85 | 0.173 | 0.437 | (B) BRAF Class 2 |
| ID3 | 1p36.12 | 8 | 0.35 | 2 | 0.39 | 5 | 0.96 | 0.173 | 0.437 | (C) BRAF Class 3 |
| PRPF40B | 12q13.12 | 10 | 2.31 | 5 | 6.02 | 2 | 2.47 | 0.173 | 0.437 | (B) BRAF Class 2 |
| DDR1 | 6p21.33 | 3 | 1.46 | 3 | 5.66 | 1 | 1.75 | 0.175 | 0.441 | (B) BRAF Class 2 |
| GAB2 | 11q14.1 | 3 | 1.81 | 0 | 0 | 2 | 6.67 | 0.176 | 0.441 | (C) BRAF Class 3 |
| SERPINA1 | 14q32.13 | 15 | 2.58 | 0 | 0 | 3 | 2.05 | 0.177 | 0.444 | (A) BRAF Class 1 v12 |
| BABAM1 | 19p13.11 | 14 | 1.28 | 0 | 0 | 3 | 1.09 | 0.178 | 0.445 | (A) BRAF Class 1 v12 |
| USP28 | 11q23.2 | 20 | 3.44 | 9 | 6.87 | 5 | 3.42 | 0.178 | 0.445 | (B) BRAF Class 2 |
| RPL10 | Xq28 | 1 | 1.05 | 1 | 7.14 | 0 | 0 | 0.181 | 0.451 | (B) BRAF Class 2 |
| COP1 | q25.1-q25. | 34 | 2.25 | 10 | 2.96 | 13 | 4 | 0.182 | 0.452 | (C) BRAF Class 3 |
| ERG | 21q22.2 | 63 | 2.96 | 12 | 2.46 | 22 | 4.33 | 0.187 | 0.462 | (C) BRAF Class 3 |
| CRTC2 | 1q21.3 | 7 | 1.98 | 3 | 4.41 | 0 | 0 | 0.187 | 0.462 | (B) BRAF Class 2 |
| UIMC1 | 5q35.2 | 14 | 2.32 | 1 | 0.72 | 6 | 3.97 | 0.188 | 0.464 | (C) BRAF Class 3 |
| INSRR | 1q23.1 | 6 | 0.18 | 0 | 0 | 3 | 0.4 | 0.192 | 0.472 | (C) BRAF Class 3 |
| DOCK8 | 9p24.3 | 48 | 8.25 | 11 | 8.4 | 19 | 13.01 | 0.192 | 0.472 | (C) BRAF Class 3 |
| TEK | 9p21.2 | 33 | 4.63 | 15 | 7.89 | 9 | 4.84 | 0.192 | 0.472 | (B) BRAF Class 2 |
| MLH3 | 14q24.3 | 28 | 3.48 | 9 | 5.26 | 12 | 6.09 | 0.193 | 0.473 | (C) BRAF Class 3 |
| GBA | 1q22 | 6 | 1.03 | 4 | 3.05 | 3 | 2.05 | 0.194 | 0.474 | (B) BRAF Class 2 |
| INHBA | 7p14.1 | 43 | 2.95 | 14 | 4.35 | 15 | 4.63 | 0.196 | 0.477 | (C) BRAF Class 3 |
| NUF2 | 1q23.3 | 10 | 1.89 | 1 | 0.72 | 0 | 0 | 0.196 | 0.477 | (A) BRAF Class 1 v12 |
| MALT1 | 18q21.32 | 17 | 1.4 | 7 | 2.55 | 7 | 2.71 | 0.197 | 0.477 | (C) BRAF Class 3 |
| SH2B3 | 12q24.12 | 61 | 2.6 | 8 | 1.54 | 9 | 1.66 | 0.197 | 0.477 | (A) BRAF Class 1 v12 |
| DNMT3B | 20q11.21 | 34 | 2.87 | 9 | 3.31 | 13 | 5.08 | 0.199 | 0.481 | (C) BRAF Class 3 |
| FGF23 | 12p13.32 | 9 | 1.27 | 4 | 3.2 | 4 | 2.6 | 0.206 | 0.497 | (B) BRAF Class 2 |
| COL7A1 | 3p21.31 | 94 | 16.15 | 26 | 19.85 | 32 | 21.92 | 0.206 | 0.497 | (C) BRAF Class 3 |
| PPP6C | 9q33.3 | 42 | 3.3 | 5 | 1.77 | 5 | 1.83 | 0.207 | 0.498 | (A) BRAF Class 1 v12 |
| PALB2 | 16p12.2 | 87 | 3.43 | 25 | 4.44 | 28 | 4.78 | 0.207 | 0.498 | (C) BRAF Class 3 |
| TYK2 | 19p13.2 | 6 | 2.41 | 3 | 6.82 | 3 | 5.77 | 0.21 | 0.502 | (B) BRAF Class 2 |
| ELANE | 19p13.3 | 14 | 2.15 | 0 | 0 | 3 | 1.92 | 0.21 | 0.502 | (A) BRAF Class 1 v12 |
| BRCA2 | 13q13.1 | 204 | 8.03 | 36 | 6.38 | 54 | 9.17 | 0.212 | 0.505 | (C) BRAF Class 3 |
| STAT4 | q32.2-q32. | 23 | 3.39 | 4 | 3.7 | 9 | 6.57 | 0.214 | 0.51 | (C) BRAF Class 3 |
| TP53BP1 | 15q15.3 | 58 | 4.91 | 11 | 3.78 | 21 | 6.84 | 0.215 | 0.51 | (C) BRAF Class 3 |
| HOXB13 | 17q21.32 | 18 | 0.94 | 7 | 1.62 | 8 | 1.78 | 0.217 | 0.513 | (C) BRAF Class 3 |

| | | | | | | | | | | |
|---|---|---|---|---|---|---|---|---|---|---|
| ARHGAP35 | 19q13.32 | 31 | 4.27 | 7 | 4.46 | 13 | 7.39 | 0.217 | 0.514 | (C) BRAF Class 3 |
| SDHA | 5p15.33 | 71 | 3.11 | 10 | 1.93 | 20 | 3.71 | 0.219 | 0.518 | (C) BRAF Class 3 |
| HIST1H3I | 6p22.1 | 6 | 0.54 | 4 | 1.57 | 2 | 0.85 | 0.22 | 0.518 | (B) BRAF Class 2 |
| RHOT1 | 17q11.2 | 6 | 1.03 | 1 | 0.76 | 4 | 2.74 | 0.221 | 0.519 | (C) BRAF Class 3 |
| KMT2A | 11q23.3 | 214 | 8.6 | 54 | 9.66 | 63 | 10.81 | 0.221 | 0.519 | (C) BRAF Class 3 |
| BCL2L12 | 19q13.33 | 11 | 1.13 | 4 | 1.93 | 6 | 2.55 | 0.225 | 0.525 | (C) BRAF Class 3 |
| CTNNA1 | 5q31.2 | 27 | 2.12 | 10 | 3.94 | 7 | 2.33 | 0.225 | 0.525 | (B) BRAF Class 2 |
| MEF2B | 19p13.11 | 28 | 1.14 | 10 | 1.81 | 11 | 1.91 | 0.226 | 0.526 | (C) BRAF Class 3 |
| GSK3B | 3q13.33 | 17 | 1.09 | 8 | 2.23 | 5 | 1.38 | 0.226 | 0.526 | (B) BRAF Class 2 |
| PTK2 | 8q24.3 | 8 | 2.27 | 0 | 0 | 3 | 4.35 | 0.228 | 0.531 | (C) BRAF Class 3 |
| ALOX12B | 17p13.1 | 37 | 2.24 | 8 | 2.1 | 14 | 3.69 | 0.229 | 0.531 | (C) BRAF Class 3 |
| MUTYH | 1p34.1 | 45 | 1.83 | 11 | 1.99 | 17 | 2.95 | 0.23 | 0.531 | (C) BRAF Class 3 |
| PRSS8 | 16p11.2 | 7 | 1.36 | 0 | 0 | 3 | 3.06 | 0.231 | 0.531 | (C) BRAF Class 3 |
| EML4 | 2p21 | 13 | 3 | 4 | 6.78 | 4 | 5.71 | 0.231 | 0.531 | (B) BRAF Class 2 |
| SNCAIP | 5q23.2 | 44 | 6.69 | 9 | 7.96 | 15 | 10.87 | 0.231 | 0.531 | (C) BRAF Class 3 |
| FGF3 | 11q13.3 | 27 | 1.64 | 8 | 2.17 | 11 | 2.96 | 0.231 | 0.531 | (C) BRAF Class 3 |
| MGMT | 10q26.3 | 1 | 0.64 | 1 | 4 | 0 | 0 | 0.232 | 0.531 | (B) BRAF Class 2 |
| RACK1 | 5q35.3 | 4 | 1.52 | 2 | 4.17 | 0 | 0 | 0.233 | 0.532 | (B) BRAF Class 2 |
| CRTC1 | 19p13.11 | 37 | 3.65 | 7 | 3.24 | 14 | 5.91 | 0.233 | 0.532 | (C) BRAF Class 3 |
| TSHZ3 | 19q12 | 2 | 2.33 | 1 | 8.33 | 2 | 10 | 0.234 | 0.532 | (C) BRAF Class 3 |
| NOTCH4 | 6p21.32 | 91 | 6.42 | 21 | 6.58 | 29 | 9.06 | 0.234 | 0.532 | (C) BRAF Class 3 |
| SHOC2 | 10q25.2 | 8 | 1.51 | 0 | 0 | 3 | 2.33 | 0.239 | 0.543 | (C) BRAF Class 3 |
| BRD4 | 19p13.12 | 126 | 5.23 | 19 | 3.49 | 28 | 4.97 | 0.24 | 0.543 | (A) BRAF Class 1 v12 |
| HIST1H3H | 6p22.1 | 4 | 0.36 | 3 | 1.18 | 1 | 0.43 | 0.241 | 0.544 | (B) BRAF Class 2 |
| PRKAR1A | 17q24.2 | 23 | 0.93 | 7 | 1.27 | 10 | 1.73 | 0.241 | 0.544 | (C) BRAF Class 3 |
| TET3 | 2p13.1 | 1 | 2.94 | 1 | 20 | 0 | 0 | 0.242 | 0.544 | (B) BRAF Class 2 |
| NECTIN4 | 1q23.3 | 8 | 1.37 | 3 | 2.29 | 5 | 3.42 | 0.242 | 0.544 | (C) BRAF Class 3 |
| SDHD | 11q23.1 | 9 | 0.37 | 5 | 0.91 | 3 | 0.53 | 0.244 | 0.548 | (B) BRAF Class 2 |
| AMER1 | Xq11.2 | 64 | 4.08 | 17 | 4.75 | 22 | 6.08 | 0.246 | 0.548 | (C) BRAF Class 3 |
| FGF4 | 11q13.3 | 29 | 1.85 | 10 | 2.79 | 4 | 1.1 | 0.247 | 0.548 | (B) BRAF Class 2 |
| HSP90AB1 | 6p21.1 | 1 | 0.4 | 1 | 2.38 | 0 | 0 | 0.247 | 0.548 | (B) BRAF Class 2 |
| MT-ND5 | - | 12 | 0.36 | 0 | 0 | 3 | 0.4 | 0.247 | 0.548 | (C) BRAF Class 3 |
| GLI2 | 2q14.2 | 79 | 7.55 | 24 | 10.91 | 21 | 8.57 | 0.248 | 0.548 | (B) BRAF Class 2 |
| RYBP | 3p13 | 8 | 0.68 | 4 | 1.47 | 4 | 1.56 | 0.248 | 0.548 | (C) BRAF Class 3 |
| PC | 11q13.2 | 1 | 1.16 | 1 | 7.14 | 0 | 0 | 0.249 | 0.548 | (B) BRAF Class 2 |
| PCLO | 7q21.11 | 1 | 1.16 | 1 | 7.14 | 0 | 0 | 0.249 | 0.548 | (B) BRAF Class 2 |
| PDCD11 | 10q24.33 | 1 | 1.16 | 1 | 7.14 | 0 | 0 | 0.249 | 0.548 | (B) BRAF Class 2 |
| CUL3 | 2q36.2 | 20 | 1.3 | 3 | 0.85 | 8 | 2.26 | 0.249 | 0.548 | (C) BRAF Class 3 |
| ESPL1 | 12q13.13 | 1 | 1.16 | 1 | 8.33 | 1 | 5 | 0.25 | 0.549 | (B) BRAF Class 2 |
| TLR2 | 4q31.3 | 1 | 2.08 | 1 | 12.5 | 0 | 0 | 0.252 | 0.55 | (B) BRAF Class 2 |
| SDHAF2 | 11q12.2 | 10 | 0.45 | 1 | 0.2 | 0 | 0 | 0.253 | 0.55 | (A) BRAF Class 1 v12 |
| U2AF1 | 21q22.3 | 38 | 1.51 | 14 | 2.5 | 11 | 1.89 | 0.253 | 0.55 | (B) BRAF Class 2 |
| PNKP | 19q13.33 | 12 | 2.06 | 0 | 0 | 3 | 2.05 | 0.253 | 0.55 | (A) BRAF Class 1 v12 |
| ACVR2A | q22.3-q23. | 1 | 0.85 | 1 | 5.26 | 1 | 4.76 | 0.253 | 0.55 | (B) BRAF Class 2 |
| TGM7 | 5q15.2-q15 | 1 | 0.85 | 1 | 5.26 | 1 | 4.76 | 0.253 | 0.55 | (B) BRAF Class 2 |

| | | | | | | | | | | |
|---|---|---|---|---|---|---|---|---|---|---|
| PPP2R2A | 8p21.2 | 8 | 2.2 | 0 | 0 | 0 | 0 | 0.255 | 0.553 | (A) BRAF Class 1 v12 |
| LAMP1 | 13q34 | 6 | 1.4 | 2 | 3.51 | 0 | 0 | 0.256 | 0.555 | (B) BRAF Class 2 |
| POLQ | 3q13.33 | 53 | 8.2 | 15 | 10.64 | 20 | 12.05 | 0.257 | 0.555 | (C) BRAF Class 3 |
| IRS1 | 2q36.3 | 63 | 5.33 | 8 | 2.94 | 13 | 5.08 | 0.258 | 0.556 | (A) BRAF Class 1 v12 |
| PPP4R2 | 3p13 | 8 | 1.51 | 5 | 3.6 | 2 | 1.55 | 0.26 | 0.56 | (B) BRAF Class 2 |
| ECT2L | 6q24.1 | 1 | 1 | 0 | 0 | 1 | 5.88 | 0.261 | 0.561 | (C) BRAF Class 3 |
| IL6ST | 5q11.2 | 1 | 0.5 | 0 | 0 | 1 | 2.94 | 0.262 | 0.562 | (C) BRAF Class 3 |
| GRM8 | 7q31.33 | 4 | 3.42 | 0 | 0 | 2 | 9.52 | 0.264 | 0.564 | (C) BRAF Class 3 |
| SMARCAL1 | 2q35 | 2 | 2.17 | 0 | 0 | 1 | 10 | 0.264 | 0.564 | (C) BRAF Class 3 |
| CCNE1 | 19q12 | 44 | 1.73 | 15 | 2.66 | 14 | 2.39 | 0.265 | 0.566 | (B) BRAF Class 2 |
| NIN | 14q22.1 | 3 | 2.46 | 2 | 9.52 | 1 | 4.55 | 0.271 | 0.577 | (B) BRAF Class 2 |
| CALR | 19p13.13 | 26 | 1.34 | 5 | 1.12 | 2 | 0.44 | 0.273 | 0.579 | (A) BRAF Class 1 v12 |
| SSX2 | Xp11.22 | 1 | 1.75 | 0 | 0 | 1 | 10 | 0.274 | 0.579 | (C) BRAF Class 3 |
| RPS6KB1 | 17q23.1 | 5 | 1.5 | 0 | 0 | 2 | 4.17 | 0.274 | 0.579 | (C) BRAF Class 3 |
| PARP1 | 1q42.12 | 44 | 3.11 | 13 | 3.92 | 6 | 1.81 | 0.275 | 0.579 | (B) BRAF Class 2 |
| CIC | 19q13.2 | 148 | 6.95 | 25 | 5.13 | 38 | 7.48 | 0.275 | 0.579 | (C) BRAF Class 3 |
| GLMN | 1p22.1 | 1 | 1.09 | 1 | 6.67 | 0 | 0 | 0.275 | 0.58 | (B) BRAF Class 2 |
| PREX2 | 8q13.2 | 139 | 11.96 | 28 | 11.72 | 40 | 15.5 | 0.276 | 0.58 | (C) BRAF Class 3 |
| SF1 | 11q13.1 | 13 | 3.01 | 0 | 0 | 2 | 2.47 | 0.277 | 0.58 | (A) BRAF Class 1 v12 |
| PNRC1 | 6q15 | 17 | 1.05 | 1 | 0.27 | 2 | 0.55 | 0.277 | 0.58 | (A) BRAF Class 1 v12 |
| MAP3K9 | 14q24.2 | 9 | 4.89 | 0 | 0 | 3 | 8.33 | 0.278 | 0.582 | (C) BRAF Class 3 |
| TET1 | 10q21.3 | 117 | 6.34 | 29 | 6.89 | 36 | 8.49 | 0.28 | 0.584 | (C) BRAF Class 3 |
| TOPBP1 | 3q22.1 | 23 | 3.89 | 9 | 6.77 | 5 | 3.38 | 0.281 | 0.586 | (B) BRAF Class 2 |
| PIK3CD | 1p36.22 | 38 | 2.97 | 5 | 1.73 | 11 | 3.97 | 0.282 | 0.588 | (C) BRAF Class 3 |
| HELQ | 4q21.23 | 17 | 2.92 | 6 | 4.58 | 2 | 1.37 | 0.284 | 0.589 | (B) BRAF Class 2 |
| NSD2 | 4p16.3 | 44 | 3.23 | 16 | 4.86 | 15 | 4.35 | 0.287 | 0.594 | (B) BRAF Class 2 |
| MAPK3 | 16p11.2 | 20 | 1.79 | 5 | 1.96 | 1 | 0.43 | 0.287 | 0.594 | (B) BRAF Class 2 |
| ESR2 | 4q23.2-q23 | 1 | 0.39 | 1 | 2 | 0 | 0 | 0.288 | 0.595 | (B) BRAF Class 2 |
| DEPDC5 | 2q12.2-q12 | 13 | 4.94 | 4 | 8.33 | 1 | 1.72 | 0.289 | 0.596 | (B) BRAF Class 2 |
| TRAF2 | 9q34.3 | 25 | 2.12 | 2 | 0.75 | 6 | 2.41 | 0.291 | 0.599 | (C) BRAF Class 3 |
| EXT1 | 8q24.11 | 21 | 2.05 | 5 | 2.29 | 9 | 3.77 | 0.292 | 0.6 | (C) BRAF Class 3 |
| HLF | 17q22 | 1 | 0.82 | 0 | 0 | 1 | 4.55 | 0.293 | 0.6 | (C) BRAF Class 3 |
| EPAS1 | 2p21 | 12 | 2.26 | 3 | 2.16 | 6 | 4.65 | 0.293 | 0.6 | (C) BRAF Class 3 |
| TAF1L | 9p21.1 | 1 | 0.85 | 0 | 0 | 1 | 4.76 | 0.295 | 0.603 | (C) BRAF Class 3 |
| RBM15 | 1p13.3 | 1 | 0.86 | 1 | 4.76 | 1 | 4.55 | 0.296 | 0.604 | (B) BRAF Class 2 |
| DKC1 | Xq28 | 8 | 1.37 | 0 | 0 | 3 | 2.04 | 0.297 | 0.604 | (C) BRAF Class 3 |
| PTEN | 10q23.31 | 350 | 10.46 | 61 | 8.7 | 69 | 9.36 | 0.3 | 0.609 | (A) BRAF Class 1 v12 |
| PDGFB | 22q13.1 | 1 | 0.39 | 1 | 2.27 | 1 | 1.92 | 0.301 | 0.611 | (B) BRAF Class 2 |
| PLAG1 | 8q12.1 | 1 | 0.38 | 1 | 2.17 | 1 | 1.85 | 0.305 | 0.617 | (B) BRAF Class 2 |
| AXIN1 | 16p13.3 | 56 | 3.68 | 8 | 2.29 | 9 | 2.59 | 0.305 | 0.617 | (A) BRAF Class 1 v12 |
| TSC1 | 9q34.13 | 81 | 3.18 | 21 | 3.72 | 26 | 4.43 | 0.309 | 0.625 | (C) BRAF Class 3 |
| RAD54L | 1p34.1 | 19 | 1.44 | 2 | 0.63 | 2 | 0.65 | 0.312 | 0.626 | (A) BRAF Class 1 v12 |
| AFDN | 6q27 | 1 | 0.86 | 0 | 0 | 1 | 4.55 | 0.312 | 0.626 | (C) BRAF Class 3 |
| ASXL2 | 2p23.3 | 58 | 4.39 | 17 | 5.74 | 9 | 3.15 | 0.312 | 0.626 | (B) BRAF Class 2 |
| UBE2T | 1q32.1 | 4 | 0.69 | 2 | 1.53 | 0 | 0 | 0.313 | 0.626 | (B) BRAF Class 2 |

| | | | | | | | | | | |
|---|---|---|---|---|---|---|---|---|---|---|
| ARFRP1 | 20q13.33 | 7 | 1 | 3 | 2.44 | 1 | 0.65 | 0.313 | 0.626 | (B) BRAF Class 2 |
| ZFHX3 | 5q22.2-q22 | 153 | 12.26 | 25 | 9.03 | 31 | 11.7 | 0.317 | 0.633 | (A) BRAF Class 1 v12 |
| PIK3CG | 7q22.3 | 76 | 4.94 | 18 | 5.28 | 24 | 6.96 | 0.317 | 0.633 | (C) BRAF Class 3 |
| DMC1 | 22q13.1 | 10 | 1.72 | 0 | 0 | 2 | 1.37 | 0.318 | 0.633 | (A) BRAF Class 1 v12 |
| ZMYM3 | Xq13.1 | 6 | 2.73 | 2 | 5.41 | 0 | 0 | 0.319 | 0.634 | (B) BRAF Class 2 |
| EME1 | 17q21.33 | 10 | 1.72 | 4 | 3.05 | 1 | 0.68 | 0.322 | 0.639 | (B) BRAF Class 2 |
| PTPN1 | 20q13.13 | 1 | 0.6 | 0 | 0 | 1 | 3.13 | 0.323 | 0.64 | (C) BRAF Class 3 |
| EIF1AX | Xp22.12 | 19 | 1.38 | 4 | 1.29 | 1 | 0.34 | 0.324 | 0.64 | (A) BRAF Class 1 v12 |
| SUFU | 10q24.32 | 45 | 1.83 | 10 | 1.81 | 16 | 2.77 | 0.325 | 0.642 | (C) BRAF Class 3 |
| MST1R | 3p21.31 | 46 | 3.57 | 15 | 4.95 | 15 | 5.08 | 0.327 | 0.645 | (C) BRAF Class 3 |
| PHF1 | 6p21.32 | 5 | 0.15 | 0 | 0 | 0 | 0 | 0.328 | 0.645 | (A) BRAF Class 1 v12 |
| CHEK1 | 11q24.2 | 27 | 1.22 | 10 | 2.01 | 9 | 1.74 | 0.328 | 0.645 | (B) BRAF Class 2 |
| DNMT3A | 2p23.3 | 126 | 5.06 | 37 | 6.61 | 33 | 5.63 | 0.329 | 0.645 | (B) BRAF Class 2 |
| MYB | 6q23.3 | 19 | 1.21 | 4 | 1.28 | 8 | 2.22 | 0.329 | 0.645 | (C) BRAF Class 3 |
| TMPRSS2 | 21q22.3 | 40 | 1.9 | 10 | 2.07 | 15 | 2.96 | 0.33 | 0.646 | (C) BRAF Class 3 |
| STAT5B | 17q21.2 | 37 | 3.1 | 4 | 1.48 | 8 | 3.17 | 0.332 | 0.648 | (C) BRAF Class 3 |
| POLB | 8p11.21 | 9 | 1.55 | 3 | 2.29 | 5 | 3.42 | 0.333 | 0.648 | (C) BRAF Class 3 |
| PTCH1 | 9q22.32 | 132 | 5.2 | 22 | 3.9 | 25 | 4.27 | 0.334 | 0.648 | (A) BRAF Class 1 v12 |
| PDE4DIP | 1q21.2 | 6 | 4.92 | 0 | 0 | 0 | 0 | 0.334 | 0.648 | (A) BRAF Class 1 v12 |
| SLX4 | 16p13.3 | 97 | 6.55 | 22 | 6.96 | 15 | 4.52 | 0.336 | 0.651 | (B) BRAF Class 2 |
| KLF2 | 19p13.11 | 8 | 1.37 | 0 | 0 | 1 | 0.68 | 0.338 | 0.653 | (A) BRAF Class 1 v12 |
| XBP1 | q12.1\|22q | 1 | 0.45 | 0 | 0 | 1 | 2.27 | 0.339 | 0.655 | (C) BRAF Class 3 |
| POLE | 12q24.33 | 161 | 7.28 | 30 | 5.99 | 43 | 8.37 | 0.343 | 0.662 | (C) BRAF Class 3 |
| SMARCD1 | 12q13.12 | 24 | 1.86 | 2 | 0.67 | 5 | 1.71 | 0.345 | 0.665 | (A) BRAF Class 1 v12 |
| TDG | 12q23.3 | 8 | 1.37 | 2 | 1.53 | 0 | 0 | 0.351 | 0.675 | (B) BRAF Class 2 |
| EXT2 | 11p11.2 | 18 | 1.76 | 7 | 3.21 | 4 | 1.67 | 0.352 | 0.675 | (B) BRAF Class 2 |
| ATF1 | 12q13.12 | 1 | 0.39 | 0 | 0 | 1 | 1.92 | 0.353 | 0.675 | (C) BRAF Class 3 |
| DDIT3 | 12q13.3 | 1 | 0.39 | 0 | 0 | 1 | 1.92 | 0.353 | 0.675 | (C) BRAF Class 3 |
| SDHB | 1p36.13 | 10 | 0.41 | 4 | 0.73 | 1 | 0.18 | 0.357 | 0.681 | (B) BRAF Class 2 |
| SMAD4 | 18q21.2 | 230 | 7.01 | 41 | 5.99 | 57 | 7.94 | 0.358 | 0.682 | (C) BRAF Class 3 |
| MYD88 | 3p22.2 | 31 | 1.21 | 10 | 1.76 | 5 | 0.84 | 0.358 | 0.682 | (B) BRAF Class 2 |
| MAPK1 | 22q11.22 | 19 | 0.79 | 7 | 1.3 | 7 | 1.28 | 0.36 | 0.685 | (B) BRAF Class 2 |
| PAXIP1 | 7q36.2 | 25 | 4.3 | 8 | 6.11 | 10 | 6.85 | 0.369 | 0.699 | (C) BRAF Class 3 |
| ERCC3 | 2q14.3 | 39 | 1.79 | 12 | 2.47 | 13 | 2.64 | 0.369 | 0.699 | (C) BRAF Class 3 |
| MC1-LUC7L | 7q34 | 1 | 0.03 | 0 | 0 | 1 | 0.13 | 0.372 | 0.702 | (C) BRAF Class 3 |
| MT-ND6 | - | 1 | 0.03 | 0 | 0 | 1 | 0.13 | 0.372 | 0.702 | (C) BRAF Class 3 |
| RHEB | 7q36.1 | 11 | 0.52 | 5 | 1.08 | 3 | 0.63 | 0.375 | 0.707 | (B) BRAF Class 2 |
| GATA4 | 8p23.1 | 32 | 2.26 | 4 | 1.4 | 10 | 3.1 | 0.376 | 0.707 | (C) BRAF Class 3 |
| BCL2L11 | 2q13 | 8 | 0.67 | 0 | 0 | 2 | 0.78 | 0.378 | 0.707 | (C) BRAF Class 3 |
| IGF2R | 6q25.3 | 8 | 2.4 | 1 | 1.59 | 4 | 4.94 | 0.378 | 0.707 | (C) BRAF Class 3 |
| HIST1H2BD | 6p22.2 | 9 | 0.76 | 2 | 0.73 | 0 | 0 | 0.378 | 0.707 | (A) BRAF Class 1 v12 |
| HIST1H3D | 6p22.2 | 8 | 0.72 | 1 | 0.39 | 0 | 0 | 0.378 | 0.707 | (A) BRAF Class 1 v12 |
| FOXL2 | 3q22.3 | 46 | 2.08 | 9 | 1.81 | 6 | 1.16 | 0.379 | 0.708 | (A) BRAF Class 1 v12 |
| NPM1 | 5q35.1 | 22 | 0.71 | 7 | 1.07 | 3 | 0.44 | 0.382 | 0.713 | (B) BRAF Class 2 |
| LZTR1 | 21\|22q11. | 25 | 3.89 | 5 | 5 | 8 | 6.61 | 0.388 | 0.72 | (C) BRAF Class 3 |

| | | | | | | | | | | |
|---|---|---|---|---|---|---|---|---|---|---|
| PRPF8 | 17p13.3 | 14 | 3.24 | 5 | 6.02 | 2 | 2.47 | 0.388 | 0.72 | (B) BRAF Class 2 |
| DDX41 | 5q35.3 | 5 | 1.07 | 2 | 3.17 | 1 | 1.28 | 0.388 | 0.72 | (B) BRAF Class 2 |
| INSR | 19p13.2 | 57 | 4.29 | 9 | 2.96 | 9 | 3.01 | 0.389 | 0.72 | (A) BRAF Class 1 v12 |
| AKT3 | 1q43-q44 | 35 | 1.36 | 10 | 1.76 | 5 | 0.84 | 0.391 | 0.724 | (B) BRAF Class 2 |
| INPP4A | 2q11.2 | 23 | 1.94 | 3 | 1.1 | 7 | 2.73 | 0.394 | 0.729 | (C) BRAF Class 3 |
| GNA11 | 19p13.3 | 24 | 0.74 | 7 | 0.95 | 9 | 1.24 | 0.396 | 0.73 | (C) BRAF Class 3 |
| PDCD1 | 2q37.3 | 33 | 2.56 | 4 | 1.32 | 8 | 2.78 | 0.397 | 0.73 | (C) BRAF Class 3 |
| ERCC2 | 19q13.32 | 77 | 3.09 | 23 | 4.23 | 18 | 3.16 | 0.399 | 0.733 | (B) BRAF Class 2 |
| DCC | 18q21.2 | 14 | 5.58 | 4 | 9.52 | 5 | 9.8 | 0.4 | 0.735 | (C) BRAF Class 3 |
| FKBP9 | 7p14.3 | 6 | 1.7 | 0 | 0 | 2 | 2.9 | 0.401 | 0.735 | (C) BRAF Class 3 |
| KLF4 | 9q31.2 | 34 | 1.75 | 11 | 2.52 | 6 | 1.35 | 0.403 | 0.736 | (B) BRAF Class 2 |
| STAG1 | 3q22.3 | 28 | 4.14 | 5 | 4.59 | 2 | 1.68 | 0.403 | 0.736 | (B) BRAF Class 2 |
| SOX9 | 17q24.3 | 109 | 4.54 | 19 | 3.52 | 29 | 5.17 | 0.405 | 0.738 | (C) BRAF Class 3 |
| FOXP1 | 3p13 | 35 | 2.31 | 12 | 3.57 | 8 | 2.37 | 0.405 | 0.738 | (B) BRAF Class 2 |
| NFKBIE | 6p21.1 | 13 | 1.82 | 4 | 2.6 | 6 | 3.41 | 0.406 | 0.738 | (C) BRAF Class 3 |
| PGBD3 | 10q11.23 | 11 | 0.33 | 5 | 0.67 | 3 | 0.4 | 0.407 | 0.738 | (B) BRAF Class 2 |
| DIS3 | 13q21.33 | 70 | 2.95 | 21 | 3.9 | 21 | 3.73 | 0.408 | 0.74 | (B) BRAF Class 2 |
| TCF7L2 | )q25.2-q25 | 76 | 3.39 | 13 | 2.65 | 21 | 4.18 | 0.41 | 0.742 | (C) BRAF Class 3 |
| RECQL4 | 8q24.3 | 121 | 5.62 | 24 | 5.01 | 34 | 6.91 | 0.411 | 0.742 | (C) BRAF Class 3 |
| TAL1 | 1p33 | 10 | 1.47 | 3 | 2 | 5 | 2.98 | 0.411 | 0.742 | (C) BRAF Class 3 |
| NAV3 | 12q21.2 | 8 | 5.13 | 3 | 12 | 2 | 6.67 | 0.412 | 0.742 | (B) BRAF Class 2 |
| RBL2 | 16q12.2 | 8 | 2.27 | 0 | 0 | 2 | 2.9 | 0.415 | 0.746 | (C) BRAF Class 3 |
| CDK8 | 13q12.13 | 36 | 1.69 | 10 | 2.05 | 13 | 2.56 | 0.415 | 0.746 | (C) BRAF Class 3 |
| CENPA | 2p23.3 | 5 | 0.45 | 2 | 0.79 | 0 | 0 | 0.416 | 0.746 | (B) BRAF Class 2 |
| GATA3 | 10p14 | 72 | 2.92 | 17 | 3.07 | 23 | 3.99 | 0.417 | 0.746 | (C) BRAF Class 3 |
| AXIN2 | 17q24.1 | 83 | 4.28 | 17 | 3.91 | 13 | 2.92 | 0.418 | 0.747 | (A) BRAF Class 1 v12 |
| SOS1 | 2p22.1 | 38 | 3.1 | 9 | 3.09 | 14 | 4.59 | 0.42 | 0.75 | (C) BRAF Class 3 |
| NSD1 | 5q35.3 | 126 | 6.03 | 32 | 6.84 | 37 | 7.57 | 0.422 | 0.75 | (C) BRAF Class 3 |
| GREM1 | 15q13.3 | 11 | 0.56 | 5 | 1.12 | 3 | 0.66 | 0.422 | 0.75 | (B) BRAF Class 2 |
| GNA13 | 17q24.1 | 3 | 0.42 | 1 | 0.79 | 2 | 1.29 | 0.423 | 0.75 | (C) BRAF Class 3 |
| HDAC9 | 7p21.1 | 3 | 3.49 | 1 | 8.33 | 2 | 10 | 0.424 | 0.75 | (C) BRAF Class 3 |
| TSHZ2 | 20q13.2 | 3 | 3.49 | 1 | 8.33 | 2 | 10 | 0.424 | 0.75 | (C) BRAF Class 3 |
| FANCI | 15q26.1 | 47 | 4.13 | 12 | 5.58 | 8 | 3.16 | 0.424 | 0.75 | (B) BRAF Class 2 |
| PIM1 | 6p21.2 | 18 | 0.82 | 7 | 1.39 | 6 | 1.18 | 0.427 | 0.753 | (B) BRAF Class 2 |
| ARID1B | 6q25.3 | 171 | 7.2 | 40 | 7.63 | 48 | 8.82 | 0.43 | 0.758 | (C) BRAF Class 3 |
| CREBBP | 16p13.3 | 192 | 7.56 | 39 | 6.93 | 52 | 8.87 | 0.431 | 0.76 | (C) BRAF Class 3 |
| RXRA | 9q34.2 | 18 | 3.4 | 2 | 1.44 | 3 | 2.33 | 0.432 | 0.76 | (A) BRAF Class 1 v12 |
| PAX3 | 2q36.1 | 11 | 1.94 | 3 | 3.66 | 1 | 1 | 0.434 | 0.762 | (B) BRAF Class 2 |
| SH2D1A | Xq25 | 11 | 0.6 | 3 | 0.72 | 5 | 1.18 | 0.435 | 0.763 | (C) BRAF Class 3 |
| PTPRO | .2.3\|12p13 | 1 | 0.86 | 1 | 3.23 | 0 | 0 | 0.437 | 0.763 | (B) BRAF Class 2 |
| RELA | 11q13.1 | 16 | 2.05 | 1 | 0.61 | 4 | 2.12 | 0.437 | 0.763 | (C) BRAF Class 3 |
| JUN | 1p32.1 | 23 | 1.6 | 3 | 0.9 | 3 | 0.9 | 0.438 | 0.763 | (A) BRAF Class 1 v12 |
| KCNQ1 | .p15.5-p15 | 25 | 4.3 | 4 | 3.05 | 9 | 6.16 | 0.438 | 0.763 | (C) BRAF Class 3 |
| RNF8 | 6p21.2 | 5 | 0.86 | 0 | 0 | 2 | 1.37 | 0.439 | 0.763 | (C) BRAF Class 3 |
| EGFL7 | 9q34.3 | 15 | 1.27 | 1 | 0.37 | 3 | 1.17 | 0.44 | 0.763 | (A) BRAF Class 1 v12 |

| | | | | | | | | | | |
|---|---|---|---|---|---|---|---|---|---|---|
| RAC2 | 22q13.1 | 6 | 1.13 | 0 | 0 | 1 | 0.78 | 0.44 | 0.763 | (A) BRAF Class 1 v12 |
| AJUBA | 14q11.2 | 2 | 1.28 | 1 | 4 | 0 | 0 | 0.44 | 0.763 | (B) BRAF Class 2 |
| SHQ1 | 3p13 | 24 | 2.03 | 9 | 3.31 | 6 | 2.34 | 0.442 | 0.765 | (B) BRAF Class 2 |
| MTOR | 1p36.22 | 145 | 5.56 | 37 | 5.89 | 42 | 6.9 | 0.445 | 0.768 | (C) BRAF Class 3 |
| EGR1 | 5q31.2 | 1 | 0.87 | 0 | 0 | 1 | 3.57 | 0.445 | 0.768 | (C) BRAF Class 3 |
| ACVR1B | 12q13.13 | 10 | 1.52 | 3 | 2.65 | 4 | 2.9 | 0.446 | 0.768 | (C) BRAF Class 3 |
| ARID5B | 10q21.2 | 50 | 3.79 | 10 | 3.38 | 15 | 5.24 | 0.447 | 0.768 | (C) BRAF Class 3 |
| TFE3 | Xp11.23 | 30 | 2.64 | 3 | 1.41 | 8 | 3.23 | 0.45 | 0.772 | (C) BRAF Class 3 |
| XRCC6 | 22q13.2 | 11 | 1.89 | 2 | 1.53 | 5 | 3.42 | 0.453 | 0.775 | (C) BRAF Class 3 |
| GRM3 | 21.11-q21. | 58 | 8.81 | 6 | 5.31 | 11 | 7.97 | 0.453 | 0.775 | (A) BRAF Class 1 v12 |
| EPHB1 | 3q22.2 | 64 | 4.08 | 17 | 4.75 | 20 | 5.52 | 0.454 | 0.775 | (C) BRAF Class 3 |
| SRP19 | 5q22.2 | 3 | 0.82 | 0 | 0 | 0 | 0 | 0.454 | 0.775 | (A) BRAF Class 1 v12 |
| MTR | 1q43 | 2 | 1.71 | 1 | 5.26 | 0 | 0 | 0.455 | 0.775 | (B) BRAF Class 2 |
| FAAP100 | 17q25.3 | 24 | 4.12 | 4 | 3.05 | 3 | 2.05 | 0.455 | 0.775 | (A) BRAF Class 1 v12 |
| PMS2 | 7p22.1 | 53 | 2.14 | 12 | 2.17 | 17 | 3 | 0.457 | 0.777 | (C) BRAF Class 3 |
| FAN1 | 15q13.3 | 15 | 2.58 | 6 | 4.58 | 5 | 3.42 | 0.459 | 0.779 | (B) BRAF Class 2 |
| BCL2L1 | 20q11.21 | 11 | 0.46 | 3 | 0.55 | 5 | 0.88 | 0.461 | 0.781 | (C) BRAF Class 3 |
| MGA | 15q15.1 | 182 | 10.05 | 35 | 8.6 | 46 | 11.19 | 0.463 | 0.783 | (C) BRAF Class 3 |
| MAP3K7 | 6q15 | 2 | 0.79 | 1 | 2.33 | 0 | 0 | 0.463 | 0.783 | (B) BRAF Class 2 |
| NCOA3 | 20q13.12 | 52 | 3.97 | 10 | 3.47 | 15 | 5.4 | 0.464 | 0.783 | (C) BRAF Class 3 |
| TRRAP | 7q22.1 | 1 | 0.45 | 0 | 0 | 1 | 1.69 | 0.465 | 0.783 | (C) BRAF Class 3 |
| KCNIP1 | 5q35.1 | 6 | 2.28 | 0 | 0 | 2 | 3.45 | 0.466 | 0.784 | (C) BRAF Class 3 |
| EPCAM | 2p21 | 18 | 0.91 | 4 | 0.89 | 7 | 1.53 | 0.468 | 0.785 | (C) BRAF Class 3 |
| MLLT3 | 9p21.3 | 9 | 2.47 | 1 | 2.08 | 3 | 5.26 | 0.468 | 0.785 | (C) BRAF Class 3 |
| PCDHAC2 | 5q31.3 | 2 | 4.17 | 1 | 12.5 | 0 | 0 | 0.469 | 0.785 | (B) BRAF Class 2 |
| FGF5 | 4q21.21 | 10 | 3 | 0 | 0 | 2 | 4.17 | 0.472 | 0.788 | (C) BRAF Class 3 |
| DMD | p21.2-p21. | 12 | 3.4 | 4 | 5.88 | 4 | 5.8 | 0.472 | 0.788 | (B) BRAF Class 2 |
| FAT1 | 4q35.2 | 262 | 12.84 | 61 | 13.32 | 71 | 14.95 | 0.473 | 0.79 | (C) BRAF Class 3 |
| FGF2 | 4q28.1 | 3 | 0.9 | 1 | 2.5 | 0 | 0 | 0.475 | 0.791 | (B) BRAF Class 2 |
| FAH | 15q25.1 | 13 | 2.23 | 1 | 0.76 | 4 | 2.74 | 0.476 | 0.791 | (C) BRAF Class 3 |
| CDK4 | 12q14.1 | 43 | 1.69 | 9 | 1.47 | 6 | 1.01 | 0.476 | 0.791 | (A) BRAF Class 1 v12 |
| ETV6 | 12p13.2 | 51 | 2.09 | 15 | 2.73 | 16 | 2.76 | 0.477 | 0.791 | (C) BRAF Class 3 |
| AR | Xq12 | 120 | 4.72 | 35 | 5.72 | 33 | 5.57 | 0.479 | 0.792 | (B) BRAF Class 2 |
| POLR2A | 17p13.1 | 4 | 4.35 | 0 | 0 | 1 | 10 | 0.479 | 0.792 | (C) BRAF Class 3 |
| RAD52 | 12p13.33 | 24 | 1.28 | 3 | 0.68 | 4 | 0.88 | 0.481 | 0.795 | (A) BRAF Class 1 v12 |
| WRN | 8p12 | 52 | 4.5 | 14 | 5.81 | 16 | 5.95 | 0.483 | 0.797 | (C) BRAF Class 3 |
| SLC34A2 | 4p15.2 | 28 | 4.12 | 7 | 4.4 | 4 | 2.27 | 0.484 | 0.797 | (B) BRAF Class 2 |
| ZBTB20 | 3q13.31 | 4 | 2.56 | 0 | 0 | 0 | 0 | 0.487 | 0.802 | (A) BRAF Class 1 v12 |
| ERF | 19q13.2 | 9 | 1.7 | 4 | 2.88 | 4 | 3.1 | 0.489 | 0.803 | (C) BRAF Class 3 |
| TAP2 | 6p21.32 | 5 | 0.94 | 3 | 2.16 | 2 | 1.55 | 0.491 | 0.805 | (B) BRAF Class 2 |
| RPTOR | 17q25.3 | 87 | 4.1 | 25 | 5.13 | 25 | 4.92 | 0.492 | 0.805 | (B) BRAF Class 2 |
| IKBKE | 1q32.1 | 22 | 1.4 | 8 | 2.23 | 5 | 1.38 | 0.495 | 0.807 | (B) BRAF Class 2 |
| CTLA4 | 2q33.2 | 26 | 1.49 | 3 | 0.74 | 5 | 1.24 | 0.495 | 0.807 | (A) BRAF Class 1 v12 |
| CDK6 | 7q21.2 | 19 | 0.75 | 6 | 1.06 | 7 | 1.19 | 0.496 | 0.807 | (C) BRAF Class 3 |
| PTCH2 | 1p34.1 | 6 | 2.51 | 0 | 0 | 2 | 3.33 | 0.496 | 0.807 | (C) BRAF Class 3 |

| | | | | | | | | | | |
|---|---|---|---|---|---|---|---|---|---|---|
| PMAIP1 | 18q21.32 | 6 | 0.51 | 0 | 0 | 1 | 0.39 | 0.497 | 0.808 | (A) BRAF Class 1 v12 |
| PDPK1 | 16p13.3 | 6 | 0.48 | 0 | 0 | 1 | 0.38 | 0.502 | 0.815 | (A) BRAF Class 1 v12 |
| CDC42 | 1p36.12 | 3 | 0.45 | 1 | 0.62 | 2 | 1.26 | 0.502 | 0.815 | (C) BRAF Class 3 |
| UBR5 | 8q22.3 | 2 | 1.49 | 0 | 0 | 1 | 4.17 | 0.506 | 0.817 | (C) BRAF Class 3 |
| CSF3R | 1p34.3 | 52 | 2.35 | 15 | 3.01 | 16 | 3.08 | 0.506 | 0.817 | (C) BRAF Class 3 |
| HIST1H3G | 6p22.2 | 6 | 0.54 | 3 | 1.18 | 2 | 0.85 | 0.506 | 0.817 | (B) BRAF Class 2 |
| FANCB | Xp22.2 | 20 | 3 | 2 | 1.33 | 4 | 2.44 | 0.508 | 0.817 | (A) BRAF Class 1 v12 |
| BCL9 | 1q21.2 | 2 | 1.64 | 0 | 0 | 1 | 4.55 | 0.508 | 0.817 | (C) BRAF Class 3 |
| LMO1 | 11p15.4 | 17 | 0.75 | 6 | 1.18 | 3 | 0.57 | 0.508 | 0.817 | (B) BRAF Class 2 |
| DYNC1I1 | 7q21.3 | 2 | 2.33 | 1 | 8.33 | 1 | 5 | 0.509 | 0.817 | (B) BRAF Class 2 |
| TGFBR1 | 9q22.33 | 20 | 1.69 | 6 | 2.21 | 7 | 2.73 | 0.511 | 0.818 | (C) BRAF Class 3 |
| BCR | 22q11.23 | 22 | 3.82 | 2 | 1.96 | 3 | 2.4 | 0.511 | 0.818 | (A) BRAF Class 1 v12 |
| CSNK1A1 | 5q32 | 5 | 1.5 | 0 | 0 | 0 | 0 | 0.512 | 0.818 | (A) BRAF Class 1 v12 |
| FGF9 | 13q12.11 | 5 | 1.5 | 0 | 0 | 0 | 0 | 0.512 | 0.818 | (A) BRAF Class 1 v12 |
| MTA1 | 14q32.33 | 25 | 4.3 | 3 | 2.29 | 7 | 4.79 | 0.513 | 0.818 | (C) BRAF Class 3 |
| CD276 | 15q24.1 | 19 | 1.61 | 2 | 0.74 | 3 | 1.17 | 0.515 | 0.818 | (A) BRAF Class 1 v12 |
| CDH11 | 16q21 | 2 | 1.64 | 0 | 0 | 1 | 4.55 | 0.515 | 0.818 | (C) BRAF Class 3 |
| TRIM33 | 1p13.2 | 2 | 1.64 | 0 | 0 | 1 | 4.55 | 0.515 | 0.818 | (C) BRAF Class 3 |
| KNSTRN | 15q15.1 | 20 | 2.69 | 4 | 2.31 | 2 | 1.19 | 0.515 | 0.818 | (A) BRAF Class 1 v12 |
| CAMTA1 | 36.31-p36. | 2 | 1.25 | 1 | 3.57 | 0 | 0 | 0.52 | 0.824 | (B) BRAF Class 2 |
| TYRO3 | 15q15.1 | 2 | 1.48 | 0 | 0 | 0 | 0 | 0.522 | 0.826 | (A) BRAF Class 1 v12 |
| SPEN | 36.21-p36. | 142 | 9.08 | 26 | 7.26 | 30 | 8.29 | 0.524 | 0.828 | (A) BRAF Class 1 v12 |
| ETV4 | 17q21.31 | 19 | 1.3 | 3 | 1.01 | 2 | 0.59 | 0.527 | 0.832 | (A) BRAF Class 1 v12 |
| MSH2 | 2p21-p16.3 | 72 | 2.91 | 17 | 3.09 | 22 | 3.82 | 0.528 | 0.832 | (C) BRAF Class 3 |
| SLITRK6 | 13q31.1 | 12 | 3.67 | 2 | 3.45 | 5 | 6.41 | 0.529 | 0.832 | (C) BRAF Class 3 |
| NFKBIA | 14q13.2 | 26 | 1.09 | 3 | 0.56 | 6 | 1.08 | 0.532 | 0.835 | (A) BRAF Class 1 v12 |
| NKX2-1 | 14q13.3 | 58 | 2.36 | 15 | 2.71 | 10 | 1.73 | 0.532 | 0.835 | (B) BRAF Class 2 |
| ZBTB2 | 6q25.1 | 3 | 0.61 | 1 | 1.37 | 0 | 0 | 0.533 | 0.835 | (B) BRAF Class 2 |
| ADGRA2 | 8p11.23 | 35 | 5.64 | 7 | 7.22 | 10 | 8.06 | 0.533 | 0.835 | (C) BRAF Class 3 |
| ID4 | 6p22.3 | 5 | 0.86 | 1 | 0.76 | 0 | 0 | 0.535 | 0.836 | (A) BRAF Class 1 v12 |
| TTYH1 | 19q13.42 | 2 | 1.28 | 1 | 4 | 1 | 3.33 | 0.537 | 0.837 | (B) BRAF Class 2 |
| RAC1 | 7p22.1 | 50 | 2.17 | 12 | 2.29 | 16 | 2.97 | 0.537 | 0.837 | (C) BRAF Class 3 |
| RAD51 | 15q15.1 | 19 | 0.86 | 4 | 0.81 | 2 | 0.39 | 0.539 | 0.84 | (A) BRAF Class 1 v12 |
| CADM2 | 3p12.1 | 8 | 3.04 | 3 | 6.25 | 2 | 3.45 | 0.541 | 0.841 | (B) BRAF Class 2 |
| CDK1 | 10q21.2 | 2 | 0.57 | 0 | 0 | 1 | 1.45 | 0.542 | 0.841 | (C) BRAF Class 3 |
| HSP90AA1 | 14q32.31 | 8 | 1.5 | 1 | 1.22 | 3 | 2.97 | 0.542 | 0.841 | (C) BRAF Class 3 |
| WT1 | 11p13 | 68 | 2.73 | 17 | 3.04 | 12 | 2.05 | 0.549 | 0.849 | (B) BRAF Class 2 |
| ENG | 9q34.11 | 13 | 2.23 | 1 | 0.76 | 3 | 2.05 | 0.55 | 0.849 | (A) BRAF Class 1 v12 |
| CDKN1B | 12p13.1 | 35 | 1.38 | 5 | 0.89 | 6 | 1.02 | 0.55 | 0.849 | (A) BRAF Class 1 v12 |
| KDM5C | Xp11.22 | 80 | 3.37 | 21 | 3.94 | 24 | 4.24 | 0.55 | 0.849 | (C) BRAF Class 3 |
| FUS | 16p11.2 | 16 | 1.51 | 5 | 2.26 | 6 | 2.33 | 0.552 | 0.85 | (C) BRAF Class 3 |
| CARM1 | 19p13.2 | 9 | 1.7 | 1 | 0.72 | 1 | 0.78 | 0.552 | 0.85 | (A) BRAF Class 1 v12 |
| MAP3K2 | 2q14.3 | 3 | 1.92 | 1 | 4 | 0 | 0 | 0.556 | 0.852 | (B) BRAF Class 2 |
| CDK9 | 9q34.11 | 3 | 0.85 | 0 | 0 | 0 | 0 | 0.557 | 0.852 | (A) BRAF Class 1 v12 |
| PSMD13 | 11p15.5 | 3 | 0.85 | 0 | 0 | 0 | 0 | 0.557 | 0.852 | (A) BRAF Class 1 v12 |

| | | | | | | | | | | |
|---|---|---|---|---|---|---|---|---|---|---|
| PHF6 | Xq26.2 | 20 | 1.33 | 6 | 2.07 | 6 | 1.83 | 0.557 | 0.852 | (B) BRAF Class 2 |
| UPF1 | 19p13.11 | 15 | 2.83 | 4 | 2.88 | 6 | 4.65 | 0.557 | 0.852 | (C) BRAF Class 3 |
| RAD21 | 8q24.11 | 57 | 2.5 | 9 | 1.76 | 11 | 2.11 | 0.561 | 0.856 | (A) BRAF Class 1 v12 |
| FRS2 | 12q15 | 6 | 0.95 | 2 | 2.06 | 2 | 1.64 | 0.564 | 0.86 | (B) BRAF Class 2 |
| CYSLTR2 | 13q14.2 | 7 | 1.06 | 3 | 1.8 | 3 | 1.99 | 0.566 | 0.861 | (C) BRAF Class 3 |
| MDM4 | 1q32.1 | 29 | 1.14 | 9 | 1.6 | 9 | 1.54 | 0.566 | 0.861 | (B) BRAF Class 2 |
| IDH2 | 15q26.1 | 47 | 1.48 | 7 | 0.97 | 10 | 1.4 | 0.569 | 0.865 | (A) BRAF Class 1 v12 |
| RPL22 | 1p36.31 | 3 | 5.66 | 0 | 0 | 0 | 0 | 0.571 | 0.865 | (A) BRAF Class 1 v12 |
| CCND3 | 6p21.1 | 21 | 0.83 | 6 | 1.07 | 3 | 0.51 | 0.572 | 0.866 | (B) BRAF Class 2 |
| INHA | 2q35 | 22 | 1.97 | 3 | 1.18 | 3 | 1.28 | 0.573 | 0.866 | (A) BRAF Class 1 v12 |
| FGF7 | 15q21.2 | 4 | 0.92 | 1 | 1.54 | 0 | 0 | 0.573 | 0.866 | (B) BRAF Class 2 |
| KAT6B | 10q22.2 | 43 | 6.3 | 13 | 8.67 | 11 | 6.55 | 0.576 | 0.868 | (B) BRAF Class 2 |
| FANCM | 14q21.2 | 39 | 5.22 | 9 | 5.39 | 14 | 7.14 | 0.576 | 0.868 | (C) BRAF Class 3 |
| CAMK1 | 3p25.3 | 2 | 0.06 | 0 | 0 | 1 | 0.13 | 0.577 | 0.868 | (C) BRAF Class 3 |
| TRBC2 | 7q34 | 2 | 0.06 | 0 | 0 | 1 | 0.13 | 0.577 | 0.868 | (C) BRAF Class 3 |
| RINT1 | 7q22.3 | 13 | 2.23 | 5 | 3.82 | 4 | 2.74 | 0.578 | 0.868 | (B) BRAF Class 2 |
| DDB2 | 11p11.2 | 15 | 1.47 | 2 | 0.92 | 5 | 2.09 | 0.582 | 0.871 | (C) BRAF Class 3 |
| GPS2 | 17p13.1 | 13 | 1.14 | 3 | 1.15 | 1 | 0.41 | 0.582 | 0.871 | (B) BRAF Class 2 |
| CD22 | 19q13.12 | 2 | 1.72 | 0 | 0 | 0 | 0 | 0.582 | 0.871 | (A) BRAF Class 1 v12 |
| RTEL1 | 20q13.33 | 22 | 4.13 | 7 | 5 | 8 | 6.2 | 0.585 | 0.872 | (C) BRAF Class 3 |
| FGF1 | 5q31.3 | 4 | 1.2 | 0 | 0 | 0 | 0 | 0.586 | 0.872 | (A) BRAF Class 1 v12 |
| CACNA1E | 1q25.3 | 3 | 6.25 | 1 | 12.5 | 0 | 0 | 0.587 | 0.872 | (B) BRAF Class 2 |
| TGFB1 | 19q13.2 | 3 | 6.25 | 0 | 0 | 1 | 12.5 | 0.587 | 0.872 | (C) BRAF Class 3 |
| GATA2 | 3q21.3 | 50 | 2.29 | 8 | 1.66 | 9 | 1.79 | 0.587 | 0.872 | (A) BRAF Class 1 v12 |
| MED12 | Xq13.1 | 118 | 5.36 | 27 | 4.95 | 33 | 6.31 | 0.589 | 0.872 | (C) BRAF Class 3 |
| CD274 | 9p24.1 | 20 | 0.83 | 4 | 0.73 | 7 | 1.24 | 0.59 | 0.872 | (C) BRAF Class 3 |
| BABAM2 | 2p23.2 | 14 | 2.41 | 2 | 1.53 | 5 | 3.42 | 0.59 | 0.872 | (C) BRAF Class 3 |
| FIP1L1 | 4q12 | 2 | 1.31 | 1 | 3.03 | 1 | 3.85 | 0.592 | 0.872 | (C) BRAF Class 3 |
| UMODL1 | 21q22.3 | 3 | 6.25 | 0 | 0 | 0 | 0 | 0.592 | 0.872 | (A) BRAF Class 1 v12 |
| PRF1 | 10q22.1 | 29 | 3.13 | 4 | 1.99 | 5 | 2.29 | 0.593 | 0.872 | (A) BRAF Class 1 v12 |
| RPS6KA2 | 6q27 | 3 | 2.56 | 0 | 0 | 0 | 0 | 0.593 | 0.872 | (A) BRAF Class 1 v12 |
| TNK2 | 3q29 | 3 | 2.56 | 0 | 0 | 0 | 0 | 0.593 | 0.872 | (A) BRAF Class 1 v12 |
| LTK | 15q15.1 | 3 | 1.3 | 1 | 1.75 | 0 | 0 | 0.594 | 0.872 | (B) BRAF Class 2 |
| OGG1 | 3p25.3 | 4 | 0.69 | 1 | 0.76 | 0 | 0 | 0.594 | 0.872 | (B) BRAF Class 2 |
| KMT2B | 19q13.12 | 121 | 11.62 | 20 | 9.43 | 27 | 12.27 | 0.598 | 0.876 | (C) BRAF Class 3 |
| HIST1H3A | 6p22.2 | 2 | 0.18 | 1 | 0.39 | 0 | 0 | 0.599 | 0.877 | (B) BRAF Class 2 |
| NEIL1 | 15q24.2 | 9 | 1.55 | 1 | 0.76 | 1 | 0.68 | 0.603 | 0.881 | (A) BRAF Class 1 v12 |
| MITF | 3p13 | 64 | 2.6 | 12 | 2.17 | 18 | 3.12 | 0.604 | 0.882 | (C) BRAF Class 3 |
| XPA | 9q22.33 | 12 | 1.17 | 1 | 0.46 | 2 | 0.84 | 0.605 | 0.882 | (A) BRAF Class 1 v12 |
| MYOD1 | 11p15.1 | 29 | 2.31 | 4 | 1.4 | 5 | 1.88 | 0.606 | 0.883 | (A) BRAF Class 1 v12 |
| CHD1 | 5q15-q21.1 | 6 | 3.85 | 0 | 0 | 1 | 3.33 | 0.608 | 0.885 | (A) BRAF Class 1 v12 |
| DOT1L | 19p13.3 | 78 | 4.99 | 15 | 4.19 | 21 | 5.8 | 0.611 | 0.888 | (C) BRAF Class 3 |
| HIST1H1C | 6p22.2 | 25 | 2 | 5 | 1.75 | 3 | 1.11 | 0.613 | 0.89 | (A) BRAF Class 1 v12 |
| BTG1 | 12q21.33 | 2 | 0.3 | 1 | 0.84 | 1 | 0.68 | 0.617 | 0.892 | (B) BRAF Class 2 |
| ERCC5 | 13q33.1 | 73 | 3.36 | 17 | 3.5 | 21 | 4.26 | 0.618 | 0.892 | (C) BRAF Class 3 |

| | | | | | | | | | | |
|---|---|---|---|---|---|---|---|---|---|---|
| THADA | 2p21 | 3 | 1.94 | 0 | 0 | 0 | 0 | 0.618 | 0.892 | (A) BRAF Class 1 v12 |
| BRCC3 | Xq28 | 12 | 2.03 | 4 | 3.01 | 2 | 1.35 | 0.619 | 0.892 | (B) BRAF Class 2 |
| ACVR1 | 2q24.1 | 22 | 1.17 | 3 | 0.72 | 6 | 1.42 | 0.62 | 0.892 | (C) BRAF Class 3 |
| FAAP24 | 19q13.11 | 2 | 0.34 | 0 | 0 | 0 | 0 | 0.621 | 0.892 | (A) BRAF Class 1 v12 |
| TAZ | Xq28 | 2 | 0.34 | 0 | 0 | 0 | 0 | 0.621 | 0.892 | (A) BRAF Class 1 v12 |
| FUBP1 | 1p31.1 | 28 | 1.85 | 5 | 1.45 | 4 | 1.16 | 0.623 | 0.892 | (A) BRAF Class 1 v12 |
| CRLF2 | .33 and Yp | 31 | 1.31 | 7 | 1.33 | 10 | 1.85 | 0.623 | 0.892 | (C) BRAF Class 3 |
| FAT3 | 11q14.3 | 21 | 8.14 | 6 | 12 | 7 | 10.45 | 0.623 | 0.892 | (B) BRAF Class 2 |
| ABL2 | 1q25.2 | 20 | 3.01 | 5 | 4.81 | 4 | 3.08 | 0.624 | 0.892 | (B) BRAF Class 2 |
| BCORL1 | Xq26.1 | 114 | 7.06 | 18 | 5.57 | 26 | 6.95 | 0.624 | 0.892 | (A) BRAF Class 1 v12 |
| PAK3 | Xq23 | 17 | 2.49 | 4 | 3.7 | 5 | 3.65 | 0.626 | 0.892 | (B) BRAF Class 2 |
| PIK3CB | 3q22.3 | 48 | 3.23 | 8 | 2.37 | 12 | 3.6 | 0.626 | 0.892 | (C) BRAF Class 3 |
| CD58 | 1p13.1 | 3 | 0.7 | 1 | 1.2 | 0 | 0 | 0.626 | 0.892 | (B) BRAF Class 2 |
| VHL | 3p25.3 | 26 | 0.83 | 8 | 1.21 | 6 | 0.86 | 0.63 | 0.896 | (B) BRAF Class 2 |
| AKT2 | 19q13.2 | 50 | 1.94 | 13 | 2.29 | 9 | 1.52 | 0.631 | 0.896 | (B) BRAF Class 2 |
| HMBS | 11q23.3 | 7 | 1.2 | 3 | 2.29 | 2 | 1.37 | 0.632 | 0.897 | (B) BRAF Class 2 |
| FN1 | 2q35 | 3 | 2.56 | 0 | 0 | 1 | 4.76 | 0.634 | 0.898 | (C) BRAF Class 3 |
| FOXP4 | 6p21.1 | 3 | 2.56 | 0 | 0 | 1 | 4.76 | 0.634 | 0.898 | (C) BRAF Class 3 |
| SAMD9 | 7q21.2 | 4 | 2.14 | 0 | 0 | 1 | 3.23 | 0.635 | 0.898 | (C) BRAF Class 3 |
| FYN | 6q21 | 26 | 2.06 | 6 | 2.14 | 8 | 3 | 0.64 | 0.902 | (C) BRAF Class 3 |
| EPB41L4A | q22.1-q22. | 2 | 0.06 | 0 | 0 | 0 | 0 | 0.64 | 0.902 | (A) BRAF Class 1 v12 |
| H19 | 11p15.5 | 2 | 0.06 | 0 | 0 | 0 | 0 | 0.64 | 0.902 | (A) BRAF Class 1 v12 |
| NUTM1 | 15q14 | 29 | 5.06 | 5 | 5.38 | 8 | 7.27 | 0.643 | 0.906 | (C) BRAF Class 3 |
| PAK1 | .q13.5-q14 | 14 | 1.06 | 5 | 1.69 | 3 | 1.05 | 0.645 | 0.907 | (B) BRAF Class 2 |
| HIST1H3F | 6p22.2 | 4 | 0.36 | 2 | 0.79 | 1 | 0.43 | 0.646 | 0.907 | (B) BRAF Class 2 |
| GSTM5 | 1p13.3 | 4 | 1.13 | 0 | 0 | 1 | 1.45 | 0.647 | 0.907 | (C) BRAF Class 3 |
| NAB2 | 12q13.3 | 13 | 3.9 | 1 | 2.5 | 3 | 6.25 | 0.649 | 0.909 | (C) BRAF Class 3 |
| IRF2 | 4q35.1 | 6 | 1.15 | 2 | 2.22 | 2 | 1.85 | 0.652 | 0.912 | (B) BRAF Class 2 |
| CDH20 | 18q21.33 | 2 | 1 | 0 | 0 | 1 | 1.96 | 0.654 | 0.913 | (C) BRAF Class 3 |
| ABRAXAS1 | 4q21.23 | 17 | 0.89 | 2 | 0.46 | 4 | 0.88 | 0.655 | 0.913 | (A) BRAF Class 1 v12 |
| TCF3 | 19p13.3 | 65 | 3.07 | 12 | 2.55 | 17 | 3.58 | 0.655 | 0.913 | (C) BRAF Class 3 |
| ZFPM1 | 16q24.2 | 3 | 3.26 | 1 | 6.67 | 0 | 0 | 0.657 | 0.913 | (B) BRAF Class 2 |
| NRIP1 | .q11.2-q21 | 3 | 3.26 | 0 | 0 | 0 | 0 | 0.658 | 0.913 | (A) BRAF Class 1 v12 |
| PHOX2B | 4p13 | 48 | 2.08 | 10 | 1.96 | 14 | 2.68 | 0.66 | 0.913 | (C) BRAF Class 3 |
| FGFR3 | 4p16.3 | 90 | 2.69 | 24 | 3.2 | 23 | 3.12 | 0.66 | 0.913 | (B) BRAF Class 2 |
| PIK3R2 | 19p13.11 | 34 | 2.21 | 5 | 1.47 | 8 | 2.32 | 0.661 | 0.913 | (C) BRAF Class 3 |
| XIAP | Xq25 | 13 | 1.1 | 2 | 0.74 | 4 | 1.56 | 0.662 | 0.913 | (C) BRAF Class 3 |
| CBFB | 16q22.1 | 16 | 0.76 | 3 | 0.62 | 2 | 0.4 | 0.663 | 0.913 | (A) BRAF Class 1 v12 |
| BRINP3 | 1q31.1 | 5 | 4.24 | 1 | 4.76 | 0 | 0 | 0.663 | 0.913 | (B) BRAF Class 2 |
| HDAC4 | 2q37.3 | 3 | 1.35 | 0 | 0 | 1 | 2.27 | 0.663 | 0.913 | (C) BRAF Class 3 |
| STAT3 | 17q21.2 | 46 | 1.88 | 8 | 1.47 | 8 | 1.43 | 0.663 | 0.913 | (A) BRAF Class 1 v12 |
| KMT5A | 12q24.31 | 3 | 0.54 | 1 | 0.68 | 0 | 0 | 0.667 | 0.913 | (B) BRAF Class 2 |
| ABL1 | 9q34.12 | 88 | 2.8 | 19 | 2.88 | 24 | 3.43 | 0.669 | 0.913 | (C) BRAF Class 3 |
| EIF4A2 | 3q27.3 | 11 | 0.98 | 4 | 1.56 | 2 | 0.85 | 0.676 | 0.913 | (B) BRAF Class 2 |
| RPL26 | 17p13.1 | 2 | 0.57 | 0 | 0 | 0 | 0 | 0.677 | 0.913 | (A) BRAF Class 1 v12 |

| | | | | | | | | | | |
|---|---|---|---|---|---|---|---|---|---|---|
| SRSF2 | 17q25.1 | 27 | 1.2 | 4 | 0.81 | 7 | 1.38 | 0.68 | 0.913 | (C) BRAF Class 3 |
| ARHGEF12 | 11q23.3 | 25 | 4.26 | 8 | 6.02 | 7 | 4.76 | 0.681 | 0.913 | (B) BRAF Class 2 |
| USP7 | 16p13.2 | 2 | 2.33 | 0 | 0 | 0 | 0 | 0.685 | 0.913 | (A) BRAF Class 1 v12 |
| NPRL2 | 3p21.31 | 3 | 1.14 | 0 | 0 | 1 | 1.72 | 0.685 | 0.913 | (C) BRAF Class 3 |
| CLDN18 | 3q22.3 | 4 | 2.56 | 0 | 0 | 1 | 3.33 | 0.686 | 0.913 | (C) BRAF Class 3 |
| ASPSCR1 | 17q25.3 | 2 | 1.72 | 0 | 0 | 0 | 0 | 0.687 | 0.913 | (A) BRAF Class 1 v12 |
| SS18 | 18q11.2 | 8 | 1.01 | 3 | 1.78 | 2 | 1.04 | 0.688 | 0.913 | (B) BRAF Class 2 |
| MAP2K4 | 17p12 | 41 | 1.62 | 12 | 2.14 | 10 | 1.71 | 0.69 | 0.913 | (B) BRAF Class 2 |
| DUSP2 | 2q11.2 | 2 | 0.91 | 0 | 0 | 0 | 0 | 0.69 | 0.913 | (A) BRAF Class 1 v12 |
| ICOSLG | 21q22.3 | 16 | 1.35 | 5 | 1.84 | 5 | 1.95 | 0.695 | 0.913 | (C) BRAF Class 3 |
| SMARCE1 | 17q21.2 | 5 | 0.74 | 2 | 1.32 | 2 | 1.26 | 0.699 | 0.913 | (B) BRAF Class 2 |
| CDKN2C | 1p32.3 | 14 | 0.57 | 3 | 0.54 | 5 | 0.87 | 0.699 | 0.913 | (C) BRAF Class 3 |
| KNL1 | 15q15.1 | 2 | 1.64 | 0 | 0 | 0 | 0 | 0.7 | 0.913 | (A) BRAF Class 1 v12 |
| EPHB6 | 7q34 | 2 | 1 | 1 | 2.5 | 1 | 1.96 | 0.701 | 0.913 | (B) BRAF Class 2 |
| LATS2 | 13q12.11 | 49 | 4.07 | 10 | 3.57 | 13 | 4.98 | 0.701 | 0.913 | (C) BRAF Class 3 |
| ERCC6 | 10q11.23 | 34 | 5.84 | 10 | 7.63 | 8 | 5.48 | 0.703 | 0.913 | (B) BRAF Class 2 |
| MAX | 14q23.3 | 15 | 0.79 | 5 | 1.17 | 3 | 0.71 | 0.704 | 0.913 | (B) BRAF Class 2 |
| BCL10 | 1p22.3 | 17 | 1.4 | 3 | 1.1 | 5 | 1.95 | 0.704 | 0.913 | (C) BRAF Class 3 |
| DIS3L2 | 2q37.1 | 19 | 2.91 | 5 | 3.47 | 3 | 1.92 | 0.705 | 0.913 | (B) BRAF Class 2 |
| GUCY1A2 | 11q22.3 | 2 | 1.71 | 0 | 0 | 0 | 0 | 0.707 | 0.913 | (A) BRAF Class 1 v12 |
| NLRP1 | 17p13.2 | 2 | 1.71 | 0 | 0 | 0 | 0 | 0.707 | 0.913 | (A) BRAF Class 1 v12 |
| STK36 | 2q35 | 2 | 1.71 | 0 | 0 | 0 | 0 | 0.707 | 0.913 | (A) BRAF Class 1 v12 |
| TRIM24 | 7q33-q34 | 2 | 1.71 | 0 | 0 | 0 | 0 | 0.707 | 0.913 | (A) BRAF Class 1 v12 |
| MYCN | 2p24.3 | 43 | 1.69 | 11 | 1.8 | 13 | 2.2 | 0.708 | 0.913 | (C) BRAF Class 3 |
| FANCC | 9q22.32 | 38 | 1.54 | 6 | 1.08 | 9 | 1.56 | 0.709 | 0.913 | (C) BRAF Class 3 |
| NBN | 8q21.3 | 77 | 3.05 | 19 | 3.38 | 15 | 2.56 | 0.711 | 0.913 | (B) BRAF Class 2 |
| FAS | 10q23.31 | 18 | 1.23 | 2 | 0.67 | 4 | 1.19 | 0.711 | 0.913 | (A) BRAF Class 1 v12 |
| BUB1B | 15q15.1 | 24 | 2.35 | 7 | 3.21 | 7 | 2.93 | 0.711 | 0.913 | (B) BRAF Class 2 |
| NEIL2 | 8p23.1 | 4 | 0.69 | 1 | 0.76 | 2 | 1.37 | 0.713 | 0.913 | (C) BRAF Class 3 |
| SAMD9L | 7q21.2 | 4 | 4.35 | 1 | 6.67 | 1 | 10 | 0.713 | 0.913 | (C) BRAF Class 3 |
| PLCG2 | 16q23.3 | 51 | 3.91 | 11 | 3.73 | 14 | 4.88 | 0.721 | 0.913 | (C) BRAF Class 3 |
| DNAH9 | 17p12 | 6 | 8.7 | 2 | 12.5 | 2 | 15.38 | 0.725 | 0.913 | (C) BRAF Class 3 |
| PIK3C3 | 18q12.3 | 33 | 2.51 | 6 | 1.97 | 9 | 2.99 | 0.725 | 0.913 | (C) BRAF Class 3 |
| GLI3 | 7p14.1 | 26 | 7.37 | 5 | 7.35 | 7 | 10.14 | 0.726 | 0.913 | (C) BRAF Class 3 |
| BIRC3 | 11q22.2 | 15 | 1.23 | 5 | 1.81 | 4 | 1.55 | 0.729 | 0.913 | (B) BRAF Class 2 |
| DNMT1 | 19p13.2 | 54 | 4.1 | 15 | 5.07 | 13 | 4.66 | 0.732 | 0.913 | (B) BRAF Class 2 |
| EPHB4 | 7q22.1 | 4 | 1.33 | 1 | 1.43 | 2 | 2.56 | 0.734 | 0.913 | (C) BRAF Class 3 |
| SMARCB1 | 11.23\|22 | 41 | 1.28 | 11 | 1.65 | 10 | 1.42 | 0.739 | 0.913 | (B) BRAF Class 2 |
| EMSY | 11q13.5 | 15 | 2.12 | 4 | 3.2 | 4 | 2.6 | 0.74 | 0.913 | (B) BRAF Class 2 |
| SOCS1 | 16p13.13 | 26 | 1.06 | 4 | 0.72 | 5 | 0.87 | 0.74 | 0.913 | (A) BRAF Class 1 v12 |
| RECQL | 12p12.1 | 14 | 2.64 | 4 | 2.88 | 2 | 1.55 | 0.741 | 0.913 | (B) BRAF Class 2 |
| HNF1A | 12q24.31 | 106 | 3.35 | 24 | 3.62 | 20 | 2.89 | 0.746 | 0.913 | (B) BRAF Class 2 |
| RRAGC | 1p34.3 | 5 | 0.75 | 2 | 1.23 | 2 | 1.26 | 0.746 | 0.913 | (C) BRAF Class 3 |
| CSMD3 | 8q23.3 | 3 | 2.56 | 1 | 5.26 | 1 | 4.76 | 0.748 | 0.913 | (B) BRAF Class 2 |
| ITGA10 | 1q21.1 | 3 | 2.56 | 1 | 5.26 | 1 | 4.76 | 0.748 | 0.913 | (B) BRAF Class 2 |

| | | | | | | | | | | |
|---|---|---|---|---|---|---|---|---|---|---|
| ABCA17P | 16p13.3 | 1 | 0.03 | 0 | 0 | 0 | 0 | 0.8 | 0.913 | (A) BRAF Class 1 v12 |
| ABR | 17p13.3 | 1 | 0.03 | 0 | 0 | 0 | 0 | 0.8 | 0.913 | (A) BRAF Class 1 v12 |
| ANKLE2 | 12q24.33 | 1 | 0.03 | 0 | 0 | 0 | 0 | 0.8 | 0.913 | (A) BRAF Class 1 v12 |
| ANKRD30A | 10p11.21 | 1 | 0.03 | 0 | 0 | 0 | 0 | 0.8 | 0.913 | (A) BRAF Class 1 v12 |
| AP1G1 | 16q22.2 | 1 | 0.03 | 0 | 0 | 0 | 0 | 0.8 | 0.913 | (A) BRAF Class 1 v12 |
| APBA1 | 9q21.12 | 1 | 0.03 | 0 | 0 | 0 | 0 | 0.8 | 0.913 | (A) BRAF Class 1 v12 |
| ARHGAP4 | Xq28 | 1 | 0.03 | 0 | 0 | 0 | 0 | 0.8 | 0.913 | (A) BRAF Class 1 v12 |
| ARSB | 5q14.1 | 1 | 0.03 | 0 | 0 | 0 | 0 | 0.8 | 0.913 | (A) BRAF Class 1 v12 |
| CASZ1 | 1p36.22 | 1 | 0.03 | 0 | 0 | 0 | 0 | 0.8 | 0.913 | (A) BRAF Class 1 v12 |
| CLIP1 | 12q24.31 | 1 | 0.03 | 0 | 0 | 0 | 0 | 0.8 | 0.913 | (A) BRAF Class 1 v12 |
| CPSF1 | 8q24.3 | 1 | 0.03 | 0 | 0 | 0 | 0 | 0.8 | 0.913 | (A) BRAF Class 1 v12 |
| CTDSP1 | 2q35 | 1 | 0.03 | 0 | 0 | 0 | 0 | 0.8 | 0.913 | (A) BRAF Class 1 v12 |
| CTRC | 1p36.21 | 1 | 0.03 | 0 | 0 | 0 | 0 | 0.8 | 0.913 | (A) BRAF Class 1 v12 |
| CTSB | 8p23.1 | 1 | 0.03 | 0 | 0 | 0 | 0 | 0.8 | 0.913 | (A) BRAF Class 1 v12 |
| CUL1 | 7q36.1 | 1 | 0.03 | 0 | 0 | 0 | 0 | 0.8 | 0.913 | (A) BRAF Class 1 v12 |
| DCBLD1 | 6q22.1 | 1 | 0.03 | 0 | 0 | 0 | 0 | 0.8 | 0.913 | (A) BRAF Class 1 v12 |
| DDAH1 | 1p22.3 | 1 | 0.03 | 0 | 0 | 0 | 0 | 0.8 | 0.913 | (A) BRAF Class 1 v12 |
| DEF8 | 16q24.3 | 1 | 0.03 | 0 | 0 | 0 | 0 | 0.8 | 0.913 | (A) BRAF Class 1 v12 |
| DENND4C | 9p22.1 | 1 | 0.03 | 0 | 0 | 0 | 0 | 0.8 | 0.913 | (A) BRAF Class 1 v12 |
| DHRSX | .33 and Yp | 1 | 0.03 | 0 | 0 | 0 | 0 | 0.8 | 0.913 | (A) BRAF Class 1 v12 |
| DRICH1 | 22q11.23 | 1 | 0.03 | 0 | 0 | 0 | 0 | 0.8 | 0.913 | (A) BRAF Class 1 v12 |
| EIF3E | 8q23.1 | 1 | 0.03 | 0 | 0 | 0 | 0 | 0.8 | 0.913 | (A) BRAF Class 1 v12 |
| ELAC2 | 17p12 | 1 | 0.03 | 0 | 0 | 0 | 0 | 0.8 | 0.913 | (A) BRAF Class 1 v12 |
| EMID1 | 22q12.2 | 1 | 0.03 | 0 | 0 | 0 | 0 | 0.8 | 0.913 | (A) BRAF Class 1 v12 |
| F11-AS1 | 4q35.2 | 1 | 0.03 | 0 | 0 | 0 | 0 | 0.8 | 0.913 | (A) BRAF Class 1 v12 |
| FKBP9P1 | 7p11.2 | 1 | 0.03 | 0 | 0 | 0 | 0 | 0.8 | 0.913 | (A) BRAF Class 1 v12 |
| GALNT11 | 36.1\|7q36 | 1 | 0.03 | 0 | 0 | 0 | 0 | 0.8 | 0.913 | (A) BRAF Class 1 v12 |
| GOLT1A | 1q32.1 | 1 | 0.03 | 0 | 0 | 0 | 0 | 0.8 | 0.913 | (A) BRAF Class 1 v12 |
| GRAPL | 17p11.2 | 1 | 0.03 | 0 | 0 | 0 | 0 | 0.8 | 0.913 | (A) BRAF Class 1 v12 |
| GRIA1 | 5q33.2 | 1 | 0.03 | 0 | 0 | 0 | 0 | 0.8 | 0.913 | (A) BRAF Class 1 v12 |
| GRIK1 | 21q21.3 | 1 | 0.03 | 0 | 0 | 0 | 0 | 0.8 | 0.913 | (A) BRAF Class 1 v12 |
| HECTD4 | 12q24.13 | 1 | 0.03 | 0 | 0 | 0 | 0 | 0.8 | 0.913 | (A) BRAF Class 1 v12 |
| HLA-DOB | 6p21.32 | 1 | 0.03 | 0 | 0 | 0 | 0 | 0.8 | 0.913 | (A) BRAF Class 1 v12 |
| HOOK2 | 19p13.13 | 1 | 0.03 | 0 | 0 | 0 | 0 | 0.8 | 0.913 | (A) BRAF Class 1 v12 |
| HSD17B6 | 12q13.3 | 1 | 0.03 | 0 | 0 | 0 | 0 | 0.8 | 0.913 | (A) BRAF Class 1 v12 |
| IGHE | 14q32.33 | 1 | 0.03 | 0 | 0 | 0 | 0 | 0.8 | 0.913 | (A) BRAF Class 1 v12 |
| LDLR | 19p13.2 | 1 | 0.03 | 0 | 0 | 0 | 0 | 0.8 | 0.913 | (A) BRAF Class 1 v12 |
| C1053744 | 2p22.2 | 1 | 0.03 | 0 | 0 | 0 | 0 | 0.8 | 0.913 | (A) BRAF Class 1 v12 |
| LTBP4 | 19q13.2 | 1 | 0.03 | 0 | 0 | 0 | 0 | 0.8 | 0.913 | (A) BRAF Class 1 v12 |
| MYEOV | 11q13.3 | 1 | 0.03 | 0 | 0 | 0 | 0 | 0.8 | 0.913 | (A) BRAF Class 1 v12 |
| MYO1D | 17q11.2 | 1 | 0.03 | 0 | 0 | 0 | 0 | 0.8 | 0.913 | (A) BRAF Class 1 v12 |
| MYSM1 | 1p32.1 | 1 | 0.03 | 0 | 0 | 0 | 0 | 0.8 | 0.913 | (A) BRAF Class 1 v12 |
| NBPF20 | 1q21.1 | 1 | 0.03 | 0 | 0 | 0 | 0 | 0.8 | 0.913 | (A) BRAF Class 1 v12 |
| NDUFB2 | 7q34 | 1 | 0.03 | 0 | 0 | 0 | 0 | 0.8 | 0.913 | (A) BRAF Class 1 v12 |
| NELFA | 4p16.3 | 1 | 0.03 | 0 | 0 | 0 | 0 | 0.8 | 0.913 | (A) BRAF Class 1 v12 |

| | | | | | | | | | | |
|---|---|---|---|---|---|---|---|---|---|---|
| PADI2 | 1p36.13 | 1 | 0.03 | 0 | 0 | 0 | 0 | 0.8 | 0.913 | (A) BRAF Class 1 v12 |
| PAX9 | 14q13.3 | 1 | 0.03 | 0 | 0 | 0 | 0 | 0.8 | 0.913 | (A) BRAF Class 1 v12 |
| PFKFB4 | 3p21.31 | 1 | 0.03 | 0 | 0 | 0 | 0 | 0.8 | 0.913 | (A) BRAF Class 1 v12 |
| PKD1 | 16p13.3 | 1 | 0.03 | 0 | 0 | 0 | 0 | 0.8 | 0.913 | (A) BRAF Class 1 v12 |
| PLEKHA7 | .p15.2-p15 | 1 | 0.03 | 0 | 0 | 0 | 0 | 0.8 | 0.913 | (A) BRAF Class 1 v12 |
| PLVAP | 19p13.11 | 1 | 0.03 | 0 | 0 | 0 | 0 | 0.8 | 0.913 | (A) BRAF Class 1 v12 |
| PNPLA6 | 19p13.2 | 1 | 0.03 | 0 | 0 | 0 | 0 | 0.8 | 0.913 | (A) BRAF Class 1 v12 |
| PRMT6 | 1p13.3 | 1 | 0.03 | 0 | 0 | 0 | 0 | 0.8 | 0.913 | (A) BRAF Class 1 v12 |
| PTBP3 | 9q32 | 1 | 0.03 | 0 | 0 | 0 | 0 | 0.8 | 0.913 | (A) BRAF Class 1 v12 |
| RAB2A | q12.1-q12. | 1 | 0.03 | 0 | 0 | 0 | 0 | 0.8 | 0.913 | (A) BRAF Class 1 v12 |
| SAE1 | 19q13.32 | 1 | 0.03 | 0 | 0 | 0 | 0 | 0.8 | 0.913 | (A) BRAF Class 1 v12 |
| SCAP | 3p21.31 | 1 | 0.03 | 0 | 0 | 0 | 0 | 0.8 | 0.913 | (A) BRAF Class 1 v12 |
| SCARA5 | 8p21.1 | 1 | 0.03 | 0 | 0 | 0 | 0 | 0.8 | 0.913 | (A) BRAF Class 1 v12 |
| SDCCAG8 | 1q43-q44 | 1 | 0.03 | 0 | 0 | 0 | 0 | 0.8 | 0.913 | (A) BRAF Class 1 v12 |
| SEMA6D | 15q21.1 | 1 | 0.03 | 0 | 0 | 0 | 0 | 0.8 | 0.913 | (A) BRAF Class 1 v12 |
| SIK3 | 11q23.3 | 1 | 0.03 | 0 | 0 | 0 | 0 | 0.8 | 0.913 | (A) BRAF Class 1 v12 |
| SLC30A5 | q13.1-q13. | 1 | 0.03 | 0 | 0 | 0 | 0 | 0.8 | 0.913 | (A) BRAF Class 1 v12 |
| SND1 | 7q32.1 | 1 | 0.03 | 0 | 0 | 0 | 0 | 0.8 | 0.913 | (A) BRAF Class 1 v12 |
| SPG11 | 15q21.1 | 1 | 0.03 | 0 | 0 | 0 | 0 | 0.8 | 0.913 | (A) BRAF Class 1 v12 |
| TCAF2P1 | 7q35 | 1 | 0.03 | 0 | 0 | 0 | 0 | 0.8 | 0.913 | (A) BRAF Class 1 v12 |
| TRAJ32 | 14q11.2 | 1 | 0.03 | 0 | 0 | 0 | 0 | 0.8 | 0.913 | (A) BRAF Class 1 v12 |
| TRAJ42 | 14q11.2 | 1 | 0.03 | 0 | 0 | 0 | 0 | 0.8 | 0.913 | (A) BRAF Class 1 v12 |
| TRAJ53 | 14q11.2 | 1 | 0.03 | 0 | 0 | 0 | 0 | 0.8 | 0.913 | (A) BRAF Class 1 v12 |
| TRBJ2-2 | 7q34 | 1 | 0.03 | 0 | 0 | 0 | 0 | 0.8 | 0.913 | (A) BRAF Class 1 v12 |
| TRGJP | 7p14.1 | 1 | 0.03 | 0 | 0 | 0 | 0 | 0.8 | 0.913 | (A) BRAF Class 1 v12 |
| TSPOAP1 | 17q22 | 1 | 0.03 | 0 | 0 | 0 | 0 | 0.8 | 0.913 | (A) BRAF Class 1 v12 |
| UGT1A6 | 2q37.1 | 1 | 0.03 | 0 | 0 | 0 | 0 | 0.8 | 0.913 | (A) BRAF Class 1 v12 |
| ZNF423 | 16q12.1 | 1 | 0.03 | 0 | 0 | 0 | 0 | 0.8 | 0.913 | (A) BRAF Class 1 v12 |
| ZNF599 | 19q13.11 | 1 | 0.03 | 0 | 0 | 0 | 0 | 0.8 | 0.913 | (A) BRAF Class 1 v12 |
| HSD3B1 | 1p12 | 13 | 2.48 | 3 | 3.33 | 2 | 1.85 | 0.801 | 0.913 | (B) BRAF Class 2 |
| ZNF217 | 20q13.2 | 54 | 3.38 | 10 | 3.12 | 10 | 2.71 | 0.801 | 0.913 | (A) BRAF Class 1 v12 |
| MAP3K14 | 17q21.31 | 1 | 0.08 | 0 | 0 | 0 | 0 | 0.804 | 0.913 | (A) BRAF Class 1 v12 |
| PRDM1 | 6q21 | 90 | 3.66 | 18 | 3.25 | 23 | 3.99 | 0.806 | 0.913 | (C) BRAF Class 3 |
| NCOR2 | 12q24.31 | 3 | 2.61 | 1 | 4.76 | 1 | 4.55 | 0.807 | 0.913 | (B) BRAF Class 2 |
| XPO1 | 2p15 | 48 | 1.89 | 13 | 2.3 | 12 | 2.05 | 0.808 | 0.913 | (B) BRAF Class 2 |
| RAD51B | 14q24.1 | 13 | 0.93 | 4 | 1.22 | 4 | 1.25 | 0.811 | 0.913 | (C) BRAF Class 3 |
| CBFA2T3 | 16q24.3 | 24 | 4.09 | 7 | 5.26 | 7 | 4.76 | 0.812 | 0.913 | (B) BRAF Class 2 |
| MSI1 | 12q24.31 | 8 | 1.51 | 2 | 1.44 | 1 | 0.78 | 0.812 | 0.913 | (A) BRAF Class 1 v12 |
| MSI2 | 17q22 | 6 | 1.13 | 1 | 0.72 | 2 | 1.55 | 0.813 | 0.913 | (C) BRAF Class 3 |
| HNRNPK | 9q21.32 | 4 | 0.93 | 1 | 1.69 | 1 | 1.52 | 0.814 | 0.913 | (B) BRAF Class 2 |
| GEN1 | 2p24.2 | 24 | 2.31 | 5 | 2.49 | 4 | 1.69 | 0.815 | 0.913 | (B) BRAF Class 2 |
| XRCC2 | 7q36.1 | 23 | 1.22 | 7 | 1.61 | 6 | 1.34 | 0.815 | 0.913 | (B) BRAF Class 2 |
| TCF7L1 | 2p11.2 | 8 | 1.79 | 2 | 2.35 | 1 | 1.11 | 0.822 | 0.913 | (B) BRAF Class 2 |
| BCL2L2 | 14q11.2 | 1 | 0.14 | 0 | 0 | 0 | 0 | 0.822 | 0.913 | (A) BRAF Class 1 v12 |
| GID4 | ›11.2\|17p1 | 1 | 0.17 | 0 | 0 | 0 | 0 | 0.823 | 0.913 | (A) BRAF Class 1 v12 |

| | | | | | | | | | | |
|---|---|---|---|---|---|---|---|---|---|---|
| STK19 | 6p21.33 | 12 | 2.26 | 2 | 1.44 | 3 | 2.33 | 0.824 | 0.913 | (C) BRAF Class 3 |
| EP300 | 22q13.2 | 172 | 6.99 | 36 | 6.51 | 43 | 7.45 | 0.825 | 0.913 | (C) BRAF Class 3 |
| IKZF3 | .7q12-q21. | 7 | 1.26 | 2 | 1.92 | 2 | 1.77 | 0.826 | 0.913 | (B) BRAF Class 2 |
| IRF8 | 16q24.1 | 1 | 0.81 | 0 | 0 | 0 | 0 | 0.826 | 0.913 | (A) BRAF Class 1 v12 |
| DUSP6 | 12q21.33 | 1 | 1.16 | 0 | 0 | 0 | 0 | 0.829 | 0.913 | (A) BRAF Class 1 v12 |
| HSPA5 | 9q33.3 | 1 | 1.16 | 0 | 0 | 0 | 0 | 0.829 | 0.913 | (A) BRAF Class 1 v12 |
| RBBP8 | 18q11.2 | 30 | 5.15 | 6 | 4.58 | 9 | 6.16 | 0.829 | 0.913 | (C) BRAF Class 3 |
| MRTFA | !q13.1-q13 | 1 | 0.86 | 0 | 0 | 0 | 0 | 0.83 | 0.913 | (A) BRAF Class 1 v12 |
| PRDM16 | 1p36.32 | 1 | 0.86 | 0 | 0 | 0 | 0 | 0.83 | 0.913 | (A) BRAF Class 1 v12 |
| FOXO3 | 6q21 | 1 | 0.81 | 0 | 0 | 0 | 0 | 0.832 | 0.913 | (A) BRAF Class 1 v12 |
| CSF1R | 5q32 | 64 | 2.49 | 14 | 2.69 | 16 | 2.93 | 0.834 | 0.913 | (C) BRAF Class 3 |
| CDK5 | 7q36.1 | 3 | 0.85 | 1 | 1.47 | 1 | 1.45 | 0.834 | 0.913 | (B) BRAF Class 2 |
| MLF1 | 3q25.32 | 1 | 1.89 | 0 | 0 | 0 | 0 | 0.834 | 0.913 | (A) BRAF Class 1 v12 |
| YAP1 | 11q22.1 | 29 | 1.55 | 5 | 1.18 | 7 | 1.62 | 0.834 | 0.913 | (C) BRAF Class 3 |
| SPI1 | 11p11.2 | 1 | 0.9 | 0 | 0 | 0 | 0 | 0.834 | 0.913 | (A) BRAF Class 1 v12 |
| AGO2 | 8q24.3 | 26 | 4.91 | 7 | 5.04 | 8 | 6.2 | 0.835 | 0.913 | (C) BRAF Class 3 |
| GNAQ | 9q21.2 | 28 | 0.86 | 8 | 1.08 | 7 | 0.96 | 0.836 | 0.913 | (B) BRAF Class 2 |
| KIF3A | 5q31.1 | 1 | 2.33 | 0 | 0 | 0 | 0 | 0.837 | 0.913 | (A) BRAF Class 1 v12 |
| ARNT | 1q21.3 | 1 | 0.82 | 0 | 0 | 0 | 0 | 0.838 | 0.913 | (A) BRAF Class 1 v12 |
| MUC1 | 1q22 | 1 | 0.82 | 0 | 0 | 0 | 0 | 0.838 | 0.913 | (A) BRAF Class 1 v12 |
| NFKB2 | 10q24.32 | 1 | 0.82 | 0 | 0 | 0 | 0 | 0.838 | 0.913 | (A) BRAF Class 1 v12 |
| PER1 | 17p13.1 | 1 | 0.82 | 0 | 0 | 0 | 0 | 0.838 | 0.913 | (A) BRAF Class 1 v12 |
| PSIP1 | 9p22.3 | 1 | 0.82 | 0 | 0 | 0 | 0 | 0.838 | 0.913 | (A) BRAF Class 1 v12 |
| TLX1 | 10q24.31 | 1 | 0.82 | 0 | 0 | 0 | 0 | 0.838 | 0.913 | (A) BRAF Class 1 v12 |
| TPR | 1q31.1 | 1 | 0.82 | 0 | 0 | 0 | 0 | 0.838 | 0.913 | (A) BRAF Class 1 v12 |
| ZNF384 | 12p13.31 | 1 | 0.82 | 0 | 0 | 0 | 0 | 0.838 | 0.913 | (A) BRAF Class 1 v12 |
| SHH | 7q36.3 | 1 | 0.64 | 0 | 0 | 0 | 0 | 0.838 | 0.913 | (A) BRAF Class 1 v12 |
| SPRY2 | 13q31.1 | 1 | 0.64 | 0 | 0 | 0 | 0 | 0.838 | 0.913 | (A) BRAF Class 1 v12 |
| LCK | 1p35.2 | 1 | 0.8 | 0 | 0 | 0 | 0 | 0.838 | 0.913 | (A) BRAF Class 1 v12 |
| MLLT1 | 19p13.3 | 1 | 1.59 | 0 | 0 | 0 | 0 | 0.838 | 0.913 | (A) BRAF Class 1 v12 |
| TBL1XR1 | 3q26.32 | 1 | 0.97 | 0 | 0 | 0 | 0 | 0.839 | 0.913 | (A) BRAF Class 1 v12 |
| AURKC | 19q13.43 | 1 | 0.85 | 0 | 0 | 0 | 0 | 0.842 | 0.913 | (A) BRAF Class 1 v12 |
| BLNK | 10q24.1 | 1 | 0.85 | 0 | 0 | 0 | 0 | 0.842 | 0.913 | (A) BRAF Class 1 v12 |
| CYP2C19 | 10q23.33 | 1 | 0.85 | 0 | 0 | 0 | 0 | 0.842 | 0.913 | (A) BRAF Class 1 v12 |
| DST | 6p12.1 | 1 | 0.85 | 0 | 0 | 0 | 0 | 0.842 | 0.913 | (A) BRAF Class 1 v12 |
| GDNF | 5p13.2 | 1 | 0.85 | 0 | 0 | 0 | 0 | 0.842 | 0.913 | (A) BRAF Class 1 v12 |
| IKBKB | 8p11.21 | 1 | 0.85 | 0 | 0 | 0 | 0 | 0.842 | 0.913 | (A) BRAF Class 1 v12 |
| ITGA9 | 3p22.2 | 1 | 0.85 | 0 | 0 | 0 | 0 | 0.842 | 0.913 | (A) BRAF Class 1 v12 |
| ITGB3 | 17q21.32 | 1 | 0.85 | 0 | 0 | 0 | 0 | 0.842 | 0.913 | (A) BRAF Class 1 v12 |
| PLEKHG5 | 1p36.31 | 1 | 0.85 | 0 | 0 | 0 | 0 | 0.842 | 0.913 | (A) BRAF Class 1 v12 |
| RRM1 | 11p15.4 | 1 | 0.85 | 0 | 0 | 0 | 0 | 0.842 | 0.913 | (A) BRAF Class 1 v12 |
| DNAJB1 | 19p13.12 | 14 | 1.07 | 2 | 0.69 | 3 | 1.08 | 0.842 | 0.913 | (C) BRAF Class 3 |
| MYC | 8q24.21 | 89 | 3.46 | 20 | 3.25 | 23 | 3.85 | 0.843 | 0.913 | (C) BRAF Class 3 |
| ARFGAP3 | 22q13.2 | 1 | 2.08 | 0 | 0 | 0 | 0 | 0.844 | 0.913 | (A) BRAF Class 1 v12 |
| GNAI3 | 1p13.3 | 1 | 2.08 | 0 | 0 | 0 | 0 | 0.844 | 0.913 | (A) BRAF Class 1 v12 |

| | | | | | | | | | | |
|---|---|---|---|---|---|---|---|---|---|---|
| SNX31 | 8q22.3 | 1 | 2.08 | 0 | 0 | 0 | 0 | 0.844 | 0.913 | (A) BRAF Class 1 v12 |
| TMEM216 | 11q12.2 | 1 | 2.08 | 0 | 0 | 0 | 0 | 0.844 | 0.913 | (A) BRAF Class 1 v12 |
| SP140 | 2q37.1 | 1 | 1.96 | 0 | 0 | 0 | 0 | 0.844 | 0.913 | (A) BRAF Class 1 v12 |
| BIRC5 | 17q25.3 | 1 | 0.53 | 0 | 0 | 0 | 0 | 0.844 | 0.913 | (A) BRAF Class 1 v12 |
| POLH | 6p21.1 | 11 | 1.71 | 3 | 2.11 | 2 | 1.26 | 0.847 | 0.915 | (B) BRAF Class 2 |
| HIP1 | 7q11.23 | 1 | 1.11 | 0 | 0 | 0 | 0 | 0.85 | 0.917 | (A) BRAF Class 1 v12 |
| PIK3R3 | 1p34.1 | 13 | 1.1 | 2 | 0.74 | 3 | 1.17 | 0.851 | 0.917 | (C) BRAF Class 3 |
| ERCC1 | 19q13.32 | 7 | 0.66 | 2 | 1.01 | 2 | 0.85 | 0.851 | 0.917 | (B) BRAF Class 2 |
| ETNK1 | 12p12.1 | 1 | 0.63 | 0 | 0 | 0 | 0 | 0.853 | 0.918 | (A) BRAF Class 1 v12 |
| LMO2 | 11p13 | 6 | 0.65 | 1 | 0.5 | 2 | 0.93 | 0.862 | 0.927 | (C) BRAF Class 3 |
| BMPR1A | 10q23.2 | 48 | 2.06 | 9 | 1.73 | 10 | 1.88 | 0.87 | 0.932 | (A) BRAF Class 1 v12 |
| KBTBD4 | 11p11.2 | 1 | 0.98 | 0 | 0 | 0 | 0 | 0.871 | 0.932 | (A) BRAF Class 1 v12 |
| FOSB | 19q13.32 | 1 | 1.09 | 0 | 0 | 0 | 0 | 0.872 | 0.932 | (A) BRAF Class 1 v12 |
| MAML3 | 4q31.1 | 1 | 1.09 | 0 | 0 | 0 | 0 | 0.872 | 0.932 | (A) BRAF Class 1 v12 |
| OTX2 | 14q22.3 | 1 | 1.09 | 0 | 0 | 0 | 0 | 0.872 | 0.932 | (A) BRAF Class 1 v12 |
| TBXT | 6q27 | 1 | 1.09 | 0 | 0 | 0 | 0 | 0.872 | 0.932 | (A) BRAF Class 1 v12 |
| EWSR1 | 22q12.2 | 28 | 1.76 | 6 | 1.88 | 8 | 2.16 | 0.872 | 0.932 | (C) BRAF Class 3 |
| PRKD1 | 14q12 | 11 | 1.85 | 2 | 1.33 | 3 | 2.11 | 0.873 | 0.932 | (C) BRAF Class 3 |
| SESN1 | 6q21 | 6 | 1.13 | 1 | 0.72 | 1 | 0.78 | 0.874 | 0.932 | (A) BRAF Class 1 v12 |
| LIG4 | 13q33.3 | 20 | 3.44 | 5 | 3.82 | 4 | 2.74 | 0.876 | 0.932 | (B) BRAF Class 2 |
| CD3EAP | 19q13.32 | 1 | 0.3 | 0 | 0 | 0 | 0 | 0.876 | 0.932 | (A) BRAF Class 1 v12 |
| TSPAN31 | 12q14.1 | 1 | 0.3 | 0 | 0 | 0 | 0 | 0.876 | 0.932 | (A) BRAF Class 1 v12 |
| FH | 1q43 | 48 | 1.97 | 9 | 1.64 | 11 | 1.93 | 0.879 | 0.933 | (A) BRAF Class 1 v12 |
| RANBP2 | 2q13 | 56 | 11.34 | 9 | 12.33 | 9 | 9.89 | 0.879 | 0.933 | (B) BRAF Class 2 |
| CBLC | 19q13.32 | 1 | 2.78 | 0 | 0 | 0 | 0 | 0.88 | 0.933 | (A) BRAF Class 1 v12 |
| LRIG3 | 12q14.1 | 1 | 2.78 | 0 | 0 | 0 | 0 | 0.88 | 0.933 | (A) BRAF Class 1 v12 |
| CHD2 | 15q26.1 | 38 | 6.04 | 6 | 6.19 | 6 | 4.92 | 0.883 | 0.935 | (B) BRAF Class 2 |
| BRD3 | 9q34.2 | 30 | 4.34 | 6 | 3.95 | 6 | 3.53 | 0.885 | 0.936 | (A) BRAF Class 1 v12 |
| PDCD1LG2 | 9p24.1 | 25 | 1.21 | 5 | 1.11 | 7 | 1.45 | 0.887 | 0.937 | (C) BRAF Class 3 |
| MDC1 | 6p21.33 | 65 | 5.49 | 15 | 5.51 | 16 | 6.25 | 0.89 | 0.94 | (C) BRAF Class 3 |
| GNB1 | 1p36.33 | 1 | 2.27 | 0 | 0 | 0 | 0 | 0.891 | 0.94 | (A) BRAF Class 1 v12 |
| TNFAIP3 | 6q23.3 | 59 | 2.4 | 15 | 2.71 | 15 | 2.6 | 0.893 | 0.941 | (B) BRAF Class 2 |
| MYBL1 | 8q13.1 | 30 | 2.7 | 5 | 2.16 | 7 | 2.71 | 0.893 | 0.941 | (C) BRAF Class 3 |
| PLK2 | 5q11.2 | 27 | 2.28 | 5 | 1.84 | 6 | 2.34 | 0.895 | 0.941 | (C) BRAF Class 3 |
| DDB1 | 11q12.2 | 21 | 3.61 | 4 | 3.05 | 6 | 4.11 | 0.895 | 0.941 | (C) BRAF Class 3 |
| MAGOH | 1p32.3 | 1 | 0.99 | 0 | 0 | 0 | 0 | 0.896 | 0.941 | (A) BRAF Class 1 v12 |
| KDM6B | 17p13.1 | 20 | 5.52 | 4 | 5.71 | 3 | 4.23 | 0.897 | 0.942 | (B) BRAF Class 2 |
| ZBTB7A | 19p13.3 | 12 | 3.6 | 2 | 5 | 2 | 4.17 | 0.9 | 0.943 | (B) BRAF Class 2 |
| RPS6KA4 | 11q13.1 | 36 | 3.04 | 7 | 2.57 | 7 | 2.73 | 0.901 | 0.943 | (A) BRAF Class 1 v12 |
| XRCC5 | 2q35 | 10 | 1.66 | 3 | 2.16 | 3 | 1.99 | 0.905 | 0.946 | (B) BRAF Class 2 |
| MCM7 | 7q22.1 | 1 | 3.23 | 0 | 0 | 0 | 0 | 0.905 | 0.946 | (A) BRAF Class 1 v12 |
| UROD | 1p34.1 | 9 | 1.55 | 2 | 1.53 | 3 | 2.05 | 0.906 | 0.946 | (C) BRAF Class 3 |
| MYH9 | 22q12.3 | 8 | 3.13 | 1 | 2.27 | 2 | 3.85 | 0.907 | 0.946 | (C) BRAF Class 3 |
| OLIG2 | 21q22.11 | 1 | 3.7 | 0 | 0 | 0 | 0 | 0.909 | 0.946 | (A) BRAF Class 1 v12 |
| IRS2 | 13q34 | 78 | 4.97 | 18 | 5.03 | 20 | 5.52 | 0.909 | 0.946 | (C) BRAF Class 3 |

| | | | | | | | | | | |
|---|---|---|---|---|---|---|---|---|---|---|
| DCUN1D1 | 3q26.33 | 7 | 0.59 | 2 | 0.62 | 1 | 0.38 | 0.909 | 0.946 | (B) BRAF Class 2 |
| ZNF703 | 8p11.23 | 21 | 2.97 | 3 | 2.4 | 5 | 3.25 | 0.914 | 0.95 | (C) BRAF Class 3 |
| NCOR1 | .7p12-p11. | 79 | 5.32 | 17 | 5.12 | 16 | 4.76 | 0.915 | 0.95 | (A) BRAF Class 1 v12 |
| CYLD | 16q12.1 | 49 | 2.36 | 11 | 2.51 | 10 | 2.1 | 0.916 | 0.95 | (B) BRAF Class 2 |
| RHOA | 3p21.31 | 31 | 1.53 | 6 | 1.34 | 6 | 1.32 | 0.916 | 0.95 | (A) BRAF Class 1 v12 |
| FLT4 | 5q35.3 | 171 | 7.03 | 35 | 6.53 | 39 | 6.96 | 0.917 | 0.95 | (A) BRAF Class 1 v12 |
| CD79A | 19q13.2 | 22 | 1.46 | 6 | 1.75 | 5 | 1.46 | 0.919 | 0.951 | (B) BRAF Class 2 |
| RHBDF2 | 17q25.1 | 25 | 4.3 | 5 | 3.82 | 7 | 4.79 | 0.923 | 0.953 | (C) BRAF Class 3 |
| SBDS | 7q11.21 | 7 | 0.68 | 2 | 0.91 | 2 | 0.83 | 0.923 | 0.953 | (B) BRAF Class 2 |
| MAFB | 20q12 | 3 | 0.44 | 1 | 0.66 | 1 | 0.59 | 0.923 | 0.953 | (B) BRAF Class 2 |
| FAAP20 | 1p36.33 | 3 | 0.52 | 1 | 0.76 | 1 | 0.68 | 0.93 | 0.959 | (B) BRAF Class 2 |
| ETV1 | 7p21.2 | 58 | 2.55 | 13 | 2.54 | 15 | 2.83 | 0.932 | 0.961 | (C) BRAF Class 3 |
| RAB35 | 12q24.23 | 11 | 0.99 | 3 | 1.18 | 2 | 0.85 | 0.933 | 0.961 | (B) BRAF Class 2 |
| RCS8-MEF | 19p13.11 | 19 | 0.57 | 5 | 0.67 | 4 | 0.54 | 0.937 | 0.963 | (B) BRAF Class 2 |
| GABRA6 | 5q34 | 16 | 3.05 | 3 | 3.33 | 4 | 3.7 | 0.937 | 0.963 | (C) BRAF Class 3 |
| TMEM127 | 2q11.2 | 11 | 0.61 | 3 | 0.72 | 3 | 0.73 | 0.938 | 0.964 | (C) BRAF Class 3 |
| HIST2H3D | 1q21.2 | 6 | 0.54 | 1 | 0.4 | 1 | 0.43 | 0.942 | 0.967 | (A) BRAF Class 1 v12 |
| ARAF | Xp11.3 | 64 | 2.47 | 13 | 2.25 | 14 | 2.34 | 0.944 | 0.968 | (A) BRAF Class 1 v12 |
| RAD51D | 17q12 | 21 | 0.96 | 4 | 0.81 | 5 | 0.97 | 0.948 | 0.971 | (C) BRAF Class 3 |
| KDM5A | 12p13.33 | 120 | 5.67 | 29 | 5.98 | 28 | 5.53 | 0.952 | 0.974 | (B) BRAF Class 2 |
| FGF10 | 5p12 | 9 | 1.27 | 2 | 1.6 | 2 | 1.3 | 0.957 | 0.976 | (B) BRAF Class 2 |
| CCN6 | 6q21 | 17 | 2.51 | 3 | 2.78 | 3 | 2.19 | 0.957 | 0.976 | (B) BRAF Class 2 |
| NUP93 | 16q13 | 30 | 2.06 | 7 | 2.17 | 6 | 1.85 | 0.957 | 0.976 | (B) BRAF Class 2 |
| MRE11 | 11q21 | 57 | 2.58 | 12 | 2.41 | 14 | 2.71 | 0.957 | 0.976 | (C) BRAF Class 3 |
| SQSTM1 | 5q35.3 | 4 | 1.52 | 1 | 2.08 | 1 | 1.72 | 0.959 | 0.977 | (B) BRAF Class 2 |
| RPA1 | 17p13.3 | 10 | 1.46 | 2 | 1.28 | 3 | 1.64 | 0.963 | 0.981 | (C) BRAF Class 3 |
| RIF1 | 2q23.3 | 41 | 7.04 | 10 | 7.63 | 10 | 6.85 | 0.964 | 0.981 | (B) BRAF Class 2 |
| FLCN | 17p11.2 | 60 | 2.46 | 13 | 2.36 | 13 | 2.28 | 0.966 | 0.982 | (A) BRAF Class 1 v12 |
| TLX3 | 5q35.1 | 14 | 2.39 | 3 | 2.26 | 3 | 2.04 | 0.969 | 0.984 | (A) BRAF Class 1 v12 |
| PGR | 11q22.1 | 67 | 6.01 | 15 | 5.91 | 15 | 6.38 | 0.972 | 0.986 | (C) BRAF Class 3 |
| TRIM37 | 17q22 | 18 | 3.09 | 4 | 3.05 | 4 | 2.74 | 0.975 | 0.988 | (A) BRAF Class 1 v12 |
| TOE1 | 1p34.1 | 0 | 0 | 0 | 0 | 1 | 0.13 | 0.975 | 0.988 | (C) BRAF Class 3 |
| DPYD | 1p21.3 | 4 | 1.62 | 1 | 2 | 1 | 1.56 | 0.979 | 0.991 | (B) BRAF Class 2 |
| GNAS | 20q13.32 | 105 | 3.2 | 21 | 3.07 | 23 | 3.2 | 0.983 | 0.993 | (C) BRAF Class 3 |
| GPC3 | Xq26.2 | 20 | 1.78 | 4 | 1.7 | 5 | 1.92 | 0.983 | 0.993 | (C) BRAF Class 3 |
| HIST1H3J | 6p22.1 | 10 | 0.9 | 2 | 0.79 | 2 | 0.85 | 0.985 | 0.994 | (A) BRAF Class 1 v12 |
| BLM | 15q26.1 | 107 | 4.4 | 24 | 4.48 | 24 | 4.29 | 0.988 | 0.995 | (B) BRAF Class 2 |
| NTHL1 | 16p13.3 | 9 | 0.78 | 2 | 0.71 | 2 | 0.7 | 0.988 | 0.995 | (A) BRAF Class 1 v12 |
| NUP214 | 9q34.13 | 5 | 3.82 | 1 | 4.35 | 1 | 4.17 | 0.991 | 0.998 | (B) BRAF Class 2 |
| RAD51C | 17q22 | 25 | 1.15 | 6 | 1.21 | 6 | 1.16 | 0.992 | 0.998 | (B) BRAF Class 2 |
| HIST2H3C | 1q21.2 | 5 | 0.45 | 1 | 0.39 | 1 | 0.43 | 0.992 | 0.998 | (A) BRAF Class 1 v12 |
| PRKN | 6q26 | 79 | 3.29 | 18 | 3.33 | 19 | 3.39 | 0.993 | 0.998 | (C) BRAF Class 3 |
| TAP1 | 6p21.32 | 12 | 2.26 | 3 | 2.16 | 3 | 2.33 | 0.996 | 0.998 | (C) BRAF Class 3 |
| TAL2 | 9q31.2 | 4 | 0.68 | 1 | 0.75 | 1 | 0.68 | 0.996 | 0.998 | (B) BRAF Class 2 |
| MAF | 16q23.2 | 4 | 0.56 | 1 | 0.6 | 1 | 0.54 | 0.997 | 0.999 | (B) BRAF Class 2 |

| | | | | | | | | | | |
|---|---|---|---|---|---|---|---|---|---|---|
| IL10 | 1q32.1 | 9 | 0.76 | 2 | 0.74 | 2 | 0.78 | 0.998 | 0.999 | (C) BRAF Class 3 |
| BRAF | 7q34 | 3347 | 100 | 749 | 100 | 743 | 100 | 1 | 1 | (A) BRAF Class 1 v12 |

#### Supplementary Table 3

| Gene | Cytoband | Class 1 Occurrence | Class 1 % | Class 2 Occurrence | Class 2 % | pvalue | qvalue | Enriched In |
|---|---|---|---|---|---|---|---|---|
| STK11 | 19p13.3 | 57 | 1.74 | 80 | 11.53 | 4.95E-28 | 5.48E-25 | (B) BRAF Class 2 |
| KRAS | 12p12.1 | 37 | 1.11 | 67 | 8.83 | 1.44E-25 | 7.96E-23 | (B) BRAF Class 2 |
| NRAS | 1p13.2 | 35 | 1.05 | 48 | 6.32 | 7.92E-16 | 2.93E-13 | (B) BRAF Class 2 |
| TP53 | 17p13.1 | 892 | 26.67 | 282 | 39.66 | 8.22E-12 | 2.28E-09 | (B) BRAF Class 2 |
| KEAP1 | 19p13.2 | 53 | 2.24 | 45 | 8.33 | 2.56E-10 | 5.68E-08 | (B) BRAF Class 2 |
| RNF43 | 17q22 | 260 | 11.71 | 17 | 3.35 | 3.7E-10 | 6.84E-08 | (A) BRAF Class 1 v12 |
| NF1 | 17q11.2 | 142 | 5.59 | 82 | 13.20 | 5.67E-10 | 8.98E-08 | (B) BRAF Class 2 |
| TERT | 5p15.33 | 842 | 34.23 | 128 | 23.23 | 2.18E-07 | 3.03E-05 | (A) BRAF Class 1 v12 |
| FLT3 | 13q12.2 | 100 | 3.11 | 51 | 7.52 | 5.25E-07 | 6.47E-05 | (B) BRAF Class 2 |
| POT1 | 7q31.33 | 13 | 1.55 | 16 | 8.65 | 5.32E-06 | 0.00059 | (B) BRAF Class 2 |
| ATR | 3q23 | 126 | 5.72 | 55 | 10.91 | 4.92E-05 | 0.004963 | (B) BRAF Class 2 |
| FGFR1 | 8p11.23 | 55 | 1.65 | 30 | 3.95 | 0.000161 | 0.0148 | (B) BRAF Class 2 |
| CBLB | 3q13.11 | 16 | 1.52 | 14 | 6.19 | 0.000195 | 0.0166 | (B) BRAF Class 2 |
| TGFBR2 | 3p24.1 | 112 | 7.16 | 9 | 2.48 | 0.000249 | 0.0197 | (A) BRAF Class 1 v12 |
| NFE2L2 | 2q31.2 | 31 | 1.22 | 20 | 3.50 | 0.000363 | 0.0268 | (B) BRAF Class 2 |
| FOXA1 | 14q21.1 | 39 | 1.98 | 23 | 5.12 | 0.000399 | 0.0277 | (B) BRAF Class 2 |
| COL2A1 | 12q13.11 | 5 | 3.21 | 7 | 25.00 | 0.000428 | 0.0279 | (B) BRAF Class 2 |
| CDKN1C | 11p15.4 | 5 | 0.51 | 8 | 3.76 | 0.000506 | 0.0296 | (B) BRAF Class 2 |
| KDR | 4q12 | 163 | 5.12 | 58 | 8.61 | 0.000508 | 0.0296 | (B) BRAF Class 2 |
| KDM6A | Xp11.3 | 65 | 2.65 | 31 | 5.54 | 0.000743 | 0.0412 | (B) BRAF Class 2 |
| FANCL | 2p16.1 | 16 | 1.26 | 12 | 4.62 | 0.001061 | 0.056 | (B) BRAF Class 2 |
| CASP8 | 2q33.1 | 91 | 4.47 | 8 | 1.66 | 0.00165 | 0.0832 | (A) BRAF Class 1 v12 |
| SPTA1 | 1q23.1 | 95 | 15.13 | 28 | 28.00 | 0.001855 | 0.0873 | (B) BRAF Class 2 |
| EPHA3 | 3p11.1 | 91 | 4.80 | 37 | 8.62 | 0.002011 | 0.0873 | (B) BRAF Class 2 |
| HRAS | 11p15.5 | 18 | 0.54 | 13 | 1.72 | 0.00202 | 0.0873 | (B) BRAF Class 2 |
| ERBB4 | 2q34 | 205 | 6.33 | 70 | 9.49 | 0.002047 | 0.0873 | (B) BRAF Class 2 |
| DGKH | 13q14.11 | 0 | 0.00 | 3 | 21.43 | 0.002251 | 0.09 | (B) BRAF Class 2 |
| MSH3 | 5q14.1 | 94 | 8.83 | 8 | 3.43 | 0.002273 | 0.09 | (A) BRAF Class 1 v12 |
| ERBB2 | 17q12 | 88 | 2.64 | 36 | 4.74 | 0.002548 | 0.0974 | (B) BRAF Class 2 |
| GNA14 | 9q21.2 | 0 | 0.00 | 3 | 18.75 | 0.002743 | 0.101 | (B) BRAF Class 2 |
| EPHA7 | 6q16.1 | 108 | 6.10 | 40 | 10.28 | 0.003194 | 0.112 | (B) BRAF Class 2 |
| PRKACA | 19p13.12 | 0 | 0.00 | 3 | 7.69 | 0.00323 | 0.112 | (B) BRAF Class 2 |
| NOTCH2 | 1p12 | 109 | 4.37 | 41 | 7.32 | 0.003483 | 0.117 | (B) BRAF Class 2 |
| MPL | 1p34.2 | 52 | 1.66 | 23 | 3.44 | 0.003619 | 0.118 | (B) BRAF Class 2 |
| SMC1A | Xp11.22 | 9 | 1.00 | 7 | 4.58 | 0.004332 | 0.135 | (B) BRAF Class 2 |
| CDKN2A | 9p21.3 | 449 | 13.71 | 123 | 17.72 | 0.004378 | 0.135 | (B) BRAF Class 2 |
| SYNE1 | 6q25.2 | 11 | 2.14 | 7 | 8.97 | 0.005078 | 0.152 | (B) BRAF Class 2 |
| GATA6 | 18q11.2 | 28 | 1.98 | 14 | 4.86 | 0.006535 | 0.191 | (B) BRAF Class 2 |
| PTPN11 | 12q24.13 | 36 | 1.11 | 17 | 2.46 | 0.007012 | 0.195 | (B) BRAF Class 2 |
| MET | 7q31.2 | 120 | 3.62 | 43 | 5.72 | 0.007045 | 0.195 | (B) BRAF Class 2 |
| NEIL3 | 4q34.3 | 23 | 3.95 | 0 | 0.00 | 0.008157 | 0.218 | (A) BRAF Class 1 v12 |
| OR5L1 | 11q12.1 | 1 | 1.16 | 3 | 21.43 | 0.008238 | 0.218 | (B) BRAF Class 2 |
| RSPO2 | 8q23.1 | 12 | 1.35 | 9 | 4.35 | 0.009081 | 0.225 | (B) BRAF Class 2 |
| ARID1A | 1p36.11 | 268 | 10.57 | 81 | 14.21 | 0.009122 | 0.225 | (B) BRAF Class 2 |
| PTPRB | 12q15 | 8 | 5.13 | 6 | 21.43 | 0.009237 | 0.225 | (B) BRAF Class 2 |
| CXCR4 | 2q22.1 | 12 | 0.63 | 9 | 2.05 | 0.009345 | 0.225 | (B) BRAF Class 2 |

| | | | | | | | | |
|---|---|---|---|---|---|---|---|---|
| DDX31 | 9q34.13 | 1 | 1.09 | 3 | 18.75 | 0.009953 | 0.235 | (B) BRAF Class 2 |
| CIITA | 16p13.13 | 48 | 4.82 | 20 | 9.26 | 0.011 | 0.254 | (B) BRAF Class 2 |
| KEL | 7q34 | 30 | 5.73 | 12 | 13.19 | 0.0129 | 0.279 | (B) BRAF Class 2 |
| IRF4 | 6p25.3 | 21 | 1.34 | 12 | 3.31 | 0.013 | 0.279 | (B) BRAF Class 2 |
| FANCF | 11p14.3 | 10 | 0.63 | 7 | 2.25 | 0.0131 | 0.279 | (B) BRAF Class 2 |
| SETBP1 | 18q12.3 | 103 | 7.02 | 32 | 11.19 | 0.0133 | 0.279 | (B) BRAF Class 2 |
| HIST1H3B | 6p22.2 | 19 | 0.91 | 11 | 2.34 | 0.0137 | 0.279 | (B) BRAF Class 2 |
| POLD1 | 19q13.33 | 132 | 6.42 | 18 | 3.76 | 0.0139 | 0.279 | (A) BRAF Class 1 v12 |
| PLCB4 | 0p12.3-p12. | 5 | 2.29 | 5 | 11.36 | 0.0141 | 0.279 | (B) BRAF Class 2 |
| SF3B1 | 2q33.1 | 60 | 2.28 | 24 | 4.04 | 0.0144 | 0.279 | (B) BRAF Class 2 |
| RAF1 | 3p25.2 | 37 | 1.42 | 18 | 2.83 | 0.0145 | 0.279 | (B) BRAF Class 2 |
| XRCC3 | 14q32.33 | 20 | 2.92 | 0 | 0.00 | 0.0146 | 0.279 | (A) BRAF Class 1 v12 |
| PDGFRA | 4q12 | 102 | 3.14 | 36 | 4.88 | 0.0153 | 0.283 | (B) BRAF Class 2 |
| MAP2K1 | 15q22.31 | 72 | 2.67 | 29 | 4.43 | 0.0154 | 0.283 | (B) BRAF Class 2 |
| ELF3 | 1q32.1 | 10 | 1.52 | 8 | 4.82 | 0.0157 | 0.283 | (B) BRAF Class 2 |
| HLA-B | 6p21.33 | 25 | 4.11 | 1 | 0.62 | 0.0159 | 0.283 | (A) BRAF Class 1 v12 |
| NOTCH1 | 9q34.3 | 177 | 5.39 | 53 | 7.63 | 0.0161 | 0.283 | (B) BRAF Class 2 |
| PDGFRB | 5q32 | 93 | 3.67 | 33 | 5.79 | 0.0166 | 0.288 | (B) BRAF Class 2 |
| SRC | 20q11.23 | 22 | 0.83 | 11 | 2.01 | 0.0173 | 0.291 | (B) BRAF Class 2 |
| ESR2 | 4q23.2-q23. | 1 | 0.39 | 3 | 5.56 | 0.0173 | 0.291 | (B) BRAF Class 2 |
| QKI | 6q26 | 48 | 3.62 | 3 | 1.12 | 0.0183 | 0.3 | (A) BRAF Class 1 v12 |
| IL2RB | 22q12.3 | 0 | 0.00 | 2 | 14.29 | 0.0184 | 0.3 | (B) BRAF Class 2 |
| ZNRF3 | 22q12.1 | 42 | 6.01 | 3 | 1.86 | 0.0187 | 0.301 | (A) BRAF Class 1 v12 |
| JAK3 | 19p13.11 | 101 | 3.15 | 35 | 4.82 | 0.0199 | 0.311 | (B) BRAF Class 2 |
| DCLRE1C | 10p13 | 7 | 1.20 | 6 | 4.51 | 0.0202 | 0.311 | (B) BRAF Class 2 |
| HLA-A | 6p22.1 | 48 | 3.96 | 4 | 1.42 | 0.0203 | 0.311 | (A) BRAF Class 1 v12 |
| NCKAP5 | 2q21.2 | 5 | 5.81 | 4 | 28.57 | 0.0207 | 0.311 | (B) BRAF Class 2 |
| PBRM1 | 3p21.1 | 108 | 4.56 | 37 | 6.85 | 0.021 | 0.311 | (B) BRAF Class 2 |
| PPARG | 3p25.2 | 40 | 2.26 | 17 | 4.31 | 0.021 | 0.311 | (B) BRAF Class 2 |
| FGFR2 | 10q26.13 | 87 | 2.60 | 31 | 4.08 | 0.0218 | 0.312 | (B) BRAF Class 2 |
| B2M | 15q21.1 | 86 | 3.97 | 10 | 2.04 | 0.0218 | 0.312 | (A) BRAF Class 1 v12 |
| NTRK2 | 9q21.33 | 53 | 2.10 | 21 | 3.68 | 0.0222 | 0.312 | (B) BRAF Class 2 |
| DROSHA | 5p13.3 | 24 | 4.03 | 1 | 0.65 | 0.0222 | 0.312 | (A) BRAF Class 1 v12 |
| SPRED1 | 15q14 | 11 | 1.67 | 8 | 4.82 | 0.0225 | 0.312 | (B) BRAF Class 2 |
| HIF1A | 14q23.2 | 0 | 0.00 | 2 | 4.35 | 0.0231 | 0.312 | (B) BRAF Class 2 |
| CD79B | 17q23.3 | 15 | 0.61 | 9 | 1.61 | 0.0232 | 0.312 | (B) BRAF Class 2 |
| CREB1 | 2q33.3 | 0 | 0.00 | 2 | 4.26 | 0.0236 | 0.312 | (B) BRAF Class 2 |
| H3F3B | 17q25.1 | 5 | 0.27 | 5 | 1.18 | 0.0237 | 0.312 | (B) BRAF Class 2 |
| ZFHX4 | 8q21.13 | 18 | 11.54 | 8 | 28.57 | 0.0241 | 0.314 | (B) BRAF Class 2 |
| RARA | 17q21.2 | 43 | 1.75 | 18 | 3.21 | 0.0244 | 0.315 | (B) BRAF Class 2 |
| FGF6 | 12p13.32 | 10 | 1.41 | 6 | 4.65 | 0.0254 | 0.32 | (B) BRAF Class 2 |
| EZH2 | 7q36.1 | 46 | 1.52 | 18 | 2.77 | 0.0254 | 0.32 | (B) BRAF Class 2 |
| LATS1 | 6q25.1 | 24 | 2.04 | 12 | 4.40 | 0.0262 | 0.326 | (B) BRAF Class 2 |
| DDR2 | 1q23.3 | 56 | 2.10 | 23 | 3.53 | 0.027 | 0.326 | (B) BRAF Class 2 |
| STAT6 | 12q13.3 | 17 | 1.51 | 9 | 3.75 | 0.0272 | 0.326 | (B) BRAF Class 2 |
| PRKCA | 17q24.2 | 0 | 0.00 | 2 | 4.17 | 0.0273 | 0.326 | (B) BRAF Class 2 |
| RUNX1T1 | 8q21.3 | 56 | 4.49 | 19 | 7.79 | 0.0274 | 0.326 | (B) BRAF Class 2 |
| ANKRD26 | 10p12.1 | 45 | 13.51 | 1 | 2.50 | 0.0278 | 0.328 | (A) BRAF Class 1 v12 |

| | | | | | | | | |
|---|---|---|---|---|---|---|---|---|
| SMARCA2 | 9p24.3 | 2 | 1.20 | 3 | 9.68 | 0.0285 | 0.333 | (B) BRAF Class 2 |
| BAP1 | 3p21.1 | 62 | 2.45 | 23 | 4.03 | 0.0296 | 0.341 | (B) BRAF Class 2 |
| EPHA5 | 4q13.1-q13.2 | 89 | 4.79 | 30 | 7.28 | 0.03 | 0.343 | (B) BRAF Class 2 |
| PTPRK | 6q22.33 | 19 | 12.18 | 8 | 28.57 | 0.0305 | 0.345 | (B) BRAF Class 2 |
| BRIP1 | 17q23.2 | 65 | 2.65 | 24 | 4.29 | 0.0308 | 0.345 | (B) BRAF Class 2 |
| CDK12 | 17q12 | 72 | 3.27 | 26 | 5.16 | 0.0312 | 0.345 | (B) BRAF Class 2 |
| ERBB3 | 12q13.2 | 81 | 3.15 | 30 | 4.82 | 0.0314 | 0.345 | (B) BRAF Class 2 |
| RRAS2 | 11p15.2 | 3 | 0.45 | 4 | 2.41 | 0.0333 | 0.362 | (B) BRAF Class 2 |
| USP28 | 11q23.2 | 20 | 3.43 | 10 | 7.52 | 0.036 | 0.387 | (B) BRAF Class 2 |
| RAD54B | 8q22.1 | 17 | 2.92 | 9 | 6.77 | 0.0363 | 0.387 | (B) BRAF Class 2 |
| PDK1 | 2q31.1 | 8 | 1.13 | 5 | 3.88 | 0.0367 | 0.387 | (B) BRAF Class 2 |
| AURKA | 20q13.2 | 26 | 1.06 | 12 | 2.14 | 0.037 | 0.387 | (B) BRAF Class 2 |
| IDH1 | 2q34 | 81 | 2.43 | 28 | 3.69 | 0.0376 | 0.39 | (B) BRAF Class 2 |
| KMT2C | 7q36.1 | 184 | 12.67 | 30 | 9.06 | 0.0389 | 0.396 | (A) BRAF Class 1 v12 |
| CD74 | 5q33.1 | 3 | 0.65 | 3 | 3.95 | 0.0389 | 0.396 | (B) BRAF Class 2 |
| MAGI2 | 7q21.11 | 38 | 7.69 | 11 | 14.86 | 0.0402 | 0.397 | (B) BRAF Class 2 |
| SOX10 | 22q13.1 | 13 | 1.92 | 6 | 5.36 | 0.0403 | 0.397 | (B) BRAF Class 2 |
| SOX17 | 8q11.23 | 55 | 4.67 | 6 | 2.20 | 0.0406 | 0.397 | (A) BRAF Class 1 v12 |
| NTRK1 | 1q23.1 | 101 | 3.89 | 33 | 5.63 | 0.0407 | 0.397 | (B) BRAF Class 2 |
| CASR | q13.33-q21. | 3 | 3.26 | 3 | 18.75 | 0.0409 | 0.397 | (B) BRAF Class 2 |
| NKX3-1 | 8p21.2 | 17 | 0.98 | 9 | 2.23 | 0.0417 | 0.397 | (B) BRAF Class 2 |
| ATM | 11q22.3 | 257 | 8.08 | 69 | 10.24 | 0.042 | 0.397 | (B) BRAF Class 2 |
| GSK3B | 3q13.33 | 17 | 1.09 | 9 | 2.48 | 0.042 | 0.397 | (B) BRAF Class 2 |
| BCL11B | 14q32.2 | 31 | 4.45 | 13 | 8.33 | 0.0431 | 0.397 | (B) BRAF Class 2 |
| DUSP4 | 8p12 | 8 | 1.21 | 6 | 3.61 | 0.0434 | 0.397 | (B) BRAF Class 2 |
| CCNE1 | 19q12 | 44 | 1.74 | 17 | 2.98 | 0.0438 | 0.397 | (B) BRAF Class 2 |
| CTNNA1 | 5q31.2 | 27 | 2.12 | 11 | 4.23 | 0.044 | 0.397 | (B) BRAF Class 2 |
| MBD4 | 3q21.3 | 15 | 2.57 | 0 | 0.00 | 0.0443 | 0.397 | (A) BRAF Class 1 v12 |
| SERPINA1 | 14q32.13 | 15 | 2.57 | 0 | 0.00 | 0.0443 | 0.397 | (A) BRAF Class 1 v12 |
| SMARCA4 | 19p13.2 | 160 | 6.31 | 48 | 8.42 | 0.0448 | 0.397 | (B) BRAF Class 2 |
| NTRK3 | 15q25.3 | 110 | 4.34 | 35 | 6.14 | 0.045 | 0.397 | (B) BRAF Class 2 |
| FGF19 | 11q13.3 | 24 | 1.47 | 11 | 2.95 | 0.0457 | 0.397 | (B) BRAF Class 2 |
| CSDE1 | 1p13.2 | 10 | 1.19 | 6 | 3.37 | 0.0458 | 0.397 | (B) BRAF Class 2 |
| BRD4 | 19p13.12 | 126 | 5.24 | 19 | 3.45 | 0.0459 | 0.397 | (A) BRAF Class 1 v12 |
| ELANE | 19p13.3 | 15 | 2.30 | 0 | 0.00 | 0.0462 | 0.397 | (A) BRAF Class 1 v12 |
| FBXW7 | 4q31.3 | 186 | 5.67 | 28 | 4.03 | 0.0466 | 0.397 | (A) BRAF Class 1 v12 |
| GALNT12 | 9q22.33 | 12 | 1.63 | 7 | 4.14 | 0.0472 | 0.4 | (B) BRAF Class 2 |
| ANKRD11 | 16q24.3 | 87 | 7.85 | 12 | 4.71 | 0.0486 | 0.408 | (A) BRAF Class 1 v12 |
| NIPBL | 5p13.2 | 6 | 3.64 | 4 | 13.33 | 0.0494 | 0.412 | (B) BRAF Class 2 |
| NSD3 | 8p11.23 | 30 | 2.67 | 14 | 4.83 | 0.0503 | 0.414 | (B) BRAF Class 2 |
| ESPL1 | 12q13.13 | 1 | 1.16 | 2 | 14.29 | 0.0506 | 0.414 | (B) BRAF Class 2 |
| MLH1 | 3p22.2 | 52 | 1.62 | 18 | 2.65 | 0.051 | 0.414 | (B) BRAF Class 2 |
| PKHD1 | 6p12.3-p12.2 | 1 | 0.85 | 2 | 10.53 | 0.0512 | 0.414 | (B) BRAF Class 2 |
| NCOA2 | 8q13.3 | 6 | 2.34 | 4 | 8.51 | 0.0526 | 0.422 | (B) BRAF Class 2 |
| SLIT2 | 4p15.31 | 52 | 8.28 | 14 | 14.00 | 0.0535 | 0.427 | (B) BRAF Class 2 |
| KLLN | 10q23.31 | 14 | 2.40 | 0 | 0.00 | 0.0547 | 0.433 | (A) BRAF Class 1 v12 |
| ZNF708 | 19p12 | 3 | 0.85 | 3 | 4.41 | 0.0561 | 0.438 | (B) BRAF Class 2 |
| TYK2 | 19p13.2 | 6 | 2.41 | 4 | 8.51 | 0.0567 | 0.438 | (B) BRAF Class 2 |

| | | | | | | | | |
|---|---|---|---|---|---|---|---|---|
| BEND2 | Xp22.13 | 1 | 1.09 | 2 | 12.50 | 0.0568 | 0.438 | (B) BRAF Class 2 |
| CDKN2B | 9p21.3 | 208 | 8.22 | 59 | 10.42 | 0.0569 | 0.438 | (B) BRAF Class 2 |
| TBX3 | 12q24.21 | 69 | 4.53 | 24 | 6.76 | 0.0573 | 0.439 | (B) BRAF Class 2 |
| FGF14 | 13q33.1 | 13 | 1.84 | 6 | 4.65 | 0.0579 | 0.439 | (B) BRAF Class 2 |
| RSPO3 | 6q22.33 | 20 | 2.56 | 9 | 5.29 | 0.0582 | 0.439 | (B) BRAF Class 2 |
| FGF8 | 10q24.32 | 2 | 0.60 | 2 | 5.00 | 0.0586 | 0.439 | (B) BRAF Class 2 |
| GLI2 | 2q14.2 | 80 | 7.63 | 25 | 11.11 | 0.0598 | 0.444 | (B) BRAF Class 2 |
| RICTOR | 5p13.1 | 74 | 3.37 | 25 | 4.96 | 0.0601 | 0.444 | (B) BRAF Class 2 |
| RHPN2 | 19q13.11 | 6 | 1.70 | 4 | 5.88 | 0.0609 | 0.445 | (B) BRAF Class 2 |
| IPMK | 10q21.1 | 1 | 0.64 | 2 | 7.14 | 0.0609 | 0.445 | (B) BRAF Class 2 |
| E2F3 | 6p22.3 | 10 | 0.85 | 6 | 2.20 | 0.0627 | 0.454 | (B) BRAF Class 2 |
| PDGFB | 22q13.1 | 1 | 0.39 | 2 | 4.26 | 0.0638 | 0.454 | (B) BRAF Class 2 |
| TCF12 | 15q21.3 | 1 | 0.50 | 2 | 5.41 | 0.0638 | 0.454 | (B) BRAF Class 2 |
| DCC | 18q21.2 | 14 | 5.58 | 6 | 13.33 | 0.0642 | 0.454 | (B) BRAF Class 2 |
| RASA1 | 5q14.3 | 39 | 2.08 | 15 | 3.49 | 0.0643 | 0.454 | (B) BRAF Class 2 |
| GLI1 | 12q13.3 | 114 | 4.97 | 17 | 3.32 | 0.0651 | 0.456 | (A) BRAF Class 1 v12 |
| EBF1 | 5q33.3 | 1 | 0.44 | 2 | 4.76 | 0.0654 | 0.456 | (B) BRAF Class 2 |
| TRAF3 | 14q32.32 | 14 | 1.73 | 7 | 3.93 | 0.0666 | 0.462 | (B) BRAF Class 2 |
| TEK | 9p21.2 | 33 | 4.65 | 15 | 7.77 | 0.0672 | 0.462 | (B) BRAF Class 2 |
| PNKP | 19q13.33 | 13 | 2.23 | 0 | 0.00 | 0.0674 | 0.462 | (A) BRAF Class 1 v12 |
| AURKB | 17p13.1 | 23 | 0.94 | 10 | 1.79 | 0.0702 | 0.477 | (B) BRAF Class 2 |
| MSH6 | 2p16.3 | 139 | 5.41 | 22 | 3.83 | 0.0709 | 0.477 | (A) BRAF Class 1 v12 |
| KIT | 4q12 | 80 | 2.39 | 26 | 3.43 | 0.0712 | 0.477 | (B) BRAF Class 2 |
| LIFR | 5p13.1 | 4 | 2.15 | 3 | 9.09 | 0.0714 | 0.477 | (B) BRAF Class 2 |
| H3F3A | 1q42.12 | 14 | 0.56 | 7 | 1.26 | 0.074 | 0.491 | (B) BRAF Class 2 |
| SPOP | 17q21.33 | 20 | 0.91 | 9 | 1.78 | 0.0757 | 0.494 | (B) BRAF Class 2 |
| MCM8 | 20p12.3 | 14 | 2.40 | 7 | 5.26 | 0.0757 | 0.494 | (B) BRAF Class 2 |
| SOX2 | 3q26.33 | 33 | 1.34 | 3 | 0.53 | 0.076 | 0.494 | (A) BRAF Class 1 v12 |
| FOXP1 | 3p13 | 34 | 2.26 | 13 | 3.81 | 0.0771 | 0.494 | (B) BRAF Class 2 |
| HIST3H3 | 1q42.13 | 8 | 0.72 | 5 | 1.96 | 0.0773 | 0.494 | (B) BRAF Class 2 |
| COL1A1 | 17q21.33 | 4 | 1.56 | 3 | 6.38 | 0.0779 | 0.494 | (B) BRAF Class 2 |
| PAK5 | 20p12.2 | 79 | 6.16 | 26 | 8.67 | 0.078 | 0.494 | (B) BRAF Class 2 |
| HGF | 7q21.11 | 61 | 3.92 | 21 | 5.79 | 0.078 | 0.494 | (B) BRAF Class 2 |
| TNFRSF14 | 1p36.32 | 17 | 1.09 | 8 | 2.22 | 0.0795 | 0.499 | (B) BRAF Class 2 |
| PRPF40B | 12q13.12 | 10 | 2.31 | 5 | 5.95 | 0.0797 | 0.499 | (B) BRAF Class 2 |
| FANCA | 16q24.3 | 90 | 3.55 | 28 | 4.91 | 0.0814 | 0.507 | (B) BRAF Class 2 |
| DNMT3A | 2p23.3 | 127 | 5.11 | 38 | 6.69 | 0.0833 | 0.516 | (B) BRAF Class 2 |
| MT-ND5 | - | 12 | 0.36 | 0 | 0.00 | 0.0856 | 0.527 | (A) BRAF Class 1 v12 |
| TFRC | 3q29 | 7 | 2.07 | 3 | 7.14 | 0.0868 | 0.53 | (B) BRAF Class 2 |
| PRAME | 22q11.22 | 4 | 1.13 | 3 | 4.41 | 0.087 | 0.53 | (B) BRAF Class 2 |
| PALB2 | 16p12.2 | 87 | 3.43 | 27 | 4.74 | 0.0876 | 0.531 | (B) BRAF Class 2 |
| NAV3 | 12q21.2 | 8 | 5.13 | 4 | 14.29 | 0.0892 | 0.538 | (B) BRAF Class 2 |
| PTEN | 10q23.31 | 350 | 10.47 | 62 | 8.72 | 0.0899 | 0.538 | (A) BRAF Class 1 v12 |
| RPS6KB2 | 11q13.2 | 21 | 1.78 | 9 | 3.30 | 0.0937 | 0.538 | (B) BRAF Class 2 |
| ZFHX3 | 6q22.2-q22. | 152 | 12.24 | 26 | 9.25 | 0.0939 | 0.538 | (A) BRAF Class 1 v12 |
| PRKDC | 8q11.21 | 170 | 10.79 | 42 | 13.59 | 0.0939 | 0.538 | (B) BRAF Class 2 |
| ARID2 | 12q12 | 180 | 7.42 | 50 | 9.21 | 0.0951 | 0.538 | (B) BRAF Class 2 |
| U2AF1 | 21q22.3 | 39 | 1.55 | 14 | 2.46 | 0.0952 | 0.538 | (B) BRAF Class 2 |

| | | | | | | | | |
|---|---|---|---|---|---|---|---|---|
| HFE | 6p22.2 | 6 | 1.03 | 4 | 3.01 | 0.0957 | 0.538 | (B) BRAF Class 2 |
| INHBA | 7p14.1 | 43 | 2.96 | 15 | 4.59 | 0.0959 | 0.538 | (B) BRAF Class 2 |
| BRCA2 | 13q13.1 | 203 | 8.01 | 36 | 6.30 | 0.096 | 0.538 | (A) BRAF Class 1 v12 |
| SF1 | 11q13.1 | 13 | 3.01 | 0 | 0.00 | 0.0963 | 0.538 | (A) BRAF Class 1 v12 |
| TRAF2 | 9q34.3 | 25 | 2.13 | 2 | 0.75 | 0.0969 | 0.538 | (A) BRAF Class 1 v12 |
| CDKN1A | 6p21.2 | 15 | 0.63 | 7 | 1.28 | 0.0975 | 0.538 | (B) BRAF Class 2 |
| CHEK2 | 22q12.1 | 60 | 2.37 | 8 | 1.40 | 0.0983 | 0.538 | (A) BRAF Class 1 v12 |
| HIST1H3I | 6p22.1 | 6 | 0.54 | 4 | 1.57 | 0.0985 | 0.538 | (B) BRAF Class 2 |
| ITK | 5q33.3 | 12 | 2.01 | 6 | 4.38 | 0.0998 | 0.538 | (B) BRAF Class 2 |
| BCL2 | 18q21.33 | 12 | 0.49 | 6 | 1.08 | 0.1 | 0.538 | (B) BRAF Class 2 |
| SDHD | 11q23.1 | 9 | 0.37 | 5 | 0.90 | 0.1 | 0.538 | (B) BRAF Class 2 |
| IRS1 | 2q36.3 | 63 | 5.35 | 9 | 3.30 | 0.101 | 0.538 | (A) BRAF Class 1 v12 |
| NOTCH3 | 19p13.12 | 219 | 10.01 | 40 | 8.03 | 0.101 | 0.538 | (A) BRAF Class 1 v12 |
| FLI1 | 11q24.3 | 7 | 1.64 | 3 | 5.26 | 0.101 | 0.538 | (B) BRAF Class 2 |
| KAT6A | 8p11.21 | 68 | 5.42 | 19 | 7.76 | 0.103 | 0.538 | (B) BRAF Class 2 |
| SMARCD1 | 12q13.12 | 24 | 1.87 | 2 | 0.67 | 0.103 | 0.538 | (A) BRAF Class 1 v12 |
| SMAD2 | 18q21.1 | 42 | 1.71 | 5 | 0.89 | 0.106 | 0.538 | (A) BRAF Class 1 v12 |
| ERCC2 | 19q13.32 | 75 | 3.02 | 23 | 4.17 | 0.107 | 0.538 | (B) BRAF Class 2 |
| NKX2-1-AS1 | 14q13.3 | 0 | 0.00 | 1 | 2.50 | 0.107 | 0.538 | (B) BRAF Class 2 |
| RASA2 | 3q23 | 2 | 0.91 | 2 | 5.13 | 0.11 | 0.538 | (B) BRAF Class 2 |
| TTYH1 | 19q13.42 | 2 | 1.28 | 2 | 7.14 | 0.11 | 0.538 | (B) BRAF Class 2 |
| DDR1 | 6p21.33 | 3 | 1.46 | 3 | 5.45 | 0.111 | 0.538 | (B) BRAF Class 2 |
| IKBKE | 1q32.1 | 22 | 1.41 | 9 | 2.48 | 0.112 | 0.538 | (B) BRAF Class 2 |
| NR4A3 | 9q31.1 | 2 | 1.25 | 2 | 6.90 | 0.112 | 0.538 | (B) BRAF Class 2 |
| SYK | 9q22.2 | 44 | 2.33 | 15 | 3.50 | 0.113 | 0.538 | (B) BRAF Class 2 |
| PTP4A1 | 6q12 | 1 | 0.19 | 2 | 1.43 | 0.114 | 0.538 | (B) BRAF Class 2 |
| XPO1 | 2p15 | 48 | 1.89 | 16 | 2.80 | 0.114 | 0.538 | (B) BRAF Class 2 |
| USP6 | 17p13.2 | 2 | 1.18 | 2 | 6.45 | 0.114 | 0.538 | (B) BRAF Class 2 |
| PPP4R2 | 3p13 | 8 | 1.52 | 5 | 3.57 | 0.116 | 0.538 | (B) BRAF Class 2 |
| BTG1 | 12q21.33 | 2 | 0.30 | 2 | 1.63 | 0.116 | 0.538 | (B) BRAF Class 2 |
| AXIN1 | 16p13.3 | 56 | 3.70 | 8 | 2.26 | 0.116 | 0.538 | (A) BRAF Class 1 v12 |
| RUNX1 | 21q22.12 | 68 | 2.73 | 21 | 3.79 | 0.117 | 0.538 | (B) BRAF Class 2 |
| NSD2 | 4p16.3 | 44 | 3.24 | 16 | 4.78 | 0.118 | 0.538 | (B) BRAF Class 2 |
| CNOT3 | 19q13.42 | 2 | 1.18 | 2 | 6.25 | 0.119 | 0.538 | (B) BRAF Class 2 |
| TOPBP1 | 3q22.1 | 23 | 3.89 | 9 | 6.67 | 0.12 | 0.538 | (B) BRAF Class 2 |
| TET2 | 4q24 | 156 | 6.28 | 44 | 7.75 | 0.12 | 0.538 | (B) BRAF Class 2 |
| JAK2 | 9p24.1 | 63 | 1.98 | 20 | 2.77 | 0.121 | 0.538 | (B) BRAF Class 2 |
| KLHL6 | 3q27.1 | 21 | 2.93 | 1 | 0.76 | 0.122 | 0.538 | (A) BRAF Class 1 v12 |
| RINT1 | 7q22.3 | 13 | 2.23 | 6 | 4.51 | 0.122 | 0.538 | (B) BRAF Class 2 |
| PRKCZ | 1p36.33 | 5 | 1.42 | 3 | 4.41 | 0.123 | 0.538 | (B) BRAF Class 2 |
| FGF23 | 12p13.32 | 9 | 1.27 | 4 | 3.10 | 0.126 | 0.538 | (B) BRAF Class 2 |
| HIST1H3H | 6p22.1 | 4 | 0.36 | 3 | 1.18 | 0.126 | 0.538 | (B) BRAF Class 2 |
| DMC1 | 22q13.1 | 10 | 1.72 | 0 | 0.00 | 0.126 | 0.538 | (A) BRAF Class 1 v12 |
| CIC | 19q13.2 | 149 | 7.01 | 27 | 5.47 | 0.127 | 0.538 | (A) BRAF Class 1 v12 |
| PNRC1 | 6q15 | 17 | 1.05 | 1 | 0.27 | 0.127 | 0.538 | (A) BRAF Class 1 v12 |
| SLC25A13 | 7q21.3 | 10 | 1.72 | 5 | 3.76 | 0.127 | 0.538 | (B) BRAF Class 2 |
| RAD50 | 5q31.1 | 94 | 4.32 | 15 | 3.08 | 0.129 | 0.538 | (A) BRAF Class 1 v12 |
| ETS1 | 11q24.3 | 7 | 1.24 | 3 | 3.57 | 0.129 | 0.538 | (B) BRAF Class 2 |

| | | | | | | | | |
|---|---|---|---|---|---|---|---|---|
| PRSS1 | 7q34 | 20 | 2.74 | 8 | 4.82 | 0.129 | 0.538 | (B) BRAF Class 2 |
| MAP3K4 | 6q26 | 30 | 9.01 | 1 | 2.50 | 0.129 | 0.538 | (A) BRAF Class 1 v12 |
| PCM1 | 8p22 | 0 | 0.00 | 1 | 25.00 | 0.129 | 0.538 | (B) BRAF Class 2 |
| ADGRL3 | 4q13.1 | 0 | 0.00 | 1 | 12.50 | 0.129 | 0.538 | (B) BRAF Class 2 |
| AKT1 | 14q32.33 | 108 | 3.23 | 18 | 2.37 | 0.129 | 0.538 | (A) BRAF Class 1 v12 |
| GBA | 1q22 | 7 | 1.20 | 4 | 3.01 | 0.129 | 0.538 | (B) BRAF Class 2 |
| SDHB | 1p36.13 | 10 | 0.41 | 5 | 0.90 | 0.129 | 0.538 | (B) BRAF Class 2 |
| TOP2A | 17q21.2 | 32 | 5.10 | 2 | 2.00 | 0.13 | 0.538 | (A) BRAF Class 1 v12 |
| FLT1 | 13q12.3 | 130 | 5.29 | 37 | 6.61 | 0.131 | 0.538 | (B) BRAF Class 2 |
| AXL | 19q13.2 | 96 | 3.79 | 28 | 4.91 | 0.132 | 0.538 | (B) BRAF Class 2 |
| PDCD1 | 2q37.3 | 33 | 2.57 | 4 | 1.31 | 0.132 | 0.538 | (A) BRAF Class 1 v12 |
| CHEK1 | 11q24.2 | 27 | 1.23 | 10 | 1.98 | 0.134 | 0.538 | (B) BRAF Class 2 |
| EML4 | 2p21 | 13 | 3.00 | 4 | 6.78 | 0.134 | 0.538 | (B) BRAF Class 2 |
| PTCH1 | 9q22.32 | 133 | 5.25 | 23 | 4.03 | 0.135 | 0.538 | (A) BRAF Class 1 v12 |
| SERPINB4 | 18q21.33 | 0 | 0.00 | 1 | 20.00 | 0.135 | 0.538 | (B) BRAF Class 2 |
| SDHA | 5p15.33 | 71 | 3.12 | 11 | 2.10 | 0.135 | 0.538 | (A) BRAF Class 1 v12 |
| SH2B3 | 12q24.12 | 62 | 2.65 | 9 | 1.71 | 0.135 | 0.538 | (A) BRAF Class 1 v12 |
| EXT2 | 11p11.2 | 18 | 1.76 | 7 | 3.18 | 0.137 | 0.538 | (B) BRAF Class 2 |
| BCL2L12 | 19q13.33 | 11 | 1.12 | 5 | 2.37 | 0.138 | 0.538 | (B) BRAF Class 2 |
| FZR1 | 19p13.3 | 0 | 0.00 | 1 | 5.26 | 0.14 | 0.538 | (B) BRAF Class 2 |
| LTF | 3p21.31 | 0 | 0.00 | 1 | 5.26 | 0.14 | 0.538 | (B) BRAF Class 2 |
| SOX11 | 2p25.2 | 0 | 0.00 | 1 | 5.26 | 0.14 | 0.538 | (B) BRAF Class 2 |
| MALT1 | 18q21.32 | 17 | 1.40 | 7 | 2.54 | 0.14 | 0.538 | (B) BRAF Class 2 |
| CAD | 2p23.3 | 0 | 0.00 | 1 | 7.14 | 0.14 | 0.538 | (B) BRAF Class 2 |
| CPS1 | 2q34 | 0 | 0.00 | 1 | 7.14 | 0.14 | 0.538 | (B) BRAF Class 2 |
| EDNRB | 13q22.3 | 0 | 0.00 | 1 | 7.14 | 0.14 | 0.538 | (B) BRAF Class 2 |
| HSPA2 | 14q23.3 | 0 | 0.00 | 1 | 7.14 | 0.14 | 0.538 | (B) BRAF Class 2 |
| INPP5D | 2q37.1 | 0 | 0.00 | 1 | 7.14 | 0.14 | 0.538 | (B) BRAF Class 2 |
| MEF2C | 5q14.3 | 0 | 0.00 | 1 | 7.14 | 0.14 | 0.538 | (B) BRAF Class 2 |
| MKI67 | 10q26.2 | 0 | 0.00 | 1 | 7.14 | 0.14 | 0.538 | (B) BRAF Class 2 |
| PRKCH | 14q23.1 | 0 | 0.00 | 1 | 7.14 | 0.14 | 0.538 | (B) BRAF Class 2 |
| PTPN6 | 12p13.31 | 0 | 0.00 | 1 | 7.14 | 0.14 | 0.538 | (B) BRAF Class 2 |
| TNFRSF11A | 18q21.33 | 0 | 0.00 | 1 | 7.14 | 0.14 | 0.538 | (B) BRAF Class 2 |
| TRIM28 | 19q13.43 | 0 | 0.00 | 1 | 7.14 | 0.14 | 0.538 | (B) BRAF Class 2 |
| WDR90 | 16p13.3 | 0 | 0.00 | 1 | 7.14 | 0.14 | 0.538 | (B) BRAF Class 2 |
| NF2 | 22q12.2 | 58 | 2.26 | 18 | 3.13 | 0.14 | 0.538 | (B) BRAF Class 2 |
| PAX5 | 9p13.2 | 53 | 2.13 | 17 | 2.99 | 0.141 | 0.538 | (B) BRAF Class 2 |
| HDAC9 | 7p21.1 | 3 | 3.49 | 2 | 14.29 | 0.143 | 0.538 | (B) BRAF Class 2 |
| PLCG1 | 20q12 | 0 | 0.00 | 1 | 5.00 | 0.143 | 0.538 | (B) BRAF Class 2 |
| BOD1L1 | 4p15.33 | 0 | 0.00 | 1 | 12.50 | 0.143 | 0.538 | (B) BRAF Class 2 |
| JARID2 | 6p22.3 | 0 | 0.00 | 1 | 6.67 | 0.144 | 0.538 | (B) BRAF Class 2 |
| ICK | 6p12.1 | 0 | 0.00 | 1 | 5.00 | 0.145 | 0.538 | (B) BRAF Class 2 |
| HABP2 | 10q25.3 | 16 | 2.74 | 1 | 0.75 | 0.145 | 0.538 | (A) BRAF Class 1 v12 |
| PIM1 | 6p21.2 | 17 | 0.77 | 7 | 1.38 | 0.146 | 0.538 | (B) BRAF Class 2 |
| AKAP9 | 7q21.2 | 0 | 0.00 | 1 | 4.76 | 0.147 | 0.538 | (B) BRAF Class 2 |
| PBX1 | 1q23.3 | 0 | 0.00 | 1 | 4.76 | 0.147 | 0.538 | (B) BRAF Class 2 |
| TRIP11 | 14q32.12 | 0 | 0.00 | 1 | 4.76 | 0.147 | 0.538 | (B) BRAF Class 2 |
| POLE | 12q24.33 | 162 | 7.33 | 30 | 5.89 | 0.147 | 0.538 | (A) BRAF Class 1 v12 |

| | | | | | | | | |
|---|---|---|---|---|---|---|---|---|
| ACVR1B | 12q13.13 | 10 | 1.52 | 4 | 3.42 | 0.147 | 0.538 | (B) BRAF Class 2 |
| RHEB | 7q36.1 | 11 | 0.52 | 5 | 1.07 | 0.148 | 0.538 | (B) BRAF Class 2 |
| HTRA1 | 10q26.13 | 0 | 0.00 | 1 | 6.25 | 0.148 | 0.538 | (B) BRAF Class 2 |
| SOX9 | 17q24.3 | 110 | 4.59 | 19 | 3.47 | 0.149 | 0.538 | (A) BRAF Class 1 v12 |
| EXO1 | 1q43 | 14 | 2.40 | 6 | 4.51 | 0.149 | 0.538 | (B) BRAF Class 2 |
| SHOC2 | 10q25.2 | 8 | 1.52 | 0 | 0.00 | 0.15 | 0.538 | (A) BRAF Class 1 v12 |
| COL7A1 | 3p21.31 | 94 | 16.12 | 27 | 20.30 | 0.151 | 0.538 | (B) BRAF Class 2 |
| SHQ1 | 3p13 | 24 | 2.04 | 9 | 3.30 | 0.151 | 0.538 | (B) BRAF Class 2 |
| ASH2L | 8p11.23 | 0 | 0.00 | 1 | 3.57 | 0.152 | 0.538 | (B) BRAF Class 2 |
| CRCT1 | 1q21.3 | 0 | 0.00 | 1 | 3.57 | 0.152 | 0.538 | (B) BRAF Class 2 |
| MAP3K5 | 6q23.3 | 0 | 0.00 | 1 | 3.57 | 0.152 | 0.538 | (B) BRAF Class 2 |
| DNM2 | 19p13.2 | 0 | 0.00 | 1 | 5.56 | 0.153 | 0.538 | (B) BRAF Class 2 |
| STAG2 | Xq25 | 73 | 2.98 | 22 | 3.93 | 0.153 | 0.538 | (B) BRAF Class 2 |
| SS18 | 18q11.2 | 8 | 1.01 | 4 | 2.30 | 0.153 | 0.538 | (B) BRAF Class 2 |
| TERC | 3q26.2 | 0 | 0.00 | 1 | 0.73 | 0.153 | 0.538 | (B) BRAF Class 2 |
| SETD2 | 3p21.31 | 223 | 8.80 | 42 | 7.37 | 0.154 | 0.538 | (A) BRAF Class 1 v12 |
| MEF2B | 19p13.11 | 28 | 1.14 | 10 | 1.79 | 0.154 | 0.538 | (B) BRAF Class 2 |
| LEF1 | 4q25 | 0 | 0.00 | 1 | 2.50 | 0.154 | 0.538 | (B) BRAF Class 2 |
| KDM5C | Xp11.22 | 81 | 3.42 | 24 | 4.44 | 0.154 | 0.538 | (B) BRAF Class 2 |
| MST1 | 3p21.31 | 9 | 0.81 | 0 | 0.00 | 0.154 | 0.538 | (A) BRAF Class 1 v12 |
| MN1 | 22q12.1 | 0 | 0.00 | 1 | 2.86 | 0.154 | 0.538 | (B) BRAF Class 2 |
| HIST1H1E | 6p22.2 | 0 | 0.00 | 1 | 4.76 | 0.154 | 0.538 | (B) BRAF Class 2 |
| RPTOR | 17q25.3 | 87 | 4.11 | 26 | 5.26 | 0.155 | 0.538 | (B) BRAF Class 2 |
| MAP2K2 | 19p13.3 | 32 | 1.45 | 12 | 2.17 | 0.155 | 0.538 | (B) BRAF Class 2 |
| NIN | 14q22.1 | 3 | 2.46 | 2 | 9.52 | 0.156 | 0.538 | (B) BRAF Class 2 |
| CARD11 | 7p22.2 | 157 | 6.39 | 43 | 7.68 | 0.156 | 0.538 | (B) BRAF Class 2 |
| NRG1 | 8p12 | 52 | 5.06 | 14 | 7.14 | 0.156 | 0.538 | (B) BRAF Class 2 |
| KMT2A | 11q23.3 | 214 | 8.62 | 57 | 10.05 | 0.158 | 0.538 | (B) BRAF Class 2 |
| PGBD3 | 10q11.23 | 11 | 0.33 | 5 | 0.66 | 0.159 | 0.538 | (B) BRAF Class 2 |
| PIK3CD | 1p36.22 | 38 | 2.99 | 5 | 1.72 | 0.161 | 0.538 | (A) BRAF Class 1 v12 |
| IDH2 | 15q26.1 | 48 | 1.51 | 7 | 0.95 | 0.164 | 0.538 | (A) BRAF Class 1 v12 |
| EPHA2 | 1p36.13 | 0 | 0.00 | 1 | 2.38 | 0.164 | 0.538 | (B) BRAF Class 2 |
| ARHGEF12 | 11q23.3 | 25 | 4.25 | 9 | 6.67 | 0.165 | 0.538 | (B) BRAF Class 2 |
| GREM1 | 15q13.3 | 11 | 0.56 | 5 | 1.11 | 0.165 | 0.538 | (B) BRAF Class 2 |
| EGFL7 | 9q34.3 | 15 | 1.27 | 1 | 0.37 | 0.166 | 0.538 | (A) BRAF Class 1 v12 |
| BABAM1 | 19p13.11 | 14 | 1.29 | 1 | 0.37 | 0.167 | 0.538 | (A) BRAF Class 1 v12 |
| CRKL | 22q11.21 | 18 | 0.73 | 7 | 1.25 | 0.167 | 0.538 | (B) BRAF Class 2 |
| SOS2 | 14q21.3 | 3 | 1.92 | 2 | 7.14 | 0.167 | 0.538 | (B) BRAF Class 2 |
| NECTIN4 | 1q23.3 | 8 | 1.37 | 4 | 3.01 | 0.167 | 0.538 | (B) BRAF Class 2 |
| NT5C2 | q24.32-q24. | 11 | 1.33 | 5 | 2.59 | 0.168 | 0.538 | (B) BRAF Class 2 |
| HOXB13 | 17q21.32 | 18 | 0.94 | 7 | 1.59 | 0.171 | 0.538 | (B) BRAF Class 2 |
| MYCL | 1p34.2 | 33 | 1.30 | 11 | 1.93 | 0.172 | 0.538 | (B) BRAF Class 2 |
| EGLN1 | 1q42.2 | 5 | 0.86 | 3 | 2.26 | 0.172 | 0.538 | (B) BRAF Class 2 |
| INPP4B | 4q31.21 | 32 | 2.08 | 11 | 3.07 | 0.174 | 0.538 | (B) BRAF Class 2 |
| DIS3 | 13q21.33 | 70 | 2.96 | 21 | 3.85 | 0.174 | 0.538 | (B) BRAF Class 2 |
| RYBP | 3p13 | 8 | 0.68 | 4 | 1.47 | 0.174 | 0.538 | (B) BRAF Class 2 |
| CTNNB1 | 3p22.1 | 115 | 3.44 | 32 | 4.22 | 0.175 | 0.538 | (B) BRAF Class 2 |
| PTPRS | 19p13.3 | 110 | 9.35 | 20 | 7.33 | 0.176 | 0.538 | (A) BRAF Class 1 v12 |

| | | | | | | | | |
|---|---|---|---|---|---|---|---|---|
| MST1R | 3p21.31 | 46 | 3.58 | 15 | 4.92 | 0.176 | 0.538 | (B) BRAF Class 2 |
| STAT5B | 17q21.2 | 37 | 3.11 | 5 | 1.84 | 0.177 | 0.538 | (A) BRAF Class 1 v12 |
| FAN1 | 15q13.3 | 15 | 2.57 | 6 | 4.51 | 0.178 | 0.538 | (B) BRAF Class 2 |
| PRPF8 | 17p13.3 | 14 | 3.24 | 5 | 5.95 | 0.182 | 0.538 | (B) BRAF Class 2 |
| EMSY | 11q13.5 | 15 | 2.12 | 5 | 3.88 | 0.182 | 0.538 | (B) BRAF Class 2 |
| CHD4 | 12p13.31 | 38 | 6.05 | 9 | 9.00 | 0.182 | 0.538 | (B) BRAF Class 2 |
| ADGRV1 | 5q14.3 | 0 | 0.00 | 1 | 0.13 | 0.185 | 0.538 | (B) BRAF Class 2 |
| ALG10 | 12p11.1 | 0 | 0.00 | 1 | 0.13 | 0.185 | 0.538 | (B) BRAF Class 2 |
| C5ORF22 | 5p13.3 | 0 | 0.00 | 1 | 0.13 | 0.185 | 0.538 | (B) BRAF Class 2 |
| CALB1 | 8q21.3 | 0 | 0.00 | 1 | 0.13 | 0.185 | 0.538 | (B) BRAF Class 2 |
| CAMKMT | 2p21 | 0 | 0.00 | 1 | 0.13 | 0.185 | 0.538 | (B) BRAF Class 2 |
| CRLF3 | 17q11.2 | 0 | 0.00 | 1 | 0.13 | 0.185 | 0.538 | (B) BRAF Class 2 |
| FAM124B | 2q36.2 | 0 | 0.00 | 1 | 0.13 | 0.185 | 0.538 | (B) BRAF Class 2 |
| HEXIM2 | 17q21.31 | 0 | 0.00 | 1 | 0.13 | 0.185 | 0.538 | (B) BRAF Class 2 |
| IGHD2-2 | 14q32.33 | 0 | 0.00 | 1 | 0.13 | 0.185 | 0.538 | (B) BRAF Class 2 |
| IRX3 | 16q12.2 | 0 | 0.00 | 1 | 0.13 | 0.185 | 0.538 | (B) BRAF Class 2 |
| KCNAB1 | 3q25.31 | 0 | 0.00 | 1 | 0.13 | 0.185 | 0.538 | (B) BRAF Class 2 |
| KCNJ1 | 11q24.3 | 0 | 0.00 | 1 | 0.13 | 0.185 | 0.538 | (B) BRAF Class 2 |
| KYNU | 2q22.2 | 0 | 0.00 | 1 | 0.13 | 0.185 | 0.538 | (B) BRAF Class 2 |
| LAMC3 | 9q34.12 | 0 | 0.00 | 1 | 0.13 | 0.185 | 0.538 | (B) BRAF Class 2 |
| LGALS17A | 19q13.2 | 0 | 0.00 | 1 | 0.13 | 0.185 | 0.538 | (B) BRAF Class 2 |
| LILRB1 | 19q13.42 | 0 | 0.00 | 1 | 0.13 | 0.185 | 0.538 | (B) BRAF Class 2 |
| LINC00484 | 9q22.2 | 0 | 0.00 | 1 | 0.13 | 0.185 | 0.538 | (B) BRAF Class 2 |
| LINC02544 | 6q27 | 0 | 0.00 | 1 | 0.13 | 0.185 | 0.538 | (B) BRAF Class 2 |
| MTMR3 | 22q12.2 | 0 | 0.00 | 1 | 0.13 | 0.185 | 0.538 | (B) BRAF Class 2 |
| NBPF1 | 1p36.13 | 0 | 0.00 | 1 | 0.13 | 0.185 | 0.538 | (B) BRAF Class 2 |
| NUFIP1 | 13q14.12 | 0 | 0.00 | 1 | 0.13 | 0.185 | 0.538 | (B) BRAF Class 2 |
| PAXIP1-AS2 | 7q36.2 | 0 | 0.00 | 1 | 0.13 | 0.185 | 0.538 | (B) BRAF Class 2 |
| PEX5 | 12p13.31 | 0 | 0.00 | 1 | 0.13 | 0.185 | 0.538 | (B) BRAF Class 2 |
| PRAG1 | 8p23.1 | 0 | 0.00 | 1 | 0.13 | 0.185 | 0.538 | (B) BRAF Class 2 |
| PXMP2 | 12q24.33 | 0 | 0.00 | 1 | 0.13 | 0.185 | 0.538 | (B) BRAF Class 2 |
| RERE | 1p36.23 | 0 | 0.00 | 1 | 0.13 | 0.185 | 0.538 | (B) BRAF Class 2 |
| ROCK1 | 18q11.1 | 0 | 0.00 | 1 | 0.13 | 0.185 | 0.538 | (B) BRAF Class 2 |
| RUNX3 | 1p36.11 | 0 | 0.00 | 1 | 0.13 | 0.185 | 0.538 | (B) BRAF Class 2 |
| SBF1P1 | 8q12.1 | 0 | 0.00 | 1 | 0.13 | 0.185 | 0.538 | (B) BRAF Class 2 |
| SEC24B-AS1 | 4q25 | 0 | 0.00 | 1 | 0.13 | 0.185 | 0.538 | (B) BRAF Class 2 |
| SELENOI | 2p23.3 | 0 | 0.00 | 1 | 0.13 | 0.185 | 0.538 | (B) BRAF Class 2 |
| ST7 | 7q31.2 | 0 | 0.00 | 1 | 0.13 | 0.185 | 0.538 | (B) BRAF Class 2 |
| TECRL | 4q13.1 | 0 | 0.00 | 1 | 0.13 | 0.185 | 0.538 | (B) BRAF Class 2 |
| TMEM132C | q24.32-q24. | 0 | 0.00 | 1 | 0.13 | 0.185 | 0.538 | (B) BRAF Class 2 |
| TTC28 | 22q12.1 | 0 | 0.00 | 1 | 0.13 | 0.185 | 0.538 | (B) BRAF Class 2 |
| VPREB1 | 22q11.22 | 0 | 0.00 | 1 | 0.13 | 0.185 | 0.538 | (B) BRAF Class 2 |
| VRK2 | 2p16.1 | 0 | 0.00 | 1 | 0.13 | 0.185 | 0.538 | (B) BRAF Class 2 |
| NFKBIA | 14q13.2 | 26 | 1.09 | 3 | 0.55 | 0.186 | 0.538 | (A) BRAF Class 1 v12 |
| MAPK1 | 22q11.22 | 19 | 0.79 | 7 | 1.29 | 0.186 | 0.538 | (B) BRAF Class 2 |
| ROBO1 | 3p12.3 | 7 | 4.40 | 3 | 10.34 | 0.186 | 0.538 | (B) BRAF Class 2 |
| FGF4 | 11q13.3 | 29 | 1.86 | 10 | 2.75 | 0.187 | 0.538 | (B) BRAF Class 2 |
| ARFRP1 | 20q13.33 | 7 | 1.00 | 3 | 2.36 | 0.187 | 0.538 | (B) BRAF Class 2 |

| | | | | | | | | |
|---|---|---|---|---|---|---|---|---|
| BCL2L11 | 2q13 | 8 | 0.68 | 0 | 0.00 | 0.187 | 0.538 | (A) BRAF Class 1 v12 |
| CDK6 | 7q21.2 | 19 | 0.75 | 7 | 1.23 | 0.187 | 0.538 | (B) BRAF Class 2 |
| UIMC1 | 5q35.2 | 14 | 2.32 | 1 | 0.70 | 0.188 | 0.538 | (A) BRAF Class 1 v12 |
| CBL | 11q23.3 | 57 | 2.22 | 17 | 2.94 | 0.188 | 0.538 | (B) BRAF Class 2 |
| PPP6C | 9q33.3 | 42 | 3.31 | 6 | 2.09 | 0.188 | 0.538 | (A) BRAF Class 1 v12 |
| COP1 | .q25.1-q25.[illegible] | 34 | 2.25 | 11 | 3.24 | 0.188 | 0.538 | (B) BRAF Class 2 |
| FAT3 | 11q14.3 | 21 | 8.14 | 7 | 12.96 | 0.19 | 0.539 | (B) BRAF Class 2 |
| DKC1 | Xq28 | 8 | 1.37 | 0 | 0.00 | 0.19 | 0.539 | (A) BRAF Class 1 v12 |
| FANCB | Xp22.2 | 20 | 3.00 | 2 | 1.31 | 0.19 | 0.539 | (A) BRAF Class 1 v12 |
| KLF2 | 19p13.11 | 8 | 1.37 | 0 | 0.00 | 0.191 | 0.542 | (A) BRAF Class 1 v12 |
| RAD54L | 1p34.1 | 19 | 1.45 | 2 | 0.63 | 0.193 | 0.545 | (A) BRAF Class 1 v12 |
| BRCA1 | 17q21.31 | 101 | 3.98 | 30 | 4.85 | 0.196 | 0.549 | (B) BRAF Class 2 |
| CTLA4 | 2q33.2 | 25 | 1.43 | 3 | 0.74 | 0.196 | 0.549 | (A) BRAF Class 1 v12 |
| TFE3 | Xp11.23 | 30 | 2.64 | 3 | 1.38 | 0.196 | 0.55 | (A) BRAF Class 1 v12 |
| AR | Xq12 | 120 | 4.73 | 35 | 5.65 | 0.197 | 0.55 | (B) BRAF Class 2 |
| SMC3 | 10q25.2 | 27 | 1.85 | 8 | 2.82 | 0.197 | 0.55 | (B) BRAF Class 2 |
| DHX15 | 4p15.2 | 14 | 4.20 | 0 | 0.00 | 0.198 | 0.551 | (A) BRAF Class 1 v12 |
| SMARCB1 | q11.23\|22q | 41 | 1.28 | 12 | 1.78 | 0.199 | 0.552 | (B) BRAF Class 2 |
| KAT6B | 10q22.2 | 43 | 6.30 | 13 | 8.55 | 0.201 | 0.552 | (B) BRAF Class 2 |
| RXRA | 9q34.2 | 17 | 3.23 | 2 | 1.43 | 0.201 | 0.552 | (A) BRAF Class 1 v12 |
| RIT1 | 1q22 | 25 | 1.28 | 3 | 0.67 | 0.203 | 0.552 | (A) BRAF Class 1 v12 |
| RB1 | 13q14.2 | 105 | 3.26 | 27 | 3.98 | 0.203 | 0.552 | (B) BRAF Class 2 |
| MTA1 | 14q32.33 | 25 | 4.29 | 3 | 2.26 | 0.204 | 0.552 | (A) BRAF Class 1 v12 |
| ATP1B2 | 17p13.1 | 0 | 0.00 | 1 | 9.09 | 0.204 | 0.552 | (B) BRAF Class 2 |
| CDH12 | 5p14.3 | 0 | 0.00 | 1 | 9.09 | 0.204 | 0.552 | (B) BRAF Class 2 |
| PRKAR1A | 17q24.2 | 23 | 0.94 | 8 | 1.43 | 0.204 | 0.552 | (B) BRAF Class 2 |
| SMAD4 | 18q21.2 | 230 | 7.02 | 42 | 6.05 | 0.204 | 0.552 | (A) BRAF Class 1 v12 |
| GRM3 | [illegible]21.11-q21.[illegible] | 58 | 8.81 | 7 | 5.98 | 0.204 | 0.552 | (A) BRAF Class 1 v12 |
| RAD52 | 12p13.33 | 24 | 1.28 | 3 | 0.67 | 0.205 | 0.553 | (A) BRAF Class 1 v12 |
| AKT3 | 1q43-q44 | 35 | 1.36 | 11 | 1.92 | 0.205 | 0.553 | (B) BRAF Class 2 |
| ERCC1 | 19q13.32 | 7 | 0.66 | 3 | 1.49 | 0.207 | 0.554 | (B) BRAF Class 2 |
| CUX1 | 7q22.1 | 117 | 8.20 | 18 | 6.52 | 0.207 | 0.554 | (A) BRAF Class 1 v12 |
| MYD88 | 3p22.2 | 31 | 1.21 | 10 | 1.74 | 0.208 | 0.554 | (B) BRAF Class 2 |
| CDC73 | 1q31.2 | 42 | 1.71 | 13 | 2.32 | 0.208 | 0.554 | (B) BRAF Class 2 |
| CRTC2 | 1q21.3 | 7 | 1.98 | 3 | 4.41 | 0.209 | 0.555 | (B) BRAF Class 2 |
| JAZF1 | 'p15.2-p15.[illegible] | 6 | 0.76 | 3 | 1.74 | 0.209 | 0.555 | (B) BRAF Class 2 |
| IKZF3 | 17q12-q21.1 | 7 | 1.26 | 3 | 2.80 | 0.21 | 0.556 | (B) BRAF Class 2 |
| DDX41 | 5q35.3 | 5 | 1.07 | 2 | 3.03 | 0.211 | 0.556 | (B) BRAF Class 2 |
| NSD1 | 5q35.3 | 126 | 6.05 | 34 | 7.16 | 0.211 | 0.556 | (B) BRAF Class 2 |
| CD276 | 15q24.1 | 19 | 1.61 | 2 | 0.73 | 0.213 | 0.56 | (A) BRAF Class 1 v12 |
| ASXL2 | 2p23.3 | 58 | 4.41 | 17 | 5.67 | 0.215 | 0.563 | (B) BRAF Class 2 |
| ERCC3 | 2q14.3 | 39 | 1.80 | 12 | 2.45 | 0.215 | 0.563 | (B) BRAF Class 2 |
| GRIN2A | 16p13.2 | 157 | 10.27 | 41 | 11.85 | 0.221 | 0.575 | (B) BRAF Class 2 |
| KLF4 | 9q31.2 | 35 | 1.80 | 11 | 2.48 | 0.223 | 0.58 | (B) BRAF Class 2 |
| TP53BP1 | 15q15.3 | 59 | 5.00 | 11 | 3.73 | 0.225 | 0.582 | (A) BRAF Class 1 v12 |
| TAP2 | 6p21.32 | 5 | 0.95 | 3 | 2.14 | 0.225 | 0.582 | (B) BRAF Class 2 |
| ELOC | 8q21.11 | 6 | 0.33 | 3 | 0.73 | 0.225 | 0.582 | (B) BRAF Class 2 |
| PARP3 | 3p21.2 | 0 | 0.00 | 1 | 2.22 | 0.226 | 0.582 | (B) BRAF Class 2 |

| | | | | | | | | |
|---|---|---|---|---|---|---|---|---|
| LRP1B | !q22.1-q22.! | 146 | 21.35 | 28 | 25.00 | 0.226 | 0.582 | (B) BRAF Class 2 |
| MLH3 | 14q24.3 | 29 | 3.60 | 9 | 5.11 | 0.227 | 0.582 | (B) BRAF Class 2 |
| FANCI | 15q26.1 | 47 | 4.13 | 12 | 5.50 | 0.227 | 0.582 | (B) BRAF Class 2 |
| HIST1H3G | 6p22.2 | 6 | 0.54 | 3 | 1.17 | 0.228 | 0.583 | (B) BRAF Class 2 |
| SPEN | ›36.21-p36.: | 142 | 9.11 | 28 | 7.71 | 0.231 | 0.585 | (A) BRAF Class 1 v12 |
| CADM2 | 3p12.1 | 8 | 3.04 | 3 | 6.25 | 0.232 | 0.585 | (B) BRAF Class 2 |
| CBFA2T3 | 16q24.3 | 24 | 4.08 | 8 | 5.93 | 0.233 | 0.585 | (B) BRAF Class 2 |
| RACK1 | 5q35.3 | 4 | 1.52 | 2 | 4.17 | 0.233 | 0.585 | (B) BRAF Class 2 |
| MYOD1 | 11p15.1 | 29 | 2.32 | 4 | 1.39 | 0.233 | 0.585 | (A) BRAF Class 1 v12 |
| MAP2K4 | 17p12 | 40 | 1.58 | 12 | 2.11 | 0.233 | 0.585 | (B) BRAF Class 2 |
| ENG | 9q34.11 | 13 | 2.23 | 1 | 0.75 | 0.233 | 0.585 | (A) BRAF Class 1 v12 |
| FAH | 15q25.1 | 13 | 2.23 | 1 | 0.75 | 0.233 | 0.585 | (A) BRAF Class 1 v12 |
| KMT2D | 12q13.12 | 362 | 14.76 | 90 | 16.07 | 0.234 | 0.586 | (B) BRAF Class 2 |
| ETV6 | 12p13.2 | 51 | 2.09 | 15 | 2.69 | 0.235 | 0.586 | (B) BRAF Class 2 |
| NPM1 | 5q35.1 | 22 | 0.71 | 7 | 1.06 | 0.238 | 0.591 | (B) BRAF Class 2 |
| JUN | 1p32.1 | 23 | 1.61 | 3 | 0.89 | 0.24 | 0.591 | (A) BRAF Class 1 v12 |
| LAMP1 | 13q34 | 6 | 1.40 | 2 | 3.51 | 0.24 | 0.591 | (B) BRAF Class 2 |
| LMO1 | 11p15.4 | 17 | 0.75 | 6 | 1.17 | 0.24 | 0.591 | (B) BRAF Class 2 |
| PRF1 | 10q22.1 | 30 | 3.23 | 4 | 1.97 | 0.241 | 0.591 | (A) BRAF Class 1 v12 |
| PTK2 | 8q24.3 | 8 | 2.27 | 0 | 0.00 | 0.241 | 0.591 | (A) BRAF Class 1 v12 |
| RBL2 | 16q12.2 | 8 | 2.27 | 0 | 0.00 | 0.241 | 0.591 | (A) BRAF Class 1 v12 |
| RAC2 | 22q13.1 | 6 | 1.14 | 0 | 0.00 | 0.241 | 0.591 | (A) BRAF Class 1 v12 |
| TRAF7 | 16p13.3 | 44 | 2.27 | 7 | 1.58 | 0.241 | 0.591 | (A) BRAF Class 1 v12 |
| TET3 | 2p13.1 | 1 | 2.94 | 1 | 20.00 | 0.243 | 0.592 | (B) BRAF Class 2 |
| CCND3 | 6p21.1 | 21 | 0.83 | 7 | 1.23 | 0.243 | 0.592 | (B) BRAF Class 2 |
| KMT2B | 19q13.12 | 121 | 11.67 | 21 | 9.72 | 0.244 | 0.594 | (A) BRAF Class 1 v12 |
| PHF6 | Xq26.2 | 20 | 1.33 | 6 | 2.03 | 0.246 | 0.596 | (B) BRAF Class 2 |
| GATA4 | 8p23.1 | 32 | 2.26 | 4 | 1.39 | 0.246 | 0.596 | (A) BRAF Class 1 v12 |
| TSC1 | 9q34.13 | 81 | 3.19 | 22 | 3.85 | 0.247 | 0.597 | (B) BRAF Class 2 |
| INPP4A | 2q11.2 | 23 | 1.95 | 3 | 1.10 | 0.249 | 0.598 | (A) BRAF Class 1 v12 |
| DMD | (p21.2-p21.: | 12 | 3.40 | 4 | 5.88 | 0.249 | 0.598 | (B) BRAF Class 2 |
| AKT2 | 19q13.2 | 49 | 1.91 | 14 | 2.44 | 0.249 | 0.598 | (B) BRAF Class 2 |
| ABCB11 | 2q31.1 | 33 | 5.05 | 10 | 6.80 | 0.252 | 0.599 | (B) BRAF Class 2 |
| EME1 | 17q21.33 | 10 | 1.72 | 4 | 3.01 | 0.252 | 0.599 | (B) BRAF Class 2 |
| XRCC4 | 5q14.2 | 10 | 1.72 | 4 | 3.01 | 0.252 | 0.599 | (B) BRAF Class 2 |
| CSF3R | 1p34.3 | 52 | 2.35 | 15 | 2.97 | 0.253 | 0.599 | (B) BRAF Class 2 |
| MDM4 | 1q32.1 | 29 | 1.14 | 9 | 1.58 | 0.253 | 0.599 | (B) BRAF Class 2 |
| LZTR1 | 21\|22q11.1 | 25 | 3.89 | 6 | 5.77 | 0.254 | 0.599 | (B) BRAF Class 2 |
| WRN | 8p12 | 52 | 4.49 | 14 | 5.69 | 0.255 | 0.599 | (B) BRAF Class 2 |
| SUZ12 | 17q11.2 | 33 | 1.47 | 10 | 1.98 | 0.256 | 0.599 | (B) BRAF Class 2 |
| PAXIP1 | 7q36.2 | 25 | 4.29 | 8 | 6.02 | 0.257 | 0.599 | (B) BRAF Class 2 |
| DEPDC5 | 2q12.2-q12. | 13 | 4.94 | 4 | 8.33 | 0.258 | 0.599 | (B) BRAF Class 2 |
| MCL1 | 1q21.2 | 25 | 1.02 | 8 | 1.43 | 0.258 | 0.599 | (B) BRAF Class 2 |
| INSR | 19p13.2 | 57 | 4.31 | 10 | 3.27 | 0.258 | 0.599 | (A) BRAF Class 1 v12 |
| TLR4 | 9q33.1 | 13 | 3.08 | 4 | 5.19 | 0.259 | 0.599 | (B) BRAF Class 2 |
| XPA | 9q22.33 | 13 | 1.27 | 1 | 0.45 | 0.261 | 0.599 | (A) BRAF Class 1 v12 |
| ACVR2A | !q22.3-q23.: | 1 | 0.85 | 1 | 5.26 | 0.261 | 0.599 | (B) BRAF Class 2 |
| TGM7 | 5q15.2-q15. | 1 | 0.85 | 1 | 5.26 | 0.261 | 0.599 | (B) BRAF Class 2 |

| | | | | | | | | |
|---|---|---|---|---|---|---|---|---|
| MAP3K13 | 3q27.2 | 20 | 1.52 | 7 | 2.21 | 0.261 | 0.599 | (B) BRAF Class 2 |
| CDH1 | 16q22.1 | 72 | 2.30 | 12 | 1.80 | 0.261 | 0.599 | (A) BRAF Class 1 v12 |
| APOBEC3G | 22q13.1 | 1 | 1.16 | 1 | 7.14 | 0.262 | 0.599 | (B) BRAF Class 2 |
| PC | 11q13.2 | 1 | 1.16 | 1 | 7.14 | 0.262 | 0.599 | (B) BRAF Class 2 |
| PCLO | 7q21.11 | 1 | 1.16 | 1 | 7.14 | 0.262 | 0.599 | (B) BRAF Class 2 |
| PDCD11 | 10q24.33 | 1 | 1.16 | 1 | 7.14 | 0.262 | 0.599 | (B) BRAF Class 2 |
| VHL | 3p25.3 | 27 | 0.86 | 8 | 1.20 | 0.263 | 0.601 | (B) BRAF Class 2 |
| ABL2 | 1q25.2 | 20 | 3.01 | 5 | 4.63 | 0.263 | 0.601 | (B) BRAF Class 2 |
| FANCC | 9q22.32 | 38 | 1.55 | 6 | 1.07 | 0.265 | 0.602 | (A) BRAF Class 1 v12 |
| POLQ | 3q13.33 | 54 | 8.35 | 15 | 10.34 | 0.266 | 0.602 | (B) BRAF Class 2 |
| BCOR | Xp11.4 | 156 | 6.36 | 31 | 5.54 | 0.266 | 0.602 | (A) BRAF Class 1 v12 |
| DNMT1 | 19p13.2 | 53 | 4.05 | 15 | 5.03 | 0.266 | 0.602 | (B) BRAF Class 2 |
| BCR | 22q11.23 | 22 | 3.82 | 2 | 1.94 | 0.267 | 0.603 | (A) BRAF Class 1 v12 |
| TLR2 | 4q31.3 | 1 | 2.08 | 1 | 12.50 | 0.268 | 0.603 | (B) BRAF Class 2 |
| RPL10 | Xq28 | 1 | 1.05 | 1 | 6.25 | 0.269 | 0.603 | (B) BRAF Class 2 |
| PAK1 | 1q13.5-q14. | 14 | 1.07 | 5 | 1.67 | 0.269 | 0.603 | (B) BRAF Class 2 |
| GNA11 | 19p13.3 | 25 | 0.77 | 8 | 1.07 | 0.269 | 0.604 | (B) BRAF Class 2 |
| EPHB1 | 3q22.2 | 64 | 4.09 | 18 | 4.96 | 0.271 | 0.604 | (B) BRAF Class 2 |
| USP8 | 15q21.2 | 14 | 2.36 | 5 | 3.68 | 0.271 | 0.604 | (B) BRAF Class 2 |
| PAX3 | 2q36.1 | 11 | 1.94 | 3 | 3.53 | 0.271 | 0.604 | (B) BRAF Class 2 |
| TP63 | 3q28 | 57 | 4.57 | 16 | 5.59 | 0.275 | 0.607 | (B) BRAF Class 2 |
| WWTR1 | 3q25.1 | 3 | 0.50 | 2 | 1.28 | 0.275 | 0.607 | (B) BRAF Class 2 |
| RAD21 | 8q24.11 | 57 | 2.51 | 10 | 1.93 | 0.275 | 0.607 | (A) BRAF Class 1 v12 |
| FRK | 6q22.1 | 1 | 1.09 | 1 | 6.25 | 0.276 | 0.607 | (B) BRAF Class 2 |
| GLMN | 1p22.1 | 1 | 1.09 | 1 | 6.25 | 0.276 | 0.607 | (B) BRAF Class 2 |
| ABRAXAS1 | 4q21.23 | 17 | 0.89 | 2 | 0.45 | 0.276 | 0.607 | (A) BRAF Class 1 v12 |
| CTCF | 16q22.1 | 51 | 2.41 | 9 | 1.82 | 0.277 | 0.607 | (A) BRAF Class 1 v12 |
| HELQ | 4q21.23 | 18 | 3.09 | 6 | 4.51 | 0.277 | 0.607 | (B) BRAF Class 2 |
| HMBS | 11q23.3 | 7 | 1.20 | 3 | 2.26 | 0.279 | 0.61 | (B) BRAF Class 2 |
| HSP90AB1 | 6p21.1 | 1 | 0.40 | 1 | 2.22 | 0.281 | 0.612 | (B) BRAF Class 2 |
| MGMT | 10q26.3 | 1 | 0.64 | 1 | 3.57 | 0.282 | 0.612 | (B) BRAF Class 2 |
| SHH | 7q36.3 | 1 | 0.64 | 1 | 3.57 | 0.282 | 0.612 | (B) BRAF Class 2 |
| SPRY2 | 13q31.1 | 1 | 0.64 | 1 | 3.57 | 0.282 | 0.612 | (B) BRAF Class 2 |
| ERF | 19q13.2 | 9 | 1.71 | 4 | 2.86 | 0.283 | 0.612 | (B) BRAF Class 2 |
| RBM15 | 1p13.3 | 1 | 0.86 | 1 | 4.76 | 0.284 | 0.612 | (B) BRAF Class 2 |
| FANCD2 | 3p25.3 | 59 | 3.85 | 14 | 4.73 | 0.284 | 0.612 | (B) BRAF Class 2 |
| HNF1A | 12q24.31 | 106 | 3.36 | 26 | 3.87 | 0.285 | 0.612 | (B) BRAF Class 2 |
| NPRL3 | 16p13.3 | 1 | 0.38 | 1 | 2.08 | 0.285 | 0.612 | (B) BRAF Class 2 |
| PMAIP1 | 18q21.32 | 6 | 0.51 | 0 | 0.00 | 0.285 | 0.612 | (A) BRAF Class 1 v12 |
| PDPK1 | 16p13.3 | 6 | 0.48 | 0 | 0.00 | 0.288 | 0.614 | (A) BRAF Class 1 v12 |
| PLAG1 | 8q12.1 | 1 | 0.38 | 1 | 2.04 | 0.288 | 0.614 | (B) BRAF Class 2 |
| PARP1 | 1q42.12 | 44 | 3.13 | 13 | 3.89 | 0.288 | 0.614 | (B) BRAF Class 2 |
| ERCC6 | 10q11.23 | 34 | 5.83 | 10 | 7.52 | 0.289 | 0.614 | (B) BRAF Class 2 |
| PMS1 | 2q32.2 | 55 | 2.39 | 15 | 2.91 | 0.289 | 0.614 | (B) BRAF Class 2 |
| NUF2 | 1q23.3 | 10 | 1.90 | 1 | 0.71 | 0.291 | 0.615 | (A) BRAF Class 1 v12 |
| FGFR3 | 4p16.3 | 91 | 2.73 | 24 | 3.16 | 0.291 | 0.615 | (B) BRAF Class 2 |
| MTAP | 9p21.3 | 39 | 6.26 | 12 | 7.84 | 0.292 | 0.615 | (B) BRAF Class 2 |
| NKX2-1 | 14q13.3 | 58 | 2.36 | 16 | 2.85 | 0.292 | 0.615 | (B) BRAF Class 2 |

| | | | | | | | | |
|---|---|---|---|---|---|---|---|---|
| INSRR | 1q23.1 | 6 | 0.18 | 0 | 0.00 | 0.293 | 0.616 | (A) BRAF Class 1 v12 |
| PIK3CB | 3q22.3 | 46 | 3.11 | 8 | 2.35 | 0.295 | 0.619 | (A) BRAF Class 1 v12 |
| CHD2 | 15q26.1 | 38 | 6.04 | 8 | 7.92 | 0.296 | 0.621 | (B) BRAF Class 2 |
| PIK3C2B | 1q32.1 | 94 | 6.49 | 16 | 5.44 | 0.3 | 0.625 | (A) BRAF Class 1 v12 |
| PIK3R1 | 5q13.1 | 68 | 2.65 | 18 | 3.13 | 0.3 | 0.625 | (B) BRAF Class 2 |
| BUB1B | 15q15.1 | 24 | 2.35 | 7 | 3.18 | 0.302 | 0.625 | (B) BRAF Class 2 |
| MGA | 15q15.1 | 180 | 9.97 | 37 | 8.96 | 0.302 | 0.625 | (A) BRAF Class 1 v12 |
| FOXO1 | 13q14.11 | 27 | 1.99 | 4 | 1.32 | 0.303 | 0.625 | (A) BRAF Class 1 v12 |
| FUS | 16p11.2 | 16 | 1.51 | 5 | 2.21 | 0.303 | 0.625 | (B) BRAF Class 2 |
| FANCE | 6p21.31 | 19 | 1.20 | 2 | 0.64 | 0.303 | 0.625 | (A) BRAF Class 1 v12 |
| REL | 2p16.1 | 41 | 1.68 | 7 | 1.25 | 0.303 | 0.625 | (A) BRAF Class 1 v12 |
| EIF4A2 | 3q27.3 | 11 | 0.99 | 4 | 1.56 | 0.305 | 0.625 | (B) BRAF Class 2 |
| FRS2 | 12q15 | 6 | 0.95 | 2 | 1.98 | 0.306 | 0.625 | (B) BRAF Class 2 |
| MAPKAP1 | 9q33.3 | 5 | 0.95 | 0 | 0.00 | 0.306 | 0.625 | (A) BRAF Class 1 v12 |
| SRSF2 | 17q25.1 | 27 | 1.20 | 4 | 0.80 | 0.306 | 0.625 | (A) BRAF Class 1 v12 |
| PRKCI | 3q26.2 | 48 | 2.50 | 13 | 3.05 | 0.307 | 0.625 | (B) BRAF Class 2 |
| INPPL1 | 11q13.4 | 31 | 5.89 | 6 | 4.29 | 0.308 | 0.625 | (A) BRAF Class 1 v12 |
| MAX | 14q23.3 | 15 | 0.79 | 5 | 1.16 | 0.308 | 0.625 | (B) BRAF Class 2 |
| PPP2R2A | 8p21.2 | 8 | 2.20 | 0 | 0.00 | 0.308 | 0.625 | (A) BRAF Class 1 v12 |
| TCF7L2 | 0q25.2-q25. | 76 | 3.40 | 14 | 2.82 | 0.308 | 0.625 | (A) BRAF Class 1 v12 |
| BCORL1 | Xq26.1 | 114 | 7.05 | 20 | 6.08 | 0.308 | 0.625 | (A) BRAF Class 1 v12 |
| BIRC3 | 11q22.2 | 15 | 1.24 | 5 | 1.81 | 0.309 | 0.625 | (B) BRAF Class 2 |
| UBE2T | 1q32.1 | 4 | 0.69 | 2 | 1.50 | 0.31 | 0.625 | (B) BRAF Class 2 |
| ORCS8-MEF2 | 19p13.11 | 19 | 0.57 | 6 | 0.79 | 0.311 | 0.626 | (B) BRAF Class 2 |
| PRDM14 | 8q13.3 | 14 | 2.66 | 2 | 1.43 | 0.312 | 0.628 | (A) BRAF Class 1 v12 |
| HIST1H3F | 6p22.2 | 4 | 0.36 | 2 | 0.78 | 0.313 | 0.628 | (B) BRAF Class 2 |
| FAS | 10q23.31 | 18 | 1.23 | 2 | 0.67 | 0.314 | 0.628 | (A) BRAF Class 1 v12 |
| BTK | Xq22.1 | 27 | 1.65 | 8 | 2.15 | 0.314 | 0.628 | (B) BRAF Class 2 |
| BARD1 | 2q35 | 38 | 1.79 | 11 | 2.23 | 0.314 | 0.628 | (B) BRAF Class 2 |
| INHA | 2q35 | 21 | 1.90 | 3 | 1.18 | 0.316 | 0.629 | (A) BRAF Class 1 v12 |
| KCNQ1 | 1p15.5-p15. | 26 | 4.46 | 4 | 3.01 | 0.317 | 0.629 | (A) BRAF Class 1 v12 |
| FGF5 | 4q21.21 | 10 | 3.00 | 0 | 0.00 | 0.317 | 0.629 | (A) BRAF Class 1 v12 |
| FGF3 | 11q13.3 | 27 | 1.65 | 8 | 2.14 | 0.317 | 0.629 | (B) BRAF Class 2 |
| APC | 5q22.2 | 355 | 11.32 | 86 | 12.01 | 0.319 | 0.632 | (B) BRAF Class 2 |
| TCF3 | 19p13.3 | 65 | 3.08 | 12 | 2.53 | 0.322 | 0.637 | (A) BRAF Class 1 v12 |
| MITF | 3p13 | 64 | 2.61 | 12 | 2.14 | 0.324 | 0.639 | (A) BRAF Class 1 v12 |
| FAT1 | 4q35.2 | 262 | 12.87 | 64 | 13.76 | 0.326 | 0.642 | (B) BRAF Class 2 |
| MDM2 | 12q15 | 44 | 1.74 | 12 | 2.10 | 0.327 | 0.642 | (B) BRAF Class 2 |
| IGF1R | 15q26.3 | 111 | 4.33 | 28 | 4.84 | 0.327 | 0.642 | (B) BRAF Class 2 |
| ARID1B | 6q25.3 | 172 | 7.26 | 42 | 7.91 | 0.33 | 0.646 | (B) BRAF Class 2 |
| ASXL1 | 20q11.21 | 131 | 5.27 | 33 | 5.82 | 0.332 | 0.649 | (B) BRAF Class 2 |
| KDM5A | 12p13.33 | 120 | 5.69 | 31 | 6.30 | 0.332 | 0.649 | (B) BRAF Class 2 |
| RTEL1 | 20q13.33 | 20 | 3.78 | 7 | 4.96 | 0.334 | 0.65 | (B) BRAF Class 2 |
| TDG | 12q23.3 | 8 | 1.37 | 3 | 2.26 | 0.335 | 0.652 | (B) BRAF Class 2 |
| IRF2 | 4q35.1 | 6 | 1.15 | 2 | 2.20 | 0.336 | 0.653 | (B) BRAF Class 2 |
| GATA2 | 3q21.3 | 50 | 2.30 | 9 | 1.84 | 0.338 | 0.655 | (A) BRAF Class 1 v12 |
| ERRFI1 | 1p36.23 | 25 | 1.74 | 4 | 1.20 | 0.339 | 0.655 | (A) BRAF Class 1 v12 |
| XRCC2 | 7q36.1 | 23 | 1.23 | 7 | 1.59 | 0.339 | 0.655 | (B) BRAF Class 2 |

| | | | | | | | | |
|---|---|---|---|---|---|---|---|---|
| PAK3 | Xq23 | 17 | 2.49 | 4 | 3.57 | 0.341 | 0.657 | (B) BRAF Class 2 |
| RECQL4 | 8q24.3 | 120 | 5.58 | 24 | 4.97 | 0.341 | 0.657 | (A) BRAF Class 1 v12 |
| PIK3CA | 3q26.32 | 312 | 9.33 | 75 | 9.88 | 0.342 | 0.657 | (B) BRAF Class 2 |
| CUL3 | 2q36.2 | 20 | 1.31 | 3 | 0.84 | 0.342 | 0.657 | (A) BRAF Class 1 v12 |
| CARM1 | 19p13.2 | 9 | 1.71 | 1 | 0.71 | 0.344 | 0.657 | (A) BRAF Class 1 v12 |
| BRCC3 | Xq28 | 12 | 2.03 | 4 | 2.96 | 0.344 | 0.657 | (B) BRAF Class 2 |
| FKBP9 | 7p14.3 | 6 | 1.70 | 0 | 0.00 | 0.345 | 0.657 | (A) BRAF Class 1 v12 |
| CEBPA | 19q13.11 | 28 | 1.17 | 8 | 1.47 | 0.346 | 0.657 | (B) BRAF Class 2 |
| MECOM | 3q26.2 | 70 | 7.34 | 13 | 6.22 | 0.346 | 0.657 | (A) BRAF Class 1 v12 |
| FANCG | 9p13.3 | 27 | 1.67 | 7 | 2.13 | 0.347 | 0.657 | (B) BRAF Class 2 |
| PIK3C2G | 12p12.3 | 81 | 6.17 | 22 | 6.94 | 0.347 | 0.657 | (B) BRAF Class 2 |
| IGF2 | 11p15.5 | 24 | 1.15 | 7 | 1.47 | 0.348 | 0.657 | (B) BRAF Class 2 |
| RELA | 11q13.1 | 16 | 2.05 | 2 | 1.18 | 0.348 | 0.657 | (A) BRAF Class 1 v12 |
| ROS1 | 6q22.1 | 241 | 9.27 | 55 | 8.66 | 0.349 | 0.657 | (A) BRAF Class 1 v12 |
| CREBBP | 16p13.3 | 193 | 7.61 | 40 | 7.02 | 0.35 | 0.657 | (A) BRAF Class 1 v12 |
| CDKN1B | 12p13.1 | 35 | 1.38 | 6 | 1.05 | 0.35 | 0.657 | (A) BRAF Class 1 v12 |
| AMER1 | Xq11.2 | 64 | 4.09 | 17 | 4.68 | 0.35 | 0.658 | (B) BRAF Class 2 |
| SOCS1 | 16p13.13 | 25 | 1.02 | 4 | 0.71 | 0.352 | 0.659 | (A) BRAF Class 1 v12 |
| ERCC4 | 16p13.12 | 52 | 2.28 | 14 | 2.65 | 0.353 | 0.66 | (B) BRAF Class 2 |
| ZMYM3 | Xq13.1 | 6 | 2.73 | 2 | 5.00 | 0.355 | 0.663 | (B) BRAF Class 2 |
| PRDM1 | 6q21 | 90 | 3.66 | 18 | 3.21 | 0.356 | 0.663 | (A) BRAF Class 1 v12 |
| RNF8 | 6p21.2 | 5 | 0.86 | 0 | 0.00 | 0.357 | 0.663 | (A) BRAF Class 1 v12 |
| DDB2 | 11p11.2 | 16 | 1.56 | 2 | 0.91 | 0.357 | 0.663 | (A) BRAF Class 1 v12 |
| CCND1 | 11q13.3 | 34 | 1.35 | 10 | 1.62 | 0.358 | 0.664 | (B) BRAF Class 2 |
| PHF1 | 6p21.32 | 5 | 0.15 | 0 | 0.00 | 0.359 | 0.664 | (A) BRAF Class 1 v12 |
| ICOSLG | 21q22.3 | 16 | 1.36 | 5 | 1.83 | 0.36 | 0.664 | (B) BRAF Class 2 |
| MTOR | 1p36.22 | 144 | 5.53 | 38 | 5.97 | 0.361 | 0.664 | (B) BRAF Class 2 |
| TGFBR1 | 9q22.33 | 20 | 1.70 | 6 | 2.20 | 0.362 | 0.664 | (B) BRAF Class 2 |
| KCNIP1 | 5q35.1 | 6 | 2.28 | 0 | 0.00 | 0.362 | 0.664 | (A) BRAF Class 1 v12 |
| PIK3CG | 7q22.3 | 75 | 4.89 | 19 | 5.49 | 0.363 | 0.664 | (B) BRAF Class 2 |
| STAT4 | [illegible]q32.2-q32.3 | 23 | 3.39 | 5 | 4.46 | 0.364 | 0.664 | (B) BRAF Class 2 |
| VEGFA | 6p21.1 | 20 | 0.99 | 6 | 1.28 | 0.364 | 0.664 | (B) BRAF Class 2 |
| CDK8 | 13q12.13 | 36 | 1.69 | 10 | 2.02 | 0.364 | 0.664 | (B) BRAF Class 2 |
| MTR | 1q43 | 2 | 1.71 | 1 | 5.26 | 0.366 | 0.664 | (B) BRAF Class 2 |
| CHD1 | 5q15-q21.1 | 6 | 3.85 | 0 | 0.00 | 0.366 | 0.664 | (A) BRAF Class 1 v12 |
| FGF2 | 4q28.1 | 3 | 0.90 | 1 | 2.50 | 0.366 | 0.664 | (B) BRAF Class 2 |
| RIF1 | 2q23.3 | 41 | 7.03 | 11 | 8.27 | 0.367 | 0.664 | (B) BRAF Class 2 |
| DYNC1I1 | 7q21.3 | 2 | 2.33 | 1 | 7.14 | 0.367 | 0.664 | (B) BRAF Class 2 |
| TSHZ3 | 19q12 | 2 | 2.33 | 1 | 7.14 | 0.367 | 0.664 | (B) BRAF Class 2 |
| PRSS8 | 16p11.2 | 7 | 1.36 | 0 | 0.00 | 0.369 | 0.667 | (A) BRAF Class 1 v12 |
| PIK3C3 | 18q12.3 | 33 | 2.52 | 6 | 1.96 | 0.37 | 0.667 | (A) BRAF Class 1 v12 |
| ADGRA2 | 8p11.23 | 35 | 5.64 | 7 | 6.93 | 0.37 | 0.667 | (B) BRAF Class 2 |
| BCL6 | 3q27.3 | 57 | 2.32 | 11 | 1.96 | 0.372 | 0.668 | (A) BRAF Class 1 v12 |
| MEN1 | 11q13.1 | 57 | 2.32 | 11 | 1.96 | 0.372 | 0.668 | (A) BRAF Class 1 v12 |
| DOT1L | 19p13.3 | 78 | 5.01 | 16 | 4.41 | 0.375 | 0.669 | (A) BRAF Class 1 v12 |
| SMAD3 | 15q22.33 | 25 | 1.70 | 7 | 2.10 | 0.376 | 0.669 | (B) BRAF Class 2 |
| PIK3R2 | 19p13.11 | 34 | 2.22 | 6 | 1.73 | 0.376 | 0.669 | (A) BRAF Class 1 v12 |
| PCDHAC2 | 5q31.3 | 2 | 4.17 | 1 | 12.50 | 0.376 | 0.669 | (B) BRAF Class 2 |

| | | | | | | | | |
|---|---|---|---|---|---|---|---|---|
| TNFAIP3 | 6q23.3 | 58 | 2.36 | 15 | 2.68 | 0.376 | 0.669 | (B) BRAF Class 2 |
| PTPRO | 12.3\|12p13· | 1 | 0.86 | 1 | 3.23 | 0.378 | 0.672 | (B) BRAF Class 2 |
| PDE4DIP | 1q21.2 | 6 | 4.92 | 0 | 0.00 | 0.379 | 0.672 | (A) BRAF Class 1 v12 |
| FAAP100 | 17q25.3 | 24 | 4.12 | 4 | 3.01 | 0.381 | 0.675 | (A) BRAF Class 1 v12 |
| SMARCE1 | 17q21.2 | 5 | 0.74 | 2 | 1.30 | 0.383 | 0.677 | (B) BRAF Class 2 |
| GATA3 | 10p14 | 71 | 2.89 | 18 | 3.21 | 0.386 | 0.681 | (B) BRAF Class 2 |
| GNAQ | 9q21.2 | 29 | 0.89 | 8 | 1.07 | 0.387 | 0.681 | (B) BRAF Class 2 |
| CENPA | 2p23.3 | 5 | 0.45 | 2 | 0.78 | 0.388 | 0.681 | (B) BRAF Class 2 |
| EIF4E | 4q23 | 5 | 0.45 | 2 | 0.78 | 0.388 | 0.681 | (B) BRAF Class 2 |
| PTPRT | 0q12-q13.1 | 159 | 11.31 | 38 | 12.03 | 0.39 | 0.683 | (B) BRAF Class 2 |
| POLB | 8p11.21 | 9 | 1.54 | 3 | 2.26 | 0.391 | 0.683 | (B) BRAF Class 2 |
| UROD | 1p34.1 | 9 | 1.54 | 3 | 2.26 | 0.391 | 0.683 | (B) BRAF Class 2 |
| RAD51C | 17q22 | 25 | 1.15 | 7 | 1.40 | 0.391 | 0.683 | (B) BRAF Class 2 |
| RNF213 | 17q25.3 | 7 | 5.74 | 2 | 9.52 | 0.392 | 0.683 | (B) BRAF Class 2 |
| AJUBA | 14q11.2 | 2 | 1.28 | 1 | 3.57 | 0.392 | 0.683 | (B) BRAF Class 2 |
| CYLD | 16q12.1 | 49 | 2.36 | 12 | 2.69 | 0.393 | 0.683 | (B) BRAF Class 2 |
| SLX4 | 16p13.3 | 98 | 6.64 | 23 | 7.21 | 0.394 | 0.683 | (B) BRAF Class 2 |
| CAMTA1 | 36.31-p36.2 | 2 | 1.25 | 1 | 3.45 | 0.395 | 0.684 | (B) BRAF Class 2 |
| HIST1H3C | 6p22.2 | 9 | 0.52 | 3 | 0.75 | 0.395 | 0.684 | (B) BRAF Class 2 |
| MAP3K7 | 6q15 | 2 | 0.79 | 1 | 2.17 | 0.396 | 0.685 | (B) BRAF Class 2 |
| AXIN2 | 17q24.1 | 83 | 4.29 | 17 | 3.85 | 0.399 | 0.688 | (A) BRAF Class 1 v12 |
| H3F3C | 12p11.21 | 9 | 0.76 | 3 | 1.10 | 0.4 | 0.688 | (B) BRAF Class 2 |
| CYSLTR2 | 13q14.2 | 8 | 1.22 | 3 | 1.76 | 0.4 | 0.688 | (B) BRAF Class 2 |
| VTCN1 | 1p13.1-p12 | 13 | 1.10 | 4 | 1.47 | 0.402 | 0.689 | (B) BRAF Class 2 |
| KIF1B | 1p36.22 | 28 | 4.80 | 5 | 3.76 | 0.403 | 0.689 | (A) BRAF Class 1 v12 |
| PTPRD | 9p24.1-p23 | 184 | 10.64 | 43 | 11.20 | 0.403 | 0.689 | (B) BRAF Class 2 |
| SESN3 | 11q21 | 8 | 1.52 | 1 | 0.71 | 0.404 | 0.689 | (A) BRAF Class 1 v12 |
| BABAM2 | 2p23.2 | 14 | 2.40 | 2 | 1.50 | 0.405 | 0.689 | (A) BRAF Class 1 v12 |
| MYBL1 | 8q13.1 | 30 | 2.70 | 5 | 2.12 | 0.405 | 0.689 | (A) BRAF Class 1 v12 |
| NBN | 8q21.3 | 77 | 3.06 | 19 | 3.34 | 0.406 | 0.689 | (B) BRAF Class 2 |
| FLT4 | 5q35.3 | 171 | 7.05 | 36 | 6.63 | 0.406 | 0.689 | (A) BRAF Class 1 v12 |
| SNCAIP | 5q23.2 | 44 | 6.69 | 9 | 7.69 | 0.406 | 0.689 | (B) BRAF Class 2 |
| TET1 | 10q21.3 | 118 | 6.41 | 29 | 6.84 | 0.408 | 0.69 | (B) BRAF Class 2 |
| STK19 | 6p21.33 | 12 | 2.28 | 2 | 1.43 | 0.409 | 0.69 | (A) BRAF Class 1 v12 |
| IL7R | 5p13.2 | 98 | 4.68 | 24 | 5.03 | 0.409 | 0.69 | (B) BRAF Class 2 |
| RAD51B | 14q24.1 | 13 | 0.93 | 4 | 1.22 | 0.412 | 0.694 | (B) BRAF Class 2 |
| WT1 | 11p13 | 68 | 2.74 | 17 | 2.99 | 0.413 | 0.694 | (B) BRAF Class 2 |
| CDK4 | 12q14.1 | 43 | 1.70 | 9 | 1.45 | 0.415 | 0.696 | (A) BRAF Class 1 v12 |
| MAP3K1 | 5q11.2 | 98 | 4.00 | 24 | 4.29 | 0.416 | 0.696 | (B) BRAF Class 2 |
| PTCH2 | 1p34.1 | 6 | 2.51 | 2 | 4.00 | 0.416 | 0.696 | (B) BRAF Class 2 |
| SMYD3 | 1q44 | 8 | 1.52 | 3 | 2.14 | 0.417 | 0.696 | (B) BRAF Class 2 |
| EGFR | 7p11.2 | 132 | 3.95 | 28 | 3.69 | 0.417 | 0.696 | (A) BRAF Class 1 v12 |
| NFKBIE | 6p21.1 | 14 | 1.95 | 4 | 2.52 | 0.418 | 0.697 | (B) BRAF Class 2 |
| NEIL1 | 15q24.2 | 9 | 1.54 | 1 | 0.75 | 0.419 | 0.697 | (A) BRAF Class 1 v12 |
| IFNGR1 | 6q23.3 | 18 | 1.53 | 3 | 1.10 | 0.421 | 0.699 | (A) BRAF Class 1 v12 |
| DNMT3B | 20q11.21 | 34 | 2.89 | 9 | 3.30 | 0.422 | 0.699 | (B) BRAF Class 2 |
| PLK2 | 5q11.2 | 27 | 2.29 | 5 | 1.83 | 0.423 | 0.699 | (A) BRAF Class 1 v12 |
| FUBP1 | 1p31.1 | 27 | 1.79 | 5 | 1.43 | 0.423 | 0.699 | (A) BRAF Class 1 v12 |

| | | | | | | | | |
|---|---|---|---|---|---|---|---|---|
| PAX7 | 1p36.13 | 15 | 2.65 | 3 | 3.53 | 0.423 | 0.699 | (B) BRAF Class 2 |
| IKZF1 | 7p12.2 | 122 | 4.99 | 26 | 4.66 | 0.423 | 0.699 | (A) BRAF Class 1 v12 |
| PML | 15q24.1 | 28 | 4.05 | 5 | 3.25 | 0.425 | 0.7 | (A) BRAF Class 1 v12 |
| ARID5B | 10q21.2 | 50 | 3.81 | 10 | 3.33 | 0.425 | 0.7 | (A) BRAF Class 1 v12 |
| FGFR4 | 5q35.2 | 72 | 2.81 | 19 | 3.03 | 0.426 | 0.7 | (B) BRAF Class 2 |
| RRAGC | 1p34.3 | 5 | 0.75 | 2 | 1.20 | 0.427 | 0.7 | (B) BRAF Class 2 |
| ZBTB2 | 6q25.1 | 3 | 0.61 | 1 | 1.35 | 0.429 | 0.7 | (B) BRAF Class 2 |
| SDHC | 1q23.3 | 18 | 0.76 | 3 | 0.55 | 0.429 | 0.7 | (A) BRAF Class 1 v12 |
| MRE11 | 11q21 | 56 | 2.54 | 14 | 2.78 | 0.43 | 0.7 | (B) BRAF Class 2 |
| EPHB6 | 7q34 | 2 | 1.00 | 1 | 2.44 | 0.43 | 0.7 | (B) BRAF Class 2 |
| HSD3B1 | 1p12 | 13 | 2.48 | 3 | 3.30 | 0.43 | 0.7 | (B) BRAF Class 2 |
| TAL1 | 1p33 | 10 | 1.46 | 3 | 1.97 | 0.431 | 0.701 | (B) BRAF Class 2 |
| CD79A | 19q13.2 | 22 | 1.47 | 6 | 1.73 | 0.434 | 0.702 | (B) BRAF Class 2 |
| NFKBIZ | 3q12.3 | 14 | 1.53 | 2 | 1.01 | 0.434 | 0.702 | (A) BRAF Class 1 v12 |
| MYC | 8q24.21 | 89 | 3.46 | 20 | 3.21 | 0.434 | 0.702 | (A) BRAF Class 1 v12 |
| NOTCH4 | 6p21.32 | 91 | 6.44 | 22 | 6.85 | 0.434 | 0.702 | (B) BRAF Class 2 |
| ERG | 21q22.2 | 62 | 2.92 | 13 | 2.63 | 0.435 | 0.702 | (A) BRAF Class 1 v12 |
| ETV4 | 17q21.31 | 20 | 1.37 | 3 | 1.00 | 0.436 | 0.702 | (A) BRAF Class 1 v12 |
| MED12 | Xq13.1 | 118 | 5.37 | 28 | 5.06 | 0.436 | 0.702 | (A) BRAF Class 1 v12 |
| LATS2 | 13q12.11 | 48 | 4.01 | 10 | 3.55 | 0.438 | 0.703 | (A) BRAF Class 1 v12 |
| PPP2R1A | 19q13.41 | 35 | 1.59 | 9 | 1.78 | 0.438 | 0.703 | (B) BRAF Class 2 |
| CDKN2C | 1p32.3 | 14 | 0.57 | 4 | 0.72 | 0.439 | 0.704 | (B) BRAF Class 2 |
| SOS1 | 2p22.1 | 37 | 3.03 | 10 | 3.37 | 0.441 | 0.704 | (B) BRAF Class 2 |
| TLX3 | 5q35.1 | 14 | 2.38 | 4 | 2.96 | 0.442 | 0.704 | (B) BRAF Class 2 |
| PIK3R3 | 1p34.1 | 13 | 1.10 | 2 | 0.73 | 0.442 | 0.704 | (A) BRAF Class 1 v12 |
| XIAP | Xq25 | 13 | 1.10 | 2 | 0.73 | 0.442 | 0.704 | (A) BRAF Class 1 v12 |
| ATRX | Xq21.1 | 167 | 6.59 | 39 | 6.84 | 0.443 | 0.704 | (B) BRAF Class 2 |
| ZNF217 | 20q13.2 | 55 | 3.44 | 10 | 3.06 | 0.443 | 0.704 | (A) BRAF Class 1 v12 |
| PTK2B | 8p21.2 | 23 | 3.55 | 6 | 4.14 | 0.444 | 0.704 | (B) BRAF Class 2 |
| CSF1R | 5q32 | 63 | 2.46 | 14 | 2.66 | 0.444 | 0.704 | (B) BRAF Class 2 |
| ALK | p23.2-p23. | 183 | 5.48 | 40 | 5.27 | 0.449 | 0.711 | (A) BRAF Class 1 v12 |
| RPS6KA4 | 11q13.1 | 35 | 2.97 | 7 | 2.56 | 0.451 | 0.712 | (A) BRAF Class 1 v12 |
| SESN2 | 1p35.3 | 5 | 0.95 | 2 | 1.43 | 0.452 | 0.712 | (B) BRAF Class 2 |
| STAT3 | 17q21.2 | 45 | 1.84 | 9 | 1.63 | 0.453 | 0.712 | (A) BRAF Class 1 v12 |
| DDB1 | 11q12.2 | 22 | 3.77 | 4 | 3.01 | 0.453 | 0.712 | (A) BRAF Class 1 v12 |
| ZBTB7A | 19p13.3 | 12 | 3.60 | 2 | 5.00 | 0.455 | 0.712 | (B) BRAF Class 2 |
| GPC3 | Xq26.2 | 20 | 1.78 | 5 | 2.08 | 0.456 | 0.712 | (B) BRAF Class 2 |
| CSMD3 | 8q23.3 | 3 | 2.56 | 1 | 5.26 | 0.456 | 0.712 | (B) BRAF Class 2 |
| ITGA10 | 1q21.1 | 3 | 2.56 | 1 | 5.26 | 0.456 | 0.712 | (B) BRAF Class 2 |
| EP400 | 12q24.33 | 5 | 4.17 | 0 | 0.00 | 0.457 | 0.712 | (A) BRAF Class 1 v12 |
| ZRSR2 | Xp22.2 | 41 | 1.83 | 10 | 2.00 | 0.458 | 0.712 | (B) BRAF Class 2 |
| BLM | 15q26.1 | 107 | 4.41 | 25 | 4.60 | 0.458 | 0.712 | (B) BRAF Class 2 |
| TSHZ2 | 20q13.2 | 3 | 3.49 | 1 | 7.14 | 0.458 | 0.712 | (B) BRAF Class 2 |
| DIS3L2 | 2q37.1 | 19 | 2.91 | 5 | 3.40 | 0.459 | 0.712 | (B) BRAF Class 2 |
| ACVR1 | 2q24.1 | 22 | 1.18 | 4 | 0.95 | 0.461 | 0.712 | (A) BRAF Class 1 v12 |
| PRKN | 6q26 | 79 | 3.30 | 19 | 3.47 | 0.461 | 0.712 | (B) BRAF Class 2 |
| NEGR1 | 1p31.1 | 28 | 2.08 | 7 | 2.33 | 0.462 | 0.712 | (B) BRAF Class 2 |
| ESR1 | q25.1-q25. | 54 | 2.05 | 12 | 1.88 | 0.463 | 0.712 | (A) BRAF Class 1 v12 |

| | | | | | | | | |
|---|---|---|---|---|---|---|---|---|
| FIP1L1 | 4q12 | 2 | 1.31 | 1 | 2.86 | 0.463 | 0.712 | (B) BRAF Class 2 |
| HIST1H3A | 6p22.2 | 2 | 0.18 | 1 | 0.39 | 0.463 | 0.712 | (B) BRAF Class 2 |
| XRCC5 | 2q35 | 10 | 1.66 | 3 | 2.11 | 0.463 | 0.712 | (B) BRAF Class 2 |
| ETV1 | 7p21.2 | 58 | 2.55 | 14 | 2.72 | 0.464 | 0.713 | (B) BRAF Class 2 |
| IRS2 | 13q34 | 78 | 4.99 | 19 | 5.23 | 0.466 | 0.714 | (B) BRAF Class 2 |
| CDH4 | 20q13.33 | 35 | 6.00 | 7 | 5.26 | 0.466 | 0.714 | (A) BRAF Class 1 v12 |
| TOP1 | 20q12 | 27 | 1.68 | 5 | 1.41 | 0.467 | 0.714 | (A) BRAF Class 1 v12 |
| KIF5B | 10p11.22 | 11 | 2.68 | 2 | 3.64 | 0.468 | 0.714 | (B) BRAF Class 2 |
| RHOA | 3p21.31 | 31 | 1.53 | 6 | 1.32 | 0.469 | 0.714 | (A) BRAF Class 1 v12 |
| RHBDF2 | 17q25.1 | 26 | 4.46 | 5 | 3.76 | 0.47 | 0.714 | (A) BRAF Class 1 v12 |
| CACNA1E | 1q25.3 | 3 | 6.25 | 1 | 12.50 | 0.47 | 0.714 | (B) BRAF Class 2 |
| CRTC1 | 19p13.11 | 37 | 3.65 | 7 | 3.21 | 0.47 | 0.714 | (A) BRAF Class 1 v12 |
| RAD51 | 15q15.1 | 19 | 0.87 | 5 | 1.00 | 0.472 | 0.714 | (B) BRAF Class 2 |
| MUTYH | 1p34.1 | 45 | 1.83 | 11 | 1.96 | 0.472 | 0.714 | (B) BRAF Class 2 |
| BBC3 | 19q13.32 | 8 | 0.68 | 1 | 0.37 | 0.472 | 0.714 | (A) BRAF Class 1 v12 |
| TSC2 | 16p13.3 | 134 | 5.28 | 29 | 5.08 | 0.472 | 0.714 | (A) BRAF Class 1 v12 |
| PTPN14 | 1q32.3-q41 | 17 | 2.92 | 3 | 2.26 | 0.474 | 0.715 | (A) BRAF Class 1 v12 |
| FH | 1q43 | 48 | 1.97 | 10 | 1.80 | 0.476 | 0.716 | (A) BRAF Class 1 v12 |
| MYH9 | 22q12.3 | 8 | 3.13 | 2 | 4.26 | 0.477 | 0.716 | (B) BRAF Class 2 |
| BCL10 | 1p22.3 | 17 | 1.41 | 3 | 1.09 | 0.478 | 0.716 | (A) BRAF Class 1 v12 |
| MSH2 | 2p21-p16.3 | 72 | 2.92 | 17 | 3.05 | 0.478 | 0.716 | (B) BRAF Class 2 |
| NRIP1 | 1q11.2-q21. | 3 | 3.26 | 1 | 6.25 | 0.479 | 0.716 | (B) BRAF Class 2 |
| ZFPM1 | 16q24.2 | 3 | 3.26 | 1 | 6.25 | 0.479 | 0.716 | (B) BRAF Class 2 |
| RANBP2 | 2q13 | 56 | 11.34 | 9 | 12.16 | 0.48 | 0.716 | (B) BRAF Class 2 |
| ZNF703 | 8p11.23 | 21 | 2.97 | 3 | 2.33 | 0.48 | 0.716 | (A) BRAF Class 1 v12 |
| TMPRSS2 | 21q22.3 | 40 | 1.91 | 10 | 2.04 | 0.481 | 0.716 | (B) BRAF Class 2 |
| HNRNPK | 9q21.32 | 4 | 0.93 | 1 | 1.67 | 0.482 | 0.716 | (B) BRAF Class 2 |
| POLH | 6p21.1 | 11 | 1.70 | 3 | 2.08 | 0.484 | 0.716 | (B) BRAF Class 2 |
| SRP19 | 5q22.2 | 3 | 0.82 | 0 | 0.00 | 0.484 | 0.716 | (A) BRAF Class 1 v12 |
| PREX2 | 8q13.2 | 139 | 12.00 | 30 | 12.30 | 0.485 | 0.716 | (B) BRAF Class 2 |
| PRKD1 | 14q12 | 11 | 1.87 | 2 | 1.32 | 0.486 | 0.716 | (A) BRAF Class 1 v12 |
| GNA13 | 17q24.1 | 3 | 0.42 | 1 | 0.77 | 0.486 | 0.716 | (B) BRAF Class 2 |
| CDK2 | 12q13.2 | 3 | 0.69 | 1 | 1.28 | 0.486 | 0.716 | (B) BRAF Class 2 |
| MAP3K2 | 2q14.3 | 3 | 1.92 | 1 | 3.57 | 0.486 | 0.716 | (B) BRAF Class 2 |
| SIN3A | 15q24.2 | 3 | 1.92 | 1 | 3.57 | 0.486 | 0.716 | (B) BRAF Class 2 |
| BMPR1A | 10q23.2 | 48 | 2.07 | 10 | 1.90 | 0.487 | 0.717 | (A) BRAF Class 1 v12 |
| NTHL1 | 16p13.3 | 10 | 0.86 | 3 | 1.05 | 0.49 | 0.72 | (B) BRAF Class 2 |
| TCF7L1 | 2p11.2 | 8 | 1.79 | 2 | 2.35 | 0.492 | 0.72 | (B) BRAF Class 2 |
| GSTM5 | 1p13.3 | 4 | 1.13 | 0 | 0.00 | 0.493 | 0.72 | (A) BRAF Class 1 v12 |
| NCOR2 | 12q24.31 | 3 | 2.61 | 1 | 4.76 | 0.493 | 0.72 | (B) BRAF Class 2 |
| STAG1 | 3q22.3 | 28 | 4.14 | 5 | 4.59 | 0.494 | 0.72 | (B) BRAF Class 2 |
| SBDS | 7q11.21 | 7 | 0.68 | 2 | 0.90 | 0.494 | 0.72 | (B) BRAF Class 2 |
| ERCC5 | 13q33.1 | 73 | 3.37 | 17 | 3.48 | 0.495 | 0.722 | (B) BRAF Class 2 |
| DNAH9 | 17p12 | 6 | 8.70 | 2 | 11.76 | 0.497 | 0.722 | (B) BRAF Class 2 |
| BCL2L1 | 20q11.21 | 11 | 0.46 | 3 | 0.55 | 0.498 | 0.722 | (B) BRAF Class 2 |
| XRCC6 | 22q13.2 | 11 | 1.89 | 3 | 2.26 | 0.498 | 0.722 | (B) BRAF Class 2 |
| EXT1 | 8q24.11 | 21 | 2.05 | 5 | 2.27 | 0.499 | 0.722 | (B) BRAF Class 2 |
| MYCN | 2p24.3 | 43 | 1.70 | 11 | 1.78 | 0.5 | 0.722 | (B) BRAF Class 2 |

| | | | | | | | | |
|---|---|---|---|---|---|---|---|---|
| ARAF | Xp11.3 | 65 | 2.51 | 14 | 2.39 | 0.501 | 0.722 | (A) BRAF Class 1 v12 |
| HIST1H1C | 6p22.2 | 25 | 2.00 | 5 | 1.75 | 0.501 | 0.722 | (A) BRAF Class 1 v12 |
| SH2D1A | Xq25 | 11 | 0.60 | 3 | 0.71 | 0.502 | 0.722 | (B) BRAF Class 2 |
| MUS81 | 11q13.1 | 12 | 2.06 | 2 | 1.50 | 0.502 | 0.722 | (A) BRAF Class 1 v12 |
| PHOX2B | 4p13 | 48 | 2.08 | 10 | 1.94 | 0.502 | 0.722 | (A) BRAF Class 1 v12 |
| RAB35 | 12q24.23 | 11 | 0.99 | 3 | 1.18 | 0.503 | 0.722 | (B) BRAF Class 2 |
| CBFB | 16q22.1 | 16 | 0.76 | 3 | 0.61 | 0.503 | 0.722 | (A) BRAF Class 1 v12 |
| MAP3K9 | 14q24.2 | 9 | 4.89 | 1 | 2.86 | 0.505 | 0.722 | (A) BRAF Class 1 v12 |
| TMEM127 | 2q11.2 | 11 | 0.61 | 3 | 0.72 | 0.506 | 0.722 | (B) BRAF Class 2 |
| CDK5 | 7q36.1 | 3 | 0.85 | 1 | 1.47 | 0.507 | 0.722 | (B) BRAF Class 2 |
| PLCG2 | 16q23.3 | 51 | 3.93 | 11 | 3.70 | 0.508 | 0.722 | (A) BRAF Class 1 v12 |
| FGF7 | 15q21.2 | 4 | 0.92 | 1 | 1.52 | 0.508 | 0.722 | (B) BRAF Class 2 |
| BRD3 | 9q34.2 | 30 | 4.34 | 6 | 3.90 | 0.508 | 0.722 | (A) BRAF Class 1 v12 |
| CD58 | 1p13.1 | 3 | 0.70 | 1 | 1.20 | 0.508 | 0.722 | (B) BRAF Class 2 |
| RAC1 | 7p22.1 | 50 | 2.17 | 12 | 2.25 | 0.508 | 0.722 | (B) BRAF Class 2 |
| LIG4 | 13q33.3 | 20 | 3.43 | 5 | 3.76 | 0.509 | 0.722 | (B) BRAF Class 2 |
| NCOA3 | 20q13.12 | 52 | 3.98 | 11 | 3.77 | 0.511 | 0.723 | (A) BRAF Class 1 v12 |
| DICER1 | 14q32.13 | 92 | 3.83 | 20 | 3.72 | 0.512 | 0.724 | (A) BRAF Class 1 v12 |
| JAK1 | 1p31.3 | 79 | 3.53 | 20 | 3.59 | 0.513 | 0.725 | (B) BRAF Class 2 |
| CLDN18 | 3q22.3 | 4 | 2.56 | 0 | 0.00 | 0.514 | 0.725 | (A) BRAF Class 1 v12 |
| ALOX12B | 17p13.1 | 37 | 2.25 | 8 | 2.09 | 0.514 | 0.725 | (A) BRAF Class 1 v12 |
| XRCC1 | 19q13.31 | 12 | 1.86 | 2 | 1.39 | 0.516 | 0.725 | (A) BRAF Class 1 v12 |
| MAPK3 | 16p11.2 | 20 | 1.80 | 5 | 1.95 | 0.516 | 0.725 | (B) BRAF Class 2 |
| DAXX | 6p21.32 | 41 | 1.99 | 9 | 1.87 | 0.518 | 0.725 | (A) BRAF Class 1 v12 |
| ABL1 | 9q34.12 | 88 | 2.80 | 19 | 2.84 | 0.518 | 0.725 | (B) BRAF Class 2 |
| CCND2 | 12p13.32 | 25 | 0.99 | 5 | 0.88 | 0.519 | 0.725 | (A) BRAF Class 1 v12 |
| IGF2R | 6q25.3 | 8 | 2.40 | 2 | 2.99 | 0.519 | 0.725 | (B) BRAF Class 2 |
| SAMD9 | 7q21.2 | 4 | 2.14 | 0 | 0.00 | 0.519 | 0.725 | (A) BRAF Class 1 v12 |
| FLCN | 17p11.2 | 60 | 2.46 | 14 | 2.51 | 0.52 | 0.725 | (B) BRAF Class 2 |
| POLR2A | 17p13.1 | 4 | 4.35 | 0 | 0.00 | 0.521 | 0.726 | (A) BRAF Class 1 v12 |
| GNAS | 20q13.32 | 106 | 3.23 | 22 | 3.17 | 0.522 | 0.726 | (A) BRAF Class 1 v12 |
| FOXL2 | 3q22.3 | 46 | 2.09 | 10 | 1.98 | 0.522 | 0.726 | (A) BRAF Class 1 v12 |
| FGF10 | 5p12 | 9 | 1.27 | 2 | 1.55 | 0.525 | 0.728 | (B) BRAF Class 2 |
| EIF1AX | Xp22.12 | 20 | 1.46 | 4 | 1.27 | 0.526 | 0.729 | (A) BRAF Class 1 v12 |
| CD274 | 9p24.1 | 20 | 0.83 | 4 | 0.72 | 0.527 | 0.729 | (A) BRAF Class 1 v12 |
| CCNQ | Xq28 | 6 | 1.14 | 2 | 1.43 | 0.527 | 0.729 | (B) BRAF Class 2 |
| EWSR1 | 22q12.2 | 28 | 1.76 | 6 | 1.85 | 0.528 | 0.729 | (B) BRAF Class 2 |
| RET | 10q11.21 | 123 | 3.68 | 28 | 3.69 | 0.529 | 0.729 | (B) BRAF Class 2 |
| PGR | 11q22.1 | 67 | 6.05 | 15 | 5.88 | 0.529 | 0.729 | (A) BRAF Class 1 v12 |
| GEN1 | 2p24.2 | 24 | 2.31 | 5 | 2.45 | 0.53 | 0.729 | (B) BRAF Class 2 |
| EP300 | 22q13.2 | 172 | 7.00 | 39 | 6.96 | 0.53 | 0.729 | (A) BRAF Class 1 v12 |
| STAT5A | 17q21.2 | 20 | 1.70 | 4 | 1.49 | 0.531 | 0.729 | (A) BRAF Class 1 v12 |
| PDCD1LG2 | 9p24.1 | 25 | 1.21 | 5 | 1.10 | 0.532 | 0.729 | (A) BRAF Class 1 v12 |
| NUP93 | 16q13 | 30 | 2.06 | 7 | 2.14 | 0.533 | 0.73 | (B) BRAF Class 2 |
| XPC | 3p25.1 | 22 | 2.15 | 5 | 2.27 | 0.535 | 0.731 | (B) BRAF Class 2 |
| TENT5C | 1p12 | 37 | 1.51 | 8 | 1.43 | 0.535 | 0.731 | (A) BRAF Class 1 v12 |
| SMO | 7q32.1 | 98 | 3.08 | 22 | 3.05 | 0.538 | 0.734 | (A) BRAF Class 1 v12 |
| HIST1H3D | 6p22.2 | 7 | 0.63 | 1 | 0.39 | 0.541 | 0.735 | (A) BRAF Class 1 v12 |

| | | | | | | | | |
|---|---|---|---|---|---|---|---|---|
| GRM8 | 7q31.33 | 4 | 3.42 | 0 | 0.00 | 0.544 | 0.735 | (A) BRAF Class 1 v12 |
| RECQL | 12p12.1 | 14 | 2.66 | 4 | 2.86 | 0.544 | 0.735 | (B) BRAF Class 2 |
| YAP1 | 11q22.1 | 28 | 1.50 | 6 | 1.40 | 0.544 | 0.735 | (A) BRAF Class 1 v12 |
| RPA1 | 17p13.3 | 11 | 1.61 | 2 | 1.26 | 0.544 | 0.735 | (A) BRAF Class 1 v12 |
| NAB2 | 12q13.3 | 13 | 3.90 | 1 | 2.50 | 0.545 | 0.735 | (A) BRAF Class 1 v12 |
| RHOH | 4p14 | 3 | 0.44 | 0 | 0.00 | 0.547 | 0.735 | (A) BRAF Class 1 v12 |
| KNSTRN | 15q15.1 | 20 | 2.71 | 5 | 2.82 | 0.547 | 0.735 | (B) BRAF Class 2 |
| RBBP8 | 18q11.2 | 30 | 5.15 | 7 | 5.26 | 0.548 | 0.735 | (B) BRAF Class 2 |
| GATA1 | Xp11.23 | 20 | 1.27 | 4 | 1.14 | 0.548 | 0.735 | (A) BRAF Class 1 v12 |
| MSI2 | 17q22 | 6 | 1.14 | 1 | 0.71 | 0.548 | 0.735 | (A) BRAF Class 1 v12 |
| SESN1 | 6q21 | 6 | 1.14 | 1 | 0.71 | 0.548 | 0.735 | (A) BRAF Class 1 v12 |
| PMS2 | 7p22.1 | 53 | 2.15 | 12 | 2.15 | 0.55 | 0.735 | (B) BRAF Class 2 |
| SLC34A2 | 4p15.2 | 29 | 4.26 | 7 | 4.35 | 0.551 | 0.735 | (B) BRAF Class 2 |
| RBM10 | Xp11.3 | 82 | 3.97 | 19 | 3.95 | 0.551 | 0.735 | (A) BRAF Class 1 v12 |
| GABRA6 | 5q34 | 16 | 3.05 | 3 | 3.30 | 0.551 | 0.735 | (B) BRAF Class 2 |
| DOCK8 | 9p24.3 | 48 | 8.23 | 11 | 8.27 | 0.552 | 0.735 | (B) BRAF Class 2 |
| RAD51D | 17q12 | 21 | 0.97 | 5 | 1.00 | 0.553 | 0.735 | (B) BRAF Class 2 |
| PPM1D | 17q23.2 | 45 | 2.49 | 10 | 2.43 | 0.553 | 0.735 | (A) BRAF Class 1 v12 |
| SUFU | 10q24.32 | 45 | 1.83 | 10 | 1.79 | 0.554 | 0.735 | (A) BRAF Class 1 v12 |
| NUTM1 | 15q14 | 29 | 5.06 | 5 | 5.21 | 0.554 | 0.735 | (B) BRAF Class 2 |
| NR0B1 | Xp21.2 | 11 | 1.89 | 2 | 1.50 | 0.554 | 0.735 | (A) BRAF Class 1 v12 |
| MAFB | 20q12 | 3 | 0.44 | 1 | 0.65 | 0.555 | 0.735 | (B) BRAF Class 2 |
| ARHGAP35 | 19q13.32 | 31 | 4.26 | 7 | 4.32 | 0.556 | 0.735 | (B) BRAF Class 2 |
| FYN | 6q21 | 26 | 2.07 | 6 | 2.11 | 0.556 | 0.735 | (B) BRAF Class 2 |
| CCN6 | 6q21 | 17 | 2.51 | 3 | 2.68 | 0.556 | 0.735 | (B) BRAF Class 2 |
| STK40 | 1p34.3 | 12 | 1.02 | 3 | 1.10 | 0.558 | 0.735 | (B) BRAF Class 2 |
| RPS6KB1 | 17q23.1 | 5 | 1.50 | 0 | 0.00 | 0.558 | 0.735 | (A) BRAF Class 1 v12 |
| SAMD9L | 7q21.2 | 4 | 4.35 | 1 | 6.25 | 0.559 | 0.735 | (B) BRAF Class 2 |
| YES1 | 18p11.32 | 21 | 1.78 | 5 | 1.83 | 0.559 | 0.735 | (B) BRAF Class 2 |
| MYB | 6q23.3 | 19 | 1.21 | 4 | 1.25 | 0.559 | 0.735 | (B) BRAF Class 2 |
| AGO2 | 8q24.3 | 26 | 4.94 | 7 | 5.00 | 0.561 | 0.735 | (B) BRAF Class 2 |
| FAAP20 | 1p36.33 | 3 | 0.51 | 1 | 0.75 | 0.561 | 0.735 | (B) BRAF Class 2 |
| MDC1 | 6p21.33 | 65 | 5.52 | 15 | 5.49 | 0.562 | 0.735 | (A) BRAF Class 1 v12 |
| NCOR1 | 17p12-p11.2 | 79 | 5.34 | 18 | 5.34 | 0.563 | 0.735 | (A) BRAF Class 1 v12 |
| ETV5 | 3q27.2 | 24 | 1.64 | 5 | 1.67 | 0.564 | 0.735 | (B) BRAF Class 2 |
| CRLF2 | 2.33 and Yp1 | 31 | 1.32 | 7 | 1.32 | 0.565 | 0.735 | (B) BRAF Class 2 |
| KDM6B | 17p13.1 | 20 | 5.52 | 4 | 5.71 | 0.565 | 0.735 | (B) BRAF Class 2 |
| CSNK1A1 | 5q32 | 5 | 1.50 | 0 | 0.00 | 0.565 | 0.735 | (A) BRAF Class 1 v12 |
| FGF9 | 13q12.11 | 5 | 1.50 | 0 | 0.00 | 0.565 | 0.735 | (A) BRAF Class 1 v12 |
| ZBTB20 | 3q13.31 | 4 | 2.56 | 1 | 3.57 | 0.566 | 0.735 | (B) BRAF Class 2 |
| EZH1 | 17q21.2 | 7 | 1.06 | 2 | 1.20 | 0.568 | 0.737 | (B) BRAF Class 2 |
| IKZF2 | 2q34 | 4 | 1.82 | 1 | 2.50 | 0.569 | 0.738 | (B) BRAF Class 2 |
| SQSTM1 | 5q35.3 | 4 | 1.52 | 1 | 2.08 | 0.57 | 0.738 | (B) BRAF Class 2 |
| TSHR | 14q31.1 | 57 | 2.64 | 13 | 2.64 | 0.571 | 0.738 | (A) BRAF Class 1 v12 |
| CHD5 | 1p36.31 | 10 | 6.41 | 2 | 7.14 | 0.572 | 0.739 | (B) BRAF Class 2 |
| TAF1 | Xq13.1 | 28 | 5.31 | 4 | 4.94 | 0.573 | 0.739 | (A) BRAF Class 1 v12 |
| FANCM | 14q21.2 | 40 | 5.35 | 9 | 5.29 | 0.578 | 0.744 | (A) BRAF Class 1 v12 |
| ID3 | 1p36.12 | 8 | 0.35 | 2 | 0.39 | 0.579 | 0.745 | (B) BRAF Class 2 |

| | | | | | | | | |
|---|---|---|---|---|---|---|---|---|
| TYRO3 | 15q15.1 | 2 | 1.48 | 0 | 0.00 | 0.587 | 0.755 | (A) BRAF Class 1 v12 |
| CDK9 | 9q34.11 | 3 | 0.85 | 0 | 0.00 | 0.589 | 0.755 | (A) BRAF Class 1 v12 |
| PSMD13 | 11p15.5 | 3 | 0.85 | 0 | 0.00 | 0.589 | 0.755 | (A) BRAF Class 1 v12 |
| RPL22 | 1p36.31 | 3 | 5.66 | 0 | 0.00 | 0.59 | 0.755 | (A) BRAF Class 1 v12 |
| UPF1 | 19p13.11 | 15 | 2.85 | 4 | 2.86 | 0.591 | 0.756 | (B) BRAF Class 2 |
| CDC42 | 1p36.12 | 3 | 0.45 | 1 | 0.60 | 0.593 | 0.757 | (B) BRAF Class 2 |
| RRAS | 19q13.33 | 3 | 0.45 | 1 | 0.60 | 0.593 | 0.757 | (B) BRAF Class 2 |
| DDX3X | Xp11.4 | 3 | 1.20 | 0 | 0.00 | 0.594 | 0.757 | (A) BRAF Class 1 v12 |
| LTK | 15q15.1 | 3 | 1.30 | 1 | 1.72 | 0.595 | 0.758 | (B) BRAF Class 2 |
| GAB2 | 11q14.1 | 3 | 1.81 | 0 | 0.00 | 0.597 | 0.759 | (A) BRAF Class 1 v12 |
| GPS2 | 17p13.1 | 13 | 1.15 | 3 | 1.15 | 0.6 | 0.762 | (B) BRAF Class 2 |
| CALR | 19p13.13 | 26 | 1.34 | 6 | 1.33 | 0.601 | 0.762 | (A) BRAF Class 1 v12 |
| HDAC4 | 2q37.3 | 3 | 1.35 | 0 | 0.00 | 0.601 | 0.762 | (A) BRAF Class 1 v12 |
| NPRL2 | 3p21.31 | 3 | 1.14 | 0 | 0.00 | 0.604 | 0.762 | (A) BRAF Class 1 v12 |
| HIST1H3E | 6p22.2 | 10 | 0.90 | 2 | 0.78 | 0.604 | 0.762 | (A) BRAF Class 1 v12 |
| HIST1H3J | 6p22.1 | 10 | 0.90 | 2 | 0.78 | 0.604 | 0.762 | (A) BRAF Class 1 v12 |
| EPCAM | 2p21 | 18 | 0.91 | 4 | 0.88 | 0.605 | 0.762 | (A) BRAF Class 1 v12 |
| LYN | 8q12.1 | 27 | 2.63 | 6 | 2.62 | 0.605 | 0.762 | (A) BRAF Class 1 v12 |
| IGF1 | 12q23.2 | 6 | 0.47 | 1 | 0.33 | 0.605 | 0.762 | (A) BRAF Class 1 v12 |
| EED | 11q14.2 | 14 | 0.88 | 3 | 0.83 | 0.608 | 0.764 | (A) BRAF Class 1 v12 |
| DCUN1D1 | 3q26.33 | 7 | 0.59 | 2 | 0.62 | 0.608 | 0.764 | (B) BRAF Class 2 |
| EPAS1 | 2p21 | 12 | 2.28 | 3 | 2.14 | 0.61 | 0.764 | (A) BRAF Class 1 v12 |
| TAP1 | 6p21.32 | 12 | 2.28 | 3 | 2.14 | 0.61 | 0.764 | (A) BRAF Class 1 v12 |
| TRIM37 | 17q22 | 18 | 3.09 | 4 | 3.01 | 0.611 | 0.765 | (A) BRAF Class 1 v12 |
| HIST2H3D | 1q21.2 | 6 | 0.55 | 1 | 0.40 | 0.614 | 0.767 | (A) BRAF Class 1 v12 |
| THADA | 2p21 | 3 | 1.94 | 0 | 0.00 | 0.616 | 0.767 | (A) BRAF Class 1 v12 |
| RHOT1 | 17q11.2 | 6 | 1.03 | 1 | 0.75 | 0.616 | 0.767 | (A) BRAF Class 1 v12 |
| SDHAF2 | 11q12.2 | 10 | 0.45 | 2 | 0.40 | 0.616 | 0.767 | (A) BRAF Class 1 v12 |
| GLI3 | 7p14.1 | 26 | 7.37 | 5 | 7.35 | 0.616 | 0.767 | (A) BRAF Class 1 v12 |
| DPYD | 1p21.3 | 4 | 1.62 | 1 | 1.92 | 0.618 | 0.768 | (B) BRAF Class 2 |
| KMT5A | 12q24.31 | 3 | 0.55 | 1 | 0.67 | 0.62 | 0.77 | (B) BRAF Class 2 |
| DNAJB1 | 19p13.12 | 14 | 1.07 | 3 | 1.03 | 0.621 | 0.77 | (A) BRAF Class 1 v12 |
| CD22 | 19q13.12 | 2 | 1.72 | 0 | 0.00 | 0.622 | 0.77 | (A) BRAF Class 1 v12 |
| WAS | Xp11.23 | 10 | 1.45 | 2 | 1.30 | 0.622 | 0.77 | (A) BRAF Class 1 v12 |
| CDH23 | 10q22.1 | 3 | 6.25 | 0 | 0.00 | 0.624 | 0.77 | (A) BRAF Class 1 v12 |
| TGFB1 | 19q13.2 | 3 | 6.25 | 0 | 0.00 | 0.624 | 0.77 | (A) BRAF Class 1 v12 |
| UMODL1 | 21q22.3 | 3 | 6.25 | 0 | 0.00 | 0.624 | 0.77 | (A) BRAF Class 1 v12 |
| SLITRK6 | 13q31.1 | 12 | 3.67 | 2 | 3.33 | 0.627 | 0.773 | (A) BRAF Class 1 v12 |
| NUP214 | 9q34.13 | 5 | 3.82 | 1 | 4.35 | 0.628 | 0.773 | (B) BRAF Class 2 |
| LMO2 | 11p13 | 6 | 0.65 | 1 | 0.50 | 0.634 | 0.775 | (A) BRAF Class 1 v12 |
| FGF1 | 5q31.3 | 4 | 1.20 | 0 | 0.00 | 0.634 | 0.775 | (A) BRAF Class 1 v12 |
| ADAMTS20 | 12q12 | 3 | 2.56 | 0 | 0.00 | 0.634 | 0.775 | (A) BRAF Class 1 v12 |
| FN1 | 2q35 | 3 | 2.56 | 0 | 0.00 | 0.634 | 0.775 | (A) BRAF Class 1 v12 |
| FOXP4 | 6p21.1 | 3 | 2.56 | 0 | 0.00 | 0.634 | 0.775 | (A) BRAF Class 1 v12 |
| RPS6KA2 | 6q27 | 3 | 2.56 | 0 | 0.00 | 0.634 | 0.775 | (A) BRAF Class 1 v12 |
| TNK2 | 3q29 | 3 | 2.56 | 0 | 0.00 | 0.634 | 0.775 | (A) BRAF Class 1 v12 |
| NEIL2 | 8p23.1 | 4 | 0.69 | 1 | 0.75 | 0.643 | 0.784 | (B) BRAF Class 2 |
| OGG1 | 3p25.3 | 4 | 0.69 | 1 | 0.75 | 0.643 | 0.784 | (B) BRAF Class 2 |

| | | | | | | | | |
|---|---|---|---|---|---|---|---|---|
| TAL2 | 9q31.2 | 4 | 0.68 | 1 | 0.74 | 0.645 | 0.786 | (B) BRAF Class 2 |
| MSI1 | 12q24.31 | 8 | 1.52 | 2 | 1.43 | 0.647 | 0.787 | (A) BRAF Class 1 v12 |
| BRINP3 | 1q31.1 | 5 | 4.24 | 1 | 4.55 | 0.649 | 0.788 | (B) BRAF Class 2 |
| HSP90AA1 | 14q32.31 | 8 | 1.50 | 1 | 1.20 | 0.652 | 0.791 | (A) BRAF Class 1 v12 |
| ADGRB3 | 6q12-q13 | 3 | 5.56 | 0 | 0.00 | 0.656 | 0.794 | (A) BRAF Class 1 v12 |
| HIST1H2BD | 6p22.2 | 9 | 0.76 | 2 | 0.73 | 0.656 | 0.794 | (A) BRAF Class 1 v12 |
| IL10 | 1q32.1 | 9 | 0.76 | 2 | 0.73 | 0.656 | 0.794 | (A) BRAF Class 1 v12 |
| MAF | 16q23.2 | 4 | 0.56 | 1 | 0.59 | 0.657 | 0.794 | (B) BRAF Class 2 |
| EPHB4 | 7q22.1 | 4 | 1.33 | 1 | 1.39 | 0.661 | 0.796 | (B) BRAF Class 2 |
| FAAP24 | 19q13.11 | 2 | 0.34 | 0 | 0.00 | 0.663 | 0.796 | (A) BRAF Class 1 v12 |
| TAZ | Xq28 | 2 | 0.34 | 0 | 0.00 | 0.663 | 0.796 | (A) BRAF Class 1 v12 |
| CAMK1 | 3p25.3 | 2 | 0.06 | 0 | 0.00 | 0.664 | 0.796 | (A) BRAF Class 1 v12 |
| EPB41L4A | ;q22.1-q22.: | 2 | 0.06 | 0 | 0.00 | 0.664 | 0.796 | (A) BRAF Class 1 v12 |
| H19 | 11p15.5 | 2 | 0.06 | 0 | 0.00 | 0.664 | 0.796 | (A) BRAF Class 1 v12 |
| TRBC2 | 7q34 | 2 | 0.06 | 0 | 0.00 | 0.664 | 0.796 | (A) BRAF Class 1 v12 |
| PAX8 | 2q14.1 | 8 | 1.39 | 1 | 1.15 | 0.665 | 0.797 | (A) BRAF Class 1 v12 |
| MLLT3 | 9p21.3 | 9 | 2.47 | 1 | 2.08 | 0.672 | 0.804 | (A) BRAF Class 1 v12 |
| HIST2H3C | 1q21.2 | 5 | 0.45 | 1 | 0.39 | 0.687 | 0.821 | (A) BRAF Class 1 v12 |
| CDH20 | 18q21.33 | 2 | 1.00 | 0 | 0.00 | 0.688 | 0.821 | (A) BRAF Class 1 v12 |
| ID4 | 6p22.3 | 5 | 0.86 | 1 | 0.75 | 0.69 | 0.823 | (A) BRAF Class 1 v12 |
| CDK1 | 10q21.2 | 2 | 0.57 | 0 | 0.00 | 0.703 | 0.836 | (A) BRAF Class 1 v12 |
| RPL26 | 17p13.1 | 2 | 0.57 | 0 | 0.00 | 0.703 | 0.836 | (A) BRAF Class 1 v12 |
| DUSP2 | 2q11.2 | 2 | 0.91 | 0 | 0.00 | 0.715 | 0.85 | (A) BRAF Class 1 v12 |
| ASPSCR1 | 17q25.3 | 2 | 1.72 | 0 | 0.00 | 0.716 | 0.85 | (A) BRAF Class 1 v12 |
| BCL9 | 1q21.2 | 2 | 1.64 | 0 | 0.00 | 0.717 | 0.85 | (A) BRAF Class 1 v12 |
| UBR5 | 8q22.3 | 2 | 1.49 | 0 | 0.00 | 0.718 | 0.851 | (A) BRAF Class 1 v12 |
| ABCB4 | 7q21.12 | 2 | 2.17 | 0 | 0.00 | 0.724 | 0.855 | (A) BRAF Class 1 v12 |
| OGA | 10q24.32 | 2 | 2.17 | 0 | 0.00 | 0.724 | 0.855 | (A) BRAF Class 1 v12 |
| SMARCAL1 | 2q35 | 2 | 2.17 | 0 | 0.00 | 0.724 | 0.855 | (A) BRAF Class 1 v12 |
| AFF1 | ·q21.3-q22.: | 2 | 1.64 | 0 | 0.00 | 0.727 | 0.855 | (A) BRAF Class 1 v12 |
| CDH11 | 16q21 | 2 | 1.64 | 0 | 0.00 | 0.727 | 0.855 | (A) BRAF Class 1 v12 |
| KNL1 | 15q15.1 | 2 | 1.64 | 0 | 0.00 | 0.727 | 0.855 | (A) BRAF Class 1 v12 |
| TRIM33 | 1p13.2 | 2 | 1.64 | 0 | 0.00 | 0.727 | 0.855 | (A) BRAF Class 1 v12 |
| USP7 | 16p13.2 | 2 | 2.33 | 0 | 0.00 | 0.738 | 0.865 | (A) BRAF Class 1 v12 |
| GUCY1A2 | 11q22.3 | 2 | 1.71 | 0 | 0.00 | 0.739 | 0.865 | (A) BRAF Class 1 v12 |
| NLRP1 | 17p13.2 | 2 | 1.71 | 0 | 0.00 | 0.739 | 0.865 | (A) BRAF Class 1 v12 |
| STK36 | 2q35 | 2 | 1.71 | 0 | 0.00 | 0.739 | 0.865 | (A) BRAF Class 1 v12 |
| TRIM24 | 7q33-q34 | 2 | 1.71 | 0 | 0.00 | 0.739 | 0.865 | (A) BRAF Class 1 v12 |
| CDKN2B-AS1 | 9p21.3 | 1 | 0.26 | 0 | 0.00 | 0.781 | 0.869 | (A) BRAF Class 1 v12 |
| KIF3A | 5q31.1 | 1 | 2.33 | 0 | 0.00 | 0.796 | 0.869 | (A) BRAF Class 1 v12 |
| SGK1 | 6q23.2 | 1 | 0.66 | 0 | 0.00 | 0.799 | 0.869 | (A) BRAF Class 1 v12 |
| HLA-C | 6p21.33 | 1 | 0.87 | 0 | 0.00 | 0.81 | 0.869 | (A) BRAF Class 1 v12 |
| TPMT | 6p22.3 | 1 | 0.66 | 0 | 0.00 | 0.813 | 0.869 | (A) BRAF Class 1 v12 |
| PHLPP2 | 16q22.2 | 1 | 0.76 | 0 | 0.00 | 0.814 | 0.869 | (A) BRAF Class 1 v12 |
| SLX1B | 16p11.2 | 1 | 0.17 | 0 | 0.00 | 0.814 | 0.869 | (A) BRAF Class 1 v12 |
| MAP3K14 | 17q21.31 | 1 | 0.09 | 0 | 0.00 | 0.814 | 0.869 | (A) BRAF Class 1 v12 |
| ABCA17P | 16p13.3 | 1 | 0.03 | 0 | 0.00 | 0.815 | 0.869 | (A) BRAF Class 1 v12 |
| ABR | 17p13.3 | 1 | 0.03 | 0 | 0.00 | 0.815 | 0.869 | (A) BRAF Class 1 v12 |

| | | | | | | | | |
|---|---|---|---|---|---|---|---|---|
| ANKLE2 | 12q24.33 | 1 | 0.03 | 0 | 0.00 | 0.815 | 0.869 | (A) BRAF Class 1 v12 |
| ANKRD30A | 10p11.21 | 1 | 0.03 | 0 | 0.00 | 0.815 | 0.869 | (A) BRAF Class 1 v12 |
| AP1G1 | 16q22.2 | 1 | 0.03 | 0 | 0.00 | 0.815 | 0.869 | (A) BRAF Class 1 v12 |
| APBA1 | 9q21.12 | 1 | 0.03 | 0 | 0.00 | 0.815 | 0.869 | (A) BRAF Class 1 v12 |
| ARHGAP4 | Xq28 | 1 | 0.03 | 0 | 0.00 | 0.815 | 0.869 | (A) BRAF Class 1 v12 |
| ARSB | 5q14.1 | 1 | 0.03 | 0 | 0.00 | 0.815 | 0.869 | (A) BRAF Class 1 v12 |
| CASZ1 | 1p36.22 | 1 | 0.03 | 0 | 0.00 | 0.815 | 0.869 | (A) BRAF Class 1 v12 |
| CLIP1 | 12q24.31 | 1 | 0.03 | 0 | 0.00 | 0.815 | 0.869 | (A) BRAF Class 1 v12 |
| CPSF1 | 8q24.3 | 1 | 0.03 | 0 | 0.00 | 0.815 | 0.869 | (A) BRAF Class 1 v12 |
| CTDSP1 | 2q35 | 1 | 0.03 | 0 | 0.00 | 0.815 | 0.869 | (A) BRAF Class 1 v12 |
| CTRC | 1p36.21 | 1 | 0.03 | 0 | 0.00 | 0.815 | 0.869 | (A) BRAF Class 1 v12 |
| CTSB | 8p23.1 | 1 | 0.03 | 0 | 0.00 | 0.815 | 0.869 | (A) BRAF Class 1 v12 |
| CUL1 | 7q36.1 | 1 | 0.03 | 0 | 0.00 | 0.815 | 0.869 | (A) BRAF Class 1 v12 |
| DCBLD1 | 6q22.1 | 1 | 0.03 | 0 | 0.00 | 0.815 | 0.869 | (A) BRAF Class 1 v12 |
| DDAH1 | 1p22.3 | 1 | 0.03 | 0 | 0.00 | 0.815 | 0.869 | (A) BRAF Class 1 v12 |
| DEF8 | 16q24.3 | 1 | 0.03 | 0 | 0.00 | 0.815 | 0.869 | (A) BRAF Class 1 v12 |
| DENND4C | 9p22.1 | 1 | 0.03 | 0 | 0.00 | 0.815 | 0.869 | (A) BRAF Class 1 v12 |
| DHRSX | 2.33 and Yp1 | 1 | 0.03 | 0 | 0.00 | 0.815 | 0.869 | (A) BRAF Class 1 v12 |
| DRICH1 | 22q11.23 | 1 | 0.03 | 0 | 0.00 | 0.815 | 0.869 | (A) BRAF Class 1 v12 |
| EIF3E | 8q23.1 | 1 | 0.03 | 0 | 0.00 | 0.815 | 0.869 | (A) BRAF Class 1 v12 |
| ELAC2 | 17p12 | 1 | 0.03 | 0 | 0.00 | 0.815 | 0.869 | (A) BRAF Class 1 v12 |
| EMID1 | 22q12.2 | 1 | 0.03 | 0 | 0.00 | 0.815 | 0.869 | (A) BRAF Class 1 v12 |
| F11-AS1 | 4q35.2 | 1 | 0.03 | 0 | 0.00 | 0.815 | 0.869 | (A) BRAF Class 1 v12 |
| FKBP9P1 | 7p11.2 | 1 | 0.03 | 0 | 0.00 | 0.815 | 0.869 | (A) BRAF Class 1 v12 |
| MC1-LUC7L | 7q34 | 1 | 0.03 | 0 | 0.00 | 0.815 | 0.869 | (A) BRAF Class 1 v12 |
| GALNT11 | q36.1\|7q36. | 1 | 0.03 | 0 | 0.00 | 0.815 | 0.869 | (A) BRAF Class 1 v12 |
| GOLT1A | 1q32.1 | 1 | 0.03 | 0 | 0.00 | 0.815 | 0.869 | (A) BRAF Class 1 v12 |
| GRAPL | 17p11.2 | 1 | 0.03 | 0 | 0.00 | 0.815 | 0.869 | (A) BRAF Class 1 v12 |
| GRIA1 | 5q33.2 | 1 | 0.03 | 0 | 0.00 | 0.815 | 0.869 | (A) BRAF Class 1 v12 |
| GRIK1 | 21q21.3 | 1 | 0.03 | 0 | 0.00 | 0.815 | 0.869 | (A) BRAF Class 1 v12 |
| HECTD4 | 12q24.13 | 1 | 0.03 | 0 | 0.00 | 0.815 | 0.869 | (A) BRAF Class 1 v12 |
| HLA-DOB | 6p21.32 | 1 | 0.03 | 0 | 0.00 | 0.815 | 0.869 | (A) BRAF Class 1 v12 |
| HOOK2 | 19p13.13 | 1 | 0.03 | 0 | 0.00 | 0.815 | 0.869 | (A) BRAF Class 1 v12 |
| HSD17B6 | 12q13.3 | 1 | 0.03 | 0 | 0.00 | 0.815 | 0.869 | (A) BRAF Class 1 v12 |
| IGHE | 14q32.33 | 1 | 0.03 | 0 | 0.00 | 0.815 | 0.869 | (A) BRAF Class 1 v12 |
| LDLR | 19p13.2 | 1 | 0.03 | 0 | 0.00 | 0.815 | 0.869 | (A) BRAF Class 1 v12 |
| OC10537446 | 2p22.2 | 1 | 0.03 | 0 | 0.00 | 0.815 | 0.869 | (A) BRAF Class 1 v12 |
| LTBP4 | 19q13.2 | 1 | 0.03 | 0 | 0.00 | 0.815 | 0.869 | (A) BRAF Class 1 v12 |
| MT-ND6 | - | 1 | 0.03 | 0 | 0.00 | 0.815 | 0.869 | (A) BRAF Class 1 v12 |
| MYEOV | 11q13.3 | 1 | 0.03 | 0 | 0.00 | 0.815 | 0.869 | (A) BRAF Class 1 v12 |
| MYO1D | 17q11.2 | 1 | 0.03 | 0 | 0.00 | 0.815 | 0.869 | (A) BRAF Class 1 v12 |
| MYSM1 | 1p32.1 | 1 | 0.03 | 0 | 0.00 | 0.815 | 0.869 | (A) BRAF Class 1 v12 |
| NBPF20 | 1q21.1 | 1 | 0.03 | 0 | 0.00 | 0.815 | 0.869 | (A) BRAF Class 1 v12 |
| NDUFB2 | 7q34 | 1 | 0.03 | 0 | 0.00 | 0.815 | 0.869 | (A) BRAF Class 1 v12 |
| NELFA | 4p16.3 | 1 | 0.03 | 0 | 0.00 | 0.815 | 0.869 | (A) BRAF Class 1 v12 |
| PADI2 | 1p36.13 | 1 | 0.03 | 0 | 0.00 | 0.815 | 0.869 | (A) BRAF Class 1 v12 |
| PAX9 | 14q13.3 | 1 | 0.03 | 0 | 0.00 | 0.815 | 0.869 | (A) BRAF Class 1 v12 |
| PFKFB4 | 3p21.31 | 1 | 0.03 | 0 | 0.00 | 0.815 | 0.869 | (A) BRAF Class 1 v12 |

| | | | | | | | | |
|---|---|---|---|---|---|---|---|---|
| PKD1 | 16p13.3 | 1 | 0.03 | 0 | 0.00 | 0.815 | 0.869 | (A) BRAF Class 1 v12 |
| PLEKHA7 | 1p15.2-p15. | 1 | 0.03 | 0 | 0.00 | 0.815 | 0.869 | (A) BRAF Class 1 v12 |
| PLVAP | 19p13.11 | 1 | 0.03 | 0 | 0.00 | 0.815 | 0.869 | (A) BRAF Class 1 v12 |
| PNPLA6 | 19p13.2 | 1 | 0.03 | 0 | 0.00 | 0.815 | 0.869 | (A) BRAF Class 1 v12 |
| PRMT6 | 1p13.3 | 1 | 0.03 | 0 | 0.00 | 0.815 | 0.869 | (A) BRAF Class 1 v12 |
| PTBP3 | 9q32 | 1 | 0.03 | 0 | 0.00 | 0.815 | 0.869 | (A) BRAF Class 1 v12 |
| RAB2A | q12.1-q12. | 1 | 0.03 | 0 | 0.00 | 0.815 | 0.869 | (A) BRAF Class 1 v12 |
| SAE1 | 19q13.32 | 1 | 0.03 | 0 | 0.00 | 0.815 | 0.869 | (A) BRAF Class 1 v12 |
| SCAP | 3p21.31 | 1 | 0.03 | 0 | 0.00 | 0.815 | 0.869 | (A) BRAF Class 1 v12 |
| SCARA5 | 8p21.1 | 1 | 0.03 | 0 | 0.00 | 0.815 | 0.869 | (A) BRAF Class 1 v12 |
| SDCCAG8 | 1q43-q44 | 1 | 0.03 | 0 | 0.00 | 0.815 | 0.869 | (A) BRAF Class 1 v12 |
| SEMA6D | 15q21.1 | 1 | 0.03 | 0 | 0.00 | 0.815 | 0.869 | (A) BRAF Class 1 v12 |
| SIK3 | 11q23.3 | 1 | 0.03 | 0 | 0.00 | 0.815 | 0.869 | (A) BRAF Class 1 v12 |
| SLC30A5 | q13.1-q13. | 1 | 0.03 | 0 | 0.00 | 0.815 | 0.869 | (A) BRAF Class 1 v12 |
| SND1 | 7q32.1 | 1 | 0.03 | 0 | 0.00 | 0.815 | 0.869 | (A) BRAF Class 1 v12 |
| SPG11 | 15q21.1 | 1 | 0.03 | 0 | 0.00 | 0.815 | 0.869 | (A) BRAF Class 1 v12 |
| TCAF2P1 | 7q35 | 1 | 0.03 | 0 | 0.00 | 0.815 | 0.869 | (A) BRAF Class 1 v12 |
| TRAJ32 | 14q11.2 | 1 | 0.03 | 0 | 0.00 | 0.815 | 0.869 | (A) BRAF Class 1 v12 |
| TRAJ42 | 14q11.2 | 1 | 0.03 | 0 | 0.00 | 0.815 | 0.869 | (A) BRAF Class 1 v12 |
| TRAJ53 | 14q11.2 | 1 | 0.03 | 0 | 0.00 | 0.815 | 0.869 | (A) BRAF Class 1 v12 |
| TRBJ2-2 | 7q34 | 1 | 0.03 | 0 | 0.00 | 0.815 | 0.869 | (A) BRAF Class 1 v12 |
| TRGJP | 7p14.1 | 1 | 0.03 | 0 | 0.00 | 0.815 | 0.869 | (A) BRAF Class 1 v12 |
| TSPOAP1 | 17q22 | 1 | 0.03 | 0 | 0.00 | 0.815 | 0.869 | (A) BRAF Class 1 v12 |
| UGT1A6 | 2q37.1 | 1 | 0.03 | 0 | 0.00 | 0.815 | 0.869 | (A) BRAF Class 1 v12 |
| ZNF423 | 16q12.1 | 1 | 0.03 | 0 | 0.00 | 0.815 | 0.869 | (A) BRAF Class 1 v12 |
| ZNF599 | 19q13.11 | 1 | 0.03 | 0 | 0.00 | 0.815 | 0.869 | (A) BRAF Class 1 v12 |
| UGT1A1 | 2q37.1 | 1 | 0.40 | 0 | 0.00 | 0.826 | 0.869 | (A) BRAF Class 1 v12 |
| TRRAP | 7q22.1 | 1 | 0.45 | 0 | 0.00 | 0.827 | 0.869 | (A) BRAF Class 1 v12 |
| MLLT1 | 19p13.3 | 1 | 1.59 | 0 | 0.00 | 0.829 | 0.869 | (A) BRAF Class 1 v12 |
| CDH5 | 16q21 | 1 | 0.50 | 0 | 0.00 | 0.83 | 0.869 | (A) BRAF Class 1 v12 |
| MLF1 | 3q25.32 | 1 | 1.89 | 0 | 0.00 | 0.841 | 0.869 | (A) BRAF Class 1 v12 |
| KBTBD4 | 11p11.2 | 1 | 0.98 | 0 | 0.00 | 0.843 | 0.869 | (A) BRAF Class 1 v12 |
| IRF8 | 16q24.1 | 1 | 0.81 | 0 | 0.00 | 0.844 | 0.869 | (A) BRAF Class 1 v12 |
| PTPN1 | 20q13.13 | 1 | 0.60 | 0 | 0.00 | 0.844 | 0.869 | (A) BRAF Class 1 v12 |
| TBL1XR1 | 3q26.32 | 1 | 0.97 | 0 | 0.00 | 0.844 | 0.869 | (A) BRAF Class 1 v12 |
| IL6ST | 5q11.2 | 1 | 0.50 | 0 | 0.00 | 0.845 | 0.869 | (A) BRAF Class 1 v12 |
| XBP1 | 2q12.1\|22q | 1 | 0.45 | 0 | 0.00 | 0.845 | 0.869 | (A) BRAF Class 1 v12 |
| ATF1 | 12q13.12 | 1 | 0.39 | 0 | 0.00 | 0.845 | 0.869 | (A) BRAF Class 1 v12 |
| DDIT3 | 12q13.3 | 1 | 0.39 | 0 | 0.00 | 0.845 | 0.869 | (A) BRAF Class 1 v12 |
| MAML2 | 11q21 | 1 | 0.39 | 0 | 0.00 | 0.845 | 0.869 | (A) BRAF Class 1 v12 |
| TFEB | 6p21.1 | 1 | 0.45 | 0 | 0.00 | 0.845 | 0.869 | (A) BRAF Class 1 v12 |
| EGR1 | 5q31.2 | 1 | 0.87 | 0 | 0.00 | 0.846 | 0.869 | (A) BRAF Class 1 v12 |
| GNB1 | 1p36.33 | 1 | 2.27 | 0 | 0.00 | 0.846 | 0.869 | (A) BRAF Class 1 v12 |
| AFDN | 6q27 | 1 | 0.86 | 0 | 0.00 | 0.847 | 0.869 | (A) BRAF Class 1 v12 |
| MRTFA | 2q13.1-q13. | 1 | 0.86 | 0 | 0.00 | 0.847 | 0.869 | (A) BRAF Class 1 v12 |
| PRDM16 | 1p36.32 | 1 | 0.86 | 0 | 0.00 | 0.847 | 0.869 | (A) BRAF Class 1 v12 |
| BCL2L2 | 14q11.2 | 1 | 0.14 | 0 | 0.00 | 0.847 | 0.869 | (A) BRAF Class 1 v12 |
| ECT2L | 6q24.1 | 1 | 1.00 | 0 | 0.00 | 0.847 | 0.869 | (A) BRAF Class 1 v12 |

| | | | | | | | | |
|---|---|---|---|---|---|---|---|---|
| FOXO3 | 6q21 | 1 | 0.81 | 0 | 0.00 | 0.848 | 0.869 | (A) BRAF Class 1 v12 |
| ETNK1 | 12p12.1 | 1 | 0.63 | 0 | 0.00 | 0.849 | 0.869 | (A) BRAF Class 1 v12 |
| BIRC5 | 17q25.3 | 1 | 0.53 | 0 | 0.00 | 0.85 | 0.869 | (A) BRAF Class 1 v12 |
| SP140 | 2q37.1 | 1 | 1.96 | 0 | 0.00 | 0.85 | 0.869 | (A) BRAF Class 1 v12 |
| GID4 | [illegible]11.2\|17p1[illegible] | 1 | 0.17 | 0 | 0.00 | 0.85 | 0.869 | (A) BRAF Class 1 v12 |
| LCK | 1p35.2 | 1 | 0.80 | 0 | 0.00 | 0.85 | 0.869 | (A) BRAF Class 1 v12 |
| SSX2 | Xp11.22 | 1 | 1.75 | 0 | 0.00 | 0.851 | 0.869 | (A) BRAF Class 1 v12 |
| FOSB | 19q13.32 | 1 | 1.09 | 0 | 0.00 | 0.852 | 0.869 | (A) BRAF Class 1 v12 |
| KRIT1 | 7q21.2 | 1 | 1.09 | 0 | 0.00 | 0.852 | 0.869 | (A) BRAF Class 1 v12 |
| MAML3 | 4q31.1 | 1 | 1.09 | 0 | 0.00 | 0.852 | 0.869 | (A) BRAF Class 1 v12 |
| OTX2 | 14q22.3 | 1 | 1.09 | 0 | 0.00 | 0.852 | 0.869 | (A) BRAF Class 1 v12 |
| TBXT | 6q27 | 1 | 1.09 | 0 | 0.00 | 0.852 | 0.869 | (A) BRAF Class 1 v12 |
| ARNT | 1q21.3 | 1 | 0.82 | 0 | 0.00 | 0.853 | 0.869 | (A) BRAF Class 1 v12 |
| HLF | 17q22 | 1 | 0.82 | 0 | 0.00 | 0.853 | 0.869 | (A) BRAF Class 1 v12 |
| MUC1 | 1q22 | 1 | 0.82 | 0 | 0.00 | 0.853 | 0.869 | (A) BRAF Class 1 v12 |
| NFKB2 | 10q24.32 | 1 | 0.82 | 0 | 0.00 | 0.853 | 0.869 | (A) BRAF Class 1 v12 |
| PER1 | 17p13.1 | 1 | 0.82 | 0 | 0.00 | 0.853 | 0.869 | (A) BRAF Class 1 v12 |
| PSIP1 | 9p22.3 | 1 | 0.82 | 0 | 0.00 | 0.853 | 0.869 | (A) BRAF Class 1 v12 |
| TLX1 | 10q24.31 | 1 | 0.82 | 0 | 0.00 | 0.853 | 0.869 | (A) BRAF Class 1 v12 |
| TPR | 1q31.1 | 1 | 0.82 | 0 | 0.00 | 0.853 | 0.869 | (A) BRAF Class 1 v12 |
| ZNF384 | 12p13.31 | 1 | 0.82 | 0 | 0.00 | 0.853 | 0.869 | (A) BRAF Class 1 v12 |
| ZNF521 | 18q11.2 | 1 | 0.82 | 0 | 0.00 | 0.853 | 0.869 | (A) BRAF Class 1 v12 |
| SPI1 | 11p11.2 | 1 | 0.90 | 0 | 0.00 | 0.854 | 0.869 | (A) BRAF Class 1 v12 |
| HIP1 | 7q11.23 | 1 | 1.11 | 0 | 0.00 | 0.857 | 0.869 | (A) BRAF Class 1 v12 |
| ARFGAP3 | 22q13.2 | 1 | 2.08 | 0 | 0.00 | 0.857 | 0.869 | (A) BRAF Class 1 v12 |
| CBLC | 19q13.32 | 1 | 2.78 | 0 | 0.00 | 0.857 | 0.869 | (A) BRAF Class 1 v12 |
| GNAI3 | 1p13.3 | 1 | 2.08 | 0 | 0.00 | 0.857 | 0.869 | (A) BRAF Class 1 v12 |
| LRIG3 | 12q14.1 | 1 | 2.78 | 0 | 0.00 | 0.857 | 0.869 | (A) BRAF Class 1 v12 |
| SNX31 | 8q22.3 | 1 | 2.08 | 0 | 0.00 | 0.857 | 0.869 | (A) BRAF Class 1 v12 |
| TMEM216 | 11q12.2 | 1 | 2.08 | 0 | 0.00 | 0.857 | 0.869 | (A) BRAF Class 1 v12 |
| DUSP6 | 12q21.33 | 1 | 1.16 | 0 | 0.00 | 0.86 | 0.869 | (A) BRAF Class 1 v12 |
| HSPA5 | 9q33.3 | 1 | 1.16 | 0 | 0.00 | 0.86 | 0.869 | (A) BRAF Class 1 v12 |
| AURKC | 19q13.43 | 1 | 0.85 | 0 | 0.00 | 0.86 | 0.869 | (A) BRAF Class 1 v12 |
| BLNK | 10q24.1 | 1 | 0.85 | 0 | 0.00 | 0.86 | 0.869 | (A) BRAF Class 1 v12 |
| CYP2C19 | 10q23.33 | 1 | 0.85 | 0 | 0.00 | 0.86 | 0.869 | (A) BRAF Class 1 v12 |
| DST | 6p12.1 | 1 | 0.85 | 0 | 0.00 | 0.86 | 0.869 | (A) BRAF Class 1 v12 |
| GDNF | 5p13.2 | 1 | 0.85 | 0 | 0.00 | 0.86 | 0.869 | (A) BRAF Class 1 v12 |
| IKBKB | 8p11.21 | 1 | 0.85 | 0 | 0.00 | 0.86 | 0.869 | (A) BRAF Class 1 v12 |
| ITGA9 | 3p22.2 | 1 | 0.85 | 0 | 0.00 | 0.86 | 0.869 | (A) BRAF Class 1 v12 |
| ITGB3 | 17q21.32 | 1 | 0.85 | 0 | 0.00 | 0.86 | 0.869 | (A) BRAF Class 1 v12 |
| PLEKHG5 | 1p36.31 | 1 | 0.85 | 0 | 0.00 | 0.86 | 0.869 | (A) BRAF Class 1 v12 |
| RRM1 | 11p15.4 | 1 | 0.85 | 0 | 0.00 | 0.86 | 0.869 | (A) BRAF Class 1 v12 |
| TAF1L | 9p21.1 | 1 | 0.85 | 0 | 0.00 | 0.86 | 0.869 | (A) BRAF Class 1 v12 |
| THBS1 | 15q14 | 1 | 0.85 | 0 | 0.00 | 0.86 | 0.869 | (A) BRAF Class 1 v12 |
| ETAA1 | 2p14 | 1 | 3.13 | 0 | 0.00 | 0.865 | 0.871 | (A) BRAF Class 1 v12 |
| FOXF1 | 16q24.1 | 1 | 3.13 | 0 | 0.00 | 0.865 | 0.871 | (A) BRAF Class 1 v12 |
| SLFN11 | 17q12 | 1 | 3.13 | 0 | 0.00 | 0.865 | 0.871 | (A) BRAF Class 1 v12 |
| OLIG2 | 21q22.11 | 1 | 3.70 | 0 | 0.00 | 0.871 | 0.877 | (A) BRAF Class 1 v12 |

| | | | | | | | | |
|---|---|---|---|---|---|---|---|---|
| MCM7 | 7q22.1 | 1 | 3.23 | 0 | 0.00 | 0.886 | 0.89 | (A) BRAF Class 1 v12 |
| MAGOH | 1p32.3 | 1 | 0.99 | 0 | 0.00 | 0.886 | 0.89 | (A) BRAF Class 1 v12 |
| CD3EAP | 19q13.32 | 1 | 0.30 | 0 | 0.00 | 0.893 | 0.894 | (A) BRAF Class 1 v12 |
| TSPAN31 | 12q14.1 | 1 | 0.30 | 0 | 0.00 | 0.893 | 0.894 | (A) BRAF Class 1 v12 |
| ATXN2 | 12q24.12 | 1 | 4.00 | 0 | 0.00 | 0.893 | 0.894 | (A) BRAF Class 1 v12 |
| DUSP22 | 6p25.3 | 1 | 4.00 | 0 | 0.00 | 0.893 | 0.894 | (A) BRAF Class 1 v12 |
| BRAF | 7q34 | 3344 | 100.00 | 759 | 100.00 | 1 | 1 | (B) BRAF Class 2 |

#### Supplementary Table 4

| Gene | Cytoband | Class 1 Occurrence | Class 1 % | Class 2 Occurrence | Class 2 % | SumClass | pvalue | qvalue | Enriched In |
|---|---|---|---|---|---|---|---|---|---|
| KDM6A | Xp11.3 | 65 | 2.65 | 31 | 5.54 | 96 | 0.0007 | 0.0412 | (B) BRAF Class 2 |
| KDR | 4q12 | 163 | 5.12 | 58 | 8.61 | 221 | 0.0005 | 0.0296 | (B) BRAF Class 2 |
| FOXA1 | 14q21.1 | 39 | 1.98 | 23 | 5.12 | 62 | 0.0004 | 0.0277 | (B) BRAF Class 2 |
| NFE2L2 | 2q31.2 | 31 | 1.22 | 20 | 3.50 | 51 | 0.0004 | 0.0268 | (B) BRAF Class 2 |
| TGFBR2 | 3p24.1 | 112 | 7.16 | 9 | 2.48 | 121 | 0.0002 | 0.0197 | (A) BRAF Class 1 v12 |
| CBLB | 3q13.11 | 16 | 1.52 | 14 | 6.19 | 30 | 0.0002 | 0.0166 | (B) BRAF Class 2 |
| FGFR1 | 8p11.23 | 55 | 1.65 | 30 | 3.95 | 85 | 0.0002 | 0.0148 | (B) BRAF Class 2 |
| ATR | 3q23 | 126 | 5.72 | 55 | 10.91 | 181 | 5E-05 | 0.005 | (B) BRAF Class 2 |
| POT1 | 7q31.33 | 13 | 1.55 | 16 | 8.65 | 29 | 5E-06 | 0.0006 | (B) BRAF Class 2 |
| FLT3 | 13q12.2 | 100 | 3.11 | 51 | 7.52 | 151 | 5E-07 | 6E-05 | (B) BRAF Class 2 |
| TERT | 5p15.33 | 842 | 34.23 | 128 | 23.23 | 970 | 2E-07 | 3E-05 | (A) BRAF Class 1 v12 |
| NF1 | 17q11.2 | 142 | 5.59 | 82 | 13.20 | 224 | 6E-10 | 9E-08 | (B) BRAF Class 2 |
| RNF43 | 17q22 | 260 | 11.71 | 17 | 3.35 | 277 | 4E-10 | 7E-08 | (A) BRAF Class 1 v12 |
| KEAP1 | 19p13.2 | 53 | 2.24 | 45 | 8.33 | 98 | 3E-10 | 6E-08 | (B) BRAF Class 2 |
| TP53 | 17p13.1 | 892 | 26.67 | 282 | 39.66 | 1174 | 8E-12 | 2E-09 | (B) BRAF Class 2 |
| NRAS | 1p13.2 | 35 | 1.05 | 48 | 6.32 | 83 | 8E-16 | 3E-13 | (B) BRAF Class 2 |
| KRAS | 12p12.1 | 37 | 1.11 | 67 | 8.83 | 104 | 1E-25 | 8E-23 | (B) BRAF Class 2 |
| STK11 | 19p13.3 | 57 | 1.74 | 80 | 11.53 | 137 | 5E-28 | 5E-25 | (B) BRAF Class 2 |

#### Supplementary Table 5

| Gene | cytoband | Class 1 occurrence | Class 1 % | Class 3 occurrence | Class 3 % | pvalue | qvalue | Enriched In |
|---|---|---|---|---|---|---|---|---|
| KRAS | 12p12.1 | 36 | 1.08 | 129 | 17.22 | 2.62E-66 | 2.9E-63 | (B) BRAF Class 3 |
| NRAS | 1p13.2 | 34 | 1.02 | 94 | 12.55 | 8.13E-44 | 4.5E-41 | (B) BRAF Class 3 |
| NF1 | 17q11.2 | 144 | 5.66 | 141 | 23.62 | 7.45E-35 | 2.75E-32 | (B) BRAF Class 3 |
| STK11 | 19p13.3 | 57 | 1.74 | 79 | 10.93 | 1.91E-26 | 5.29E-24 | (B) BRAF Class 3 |
| TP53 | 17p13.1 | 892 | 26.64 | 328 | 44.15 | 2.97E-20 | 6.58E-18 | (B) BRAF Class 3 |
| KEAP1 | 19p13.2 | 53 | 2.24 | 63 | 11.07 | 1.03E-17 | 1.91E-15 | (B) BRAF Class 3 |
| APC | 5q22.2 | 356 | 11.33 | 160 | 22.63 | 3.86E-14 | 6.1E-12 | (B) BRAF Class 3 |
| HRAS | 11p15.5 | 17 | 0.51 | 26 | 3.48 | 7.73E-10 | 1.07E-07 | (B) BRAF Class 3 |
| RNF43 | 17q22 | 259 | 11.64 | 21 | 4.02 | 1.14E-08 | 1.41E-06 | (A) BRAF Class 1 v12 |
| SMARCA4 | 19p13.2 | 159 | 6.26 | 76 | 12.90 | 1.73E-07 | 1.92E-05 | (B) BRAF Class 3 |
| NOTCH2 | 1p12 | 109 | 4.36 | 57 | 10.00 | 4.77E-07 | 4.8E-05 | (B) BRAF Class 3 |
| PTPN11 | 12q24.13 | 35 | 1.08 | 27 | 3.75 | 2.75E-06 | 0.000253 | (B) BRAF Class 3 |
| ERBB4 | 2q34 | 206 | 6.36 | 83 | 11.53 | 3.34E-06 | 0.000284 | (B) BRAF Class 3 |
| MET | 7q31.2 | 121 | 3.65 | 57 | 7.71 | 3.88E-06 | 0.000307 | (B) BRAF Class 3 |
| ERBB2 | 17q12 | 88 | 2.63 | 46 | 6.19 | 4.21E-06 | 0.00031 | (B) BRAF Class 3 |
| PPP2R1A | 19q13.41 | 34 | 1.54 | 27 | 5.19 | 4.61E-06 | 0.000319 | (B) BRAF Class 3 |
| FLT3 | 13q12.2 | 100 | 3.10 | 49 | 6.87 | 7.08E-06 | 0.000461 | (B) BRAF Class 3 |
| MCL1 | 1q21.2 | 25 | 1.02 | 22 | 3.80 | 1.22E-05 | 0.000748 | (B) BRAF Class 3 |
| TET2 | 4q24 | 155 | 6.23 | 68 | 11.53 | 1.62E-05 | 0.000946 | (B) BRAF Class 3 |
| ARID1A | 1p36.11 | 269 | 10.59 | 99 | 16.81 | 3.24E-05 | 0.001793 | (B) BRAF Class 3 |
| ERBB3 | 12q13.2 | 81 | 3.15 | 41 | 6.82 | 6.11E-05 | 0.003221 | (B) BRAF Class 3 |
| PTPRD | 9p24.1-p23 | 184 | 10.62 | 71 | 17.93 | 7.47E-05 | 0.00369 | (B) BRAF Class 3 |
| PDGFRA | 4q12 | 102 | 3.13 | 46 | 6.33 | 8.11E-05 | 0.00369 | (B) BRAF Class 3 |
| ATM | 11q22.3 | 260 | 8.16 | 91 | 12.93 | 8.16E-05 | 0.00369 | (B) BRAF Class 3 |
| PTPRT | 0q12-q13.1 | 160 | 11.36 | 61 | 19.87 | 8.33E-05 | 0.00369 | (B) BRAF Class 3 |
| EPHA3 | 3p11.1 | 92 | 4.85 | 43 | 9.95 | 8.85E-05 | 0.003769 | (B) BRAF Class 3 |
| SYNE1 | 6q25.2 | 11 | 2.14 | 11 | 11.34 | 0.000149 | 0.005777 | (B) BRAF Class 3 |
| RAF1 | 3p25.2 | 37 | 1.42 | 24 | 3.95 | 0.000149 | 0.005777 | (B) BRAF Class 3 |
| TERT | 5p15.33 | 842 | 34.20 | 145 | 26.22 | 0.000151 | 0.005777 | (A) BRAF Class 1 v12 |
| BRIP1 | 17q23.2 | 65 | 2.64 | 34 | 5.86 | 0.00019 | 0.006998 | (B) BRAF Class 3 |
| CBLB | 3q13.11 | 16 | 1.51 | 15 | 5.88 | 0.000218 | 0.007776 | (B) BRAF Class 3 |
| FLT1 | 13q12.3 | 132 | 5.37 | 55 | 9.48 | 0.000281 | 0.009684 | (B) BRAF Class 3 |
| SRC | 20q11.23 | 22 | 0.83 | 16 | 2.86 | 0.000289 | 0.009684 | (B) BRAF Class 3 |
| RET | 10q11.21 | 123 | 3.67 | 50 | 6.70 | 0.000301 | 0.009808 | (B) BRAF Class 3 |
| PIK3CA | 3q26.32 | 311 | 9.29 | 102 | 13.67 | 0.00032 | 0.0101 | (B) BRAF Class 3 |
| FGFR2 | 10q26.13 | 89 | 2.66 | 39 | 5.23 | 0.000439 | 0.0135 | (B) BRAF Class 3 |
| RHPN2 | 19q13.11 | 6 | 1.69 | 8 | 11.59 | 0.000456 | 0.0136 | (B) BRAF Class 3 |
| ARID2 | 12q12 | 180 | 7.41 | 67 | 11.90 | 0.000525 | 0.0151 | (B) BRAF Class 3 |
| HLA-A | 6p22.1 | 48 | 3.95 | 1 | 0.37 | 0.000533 | 0.0151 | (A) BRAF Class 1 v12 |
| NOTCH1 | 9q34.3 | 178 | 5.42 | 64 | 8.80 | 0.00058 | 0.0157 | (B) BRAF Class 3 |
| CARD11 | 7p22.2 | 157 | 6.38 | 61 | 10.52 | 0.000581 | 0.0157 | (B) BRAF Class 3 |
| KMT2D | 12q13.12 | 362 | 14.73 | 118 | 20.34 | 0.000703 | 0.0184 | (B) BRAF Class 3 |
| KDR | 4q12 | 162 | 5.08 | 59 | 8.38 | 0.000714 | 0.0184 | (B) BRAF Class 3 |
| MAP3K5 | 6q23.3 | 0 | 0.00 | 4 | 12.12 | 0.000778 | 0.0196 | (B) BRAF Class 3 |
| DICER1 | 14q32.13 | 91 | 3.78 | 39 | 7.01 | 0.001033 | 0.0254 | (B) BRAF Class 3 |
| COL2A1 | 12q13.11 | 5 | 3.18 | 7 | 21.21 | 0.001095 | 0.0262 | (B) BRAF Class 3 |

| | | | | | | | | |
|---|---|---|---|---|---|---|---|---|
| PMS1 | 2q32.2 | 55 | 2.38 | 27 | 5.14 | 0.00112 | 0.0262 | (B) BRAF Class 3 |
| POT1 | 7q31.33 | 13 | 1.54 | 12 | 5.83 | 0.001137 | 0.0262 | (B) BRAF Class 3 |
| TP63 | 3q28 | 57 | 4.57 | 26 | 9.74 | 0.001246 | 0.0282 | (B) BRAF Class 3 |
| CBL | 11q23.3 | 57 | 2.22 | 28 | 4.67 | 0.001277 | 0.0282 | (B) BRAF Class 3 |
| ASXL1 | 20q11.21 | 132 | 5.30 | 52 | 8.81 | 0.0013 | 0.0282 | (B) BRAF Class 3 |
| SMAD3 | 15q22.33 | 25 | 1.70 | 16 | 4.85 | 0.001331 | 0.0283 | (B) BRAF Class 3 |
| NTRK3 | 15q25.3 | 112 | 4.41 | 45 | 7.64 | 0.001392 | 0.0291 | (B) BRAF Class 3 |
| BCOR | Xp11.4 | 157 | 6.40 | 19 | 3.26 | 0.001469 | 0.0301 | (A) BRAF Class 1 v12 |
| ANKRD26 | 10p12.1 | 45 | 13.51 | 0 | 0.00 | 0.001551 | 0.0312 | (A) BRAF Class 1 v12 |
| MSH3 | 5q14.1 | 94 | 8.82 | 8 | 3.35 | 0.001712 | 0.0338 | (A) BRAF Class 1 v12 |
| RUNX1T1 | 8q21.3 | 57 | 4.56 | 27 | 9.34 | 0.001796 | 0.0348 | (B) BRAF Class 3 |
| ZRSR2 | Xp22.2 | 41 | 1.83 | 1 | 0.20 | 0.001821 | 0.0348 | (A) BRAF Class 1 v12 |
| FGFR4 | 5q35.2 | 71 | 2.76 | 32 | 5.30 | 0.001996 | 0.0374 | (B) BRAF Class 3 |
| MAP3K13 | 3q27.2 | 20 | 1.52 | 14 | 4.55 | 0.002103 | 0.0388 | (B) BRAF Class 3 |
| EGFR | 7p11.2 | 134 | 4.00 | 49 | 6.57 | 0.002211 | 0.0401 | (B) BRAF Class 3 |
| RRAS2 | 11p15.2 | 3 | 0.45 | 6 | 3.75 | 0.002513 | 0.0449 | (B) BRAF Class 3 |
| KIT | 4q12 | 80 | 2.39 | 33 | 4.41 | 0.002742 | 0.0482 | (B) BRAF Class 3 |
| RHOH | 4p14 | 3 | 0.44 | 6 | 3.53 | 0.00283 | 0.049 | (B) BRAF Class 3 |
| FGFR1 | 8p11.23 | 55 | 1.65 | 25 | 3.36 | 0.00303 | 0.0514 | (B) BRAF Class 3 |
| DDR2 | 1q23.3 | 55 | 2.06 | 26 | 4.16 | 0.003064 | 0.0514 | (B) BRAF Class 3 |
| EZH1 | 17q21.2 | 7 | 1.06 | 8 | 5.00 | 0.003271 | 0.0541 | (B) BRAF Class 3 |
| ROS1 | 6q22.1 | 244 | 9.37 | 81 | 13.24 | 0.00338 | 0.0549 | (B) BRAF Class 3 |
| JAK3 | 19p13.11 | 103 | 3.20 | 39 | 5.47 | 0.003443 | 0.0549 | (B) BRAF Class 3 |
| JAK2 | 9p24.1 | 62 | 1.95 | 27 | 3.80 | 0.003497 | 0.0549 | (B) BRAF Class 3 |
| PBRM1 | 3p21.1 | 108 | 4.56 | 43 | 7.56 | 0.003524 | 0.0549 | (B) BRAF Class 3 |
| KDM6A | Xp11.3 | 65 | 2.64 | 29 | 5.00 | 0.003694 | 0.0564 | (B) BRAF Class 3 |
| SETBP1 | 18q12.3 | 105 | 7.14 | 39 | 11.96 | 0.003719 | 0.0564 | (B) BRAF Class 3 |
| GRIN2A | 16p13.2 | 157 | 10.26 | 54 | 15.61 | 0.003868 | 0.0579 | (B) BRAF Class 3 |
| PTK2B | 8p21.2 | 22 | 3.40 | 15 | 8.82 | 0.004048 | 0.0597 | (B) BRAF Class 3 |
| IGF1 | 12q23.2 | 6 | 0.47 | 7 | 2.41 | 0.004268 | 0.0618 | (B) BRAF Class 3 |
| MAP3K1 | 5q11.2 | 98 | 3.99 | 39 | 6.72 | 0.004299 | 0.0618 | (B) BRAF Class 3 |
| H3F3B | 17q25.1 | 6 | 0.32 | 7 | 1.64 | 0.004467 | 0.0625 | (B) BRAF Class 3 |
| PTPRS | 19p13.3 | 110 | 9.34 | 11 | 4.35 | 0.004484 | 0.0625 | (A) BRAF Class 1 v12 |
| ELOC | 8q21.11 | 6 | 0.33 | 7 | 1.69 | 0.004543 | 0.0625 | (B) BRAF Class 3 |
| ZFHX4 | 8q21.13 | 19 | 12.10 | 11 | 33.33 | 0.004576 | 0.0625 | (B) BRAF Class 3 |
| NFE2L2 | 2q31.2 | 31 | 1.22 | 17 | 2.89 | 0.004696 | 0.0629 | (B) BRAF Class 3 |
| ABCB11 | 2q31.1 | 33 | 5.04 | 18 | 11.32 | 0.004715 | 0.0629 | (B) BRAF Class 3 |
| PLCB4 | 0p12.3-p12. | 5 | 2.30 | 6 | 13.04 | 0.004874 | 0.0642 | (B) BRAF Class 3 |
| HIF1A | 14q23.2 | 0 | 0.00 | 3 | 5.56 | 0.005094 | 0.0663 | (B) BRAF Class 3 |
| SPOP | 17q21.33 | 20 | 0.91 | 13 | 2.50 | 0.005236 | 0.0674 | (B) BRAF Class 3 |
| PRKCA | 17q24.2 | 0 | 0.00 | 3 | 5.77 | 0.005326 | 0.0678 | (B) BRAF Class 3 |
| EPHA7 | 6q16.1 | 109 | 6.15 | 39 | 10.05 | 0.005445 | 0.0685 | (B) BRAF Class 3 |
| TRAF7 | 16p13.3 | 44 | 2.26 | 21 | 4.69 | 0.005627 | 0.07 | (B) BRAF Class 3 |
| OR5L1 | 11q12.1 | 1 | 1.16 | 4 | 18.18 | 0.00588 | 0.0723 | (B) BRAF Class 3 |
| QKI | 6q26 | 48 | 3.62 | 3 | 0.95 | 0.006342 | 0.0771 | (A) BRAF Class 1 v12 |
| EPHA2 | 1p36.13 | 0 | 0.00 | 3 | 5.77 | 0.007125 | 0.0857 | (B) BRAF Class 3 |
| DGKH | 13q14.11 | 0 | 0.00 | 3 | 13.64 | 0.007543 | 0.0895 | (B) BRAF Class 3 |
| YES1 | 18p11.32 | 21 | 1.78 | 12 | 4.74 | 0.007603 | 0.0895 | (B) BRAF Class 3 |

| | | | | | | | | |
|---|---|---|---|---|---|---|---|---|
| ESR1 | 6q25.1-q25.2 | 54 | 2.05 | 24 | 3.88 | 0.0077 | 0.0895 | (B) BRAF Class 3 |
| XRCC3 | 14q32.33 | 20 | 2.92 | 0 | 0.00 | 0.007763 | 0.0895 | (A) BRAF Class 1 v12 |
| BRCA1 | 17q21.31 | 103 | 4.06 | 39 | 6.52 | 0.007961 | 0.0909 | (B) BRAF Class 3 |
| BARD1 | 2q35 | 38 | 1.79 | 19 | 3.72 | 0.0085 | 0.096 | (B) BRAF Class 3 |
| NCKAP5 | 2q21.2 | 5 | 5.81 | 6 | 27.27 | 0.008682 | 0.0971 | (B) BRAF Class 3 |
| RASA2 | 3q23 | 2 | 0.91 | 4 | 8.70 | 0.009126 | 0.101 | (B) BRAF Class 3 |
| RBM10 | Xp11.3 | 83 | 4.02 | 33 | 6.67 | 0.009728 | 0.106 | (B) BRAF Class 3 |
| ATRX | Xq21.1 | 167 | 6.58 | 56 | 9.51 | 0.009736 | 0.106 | (B) BRAF Class 3 |
| PRSS1 | 7q34 | 20 | 2.74 | 13 | 6.77 | 0.0102 | 0.11 | (B) BRAF Class 3 |
| NTRK2 | 9q21.33 | 53 | 2.09 | 23 | 3.90 | 0.0104 | 0.111 | (B) BRAF Class 3 |
| NRG1 | 8p12 | 52 | 5.05 | 21 | 9.55 | 0.0105 | 0.111 | (B) BRAF Class 3 |
| IL7R | 5p13.2 | 99 | 4.72 | 37 | 7.49 | 0.0109 | 0.114 | (B) BRAF Class 3 |
| HIST1H3C | 6p22.2 | 8 | 0.46 | 7 | 1.79 | 0.0111 | 0.114 | (B) BRAF Class 3 |
| IKZF1 | 7p12.2 | 123 | 5.02 | 44 | 7.61 | 0.0111 | 0.114 | (B) BRAF Class 3 |
| PML | 15q24.1 | 28 | 4.04 | 1 | 0.58 | 0.0122 | 0.124 | (A) BRAF Class 1 v12 |
| SPTA1 | 1q23.1 | 95 | 15.10 | 30 | 24.00 | 0.0125 | 0.126 | (B) BRAF Class 3 |
| MECOM | 3q26.2 | 70 | 7.32 | 28 | 12.28 | 0.0129 | 0.129 | (B) BRAF Class 3 |
| NF2 | 22q12.2 | 58 | 2.25 | 24 | 4.03 | 0.013 | 0.129 | (B) BRAF Class 3 |
| DAXX | 6p21.32 | 41 | 1.99 | 19 | 3.89 | 0.0133 | 0.13 | (B) BRAF Class 3 |
| LRP1B | 2q22.1-q22.2 | 146 | 21.31 | 43 | 30.50 | 0.0136 | 0.132 | (B) BRAF Class 3 |
| TLR4 | 9q33.1 | 13 | 3.08 | 9 | 8.91 | 0.0142 | 0.136 | (B) BRAF Class 3 |
| LIFR | 5p13.1 | 4 | 2.15 | 5 | 11.36 | 0.0143 | 0.136 | (B) BRAF Class 3 |
| AXL | 19q13.2 | 96 | 3.78 | 35 | 5.94 | 0.0149 | 0.141 | (B) BRAF Class 3 |
| CASR | q13.33-q21. | 3 | 3.23 | 3 | 27.27 | 0.0151 | 0.141 | (B) BRAF Class 3 |
| SLIT2 | 4p15.31 | 52 | 8.27 | 19 | 15.20 | 0.0154 | 0.143 | (B) BRAF Class 3 |
| PAX7 | 1p36.13 | 15 | 2.64 | 8 | 7.77 | 0.0156 | 0.144 | (B) BRAF Class 3 |
| FOXO1 | 13q14.11 | 27 | 1.99 | 13 | 4.47 | 0.0157 | 0.144 | (B) BRAF Class 3 |
| CCNQ | Xq28 | 6 | 1.14 | 6 | 4.72 | 0.016 | 0.145 | (B) BRAF Class 3 |
| XRCC4 | 5q14.2 | 10 | 1.71 | 8 | 5.41 | 0.0161 | 0.145 | (B) BRAF Class 3 |
| EZH2 | 7q36.1 | 48 | 1.59 | 20 | 2.95 | 0.0165 | 0.147 | (B) BRAF Class 3 |
| FANCA | 16q24.3 | 90 | 3.54 | 33 | 5.60 | 0.0167 | 0.148 | (B) BRAF Class 3 |
| CCND2 | 12p13.32 | 25 | 0.99 | 13 | 2.21 | 0.0175 | 0.153 | (B) BRAF Class 3 |
| HLA-B | 6p21.33 | 25 | 4.11 | 1 | 0.63 | 0.0178 | 0.154 | (A) BRAF Class 1 v12 |
| EBF1 | 5q33.3 | 1 | 0.44 | 3 | 6.25 | 0.0179 | 0.154 | (B) BRAF Class 3 |
| PTPRB | 12q15 | 8 | 5.10 | 6 | 18.18 | 0.0187 | 0.16 | (B) BRAF Class 3 |
| B2M | 15q21.1 | 87 | 4.01 | 10 | 2.03 | 0.0187 | 0.16 | (A) BRAF Class 1 v12 |
| PAK5 | 20p12.2 | 80 | 6.24 | 29 | 9.97 | 0.0193 | 0.163 | (B) BRAF Class 3 |
| MTAP | 9p21.3 | 39 | 6.25 | 19 | 11.52 | 0.0197 | 0.165 | (B) BRAF Class 3 |
| NFKBIZ | 3q12.3 | 14 | 1.53 | 9 | 4.15 | 0.0198 | 0.165 | (B) BRAF Class 3 |
| PPM1D | 17q23.2 | 45 | 2.49 | 19 | 4.59 | 0.02 | 0.165 | (B) BRAF Class 3 |
| PIK3C2G | 12p12.3 | 81 | 6.17 | 30 | 9.74 | 0.0205 | 0.168 | (B) BRAF Class 3 |
| PRDM14 | 8q13.3 | 14 | 2.66 | 9 | 7.09 | 0.021 | 0.171 | (B) BRAF Class 3 |
| RB1 | 13q14.2 | 105 | 3.26 | 35 | 4.93 | 0.0226 | 0.182 | (B) BRAF Class 3 |
| FBXW7 | 4q31.3 | 185 | 5.64 | 56 | 7.71 | 0.0227 | 0.182 | (B) BRAF Class 3 |
| SF3B1 | 2q33.1 | 58 | 2.21 | 23 | 3.76 | 0.023 | 0.182 | (B) BRAF Class 3 |
| CXCR4 | 2q22.1 | 12 | 0.63 | 8 | 1.80 | 0.0231 | 0.182 | (B) BRAF Class 3 |
| PAX5 | 9p13.2 | 54 | 2.17 | 22 | 3.75 | 0.0232 | 0.182 | (B) BRAF Class 3 |
| STAG2 | Xq25 | 74 | 3.02 | 28 | 4.83 | 0.0236 | 0.184 | (B) BRAF Class 3 |

| | | | | | | | | |
|---|---|---|---|---|---|---|---|---|
| PIK3R1 | 5q13.1 | 69 | 2.68 | 26 | 4.37 | 0.024 | 0.185 | (B) BRAF Class 3 |
| FGF14 | 13q33.1 | 13 | 1.83 | 8 | 5.06 | 0.0244 | 0.187 | (B) BRAF Class 3 |
| VEGFA | 6p21.1 | 20 | 0.99 | 11 | 2.28 | 0.0248 | 0.188 | (B) BRAF Class 3 |
| BCL2 | 18q21.33 | 12 | 0.49 | 8 | 1.39 | 0.0249 | 0.188 | (B) BRAF Class 3 |
| USP8 | 15q21.2 | 14 | 2.36 | 9 | 6.08 | 0.0249 | 0.188 | (B) BRAF Class 3 |
| CTNNB1 | 3p22.1 | 116 | 3.47 | 38 | 5.09 | 0.0254 | 0.19 | (B) BRAF Class 3 |
| ETS1 | 11q24.3 | 7 | 1.24 | 5 | 4.85 | 0.0257 | 0.191 | (B) BRAF Class 3 |
| ERCC4 | 16p13.12 | 52 | 2.28 | 5 | 0.93 | 0.0258 | 0.191 | (A) BRAF Class 1 v12 |
| MYCL | 1p34.2 | 33 | 1.30 | 15 | 2.55 | 0.0262 | 0.192 | (B) BRAF Class 3 |
| CDKN1A | 6p21.2 | 15 | 0.63 | 9 | 1.60 | 0.0265 | 0.193 | (B) BRAF Class 3 |
| SMAD2 | 18q21.1 | 42 | 1.71 | 18 | 3.10 | 0.0267 | 0.194 | (B) BRAF Class 3 |
| SESN2 | 1p35.3 | 5 | 0.95 | 5 | 3.94 | 0.0282 | 0.203 | (B) BRAF Class 3 |
| CRKL | 22q11.21 | 18 | 0.73 | 10 | 1.72 | 0.0284 | 0.203 | (B) BRAF Class 3 |
| CDH4 | 20q13.33 | 34 | 5.82 | 16 | 10.81 | 0.0289 | 0.205 | (B) BRAF Class 3 |
| JAK1 | 1p31.3 | 79 | 3.52 | 29 | 5.45 | 0.0294 | 0.206 | (B) BRAF Class 3 |
| IFNGR1 | 6q23.3 | 18 | 1.53 | 0 | 0.00 | 0.0294 | 0.206 | (A) BRAF Class 1 v12 |
| FANCE | 6p21.31 | 19 | 1.20 | 10 | 2.78 | 0.0298 | 0.207 | (B) BRAF Class 3 |
| RRAS | 19q13.33 | 3 | 0.45 | 4 | 2.50 | 0.0299 | 0.207 | (B) BRAF Class 3 |
| GATA6 | 18q11.2 | 28 | 1.97 | 13 | 3.99 | 0.0304 | 0.209 | (B) BRAF Class 3 |
| CREB1 | 2q33.3 | 0 | 0.00 | 2 | 3.64 | 0.0306 | 0.209 | (B) BRAF Class 3 |
| COP1 | .q25.1-q25.: | 34 | 2.25 | 14 | 4.35 | 0.0312 | 0.212 | (B) BRAF Class 3 |
| TERC | 3q26.2 | 0 | 0.00 | 2 | 1.22 | 0.0315 | 0.212 | (B) BRAF Class 3 |
| ITK | 5q33.3 | 12 | 2.01 | 8 | 5.30 | 0.0316 | 0.212 | (B) BRAF Class 3 |
| ATXN2 | 12q24.12 | 1 | 4.00 | 0 | 0.00 | 0.0323 | 0.214 | (A) BRAF Class 1 v12 |
| DUSP22 | 6p25.3 | 1 | 4.00 | 0 | 0.00 | 0.0323 | 0.214 | (A) BRAF Class 1 v12 |
| SMARCA2 | 9p24.3 | 2 | 1.20 | 3 | 9.09 | 0.0326 | 0.214 | (B) BRAF Class 3 |
| SYK | 9q22.2 | 45 | 2.37 | 18 | 4.17 | 0.0329 | 0.214 | (B) BRAF Class 3 |
| EED | 11q14.2 | 14 | 0.88 | 8 | 2.27 | 0.033 | 0.214 | (B) BRAF Class 3 |
| ALK | !p23.2-p23.: | 181 | 5.42 | 54 | 7.27 | 0.0331 | 0.214 | (B) BRAF Class 3 |
| MKRN1 | 7q34 | 0 | 0.00 | 2 | 0.27 | 0.0334 | 0.215 | (B) BRAF Class 3 |
| NTRK1 | 1q23.1 | 100 | 3.85 | 34 | 5.65 | 0.0337 | 0.215 | (B) BRAF Class 3 |
| KMT2A | 11q23.3 | 216 | 8.68 | 66 | 11.24 | 0.0339 | 0.215 | (B) BRAF Class 3 |
| MSH6 | 2p16.3 | 139 | 5.40 | 21 | 3.53 | 0.0342 | 0.215 | (A) BRAF Class 1 v12 |
| TSC2 | 16p13.3 | 133 | 5.23 | 43 | 7.29 | 0.0344 | 0.215 | (B) BRAF Class 3 |
| RASA1 | 5q14.3 | 38 | 2.03 | 16 | 3.69 | 0.0346 | 0.215 | (B) BRAF Class 3 |
| H3F3C | 12p11.21 | 9 | 0.76 | 6 | 2.37 | 0.0349 | 0.215 | (B) BRAF Class 3 |
| XPC | 3p25.1 | 22 | 2.15 | 11 | 4.56 | 0.035 | 0.215 | (B) BRAF Class 3 |
| ROBO1 | 3p12.3 | 7 | 4.38 | 5 | 15.15 | 0.0351 | 0.215 | (B) BRAF Class 3 |
| WAS | Xp11.23 | 10 | 1.44 | 7 | 4.07 | 0.0351 | 0.215 | (B) BRAF Class 3 |
| SMC3 | 10q25.2 | 27 | 1.84 | 12 | 3.73 | 0.0358 | 0.218 | (B) BRAF Class 3 |
| PAX8 | 2q14.1 | 8 | 1.39 | 5 | 4.76 | 0.0363 | 0.22 | (B) BRAF Class 3 |
| NOTCH3 | 19p13.12 | 219 | 9.99 | 66 | 12.84 | 0.0369 | 0.222 | (B) BRAF Class 3 |
| PRKDC | 8q11.21 | 169 | 10.70 | 51 | 14.25 | 0.0371 | 0.222 | (B) BRAF Class 3 |
| SUZ12 | 17q11.2 | 33 | 1.47 | 14 | 2.78 | 0.0373 | 0.222 | (B) BRAF Class 3 |
| FANCD2 | 3p25.3 | 59 | 3.84 | 21 | 6.25 | 0.0383 | 0.226 | (B) BRAF Class 3 |
| NEGR1 | 1p31.1 | 28 | 2.07 | 12 | 4.12 | 0.0384 | 0.226 | (B) BRAF Class 3 |
| CTCF | 16q22.1 | 50 | 2.36 | 20 | 3.91 | 0.0394 | 0.23 | (B) BRAF Class 3 |
| PALB2 | 16p12.2 | 87 | 3.43 | 30 | 5.09 | 0.0395 | 0.23 | (B) BRAF Class 3 |

| | | | | | | | | |
|---|---|---|---|---|---|---|---|---|
| SETD2 | 3p21.31 | 226 | 8.90 | 67 | 11.38 | 0.0398 | 0.231 | (B) BRAF Class 3 |
| TRIM28 | 19q13.43 | 0 | 0.00 | 2 | 9.09 | 0.04 | 0.231 | (B) BRAF Class 3 |
| ETAA1 | 2p14 | 1 | 3.13 | 0 | 0.00 | 0.041 | 0.233 | (A) BRAF Class 1 v12 |
| FOXF1 | 16q24.1 | 1 | 3.13 | 0 | 0.00 | 0.041 | 0.233 | (A) BRAF Class 1 v12 |
| SLFN11 | 17q12 | 1 | 3.13 | 0 | 0.00 | 0.041 | 0.233 | (A) BRAF Class 1 v12 |
| CDKN2A | 9p21.3 | 450 | 13.73 | 118 | 16.32 | 0.0416 | 0.233 | (B) BRAF Class 3 |
| RMRP | 9p13.3 | 0 | 0.00 | 1 | 100.00 | 0.0417 | 0.233 | (B) BRAF Class 3 |
| MPL | 1p34.2 | 51 | 1.62 | 19 | 2.71 | 0.0417 | 0.233 | (B) BRAF Class 3 |
| AKT1 | 14q32.33 | 109 | 3.26 | 15 | 2.01 | 0.0419 | 0.233 | (A) BRAF Class 1 v12 |
| STAT4 | !q32.2-q32.3 | 23 | 3.39 | 10 | 7.09 | 0.0423 | 0.234 | (B) BRAF Class 3 |
| RPS6KB2 | 11q13.2 | 22 | 1.87 | 10 | 3.95 | 0.0425 | 0.234 | (B) BRAF Class 3 |
| CSDE1 | 1p13.2 | 10 | 1.19 | 6 | 3.43 | 0.0431 | 0.235 | (B) BRAF Class 3 |
| RUNX1 | 21q22.12 | 70 | 2.81 | 25 | 4.32 | 0.0431 | 0.235 | (B) BRAF Class 3 |
| STAT6 | 12q13.3 | 17 | 1.51 | 9 | 3.38 | 0.0442 | 0.239 | (B) BRAF Class 3 |
| NECTIN4 | 1q23.3 | 8 | 1.37 | 6 | 4.05 | 0.0444 | 0.239 | (B) BRAF Class 3 |
| LYN | 8q12.1 | 27 | 2.63 | 12 | 5.08 | 0.0449 | 0.239 | (B) BRAF Class 3 |
| MAP2K2 | 19p13.3 | 32 | 1.45 | 14 | 2.66 | 0.0452 | 0.239 | (B) BRAF Class 3 |
| PRKAR1A | 17q24.2 | 23 | 0.93 | 11 | 1.90 | 0.0452 | 0.239 | (B) BRAF Class 3 |
| ADAMTS20 | 12q12 | 3 | 2.56 | 3 | 14.29 | 0.0453 | 0.239 | (B) BRAF Class 3 |
| BCL2L12 | 19q13.33 | 11 | 1.12 | 7 | 2.93 | 0.0453 | 0.239 | (B) BRAF Class 3 |
| PREX2 | 8q13.2 | 139 | 11.99 | 42 | 16.15 | 0.0458 | 0.239 | (B) BRAF Class 3 |
| CDKN2B | 9p21.3 | 209 | 8.25 | 62 | 10.56 | 0.0459 | 0.239 | (B) BRAF Class 3 |
| NOTCH4 | 6p21.32 | 92 | 6.51 | 30 | 9.43 | 0.0459 | 0.239 | (B) BRAF Class 3 |
| DROSHA | 5p13.3 | 24 | 4.02 | 11 | 7.97 | 0.0466 | 0.241 | (B) BRAF Class 3 |
| E2F3 | 6p22.3 | 10 | 0.85 | 6 | 2.37 | 0.0479 | 0.246 | (B) BRAF Class 3 |
| KAT6A | 8p11.21 | 68 | 5.41 | 24 | 8.25 | 0.048 | 0.246 | (B) BRAF Class 3 |
| XRCC1 | 19q13.31 | 11 | 1.70 | 7 | 4.35 | 0.0484 | 0.247 | (B) BRAF Class 3 |
| PIK3C2B | 1q32.1 | 94 | 6.48 | 31 | 9.28 | 0.0491 | 0.249 | (B) BRAF Class 3 |
| SOX17 | 8q11.23 | 55 | 4.67 | 19 | 7.51 | 0.0495 | 0.25 | (B) BRAF Class 3 |
| COL7A1 | 3p21.31 | 94 | 16.10 | 33 | 22.30 | 0.0511 | 0.257 | (B) BRAF Class 3 |
| NR0B1 | Xp21.2 | 11 | 1.88 | 7 | 4.73 | 0.0518 | 0.259 | (B) BRAF Class 3 |
| CEBPA | 19q13.11 | 27 | 1.13 | 12 | 2.14 | 0.0521 | 0.26 | (B) BRAF Class 3 |
| RICTOR | 5p13.1 | 74 | 3.36 | 26 | 5.00 | 0.0526 | 0.261 | (B) BRAF Class 3 |
| BCL6 | 3q27.3 | 57 | 2.32 | 21 | 3.62 | 0.0547 | 0.27 | (B) BRAF Class 3 |
| DCC | 18q21.2 | 14 | 5.56 | 7 | 12.96 | 0.0558 | 0.274 | (B) BRAF Class 3 |
| PIK3CG | 7q22.3 | 75 | 4.88 | 25 | 7.23 | 0.0561 | 0.274 | (B) BRAF Class 3 |
| ERG | 21q22.2 | 63 | 2.96 | 23 | 4.50 | 0.0563 | 0.274 | (B) BRAF Class 3 |
| EXO1 | 1q43 | 14 | 2.40 | 8 | 5.41 | 0.0565 | 0.274 | (B) BRAF Class 3 |
| CUX1 | 7q22.1 | 117 | 8.19 | 35 | 11.22 | 0.057 | 0.275 | (B) BRAF Class 3 |
| DNMT3B | 20q11.21 | 34 | 2.89 | 13 | 5.14 | 0.0571 | 0.275 | (B) BRAF Class 3 |
| TENT5C | 1p12 | 37 | 1.51 | 15 | 2.59 | 0.0574 | 0.275 | (B) BRAF Class 3 |
| HGF | 7q21.11 | 62 | 3.97 | 22 | 6.06 | 0.0578 | 0.276 | (B) BRAF Class 3 |
| FANCF | 11p14.3 | 10 | 0.63 | 6 | 1.67 | 0.0582 | 0.276 | (B) BRAF Class 3 |
| INHBA | 7p14.1 | 43 | 2.96 | 16 | 4.91 | 0.0586 | 0.277 | (B) BRAF Class 3 |
| RIT1 | 1q22 | 25 | 1.28 | 11 | 2.43 | 0.0598 | 0.282 | (B) BRAF Class 3 |
| THBS1 | 15q14 | 1 | 0.85 | 2 | 9.52 | 0.0604 | 0.283 | (B) BRAF Class 3 |
| DOCK8 | 9p24.3 | 48 | 8.22 | 19 | 12.84 | 0.0607 | 0.283 | (B) BRAF Class 3 |
| ZNF521 | 18q11.2 | 1 | 0.82 | 2 | 9.09 | 0.061 | 0.284 | (B) BRAF Class 3 |

| | | | | | | | | |
|---|---|---|---|---|---|---|---|---|
| TSC1 | 9q34.13 | 81 | 3.18 | 27 | 4.58 | 0.0646 | 0.298 | (B) BRAF Class 3 |
| INPP4B | 4q31.21 | 32 | 2.08 | 13 | 3.65 | 0.0647 | 0.298 | (B) BRAF Class 3 |
| TOP1 | 20q12 | 27 | 1.68 | 11 | 3.12 | 0.0655 | 0.301 | (B) BRAF Class 3 |
| GNA11 | 19p13.3 | 23 | 0.71 | 10 | 1.37 | 0.0675 | 0.309 | (B) BRAF Class 3 |
| MUTYH | 1p34.1 | 45 | 1.83 | 17 | 2.93 | 0.068 | 0.309 | (B) BRAF Class 3 |
| H3F3A | 1q42.12 | 13 | 0.52 | 0 | 0.00 | 0.0681 | 0.309 | (A) BRAF Class 1 v12 |
| CDH1 | 16q22.1 | 74 | 2.36 | 24 | 3.44 | 0.0701 | 0.315 | (B) BRAF Class 3 |
| AMER1 | Xq11.2 | 64 | 4.09 | 22 | 6.06 | 0.0704 | 0.315 | (B) BRAF Class 3 |
| POLD1 | 19q13.33 | 131 | 6.37 | 22 | 4.51 | 0.0704 | 0.315 | (A) BRAF Class 1 v12 |
| PTPN14 | 1q32.3-q41 | 18 | 3.08 | 9 | 6.08 | 0.0741 | 0.328 | (B) BRAF Class 3 |
| DDX3X | Xp11.4 | 3 | 1.20 | 3 | 5.45 | 0.0743 | 0.328 | (B) BRAF Class 3 |
| EIF1AX | Xp22.12 | 21 | 1.53 | 1 | 0.33 | 0.0746 | 0.328 | (A) BRAF Class 1 v12 |
| NIPBL | 5p13.2 | 6 | 3.61 | 4 | 11.43 | 0.0747 | 0.328 | (B) BRAF Class 3 |
| ARID1B | 6q25.3 | 171 | 7.20 | 50 | 9.14 | 0.075 | 0.328 | (B) BRAF Class 3 |
| MLH1 | 3p22.2 | 53 | 1.65 | 18 | 2.55 | 0.0753 | 0.328 | (B) BRAF Class 3 |
| MAP3K4 | 6q26 | 30 | 9.01 | 1 | 2.08 | 0.0755 | 0.328 | (A) BRAF Class 1 v12 |
| KEL | 7q34 | 30 | 5.73 | 11 | 10.09 | 0.0756 | 0.328 | (B) BRAF Class 3 |
| TET1 | 10q21.3 | 118 | 6.40 | 36 | 8.51 | 0.0766 | 0.33 | (B) BRAF Class 3 |
| FGF3 | 11q13.3 | 27 | 1.65 | 11 | 2.96 | 0.0767 | 0.33 | (B) BRAF Class 3 |
| ALOX12B | 17p13.1 | 37 | 2.25 | 14 | 3.71 | 0.0775 | 0.33 | (B) BRAF Class 3 |
| ARHGAP35 | 19q13.32 | 31 | 4.25 | 13 | 7.18 | 0.0776 | 0.33 | (B) BRAF Class 3 |
| NSD1 | 5q35.3 | 126 | 6.04 | 39 | 7.93 | 0.078 | 0.33 | (B) BRAF Class 3 |
| KDM5C | Xp11.22 | 80 | 3.38 | 27 | 4.75 | 0.078 | 0.33 | (B) BRAF Class 3 |
| TFEB | 6p21.1 | 1 | 0.44 | 2 | 4.26 | 0.0782 | 0.33 | (B) BRAF Class 3 |
| ID3 | 1p36.12 | 8 | 0.35 | 5 | 0.95 | 0.0785 | 0.33 | (B) BRAF Class 3 |
| CDK12 | 17q12 | 71 | 3.22 | 24 | 4.62 | 0.0794 | 0.332 | (B) BRAF Class 3 |
| FAT1 | 4q35.2 | 263 | 12.90 | 74 | 15.48 | 0.0796 | 0.332 | (B) BRAF Class 3 |
| GATA3 | 10p14 | 71 | 2.89 | 24 | 4.14 | 0.0799 | 0.332 | (B) BRAF Class 3 |
| SNCAIP | 5q23.2 | 44 | 6.68 | 15 | 10.56 | 0.0801 | 0.332 | (B) BRAF Class 3 |
| MAML2 | 11q21 | 1 | 0.39 | 2 | 3.64 | 0.0814 | 0.335 | (B) BRAF Class 3 |
| PDGFB | 22q13.1 | 1 | 0.39 | 2 | 3.64 | 0.0814 | 0.335 | (B) BRAF Class 3 |
| TBX3 | 12q24.21 | 69 | 4.52 | 23 | 6.50 | 0.0816 | 0.335 | (B) BRAF Class 3 |
| GLI1 | 12q13.3 | 115 | 5.00 | 35 | 6.64 | 0.0824 | 0.336 | (B) BRAF Class 3 |
| MLH3 | 14q24.3 | 28 | 3.47 | 12 | 5.94 | 0.0832 | 0.338 | (B) BRAF Class 3 |
| WWTR1 | 3q25.1 | 3 | 0.50 | 3 | 2.16 | 0.0834 | 0.338 | (B) BRAF Class 3 |
| LZTR1 | 21\|22q11.1 | 25 | 3.88 | 9 | 7.20 | 0.084 | 0.339 | (B) BRAF Class 3 |
| CCNE1 | 19q12 | 44 | 1.73 | 16 | 2.72 | 0.0844 | 0.34 | (B) BRAF Class 3 |
| NPRL3 | 16p13.3 | 1 | 0.38 | 2 | 3.45 | 0.0853 | 0.341 | (B) BRAF Class 3 |
| TYK2 | 19p13.2 | 6 | 2.40 | 4 | 7.27 | 0.0854 | 0.341 | (B) BRAF Class 3 |
| AURKB | 17p13.1 | 25 | 1.02 | 2 | 0.34 | 0.0876 | 0.347 | (A) BRAF Class 1 v12 |
| CRTC1 | 19p13.11 | 37 | 3.64 | 14 | 5.86 | 0.0877 | 0.347 | (B) BRAF Class 3 |
| SESN3 | 11q21 | 8 | 1.52 | 5 | 3.94 | 0.0878 | 0.347 | (B) BRAF Class 3 |
| NSD3 | 8p11.23 | 30 | 2.67 | 13 | 4.44 | 0.088 | 0.347 | (B) BRAF Class 3 |
| USP6 | 17p13.2 | 2 | 1.18 | 2 | 7.41 | 0.0913 | 0.358 | (B) BRAF Class 3 |
| SOS1 | 2p22.1 | 38 | 3.11 | 15 | 4.87 | 0.0937 | 0.366 | (B) BRAF Class 3 |
| MTOR | 1p36.22 | 144 | 5.52 | 43 | 7.01 | 0.0939 | 0.366 | (B) BRAF Class 3 |
| EP400 | 12q24.33 | 5 | 4.17 | 3 | 14.29 | 0.0975 | 0.376 | (B) BRAF Class 3 |
| HDAC9 | 7p21.1 | 3 | 3.49 | 3 | 13.64 | 0.0976 | 0.376 | (B) BRAF Class 3 |

| | | | | | | | | |
|---|---|---|---|---|---|---|---|---|
| MAPK3 | 16p11.2 | 20 | 1.80 | 1 | 0.43 | 0.0976 | 0.376 | (A) BRAF Class 1 v12 |
| POLQ | 3q13.33 | 53 | 8.18 | 20 | 11.76 | 0.0978 | 0.376 | (B) BRAF Class 3 |
| EXT1 | 8q24.11 | 21 | 2.05 | 9 | 3.73 | 0.0987 | 0.378 | (B) BRAF Class 3 |
| IGF2R | 6q25.3 | 8 | 2.39 | 5 | 5.88 | 0.0999 | 0.381 | (B) BRAF Class 3 |
| TMPRSS2 | 21q22.3 | 40 | 1.90 | 15 | 2.94 | 0.102 | 0.386 | (B) BRAF Class 3 |
| CASP8 | 2q33.1 | 91 | 4.46 | 15 | 3.07 | 0.102 | 0.386 | (A) BRAF Class 1 v12 |
| RAD50 | 5q31.1 | 94 | 4.32 | 15 | 2.98 | 0.104 | 0.393 | (A) BRAF Class 1 v12 |
| SUFU | 10q24.32 | 45 | 1.83 | 16 | 2.76 | 0.105 | 0.393 | (B) BRAF Class 3 |
| ESPL1 | 12q13.13 | 1 | 1.16 | 2 | 9.09 | 0.105 | 0.393 | (B) BRAF Class 3 |
| IKZF2 | 2q34 | 4 | 1.81 | 3 | 6.38 | 0.106 | 0.394 | (B) BRAF Class 3 |
| AIP | 11q13.2 | 0 | 0.00 | 1 | 9.09 | 0.106 | 0.394 | (B) BRAF Class 3 |
| HOXB13 | 17q21.32 | 18 | 0.94 | 8 | 1.77 | 0.106 | 0.395 | (B) BRAF Class 3 |
| COL1A1 | 17q21.33 | 4 | 1.56 | 3 | 5.45 | 0.107 | 0.397 | (B) BRAF Class 3 |
| XRCC6 | 22q13.2 | 11 | 1.88 | 6 | 4.05 | 0.108 | 0.397 | (B) BRAF Class 3 |
| MALT1 | 18q21.32 | 17 | 1.40 | 7 | 2.73 | 0.108 | 0.398 | (B) BRAF Class 3 |
| ACVR1B | 12q13.13 | 10 | 1.52 | 5 | 3.52 | 0.109 | 0.399 | (B) BRAF Class 3 |
| MEF2B | 19p13.11 | 28 | 1.14 | 11 | 1.90 | 0.109 | 0.399 | (B) BRAF Class 3 |
| AFF1 | ₄q21.3-q22.₁ | 2 | 1.64 | 2 | 9.09 | 0.111 | 0.403 | (B) BRAF Class 3 |
| AXIN2 | 17q24.1 | 83 | 4.28 | 13 | 2.91 | 0.113 | 0.408 | (A) BRAF Class 1 v12 |
| NUF2 | 1q23.3 | 10 | 1.90 | 0 | 0.00 | 0.113 | 0.409 | (A) BRAF Class 1 v12 |
| TGFBR2 | 3p24.1 | 112 | 7.15 | 19 | 5.23 | 0.114 | 0.413 | (A) BRAF Class 1 v12 |
| NT5C2 | q24.32-q24. | 11 | 1.33 | 6 | 2.79 | 0.116 | 0.415 | (B) BRAF Class 3 |
| CIITA | 16p13.13 | 50 | 5.02 | 17 | 7.26 | 0.116 | 0.415 | (B) BRAF Class 3 |
| FGA | 4q31.3 | 0 | 0.00 | 1 | 20.00 | 0.116 | 0.415 | (B) BRAF Class 3 |
| EPAS1 | 2p21 | 12 | 2.28 | 6 | 4.72 | 0.117 | 0.416 | (B) BRAF Class 3 |
| ESR2 | 4q23.2-q23. | 1 | 0.39 | 2 | 2.82 | 0.118 | 0.419 | (B) BRAF Class 3 |
| CDK6 | 7q21.2 | 19 | 0.75 | 8 | 1.36 | 0.119 | 0.42 | (B) BRAF Class 3 |
| MYB | 6q23.3 | 19 | 1.20 | 8 | 2.18 | 0.12 | 0.421 | (B) BRAF Class 3 |
| TP53BP1 | 15q15.3 | 58 | 4.92 | 21 | 6.82 | 0.12 | 0.421 | (B) BRAF Class 3 |
| ADGRB3 | 6q12-q13 | 3 | 5.56 | 2 | 25.00 | 0.12 | 0.421 | (B) BRAF Class 3 |
| EFNB3 | 17p13.1 | 0 | 0.00 | 1 | 16.67 | 0.122 | 0.428 | (B) BRAF Class 3 |
| RHOT1 | 17q11.2 | 6 | 1.03 | 4 | 2.70 | 0.123 | 0.429 | (B) BRAF Class 3 |
| NCOA3 | 20q13.12 | 52 | 3.98 | 16 | 5.73 | 0.126 | 0.436 | (B) BRAF Class 3 |
| PRKCZ | 1p36.33 | 5 | 1.41 | 3 | 4.35 | 0.126 | 0.436 | (B) BRAF Class 3 |
| MAGI2 | 7q21.11 | 38 | 7.69 | 11 | 11.96 | 0.127 | 0.436 | (B) BRAF Class 3 |
| POLB | 8p11.21 | 9 | 1.54 | 5 | 3.38 | 0.133 | 0.454 | (B) BRAF Class 3 |
| SLC25A13 | 7q21.3 | 9 | 1.54 | 5 | 3.38 | 0.133 | 0.454 | (B) BRAF Class 3 |
| MCM8 | 20p12.3 | 15 | 2.57 | 7 | 4.73 | 0.136 | 0.463 | (B) BRAF Class 3 |
| PARP1 | 1q42.12 | 44 | 3.13 | 6 | 1.82 | 0.136 | 0.463 | (A) BRAF Class 1 v12 |
| PTCH2 | 1p34.1 | 6 | 2.50 | 4 | 6.25 | 0.137 | 0.463 | (B) BRAF Class 3 |
| CDK8 | 13q12.13 | 36 | 1.69 | 13 | 2.54 | 0.137 | 0.463 | (B) BRAF Class 3 |
| CUL3 | 2q36.2 | 20 | 1.31 | 8 | 2.25 | 0.14 | 0.471 | (B) BRAF Class 3 |
| TTYH1 | 19q13.42 | 2 | 1.27 | 2 | 6.06 | 0.14 | 0.471 | (B) BRAF Class 3 |
| BOD1L1 | 4p15.33 | 0 | 0.00 | 1 | 12.50 | 0.143 | 0.474 | (B) BRAF Class 3 |
| PMS2 | 7p22.1 | 53 | 2.14 | 17 | 3.00 | 0.143 | 0.474 | (B) BRAF Class 3 |
| SLX4 | 16p13.3 | 97 | 6.56 | 16 | 4.82 | 0.143 | 0.474 | (A) BRAF Class 1 v12 |
| CDH23 | 10q22.1 | 3 | 6.25 | 2 | 25.00 | 0.144 | 0.474 | (B) BRAF Class 3 |
| CHEK2 | 22q12.1 | 60 | 2.36 | 19 | 3.23 | 0.146 | 0.474 | (B) BRAF Class 3 |

| | | | | | | | | |
|---|---|---|---|---|---|---|---|---|
| MST1R | 3p21.31 | 46 | 3.58 | 15 | 5.10 | 0.146 | 0.474 | (B) BRAF Class 3 |
| RYBP | 3p13 | 8 | 0.68 | 4 | 1.58 | 0.147 | 0.474 | (B) BRAF Class 3 |
| STAG1 | 3q22.3 | 28 | 4.14 | 2 | 1.68 | 0.148 | 0.474 | (A) BRAF Class 1 v12 |
| PAXIP1 | 7q36.2 | 25 | 4.28 | 10 | 6.76 | 0.148 | 0.474 | (B) BRAF Class 3 |
| ERCC3 | 2q14.3 | 39 | 1.80 | 13 | 2.64 | 0.148 | 0.474 | (B) BRAF Class 3 |
| ELF3 | 1q32.1 | 10 | 1.51 | 5 | 3.13 | 0.149 | 0.474 | (B) BRAF Class 3 |
| FLI1 | 11q24.3 | 7 | 1.64 | 3 | 4.35 | 0.15 | 0.474 | (B) BRAF Class 3 |
| CDK4 | 12q14.1 | 43 | 1.69 | 6 | 1.01 | 0.151 | 0.474 | (A) BRAF Class 1 v12 |
| RECQL4 | 8q24.3 | 120 | 5.58 | 34 | 6.91 | 0.151 | 0.474 | (B) BRAF Class 3 |
| G6PD | Xq28 | 0 | 0.00 | 1 | 4.76 | 0.152 | 0.474 | (B) BRAF Class 3 |
| MAGEA1 | Xq28 | 0 | 0.00 | 1 | 4.76 | 0.152 | 0.474 | (B) BRAF Class 3 |
| MAGI1 | 3p14.1 | 0 | 0.00 | 1 | 4.76 | 0.152 | 0.474 | (B) BRAF Class 3 |
| MMP2 | 16q12.2 | 0 | 0.00 | 1 | 4.76 | 0.152 | 0.474 | (B) BRAF Class 3 |
| USP9X | Xp11.4 | 0 | 0.00 | 1 | 4.76 | 0.152 | 0.474 | (B) BRAF Class 3 |
| AKAP9 | 7q21.2 | 0 | 0.00 | 1 | 4.55 | 0.153 | 0.474 | (B) BRAF Class 3 |
| BCL3 | 19q13.32 | 0 | 0.00 | 1 | 4.55 | 0.153 | 0.474 | (B) BRAF Class 3 |
| HOOK3 | 8p11.21 | 0 | 0.00 | 1 | 4.55 | 0.153 | 0.474 | (B) BRAF Class 3 |
| LPP | 3q27.3-q28 | 0 | 0.00 | 1 | 4.55 | 0.153 | 0.474 | (B) BRAF Class 3 |
| NUMA1 | 11q13.4 | 0 | 0.00 | 1 | 4.55 | 0.153 | 0.474 | (B) BRAF Class 3 |
| PBX1 | 1q23.3 | 0 | 0.00 | 1 | 4.55 | 0.153 | 0.474 | (B) BRAF Class 3 |
| RALGDS | q34.13-q34. | 0 | 0.00 | 1 | 4.55 | 0.153 | 0.474 | (B) BRAF Class 3 |
| RTEL1 | 20q13.33 | 20 | 3.78 | 8 | 6.30 | 0.154 | 0.474 | (B) BRAF Class 3 |
| KLF6 | 10p15.2 | 0 | 0.00 | 1 | 4.35 | 0.154 | 0.474 | (B) BRAF Class 3 |
| BTG1 | 12q21.33 | 2 | 0.30 | 2 | 1.33 | 0.154 | 0.474 | (B) BRAF Class 3 |
| NUP98 | 11p15.4 | 0 | 0.00 | 1 | 4.00 | 0.159 | 0.474 | (B) BRAF Class 3 |
| ZMYM2 | 13q12.11 | 0 | 0.00 | 1 | 4.55 | 0.159 | 0.474 | (B) BRAF Class 3 |
| EPHB1 | 3q22.2 | 65 | 4.15 | 20 | 5.51 | 0.16 | 0.474 | (B) BRAF Class 3 |
| LMO3 | 12p12.3 | 0 | 0.00 | 1 | 1.45 | 0.163 | 0.474 | (B) BRAF Class 3 |
| RSPO2 | 8q23.1 | 12 | 1.34 | 6 | 2.49 | 0.163 | 0.474 | (B) BRAF Class 3 |
| PDGFRB | 5q32 | 92 | 3.62 | 27 | 4.58 | 0.164 | 0.474 | (B) BRAF Class 3 |
| SH2D1A | Xq25 | 11 | 0.60 | 5 | 1.18 | 0.164 | 0.474 | (B) BRAF Class 3 |
| NFKBIE | 6p21.1 | 13 | 1.81 | 6 | 3.31 | 0.164 | 0.474 | (B) BRAF Class 3 |
| CALR | 19p13.13 | 26 | 1.34 | 3 | 0.66 | 0.167 | 0.474 | (A) BRAF Class 1 v12 |
| ARID5B | 10q21.2 | 50 | 3.80 | 15 | 5.24 | 0.168 | 0.474 | (B) BRAF Class 3 |
| CDKN2C | 1p32.3 | 14 | 0.57 | 6 | 1.03 | 0.169 | 0.474 | (B) BRAF Class 3 |
| FAAP100 | 17q25.3 | 24 | 4.11 | 3 | 2.03 | 0.17 | 0.474 | (A) BRAF Class 1 v12 |
| CREBBP | 16p13.3 | 192 | 7.56 | 52 | 8.83 | 0.171 | 0.474 | (B) BRAF Class 3 |
| PPARG | 3p25.2 | 39 | 2.20 | 13 | 3.13 | 0.172 | 0.474 | (B) BRAF Class 3 |
| SLC34A2 | 4p15.2 | 28 | 4.11 | 4 | 2.25 | 0.173 | 0.474 | (A) BRAF Class 1 v12 |
| PRKACA | 19p13.12 | 0 | 0.00 | 1 | 2.17 | 0.173 | 0.474 | (B) BRAF Class 3 |
| HIST1H2BD | 6p22.2 | 9 | 0.76 | 0 | 0.00 | 0.173 | 0.474 | (A) BRAF Class 1 v12 |
| ASH2L | 8p11.23 | 0 | 0.00 | 1 | 3.03 | 0.174 | 0.474 | (B) BRAF Class 3 |
| CRCT1 | 1q21.3 | 0 | 0.00 | 1 | 3.03 | 0.174 | 0.474 | (B) BRAF Class 3 |
| LEF1 | 4q25 | 0 | 0.00 | 1 | 2.13 | 0.175 | 0.474 | (B) BRAF Class 3 |
| BCL2L1 | 20q11.21 | 11 | 0.46 | 5 | 0.88 | 0.176 | 0.474 | (B) BRAF Class 3 |
| EPHA5 | 4q13.1-q13.2 | 89 | 4.78 | 25 | 6.02 | 0.176 | 0.474 | (B) BRAF Class 3 |
| EPCAM | 2p21 | 18 | 0.91 | 7 | 1.52 | 0.176 | 0.474 | (B) BRAF Class 3 |
| MSH2 | 2p21-p16.3 | 73 | 2.96 | 22 | 3.81 | 0.177 | 0.474 | (B) BRAF Class 3 |

| | | | | | | | | |
|---|---|---|---|---|---|---|---|---|
| PTEN | 10q23.31 | 352 | 10.51 | 69 | 9.29 | 0.177 | 0.474 | (A) BRAF Class 1 v12 |
| TCF7L2 | 0q25.2-q25. | 76 | 3.40 | 22 | 4.36 | 0.178 | 0.474 | (B) BRAF Class 3 |
| MST1 | 3p21.31 | 9 | 0.81 | 0 | 0.00 | 0.179 | 0.474 | (A) BRAF Class 1 v12 |
| DHX15 | 4p15.2 | 14 | 4.20 | 4 | 8.33 | 0.179 | 0.474 | (B) BRAF Class 3 |
| FOXL2 | 3q22.3 | 46 | 2.08 | 7 | 1.35 | 0.18 | 0.474 | (A) BRAF Class 1 v12 |
| INSIG1 | 7q36.3 | 0 | 0.00 | 1 | 1.72 | 0.181 | 0.474 | (B) BRAF Class 3 |
| MGA | 15q15.1 | 180 | 9.95 | 48 | 11.59 | 0.182 | 0.474 | (B) BRAF Class 3 |
| MAP3K9 | 14q24.2 | 9 | 4.86 | 4 | 10.00 | 0.182 | 0.474 | (B) BRAF Class 3 |
| ACP2 | 11.2\|11p12- | 0 | 0.00 | 1 | 0.13 | 0.183 | 0.474 | (B) BRAF Class 3 |
| ARHGEF38 | 4q24 | 0 | 0.00 | 1 | 0.13 | 0.183 | 0.474 | (B) BRAF Class 3 |
| C1ORF147 | 1q32.1 | 0 | 0.00 | 1 | 0.13 | 0.183 | 0.474 | (B) BRAF Class 3 |
| CACNA1C | 12p13.33 | 0 | 0.00 | 1 | 0.13 | 0.183 | 0.474 | (B) BRAF Class 3 |
| CCHCR1 | 6p21.33 | 0 | 0.00 | 1 | 0.13 | 0.183 | 0.474 | (B) BRAF Class 3 |
| DAP | 5p15.2 | 0 | 0.00 | 1 | 0.13 | 0.183 | 0.474 | (B) BRAF Class 3 |
| DOCK8-AS1 | 9p24.3 | 0 | 0.00 | 1 | 0.13 | 0.183 | 0.474 | (B) BRAF Class 3 |
| EIF2S3 | Xp22.11 | 0 | 0.00 | 1 | 0.13 | 0.183 | 0.474 | (B) BRAF Class 3 |
| ENSA | 1q21.3 | 0 | 0.00 | 1 | 0.13 | 0.183 | 0.474 | (B) BRAF Class 3 |
| FNDC3B | 3q26.31 | 0 | 0.00 | 1 | 0.13 | 0.183 | 0.474 | (B) BRAF Class 3 |
| GON4L | 1q22 | 0 | 0.00 | 1 | 0.13 | 0.183 | 0.474 | (B) BRAF Class 3 |
| GPD1 | 12q13.12 | 0 | 0.00 | 1 | 0.13 | 0.183 | 0.474 | (B) BRAF Class 3 |
| HIPK2 | 7q34 | 0 | 0.00 | 1 | 0.13 | 0.183 | 0.474 | (B) BRAF Class 3 |
| IGLJ5 | 22q11.22 | 0 | 0.00 | 1 | 0.13 | 0.183 | 0.474 | (B) BRAF Class 3 |
| INHBE | 12q13.3 | 0 | 0.00 | 1 | 0.13 | 0.183 | 0.474 | (B) BRAF Class 3 |
| LINC00996 | 7q36.1 | 0 | 0.00 | 1 | 0.13 | 0.183 | 0.474 | (B) BRAF Class 3 |
| LINC01514 | 10q24.31 | 0 | 0.00 | 1 | 0.13 | 0.183 | 0.474 | (B) BRAF Class 3 |
| MIR99AHG | 21q21.1 | 0 | 0.00 | 1 | 0.13 | 0.183 | 0.474 | (B) BRAF Class 3 |
| NSRP1 | 17q11.2 | 0 | 0.00 | 1 | 0.13 | 0.183 | 0.474 | (B) BRAF Class 3 |
| PAX8-AS1 | 2q14.1 | 0 | 0.00 | 1 | 0.13 | 0.183 | 0.474 | (B) BRAF Class 3 |
| PCMT1 | 6q25.1 | 0 | 0.00 | 1 | 0.13 | 0.183 | 0.474 | (B) BRAF Class 3 |
| PDE4D | ;q11.2-q12.: | 0 | 0.00 | 1 | 0.13 | 0.183 | 0.474 | (B) BRAF Class 3 |
| PTTG1IP | 21q22.3 | 0 | 0.00 | 1 | 0.13 | 0.183 | 0.474 | (B) BRAF Class 3 |
| RAB11FIP4 | 17q11.2 | 0 | 0.00 | 1 | 0.13 | 0.183 | 0.474 | (B) BRAF Class 3 |
| RLF | 1p34.2 | 0 | 0.00 | 1 | 0.13 | 0.183 | 0.474 | (B) BRAF Class 3 |
| SDHAP3 | 5p15.33 | 0 | 0.00 | 1 | 0.13 | 0.183 | 0.474 | (B) BRAF Class 3 |
| SEPTIN14 | 7p11.2 | 0 | 0.00 | 1 | 0.13 | 0.183 | 0.474 | (B) BRAF Class 3 |
| SNTG1 | 8q11.21 | 0 | 0.00 | 1 | 0.13 | 0.183 | 0.474 | (B) BRAF Class 3 |
| TANC2 | 7q23.2-q23. | 0 | 0.00 | 1 | 0.13 | 0.183 | 0.474 | (B) BRAF Class 3 |
| TMIE | 3p21.31 | 0 | 0.00 | 1 | 0.13 | 0.183 | 0.474 | (B) BRAF Class 3 |
| TRAJ22 | 14q11.2 | 0 | 0.00 | 1 | 0.13 | 0.183 | 0.474 | (B) BRAF Class 3 |
| TTF1 | 9q34.13 | 0 | 0.00 | 1 | 0.13 | 0.183 | 0.474 | (B) BRAF Class 3 |
| TTF2 | 1p13.1 | 0 | 0.00 | 1 | 0.13 | 0.183 | 0.474 | (B) BRAF Class 3 |
| UAP1L1 | 9q34.3 | 0 | 0.00 | 1 | 0.13 | 0.183 | 0.474 | (B) BRAF Class 3 |
| ZFP30 | 19q13.12 | 0 | 0.00 | 1 | 0.13 | 0.183 | 0.474 | (B) BRAF Class 3 |
| ZNF135 | 19q13.43 | 0 | 0.00 | 1 | 0.13 | 0.183 | 0.474 | (B) BRAF Class 3 |
| ZNF284 | 19q13.31 | 0 | 0.00 | 1 | 0.13 | 0.183 | 0.474 | (B) BRAF Class 3 |
| TSHZ3 | 19q12 | 2 | 2.33 | 2 | 9.09 | 0.184 | 0.474 | (B) BRAF Class 3 |
| CDC73 | 1q31.2 | 43 | 1.75 | 14 | 2.41 | 0.184 | 0.474 | (B) BRAF Class 3 |
| SPRED1 | 15q14 | 11 | 1.66 | 5 | 3.13 | 0.185 | 0.476 | (B) BRAF Class 3 |

| | | | | | | | | |
|---|---|---|---|---|---|---|---|---|
| MED12 | Xq13.1 | 118 | 5.36 | 34 | 6.46 | 0.187 | 0.48 | (B) BRAF Class 3 |
| BIRC2 | 11q22.2 | 0 | 0.00 | 1 | 3.70 | 0.188 | 0.48 | (B) BRAF Class 3 |
| ZNF708 | 19p12 | 3 | 0.85 | 2 | 2.90 | 0.189 | 0.482 | (B) BRAF Class 3 |
| SH2B3 | 12q24.12 | 61 | 2.60 | 10 | 1.83 | 0.189 | 0.482 | (A) BRAF Class 1 v12 |
| SMAD4 | 18q21.2 | 230 | 7.01 | 58 | 8.02 | 0.19 | 0.482 | (B) BRAF Class 3 |
| MAPKAP1 | 9q33.3 | 5 | 0.95 | 3 | 2.36 | 0.19 | 0.482 | (B) BRAF Class 3 |
| RPTOR | 17q25.3 | 87 | 4.10 | 26 | 5.09 | 0.191 | 0.484 | (B) BRAF Class 3 |
| GAB2 | 11q14.1 | 3 | 1.80 | 2 | 6.06 | 0.191 | 0.484 | (B) BRAF Class 3 |
| XPO1 | 2p15 | 48 | 1.89 | 15 | 2.55 | 0.193 | 0.486 | (B) BRAF Class 3 |
| HABP2 | 10q25.3 | 17 | 2.91 | 7 | 4.73 | 0.193 | 0.486 | (B) BRAF Class 3 |
| MAPK1 | 22q11.22 | 19 | 0.79 | 7 | 1.27 | 0.194 | 0.486 | (B) BRAF Class 3 |
| RAC1 | 7p22.1 | 51 | 2.22 | 16 | 2.95 | 0.195 | 0.488 | (B) BRAF Class 3 |
| ERCC5 | 13q33.1 | 73 | 3.36 | 21 | 4.27 | 0.196 | 0.489 | (B) BRAF Class 3 |
| AXIN1 | 16p13.3 | 56 | 3.69 | 9 | 2.58 | 0.196 | 0.489 | (A) BRAF Class 1 v12 |
| TAL1 | 1p33 | 11 | 1.61 | 5 | 2.94 | 0.197 | 0.489 | (B) BRAF Class 3 |
| FGF23 | 12p13.32 | 9 | 1.27 | 4 | 2.53 | 0.199 | 0.489 | (B) BRAF Class 3 |
| NCOA2 | 8q13.3 | 6 | 2.33 | 3 | 5.45 | 0.199 | 0.489 | (B) BRAF Class 3 |
| MAP2K1 | 15q22.31 | 71 | 2.63 | 21 | 3.33 | 0.2 | 0.489 | (B) BRAF Class 3 |
| CD79B | 17q23.3 | 15 | 0.61 | 6 | 1.03 | 0.201 | 0.489 | (B) BRAF Class 3 |
| EML4 | 2p21 | 13 | 3.00 | 4 | 5.71 | 0.201 | 0.489 | (B) BRAF Class 3 |
| DDX31 | 9q34.13 | 1 | 1.08 | 1 | 9.09 | 0.201 | 0.489 | (B) BRAF Class 3 |
| FRK | 6q22.1 | 1 | 1.08 | 1 | 9.09 | 0.201 | 0.489 | (B) BRAF Class 3 |
| KRIT1 | 7q21.2 | 1 | 1.08 | 1 | 9.09 | 0.201 | 0.489 | (B) BRAF Class 3 |
| FANCG | 9p13.3 | 26 | 1.61 | 9 | 2.39 | 0.201 | 0.489 | (B) BRAF Class 3 |
| ATR | 3q23 | 125 | 5.66 | 35 | 6.73 | 0.202 | 0.489 | (B) BRAF Class 3 |
| UIMC1 | 5q35.2 | 14 | 2.31 | 6 | 3.90 | 0.203 | 0.489 | (B) BRAF Class 3 |
| TEX12 | 11q23.1 | 0 | 0.00 | 1 | 1.06 | 0.203 | 0.489 | (B) BRAF Class 3 |
| HSPA2 | 14q23.3 | 0 | 0.00 | 1 | 4.55 | 0.204 | 0.489 | (B) BRAF Class 3 |
| SOX2 | 3q26.33 | 33 | 1.34 | 11 | 1.90 | 0.204 | 0.489 | (B) BRAF Class 3 |
| SRY | Yp11.2 | 0 | 0.00 | 1 | 0.65 | 0.204 | 0.489 | (B) BRAF Class 3 |
| NSD2 | 4p16.3 | 44 | 3.24 | 15 | 4.31 | 0.204 | 0.489 | (B) BRAF Class 3 |
| BTK | Xq22.1 | 27 | 1.65 | 9 | 2.43 | 0.205 | 0.489 | (B) BRAF Class 3 |
| ETV6 | 12p13.2 | 51 | 2.09 | 16 | 2.75 | 0.205 | 0.489 | (B) BRAF Class 3 |
| PPP6C | 9q33.3 | 43 | 3.39 | 6 | 2.19 | 0.205 | 0.489 | (A) BRAF Class 1 v12 |
| ANKRD11 | 16q24.3 | 87 | 7.84 | 14 | 6.01 | 0.206 | 0.489 | (A) BRAF Class 1 v12 |
| BRCA2 | 13q13.1 | 203 | 8.00 | 54 | 9.12 | 0.206 | 0.489 | (B) BRAF Class 3 |
| RAD54L | 1p34.1 | 19 | 1.45 | 2 | 0.65 | 0.207 | 0.491 | (A) BRAF Class 1 v12 |
| UPF1 | 19p13.11 | 15 | 2.85 | 6 | 4.72 | 0.208 | 0.491 | (B) BRAF Class 3 |
| SOS2 | 14q21.3 | 3 | 1.91 | 2 | 6.06 | 0.208 | 0.492 | (B) BRAF Class 3 |
| ETV4 | 17q21.31 | 19 | 1.30 | 2 | 0.58 | 0.209 | 0.493 | (A) BRAF Class 1 v12 |
| PRSS8 | 16p11.2 | 7 | 1.36 | 3 | 3.03 | 0.21 | 0.494 | (B) BRAF Class 3 |
| WT1 | 11p13 | 68 | 2.73 | 12 | 2.03 | 0.21 | 0.494 | (A) BRAF Class 1 v12 |
| CD70 | 19p13.3 | 0 | 0.00 | 1 | 3.23 | 0.211 | 0.494 | (B) BRAF Class 3 |
| MLLT3 | 9p21.3 | 9 | 2.47 | 3 | 5.26 | 0.213 | 0.498 | (B) BRAF Class 3 |
| SDHA | 5p15.33 | 71 | 3.11 | 21 | 3.89 | 0.215 | 0.499 | (B) BRAF Class 3 |
| ASXL2 | 2p23.3 | 58 | 4.41 | 9 | 3.15 | 0.215 | 0.499 | (A) BRAF Class 1 v12 |
| CSF3R | 1p34.3 | 52 | 2.35 | 16 | 3.05 | 0.216 | 0.501 | (B) BRAF Class 3 |
| RPS6KB1 | 17q23.1 | 5 | 1.50 | 2 | 4.17 | 0.216 | 0.501 | (B) BRAF Class 3 |

| | | | | | | | | |
|---|---|---|---|---|---|---|---|---|
| INSRR | 1q23.1 | 6 | 0.18 | 3 | 0.40 | 0.217 | 0.502 | (B) BRAF Class 3 |
| CRLF2 | 2.33 and Yp1 | 31 | 1.31 | 10 | 1.85 | 0.222 | 0.511 | (B) BRAF Class 3 |
| ADGRA2 | 8p11.23 | 35 | 5.63 | 10 | 7.81 | 0.223 | 0.513 | (B) BRAF Class 3 |
| GNA13 | 17q24.1 | 3 | 0.42 | 2 | 1.26 | 0.224 | 0.513 | (B) BRAF Class 3 |
| FANCM | 14q21.2 | 40 | 5.34 | 14 | 7.04 | 0.224 | 0.513 | (B) BRAF Class 3 |
| ABL1 | 9q34.12 | 88 | 2.80 | 24 | 3.41 | 0.224 | 0.513 | (B) BRAF Class 3 |
| FGF4 | 11q13.3 | 29 | 1.86 | 4 | 1.10 | 0.225 | 0.513 | (A) BRAF Class 1 v12 |
| HELQ | 4q21.23 | 17 | 2.91 | 2 | 1.35 | 0.226 | 0.513 | (A) BRAF Class 1 v12 |
| DIS3 | 13q21.33 | 71 | 3.00 | 21 | 3.71 | 0.226 | 0.513 | (B) BRAF Class 3 |
| GRM8 | 7q31.33 | 4 | 3.42 | 2 | 9.52 | 0.226 | 0.513 | (B) BRAF Class 3 |
| SLITRK6 | 13q31.1 | 12 | 3.67 | 5 | 6.25 | 0.227 | 0.513 | (B) BRAF Class 3 |
| AR | Xq12 | 120 | 4.73 | 33 | 5.55 | 0.23 | 0.52 | (B) BRAF Class 3 |
| ERF | 19q13.2 | 9 | 1.71 | 4 | 3.15 | 0.234 | 0.528 | (B) BRAF Class 3 |
| CHEK1 | 11q24.2 | 27 | 1.22 | 9 | 1.73 | 0.236 | 0.53 | (B) BRAF Class 3 |
| FYN | 6q21 | 26 | 2.07 | 8 | 2.99 | 0.237 | 0.531 | (B) BRAF Class 3 |
| PIK3CD | 1p36.22 | 38 | 2.99 | 11 | 4.01 | 0.238 | 0.533 | (B) BRAF Class 3 |
| EP300 | 22q13.2 | 172 | 6.99 | 46 | 7.93 | 0.24 | 0.535 | (B) BRAF Class 3 |
| POLE | 12q24.33 | 161 | 7.29 | 43 | 8.30 | 0.24 | 0.535 | (B) BRAF Class 3 |
| IKZF3 | 17q12-q21.1 | 7 | 1.26 | 3 | 2.59 | 0.241 | 0.536 | (B) BRAF Class 3 |
| IDH1 | 2q34 | 80 | 2.39 | 14 | 1.88 | 0.241 | 0.536 | (A) BRAF Class 1 v12 |
| DOT1L | 19p13.3 | 78 | 5.00 | 22 | 6.06 | 0.242 | 0.536 | (B) BRAF Class 3 |
| CD274 | 9p24.1 | 20 | 0.83 | 7 | 1.24 | 0.242 | 0.536 | (B) BRAF Class 3 |
| DEPDC5 | 2q12.2-q12. | 13 | 4.94 | 1 | 1.72 | 0.245 | 0.54 | (A) BRAF Class 1 v12 |
| HIST1H1C | 6p22.2 | 25 | 2.00 | 3 | 1.12 | 0.245 | 0.54 | (A) BRAF Class 1 v12 |
| PHOX2B | 4p13 | 48 | 2.08 | 14 | 2.67 | 0.246 | 0.542 | (B) BRAF Class 3 |
| GATA4 | 8p23.1 | 32 | 2.26 | 10 | 3.07 | 0.247 | 0.543 | (B) BRAF Class 3 |
| WRN | 8p12 | 54 | 4.66 | 16 | 5.84 | 0.249 | 0.545 | (B) BRAF Class 3 |
| NUTM1 | 15q14 | 29 | 5.05 | 8 | 7.08 | 0.251 | 0.549 | (B) BRAF Class 3 |
| CDC42 | 1p36.12 | 3 | 0.45 | 2 | 1.25 | 0.252 | 0.549 | (B) BRAF Class 3 |
| PTCH1 | 9q22.32 | 132 | 5.20 | 26 | 4.41 | 0.252 | 0.549 | (A) BRAF Class 1 v12 |
| JUN | 1p32.1 | 23 | 1.61 | 3 | 0.91 | 0.253 | 0.55 | (A) BRAF Class 1 v12 |
| HSP90AA1 | 14q32.31 | 8 | 1.50 | 3 | 2.94 | 0.254 | 0.55 | (B) BRAF Class 3 |
| MYCN | 2p24.3 | 43 | 1.69 | 13 | 2.18 | 0.254 | 0.55 | (B) BRAF Class 3 |
| PRAME | 22q11.22 | 4 | 1.13 | 2 | 2.90 | 0.254 | 0.55 | (B) BRAF Class 3 |
| GSK3B | 3q13.33 | 17 | 1.09 | 6 | 1.65 | 0.257 | 0.554 | (B) BRAF Class 3 |
| TOP2A | 17q21.2 | 32 | 5.09 | 4 | 3.20 | 0.259 | 0.558 | (A) BRAF Class 1 v12 |
| FUS | 16p11.2 | 16 | 1.51 | 6 | 2.28 | 0.259 | 0.558 | (B) BRAF Class 3 |
| PTK2 | 8q24.3 | 8 | 2.26 | 3 | 4.35 | 0.261 | 0.559 | (B) BRAF Class 3 |
| KCNQ1 | 1p15.5-p15. | 26 | 4.45 | 9 | 6.08 | 0.262 | 0.56 | (B) BRAF Class 3 |
| HIST1H3D | 6p22.2 | 7 | 0.63 | 0 | 0.00 | 0.262 | 0.56 | (A) BRAF Class 1 v12 |
| FGF12 | 3q28-q29 | 0 | 0.00 | 1 | 1.82 | 0.263 | 0.56 | (B) BRAF Class 3 |
| TIPARP | 3q25.31 | 0 | 0.00 | 1 | 1.82 | 0.263 | 0.56 | (B) BRAF Class 3 |
| IRF4 | 6p25.3 | 21 | 1.34 | 7 | 1.93 | 0.264 | 0.561 | (B) BRAF Class 3 |
| PPP2R2A | 8p21.2 | 8 | 2.20 | 0 | 0.00 | 0.265 | 0.561 | (A) BRAF Class 1 v12 |
| GPS2 | 17p13.1 | 13 | 1.15 | 1 | 0.41 | 0.267 | 0.564 | (A) BRAF Class 1 v12 |
| GBA | 1q22 | 6 | 1.03 | 3 | 2.03 | 0.267 | 0.564 | (B) BRAF Class 3 |
| FAT3 | 11q14.3 | 21 | 8.11 | 8 | 11.27 | 0.268 | 0.564 | (B) BRAF Class 3 |
| TSHZ2 | 20q13.2 | 3 | 3.49 | 2 | 9.09 | 0.268 | 0.565 | (B) BRAF Class 3 |

| | | | | | | | | |
|---|---|---|---|---|---|---|---|---|
| INPP4A | 2q11.2 | 23 | 1.95 | 7 | 2.77 | 0.27 | 0.567 | (B) BRAF Class 3 |
| ECT2L | 6q24.1 | 1 | 1.00 | 1 | 5.88 | 0.271 | 0.567 | (B) BRAF Class 3 |
| PLCG2 | 16q23.3 | 51 | 3.92 | 14 | 4.90 | 0.272 | 0.569 | (B) BRAF Class 3 |
| CIC | 19q13.2 | 148 | 6.95 | 40 | 7.83 | 0.273 | 0.57 | (B) BRAF Class 3 |
| IL6ST | 5q11.2 | 1 | 0.50 | 1 | 2.86 | 0.274 | 0.571 | (B) BRAF Class 3 |
| DNMT3A | 2p23.3 | 126 | 5.06 | 34 | 5.76 | 0.275 | 0.572 | (B) BRAF Class 3 |
| ERCC1 | 19q13.32 | 7 | 0.66 | 3 | 1.25 | 0.276 | 0.574 | (B) BRAF Class 3 |
| LATS2 | 13q12.11 | 48 | 4.00 | 13 | 5.02 | 0.277 | 0.574 | (B) BRAF Class 3 |
| MDM4 | 1q32.1 | 29 | 1.14 | 9 | 1.53 | 0.278 | 0.574 | (B) BRAF Class 3 |
| SSX2 | Xp11.22 | 1 | 1.75 | 1 | 10.00 | 0.278 | 0.574 | (B) BRAF Class 3 |
| GLI3 | 7p14.1 | 26 | 7.34 | 7 | 10.14 | 0.281 | 0.578 | (B) BRAF Class 3 |
| ACVR2A | :q22.3-q23.: | 1 | 0.85 | 1 | 4.76 | 0.282 | 0.578 | (B) BRAF Class 3 |
| TAF1L | 9p21.1 | 1 | 0.85 | 1 | 4.76 | 0.282 | 0.578 | (B) BRAF Class 3 |
| TGM7 | 5q15.2-q15. | 1 | 0.85 | 1 | 4.76 | 0.282 | 0.578 | (B) BRAF Class 3 |
| CYSLTR2 | 13q14.2 | 7 | 1.07 | 3 | 1.99 | 0.282 | 0.578 | (B) BRAF Class 3 |
| HLF | 17q22 | 1 | 0.82 | 1 | 4.55 | 0.283 | 0.578 | (B) BRAF Class 3 |
| ACVR1 | 2q24.1 | 22 | 1.18 | 7 | 1.64 | 0.284 | 0.578 | (B) BRAF Class 3 |
| CRTC2 | 1q21.3 | 7 | 1.98 | 0 | 0.00 | 0.285 | 0.578 | (A) BRAF Class 1 v12 |
| PRF1 | 10q22.1 | 31 | 3.33 | 5 | 2.27 | 0.285 | 0.578 | (A) BRAF Class 1 v12 |
| FUBP1 | 1p31.1 | 27 | 1.79 | 4 | 1.15 | 0.286 | 0.578 | (A) BRAF Class 1 v12 |
| GLI2 | 2q14.2 | 79 | 7.52 | 22 | 8.80 | 0.286 | 0.578 | (B) BRAF Class 3 |
| SMARCAL1 | 2q35 | 2 | 2.15 | 1 | 9.09 | 0.287 | 0.58 | (B) BRAF Class 3 |
| MRE11 | 11q21 | 56 | 2.54 | 16 | 3.08 | 0.288 | 0.58 | (B) BRAF Class 3 |
| ZNRF3 | 22q12.1 | 41 | 5.85 | 8 | 4.40 | 0.288 | 0.58 | (A) BRAF Class 1 v12 |
| RINT1 | 7q22.3 | 13 | 2.23 | 5 | 3.38 | 0.29 | 0.583 | (B) BRAF Class 3 |
| INSR | 19p13.2 | 57 | 4.31 | 10 | 3.37 | 0.291 | 0.583 | (A) BRAF Class 1 v12 |
| CBFB | 16q22.1 | 16 | 0.76 | 2 | 0.39 | 0.291 | 0.583 | (A) BRAF Class 1 v12 |
| MITF | 3p13 | 64 | 2.60 | 18 | 3.10 | 0.292 | 0.583 | (B) BRAF Class 3 |
| EMSY | 11q13.5 | 15 | 2.12 | 5 | 3.16 | 0.293 | 0.583 | (B) BRAF Class 3 |
| NAV3 | 12q21.2 | 8 | 5.10 | 3 | 9.09 | 0.293 | 0.583 | (B) BRAF Class 3 |
| FANCI | 15q26.1 | 47 | 4.13 | 8 | 3.13 | 0.294 | 0.583 | (A) BRAF Class 1 v12 |
| AFDN | 6q27 | 1 | 0.86 | 1 | 4.55 | 0.294 | 0.583 | (B) BRAF Class 3 |
| RBM15 | 1p13.3 | 1 | 0.86 | 1 | 4.55 | 0.294 | 0.583 | (B) BRAF Class 3 |
| DMD | (p21.2-p21.: | 13 | 3.67 | 4 | 5.80 | 0.295 | 0.583 | (B) BRAF Class 3 |
| MYD88 | 3p22.2 | 31 | 1.21 | 5 | 0.84 | 0.298 | 0.587 | (A) BRAF Class 1 v12 |
| RSPO3 | 6q22.33 | 20 | 2.56 | 3 | 1.55 | 0.299 | 0.587 | (A) BRAF Class 1 v12 |
| PNRC1 | 6q15 | 17 | 1.05 | 2 | 0.56 | 0.299 | 0.587 | (A) BRAF Class 1 v12 |
| MBD4 | 3q21.3 | 15 | 2.57 | 2 | 1.35 | 0.299 | 0.587 | (A) BRAF Class 1 v12 |
| SMO | 7q32.1 | 102 | 3.20 | 26 | 3.66 | 0.301 | 0.588 | (B) BRAF Class 3 |
| LATS1 | 6q25.1 | 24 | 2.04 | 7 | 2.77 | 0.301 | 0.588 | (B) BRAF Class 3 |
| BAP1 | 3p21.1 | 64 | 2.52 | 12 | 2.04 | 0.303 | 0.589 | (A) BRAF Class 1 v12 |
| MYH9 | 22q12.3 | 8 | 3.11 | 3 | 5.45 | 0.303 | 0.589 | (B) BRAF Class 3 |
| NPM1 | 5q35.1 | 22 | 0.71 | 3 | 0.43 | 0.303 | 0.589 | (A) BRAF Class 1 v12 |
| IRS2 | 13q34 | 78 | 4.98 | 21 | 5.79 | 0.304 | 0.589 | (B) BRAF Class 3 |
| RBBP8 | 18q11.2 | 31 | 5.31 | 10 | 6.76 | 0.305 | 0.589 | (B) BRAF Class 3 |
| TGFBR1 | 9q22.33 | 20 | 1.70 | 6 | 2.37 | 0.305 | 0.589 | (B) BRAF Class 3 |
| TSHR | 14q31.1 | 57 | 2.64 | 16 | 3.16 | 0.305 | 0.589 | (B) BRAF Class 3 |
| SOX9 | 17q24.3 | 109 | 4.55 | 29 | 5.14 | 0.305 | 0.589 | (B) BRAF Class 3 |

| | | | | | | | | |
|---|---|---|---|---|---|---|---|---|
| ZNF217 | 20q13.2 | 54 | 3.37 | 10 | 2.67 | 0.306 | 0.589 | (A) BRAF Class 1 v12 |
| ICOSLG | 21q22.3 | 16 | 1.36 | 5 | 1.98 | 0.308 | 0.589 | (B) BRAF Class 3 |
| UROD | 1p34.1 | 10 | 1.71 | 4 | 2.70 | 0.308 | 0.589 | (B) BRAF Class 3 |
| NBN | 8q21.3 | 77 | 3.05 | 15 | 2.55 | 0.309 | 0.589 | (A) BRAF Class 1 v12 |
| CBFA2T3 | 16q24.3 | 24 | 4.07 | 8 | 5.37 | 0.309 | 0.589 | (B) BRAF Class 3 |
| CSF1R | 5q32 | 63 | 2.46 | 16 | 2.92 | 0.309 | 0.589 | (B) BRAF Class 3 |
| STK40 | 1p34.3 | 12 | 1.02 | 4 | 1.58 | 0.309 | 0.589 | (B) BRAF Class 3 |
| ZMYM3 | Xq13.1 | 6 | 2.71 | 0 | 0.00 | 0.311 | 0.591 | (A) BRAF Class 1 v12 |
| NKX2-1 | 14q13.3 | 58 | 2.36 | 11 | 1.90 | 0.311 | 0.591 | (A) BRAF Class 1 v12 |
| U2AF1 | 21q22.3 | 38 | 1.51 | 11 | 1.88 | 0.313 | 0.593 | (B) BRAF Class 3 |
| EME1 | 17q21.33 | 10 | 1.71 | 1 | 0.68 | 0.314 | 0.593 | (A) BRAF Class 1 v12 |
| PTPN1 | 20q13.13 | 1 | 0.59 | 1 | 2.86 | 0.314 | 0.593 | (B) BRAF Class 3 |
| BCR | 22q11.23 | 22 | 3.82 | 3 | 2.38 | 0.315 | 0.593 | (A) BRAF Class 1 v12 |
| AURKA | 20q13.2 | 26 | 1.06 | 8 | 1.38 | 0.316 | 0.593 | (B) BRAF Class 3 |
| ETV1 | 7p21.2 | 58 | 2.55 | 16 | 3.02 | 0.316 | 0.593 | (B) BRAF Class 3 |
| PAK3 | Xq23 | 17 | 2.48 | 5 | 3.55 | 0.317 | 0.593 | (B) BRAF Class 3 |
| XBP1 | 2q12.1\|22q1 | 1 | 0.45 | 1 | 2.13 | 0.317 | 0.593 | (B) BRAF Class 3 |
| IPMK | 10q21.1 | 1 | 0.64 | 1 | 3.03 | 0.318 | 0.593 | (B) BRAF Class 3 |
| SHH | 7q36.3 | 1 | 0.64 | 1 | 3.03 | 0.318 | 0.593 | (B) BRAF Class 3 |
| SPRY2 | 13q31.1 | 1 | 0.64 | 1 | 3.03 | 0.318 | 0.593 | (B) BRAF Class 3 |
| ATF1 | 12q13.12 | 1 | 0.39 | 1 | 1.82 | 0.322 | 0.596 | (B) BRAF Class 3 |
| DDIT3 | 12q13.3 | 1 | 0.39 | 1 | 1.82 | 0.322 | 0.596 | (B) BRAF Class 3 |
| EGLN1 | 1q42.2 | 5 | 0.86 | 0 | 0.00 | 0.322 | 0.596 | (A) BRAF Class 1 v12 |
| ID4 | 6p22.3 | 5 | 0.86 | 0 | 0.00 | 0.322 | 0.596 | (A) BRAF Class 1 v12 |
| PLAG1 | 8q12.1 | 1 | 0.38 | 1 | 1.75 | 0.322 | 0.596 | (B) BRAF Class 3 |
| PIM1 | 6p21.2 | 19 | 0.86 | 6 | 1.18 | 0.323 | 0.596 | (B) BRAF Class 3 |
| FGFR3 | 4p16.3 | 91 | 2.72 | 23 | 3.10 | 0.325 | 0.598 | (B) BRAF Class 3 |
| PHF6 | Xq26.2 | 20 | 1.33 | 6 | 1.80 | 0.325 | 0.598 | (B) BRAF Class 3 |
| TAF1 | Xq13.1 | 28 | 5.31 | 7 | 6.93 | 0.326 | 0.598 | (B) BRAF Class 3 |
| NAB2 | 12q13.3 | 13 | 3.90 | 3 | 6.25 | 0.327 | 0.599 | (B) BRAF Class 3 |
| AKT3 | 1q43-q44 | 35 | 1.36 | 6 | 1.01 | 0.327 | 0.599 | (A) BRAF Class 1 v12 |
| DDB2 | 11p11.2 | 15 | 1.46 | 5 | 2.07 | 0.328 | 0.599 | (B) BRAF Class 3 |
| SMARCB1 | q11.23\|22q | 41 | 1.27 | 11 | 1.56 | 0.328 | 0.599 | (B) BRAF Class 3 |
| MC1-LUC7L | 7q34 | 1 | 0.03 | 1 | 0.13 | 0.332 | 0.602 | (B) BRAF Class 3 |
| MT-ND6 | - | 1 | 0.03 | 1 | 0.13 | 0.332 | 0.602 | (B) BRAF Class 3 |
| SOX10 | 22q13.1 | 13 | 1.91 | 4 | 2.84 | 0.333 | 0.602 | (B) BRAF Class 3 |
| FGF8 | 10q24.32 | 2 | 0.60 | 1 | 2.08 | 0.333 | 0.602 | (B) BRAF Class 3 |
| AGO2 | 8q24.3 | 26 | 4.94 | 8 | 6.30 | 0.334 | 0.602 | (B) BRAF Class 3 |
| RAD52 | 12p13.33 | 24 | 1.28 | 4 | 0.88 | 0.334 | 0.602 | (A) BRAF Class 1 v12 |
| BCL10 | 1p22.3 | 17 | 1.41 | 5 | 1.96 | 0.334 | 0.602 | (B) BRAF Class 3 |
| BABAM2 | 2p23.2 | 14 | 2.40 | 5 | 3.38 | 0.335 | 0.602 | (B) BRAF Class 3 |
| GATA1 | Xp11.23 | 20 | 1.27 | 6 | 1.70 | 0.336 | 0.603 | (B) BRAF Class 3 |
| RARA | 17q21.2 | 43 | 1.75 | 8 | 1.38 | 0.34 | 0.61 | (A) BRAF Class 1 v12 |
| KLLN | 10q23.31 | 14 | 2.40 | 2 | 1.35 | 0.341 | 0.611 | (A) BRAF Class 1 v12 |
| ARHGEF12 | 11q23.3 | 25 | 4.24 | 8 | 5.37 | 0.342 | 0.612 | (B) BRAF Class 3 |
| KIF1B | 1p36.22 | 27 | 4.62 | 5 | 3.38 | 0.344 | 0.615 | (A) BRAF Class 1 v12 |
| INHA | 2q35 | 22 | 1.98 | 3 | 1.29 | 0.345 | 0.615 | (A) BRAF Class 1 v12 |
| TCF3 | 19p13.3 | 66 | 3.12 | 17 | 3.58 | 0.349 | 0.62 | (B) BRAF Class 3 |

| | | | | | | | | |
|---|---|---|---|---|---|---|---|---|
| DIS3L2 | 2q37.1 | 19 | 2.90 | 3 | 1.89 | 0.349 | 0.62 | (A) BRAF Class 1 v12 |
| NEIL2 | 8p23.1 | 4 | 0.68 | 2 | 1.35 | 0.35 | 0.621 | (B) BRAF Class 3 |
| NR4A3 | 9q31.1 | 2 | 1.24 | 1 | 4.00 | 0.353 | 0.625 | (B) BRAF Class 3 |
| XIAP | Xq25 | 13 | 1.10 | 4 | 1.58 | 0.353 | 0.625 | (B) BRAF Class 3 |
| KNSTRN | 15q15.1 | 20 | 2.70 | 3 | 1.78 | 0.355 | 0.626 | (A) BRAF Class 1 v12 |
| KLF4 | 9q31.2 | 34 | 1.75 | 6 | 1.34 | 0.356 | 0.627 | (A) BRAF Class 1 v12 |
| DNMT1 | 19p13.2 | 53 | 4.04 | 13 | 4.69 | 0.36 | 0.633 | (B) BRAF Class 3 |
| YAP1 | 11q22.1 | 28 | 1.49 | 8 | 1.84 | 0.362 | 0.635 | (B) BRAF Class 3 |
| PDE4DIP | 1q21.2 | 6 | 4.92 | 0 | 0.00 | 0.363 | 0.636 | (A) BRAF Class 1 v12 |
| SHOC2 | 10q25.2 | 8 | 1.52 | 3 | 2.36 | 0.364 | 0.636 | (B) BRAF Class 3 |
| SMYD3 | 1q44 | 8 | 1.52 | 3 | 2.36 | 0.364 | 0.636 | (B) BRAF Class 3 |
| PHF1 | 6p21.32 | 5 | 0.15 | 0 | 0.00 | 0.364 | 0.636 | (A) BRAF Class 1 v12 |
| NRIP1 | 1q11.2-q21. | 3 | 3.23 | 1 | 9.09 | 0.365 | 0.636 | (B) BRAF Class 3 |
| MYC | 8q24.21 | 89 | 3.46 | 23 | 3.83 | 0.366 | 0.636 | (B) BRAF Class 3 |
| PIK3CB | 3q22.3 | 46 | 3.11 | 12 | 3.63 | 0.366 | 0.636 | (B) BRAF Class 3 |
| KLHL6 | 3q27.1 | 21 | 2.92 | 6 | 3.75 | 0.367 | 0.636 | (B) BRAF Class 3 |
| NEIL1 | 15q24.2 | 9 | 1.54 | 1 | 0.68 | 0.368 | 0.636 | (A) BRAF Class 1 v12 |
| TDG | 12q23.3 | 9 | 1.54 | 1 | 0.68 | 0.368 | 0.636 | (A) BRAF Class 1 v12 |
| EPHB4 | 7q22.1 | 4 | 1.33 | 2 | 2.50 | 0.369 | 0.637 | (B) BRAF Class 3 |
| RAD51 | 15q15.1 | 19 | 0.87 | 3 | 0.58 | 0.37 | 0.637 | (A) BRAF Class 1 v12 |
| RECQL | 12p12.1 | 14 | 2.66 | 2 | 1.57 | 0.37 | 0.637 | (A) BRAF Class 1 v12 |
| EGR1 | 5q31.2 | 1 | 0.87 | 1 | 3.33 | 0.372 | 0.638 | (B) BRAF Class 3 |
| KMT2B | 19q13.12 | 121 | 11.66 | 28 | 12.67 | 0.372 | 0.638 | (B) BRAF Class 3 |
| MDC1 | 6p21.33 | 66 | 5.60 | 16 | 6.32 | 0.373 | 0.638 | (B) BRAF Class 3 |
| GEN1 | 2p24.2 | 24 | 2.31 | 4 | 1.67 | 0.375 | 0.638 | (A) BRAF Class 1 v12 |
| SDHAF2 | 11q12.2 | 10 | 0.45 | 1 | 0.20 | 0.375 | 0.638 | (A) BRAF Class 1 v12 |
| BUB1B | 15q15.1 | 24 | 2.34 | 7 | 2.90 | 0.375 | 0.638 | (B) BRAF Class 3 |
| GPC3 | Xq26.2 | 20 | 1.77 | 6 | 2.26 | 0.375 | 0.638 | (B) BRAF Class 3 |
| FGF6 | 12p13.32 | 10 | 1.41 | 1 | 0.63 | 0.376 | 0.638 | (A) BRAF Class 1 v12 |
| PRKCI | 3q26.2 | 48 | 2.50 | 13 | 2.87 | 0.376 | 0.638 | (B) BRAF Class 3 |
| PIK3C3 | 18q12.3 | 33 | 2.52 | 9 | 3.01 | 0.377 | 0.638 | (B) BRAF Class 3 |
| SS18 | 18q11.2 | 8 | 1.01 | 3 | 1.52 | 0.377 | 0.638 | (B) BRAF Class 3 |
| EWSR1 | 22q12.2 | 28 | 1.75 | 8 | 2.13 | 0.378 | 0.638 | (B) BRAF Class 3 |
| TRRAP | 7q22.1 | 1 | 0.45 | 1 | 1.67 | 0.378 | 0.639 | (B) BRAF Class 3 |
| FKBP9 | 7p14.3 | 6 | 1.69 | 2 | 2.90 | 0.385 | 0.646 | (B) BRAF Class 3 |
| TFE3 | Xp11.23 | 30 | 2.63 | 8 | 3.16 | 0.385 | 0.646 | (B) BRAF Class 3 |
| CENPA | 2p23.3 | 5 | 0.45 | 0 | 0.00 | 0.385 | 0.646 | (A) BRAF Class 1 v12 |
| CDKN1C | 11p15.4 | 5 | 0.51 | 2 | 0.88 | 0.389 | 0.651 | (B) BRAF Class 3 |
| RAD51B | 14q24.1 | 13 | 0.93 | 4 | 1.26 | 0.39 | 0.651 | (B) BRAF Class 3 |
| CARM1 | 19p13.2 | 9 | 1.71 | 1 | 0.79 | 0.391 | 0.651 | (A) BRAF Class 1 v12 |
| PIK3R2 | 19p13.11 | 34 | 2.21 | 9 | 2.60 | 0.392 | 0.651 | (B) BRAF Class 3 |
| UBR5 | 8q22.3 | 2 | 1.49 | 1 | 4.17 | 0.392 | 0.651 | (B) BRAF Class 3 |
| DKC1 | Xq28 | 8 | 1.37 | 3 | 2.01 | 0.393 | 0.651 | (B) BRAF Class 3 |
| DNAJB1 | 19p13.12 | 14 | 1.07 | 4 | 1.43 | 0.393 | 0.651 | (B) BRAF Class 3 |
| CTNNA1 | 5q31.2 | 28 | 2.19 | 8 | 2.61 | 0.393 | 0.651 | (B) BRAF Class 3 |
| BCL9 | 1q21.2 | 2 | 1.64 | 1 | 4.55 | 0.394 | 0.651 | (B) BRAF Class 3 |
| CDH11 | 16q21 | 2 | 1.64 | 1 | 4.55 | 0.394 | 0.651 | (B) BRAF Class 3 |
| TRIM33 | 1p13.2 | 2 | 1.64 | 1 | 4.55 | 0.394 | 0.651 | (B) BRAF Class 3 |

| | | | | | | | | |
|---|---|---|---|---|---|---|---|---|
| FRS2 | 12q15 | 6 | 0.95 | 2 | 1.59 | 0.396 | 0.652 | (B) BRAF Class 3 |
| FIP1L1 | 4q12 | 2 | 1.30 | 1 | 3.57 | 0.396 | 0.652 | (B) BRAF Class 3 |
| CHD5 | 1p36.31 | 10 | 6.37 | 3 | 9.09 | 0.4 | 0.657 | (B) BRAF Class 3 |
| PRDM1 | 6q21 | 90 | 3.66 | 23 | 3.97 | 0.401 | 0.657 | (B) BRAF Class 3 |
| SMARCE1 | 17q21.2 | 5 | 0.74 | 2 | 1.23 | 0.403 | 0.657 | (B) BRAF Class 3 |
| BRD3 | 9q34.2 | 30 | 4.33 | 6 | 3.49 | 0.404 | 0.657 | (A) BRAF Class 1 v12 |
| OGG1 | 3p25.3 | 4 | 0.68 | 0 | 0.00 | 0.404 | 0.657 | (A) BRAF Class 1 v12 |
| UBE2T | 1q32.1 | 4 | 0.68 | 0 | 0.00 | 0.404 | 0.657 | (A) BRAF Class 1 v12 |
| STAT5B | 17q21.2 | 37 | 3.11 | 9 | 3.60 | 0.404 | 0.657 | (B) BRAF Class 3 |
| DNAH9 | 17p12 | 6 | 8.70 | 2 | 14.29 | 0.405 | 0.657 | (B) BRAF Class 3 |
| HIST1H3E | 6p22.2 | 10 | 0.90 | 1 | 0.43 | 0.405 | 0.657 | (A) BRAF Class 1 v12 |
| GATA2 | 3q21.3 | 50 | 2.30 | 10 | 1.98 | 0.406 | 0.657 | (A) BRAF Class 1 v12 |
| LAMP1 | 13q34 | 6 | 1.40 | 0 | 0.00 | 0.406 | 0.657 | (A) BRAF Class 1 v12 |
| SDHD | 11q23.1 | 9 | 0.37 | 3 | 0.52 | 0.406 | 0.657 | (B) BRAF Class 3 |
| LIG4 | 13q33.3 | 21 | 3.60 | 4 | 2.70 | 0.407 | 0.657 | (A) BRAF Class 1 v12 |
| RRAGC | 1p34.3 | 5 | 0.75 | 2 | 1.25 | 0.407 | 0.657 | (B) BRAF Class 3 |
| RELA | 11q13.1 | 16 | 2.05 | 5 | 2.58 | 0.408 | 0.657 | (B) BRAF Class 3 |
| TAP2 | 6p21.32 | 5 | 0.95 | 2 | 1.57 | 0.408 | 0.657 | (B) BRAF Class 3 |
| MUS81 | 11q13.1 | 12 | 2.05 | 4 | 2.70 | 0.411 | 0.66 | (B) BRAF Class 3 |
| RANBP2 | 2q13 | 56 | 11.34 | 9 | 9.78 | 0.412 | 0.66 | (A) BRAF Class 1 v12 |
| IRF2 | 4q35.1 | 6 | 1.15 | 2 | 1.83 | 0.413 | 0.662 | (B) BRAF Class 3 |
| CDK1 | 10q21.2 | 2 | 0.56 | 1 | 1.45 | 0.415 | 0.662 | (B) BRAF Class 3 |
| PDCD1LG2 | 9p24.1 | 25 | 1.21 | 7 | 1.43 | 0.415 | 0.662 | (B) BRAF Class 3 |
| FOXP1 | 3p13 | 35 | 2.32 | 9 | 2.66 | 0.415 | 0.662 | (B) BRAF Class 3 |
| HIST1H3G | 6p22.2 | 6 | 0.54 | 2 | 0.86 | 0.416 | 0.663 | (B) BRAF Class 3 |
| HIST1H3I | 6p22.1 | 6 | 0.54 | 2 | 0.86 | 0.417 | 0.664 | (B) BRAF Class 3 |
| TNFAIP3 | 6q23.3 | 58 | 2.36 | 15 | 2.59 | 0.42 | 0.667 | (B) BRAF Class 3 |
| RAD51C | 17q22 | 25 | 1.15 | 7 | 1.35 | 0.421 | 0.667 | (B) BRAF Class 3 |
| SDHC | 1q23.3 | 18 | 0.76 | 3 | 0.54 | 0.422 | 0.667 | (A) BRAF Class 1 v12 |
| RAD51D | 17q12 | 21 | 0.96 | 6 | 1.16 | 0.422 | 0.667 | (B) BRAF Class 3 |
| PRKN | 6q26 | 79 | 3.29 | 20 | 3.55 | 0.423 | 0.668 | (B) BRAF Class 3 |
| FAN1 | 15q13.3 | 16 | 2.74 | 5 | 3.38 | 0.424 | 0.668 | (B) BRAF Class 3 |
| IGF1R | 15q26.3 | 111 | 4.32 | 24 | 4.03 | 0.425 | 0.668 | (A) BRAF Class 1 v12 |
| KMT2C | 7q36.1 | 184 | 12.65 | 42 | 13.21 | 0.425 | 0.668 | (B) BRAF Class 3 |
| PAX3 | 2q36.1 | 11 | 1.94 | 1 | 0.97 | 0.428 | 0.67 | (A) BRAF Class 1 v12 |
| STAT3 | 17q21.2 | 45 | 1.84 | 9 | 1.59 | 0.428 | 0.67 | (A) BRAF Class 1 v12 |
| KLF2 | 19p13.11 | 8 | 1.37 | 1 | 0.68 | 0.428 | 0.67 | (A) BRAF Class 1 v12 |
| BIRC3 | 11q22.2 | 15 | 1.24 | 4 | 1.56 | 0.429 | 0.67 | (B) BRAF Class 3 |
| RNF8 | 6p21.2 | 5 | 0.86 | 2 | 1.35 | 0.43 | 0.671 | (B) BRAF Class 3 |
| RXRA | 9q34.2 | 17 | 3.23 | 3 | 2.36 | 0.433 | 0.674 | (A) BRAF Class 1 v12 |
| USP28 | 11q23.2 | 20 | 3.42 | 6 | 4.05 | 0.434 | 0.674 | (B) BRAF Class 3 |
| NTHL1 | 16p13.3 | 9 | 0.78 | 3 | 1.05 | 0.435 | 0.674 | (B) BRAF Class 3 |
| POLR2A | 17p13.1 | 4 | 4.30 | 1 | 9.09 | 0.435 | 0.674 | (B) BRAF Class 3 |
| SAMD9L | 7q21.2 | 4 | 4.30 | 1 | 9.09 | 0.435 | 0.674 | (B) BRAF Class 3 |
| AKT2 | 19q13.2 | 49 | 1.90 | 10 | 1.68 | 0.437 | 0.675 | (A) BRAF Class 1 v12 |
| CD276 | 15q24.1 | 19 | 1.61 | 3 | 1.19 | 0.437 | 0.675 | (A) BRAF Class 1 v12 |
| BRCC3 | Xq28 | 12 | 2.02 | 2 | 1.33 | 0.439 | 0.678 | (A) BRAF Class 1 v12 |
| KCNIP1 | 5q35.1 | 6 | 2.28 | 2 | 3.45 | 0.44 | 0.678 | (B) BRAF Class 3 |

| | | | | | | | | |
|---|---|---|---|---|---|---|---|---|
| BRD4 | 19p13.12 | 126 | 5.23 | 28 | 4.95 | 0.442 | 0.679 | (A) BRAF Class 1 v12 |
| SHQ1 | 3p13 | 24 | 2.04 | 6 | 2.37 | 0.442 | 0.679 | (B) BRAF Class 3 |
| MYOD1 | 11p15.1 | 29 | 2.32 | 5 | 1.89 | 0.442 | 0.679 | (A) BRAF Class 1 v12 |
| SOCS1 | 16p13.13 | 26 | 1.06 | 5 | 0.86 | 0.443 | 0.679 | (A) BRAF Class 1 v12 |
| STAT5A | 17q21.2 | 20 | 1.70 | 5 | 2.02 | 0.443 | 0.679 | (B) BRAF Class 3 |
| ZFHX3 | 6q22.2-q22. | 152 | 12.22 | 31 | 11.65 | 0.446 | 0.682 | (A) BRAF Class 1 v12 |
| SRSF2 | 17q25.1 | 27 | 1.20 | 7 | 1.37 | 0.447 | 0.682 | (B) BRAF Class 3 |
| RACK1 | 5q35.3 | 4 | 1.52 | 0 | 0.00 | 0.449 | 0.684 | (A) BRAF Class 1 v12 |
| CDKN1B | 12p13.1 | 35 | 1.38 | 7 | 1.19 | 0.449 | 0.684 | (A) BRAF Class 1 v12 |
| DUSP4 | 8p12 | 8 | 1.21 | 1 | 0.63 | 0.451 | 0.685 | (A) BRAF Class 1 v12 |
| MSI1 | 12q24.31 | 8 | 1.52 | 1 | 0.79 | 0.452 | 0.686 | (A) BRAF Class 1 v12 |
| LTK | 15q15.1 | 3 | 1.30 | 0 | 0.00 | 0.454 | 0.686 | (A) BRAF Class 1 v12 |
| ORCS8-MEF2 | 19p13.11 | 19 | 0.57 | 5 | 0.67 | 0.454 | 0.686 | (B) BRAF Class 3 |
| CAMK1 | 3p25.3 | 2 | 0.06 | 1 | 0.13 | 0.454 | 0.686 | (B) BRAF Class 3 |
| TRBC2 | 7q34 | 2 | 0.06 | 1 | 0.13 | 0.454 | 0.686 | (B) BRAF Class 3 |
| TRAF2 | 9q34.3 | 25 | 2.13 | 6 | 2.43 | 0.457 | 0.688 | (B) BRAF Class 3 |
| HNF1A | 12q24.31 | 106 | 3.35 | 22 | 3.17 | 0.457 | 0.688 | (A) BRAF Class 1 v12 |
| RAD21 | 8q24.11 | 57 | 2.51 | 12 | 2.29 | 0.458 | 0.688 | (A) BRAF Class 1 v12 |
| FANCB | Xp22.2 | 20 | 2.99 | 4 | 2.40 | 0.458 | 0.688 | (A) BRAF Class 1 v12 |
| FGF5 | 4q21.21 | 10 | 3.00 | 2 | 4.17 | 0.46 | 0.688 | (B) BRAF Class 3 |
| HIST1H3B | 6p22.2 | 19 | 0.91 | 5 | 1.06 | 0.46 | 0.688 | (B) BRAF Class 3 |
| KDM5A | 12p13.33 | 120 | 5.68 | 30 | 5.89 | 0.46 | 0.688 | (B) BRAF Class 3 |
| FAH | 15q25.1 | 13 | 2.23 | 4 | 2.70 | 0.461 | 0.688 | (B) BRAF Class 3 |
| GABRA6 | 5q34 | 16 | 3.05 | 4 | 3.67 | 0.461 | 0.688 | (B) BRAF Class 3 |
| KDM6B | 17p13.1 | 20 | 5.51 | 3 | 4.23 | 0.463 | 0.69 | (A) BRAF Class 1 v12 |
| FOXA1 | 14q21.1 | 39 | 1.98 | 8 | 1.75 | 0.464 | 0.691 | (A) BRAF Class 1 v12 |
| TNFRSF14 | 1p36.32 | 17 | 1.09 | 3 | 0.83 | 0.465 | 0.691 | (A) BRAF Class 1 v12 |
| LMO1 | 11p15.4 | 17 | 0.75 | 3 | 0.57 | 0.466 | 0.692 | (A) BRAF Class 1 v12 |
| TGFB1 | 19q13.2 | 3 | 6.25 | 1 | 12.50 | 0.47 | 0.695 | (B) BRAF Class 3 |
| GNAQ | 9q21.2 | 28 | 0.86 | 7 | 0.95 | 0.471 | 0.695 | (B) BRAF Class 3 |
| CCND1 | 11q13.3 | 34 | 1.34 | 7 | 1.18 | 0.471 | 0.695 | (A) BRAF Class 1 v12 |
| BRINP3 | 1q31.1 | 5 | 4.20 | 0 | 0.00 | 0.471 | 0.695 | (A) BRAF Class 1 v12 |
| LMO2 | 11p13 | 6 | 0.65 | 2 | 0.92 | 0.472 | 0.695 | (B) BRAF Class 3 |
| IKBKE | 1q32.1 | 23 | 1.47 | 6 | 1.65 | 0.472 | 0.695 | (B) BRAF Class 3 |
| MAP2K4 | 17p12 | 40 | 1.58 | 10 | 1.70 | 0.473 | 0.695 | (B) BRAF Class 3 |
| BCL11B | 14q32.2 | 32 | 4.59 | 7 | 4.05 | 0.473 | 0.695 | (A) BRAF Class 1 v12 |
| DDB1 | 11q12.2 | 21 | 3.60 | 6 | 4.05 | 0.474 | 0.695 | (B) BRAF Class 3 |
| PDCD1 | 2q37.3 | 33 | 2.57 | 8 | 2.81 | 0.475 | 0.695 | (B) BRAF Class 3 |
| ERCC2 | 19q13.32 | 75 | 3.01 | 18 | 3.15 | 0.477 | 0.695 | (B) BRAF Class 3 |
| FH | 1q43 | 48 | 1.97 | 12 | 2.09 | 0.477 | 0.695 | (B) BRAF Class 3 |
| MTA1 | 14q32.33 | 25 | 4.28 | 7 | 4.73 | 0.478 | 0.695 | (B) BRAF Class 3 |
| RHBDF2 | 17q25.1 | 25 | 4.28 | 7 | 4.73 | 0.478 | 0.695 | (B) BRAF Class 3 |
| CCND3 | 6p21.1 | 21 | 0.83 | 4 | 0.68 | 0.478 | 0.695 | (A) BRAF Class 1 v12 |
| NCOR1 | 17p12-p11.2 | 79 | 5.33 | 17 | 5.04 | 0.478 | 0.695 | (A) BRAF Class 1 v12 |
| SPEN | 36.21-p36.1 | 142 | 9.10 | 32 | 8.82 | 0.479 | 0.695 | (A) BRAF Class 1 v12 |
| FANCL | 2p16.1 | 16 | 1.26 | 3 | 0.98 | 0.481 | 0.695 | (A) BRAF Class 1 v12 |
| BCORL1 | Xq26.1 | 116 | 7.16 | 28 | 7.37 | 0.481 | 0.695 | (B) BRAF Class 3 |
| MSI2 | 17q22 | 6 | 1.14 | 2 | 1.57 | 0.481 | 0.695 | (B) BRAF Class 3 |

| | | | | | | | | |
|---|---|---|---|---|---|---|---|---|
| XPA | 9q22.33 | 12 | 1.17 | 2 | 0.83 | 0.484 | 0.695 | (A) BRAF Class 1 v12 |
| FGF7 | 15q21.2 | 4 | 0.92 | 0 | 0.00 | 0.485 | 0.695 | (A) BRAF Class 1 v12 |
| CDK2 | 12q13.2 | 3 | 0.69 | 1 | 1.28 | 0.486 | 0.695 | (B) BRAF Class 3 |
| PGBD3 | 10q11.23 | 11 | 0.33 | 3 | 0.40 | 0.486 | 0.695 | (B) BRAF Class 3 |
| NEIL3 | 4q34.3 | 23 | 3.94 | 5 | 3.38 | 0.487 | 0.695 | (A) BRAF Class 1 v12 |
| CSMD3 | 8q23.3 | 3 | 2.56 | 1 | 4.76 | 0.487 | 0.695 | (B) BRAF Class 3 |
| FN1 | 2q35 | 3 | 2.56 | 1 | 4.76 | 0.487 | 0.695 | (B) BRAF Class 3 |
| FOXP4 | 6p21.1 | 3 | 2.56 | 1 | 4.76 | 0.487 | 0.695 | (B) BRAF Class 3 |
| ITGA10 | 1q21.1 | 3 | 2.56 | 1 | 4.76 | 0.487 | 0.695 | (B) BRAF Class 3 |
| IDH2 | 15q26.1 | 48 | 1.51 | 10 | 1.39 | 0.488 | 0.695 | (A) BRAF Class 1 v12 |
| NIN | 14q22.1 | 3 | 2.46 | 1 | 4.55 | 0.489 | 0.695 | (B) BRAF Class 3 |
| TOPBP1 | 3q22.1 | 23 | 3.88 | 5 | 3.33 | 0.489 | 0.695 | (A) BRAF Class 1 v12 |
| VTCN1 | 1p13.1-p12 | 13 | 1.10 | 2 | 0.79 | 0.489 | 0.695 | (A) BRAF Class 1 v12 |
| RIF1 | 2q23.3 | 41 | 7.02 | 11 | 7.43 | 0.49 | 0.695 | (B) BRAF Class 3 |
| SERPINA1 | 14q32.13 | 15 | 2.57 | 3 | 2.03 | 0.49 | 0.695 | (A) BRAF Class 1 v12 |
| RHEB | 7q36.1 | 11 | 0.52 | 3 | 0.62 | 0.493 | 0.696 | (B) BRAF Class 3 |
| JAZF1 | 'p15.2-p15.: | 6 | 0.76 | 2 | 1.03 | 0.493 | 0.696 | (B) BRAF Class 3 |
| MEN1 | 11q13.1 | 57 | 2.32 | 14 | 2.41 | 0.493 | 0.696 | (B) BRAF Class 3 |
| RHOA | 3p21.31 | 30 | 1.48 | 6 | 1.31 | 0.495 | 0.696 | (A) BRAF Class 1 v12 |
| TMEM127 | 2q11.2 | 11 | 0.61 | 3 | 0.73 | 0.496 | 0.696 | (B) BRAF Class 3 |
| DCLRE1C | 10p13 | 7 | 1.20 | 1 | 0.68 | 0.496 | 0.696 | (A) BRAF Class 1 v12 |
| CTLA4 | 2q33.2 | 25 | 1.43 | 5 | 1.24 | 0.498 | 0.696 | (A) BRAF Class 1 v12 |
| XRCC2 | 7q36.1 | 23 | 1.22 | 6 | 1.34 | 0.499 | 0.696 | (B) BRAF Class 3 |
| DYNC1I1 | 7q21.3 | 2 | 2.33 | 1 | 4.55 | 0.499 | 0.696 | (B) BRAF Class 3 |
| RPS6KA4 | 11q13.1 | 36 | 3.06 | 7 | 2.77 | 0.5 | 0.696 | (A) BRAF Class 1 v12 |
| REL | 2p16.1 | 41 | 1.68 | 9 | 1.55 | 0.501 | 0.696 | (A) BRAF Class 1 v12 |
| CDH20 | 18q21.33 | 2 | 1.00 | 1 | 1.92 | 0.502 | 0.696 | (B) BRAF Class 3 |
| EPHB6 | 7q34 | 2 | 1.00 | 1 | 1.92 | 0.502 | 0.696 | (B) BRAF Class 3 |
| KIF5B | 10p11.22 | 11 | 2.68 | 1 | 1.56 | 0.502 | 0.696 | (A) BRAF Class 1 v12 |
| IRS1 | 2q36.3 | 63 | 5.35 | 14 | 5.53 | 0.502 | 0.696 | (B) BRAF Class 3 |
| NUP93 | 16q13 | 30 | 2.06 | 6 | 1.84 | 0.502 | 0.696 | (A) BRAF Class 1 v12 |
| POLH | 6p21.1 | 11 | 1.70 | 2 | 1.24 | 0.503 | 0.696 | (A) BRAF Class 1 v12 |
| CHD4 | 12p13.31 | 38 | 6.04 | 8 | 6.40 | 0.504 | 0.696 | (B) BRAF Class 3 |
| RBL2 | 16q12.2 | 8 | 2.26 | 2 | 2.90 | 0.505 | 0.696 | (B) BRAF Class 3 |
| SRP19 | 5q22.2 | 3 | 0.81 | 0 | 0.00 | 0.505 | 0.696 | (A) BRAF Class 1 v12 |
| FLT4 | 5q35.3 | 171 | 7.04 | 39 | 6.93 | 0.506 | 0.696 | (A) BRAF Class 1 v12 |
| BABAM1 | 19p13.11 | 14 | 1.29 | 4 | 1.45 | 0.507 | 0.696 | (B) BRAF Class 3 |
| HSD3B1 | 1p12 | 13 | 2.48 | 2 | 1.83 | 0.508 | 0.696 | (A) BRAF Class 1 v12 |
| NCOR2 | 12q24.31 | 3 | 2.61 | 1 | 4.55 | 0.508 | 0.696 | (B) BRAF Class 3 |
| CSNK1A1 | 5q32 | 5 | 1.50 | 0 | 0.00 | 0.508 | 0.696 | (A) BRAF Class 1 v12 |
| FGF9 | 13q12.11 | 5 | 1.50 | 0 | 0.00 | 0.508 | 0.696 | (A) BRAF Class 1 v12 |
| TLX3 | 5q35.1 | 14 | 2.38 | 4 | 2.68 | 0.508 | 0.696 | (B) BRAF Class 3 |
| BBC3 | 19q13.32 | 8 | 0.68 | 1 | 0.40 | 0.509 | 0.696 | (A) BRAF Class 1 v12 |
| GREM1 | 15q13.3 | 11 | 0.56 | 3 | 0.66 | 0.51 | 0.696 | (B) BRAF Class 3 |
| FGF19 | 11q13.3 | 25 | 1.53 | 5 | 1.34 | 0.51 | 0.696 | (A) BRAF Class 1 v12 |
| CDK5 | 7q36.1 | 3 | 0.85 | 1 | 1.45 | 0.511 | 0.696 | (B) BRAF Class 3 |
| CHD2 | 15q26.1 | 38 | 6.03 | 8 | 6.35 | 0.511 | 0.696 | (B) BRAF Class 3 |
| TEK | 9p21.2 | 33 | 4.65 | 8 | 4.30 | 0.512 | 0.697 | (A) BRAF Class 1 v12 |

| | | | | | | | | |
|---|---|---|---|---|---|---|---|---|
| IGF2 | 11p15.5 | 24 | 1.15 | 5 | 1.01 | 0.513 | 0.697 | (A) BRAF Class 1 v12 |
| ERCC6 | 10q11.23 | 34 | 5.82 | 8 | 5.41 | 0.515 | 0.699 | (A) BRAF Class 1 v12 |
| HNRNPK | 9q21.32 | 4 | 0.93 | 1 | 1.49 | 0.517 | 0.699 | (B) BRAF Class 3 |
| ETV5 | 3q27.2 | 24 | 1.64 | 5 | 1.46 | 0.517 | 0.699 | (A) BRAF Class 1 v12 |
| PRPF8 | 17p13.3 | 14 | 3.23 | 2 | 2.44 | 0.519 | 0.702 | (A) BRAF Class 1 v12 |
| KMT5A | 12q24.31 | 3 | 0.55 | 0 | 0.00 | 0.522 | 0.704 | (A) BRAF Class 1 v12 |
| KAT6B | 10q22.2 | 43 | 6.29 | 11 | 6.47 | 0.523 | 0.704 | (B) BRAF Class 3 |
| PRKD1 | 14q12 | 11 | 1.87 | 3 | 2.14 | 0.523 | 0.704 | (B) BRAF Class 3 |
| GRM3 | q21.11-q21.1 | 58 | 8.80 | 12 | 8.45 | 0.524 | 0.704 | (A) BRAF Class 1 v12 |
| ERRFI1 | 1p36.23 | 25 | 1.74 | 6 | 1.83 | 0.528 | 0.708 | (B) BRAF Class 3 |
| ZNF703 | 8p11.23 | 21 | 2.96 | 5 | 3.16 | 0.528 | 0.708 | (B) BRAF Class 3 |
| GNAS | 20q13.32 | 104 | 3.17 | 23 | 3.18 | 0.531 | 0.711 | (B) BRAF Class 3 |
| TRIM37 | 17q22 | 18 | 3.08 | 4 | 2.70 | 0.532 | 0.712 | (A) BRAF Class 1 v12 |
| ARAF | Xp11.3 | 64 | 2.47 | 15 | 2.49 | 0.533 | 0.713 | (B) BRAF Class 3 |
| INPPL1 | 11q13.4 | 31 | 5.89 | 7 | 5.51 | 0.534 | 0.713 | (A) BRAF Class 1 v12 |
| MT-ND5 | - | 12 | 0.36 | 3 | 0.40 | 0.535 | 0.713 | (B) BRAF Class 3 |
| HDAC4 | 2q37.3 | 3 | 1.34 | 1 | 2.13 | 0.535 | 0.713 | (B) BRAF Class 3 |
| SBDS | 7q11.21 | 7 | 0.68 | 2 | 0.83 | 0.536 | 0.714 | (B) BRAF Class 3 |
| SIN3A | 15q24.2 | 3 | 1.91 | 1 | 3.03 | 0.537 | 0.714 | (B) BRAF Class 3 |
| HIST3H3 | 1q42.13 | 8 | 0.72 | 2 | 0.86 | 0.54 | 0.716 | (B) BRAF Class 3 |
| CCN6 | 6q21 | 17 | 2.50 | 3 | 2.13 | 0.54 | 0.716 | (A) BRAF Class 1 v12 |
| TCF7L1 | 2p11.2 | 8 | 1.78 | 1 | 1.11 | 0.541 | 0.716 | (A) BRAF Class 1 v12 |
| VHL | 3p25.3 | 26 | 0.83 | 6 | 0.86 | 0.542 | 0.716 | (B) BRAF Class 3 |
| PLK2 | 5q11.2 | 27 | 2.29 | 6 | 2.37 | 0.542 | 0.716 | (B) BRAF Class 3 |
| TYRO3 | 15q15.1 | 2 | 1.48 | 0 | 0.00 | 0.543 | 0.717 | (A) BRAF Class 1 v12 |
| CYLD | 16q12.1 | 49 | 2.36 | 11 | 2.29 | 0.544 | 0.717 | (A) BRAF Class 1 v12 |
| ZBTB7A | 19p13.3 | 12 | 3.60 | 2 | 4.17 | 0.546 | 0.718 | (B) BRAF Class 3 |
| SMARCD1 | 12q13.12 | 24 | 1.87 | 5 | 1.72 | 0.546 | 0.718 | (A) BRAF Class 1 v12 |
| RPA1 | 17p13.3 | 10 | 1.46 | 3 | 1.61 | 0.548 | 0.719 | (B) BRAF Class 3 |
| SAMD9 | 7q21.2 | 4 | 2.13 | 1 | 3.13 | 0.548 | 0.719 | (B) BRAF Class 3 |
| DMC1 | 22q13.1 | 10 | 1.71 | 2 | 1.35 | 0.55 | 0.72 | (A) BRAF Class 1 v12 |
| BCL2L11 | 2q13 | 8 | 0.68 | 2 | 0.79 | 0.551 | 0.72 | (B) BRAF Class 3 |
| NPRL2 | 3p21.31 | 3 | 1.14 | 1 | 1.72 | 0.551 | 0.72 | (B) BRAF Class 3 |
| BLM | 15q26.1 | 108 | 4.44 | 25 | 4.44 | 0.552 | 0.72 | (A) BRAF Class 1 v12 |
| SMC1A | Xp11.22 | 9 | 1.00 | 2 | 1.15 | 0.553 | 0.72 | (B) BRAF Class 3 |
| MDM2 | 12q15 | 43 | 1.69 | 10 | 1.70 | 0.553 | 0.721 | (B) BRAF Class 3 |
| FANCC | 9q22.32 | 38 | 1.54 | 9 | 1.55 | 0.555 | 0.721 | (B) BRAF Class 3 |
| ARFRP1 | 20q13.33 | 7 | 0.99 | 1 | 0.64 | 0.556 | 0.722 | (A) BRAF Class 1 v12 |
| PIK3R3 | 1p34.1 | 13 | 1.10 | 3 | 1.19 | 0.558 | 0.724 | (B) BRAF Class 3 |
| FLCN | 17p11.2 | 60 | 2.46 | 14 | 2.44 | 0.561 | 0.727 | (A) BRAF Class 1 v12 |
| MAP3K2 | 2q14.3 | 3 | 1.91 | 0 | 0.00 | 0.562 | 0.727 | (A) BRAF Class 1 v12 |
| DCUN1D1 | 3q26.33 | 7 | 0.59 | 1 | 0.39 | 0.562 | 0.727 | (A) BRAF Class 1 v12 |
| PGR | 11q22.1 | 67 | 6.04 | 14 | 6.01 | 0.564 | 0.728 | (A) BRAF Class 1 v12 |
| XRCC5 | 2q35 | 11 | 1.82 | 3 | 1.95 | 0.564 | 0.728 | (B) BRAF Class 3 |
| SF1 | 11q13.1 | 13 | 3.00 | 2 | 2.44 | 0.565 | 0.728 | (A) BRAF Class 1 v12 |
| CADM2 | 3p12.1 | 8 | 3.04 | 2 | 3.45 | 0.566 | 0.729 | (B) BRAF Class 3 |
| ELANE | 19p13.3 | 14 | 2.14 | 3 | 1.89 | 0.568 | 0.73 | (A) BRAF Class 1 v12 |
| PDK1 | 2q31.1 | 8 | 1.13 | 2 | 1.27 | 0.569 | 0.731 | (B) BRAF Class 3 |

| | | | | | | | | |
|---|---|---|---|---|---|---|---|---|
| HFE | 6p22.2 | 6 | 1.03 | 1 | 0.68 | 0.57 | 0.732 | (A) BRAF Class 1 v12 |
| HMBS | 11q23.3 | 7 | 1.20 | 2 | 1.35 | 0.572 | 0.732 | (B) BRAF Class 3 |
| NKX3-1 | 8p21.2 | 17 | 0.98 | 4 | 1.00 | 0.574 | 0.734 | (B) BRAF Class 3 |
| RAD54B | 8q22.1 | 17 | 2.91 | 4 | 2.70 | 0.576 | 0.736 | (A) BRAF Class 1 v12 |
| BMPR1A | 10q23.2 | 48 | 2.06 | 11 | 2.06 | 0.578 | 0.738 | (A) BRAF Class 1 v12 |
| GALNT12 | 9q22.33 | 13 | 1.76 | 3 | 1.58 | 0.579 | 0.738 | (A) BRAF Class 1 v12 |
| MAX | 14q23.3 | 15 | 0.79 | 3 | 0.71 | 0.579 | 0.738 | (A) BRAF Class 1 v12 |
| CD58 | 1p13.1 | 3 | 0.70 | 0 | 0.00 | 0.581 | 0.739 | (A) BRAF Class 1 v12 |
| FGF1 | 5q31.3 | 4 | 1.20 | 0 | 0.00 | 0.582 | 0.739 | (A) BRAF Class 1 v12 |
| STK19 | 6p21.33 | 12 | 2.28 | 3 | 2.36 | 0.583 | 0.739 | (B) BRAF Class 3 |
| TAP1 | 6p21.32 | 12 | 2.28 | 3 | 2.36 | 0.583 | 0.739 | (B) BRAF Class 3 |
| CDK9 | 9q34.11 | 3 | 0.85 | 0 | 0.00 | 0.585 | 0.741 | (A) BRAF Class 1 v12 |
| PSMD13 | 11p15.5 | 3 | 0.85 | 0 | 0.00 | 0.585 | 0.741 | (A) BRAF Class 1 v12 |
| MYBL1 | 8q13.1 | 30 | 2.69 | 7 | 2.66 | 0.588 | 0.742 | (A) BRAF Class 1 v12 |
| EXT2 | 11p11.2 | 18 | 1.76 | 4 | 1.66 | 0.589 | 0.742 | (A) BRAF Class 1 v12 |
| ENG | 9q34.11 | 13 | 2.23 | 3 | 2.03 | 0.589 | 0.742 | (A) BRAF Class 1 v12 |
| NFKBIA | 14q13.2 | 26 | 1.09 | 6 | 1.07 | 0.59 | 0.742 | (A) BRAF Class 1 v12 |
| MAFB | 20q12 | 3 | 0.44 | 1 | 0.58 | 0.591 | 0.742 | (B) BRAF Class 3 |
| GSTM5 | 1p13.3 | 4 | 1.13 | 1 | 1.45 | 0.591 | 0.742 | (B) BRAF Class 3 |
| RAC2 | 22q13.1 | 6 | 1.14 | 1 | 0.79 | 0.592 | 0.742 | (A) BRAF Class 1 v12 |
| SESN1 | 6q21 | 6 | 1.14 | 1 | 0.79 | 0.592 | 0.742 | (A) BRAF Class 1 v12 |
| SDHB | 1p36.13 | 10 | 0.41 | 2 | 0.35 | 0.593 | 0.742 | (A) BRAF Class 1 v12 |
| PRPF40B | 12q13.12 | 10 | 2.30 | 2 | 2.44 | 0.593 | 0.742 | (B) BRAF Class 3 |
| CD74 | 5q33.1 | 3 | 0.65 | 0 | 0.00 | 0.594 | 0.742 | (A) BRAF Class 1 v12 |
| FAAP20 | 1p36.33 | 3 | 0.51 | 1 | 0.68 | 0.596 | 0.743 | (B) BRAF Class 3 |
| PTPRK | 6q22.33 | 19 | 12.10 | 4 | 12.12 | 0.596 | 0.743 | (B) BRAF Class 3 |
| TRAF3 | 14q32.32 | 13 | 1.60 | 3 | 1.48 | 0.598 | 0.744 | (A) BRAF Class 1 v12 |
| ZBTB2 | 6q25.1 | 3 | 0.61 | 0 | 0.00 | 0.599 | 0.744 | (A) BRAF Class 1 v12 |
| RAB35 | 12q24.23 | 11 | 0.99 | 2 | 0.86 | 0.601 | 0.746 | (A) BRAF Class 1 v12 |
| EIF4A2 | 3q27.3 | 11 | 0.99 | 2 | 0.85 | 0.601 | 0.746 | (A) BRAF Class 1 v12 |
| EGFL7 | 9q34.3 | 15 | 1.27 | 3 | 1.19 | 0.603 | 0.747 | (A) BRAF Class 1 v12 |
| FAS | 10q23.31 | 18 | 1.23 | 4 | 1.18 | 0.604 | 0.748 | (A) BRAF Class 1 v12 |
| IL10 | 1q32.1 | 9 | 0.76 | 2 | 0.79 | 0.605 | 0.749 | (B) BRAF Class 3 |
| RPS6KA2 | 6q27 | 3 | 2.56 | 0 | 0.00 | 0.607 | 0.749 | (A) BRAF Class 1 v12 |
| TNK2 | 3q29 | 3 | 2.56 | 0 | 0.00 | 0.607 | 0.749 | (A) BRAF Class 1 v12 |
| CD79A | 19q13.2 | 22 | 1.46 | 5 | 1.45 | 0.609 | 0.75 | (A) BRAF Class 1 v12 |
| PPP4R2 | 3p13 | 8 | 1.52 | 2 | 1.57 | 0.609 | 0.75 | (B) BRAF Class 3 |
| HIST1H3F | 6p22.2 | 4 | 0.36 | 1 | 0.43 | 0.615 | 0.756 | (B) BRAF Class 3 |
| HIST1H3H | 6p22.1 | 4 | 0.36 | 1 | 0.43 | 0.615 | 0.756 | (B) BRAF Class 3 |
| DDX41 | 5q35.3 | 5 | 1.07 | 1 | 1.23 | 0.618 | 0.758 | (B) BRAF Class 3 |
| CLDN18 | 3q22.3 | 4 | 2.55 | 1 | 3.03 | 0.619 | 0.758 | (B) BRAF Class 3 |
| ZBTB20 | 3q13.31 | 4 | 2.55 | 1 | 3.03 | 0.619 | 0.758 | (B) BRAF Class 3 |
| RPL22 | 1p36.31 | 3 | 5.66 | 0 | 0.00 | 0.619 | 0.758 | (A) BRAF Class 1 v12 |
| CD22 | 19q13.12 | 2 | 1.72 | 0 | 0.00 | 0.622 | 0.759 | (A) BRAF Class 1 v12 |
| CACNA1E | 1q25.3 | 3 | 6.25 | 0 | 0.00 | 0.624 | 0.761 | (A) BRAF Class 1 v12 |
| UMODL1 | 21q22.3 | 3 | 6.25 | 0 | 0.00 | 0.624 | 0.761 | (A) BRAF Class 1 v12 |
| ABRAXAS1 | 4q21.23 | 17 | 0.89 | 4 | 0.88 | 0.625 | 0.761 | (A) BRAF Class 1 v12 |
| ABL2 | 1q25.2 | 20 | 3.00 | 4 | 2.99 | 0.626 | 0.761 | (A) BRAF Class 1 v12 |

| | | | | | | | | |
|---|---|---|---|---|---|---|---|---|
| USP7 | 16p13.2 | 2 | 2.33 | 0 | 0.00 | 0.633 | 0.769 | (A) BRAF Class 1 v12 |
| SQSTM1 | 5q35.3 | 4 | 1.52 | 1 | 1.72 | 0.633 | 0.769 | (B) BRAF Class 3 |
| FAAP24 | 19q13.11 | 2 | 0.34 | 0 | 0.00 | 0.636 | 0.771 | (A) BRAF Class 1 v12 |
| TAZ | Xq28 | 2 | 0.34 | 0 | 0.00 | 0.636 | 0.771 | (A) BRAF Class 1 v12 |
| DDR1 | 6p21.33 | 3 | 1.46 | 1 | 1.69 | 0.637 | 0.771 | (B) BRAF Class 3 |
| PAK1 | 1q13.5-q14. | 14 | 1.07 | 3 | 1.05 | 0.638 | 0.771 | (A) BRAF Class 1 v12 |
| PNKP | 19q13.33 | 12 | 2.05 | 3 | 2.03 | 0.64 | 0.773 | (A) BRAF Class 1 v12 |
| PMAIP1 | 18q21.32 | 6 | 0.51 | 1 | 0.40 | 0.641 | 0.773 | (A) BRAF Class 1 v12 |
| NUP214 | 9q34.13 | 5 | 3.82 | 1 | 4.17 | 0.642 | 0.773 | (B) BRAF Class 3 |
| HIST2H3D | 1q21.2 | 6 | 0.55 | 1 | 0.43 | 0.647 | 0.778 | (A) BRAF Class 1 v12 |
| RNF213 | 17q25.3 | 7 | 5.74 | 1 | 4.55 | 0.648 | 0.778 | (A) BRAF Class 1 v12 |
| PDPK1 | 16p13.3 | 6 | 0.48 | 1 | 0.38 | 0.649 | 0.778 | (A) BRAF Class 1 v12 |
| THADA | 2p21 | 3 | 1.92 | 0 | 0.00 | 0.649 | 0.778 | (A) BRAF Class 1 v12 |
| CHD1 | 5q15-q21.1 | 6 | 3.82 | 1 | 3.03 | 0.65 | 0.778 | (A) BRAF Class 1 v12 |
| HIST1H3J | 6p22.1 | 10 | 0.90 | 2 | 0.86 | 0.653 | 0.781 | (A) BRAF Class 1 v12 |
| FGF2 | 4q28.1 | 3 | 0.90 | 0 | 0.00 | 0.667 | 0.797 | (A) BRAF Class 1 v12 |
| EPB41L4A | ;q22.1-q22.2 | 2 | 0.06 | 0 | 0.00 | 0.668 | 0.797 | (A) BRAF Class 1 v12 |
| H19 | 11p15.5 | 2 | 0.06 | 0 | 0.00 | 0.668 | 0.797 | (A) BRAF Class 1 v12 |
| MAP3K7 | 6q15 | 2 | 0.79 | 0 | 0.00 | 0.674 | 0.803 | (A) BRAF Class 1 v12 |
| FGF10 | 5p12 | 9 | 1.27 | 2 | 1.27 | 0.677 | 0.805 | (A) BRAF Class 1 v12 |
| DUSP2 | 2q11.2 | 2 | 0.90 | 0 | 0.00 | 0.679 | 0.808 | (A) BRAF Class 1 v12 |
| CNOT3 | 19q13.42 | 2 | 1.17 | 0 | 0.00 | 0.682 | 0.809 | (A) BRAF Class 1 v12 |
| AJUBA | 14q11.2 | 2 | 1.27 | 0 | 0.00 | 0.682 | 0.809 | (A) BRAF Class 1 v12 |
| HIST1H3A | 6p22.2 | 2 | 0.18 | 0 | 0.00 | 0.683 | 0.809 | (A) BRAF Class 1 v12 |
| RPL26 | 17p13.1 | 2 | 0.56 | 0 | 0.00 | 0.7 | 0.829 | (A) BRAF Class 1 v12 |
| ASPSCR1 | 17q25.3 | 2 | 1.72 | 0 | 0.00 | 0.706 | 0.835 | (A) BRAF Class 1 v12 |
| OGA | 10q24.32 | 3 | 3.23 | 0 | 0.00 | 0.713 | 0.841 | (A) BRAF Class 1 v12 |
| ZFPM1 | 16q24.2 | 3 | 3.23 | 0 | 0.00 | 0.713 | 0.841 | (A) BRAF Class 1 v12 |
| DPYD | 1p21.3 | 4 | 1.62 | 1 | 1.52 | 0.715 | 0.841 | (A) BRAF Class 1 v12 |
| KNL1 | 15q15.1 | 2 | 1.64 | 0 | 0.00 | 0.717 | 0.841 | (A) BRAF Class 1 v12 |
| GUCY1A2 | 11q22.3 | 2 | 1.71 | 0 | 0.00 | 0.718 | 0.841 | (A) BRAF Class 1 v12 |
| MTR | 1q43 | 2 | 1.71 | 0 | 0.00 | 0.718 | 0.841 | (A) BRAF Class 1 v12 |
| NLRP1 | 17p13.2 | 2 | 1.71 | 0 | 0.00 | 0.718 | 0.841 | (A) BRAF Class 1 v12 |
| STK36 | 2q35 | 2 | 1.71 | 0 | 0.00 | 0.718 | 0.841 | (A) BRAF Class 1 v12 |
| TRIM24 | 7q33-q34 | 2 | 1.71 | 0 | 0.00 | 0.718 | 0.841 | (A) BRAF Class 1 v12 |
| EIF4E | 4q23 | 5 | 0.45 | 1 | 0.43 | 0.72 | 0.841 | (A) BRAF Class 1 v12 |
| HIST2H3C | 1q21.2 | 5 | 0.45 | 1 | 0.43 | 0.72 | 0.841 | (A) BRAF Class 1 v12 |
| MAF | 16q23.2 | 4 | 0.56 | 1 | 0.53 | 0.721 | 0.841 | (A) BRAF Class 1 v12 |
| TFRC | 3q29 | 7 | 2.07 | 1 | 2.04 | 0.732 | 0.854 | (A) BRAF Class 1 v12 |
| PCDHAC2 | 5q31.3 | 2 | 4.17 | 0 | 0.00 | 0.732 | 0.854 | (A) BRAF Class 1 v12 |
| TAL2 | 9q31.2 | 4 | 0.68 | 1 | 0.67 | 0.734 | 0.854 | (A) BRAF Class 1 v12 |
| CAMTA1 | )36.31-p36.2 | 2 | 1.24 | 0 | 0.00 | 0.749 | 0.867 | (A) BRAF Class 1 v12 |
| PHLPP2 | 16q22.2 | 1 | 0.76 | 0 | 0.00 | 0.771 | 0.867 | (A) BRAF Class 1 v12 |
| TPMT | 6p22.3 | 1 | 0.66 | 0 | 0.00 | 0.772 | 0.867 | (A) BRAF Class 1 v12 |
| HLA-C | 6p21.33 | 1 | 0.87 | 0 | 0.00 | 0.788 | 0.867 | (A) BRAF Class 1 v12 |
| PTPRO | 12.3\|12p13- | 1 | 0.86 | 0 | 0.00 | 0.789 | 0.867 | (A) BRAF Class 1 v12 |
| UGT1A1 | 2q37.1 | 1 | 0.40 | 0 | 0.00 | 0.789 | 0.867 | (A) BRAF Class 1 v12 |
| CDH5 | 16q21 | 1 | 0.50 | 0 | 0.00 | 0.794 | 0.867 | (A) BRAF Class 1 v12 |

| | | | | | | | | |
|---|---|---|---|---|---|---|---|---|
| SGK1 | 6q23.2 | 1 | 0.66 | 0 | 0.00 | 0.795 | 0.867 | (A) BRAF Class 1 v12 |
| CDKN2B-AS1 | 9p21.3 | 1 | 0.26 | 0 | 0.00 | 0.796 | 0.867 | (A) BRAF Class 1 v12 |
| APOBEC3G | 22q13.1 | 1 | 1.16 | 0 | 0.00 | 0.796 | 0.867 | (A) BRAF Class 1 v12 |
| DUSP6 | 12q21.33 | 1 | 1.16 | 0 | 0.00 | 0.796 | 0.867 | (A) BRAF Class 1 v12 |
| HSPA5 | 9q33.3 | 1 | 1.16 | 0 | 0.00 | 0.796 | 0.867 | (A) BRAF Class 1 v12 |
| SLX1B | 16p11.2 | 1 | 0.17 | 0 | 0.00 | 0.798 | 0.867 | (A) BRAF Class 1 v12 |
| RPL10 | Xq28 | 1 | 1.05 | 0 | 0.00 | 0.798 | 0.867 | (A) BRAF Class 1 v12 |
| ABCB4 | 7q21.12 | 2 | 2.15 | 0 | 0.00 | 0.799 | 0.867 | (A) BRAF Class 1 v12 |
| MAML3 | 4q31.1 | 2 | 2.15 | 0 | 0.00 | 0.799 | 0.867 | (A) BRAF Class 1 v12 |
| PTP4A1 | 6q12 | 1 | 0.19 | 0 | 0.00 | 0.806 | 0.867 | (A) BRAF Class 1 v12 |
| ABCA17P | 16p13.3 | 1 | 0.03 | 0 | 0.00 | 0.817 | 0.867 | (A) BRAF Class 1 v12 |
| ABR | 17p13.3 | 1 | 0.03 | 0 | 0.00 | 0.817 | 0.867 | (A) BRAF Class 1 v12 |
| ANKLE2 | 12q24.33 | 1 | 0.03 | 0 | 0.00 | 0.817 | 0.867 | (A) BRAF Class 1 v12 |
| ANKRD30A | 10p11.21 | 1 | 0.03 | 0 | 0.00 | 0.817 | 0.867 | (A) BRAF Class 1 v12 |
| AP1G1 | 16q22.2 | 1 | 0.03 | 0 | 0.00 | 0.817 | 0.867 | (A) BRAF Class 1 v12 |
| APBA1 | 9q21.12 | 1 | 0.03 | 0 | 0.00 | 0.817 | 0.867 | (A) BRAF Class 1 v12 |
| ARHGAP4 | Xq28 | 1 | 0.03 | 0 | 0.00 | 0.817 | 0.867 | (A) BRAF Class 1 v12 |
| ARSB | 5q14.1 | 1 | 0.03 | 0 | 0.00 | 0.817 | 0.867 | (A) BRAF Class 1 v12 |
| C2ORF83 | 2q36.3 | 1 | 0.03 | 0 | 0.00 | 0.817 | 0.867 | (A) BRAF Class 1 v12 |
| CASZ1 | 1p36.22 | 1 | 0.03 | 0 | 0.00 | 0.817 | 0.867 | (A) BRAF Class 1 v12 |
| CLIP1 | 12q24.31 | 1 | 0.03 | 0 | 0.00 | 0.817 | 0.867 | (A) BRAF Class 1 v12 |
| CPSF1 | 8q24.3 | 1 | 0.03 | 0 | 0.00 | 0.817 | 0.867 | (A) BRAF Class 1 v12 |
| CTDSP1 | 2q35 | 1 | 0.03 | 0 | 0.00 | 0.817 | 0.867 | (A) BRAF Class 1 v12 |
| CTRC | 1p36.21 | 1 | 0.03 | 0 | 0.00 | 0.817 | 0.867 | (A) BRAF Class 1 v12 |
| CTSB | 8p23.1 | 1 | 0.03 | 0 | 0.00 | 0.817 | 0.867 | (A) BRAF Class 1 v12 |
| CUL1 | 7q36.1 | 1 | 0.03 | 0 | 0.00 | 0.817 | 0.867 | (A) BRAF Class 1 v12 |
| DCBLD1 | 6q22.1 | 1 | 0.03 | 0 | 0.00 | 0.817 | 0.867 | (A) BRAF Class 1 v12 |
| DDAH1 | 1p22.3 | 1 | 0.03 | 0 | 0.00 | 0.817 | 0.867 | (A) BRAF Class 1 v12 |
| DEF8 | 16q24.3 | 1 | 0.03 | 0 | 0.00 | 0.817 | 0.867 | (A) BRAF Class 1 v12 |
| DENND4C | 9p22.1 | 1 | 0.03 | 0 | 0.00 | 0.817 | 0.867 | (A) BRAF Class 1 v12 |
| DHRSX | 2.33 and Yp1 | 1 | 0.03 | 0 | 0.00 | 0.817 | 0.867 | (A) BRAF Class 1 v12 |
| DRICH1 | 22q11.23 | 1 | 0.03 | 0 | 0.00 | 0.817 | 0.867 | (A) BRAF Class 1 v12 |
| EIF3E | 8q23.1 | 1 | 0.03 | 0 | 0.00 | 0.817 | 0.867 | (A) BRAF Class 1 v12 |
| ELAC2 | 17p12 | 1 | 0.03 | 0 | 0.00 | 0.817 | 0.867 | (A) BRAF Class 1 v12 |
| EMID1 | 22q12.2 | 1 | 0.03 | 0 | 0.00 | 0.817 | 0.867 | (A) BRAF Class 1 v12 |
| F11-AS1 | 4q35.2 | 1 | 0.03 | 0 | 0.00 | 0.817 | 0.867 | (A) BRAF Class 1 v12 |
| FKBP9P1 | 7p11.2 | 1 | 0.03 | 0 | 0.00 | 0.817 | 0.867 | (A) BRAF Class 1 v12 |
| GALNT11 | q36.1\|7q36. | 1 | 0.03 | 0 | 0.00 | 0.817 | 0.867 | (A) BRAF Class 1 v12 |
| GOLT1A | 1q32.1 | 1 | 0.03 | 0 | 0.00 | 0.817 | 0.867 | (A) BRAF Class 1 v12 |
| GRAPL | 17p11.2 | 1 | 0.03 | 0 | 0.00 | 0.817 | 0.867 | (A) BRAF Class 1 v12 |
| GRIA1 | 5q33.2 | 1 | 0.03 | 0 | 0.00 | 0.817 | 0.867 | (A) BRAF Class 1 v12 |
| GRIK1 | 21q21.3 | 1 | 0.03 | 0 | 0.00 | 0.817 | 0.867 | (A) BRAF Class 1 v12 |
| HECTD4 | 12q24.13 | 1 | 0.03 | 0 | 0.00 | 0.817 | 0.867 | (A) BRAF Class 1 v12 |
| HLA-DOB | 6p21.32 | 1 | 0.03 | 0 | 0.00 | 0.817 | 0.867 | (A) BRAF Class 1 v12 |
| HOOK2 | 19p13.13 | 1 | 0.03 | 0 | 0.00 | 0.817 | 0.867 | (A) BRAF Class 1 v12 |
| HSD17B6 | 12q13.3 | 1 | 0.03 | 0 | 0.00 | 0.817 | 0.867 | (A) BRAF Class 1 v12 |
| IGHE | 14q32.33 | 1 | 0.03 | 0 | 0.00 | 0.817 | 0.867 | (A) BRAF Class 1 v12 |
| LDLR | 19p13.2 | 1 | 0.03 | 0 | 0.00 | 0.817 | 0.867 | (A) BRAF Class 1 v12 |

| | | | | | | | | |
|---|---|---|---|---|---|---|---|---|
| OC1053744€ | 2p22.2 | 1 | 0.03 | 0 | 0.00 | 0.817 | 0.867 | (A) BRAF Class 1 v12 |
| LTBP4 | 19q13.2 | 1 | 0.03 | 0 | 0.00 | 0.817 | 0.867 | (A) BRAF Class 1 v12 |
| MYEOV | 11q13.3 | 1 | 0.03 | 0 | 0.00 | 0.817 | 0.867 | (A) BRAF Class 1 v12 |
| MYO1D | 17q11.2 | 1 | 0.03 | 0 | 0.00 | 0.817 | 0.867 | (A) BRAF Class 1 v12 |
| MYSM1 | 1p32.1 | 1 | 0.03 | 0 | 0.00 | 0.817 | 0.867 | (A) BRAF Class 1 v12 |
| NBPF20 | 1q21.1 | 1 | 0.03 | 0 | 0.00 | 0.817 | 0.867 | (A) BRAF Class 1 v12 |
| NDUFB2 | 7q34 | 1 | 0.03 | 0 | 0.00 | 0.817 | 0.867 | (A) BRAF Class 1 v12 |
| NELFA | 4p16.3 | 1 | 0.03 | 0 | 0.00 | 0.817 | 0.867 | (A) BRAF Class 1 v12 |
| PADI2 | 1p36.13 | 1 | 0.03 | 0 | 0.00 | 0.817 | 0.867 | (A) BRAF Class 1 v12 |
| PAX9 | 14q13.3 | 1 | 0.03 | 0 | 0.00 | 0.817 | 0.867 | (A) BRAF Class 1 v12 |
| PFKFB4 | 3p21.31 | 1 | 0.03 | 0 | 0.00 | 0.817 | 0.867 | (A) BRAF Class 1 v12 |
| PKD1 | 16p13.3 | 1 | 0.03 | 0 | 0.00 | 0.817 | 0.867 | (A) BRAF Class 1 v12 |
| PLEKHA7 | 1p15.2-p15. | 1 | 0.03 | 0 | 0.00 | 0.817 | 0.867 | (A) BRAF Class 1 v12 |
| PLVAP | 19p13.11 | 1 | 0.03 | 0 | 0.00 | 0.817 | 0.867 | (A) BRAF Class 1 v12 |
| PNPLA6 | 19p13.2 | 1 | 0.03 | 0 | 0.00 | 0.817 | 0.867 | (A) BRAF Class 1 v12 |
| PRMT6 | 1p13.3 | 1 | 0.03 | 0 | 0.00 | 0.817 | 0.867 | (A) BRAF Class 1 v12 |
| PTBP3 | 9q32 | 1 | 0.03 | 0 | 0.00 | 0.817 | 0.867 | (A) BRAF Class 1 v12 |
| RAB2A | 8q12.1-q12.2 | 1 | 0.03 | 0 | 0.00 | 0.817 | 0.867 | (A) BRAF Class 1 v12 |
| SAE1 | 19q13.32 | 1 | 0.03 | 0 | 0.00 | 0.817 | 0.867 | (A) BRAF Class 1 v12 |
| SCAP | 3p21.31 | 1 | 0.03 | 0 | 0.00 | 0.817 | 0.867 | (A) BRAF Class 1 v12 |
| SCARA5 | 8p21.1 | 1 | 0.03 | 0 | 0.00 | 0.817 | 0.867 | (A) BRAF Class 1 v12 |
| SDCCAG8 | 1q43-q44 | 1 | 0.03 | 0 | 0.00 | 0.817 | 0.867 | (A) BRAF Class 1 v12 |
| SEMA6D | 15q21.1 | 1 | 0.03 | 0 | 0.00 | 0.817 | 0.867 | (A) BRAF Class 1 v12 |
| SIK3 | 11q23.3 | 1 | 0.03 | 0 | 0.00 | 0.817 | 0.867 | (A) BRAF Class 1 v12 |
| SLC30A5 | 5q13.1-q13.2 | 1 | 0.03 | 0 | 0.00 | 0.817 | 0.867 | (A) BRAF Class 1 v12 |
| SND1 | 7q32.1 | 1 | 0.03 | 0 | 0.00 | 0.817 | 0.867 | (A) BRAF Class 1 v12 |
| SPG11 | 15q21.1 | 1 | 0.03 | 0 | 0.00 | 0.817 | 0.867 | (A) BRAF Class 1 v12 |
| TCAF2P1 | 7q35 | 1 | 0.03 | 0 | 0.00 | 0.817 | 0.867 | (A) BRAF Class 1 v12 |
| TRAJ32 | 14q11.2 | 1 | 0.03 | 0 | 0.00 | 0.817 | 0.867 | (A) BRAF Class 1 v12 |
| TRAJ42 | 14q11.2 | 1 | 0.03 | 0 | 0.00 | 0.817 | 0.867 | (A) BRAF Class 1 v12 |
| TRAJ53 | 14q11.2 | 1 | 0.03 | 0 | 0.00 | 0.817 | 0.867 | (A) BRAF Class 1 v12 |
| TRBJ2-2 | 7q34 | 1 | 0.03 | 0 | 0.00 | 0.817 | 0.867 | (A) BRAF Class 1 v12 |
| TRGJP | 7p14.1 | 1 | 0.03 | 0 | 0.00 | 0.817 | 0.867 | (A) BRAF Class 1 v12 |
| TSPOAP1 | 17q22 | 1 | 0.03 | 0 | 0.00 | 0.817 | 0.867 | (A) BRAF Class 1 v12 |
| UGT1A6 | 2q37.1 | 1 | 0.03 | 0 | 0.00 | 0.817 | 0.867 | (A) BRAF Class 1 v12 |
| ZNF423 | 16q12.1 | 1 | 0.03 | 0 | 0.00 | 0.817 | 0.867 | (A) BRAF Class 1 v12 |
| ZNF599 | 19q13.11 | 1 | 0.03 | 0 | 0.00 | 0.817 | 0.867 | (A) BRAF Class 1 v12 |
| BCL2L2 | 14q11.2 | 1 | 0.14 | 0 | 0.00 | 0.819 | 0.868 | (A) BRAF Class 1 v12 |
| GID4 | p11.2\|17p1 | 1 | 0.17 | 0 | 0.00 | 0.822 | 0.871 | (A) BRAF Class 1 v12 |
| HSP90AB1 | 6p21.1 | 1 | 0.40 | 0 | 0.00 | 0.824 | 0.872 | (A) BRAF Class 1 v12 |
| MGMT | 10q26.3 | 1 | 0.64 | 0 | 0.00 | 0.826 | 0.873 | (A) BRAF Class 1 v12 |
| MAP3K14 | 17q21.31 | 1 | 0.09 | 0 | 0.00 | 0.826 | 0.873 | (A) BRAF Class 1 v12 |
| IRF8 | 16q24.1 | 1 | 0.81 | 0 | 0.00 | 0.838 | 0.876 | (A) BRAF Class 1 v12 |
| MRTFA | 2q13.1-q13. | 1 | 0.86 | 0 | 0.00 | 0.841 | 0.876 | (A) BRAF Class 1 v12 |
| PRDM16 | 1p36.32 | 1 | 0.86 | 0 | 0.00 | 0.841 | 0.876 | (A) BRAF Class 1 v12 |
| SPI1 | 11p11.2 | 1 | 0.90 | 0 | 0.00 | 0.841 | 0.876 | (A) BRAF Class 1 v12 |
| FOXO3 | 6q21 | 1 | 0.81 | 0 | 0.00 | 0.842 | 0.876 | (A) BRAF Class 1 v12 |
| ARNT | 1q21.3 | 1 | 0.82 | 0 | 0.00 | 0.847 | 0.876 | (A) BRAF Class 1 v12 |

| | | | | | | | | |
|---|---|---|---|---|---|---|---|---|
| MUC1 | 1q22 | 1 | 0.82 | 0 | 0.00 | 0.847 | 0.876 | (A) BRAF Class 1 v12 |
| NFKB2 | 10q24.32 | 1 | 0.82 | 0 | 0.00 | 0.847 | 0.876 | (A) BRAF Class 1 v12 |
| PER1 | 17p13.1 | 1 | 0.82 | 0 | 0.00 | 0.847 | 0.876 | (A) BRAF Class 1 v12 |
| PSIP1 | 9p22.3 | 1 | 0.82 | 0 | 0.00 | 0.847 | 0.876 | (A) BRAF Class 1 v12 |
| TLX1 | 10q24.31 | 1 | 0.82 | 0 | 0.00 | 0.847 | 0.876 | (A) BRAF Class 1 v12 |
| TPR | 1q31.1 | 1 | 0.82 | 0 | 0.00 | 0.847 | 0.876 | (A) BRAF Class 1 v12 |
| ZNF384 | 12p13.31 | 1 | 0.82 | 0 | 0.00 | 0.847 | 0.876 | (A) BRAF Class 1 v12 |
| AURKC | 19q13.43 | 1 | 0.85 | 0 | 0.00 | 0.848 | 0.876 | (A) BRAF Class 1 v12 |
| BLNK | 10q24.1 | 1 | 0.85 | 0 | 0.00 | 0.848 | 0.876 | (A) BRAF Class 1 v12 |
| CYP2C19 | 10q23.33 | 1 | 0.85 | 0 | 0.00 | 0.848 | 0.876 | (A) BRAF Class 1 v12 |
| DST | 6p12.1 | 1 | 0.85 | 0 | 0.00 | 0.848 | 0.876 | (A) BRAF Class 1 v12 |
| GDNF | 5p13.2 | 1 | 0.85 | 0 | 0.00 | 0.848 | 0.876 | (A) BRAF Class 1 v12 |
| IKBKB | 8p11.21 | 1 | 0.85 | 0 | 0.00 | 0.848 | 0.876 | (A) BRAF Class 1 v12 |
| ITGA9 | 3p22.2 | 1 | 0.85 | 0 | 0.00 | 0.848 | 0.876 | (A) BRAF Class 1 v12 |
| ITGB3 | 17q21.32 | 1 | 0.85 | 0 | 0.00 | 0.848 | 0.876 | (A) BRAF Class 1 v12 |
| PKHD1 | [illegible]p12.3-p12.[illegible] | 1 | 0.85 | 0 | 0.00 | 0.848 | 0.876 | (A) BRAF Class 1 v12 |
| PLEKHG5 | 1p36.31 | 1 | 0.85 | 0 | 0.00 | 0.848 | 0.876 | (A) BRAF Class 1 v12 |
| RRM1 | 11p15.4 | 1 | 0.85 | 0 | 0.00 | 0.848 | 0.876 | (A) BRAF Class 1 v12 |
| LCK | 1p35.2 | 1 | 0.80 | 0 | 0.00 | 0.85 | 0.877 | (A) BRAF Class 1 v12 |
| TCF12 | 15q21.3 | 1 | 0.50 | 0 | 0.00 | 0.852 | 0.877 | (A) BRAF Class 1 v12 |
| BIRC5 | 17q25.3 | 1 | 0.53 | 0 | 0.00 | 0.855 | 0.877 | (A) BRAF Class 1 v12 |
| MLF1 | 3q25.32 | 1 | 1.89 | 0 | 0.00 | 0.855 | 0.877 | (A) BRAF Class 1 v12 |
| ARFGAP3 | 22q13.2 | 1 | 2.08 | 0 | 0.00 | 0.857 | 0.877 | (A) BRAF Class 1 v12 |
| GNAI3 | 1p13.3 | 1 | 2.08 | 0 | 0.00 | 0.857 | 0.877 | (A) BRAF Class 1 v12 |
| SNX31 | 8q22.3 | 1 | 2.08 | 0 | 0.00 | 0.857 | 0.877 | (A) BRAF Class 1 v12 |
| TLR2 | 4q31.3 | 1 | 2.08 | 0 | 0.00 | 0.857 | 0.877 | (A) BRAF Class 1 v12 |
| TMEM216 | 11q12.2 | 1 | 2.08 | 0 | 0.00 | 0.857 | 0.877 | (A) BRAF Class 1 v12 |
| TBL1XR1 | 3q26.32 | 1 | 0.97 | 0 | 0.00 | 0.858 | 0.877 | (A) BRAF Class 1 v12 |
| PC | 11q13.2 | 1 | 1.16 | 0 | 0.00 | 0.86 | 0.877 | (A) BRAF Class 1 v12 |
| PCLO | 7q21.11 | 1 | 1.16 | 0 | 0.00 | 0.86 | 0.877 | (A) BRAF Class 1 v12 |
| PDCD11 | 10q24.33 | 1 | 1.16 | 0 | 0.00 | 0.86 | 0.877 | (A) BRAF Class 1 v12 |
| SP140 | 2q37.1 | 1 | 1.96 | 0 | 0.00 | 0.864 | 0.881 | (A) BRAF Class 1 v12 |
| HIP1 | 7q11.23 | 1 | 1.11 | 0 | 0.00 | 0.865 | 0.881 | (A) BRAF Class 1 v12 |
| ETNK1 | 12p12.1 | 1 | 0.63 | 0 | 0.00 | 0.869 | 0.884 | (A) BRAF Class 1 v12 |
| CD3EAP | 19q13.32 | 1 | 0.30 | 0 | 0.00 | 0.874 | 0.888 | (A) BRAF Class 1 v12 |
| TSPAN31 | 12q14.1 | 1 | 0.30 | 0 | 0.00 | 0.874 | 0.888 | (A) BRAF Class 1 v12 |
| MLLT1 | 19p13.3 | 1 | 1.59 | 0 | 0.00 | 0.875 | 0.888 | (A) BRAF Class 1 v12 |
| KIF3A | 5q31.1 | 1 | 2.33 | 0 | 0.00 | 0.878 | 0.89 | (A) BRAF Class 1 v12 |
| BEND2 | Xp22.13 | 1 | 1.08 | 0 | 0.00 | 0.894 | 0.902 | (A) BRAF Class 1 v12 |
| FOSB | 19q13.32 | 1 | 1.08 | 0 | 0.00 | 0.894 | 0.902 | (A) BRAF Class 1 v12 |
| GLMN | 1p22.1 | 1 | 1.08 | 0 | 0.00 | 0.894 | 0.902 | (A) BRAF Class 1 v12 |
| OTX2 | 14q22.3 | 1 | 1.08 | 0 | 0.00 | 0.894 | 0.902 | (A) BRAF Class 1 v12 |
| TBXT | 6q27 | 1 | 1.08 | 0 | 0.00 | 0.894 | 0.902 | (A) BRAF Class 1 v12 |
| KBTBD4 | 11p11.2 | 1 | 0.97 | 0 | 0.00 | 0.904 | 0.911 | (A) BRAF Class 1 v12 |
| MAGOH | 1p32.3 | 1 | 0.99 | 0 | 0.00 | 0.918 | 0.925 | (A) BRAF Class 1 v12 |
| CBLC | 19q13.32 | 1 | 2.78 | 0 | 0.00 | 0.923 | 0.928 | (A) BRAF Class 1 v12 |
| LRIG3 | 12q14.1 | 1 | 2.78 | 0 | 0.00 | 0.923 | 0.928 | (A) BRAF Class 1 v12 |
| MCM7 | 7q22.1 | 1 | 3.23 | 0 | 0.00 | 0.939 | 0.944 | (A) BRAF Class 1 v12 |

| | | | | | | | | |
|---|---|---|---|---|---|---|---|---|
| TET3 | 2p13.1 | 1 | 2.94 | 0 | 0.00 | 0.944 | 0.948 | (A) BRAF Class 1 v12 |
| GNB1 | 1p36.33 | 1 | 2.27 | 0 | 0.00 | 0.957 | 0.959 | (A) BRAF Class 1 v12 |
| TOE1 | 1p34.1 | 0 | 0.00 | 1 | 0.13 | 0.959 | 0.961 | (B) BRAF Class 3 |
| OLIG2 | 21q22.11 | 1 | 3.70 | 0 | 0.00 | 0.964 | 0.965 | (A) BRAF Class 1 v12 |
| BRAF | 7q34 | 3348 | 100.00 | 749 | 100.00 | 1 | 1 | (B) BRAF Class 3 |

#### Supplementary Table 6

| Gene | cytoband | Class 1 occurrence | Class 1 % | Class 3 occurrence | Class 3 % | SumClass | pvalue | qvalue | Enriched In |
|---|---|---|---|---|---|---|---|---|---|
| KIT | 4q12 | 80 | 2.39 | 33 | 4.41 | 113 | 0.002742 | 0.0482 | (B) BRAF Class 3 |
| EGFR | 7p11.2 | 134 | 4.00 | 49 | 6.57 | 183 | 0.002211 | 0.0401 | (B) BRAF Class 3 |
| MAP3K13 | 3q27.2 | 20 | 1.52 | 14 | 4.55 | 34 | 0.002103 | 0.0388 | (B) BRAF Class 3 |
| FGFR4 | 5q35.2 | 71 | 2.76 | 32 | 5.30 | 103 | 0.001996 | 0.0374 | (B) BRAF Class 3 |
| ZRSR2 | Xp22.2 | 41 | 1.83 | 1 | 0.20 | 42 | 0.001821 | 0.0348 | (A) BRAF Class 1 v12 |
| RUNX1T1 | 8q21.3 | 57 | 4.56 | 27 | 9.34 | 84 | 0.001796 | 0.0348 | (B) BRAF Class 3 |
| MSH3 | 5q14.1 | 94 | 8.82 | 8 | 3.35 | 102 | 0.001712 | 0.0338 | (A) BRAF Class 1 v12 |
| ANKRD26 | 10p12.1 | 45 | 13.51 | 0 | 0.00 | 45 | 0.001551 | 0.0312 | (A) BRAF Class 1 v12 |
| BCOR | Xp11.4 | 157 | 6.40 | 19 | 3.26 | 176 | 0.001469 | 0.0301 | (A) BRAF Class 1 v12 |
| NTRK3 | 15q25.3 | 112 | 4.41 | 45 | 7.64 | 157 | 0.001392 | 0.0291 | (B) BRAF Class 3 |
| SMAD3 | 15q22.33 | 25 | 1.70 | 16 | 4.85 | 41 | 0.001331 | 0.0283 | (B) BRAF Class 3 |
| TP63 | 3q28 | 57 | 4.57 | 26 | 9.74 | 83 | 0.001246 | 0.0282 | (B) BRAF Class 3 |
| CBL | 11q23.3 | 57 | 2.22 | 28 | 4.67 | 85 | 0.001277 | 0.0282 | (B) BRAF Class 3 |
| ASXL1 | 20q11.21 | 132 | 5.30 | 52 | 8.81 | 184 | 0.0013 | 0.0282 | (B) BRAF Class 3 |
| POT1 | 7q31.33 | 13 | 1.54 | 12 | 5.83 | 25 | 0.001137 | 0.0262 | (B) BRAF Class 3 |
| PMS1 | 2q32.2 | 55 | 2.38 | 27 | 5.14 | 82 | 0.00112 | 0.0262 | (B) BRAF Class 3 |
| DICER1 | 14q32.13 | 91 | 3.78 | 39 | 7.01 | 130 | 0.001033 | 0.0254 | (B) BRAF Class 3 |
| KDR | 4q12 | 162 | 5.08 | 59 | 8.38 | 221 | 0.000714 | 0.0184 | (B) BRAF Class 3 |
| KMT2D | 12q13.12 | 362 | 14.73 | 118 | 20.34 | 480 | 0.000703 | 0.0184 | (B) BRAF Class 3 |
| CARD11 | 7p22.2 | 157 | 6.38 | 61 | 10.52 | 218 | 0.000581 | 0.0157 | (B) BRAF Class 3 |
| NOTCH1 | 9q34.3 | 178 | 5.42 | 64 | 8.80 | 242 | 0.00058 | 0.0157 | (B) BRAF Class 3 |
| HLA-A | 6p22.1 | 48 | 3.95 | 1 | 0.37 | 49 | 0.000533 | 0.0151 | (A) BRAF Class 1 v12 |
| ARID2 | 12q12 | 180 | 7.41 | 67 | 11.90 | 247 | 0.000525 | 0.0151 | (B) BRAF Class 3 |
| FGFR2 | 10q26.13 | 89 | 2.66 | 39 | 5.23 | 128 | 0.000439 | 0.0135 | (B) BRAF Class 3 |
| PIK3CA | 3q26.32 | 311 | 9.29 | 102 | 13.67 | 413 | 0.00032 | 0.0101 | (B) BRAF Class 3 |
| RET | 10q11.21 | 123 | 3.67 | 50 | 6.70 | 173 | 0.000301 | 0.009808 | (B) BRAF Class 3 |
| SRC | 20q11.23 | 22 | 0.83 | 16 | 2.86 | 38 | 0.000289 | 0.009684 | (B) BRAF Class 3 |
| FLT1 | 13q12.3 | 132 | 5.37 | 55 | 9.48 | 187 | 0.000281 | 0.009684 | (B) BRAF Class 3 |
| CBLB | 3q13.11 | 16 | 1.51 | 15 | 5.88 | 31 | 0.000218 | 0.007776 | (B) BRAF Class 3 |
| BRIP1 | 17q23.2 | 65 | 2.64 | 34 | 5.86 | 99 | 0.00019 | 0.006998 | (B) BRAF Class 3 |
| SYNE1 | 6q25.2 | 11 | 2.14 | 11 | 11.34 | 22 | 0.000149 | 0.005777 | (B) BRAF Class 3 |
| RAF1 | 3p25.2 | 37 | 1.42 | 24 | 3.95 | 61 | 0.000149 | 0.005777 | (B) BRAF Class 3 |
| TERT | 5p15.33 | 842 | 34.20 | 145 | 26.22 | 987 | 0.000151 | 0.005777 | (A) BRAF Class 1 v12 |
| EPHA3 | 3p11.1 | 92 | 4.85 | 43 | 9.95 | 135 | 8.85E-05 | 0.003769 | (B) BRAF Class 3 |
| PDGFRA | 4q12 | 102 | 3.13 | 46 | 6.33 | 148 | 8.11E-05 | 0.00369 | (B) BRAF Class 3 |
| PTPRT | [illegible]q12-q13.[illegible] | 160 | 11.36 | 61 | 19.87 | 221 | 8.33E-05 | 0.00369 | (B) BRAF Class 3 |
| PTPRD | [illegible]p24.1-p2[illegible] | 184 | 10.62 | 71 | 17.93 | 255 | 7.47E-05 | 0.00369 | (B) BRAF Class 3 |
| ATM | 11q22.3 | 260 | 8.16 | 91 | 12.93 | 351 | 8.16E-05 | 0.00369 | (B) BRAF Class 3 |
| ERBB3 | 12q13.2 | 81 | 3.15 | 41 | 6.82 | 122 | 6.11E-05 | 0.003221 | (B) BRAF Class 3 |
| ARID1A | 1p36.11 | 269 | 10.59 | 99 | 16.81 | 368 | 3.24E-05 | 0.001793 | (B) BRAF Class 3 |
| TET2 | 4q24 | 155 | 6.23 | 68 | 11.53 | 223 | 1.62E-05 | 0.000946 | (B) BRAF Class 3 |
| MCL1 | 1q21.2 | 25 | 1.02 | 22 | 3.80 | 47 | 1.22E-05 | 0.000748 | (B) BRAF Class 3 |
| FLT3 | 13q12.2 | 100 | 3.10 | 49 | 6.87 | 149 | 7.08E-06 | 0.000461 | (B) BRAF Class 3 |
| PPP2R1A | 19q13.41 | 34 | 1.54 | 27 | 5.19 | 61 | 4.61E-06 | 0.000319 | (B) BRAF Class 3 |
| ERBB2 | 17q12 | 88 | 2.63 | 46 | 6.19 | 134 | 4.21E-06 | 0.00031 | (B) BRAF Class 3 |
| MET | 7q31.2 | 121 | 3.65 | 57 | 7.71 | 178 | 3.88E-06 | 0.000307 | (B) BRAF Class 3 |

| | | | | | | | | | |
|---|---|---|---|---|---|---|---|---|---|
| ERBB4 | 2q34 | 206 | 6.36 | 83 | 11.53 | 289 | 3.34E-06 | 0.000284 | (B) BRAF Class 3 |
| PTPN11 | 12q24.13 | 35 | 1.08 | 27 | 3.75 | 62 | 2.75E-06 | 0.000253 | (B) BRAF Class 3 |
| NOTCH2 | 1p12 | 109 | 4.36 | 57 | 10.00 | 166 | 4.77E-07 | 4.8E-05 | (B) BRAF Class 3 |
| SMARCA4 | 19p13.2 | 159 | 6.26 | 76 | 12.90 | 235 | 1.73E-07 | 1.92E-05 | (B) BRAF Class 3 |
| RNF43 | 17q22 | 259 | 11.64 | 21 | 4.02 | 280 | 1.14E-08 | 1.41E-06 | (A) BRAF Class 1 v12 |
| HRAS | 11p15.5 | 17 | 0.51 | 26 | 3.48 | 43 | 7.73E-10 | 1.07E-07 | (B) BRAF Class 3 |
| APC | 5q22.2 | 356 | 11.33 | 160 | 22.63 | 516 | 3.86E-14 | 6.1E-12 | (B) BRAF Class 3 |
| KEAP1 | 19p13.2 | 53 | 2.24 | 63 | 11.07 | 116 | 1.03E-17 | 1.91E-15 | (B) BRAF Class 3 |
| TP53 | 17p13.1 | 892 | 26.64 | 328 | 44.15 | 1220 | 2.97E-20 | 6.58E-18 | (B) BRAF Class 3 |
| STK11 | 19p13.3 | 57 | 1.74 | 79 | 10.93 | 136 | 1.91E-26 | 5.29E-24 | (B) BRAF Class 3 |
| NF1 | 17q11.2 | 144 | 5.66 | 141 | 23.62 | 285 | 7.45E-35 | 2.75E-32 | (B) BRAF Class 3 |
| NRAS | 1p13.2 | 34 | 1.02 | 94 | 12.55 | 128 | 8.13E-44 | 4.5E-41 | (B) BRAF Class 3 |
| KRAS | 12p12.1 | 36 | 1.08 | 129 | 17.22 | 165 | 2.62E-66 | 2.9E-63 | (B) BRAF Class 3 |

#### Supplementary Table 7

| Gene | Cytoband | Class 1 Occurrence | Class 1 % | Class 2 occurrence | Class 2 % | pvalue | qvalue | Enriched In |
|---|---|---|---|---|---|---|---|---|
| NF1 | 17q11.2 | 50 | 6.19 | 36 | 27.07 | 2.1E-11 | 1.9E-08 | (B) BRAF Class 2 Melanoma |
| FLT3 | 13q12.2 | 41 | 3.70 | 28 | 19.05 | 2E-10 | 9.1E-08 | (B) BRAF Class 2 Melanoma |
| HNF1A | 12q24.31 | 18 | 1.63 | 17 | 11.64 | 4.3E-08 | 1.3E-05 | (B) BRAF Class 2 Melanoma |
| NRAS | 1p13.2 | 10 | 0.85 | 14 | 8.00 | 1.4E-07 | 3E-05 | (B) BRAF Class 2 Melanoma |
| CARD11 | 7p22.2 | 61 | 7.92 | 27 | 24.32 | 1.5E-06 | 0.00028 | (B) BRAF Class 2 Melanoma |
| KMT2D | 12q13.12 | 118 | 15.32 | 38 | 34.23 | 4.8E-06 | 0.00072 | (B) BRAF Class 2 Melanoma |
| KMT2A | 11q23.3 | 67 | 8.61 | 27 | 23.89 | 7.8E-06 | 0.001 | (B) BRAF Class 2 Melanoma |
| ATR | 3q23 | 27 | 3.58 | 16 | 15.24 | 1.3E-05 | 0.00146 | (B) BRAF Class 2 Melanoma |
| MPL | 1p34.2 | 19 | 1.77 | 13 | 9.03 | 2.5E-05 | 0.00247 | (B) BRAF Class 2 Melanoma |
| EPHA7 | 6q16.1 | 57 | 10.27 | 23 | 28.40 | 3E-05 | 0.00267 | (B) BRAF Class 2 Melanoma |
| ARHGEF12 | 11q23.3 | 5 | 2.48 | 7 | 26.92 | 6.1E-05 | 0.00498 | (B) BRAF Class 2 Melanoma |
| CDKN1C | 11p15.4 | 1 | 0.37 | 5 | 13.16 | 0.00012 | 0.00892 | (B) BRAF Class 2 Melanoma |
| RPTOR | 17q25.3 | 34 | 4.73 | 16 | 15.69 | 0.00014 | 0.0096 | (B) BRAF Class 2 Melanoma |
| BLM | 15q26.1 | 23 | 3.03 | 13 | 11.82 | 0.0002 | 0.0128 | (B) BRAF Class 2 Melanoma |
| MAGI2 | 7q21.11 | 18 | 7.89 | 8 | 40.00 | 0.00027 | 0.0158 | (B) BRAF Class 2 Melanoma |
| MET | 7q31.2 | 68 | 5.80 | 24 | 13.87 | 0.00028 | 0.0158 | (B) BRAF Class 2 Melanoma |
| RNF213 | 17q25.3 | 5 | 11.11 | 0 | 0.00 | 0.0003 | 0.0158 | (A) BRAF Class 1 Melanoma |
| GRIN2A | 16p13.2 | 92 | 17.49 | 27 | 36.00 | 0.00033 | 0.0164 | (B) BRAF Class 2 Melanoma |
| FOXP1 | 3p13 | 9 | 1.74 | 8 | 10.81 | 0.0004 | 0.0185 | (B) BRAF Class 2 Melanoma |
| FLT1 | 13q12.3 | 69 | 8.96 | 23 | 20.72 | 0.00041 | 0.0185 | (B) BRAF Class 2 Melanoma |
| COL7A1 | 3p21.31 | 46 | 23.00 | 15 | 57.69 | 0.00043 | 0.0185 | (B) BRAF Class 2 Melanoma |
| ERBB4 | 2q34 | 107 | 9.54 | 32 | 18.93 | 0.00045 | 0.0185 | (B) BRAF Class 2 Melanoma |
| SF3B1 | 2q33.1 | 22 | 2.65 | 12 | 9.92 | 0.0005 | 0.0196 | (B) BRAF Class 2 Melanoma |
| SLIT2 | 4p15.31 | 28 | 10.61 | 10 | 38.46 | 0.00054 | 0.0201 | (B) BRAF Class 2 Melanoma |
| KDR | 4q12 | 87 | 7.87 | 25 | 17.01 | 0.00061 | 0.022 | (B) BRAF Class 2 Melanoma |
| XPO1 | 2p15 | 7 | 0.87 | 7 | 6.09 | 0.00065 | 0.0225 | (B) BRAF Class 2 Melanoma |
| SOX9 | 17q24.3 | 7 | 0.93 | 7 | 6.42 | 0.00068 | 0.0228 | (B) BRAF Class 2 Melanoma |
| COL2A1 | 12q13.11 | 3 | 5.56 | 4 | 66.67 | 0.00099 | 0.03 | (B) BRAF Class 2 Melanoma |
| RPS6KB2 | 11q13.2 | 3 | 0.74 | 5 | 8.77 | 0.00099 | 0.03 | (B) BRAF Class 2 Melanoma |
| JAK1 | 1p31.3 | 18 | 2.37 | 11 | 8.87 | 0.00101 | 0.03 | (B) BRAF Class 2 Melanoma |
| GATA6 | 18q11.2 | 4 | 0.85 | 5 | 8.93 | 0.00103 | 0.03 | (B) BRAF Class 2 Melanoma |
| ATM | 11q22.3 | 66 | 5.97 | 20 | 13.61 | 0.00133 | 0.036 | (B) BRAF Class 2 Melanoma |
| GRM8 | 7q31.33 | 4 | 9.30 | 0 | 0.00 | 0.00135 | 0.036 | (A) BRAF Class 1 Melanoma |
| TET2 | 4q24 | 70 | 9.00 | 22 | 19.30 | 0.00138 | 0.036 | (B) BRAF Class 2 Melanoma |
| PPARG | 3p25.2 | 17 | 2.64 | 9 | 10.71 | 0.00144 | 0.036 | (B) BRAF Class 2 Melanoma |
| POT1 | 7q31.33 | 2 | 0.74 | 4 | 12.12 | 0.0015 | 0.036 | (B) BRAF Class 2 Melanoma |
| NOTCH4 | 6p21.32 | 32 | 6.54 | 13 | 18.84 | 0.00152 | 0.036 | (B) BRAF Class 2 Melanoma |
| TP53 | 17p13.1 | 175 | 14.83 | 39 | 24.84 | 0.00154 | 0.036 | (B) BRAF Class 2 Melanoma |
| PAK5 | 20p12.2 | 62 | 13.45 | 20 | 28.99 | 0.00156 | 0.036 | (B) BRAF Class 2 Melanoma |
| RSPO2 | 8q23.1 | 4 | 1.28 | 5 | 11.90 | 0.00165 | 0.0371 | (B) BRAF Class 2 Melanoma |
| CSF3R | 1p34.3 | 13 | 1.85 | 8 | 8.33 | 0.00177 | 0.039 | (B) BRAF Class 2 Melanoma |
| SYK | 9q22.2 | 16 | 2.72 | 9 | 10.59 | 0.002 | 0.0423 | (B) BRAF Class 2 Melanoma |
| KRAS | 12p12.1 | 6 | 0.51 | 6 | 3.43 | 0.00202 | 0.0423 | (B) BRAF Class 2 Melanoma |

| | | | | | | | | |
|---|---|---|---|---|---|---|---|---|
| HIST1H3B | 6p22.2 | 7 | 1.01 | 6 | 6.19 | 0.00243 | 0.0498 | (B) BRAF Class 2 Melanoma |
| H3F3B | 17q25.1 | 2 | 0.32 | 4 | 4.60 | 0.00253 | 0.0498 | (B) BRAF Class 2 Melanoma |
| CBLB | 3q13.11 | 8 | 2.78 | 6 | 15.79 | 0.00254 | 0.0498 | (B) BRAF Class 2 Melanoma |
| FLT4 | 5q35.3 | 61 | 8.04 | 19 | 17.27 | 0.00294 | 0.0565 | (B) BRAF Class 2 Melanoma |
| TAP2 | 6p21.32 | 0 | 0.00 | 3 | 10.00 | 0.00305 | 0.0566 | (B) BRAF Class 2 Melanoma |
| FGFR2 | 10q26.13 | 45 | 3.81 | 16 | 9.14 | 0.00311 | 0.0566 | (B) BRAF Class 2 Melanoma |
| NOTCH2 | 1p12 | 44 | 5.56 | 15 | 13.51 | 0.0032 | 0.0566 | (B) BRAF Class 2 Melanoma |
| TCF12 | 15q21.3 | 1 | 1.39 | 2 | 66.67 | 0.00321 | 0.0566 | (B) BRAF Class 2 Melanoma |
| SLX4 | 16p13.3 | 29 | 5.48 | 11 | 15.94 | 0.00326 | 0.0566 | (B) BRAF Class 2 Melanoma |
| NTRK1 | 1q23.1 | 43 | 5.24 | 15 | 12.71 | 0.00336 | 0.0571 | (B) BRAF Class 2 Melanoma |
| FGFR1 | 8p11.23 | 17 | 1.44 | 9 | 5.14 | 0.00357 | 0.0594 | (B) BRAF Class 2 Melanoma |
| APC | 5q22.2 | 73 | 6.80 | 22 | 13.58 | 0.00362 | 0.0594 | (B) BRAF Class 2 Melanoma |
| CUX1 | 7q22.1 | 34 | 7.66 | 11 | 21.15 | 0.00372 | 0.06 | (B) BRAF Class 2 Melanoma |
| RUNX1T1 | 8q21.3 | 34 | 7.38 | 11 | 20.37 | 0.0038 | 0.0601 | (B) BRAF Class 2 Melanoma |
| NOTCH1 | 9q34.3 | 48 | 4.12 | 15 | 9.80 | 0.00397 | 0.0618 | (B) BRAF Class 2 Melanoma |
| TEK | 9p21.2 | 19 | 7.51 | 10 | 22.73 | 0.00433 | 0.0662 | (B) BRAF Class 2 Melanoma |
| FANCI | 15q26.1 | 11 | 2.46 | 6 | 11.76 | 0.00442 | 0.0664 | (B) BRAF Class 2 Melanoma |
| NR4A3 | 9q31.1 | 1 | 1.69 | 2 | 66.67 | 0.00471 | 0.0696 | (B) BRAF Class 2 Melanoma |
| HRAS | 11p15.5 | 5 | 0.42 | 5 | 2.87 | 0.00487 | 0.0702 | (B) BRAF Class 2 Melanoma |
| PDGFRB | 5q32 | 32 | 3.98 | 12 | 10.53 | 0.0049 | 0.0702 | (B) BRAF Class 2 Melanoma |
| RICTOR | 5p13.1 | 24 | 3.18 | 10 | 9.52 | 0.00504 | 0.0711 | (B) BRAF Class 2 Melanoma |
| RET | 10q11.21 | 44 | 3.73 | 15 | 8.57 | 0.00577 | 0.0792 | (B) BRAF Class 2 Melanoma |
| TCF7L2 | )q25.2-q25 | 9 | 1.31 | 6 | 6.25 | 0.0058 | 0.0792 | (B) BRAF Class 2 Melanoma |
| INPP4B | 4q31.21 | 9 | 1.70 | 6 | 8.00 | 0.00598 | 0.0804 | (B) BRAF Class 2 Melanoma |
| ARID2 | 12q12 | 66 | 8.70 | 19 | 17.27 | 0.00617 | 0.0818 | (B) BRAF Class 2 Melanoma |
| TYK2 | 19p13.2 | 3 | 3.85 | 3 | 42.86 | 0.00633 | 0.0823 | (B) BRAF Class 2 Melanoma |
| EIF4A2 | 3q27.3 | 1 | 0.25 | 3 | 5.45 | 0.00639 | 0.0823 | (B) BRAF Class 2 Melanoma |
| KMT2B | 19q13.12 | 40 | 10.96 | 12 | 26.09 | 0.00654 | 0.083 | (B) BRAF Class 2 Melanoma |
| FAT1 | 4q35.2 | 90 | 12.97 | 23 | 23.23 | 0.00696 | 0.0872 | (B) BRAF Class 2 Melanoma |
| DNMT1 | 19p13.2 | 14 | 3.14 | 7 | 11.67 | 0.00709 | 0.0872 | (B) BRAF Class 2 Melanoma |
| EP400 | 12q24.33 | 3 | 6.98 | 0 | 0.00 | 0.00725 | 0.0872 | (A) BRAF Class 1 Melanoma |
| RPS6KA2 | 6q27 | 3 | 6.98 | 0 | 0.00 | 0.00725 | 0.0872 | (A) BRAF Class 1 Melanoma |
| NOTCH3 | 19p13.12 | 86 | 11.54 | 22 | 20.95 | 0.00757 | 0.0899 | (B) BRAF Class 2 Melanoma |
| SOX10 | 22q13.1 | 6 | 2.16 | 4 | 14.29 | 0.0081 | 0.0948 | (B) BRAF Class 2 Melanoma |
| ZFHX3 | ;q22.2-q22 | 36 | 8.41 | 12 | 19.67 | 0.00913 | 0.104 | (B) BRAF Class 2 Melanoma |
| CREBBP | 16p13.3 | 40 | 4.97 | 13 | 11.40 | 0.00914 | 0.104 | (B) BRAF Class 2 Melanoma |
| DIS3 | 13q21.33 | 13 | 1.74 | 7 | 6.31 | 0.00922 | 0.104 | (B) BRAF Class 2 Melanoma |
| LIFR | 5p13.1 | 2 | 3.39 | 2 | 66.67 | 0.00931 | 0.104 | (B) BRAF Class 2 Melanoma |
| RARA | 17q21.2 | 10 | 1.30 | 6 | 5.41 | 0.00967 | 0.105 | (B) BRAF Class 2 Melanoma |
| CIITA | 16p13.13 | 16 | 5.84 | 7 | 19.44 | 0.00976 | 0.105 | (B) BRAF Class 2 Melanoma |
| BEND2 | Xp22.13 | 1 | 2.50 | 2 | 66.67 | 0.00981 | 0.105 | (B) BRAF Class 2 Melanoma |
| KDM5A | 12p13.33 | 22 | 3.08 | 9 | 8.82 | 0.00988 | 0.105 | (B) BRAF Class 2 Melanoma |
| ARID1A | 1p36.11 | 61 | 7.58 | 17 | 14.91 | 0.0104 | 0.109 | (B) BRAF Class 2 Melanoma |
| FANCA | 16q24.3 | 27 | 3.35 | 10 | 8.77 | 0.0109 | 0.112 | (B) BRAF Class 2 Melanoma |
| PREX2 | 8q13.2 | 75 | 17.69 | 18 | 32.14 | 0.011 | 0.112 | (B) BRAF Class 2 Melanoma |

| | | | | | | | | |
|---|---|---|---|---|---|---|---|---|
| PBRM1 | 3p21.1 | 23 | 3.02 | 9 | 8.41 | 0.0113 | 0.114 | (B) BRAF Class 2 Melanoma |
| ARAF | Xp11.3 | 7 | 0.85 | 5 | 4.20 | 0.0114 | 0.114 | (B) BRAF Class 2 Melanoma |
| LIG4 | 13q33.3 | 5 | 2.50 | 4 | 15.38 | 0.0118 | 0.117 | (B) BRAF Class 2 Melanoma |
| PMS1 | 2q32.2 | 15 | 2.14 | 7 | 7.14 | 0.012 | 0.118 | (B) BRAF Class 2 Melanoma |
| AR | Xq12 | 31 | 3.85 | 12 | 9.02 | 0.0121 | 0.118 | (B) BRAF Class 2 Melanoma |
| CDK8 | 13q12.13 | 11 | 1.53 | 6 | 5.88 | 0.0124 | 0.119 | (B) BRAF Class 2 Melanoma |
| JAK3 | 19p13.11 | 35 | 3.16 | 12 | 7.27 | 0.013 | 0.123 | (B) BRAF Class 2 Melanoma |
| DCC | 18q21.2 | 13 | 16.46 | 4 | 66.67 | 0.0134 | 0.126 | (B) BRAF Class 2 Melanoma |
| EP300 | 22q13.2 | 42 | 5.45 | 13 | 11.71 | 0.0143 | 0.13 | (B) BRAF Class 2 Melanoma |
| ICOSLG | 21q22.3 | 2 | 0.49 | 3 | 5.26 | 0.0147 | 0.13 | (B) BRAF Class 2 Melanoma |
| HIST1H3G | 6p22.2 | 2 | 0.51 | 3 | 5.45 | 0.0148 | 0.13 | (B) BRAF Class 2 Melanoma |
| HIST3H3 | 1q42.13 | 2 | 0.51 | 3 | 5.45 | 0.0148 | 0.13 | (B) BRAF Class 2 Melanoma |
| IFNGR1 | 6q23.3 | 0 | 0.00 | 2 | 3.51 | 0.0149 | 0.13 | (B) BRAF Class 2 Melanoma |
| ERCC3 | 2q14.3 | 12 | 1.80 | 6 | 6.52 | 0.015 | 0.13 | (B) BRAF Class 2 Melanoma |
| CDH23 | 10q22.1 | 2 | 8.00 | 0 | 0.00 | 0.0151 | 0.13 | (A) BRAF Class 1 Melanoma |
| PCDHAC2 | 5q31.3 | 2 | 8.00 | 0 | 0.00 | 0.0151 | 0.13 | (A) BRAF Class 1 Melanoma |
| TGFB1 | 19q13.2 | 2 | 8.00 | 0 | 0.00 | 0.0151 | 0.13 | (A) BRAF Class 1 Melanoma |
| PIK3C2G | 12p12.3 | 47 | 9.96 | 14 | 20.00 | 0.0155 | 0.13 | (B) BRAF Class 2 Melanoma |
| GALNT12 | 9q22.33 | 0 | 0.00 | 2 | 5.13 | 0.0155 | 0.13 | (B) BRAF Class 2 Melanoma |
| KEL | 7q34 | 22 | 9.21 | 6 | 28.57 | 0.0157 | 0.131 | (B) BRAF Class 2 Melanoma |
| MLH1 | 3p22.2 | 17 | 1.53 | 7 | 4.76 | 0.0161 | 0.133 | (B) BRAF Class 2 Melanoma |
| ADGRB3 | 6q12-q13 | 2 | 7.69 | 0 | 0.00 | 0.0162 | 0.133 | (A) BRAF Class 1 Melanoma |
| SPOP | 17q21.33 | 5 | 0.66 | 4 | 3.77 | 0.0168 | 0.137 | (B) BRAF Class 2 Melanoma |
| NBN | 8q21.3 | 16 | 2.00 | 7 | 6.14 | 0.0171 | 0.138 | (B) BRAF Class 2 Melanoma |
| DNAH9 | 17p12 | 2 | 7.41 | 0 | 0.00 | 0.0173 | 0.138 | (A) BRAF Class 1 Melanoma |
| PTPRT | [illegible]q12-q13.1 | 89 | 19.02 | 20 | 31.75 | 0.0175 | 0.138 | (B) BRAF Class 2 Melanoma |
| PTPN11 | 12q24.13 | 13 | 1.16 | 6 | 3.97 | 0.0183 | 0.143 | (B) BRAF Class 2 Melanoma |
| PIK3CG | 7q22.3 | 30 | 5.69 | 10 | 13.33 | 0.0184 | 0.143 | (B) BRAF Class 2 Melanoma |
| FGF14 | 13q33.1 | 4 | 1.38 | 3 | 10.34 | 0.0187 | 0.144 | (B) BRAF Class 2 Melanoma |
| DICER1 | 14q32.13 | 25 | 3.32 | 9 | 8.33 | 0.019 | 0.145 | (B) BRAF Class 2 Melanoma |
| AXL | 19q13.2 | 30 | 3.73 | 10 | 8.77 | 0.0191 | 0.145 | (B) BRAF Class 2 Melanoma |
| NTRK3 | 15q25.3 | 50 | 6.21 | 14 | 12.28 | 0.0194 | 0.146 | (B) BRAF Class 2 Melanoma |
| MAP3K13 | 3q27.2 | 12 | 2.54 | 6 | 8.57 | 0.0196 | 0.146 | (B) BRAF Class 2 Melanoma |
| GPC3 | Xq26.2 | 6 | 1.94 | 4 | 9.76 | 0.0201 | 0.148 | (B) BRAF Class 2 Melanoma |
| MST1R | 3p21.31 | 13 | 2.77 | 6 | 9.09 | 0.0207 | 0.152 | (B) BRAF Class 2 Melanoma |
| MTA1 | 14q32.33 | 3 | 1.50 | 3 | 11.54 | 0.0215 | 0.157 | (B) BRAF Class 2 Melanoma |
| IKZF1 | 7p12.2 | 60 | 7.84 | 16 | 14.41 | 0.0217 | 0.157 | (B) BRAF Class 2 Melanoma |
| TP63 | 3q28 | 38 | 8.96 | 11 | 18.97 | 0.022 | 0.157 | (B) BRAF Class 2 Melanoma |
| GLI1 | 12q13.3 | 30 | 4.12 | 10 | 9.43 | 0.0221 | 0.157 | (B) BRAF Class 2 Melanoma |
| ICK | 6p12.1 | 0 | 0.00 | 1 | 100.00 | 0.0227 | 0.16 | (B) BRAF Class 2 Melanoma |
| HIST1H1E | 6p22.2 | 0 | 0.00 | 1 | 100.00 | 0.0233 | 0.163 | (B) BRAF Class 2 Melanoma |
| ERCC1 | 19q13.32 | 1 | 0.26 | 2 | 5.13 | 0.0237 | 0.164 | (B) BRAF Class 2 Melanoma |
| KDM6A | Xp11.3 | 13 | 1.69 | 6 | 5.41 | 0.0238 | 0.164 | (B) BRAF Class 2 Melanoma |
| SPRED1 | 15q14 | 2 | 0.97 | 3 | 8.33 | 0.0243 | 0.164 | (B) BRAF Class 2 Melanoma |
| RIF1 | 2q23.3 | 11 | 5.50 | 5 | 19.23 | 0.0245 | 0.164 | (B) BRAF Class 2 Melanoma |

| | | | | | | | | |
|---|---|---|---|---|---|---|---|---|
| PLCG2 | 16q23.3 | 18 | 3.85 | 7 | 10.61 | 0.0249 | 0.164 | (B) BRAF Class 2 Melanoma |
| ABCB11 | 2q31.1 | 11 | 4.95 | 5 | 17.24 | 0.0255 | 0.164 | (B) BRAF Class 2 Melanoma |
| DNMT3A | 2p23.3 | 26 | 3.34 | 9 | 7.89 | 0.0259 | 0.164 | (B) BRAF Class 2 Melanoma |
| BRCA2 | 13q13.1 | 57 | 7.09 | 15 | 13.04 | 0.026 | 0.164 | (B) BRAF Class 2 Melanoma |
| BARD1 | 2q35 | 6 | 0.83 | 4 | 3.92 | 0.0261 | 0.164 | (B) BRAF Class 2 Melanoma |
| ARID1B | 6q25.3 | 46 | 6.18 | 13 | 12.04 | 0.0265 | 0.164 | (B) BRAF Class 2 Melanoma |
| PTPRB | 12q15 | 5 | 9.26 | 3 | 50.00 | 0.0266 | 0.164 | (B) BRAF Class 2 Melanoma |
| SNCAIP | 5q23.2 | 36 | 13.09 | 8 | 29.63 | 0.0275 | 0.164 | (B) BRAF Class 2 Melanoma |
| MEF2B | 19p13.11 | 6 | 0.78 | 4 | 3.60 | 0.0275 | 0.164 | (B) BRAF Class 2 Melanoma |
| CAD | 2p23.3 | 0 | 0.00 | 1 | 100.00 | 0.0278 | 0.164 | (B) BRAF Class 2 Melanoma |
| CPS1 | 2q34 | 0 | 0.00 | 1 | 100.00 | 0.0278 | 0.164 | (B) BRAF Class 2 Melanoma |
| INPP5D | 2q37.1 | 0 | 0.00 | 1 | 100.00 | 0.0278 | 0.164 | (B) BRAF Class 2 Melanoma |
| JARID2 | 6p22.3 | 0 | 0.00 | 1 | 100.00 | 0.0278 | 0.164 | (B) BRAF Class 2 Melanoma |
| MEF2C | 5q14.3 | 0 | 0.00 | 1 | 100.00 | 0.0278 | 0.164 | (B) BRAF Class 2 Melanoma |
| MKI67 | 10q26.2 | 0 | 0.00 | 1 | 100.00 | 0.0278 | 0.164 | (B) BRAF Class 2 Melanoma |
| PC | 11q13.2 | 0 | 0.00 | 1 | 100.00 | 0.0278 | 0.164 | (B) BRAF Class 2 Melanoma |
| PCLO | 7q21.11 | 0 | 0.00 | 1 | 100.00 | 0.0278 | 0.164 | (B) BRAF Class 2 Melanoma |
| PTPN6 | 12p13.31 | 0 | 0.00 | 1 | 100.00 | 0.0278 | 0.164 | (B) BRAF Class 2 Melanoma |
| TNFRSF11A | 18q21.33 | 0 | 0.00 | 1 | 100.00 | 0.0278 | 0.164 | (B) BRAF Class 2 Melanoma |
| WDR90 | 16p13.3 | 0 | 0.00 | 1 | 100.00 | 0.0278 | 0.164 | (B) BRAF Class 2 Melanoma |
| NSD3 | 8p11.23 | 9 | 2.46 | 5 | 8.77 | 0.0286 | 0.167 | (B) BRAF Class 2 Melanoma |
| RASA1 | 5q14.3 | 14 | 2.24 | 6 | 6.74 | 0.0288 | 0.167 | (B) BRAF Class 2 Melanoma |
| DDX41 | 5q35.3 | 2 | 1.03 | 2 | 12.50 | 0.0297 | 0.171 | (B) BRAF Class 2 Melanoma |
| ZFHX4 | 8q21.13 | 11 | 20.37 | 4 | 66.67 | 0.0298 | 0.171 | (B) BRAF Class 2 Melanoma |
| IL7R | 5p13.2 | 48 | 6.77 | 13 | 12.87 | 0.0301 | 0.172 | (B) BRAF Class 2 Melanoma |
| MYD88 | 3p22.2 | 10 | 1.23 | 5 | 4.27 | 0.0309 | 0.175 | (B) BRAF Class 2 Melanoma |
| CASR | q13.33-q21 | 3 | 7.50 | 2 | 66.67 | 0.0316 | 0.178 | (B) BRAF Class 2 Melanoma |
| ITK | 5q33.3 | 4 | 1.93 | 3 | 11.54 | 0.032 | 0.18 | (B) BRAF Class 2 Melanoma |
| SPTA1 | 1q23.1 | 53 | 20.08 | 10 | 38.46 | 0.0325 | 0.181 | (B) BRAF Class 2 Melanoma |
| PRPF8 | 17p13.3 | 1 | 1.05 | 2 | 16.67 | 0.0327 | 0.181 | (B) BRAF Class 2 Melanoma |
| LRP1B | q22.1-q22. | 86 | 30.82 | 14 | 50.00 | 0.0345 | 0.19 | (B) BRAF Class 2 Melanoma |
| BCL11B | 14q32.2 | 14 | 6.03 | 5 | 18.52 | 0.035 | 0.191 | (B) BRAF Class 2 Melanoma |
| EME1 | 17q21.33 | 1 | 0.50 | 2 | 7.69 | 0.0356 | 0.193 | (B) BRAF Class 2 Melanoma |
| CDKN2B | 9p21.3 | 113 | 14.09 | 24 | 21.24 | 0.0356 | 0.193 | (B) BRAF Class 2 Melanoma |
| COL1A1 | 17q21.33 | 3 | 3.70 | 2 | 33.33 | 0.0364 | 0.194 | (B) BRAF Class 2 Melanoma |
| NCOA2 | 8q13.3 | 3 | 3.70 | 2 | 33.33 | 0.0364 | 0.194 | (B) BRAF Class 2 Melanoma |
| NUTM1 | 15q14 | 17 | 7.42 | 4 | 25.00 | 0.0369 | 0.195 | (B) BRAF Class 2 Melanoma |
| IDH1 | 2q34 | 42 | 3.56 | 12 | 6.86 | 0.0371 | 0.195 | (B) BRAF Class 2 Melanoma |
| NFE2L2 | 2q31.2 | 7 | 0.87 | 4 | 3.48 | 0.038 | 0.199 | (B) BRAF Class 2 Melanoma |
| NLRP1 | 17p13.2 | 2 | 4.65 | 0 | 0.00 | 0.0382 | 0.199 | (A) BRAF Class 1 Melanoma |
| ETV1 | 7p21.2 | 15 | 2.08 | 6 | 5.77 | 0.0387 | 0.2 | (B) BRAF Class 2 Melanoma |
| SMC1A | Xp11.22 | 2 | 0.73 | 2 | 7.69 | 0.0389 | 0.2 | (B) BRAF Class 2 Melanoma |
| ABL1 | 9q34.12 | 26 | 2.42 | 8 | 5.56 | 0.039 | 0.2 | (B) BRAF Class 2 Melanoma |
| PRKCI | 3q26.2 | 16 | 2.56 | 6 | 6.98 | 0.0397 | 0.203 | (B) BRAF Class 2 Melanoma |
| NFKBIZ | 3q12.3 | 1 | 0.40 | 2 | 5.71 | 0.0407 | 0.205 | (B) BRAF Class 2 Melanoma |

| | | | | | | | | |
|---|---|---|---|---|---|---|---|---|
| ASXL1 | 20q11.21 | 29 | 3.73 | 9 | 7.96 | 0.0411 | 0.205 | (B) BRAF Class 2 Melanoma |
| CDH11 | 16q21 | 2 | 4.44 | 0 | 0.00 | 0.0411 | 0.205 | (A) BRAF Class 1 Melanoma |
| NIN | 14q22.1 | 2 | 4.44 | 0 | 0.00 | 0.0411 | 0.205 | (A) BRAF Class 1 Melanoma |
| SMARCB1 | ı11.23\|22ı | 8 | 0.72 | 4 | 2.74 | 0.0413 | 0.205 | (B) BRAF Class 2 Melanoma |
| PTPRO | 2.3\|12p13 | 0 | 0.00 | 1 | 50.00 | 0.0417 | 0.205 | (B) BRAF Class 2 Melanoma |
| KAT6A | 8p11.21 | 20 | 4.29 | 6 | 11.11 | 0.0423 | 0.206 | (B) BRAF Class 2 Melanoma |
| DAXX | 6p21.32 | 16 | 2.27 | 6 | 6.06 | 0.0428 | 0.206 | (B) BRAF Class 2 Melanoma |
| MN1 | 22q12.1 | 0 | 0.00 | 1 | 33.33 | 0.0429 | 0.206 | (B) BRAF Class 2 Melanoma |
| PTPRS | 19p13.3 | 21 | 5.16 | 7 | 12.28 | 0.0432 | 0.206 | (B) BRAF Class 2 Melanoma |
| SDHB | 1p36.13 | 1 | 0.13 | 2 | 1.82 | 0.0432 | 0.206 | (B) BRAF Class 2 Melanoma |
| RAD51 | 15q15.1 | 4 | 0.53 | 3 | 2.86 | 0.0433 | 0.206 | (B) BRAF Class 2 Melanoma |
| PPP4R2 | 3p13 | 3 | 1.75 | 3 | 10.00 | 0.0441 | 0.209 | (B) BRAF Class 2 Melanoma |
| XRCC2 | 7q36.1 | 12 | 1.82 | 5 | 5.49 | 0.0447 | 0.21 | (B) BRAF Class 2 Melanoma |
| USP6 | 17p13.2 | 0 | 0.00 | 1 | 33.33 | 0.0448 | 0.21 | (B) BRAF Class 2 Melanoma |
| NF2 | 22q12.2 | 16 | 1.98 | 6 | 5.22 | 0.0456 | 0.213 | (B) BRAF Class 2 Melanoma |
| ADGRA2 | 8p11.23 | 6 | 2.29 | 3 | 10.71 | 0.0457 | 0.213 | (B) BRAF Class 2 Melanoma |
| RNF43 | 17q22 | 12 | 1.59 | 5 | 4.76 | 0.046 | 0.213 | (B) BRAF Class 2 Melanoma |
| NCOR2 | 12q24.31 | 1 | 2.38 | 1 | 100.00 | 0.0465 | 0.214 | (B) BRAF Class 2 Melanoma |
| CD74 | 5q33.1 | 0 | 0.00 | 1 | 10.00 | 0.0476 | 0.218 | (B) BRAF Class 2 Melanoma |
| PIK3CA | 3q26.32 | 34 | 2.88 | 10 | 5.71 | 0.0479 | 0.218 | (B) BRAF Class 2 Melanoma |
| NT5C2 | ı24.32-q24 | 5 | 1.85 | 3 | 8.82 | 0.0481 | 0.218 | (B) BRAF Class 2 Melanoma |
| PIM1 | 6p21.2 | 8 | 1.19 | 4 | 4.26 | 0.0484 | 0.218 | (B) BRAF Class 2 Melanoma |
| ZNF217 | 20q13.2 | 14 | 2.69 | 5 | 7.69 | 0.0488 | 0.219 | (B) BRAF Class 2 Melanoma |
| RHBDF2 | 17q25.1 | 5 | 2.50 | 3 | 11.54 | 0.0514 | 0.229 | (B) BRAF Class 2 Melanoma |
| BCOR | Xp11.4 | 36 | 4.68 | 10 | 9.01 | 0.0528 | 0.235 | (B) BRAF Class 2 Melanoma |
| CDKN2A | 9p21.3 | 298 | 26.49 | 50 | 33.11 | 0.0545 | 0.241 | (B) BRAF Class 2 Melanoma |
| MAP2K1 | 15q22.31 | 51 | 5.81 | 14 | 9.86 | 0.0552 | 0.242 | (B) BRAF Class 2 Melanoma |
| PDCD11 | 10q24.33 | 1 | 2.86 | 1 | 100.00 | 0.0556 | 0.242 | (B) BRAF Class 2 Melanoma |
| WWTR1 | 3q25.1 | 1 | 0.51 | 2 | 6.06 | 0.0556 | 0.242 | (B) BRAF Class 2 Melanoma |
| ALK | p23.2-p23. | 72 | 6.10 | 17 | 9.71 | 0.056 | 0.243 | (B) BRAF Class 2 Melanoma |
| TCF3 | 19p13.3 | 13 | 1.98 | 5 | 5.49 | 0.0567 | 0.244 | (B) BRAF Class 2 Melanoma |
| ERG | 21q22.2 | 27 | 3.75 | 8 | 7.84 | 0.057 | 0.244 | (B) BRAF Class 2 Melanoma |
| PLCB4 | )p12.3-p12 | 5 | 7.14 | 3 | 30.00 | 0.0572 | 0.244 | (B) BRAF Class 2 Melanoma |
| CTNNA1 | 5q31.2 | 6 | 1.27 | 3 | 5.45 | 0.0574 | 0.244 | (B) BRAF Class 2 Melanoma |
| SESN2 | 1p35.3 | 1 | 0.58 | 2 | 6.67 | 0.0588 | 0.249 | (B) BRAF Class 2 Melanoma |
| PTCH1 | 9q22.32 | 27 | 3.35 | 8 | 6.96 | 0.0596 | 0.251 | (B) BRAF Class 2 Melanoma |
| NSD1 | 5q35.3 | 48 | 6.82 | 12 | 11.88 | 0.0597 | 0.251 | (B) BRAF Class 2 Melanoma |
| PGR | 11q22.1 | 18 | 4.59 | 6 | 10.91 | 0.0607 | 0.252 | (B) BRAF Class 2 Melanoma |
| SS18 | 18q11.2 | 2 | 0.77 | 2 | 6.25 | 0.0607 | 0.252 | (B) BRAF Class 2 Melanoma |
| SYNE1 | 6q25.2 | 5 | 2.50 | 2 | 15.38 | 0.0608 | 0.252 | (B) BRAF Class 2 Melanoma |
| RPS6KA4 | 11q13.1 | 9 | 2.21 | 4 | 7.02 | 0.0627 | 0.257 | (B) BRAF Class 2 Melanoma |
| SOX17 | 8q11.23 | 5 | 1.23 | 3 | 5.26 | 0.0628 | 0.257 | (B) BRAF Class 2 Melanoma |
| TNFRSF14 | 1p36.32 | 5 | 0.93 | 3 | 4.00 | 0.0629 | 0.257 | (B) BRAF Class 2 Melanoma |
| RSPO3 | 6q22.33 | 2 | 0.79 | 2 | 6.25 | 0.0635 | 0.258 | (B) BRAF Class 2 Melanoma |
| IRF4 | 6p25.3 | 5 | 0.93 | 3 | 3.95 | 0.0645 | 0.26 | (B) BRAF Class 2 Melanoma |

| | | | | | | | | |
|---|---|---|---|---|---|---|---|---|
| TNFAIP3 | 6q23.3 | 13 | 1.69 | 5 | 4.50 | 0.0645 | 0.26 | (B) BRAF Class 2 Melanoma |
| TLX3 | 5q35.1 | 2 | 0.99 | 2 | 7.69 | 0.0651 | 0.261 | (B) BRAF Class 2 Melanoma |
| GREM1 | 15q13.3 | 5 | 0.74 | 3 | 3.13 | 0.0653 | 0.261 | (B) BRAF Class 2 Melanoma |
| NECTIN4 | 1q23.3 | 2 | 1.00 | 2 | 7.69 | 0.0662 | 0.263 | (B) BRAF Class 2 Melanoma |
| INHBA | 7p14.1 | 23 | 4.50 | 7 | 9.59 | 0.0673 | 0.266 | (B) BRAF Class 2 Melanoma |
| TBX3 | 12q24.21 | 28 | 5.30 | 8 | 10.53 | 0.0689 | 0.27 | (B) BRAF Class 2 Melanoma |
| MGA | 15q15.1 | 67 | 10.98 | 15 | 17.24 | 0.0692 | 0.27 | (B) BRAF Class 2 Melanoma |
| MEN1 | 11q13.1 | 9 | 1.17 | 4 | 3.60 | 0.0692 | 0.27 | (B) BRAF Class 2 Melanoma |
| GNA14 | 9q21.2 | 0 | 0.00 | 1 | 33.33 | 0.0698 | 0.27 | (B) BRAF Class 2 Melanoma |
| HTRA1 | 10q26.13 | 0 | 0.00 | 1 | 33.33 | 0.0698 | 0.27 | (B) BRAF Class 2 Melanoma |
| PIK3C3 | 18q12.3 | 5 | 1.07 | 3 | 4.35 | 0.0707 | 0.272 | (B) BRAF Class 2 Melanoma |
| PAXIP1 | 7q36.2 | 6 | 3.00 | 3 | 11.54 | 0.0712 | 0.273 | (B) BRAF Class 2 Melanoma |
| FGFR3 | 4p16.3 | 32 | 2.71 | 9 | 5.14 | 0.0716 | 0.273 | (B) BRAF Class 2 Melanoma |
| EPHA5 | q13.1-q13. | 18 | 3.13 | 6 | 7.14 | 0.0718 | 0.273 | (B) BRAF Class 2 Melanoma |
| CDK12 | 17q12 | 24 | 3.18 | 7 | 6.67 | 0.0723 | 0.274 | (B) BRAF Class 2 Melanoma |
| KDM5C | Xp11.22 | 24 | 3.15 | 7 | 6.54 | 0.0751 | 0.282 | (B) BRAF Class 2 Melanoma |
| CHEK1 | 11q24.2 | 2 | 0.26 | 2 | 1.90 | 0.0751 | 0.282 | (B) BRAF Class 2 Melanoma |
| HIST1H2BD | 6p22.2 | 2 | 0.49 | 2 | 3.51 | 0.0756 | 0.283 | (B) BRAF Class 2 Melanoma |
| PRKCA | 17q24.2 | 0 | 0.00 | 1 | 16.67 | 0.0759 | 0.283 | (B) BRAF Class 2 Melanoma |
| PRKACA | 19p13.12 | 0 | 0.00 | 1 | 16.67 | 0.0779 | 0.288 | (B) BRAF Class 2 Melanoma |
| SMAD3 | 15q22.33 | 2 | 0.39 | 2 | 2.74 | 0.078 | 0.288 | (B) BRAF Class 2 Melanoma |
| RAF1 | 3p25.2 | 8 | 0.97 | 4 | 2.92 | 0.0787 | 0.29 | (B) BRAF Class 2 Melanoma |
| PIK3C2B | 1q32.1 | 24 | 5.01 | 6 | 10.71 | 0.0812 | 0.296 | (B) BRAF Class 2 Melanoma |
| ROS1 | 6q22.1 | 147 | 17.09 | 31 | 22.46 | 0.0816 | 0.296 | (B) BRAF Class 2 Melanoma |
| SETBP1 | 18q12.3 | 38 | 8.30 | 8 | 15.38 | 0.0817 | 0.296 | (B) BRAF Class 2 Melanoma |
| CHD4 | 12p13.31 | 9 | 3.41 | 3 | 11.54 | 0.0817 | 0.296 | (B) BRAF Class 2 Melanoma |
| EWSR1 | 22q12.2 | 7 | 1.34 | 3 | 4.76 | 0.0824 | 0.297 | (B) BRAF Class 2 Melanoma |
| AXIN2 | 17q24.1 | 20 | 3.12 | 6 | 6.74 | 0.0842 | 0.302 | (B) BRAF Class 2 Melanoma |
| MAP2K4 | 17p12 | 10 | 1.25 | 4 | 3.51 | 0.0844 | 0.302 | (B) BRAF Class 2 Melanoma |
| DGKH | 13q14.11 | 0 | 0.00 | 1 | 33.33 | 0.0857 | 0.304 | (B) BRAF Class 2 Melanoma |
| PRKCH | 14q23.1 | 0 | 0.00 | 1 | 33.33 | 0.0857 | 0.304 | (B) BRAF Class 2 Melanoma |
| EBF1 | 5q33.3 | 0 | 0.00 | 1 | 14.29 | 0.0875 | 0.308 | (B) BRAF Class 2 Melanoma |
| FOXL2 | 3q22.3 | 6 | 0.79 | 3 | 2.83 | 0.0876 | 0.308 | (B) BRAF Class 2 Melanoma |
| ASXL2 | 2p23.3 | 25 | 5.64 | 7 | 11.11 | 0.088 | 0.309 | (B) BRAF Class 2 Melanoma |
| WT1 | 11p13 | 24 | 3.08 | 7 | 6.14 | 0.0885 | 0.309 | (B) BRAF Class 2 Melanoma |
| HGF | 7q21.11 | 36 | 6.70 | 9 | 11.84 | 0.09 | 0.313 | (B) BRAF Class 2 Melanoma |
| BMPR1A | 10q23.2 | 6 | 0.81 | 3 | 2.86 | 0.0903 | 0.313 | (B) BRAF Class 2 Melanoma |
| NTHL1 | 16p13.3 | 2 | 0.53 | 2 | 3.39 | 0.091 | 0.314 | (B) BRAF Class 2 Melanoma |
| ATRX | Xq21.1 | 47 | 5.84 | 11 | 9.65 | 0.0915 | 0.315 | (B) BRAF Class 2 Melanoma |
| CDH1 | 16q22.1 | 16 | 1.49 | 5 | 3.47 | 0.0917 | 0.315 | (B) BRAF Class 2 Melanoma |
| MTOR | 1p36.22 | 41 | 4.75 | 11 | 7.91 | 0.0923 | 0.315 | (B) BRAF Class 2 Melanoma |
| CBL | 11q23.3 | 30 | 3.69 | 8 | 6.84 | 0.0926 | 0.315 | (B) BRAF Class 2 Melanoma |
| KIT | 4q12 | 24 | 2.03 | 7 | 4.00 | 0.0939 | 0.318 | (B) BRAF Class 2 Melanoma |
| MLH3 | 14q24.3 | 3 | 1.15 | 2 | 6.25 | 0.0944 | 0.318 | (B) BRAF Class 2 Melanoma |
| HOXB13 | 17q21.32 | 6 | 0.90 | 3 | 3.09 | 0.0945 | 0.318 | (B) BRAF Class 2 Melanoma |

| | | | | | | | | |
|---|---|---|---|---|---|---|---|---|
| CAMTA1 | 36.31-p36. | 1 | 1.69 | 1 | 33.33 | 0.0952 | 0.318 | (B) BRAF Class 2 Melanoma |
| NCOR1 | 7p12-p11. | 30 | 5.84 | 8 | 10.67 | 0.0953 | 0.318 | (B) BRAF Class 2 Melanoma |
| FGFR4 | 5q35.2 | 26 | 3.20 | 8 | 5.93 | 0.0969 | 0.323 | (B) BRAF Class 2 Melanoma |
| GATA3 | 10p14 | 20 | 2.60 | 6 | 5.36 | 0.0999 | 0.331 | (B) BRAF Class 2 Melanoma |
| RAD21 | 8q24.11 | 16 | 2.28 | 5 | 5.10 | 0.102 | 0.337 | (B) BRAF Class 2 Melanoma |
| RINT1 | 7q22.3 | 3 | 1.50 | 2 | 7.69 | 0.102 | 0.337 | (B) BRAF Class 2 Melanoma |
| PRSS1 | 7q34 | 17 | 6.39 | 5 | 13.89 | 0.105 | 0.34 | (B) BRAF Class 2 Melanoma |
| CHD5 | 1p36.31 | 4 | 7.41 | 2 | 33.33 | 0.105 | 0.34 | (B) BRAF Class 2 Melanoma |
| AXIN1 | 16p13.3 | 11 | 2.10 | 4 | 5.33 | 0.105 | 0.34 | (B) BRAF Class 2 Melanoma |
| STAT4 | q32.2-q32. | 10 | 3.60 | 3 | 10.71 | 0.105 | 0.34 | (B) BRAF Class 2 Melanoma |
| FH | 1q43 | 11 | 1.44 | 4 | 3.67 | 0.105 | 0.34 | (B) BRAF Class 2 Melanoma |
| FOXF1 | 16q24.1 | 1 | 4.76 | 0 | 0.00 | 0.107 | 0.344 | (A) BRAF Class 1 Melanoma |
| SLFN11 | 17q12 | 1 | 4.76 | 0 | 0.00 | 0.107 | 0.344 | (A) BRAF Class 1 Melanoma |
| STAT5B | 17q21.2 | 3 | 0.72 | 2 | 3.57 | 0.109 | 0.349 | (B) BRAF Class 2 Melanoma |
| MLLT3 | 9p21.3 | 1 | 0.60 | 1 | 10.00 | 0.11 | 0.349 | (B) BRAF Class 2 Melanoma |
| PRPF40B | 12q13.12 | 0 | 0.00 | 1 | 8.33 | 0.112 | 0.356 | (B) BRAF Class 2 Melanoma |
| BRCA1 | 17q21.31 | 32 | 3.98 | 9 | 6.77 | 0.113 | 0.358 | (B) BRAF Class 2 Melanoma |
| EPHB1 | 3q22.2 | 27 | 5.02 | 7 | 9.21 | 0.113 | 0.358 | (B) BRAF Class 2 Melanoma |
| FGF8 | 10q24.32 | 1 | 0.63 | 1 | 10.00 | 0.115 | 0.36 | (B) BRAF Class 2 Melanoma |
| MCM7 | 7q22.1 | 1 | 4.35 | 0 | 0.00 | 0.116 | 0.36 | (A) BRAF Class 1 Melanoma |
| E2F3 | 6p22.3 | 3 | 0.74 | 2 | 3.51 | 0.116 | 0.36 | (B) BRAF Class 2 Melanoma |
| H3F3C | 12p11.21 | 3 | 0.74 | 2 | 3.51 | 0.116 | 0.36 | (B) BRAF Class 2 Melanoma |
| CENPA | 2p23.3 | 3 | 0.77 | 2 | 3.64 | 0.117 | 0.36 | (B) BRAF Class 2 Melanoma |
| HIST1H3I | 6p22.1 | 3 | 0.77 | 2 | 3.64 | 0.117 | 0.36 | (B) BRAF Class 2 Melanoma |
| BTK | Xq22.1 | 12 | 2.10 | 4 | 5.06 | 0.117 | 0.36 | (B) BRAF Class 2 Melanoma |
| SMO | 7q32.1 | 36 | 3.25 | 9 | 5.45 | 0.118 | 0.36 | (B) BRAF Class 2 Melanoma |
| IGF2 | 11p15.5 | 7 | 0.99 | 3 | 2.97 | 0.118 | 0.36 | (B) BRAF Class 2 Melanoma |
| MYCN | 2p24.3 | 14 | 1.74 | 5 | 3.76 | 0.119 | 0.36 | (B) BRAF Class 2 Melanoma |
| MRE11 | 11q21 | 17 | 2.25 | 5 | 4.76 | 0.119 | 0.36 | (B) BRAF Class 2 Melanoma |
| USP28 | 11q23.2 | 8 | 4.00 | 3 | 11.54 | 0.119 | 0.36 | (B) BRAF Class 2 Melanoma |
| PTPRD | 9p24.1-p23 | 88 | 15.74 | 17 | 21.79 | 0.119 | 0.36 | (B) BRAF Class 2 Melanoma |
| AURKB | 17p13.1 | 7 | 0.91 | 3 | 2.70 | 0.121 | 0.362 | (B) BRAF Class 2 Melanoma |
| RECQL4 | 8q24.3 | 35 | 5.15 | 8 | 8.79 | 0.122 | 0.362 | (B) BRAF Class 2 Melanoma |
| AKT2 | 19q13.2 | 12 | 1.49 | 4 | 3.51 | 0.125 | 0.362 | (B) BRAF Class 2 Melanoma |
| CACNA1E | 1q25.3 | 1 | 4.00 | 0 | 0.00 | 0.125 | 0.362 | (A) BRAF Class 1 Melanoma |
| GNAI3 | 1p13.3 | 1 | 4.00 | 0 | 0.00 | 0.125 | 0.362 | (A) BRAF Class 1 Melanoma |
| LRIG3 | 12q14.1 | 1 | 4.00 | 0 | 0.00 | 0.125 | 0.362 | (A) BRAF Class 1 Melanoma |
| SP140 | 2q37.1 | 1 | 4.00 | 0 | 0.00 | 0.125 | 0.362 | (A) BRAF Class 1 Melanoma |
| UMODL1 | 21q22.3 | 1 | 4.00 | 0 | 0.00 | 0.125 | 0.362 | (A) BRAF Class 1 Melanoma |
| STK11 | 19p13.3 | 18 | 1.60 | 5 | 3.31 | 0.126 | 0.362 | (B) BRAF Class 2 Melanoma |
| BCL2 | 18q21.33 | 0 | 0.00 | 1 | 0.91 | 0.126 | 0.362 | (B) BRAF Class 2 Melanoma |
| FGF6 | 12p13.32 | 5 | 1.73 | 2 | 6.90 | 0.127 | 0.362 | (B) BRAF Class 2 Melanoma |
| PGBD3 | 10q11.23 | 3 | 0.25 | 2 | 1.14 | 0.127 | 0.362 | (B) BRAF Class 2 Melanoma |
| KDM6B | 17p13.1 | 4 | 5.48 | 2 | 22.22 | 0.128 | 0.362 | (B) BRAF Class 2 Melanoma |
| EPHA3 | 3p11.1 | 44 | 7.48 | 10 | 11.76 | 0.128 | 0.362 | (B) BRAF Class 2 Melanoma |

| | | | | | | | | |
|---|---|---|---|---|---|---|---|---|
| MDM2 | 12q15 | 12 | 1.49 | 4 | 3.48 | 0.128 | 0.362 | (B) BRAF Class 2 Melanoma |
| POLQ | 3q13.33 | 13 | 6.07 | 4 | 13.79 | 0.129 | 0.362 | (B) BRAF Class 2 Melanoma |
| C5ORF22 | 5p13.3 | 0 | 0.00 | 1 | 0.57 | 0.129 | 0.362 | (B) BRAF Class 2 Melanoma |
| KCNJ1 | 11q24.3 | 0 | 0.00 | 1 | 0.57 | 0.129 | 0.362 | (B) BRAF Class 2 Melanoma |
| LAMC3 | 9q34.12 | 0 | 0.00 | 1 | 0.57 | 0.129 | 0.362 | (B) BRAF Class 2 Melanoma |
| PEX5 | 12p13.31 | 0 | 0.00 | 1 | 0.57 | 0.129 | 0.362 | (B) BRAF Class 2 Melanoma |
| VPREB1 | 22q11.22 | 0 | 0.00 | 1 | 0.57 | 0.129 | 0.362 | (B) BRAF Class 2 Melanoma |
| VRK2 | 2p16.1 | 0 | 0.00 | 1 | 0.57 | 0.129 | 0.362 | (B) BRAF Class 2 Melanoma |
| GNB1 | 1p36.33 | 1 | 3.85 | 0 | 0.00 | 0.129 | 0.362 | (A) BRAF Class 1 Melanoma |
| PAX3 | 2q36.1 | 1 | 0.45 | 1 | 6.25 | 0.13 | 0.364 | (B) BRAF Class 2 Melanoma |
| NFKBIE | 6p21.1 | 8 | 3.39 | 3 | 9.38 | 0.131 | 0.365 | (B) BRAF Class 2 Melanoma |
| SETD2 | 3p21.31 | 57 | 7.08 | 12 | 10.53 | 0.133 | 0.369 | (B) BRAF Class 2 Melanoma |
| GNA11 | 19p13.3 | 7 | 0.63 | 3 | 1.78 | 0.134 | 0.369 | (B) BRAF Class 2 Melanoma |
| RPL22 | 1p36.31 | 1 | 3.70 | 0 | 0.00 | 0.134 | 0.369 | (A) BRAF Class 1 Melanoma |
| ERCC5 | 13q33.1 | 18 | 2.70 | 5 | 5.43 | 0.135 | 0.37 | (B) BRAF Class 2 Melanoma |
| FRK | 6q22.1 | 1 | 2.50 | 1 | 33.33 | 0.136 | 0.37 | (B) BRAF Class 2 Melanoma |
| GLMN | 1p22.1 | 1 | 2.50 | 1 | 33.33 | 0.136 | 0.37 | (B) BRAF Class 2 Melanoma |
| NRIP1 | .q11.2-q21 | 1 | 2.50 | 1 | 33.33 | 0.136 | 0.37 | (B) BRAF Class 2 Melanoma |
| ZFPM1 | 16q24.2 | 1 | 2.50 | 1 | 33.33 | 0.136 | 0.37 | (B) BRAF Class 2 Melanoma |
| NRG1 | 8p12 | 18 | 4.74 | 4 | 10.26 | 0.137 | 0.372 | (B) BRAF Class 2 Melanoma |
| MAP2K2 | 19p13.3 | 15 | 1.99 | 5 | 4.03 | 0.138 | 0.374 | (B) BRAF Class 2 Melanoma |
| PHOX2B | 4p13 | 18 | 2.56 | 5 | 5.10 | 0.139 | 0.375 | (B) BRAF Class 2 Melanoma |
| PTPRK | 6q22.33 | 5 | 9.26 | 2 | 33.33 | 0.14 | 0.375 | (B) BRAF Class 2 Melanoma |
| TERT | 5p15.33 | 460 | 61.01 | 71 | 66.98 | 0.14 | 0.375 | (B) BRAF Class 2 Melanoma |
| SHQ1 | 3p13 | 8 | 1.97 | 3 | 5.26 | 0.142 | 0.378 | (B) BRAF Class 2 Melanoma |
| MCM8 | 20p12.3 | 4 | 2.00 | 2 | 7.69 | 0.143 | 0.379 | (B) BRAF Class 2 Melanoma |
| XRCC4 | 5q14.2 | 4 | 2.00 | 2 | 7.69 | 0.143 | 0.379 | (B) BRAF Class 2 Melanoma |
| ZRSR2 | Xp22.2 | 8 | 1.16 | 3 | 3.09 | 0.143 | 0.379 | (B) BRAF Class 2 Melanoma |
| CBFA2T3 | 16q24.3 | 9 | 4.46 | 3 | 11.54 | 0.144 | 0.38 | (B) BRAF Class 2 Melanoma |
| ERCC4 | 16p13.12 | 18 | 2.48 | 5 | 4.85 | 0.146 | 0.384 | (B) BRAF Class 2 Melanoma |
| SOS1 | 2p22.1 | 7 | 1.80 | 3 | 4.84 | 0.146 | 0.384 | (B) BRAF Class 2 Melanoma |
| PDGFRA | 4q12 | 50 | 4.33 | 11 | 6.47 | 0.148 | 0.386 | (B) BRAF Class 2 Melanoma |
| RRAS2 | 11p15.2 | 0 | 0.00 | 1 | 2.78 | 0.148 | 0.386 | (B) BRAF Class 2 Melanoma |
| MYCL | 1p34.2 | 8 | 0.99 | 3 | 2.61 | 0.149 | 0.386 | (B) BRAF Class 2 Melanoma |
| UPF1 | 19p13.11 | 0 | 0.00 | 1 | 3.33 | 0.149 | 0.387 | (B) BRAF Class 2 Melanoma |
| RASA2 | 3q23 | 1 | 1.41 | 1 | 16.67 | 0.151 | 0.39 | (B) BRAF Class 2 Melanoma |
| SPEN | 36.21-p36. | 53 | 9.87 | 11 | 14.47 | 0.152 | 0.391 | (B) BRAF Class 2 Melanoma |
| REL | 2p16.1 | 8 | 1.04 | 3 | 2.70 | 0.152 | 0.391 | (B) BRAF Class 2 Melanoma |
| EGFR | 7p11.2 | 50 | 4.24 | 11 | 6.29 | 0.153 | 0.391 | (B) BRAF Class 2 Melanoma |
| ZBTB2 | 6q25.1 | 1 | 0.44 | 1 | 5.00 | 0.155 | 0.396 | (B) BRAF Class 2 Melanoma |
| SUFU | 10q24.32 | 13 | 1.69 | 4 | 3.60 | 0.156 | 0.397 | (B) BRAF Class 2 Melanoma |
| GABRA6 | 5q34 | 7 | 2.93 | 2 | 9.52 | 0.159 | 0.402 | (B) BRAF Class 2 Melanoma |
| CDH4 | 20q13.33 | 15 | 7.50 | 4 | 15.38 | 0.159 | 0.402 | (B) BRAF Class 2 Melanoma |
| FGF10 | 5p12 | 6 | 2.08 | 2 | 6.90 | 0.159 | 0.402 | (B) BRAF Class 2 Melanoma |
| FUBP1 | 1p31.1 | 4 | 0.76 | 2 | 2.74 | 0.16 | 0.402 | (B) BRAF Class 2 Melanoma |

| | | | | | | | | |
|---|---|---|---|---|---|---|---|---|
| SLC34A2 | 4p15.2 | 11 | 4.80 | 3 | 11.54 | 0.16 | 0.402 | (B) BRAF Class 2 Melanoma |
| MECOM | 3q26.2 | 39 | 14.77 | 8 | 22.86 | 0.161 | 0.402 | (B) BRAF Class 2 Melanoma |
| RYBP | 3p13 | 4 | 0.98 | 2 | 3.51 | 0.161 | 0.402 | (B) BRAF Class 2 Melanoma |
| CD79A | 19q13.2 | 4 | 0.76 | 2 | 2.70 | 0.161 | 0.402 | (B) BRAF Class 2 Melanoma |
| NEGR1 | 1p31.1 | 14 | 3.23 | 4 | 6.67 | 0.163 | 0.406 | (B) BRAF Class 2 Melanoma |
| LYN | 8q12.1 | 15 | 3.83 | 4 | 7.84 | 0.164 | 0.406 | (B) BRAF Class 2 Melanoma |
| TSC1 | 9q34.13 | 30 | 3.71 | 7 | 6.09 | 0.166 | 0.411 | (B) BRAF Class 2 Melanoma |
| BCL2L12 | 19q13.33 | 4 | 1.52 | 2 | 5.26 | 0.167 | 0.411 | (B) BRAF Class 2 Melanoma |
| CNOT3 | 19q13.42 | 1 | 1.64 | 1 | 16.67 | 0.172 | 0.424 | (B) BRAF Class 2 Melanoma |
| MSH3 | 5q14.1 | 10 | 2.66 | 3 | 6.25 | 0.173 | 0.424 | (B) BRAF Class 2 Melanoma |
| INPPL1 | 11q13.4 | 7 | 4.09 | 3 | 10.00 | 0.173 | 0.424 | (B) BRAF Class 2 Melanoma |
| NIPBL | 5p13.2 | 1 | 1.69 | 1 | 16.67 | 0.177 | 0.432 | (B) BRAF Class 2 Melanoma |
| PDCD1LG2 | 9p24.1 | 9 | 1.36 | 3 | 3.19 | 0.179 | 0.434 | (B) BRAF Class 2 Melanoma |
| BRCC3 | Xq28 | 5 | 2.44 | 2 | 7.69 | 0.18 | 0.435 | (B) BRAF Class 2 Melanoma |
| IGF1R | 15q26.3 | 42 | 5.17 | 9 | 7.69 | 0.18 | 0.435 | (B) BRAF Class 2 Melanoma |
| ERBB2 | 17q12 | 24 | 2.03 | 6 | 3.43 | 0.181 | 0.437 | (B) BRAF Class 2 Melanoma |
| MUTYH | 1p34.1 | 14 | 1.82 | 4 | 3.60 | 0.182 | 0.437 | (B) BRAF Class 2 Melanoma |
| HNRNPK | 9q21.32 | 2 | 1.06 | 1 | 7.69 | 0.183 | 0.438 | (B) BRAF Class 2 Melanoma |
| AURKA | 20q13.2 | 9 | 1.17 | 3 | 2.70 | 0.184 | 0.439 | (B) BRAF Class 2 Melanoma |
| JAK2 | 9p24.1 | 24 | 2.17 | 6 | 3.64 | 0.184 | 0.439 | (B) BRAF Class 2 Melanoma |
| LZTR1 | 21\|22q11. | 7 | 2.62 | 2 | 7.69 | 0.185 | 0.441 | (B) BRAF Class 2 Melanoma |
| PTPN14 | lq32.3-q41 | 5 | 2.50 | 2 | 7.69 | 0.186 | 0.442 | (B) BRAF Class 2 Melanoma |
| WAS | Xp11.23 | 6 | 2.59 | 2 | 7.69 | 0.187 | 0.443 | (B) BRAF Class 2 Melanoma |
| EPHB4 | 7q22.1 | 1 | 0.80 | 1 | 7.14 | 0.192 | 0.443 | (B) BRAF Class 2 Melanoma |
| FAT3 | 11q14.3 | 8 | 7.41 | 3 | 16.67 | 0.193 | 0.443 | (B) BRAF Class 2 Melanoma |
| TAL1 | 1p33 | 6 | 2.64 | 2 | 7.69 | 0.193 | 0.443 | (B) BRAF Class 2 Melanoma |
| BCL10 | 1p22.3 | 5 | 1.20 | 2 | 3.57 | 0.196 | 0.443 | (B) BRAF Class 2 Melanoma |
| PPP2R1A | 19q13.41 | 15 | 1.99 | 4 | 3.77 | 0.197 | 0.443 | (B) BRAF Class 2 Melanoma |
| AURKC | 19q13.43 | 1 | 2.33 | 0 | 0.00 | 0.197 | 0.443 | (A) BRAF Class 1 Melanoma |
| CSMD3 | 8q23.3 | 1 | 2.33 | 0 | 0.00 | 0.197 | 0.443 | (A) BRAF Class 1 Melanoma |
| CYP2C19 | 10q23.33 | 1 | 2.33 | 0 | 0.00 | 0.197 | 0.443 | (A) BRAF Class 1 Melanoma |
| DST | 6p12.1 | 1 | 2.33 | 0 | 0.00 | 0.197 | 0.443 | (A) BRAF Class 1 Melanoma |
| FN1 | 2q35 | 1 | 2.33 | 0 | 0.00 | 0.197 | 0.443 | (A) BRAF Class 1 Melanoma |
| FOXP4 | 6p21.1 | 1 | 2.33 | 0 | 0.00 | 0.197 | 0.443 | (A) BRAF Class 1 Melanoma |
| GDNF | 5p13.2 | 1 | 2.33 | 0 | 0.00 | 0.197 | 0.443 | (A) BRAF Class 1 Melanoma |
| GUCY1A2 | 11q22.3 | 1 | 2.33 | 0 | 0.00 | 0.197 | 0.443 | (A) BRAF Class 1 Melanoma |
| IKBKB | 8p11.21 | 1 | 2.33 | 0 | 0.00 | 0.197 | 0.443 | (A) BRAF Class 1 Melanoma |
| ITGA10 | 1q21.1 | 1 | 2.33 | 0 | 0.00 | 0.197 | 0.443 | (A) BRAF Class 1 Melanoma |
| STK36 | 2q35 | 1 | 2.33 | 0 | 0.00 | 0.197 | 0.443 | (A) BRAF Class 1 Melanoma |
| TNK2 | 3q29 | 1 | 2.33 | 0 | 0.00 | 0.197 | 0.443 | (A) BRAF Class 1 Melanoma |
| TRIM24 | 7q33-q34 | 1 | 2.33 | 0 | 0.00 | 0.197 | 0.443 | (A) BRAF Class 1 Melanoma |
| CRTC1 | 19p13.11 | 11 | 3.97 | 3 | 8.57 | 0.199 | 0.443 | (B) BRAF Class 2 Melanoma |
| SAMD9L | 7q21.2 | 2 | 5.00 | 1 | 33.33 | 0.199 | 0.443 | (B) BRAF Class 2 Melanoma |
| FANCL | 2p16.1 | 6 | 1.27 | 2 | 3.64 | 0.2 | 0.443 | (B) BRAF Class 2 Melanoma |
| TMPRSS2 | 21q22.3 | 15 | 2.10 | 4 | 3.96 | 0.2 | 0.443 | (B) BRAF Class 2 Melanoma |

| | | | | | | | | |
|---|---|---|---|---|---|---|---|---|
| ASPSCR1 | 17q25.3 | 1 | 2.27 | 0 | 0.00 | 0.201 | 0.443 | (A) BRAF Class 1 Melanoma |
| PRDM16 | 1p36.32 | 1 | 2.27 | 0 | 0.00 | 0.201 | 0.443 | (A) BRAF Class 1 Melanoma |
| RBM15 | 1p13.3 | 1 | 2.27 | 0 | 0.00 | 0.201 | 0.443 | (A) BRAF Class 1 Melanoma |
| ETV5 | 3q27.2 | 12 | 2.47 | 3 | 5.26 | 0.202 | 0.443 | (B) BRAF Class 2 Melanoma |
| CXCR4 | 2q22.1 | 5 | 0.78 | 2 | 2.27 | 0.204 | 0.443 | (B) BRAF Class 2 Melanoma |
| ARNT | 1q21.3 | 1 | 2.22 | 0 | 0.00 | 0.205 | 0.443 | (A) BRAF Class 1 Melanoma |
| KNL1 | 15q15.1 | 1 | 2.22 | 0 | 0.00 | 0.205 | 0.443 | (A) BRAF Class 1 Melanoma |
| NFKB2 | 10q24.32 | 1 | 2.22 | 0 | 0.00 | 0.205 | 0.443 | (A) BRAF Class 1 Melanoma |
| PDE4DIP | 1q21.2 | 1 | 2.22 | 0 | 0.00 | 0.205 | 0.443 | (A) BRAF Class 1 Melanoma |
| PER1 | 17p13.1 | 1 | 2.22 | 0 | 0.00 | 0.205 | 0.443 | (A) BRAF Class 1 Melanoma |
| TPR | 1q31.1 | 1 | 2.22 | 0 | 0.00 | 0.205 | 0.443 | (A) BRAF Class 1 Melanoma |
| TRIM33 | 1p13.2 | 1 | 2.22 | 0 | 0.00 | 0.205 | 0.443 | (A) BRAF Class 1 Melanoma |
| ZNF384 | 12p13.31 | 1 | 2.22 | 0 | 0.00 | 0.205 | 0.443 | (A) BRAF Class 1 Melanoma |
| GRM3 | 21.11-q21 | 31 | 11.27 | 5 | 18.52 | 0.205 | 0.443 | (B) BRAF Class 2 Melanoma |
| TP53BP1 | 15q15.3 | 18 | 4.42 | 5 | 7.58 | 0.205 | 0.443 | (B) BRAF Class 2 Melanoma |
| B2M | 15q21.1 | 21 | 3.17 | 5 | 5.38 | 0.206 | 0.444 | (B) BRAF Class 2 Melanoma |
| SMARCE1 | 17q21.2 | 1 | 0.43 | 1 | 3.45 | 0.21 | 0.452 | (B) BRAF Class 2 Melanoma |
| COP1 | q25.1-q25 | 5 | 1.09 | 2 | 3.03 | 0.217 | 0.463 | (B) BRAF Class 2 Melanoma |
| MCL1 | 1q21.2 | 5 | 0.65 | 2 | 1.80 | 0.217 | 0.463 | (B) BRAF Class 2 Melanoma |
| ID4 | 6p22.3 | 1 | 0.50 | 1 | 3.85 | 0.217 | 0.463 | (B) BRAF Class 2 Melanoma |
| CDC73 | 1q31.2 | 10 | 1.30 | 3 | 2.70 | 0.218 | 0.464 | (B) BRAF Class 2 Melanoma |
| CRKL | 22q11.21 | 10 | 1.30 | 3 | 2.70 | 0.218 | 0.464 | (B) BRAF Class 2 Melanoma |
| SMYD3 | 1q44 | 4 | 2.34 | 2 | 6.67 | 0.22 | 0.466 | (B) BRAF Class 2 Melanoma |
| PRKCZ | 1p36.33 | 1 | 1.47 | 1 | 11.11 | 0.221 | 0.468 | (B) BRAF Class 2 Melanoma |
| UBR5 | 8q22.3 | 1 | 2.00 | 0 | 0.00 | 0.222 | 0.468 | (A) BRAF Class 1 Melanoma |
| CTNNB1 | 3p22.1 | 67 | 5.68 | 13 | 7.43 | 0.223 | 0.469 | (B) BRAF Class 2 Melanoma |
| DNAJB1 | 19p13.12 | 1 | 0.22 | 1 | 1.64 | 0.227 | 0.476 | (B) BRAF Class 2 Melanoma |
| MDC1 | 6p21.33 | 28 | 6.88 | 6 | 10.53 | 0.227 | 0.476 | (B) BRAF Class 2 Melanoma |
| SDHAF2 | 11q12.2 | 1 | 0.15 | 1 | 1.05 | 0.23 | 0.481 | (B) BRAF Class 2 Melanoma |
| HIST1H3H | 6p22.1 | 1 | 0.26 | 1 | 1.82 | 0.231 | 0.482 | (B) BRAF Class 2 Melanoma |
| EML4 | 2p21 | 4 | 2.15 | 1 | 10.00 | 0.233 | 0.483 | (B) BRAF Class 2 Melanoma |
| AMER1 | Xq11.2 | 22 | 4.09 | 5 | 6.58 | 0.233 | 0.483 | (B) BRAF Class 2 Melanoma |
| FLI1 | 11q24.3 | 4 | 2.17 | 1 | 10.00 | 0.235 | 0.485 | (B) BRAF Class 2 Melanoma |
| CIC | 19q13.2 | 40 | 5.56 | 8 | 7.84 | 0.235 | 0.485 | (B) BRAF Class 2 Melanoma |
| BRD4 | 19p13.12 | 39 | 5.15 | 8 | 7.27 | 0.236 | 0.485 | (B) BRAF Class 2 Melanoma |
| ARID5B | 10q21.2 | 22 | 4.97 | 5 | 7.94 | 0.236 | 0.485 | (B) BRAF Class 2 Melanoma |
| RAD52 | 12p13.33 | 1 | 0.15 | 1 | 1.04 | 0.24 | 0.49 | (B) BRAF Class 2 Melanoma |
| TSHR | 14q31.1 | 16 | 2.21 | 4 | 3.81 | 0.24 | 0.49 | (B) BRAF Class 2 Melanoma |
| DNMT3B | 20q11.21 | 11 | 2.70 | 3 | 5.26 | 0.241 | 0.491 | (B) BRAF Class 2 Melanoma |
| PRKDC | 8q11.21 | 54 | 10.57 | 9 | 14.29 | 0.242 | 0.491 | (B) BRAF Class 2 Melanoma |
| NCKAP5 | 2q21.2 | 2 | 6.25 | 1 | 33.33 | 0.242 | 0.491 | (B) BRAF Class 2 Melanoma |
| TSHZ2 | 20q13.2 | 2 | 6.25 | 1 | 33.33 | 0.242 | 0.491 | (B) BRAF Class 2 Melanoma |
| RAC1 | 7p22.1 | 35 | 4.27 | 7 | 6.14 | 0.244 | 0.493 | (B) BRAF Class 2 Melanoma |
| KLF4 | 9q31.2 | 17 | 2.64 | 4 | 4.49 | 0.244 | 0.493 | (B) BRAF Class 2 Melanoma |
| TAF1 | Xq13.1 | 10 | 4.24 | 2 | 9.52 | 0.256 | 0.514 | (B) BRAF Class 2 Melanoma |

| | | | | | | | | |
|---|---|---|---|---|---|---|---|---|
| RHEB | 7q36.1 | 6 | 0.87 | 2 | 2.08 | 0.256 | 0.514 | (B) BRAF Class 2 Melanoma |
| PLK2 | 5q11.2 | 6 | 1.47 | 2 | 3.51 | 0.256 | 0.514 | (B) BRAF Class 2 Melanoma |
| ANKRD11 | 16q24.3 | 28 | 7.14 | 2 | 3.64 | 0.258 | 0.517 | (A) BRAF Class 1 Melanoma |
| ETS1 | 11q24.3 | 3 | 1.36 | 1 | 5.88 | 0.259 | 0.517 | (B) BRAF Class 2 Melanoma |
| IKBKE | 1q32.1 | 6 | 1.12 | 2 | 2.63 | 0.26 | 0.517 | (B) BRAF Class 2 Melanoma |
| PIK3R1 | 5q13.1 | 17 | 2.10 | 4 | 3.48 | 0.26 | 0.517 | (B) BRAF Class 2 Melanoma |
| PRDM1 | 6q21 | 53 | 6.88 | 10 | 9.01 | 0.261 | 0.517 | (B) BRAF Class 2 Melanoma |
| CASP8 | 2q33.1 | 10 | 1.42 | 0 | 0.00 | 0.265 | 0.525 | (A) BRAF Class 1 Melanoma |
| ESR2 | 4q23.2-q23 | 1 | 0.93 | 1 | 5.56 | 0.266 | 0.526 | (B) BRAF Class 2 Melanoma |
| TENT5C | 1p12 | 17 | 2.21 | 4 | 3.60 | 0.268 | 0.526 | (B) BRAF Class 2 Melanoma |
| TOPBP1 | 3q22.1 | 7 | 3.41 | 2 | 7.69 | 0.268 | 0.526 | (B) BRAF Class 2 Melanoma |
| TET1 | 10q21.3 | 31 | 5.03 | 6 | 7.23 | 0.268 | 0.526 | (B) BRAF Class 2 Melanoma |
| BRIP1 | 17q23.2 | 29 | 3.77 | 6 | 5.41 | 0.272 | 0.531 | (B) BRAF Class 2 Melanoma |
| MITF | 3p13 | 35 | 4.55 | 3 | 2.70 | 0.272 | 0.531 | (A) BRAF Class 1 Melanoma |
| PRF1 | 10q22.1 | 10 | 3.89 | 0 | 0.00 | 0.273 | 0.532 | (A) BRAF Class 1 Melanoma |
| DOCK8 | 9p24.3 | 13 | 6.50 | 3 | 11.54 | 0.274 | 0.532 | (B) BRAF Class 2 Melanoma |
| TTYH1 | 19q13.42 | 2 | 3.70 | 1 | 16.67 | 0.275 | 0.532 | (B) BRAF Class 2 Melanoma |
| EPAS1 | 2p21 | 1 | 0.58 | 1 | 3.33 | 0.277 | 0.532 | (B) BRAF Class 2 Melanoma |
| ERF | 19q13.2 | 1 | 0.58 | 1 | 3.33 | 0.277 | 0.532 | (B) BRAF Class 2 Melanoma |
| PTP4A1 | 6q12 | 1 | 0.58 | 1 | 3.33 | 0.277 | 0.532 | (B) BRAF Class 2 Melanoma |
| SESN3 | 11q21 | 1 | 0.58 | 1 | 3.33 | 0.277 | 0.532 | (B) BRAF Class 2 Melanoma |
| XRCC6 | 22q13.2 | 7 | 3.50 | 2 | 7.69 | 0.277 | 0.532 | (B) BRAF Class 2 Melanoma |
| RB1 | 13q14.2 | 32 | 2.89 | 6 | 4.08 | 0.28 | 0.536 | (B) BRAF Class 2 Melanoma |
| NCOA3 | 20q13.12 | 19 | 4.27 | 1 | 1.64 | 0.281 | 0.536 | (A) BRAF Class 1 Melanoma |
| KMT2C | 7q36.1 | 76 | 14.96 | 13 | 18.31 | 0.282 | 0.537 | (B) BRAF Class 2 Melanoma |
| NTRK2 | 9q21.33 | 18 | 2.24 | 4 | 3.51 | 0.287 | 0.547 | (B) BRAF Class 2 Melanoma |
| VHL | 3p25.3 | 7 | 0.65 | 2 | 1.39 | 0.289 | 0.548 | (B) BRAF Class 2 Melanoma |
| DCUN1D1 | 3q26.33 | 1 | 0.25 | 1 | 1.32 | 0.29 | 0.548 | (B) BRAF Class 2 Melanoma |
| FOXO1 | 13q14.11 | 7 | 1.52 | 2 | 3.23 | 0.291 | 0.548 | (B) BRAF Class 2 Melanoma |
| CHEK2 | 22q12.1 | 18 | 2.24 | 1 | 0.87 | 0.291 | 0.548 | (A) BRAF Class 1 Melanoma |
| SDHA | 5p15.33 | 27 | 3.64 | 2 | 1.94 | 0.291 | 0.548 | (A) BRAF Class 1 Melanoma |
| NFKBIA | 14q13.2 | 9 | 1.19 | 0 | 0.00 | 0.295 | 0.553 | (A) BRAF Class 1 Melanoma |
| ZNRF3 | 22q12.1 | 9 | 3.57 | 0 | 0.00 | 0.295 | 0.553 | (A) BRAF Class 1 Melanoma |
| MED12 | Xq13.1 | 37 | 4.91 | 8 | 6.45 | 0.295 | 0.553 | (B) BRAF Class 2 Melanoma |
| PAX5 | 9p13.2 | 12 | 1.54 | 3 | 2.63 | 0.298 | 0.557 | (B) BRAF Class 2 Melanoma |
| RIT1 | 1q22 | 7 | 1.09 | 2 | 2.25 | 0.3 | 0.559 | (B) BRAF Class 2 Melanoma |
| SMARCA4 | 19p13.2 | 50 | 6.21 | 9 | 7.89 | 0.303 | 0.562 | (B) BRAF Class 2 Melanoma |
| MAPK1 | 22q11.22 | 7 | 0.94 | 2 | 1.92 | 0.303 | 0.562 | (B) BRAF Class 2 Melanoma |
| SBDS | 7q11.21 | 2 | 0.71 | 1 | 2.78 | 0.303 | 0.562 | (B) BRAF Class 2 Melanoma |
| CSDE1 | 1p13.2 | 2 | 0.64 | 1 | 2.50 | 0.304 | 0.562 | (B) BRAF Class 2 Melanoma |
| LATS1 | 6q25.1 | 7 | 1.72 | 2 | 3.51 | 0.305 | 0.562 | (B) BRAF Class 2 Melanoma |
| INHA | 2q35 | 7 | 1.79 | 2 | 3.64 | 0.306 | 0.562 | (B) BRAF Class 2 Melanoma |
| KAT6B | 10q22.2 | 16 | 7.05 | 3 | 11.54 | 0.308 | 0.562 | (B) BRAF Class 2 Melanoma |
| FANCD2 | 3p25.3 | 23 | 4.47 | 4 | 6.67 | 0.308 | 0.562 | (B) BRAF Class 2 Melanoma |
| FAAP20 | 1p36.33 | 2 | 1.00 | 1 | 3.85 | 0.308 | 0.562 | (B) BRAF Class 2 Melanoma |

| | | | | | | | | |
|---|---|---|---|---|---|---|---|---|
| UROD | 1p34.1 | 2 | 1.00 | 1 | 3.85 | 0.308 | 0.562 | (B) BRAF Class 2 Melanoma |
| KIF1B | 1p36.22 | 14 | 7.00 | 3 | 11.54 | 0.308 | 0.562 | (B) BRAF Class 2 Melanoma |
| LATS2 | 13q12.11 | 13 | 3.18 | 3 | 5.26 | 0.31 | 0.563 | (B) BRAF Class 2 Melanoma |
| PMS2 | 7p22.1 | 19 | 2.40 | 4 | 3.60 | 0.311 | 0.564 | (B) BRAF Class 2 Melanoma |
| STAT6 | 12q13.3 | 9 | 2.92 | 0 | 0.00 | 0.312 | 0.565 | (A) BRAF Class 1 Melanoma |
| CDK6 | 7q21.2 | 7 | 0.87 | 2 | 1.74 | 0.313 | 0.565 | (B) BRAF Class 2 Melanoma |
| DMD | p21.2-p21. | 2 | 2.94 | 1 | 11.11 | 0.315 | 0.568 | (B) BRAF Class 2 Melanoma |
| FLCN | 17p11.2 | 7 | 0.92 | 2 | 1.82 | 0.315 | 0.568 | (B) BRAF Class 2 Melanoma |
| GATA2 | 3q21.3 | 13 | 1.75 | 3 | 2.86 | 0.316 | 0.568 | (B) BRAF Class 2 Melanoma |
| LMO1 | 11p15.4 | 7 | 0.98 | 2 | 1.92 | 0.32 | 0.574 | (B) BRAF Class 2 Melanoma |
| EIF1AX | Xp22.12 | 7 | 1.53 | 2 | 2.99 | 0.321 | 0.575 | (B) BRAF Class 2 Melanoma |
| RBBP8 | 18q11.2 | 8 | 4.00 | 2 | 7.69 | 0.323 | 0.575 | (B) BRAF Class 2 Melanoma |
| MAX | 14q23.3 | 2 | 0.31 | 1 | 1.11 | 0.323 | 0.575 | (B) BRAF Class 2 Melanoma |
| PIK3CB | 3q22.3 | 14 | 2.64 | 3 | 4.23 | 0.325 | 0.575 | (B) BRAF Class 2 Melanoma |
| TGFBR1 | 9q22.33 | 2 | 0.49 | 1 | 1.75 | 0.326 | 0.575 | (B) BRAF Class 2 Melanoma |
| HIST1H3A | 6p22.2 | 2 | 0.51 | 1 | 1.82 | 0.326 | 0.575 | (B) BRAF Class 2 Melanoma |
| HIST1H3F | 6p22.2 | 2 | 0.51 | 1 | 1.82 | 0.326 | 0.575 | (B) BRAF Class 2 Melanoma |
| HIST1H3J | 6p22.1 | 2 | 0.51 | 1 | 1.82 | 0.326 | 0.575 | (B) BRAF Class 2 Melanoma |
| CCND3 | 6p21.1 | 2 | 0.25 | 1 | 0.88 | 0.327 | 0.575 | (B) BRAF Class 2 Melanoma |
| SDHD | 11q23.1 | 2 | 0.26 | 1 | 0.92 | 0.33 | 0.58 | (B) BRAF Class 2 Melanoma |
| AKT1 | 14q32.33 | 16 | 1.36 | 1 | 0.57 | 0.334 | 0.586 | (A) BRAF Class 1 Melanoma |
| GLI2 | 2q14.2 | 32 | 11.19 | 6 | 14.63 | 0.335 | 0.587 | (B) BRAF Class 2 Melanoma |
| IGF1 | 12q23.2 | 2 | 0.43 | 1 | 1.45 | 0.342 | 0.598 | (B) BRAF Class 2 Melanoma |
| U2AF1 | 21q22.3 | 8 | 1.00 | 0 | 0.00 | 0.343 | 0.598 | (A) BRAF Class 1 Melanoma |
| XPC | 3p25.1 | 9 | 3.19 | 0 | 0.00 | 0.344 | 0.599 | (A) BRAF Class 1 Melanoma |
| INSR | 19p13.2 | 20 | 4.23 | 4 | 5.97 | 0.346 | 0.602 | (B) BRAF Class 2 Melanoma |
| RHOA | 3p21.31 | 8 | 1.18 | 2 | 2.13 | 0.349 | 0.606 | (B) BRAF Class 2 Melanoma |
| SIN3A | 15q24.2 | 3 | 5.56 | 1 | 16.67 | 0.351 | 0.607 | (B) BRAF Class 2 Melanoma |
| AKT3 | 1q43-q44 | 14 | 1.73 | 3 | 2.63 | 0.352 | 0.607 | (B) BRAF Class 2 Melanoma |
| ALOX12B | 17p13.1 | 13 | 2.49 | 3 | 3.80 | 0.353 | 0.607 | (B) BRAF Class 2 Melanoma |
| MAP3K4 | 6q26 | 6 | 3.77 | 1 | 10.00 | 0.353 | 0.607 | (B) BRAF Class 2 Melanoma |
| KEAP1 | 19p13.2 | 8 | 1.05 | 2 | 1.87 | 0.354 | 0.608 | (B) BRAF Class 2 Melanoma |
| RBM10 | Xp11.3 | 16 | 2.27 | 1 | 1.00 | 0.355 | 0.608 | (A) BRAF Class 1 Melanoma |
| PARP1 | 1q42.12 | 8 | 1.61 | 2 | 2.86 | 0.356 | 0.608 | (B) BRAF Class 2 Melanoma |
| MYB | 6q23.3 | 9 | 1.75 | 0 | 0.00 | 0.361 | 0.616 | (A) BRAF Class 1 Melanoma |
| CCNE1 | 19q12 | 8 | 0.99 | 2 | 1.74 | 0.361 | 0.616 | (B) BRAF Class 2 Melanoma |
| CRLF2 | 2.33 and Yp | 14 | 1.88 | 3 | 2.80 | 0.363 | 0.617 | (B) BRAF Class 2 Melanoma |
| ESR1 | q25.1-q25. | 18 | 2.09 | 4 | 2.88 | 0.365 | 0.619 | (B) BRAF Class 2 Melanoma |
| POLH | 6p21.1 | 3 | 1.38 | 1 | 3.85 | 0.366 | 0.62 | (B) BRAF Class 2 Melanoma |
| ERCC6 | 10q11.23 | 9 | 4.50 | 2 | 7.69 | 0.368 | 0.623 | (B) BRAF Class 2 Melanoma |
| KLLN | 10q23.31 | 8 | 4.00 | 0 | 0.00 | 0.37 | 0.625 | (A) BRAF Class 1 Melanoma |
| SMC3 | 10q25.2 | 9 | 1.97 | 0 | 0.00 | 0.375 | 0.631 | (A) BRAF Class 1 Melanoma |
| MAP3K1 | 5q11.2 | 27 | 3.51 | 5 | 4.50 | 0.378 | 0.631 | (B) BRAF Class 2 Melanoma |
| POLD1 | 19q13.33 | 27 | 3.82 | 5 | 4.90 | 0.379 | 0.631 | (B) BRAF Class 2 Melanoma |
| PAX8 | 2q14.1 | 6 | 2.64 | 1 | 6.25 | 0.383 | 0.631 | (B) BRAF Class 2 Melanoma |

| | | | | | | | | |
|---|---|---|---|---|---|---|---|---|
| ZMYM3 | Xq13.1 | 4 | 5.63 | 1 | 14.29 | 0.383 | 0.631 | (B) BRAF Class 2 Melanoma |
| ARFRP1 | 20q13.33 | 4 | 1.39 | 1 | 3.45 | 0.384 | 0.631 | (B) BRAF Class 2 Melanoma |
| KLHL6 | 3q27.1 | 10 | 3.40 | 0 | 0.00 | 0.385 | 0.631 | (A) BRAF Class 1 Melanoma |
| IDH2 | 15q26.1 | 8 | 0.72 | 2 | 1.20 | 0.386 | 0.631 | (B) BRAF Class 2 Melanoma |
| CARM1 | 19p13.2 | 2 | 1.17 | 1 | 3.33 | 0.386 | 0.631 | (B) BRAF Class 2 Melanoma |
| CCNQ | Xq28 | 2 | 1.17 | 1 | 3.33 | 0.386 | 0.631 | (B) BRAF Class 2 Melanoma |
| RXRA | 9q34.2 | 2 | 1.17 | 1 | 3.33 | 0.386 | 0.631 | (B) BRAF Class 2 Melanoma |
| TAP1 | 6p21.32 | 2 | 1.17 | 1 | 3.33 | 0.386 | 0.631 | (B) BRAF Class 2 Melanoma |
| PTK2B | 8p21.2 | 9 | 4.21 | 2 | 6.90 | 0.386 | 0.631 | (B) BRAF Class 2 Melanoma |
| MSH6 | 2p16.3 | 28 | 3.47 | 5 | 4.39 | 0.387 | 0.631 | (B) BRAF Class 2 Melanoma |
| EXT1 | 8q24.11 | 8 | 2.84 | 0 | 0.00 | 0.388 | 0.631 | (A) BRAF Class 1 Melanoma |
| EXT2 | 11p11.2 | 8 | 2.84 | 0 | 0.00 | 0.388 | 0.631 | (A) BRAF Class 1 Melanoma |
| SMAD2 | 18q21.1 | 7 | 0.91 | 0 | 0.00 | 0.388 | 0.631 | (A) BRAF Class 1 Melanoma |
| BABAM2 | 2p23.2 | 3 | 1.50 | 1 | 3.85 | 0.389 | 0.631 | (B) BRAF Class 2 Melanoma |
| DCLRE1C | 10p13 | 3 | 1.50 | 1 | 3.85 | 0.389 | 0.631 | (B) BRAF Class 2 Melanoma |
| DDB1 | 11q12.2 | 3 | 1.50 | 1 | 3.85 | 0.389 | 0.631 | (B) BRAF Class 2 Melanoma |
| HMBS | 11q23.3 | 3 | 1.50 | 1 | 3.85 | 0.389 | 0.631 | (B) BRAF Class 2 Melanoma |
| MUS81 | 11q13.1 | 3 | 1.50 | 1 | 3.85 | 0.389 | 0.631 | (B) BRAF Class 2 Melanoma |
| BUB1B | 15q15.1 | 10 | 3.55 | 2 | 5.71 | 0.39 | 0.632 | (B) BRAF Class 2 Melanoma |
| FGF7 | 15q21.2 | 4 | 1.88 | 1 | 4.55 | 0.391 | 0.632 | (B) BRAF Class 2 Melanoma |
| PRKN | 6q26 | 34 | 4.50 | 6 | 5.50 | 0.392 | 0.633 | (B) BRAF Class 2 Melanoma |
| MDM4 | 1q32.1 | 15 | 1.86 | 3 | 2.61 | 0.395 | 0.636 | (B) BRAF Class 2 Melanoma |
| NUP214 | 9q34.13 | 1 | 2.00 | 1 | 0.57 | 0.396 | 0.636 | (A) BRAF Class 1 Melanoma |
| MSH2 | 2p21-p16.3 | 15 | 1.92 | 3 | 2.68 | 0.396 | 0.636 | (B) BRAF Class 2 Melanoma |
| BAP1 | 3p21.1 | 28 | 3.48 | 5 | 4.35 | 0.397 | 0.636 | (B) BRAF Class 2 Melanoma |
| HSD3B1 | 1p12 | 5 | 2.09 | 1 | 4.76 | 0.4 | 0.639 | (B) BRAF Class 2 Melanoma |
| FBXW7 | 4q31.3 | 30 | 2.67 | 5 | 3.31 | 0.4 | 0.639 | (B) BRAF Class 2 Melanoma |
| STAT5A | 17q21.2 | 3 | 0.73 | 1 | 1.79 | 0.403 | 0.642 | (B) BRAF Class 2 Melanoma |
| GSK3B | 3q13.33 | 9 | 1.68 | 2 | 2.63 | 0.405 | 0.644 | (B) BRAF Class 2 Melanoma |
| ERRFI1 | 1p36.23 | 9 | 1.77 | 2 | 2.78 | 0.405 | 0.644 | (B) BRAF Class 2 Melanoma |
| MYOD1 | 11p15.1 | 3 | 0.70 | 1 | 1.67 | 0.409 | 0.646 | (B) BRAF Class 2 Melanoma |
| EIF4E | 4q23 | 3 | 0.77 | 1 | 1.82 | 0.41 | 0.646 | (B) BRAF Class 2 Melanoma |
| HIST1H3D | 6p22.2 | 3 | 0.77 | 1 | 1.82 | 0.41 | 0.646 | (B) BRAF Class 2 Melanoma |
| TMEM127 | 2q11.2 | 3 | 0.49 | 1 | 1.16 | 0.41 | 0.646 | (B) BRAF Class 2 Melanoma |
| CALR | 19p13.13 | 7 | 1.07 | 0 | 0.00 | 0.411 | 0.646 | (A) BRAF Class 1 Melanoma |
| EED | 11q14.2 | 7 | 1.40 | 0 | 0.00 | 0.412 | 0.646 | (A) BRAF Class 1 Melanoma |
| RAD54B | 8q22.1 | 10 | 5.00 | 2 | 7.69 | 0.412 | 0.646 | (B) BRAF Class 2 Melanoma |
| ELOC | 8q21.11 | 3 | 0.49 | 1 | 1.15 | 0.414 | 0.648 | (B) BRAF Class 2 Melanoma |
| CDKN1A | 6p21.2 | 3 | 0.40 | 1 | 0.92 | 0.417 | 0.653 | (B) BRAF Class 2 Melanoma |
| HABP2 | 10q25.3 | 7 | 3.50 | 0 | 0.00 | 0.42 | 0.653 | (A) BRAF Class 1 Melanoma |
| PNKP | 19q13.33 | 7 | 3.50 | 0 | 0.00 | 0.42 | 0.653 | (A) BRAF Class 1 Melanoma |
| TRIM37 | 17q22 | 7 | 3.50 | 0 | 0.00 | 0.42 | 0.653 | (A) BRAF Class 1 Melanoma |
| PPM1D | 17q23.2 | 14 | 2.30 | 1 | 1.15 | 0.422 | 0.656 | (A) BRAF Class 1 Melanoma |
| RUNX1 | 21q22.12 | 22 | 2.86 | 4 | 3.57 | 0.425 | 0.658 | (B) BRAF Class 2 Melanoma |
| STAT3 | 17q21.2 | 14 | 1.77 | 1 | 0.89 | 0.425 | 0.658 | (A) BRAF Class 1 Melanoma |

| | | | | | | | | |
|---|---|---|---|---|---|---|---|---|
| RAD50 | 5q31.1 | 23 | 3.09 | 4 | 3.85 | 0.427 | 0.66 | (B) BRAF Class 2 Melanoma |
| EZH2 | 7q36.1 | 31 | 3.01 | 5 | 3.60 | 0.429 | 0.661 | (B) BRAF Class 2 Melanoma |
| ACVR1B | 12q13.13 | 5 | 1.82 | 1 | 3.70 | 0.433 | 0.666 | (B) BRAF Class 2 Melanoma |
| BCL6 | 3q27.3 | 16 | 2.08 | 3 | 2.70 | 0.436 | 0.67 | (B) BRAF Class 2 Melanoma |
| PPP6C | 9q33.3 | 36 | 8.28 | 4 | 6.56 | 0.438 | 0.671 | (A) BRAF Class 1 Melanoma |
| FGF19 | 11q13.3 | 10 | 1.75 | 2 | 2.53 | 0.439 | 0.672 | (B) BRAF Class 2 Melanoma |
| EMSY | 11q13.5 | 5 | 1.73 | 1 | 3.45 | 0.439 | 0.672 | (B) BRAF Class 2 Melanoma |
| ACVR1 | 2q24.1 | 10 | 1.59 | 2 | 2.30 | 0.441 | 0.672 | (B) BRAF Class 2 Melanoma |
| DDR2 | 1q23.3 | 20 | 2.28 | 4 | 2.80 | 0.441 | 0.672 | (B) BRAF Class 2 Melanoma |
| PTCH2 | 1p34.1 | 3 | 2.91 | 1 | 6.25 | 0.443 | 0.672 | (B) BRAF Class 2 Melanoma |
| FANCE | 6p21.31 | 7 | 1.36 | 0 | 0.00 | 0.444 | 0.672 | (A) BRAF Class 1 Melanoma |
| PTEN | 10q23.31 | 172 | 14.58 | 24 | 15.29 | 0.445 | 0.672 | (B) BRAF Class 2 Melanoma |
| ELF3 | 1q32.1 | 5 | 2.42 | 0 | 0.00 | 0.445 | 0.672 | (A) BRAF Class 1 Melanoma |
| TFE3 | Xp11.23 | 5 | 1.22 | 1 | 2.38 | 0.446 | 0.672 | (B) BRAF Class 2 Melanoma |
| SRSF2 | 17q25.1 | 10 | 1.45 | 2 | 2.06 | 0.446 | 0.672 | (B) BRAF Class 2 Melanoma |
| TSC2 | 16p13.3 | 30 | 3.71 | 5 | 4.35 | 0.447 | 0.672 | (B) BRAF Class 2 Melanoma |
| NUP93 | 16q13 | 6 | 1.17 | 0 | 0.00 | 0.447 | 0.672 | (A) BRAF Class 1 Melanoma |
| CHD2 | 15q26.1 | 14 | 5.30 | 2 | 7.41 | 0.449 | 0.674 | (B) BRAF Class 2 Melanoma |
| FUS | 16p11.2 | 7 | 2.27 | 0 | 0.00 | 0.45 | 0.674 | (A) BRAF Class 1 Melanoma |
| TRAF3 | 14q32.32 | 4 | 1.54 | 1 | 3.03 | 0.452 | 0.676 | (B) BRAF Class 2 Melanoma |
| GNAS | 20q13.32 | 14 | 1.24 | 1 | 0.66 | 0.454 | 0.678 | (A) BRAF Class 1 Melanoma |
| CUL3 | 2q36.2 | 6 | 1.13 | 0 | 0.00 | 0.455 | 0.678 | (A) BRAF Class 1 Melanoma |
| RTEL1 | 20q13.33 | 8 | 4.68 | 2 | 6.67 | 0.456 | 0.679 | (B) BRAF Class 2 Melanoma |
| SH2B3 | 12q24.12 | 20 | 2.68 | 2 | 1.85 | 0.458 | 0.679 | (A) BRAF Class 1 Melanoma |
| PRKAR1A | 17q24.2 | 10 | 1.30 | 2 | 1.80 | 0.459 | 0.679 | (B) BRAF Class 2 Melanoma |
| EXO1 | 1q43 | 4 | 2.00 | 1 | 3.85 | 0.46 | 0.679 | (B) BRAF Class 2 Melanoma |
| HFE | 6p22.2 | 4 | 2.00 | 1 | 3.85 | 0.46 | 0.679 | (B) BRAF Class 2 Melanoma |
| SLC25A13 | 7q21.3 | 4 | 2.00 | 1 | 3.85 | 0.46 | 0.679 | (B) BRAF Class 2 Melanoma |
| TRAF2 | 9q34.3 | 6 | 1.47 | 0 | 0.00 | 0.461 | 0.679 | (A) BRAF Class 1 Melanoma |
| GLI3 | 7p14.1 | 6 | 8.82 | 0 | 0.00 | 0.462 | 0.679 | (A) BRAF Class 1 Melanoma |
| PALB2 | 16p12.2 | 24 | 2.98 | 4 | 3.51 | 0.466 | 0.684 | (B) BRAF Class 2 Melanoma |
| NPM1 | 5q35.1 | 6 | 0.56 | 0 | 0.00 | 0.47 | 0.689 | (A) BRAF Class 1 Melanoma |
| ABL2 | 1q25.2 | 6 | 2.19 | 1 | 3.85 | 0.473 | 0.691 | (B) BRAF Class 2 Melanoma |
| EPCAM | 2p21 | 4 | 0.61 | 1 | 1.12 | 0.474 | 0.691 | (B) BRAF Class 2 Melanoma |
| CYLD | 16q12.1 | 18 | 2.71 | 3 | 3.30 | 0.474 | 0.691 | (B) BRAF Class 2 Melanoma |
| DIS3L2 | 2q37.1 | 6 | 2.70 | 0 | 0.00 | 0.475 | 0.691 | (A) BRAF Class 1 Melanoma |
| ELANE | 19p13.3 | 6 | 2.70 | 0 | 0.00 | 0.475 | 0.691 | (A) BRAF Class 1 Melanoma |
| NEIL3 | 4q34.3 | 6 | 3.00 | 0 | 0.00 | 0.476 | 0.692 | (A) BRAF Class 1 Melanoma |
| GPS2 | 17p13.1 | 4 | 1.00 | 1 | 1.82 | 0.477 | 0.692 | (B) BRAF Class 2 Melanoma |
| MSI1 | 12q24.31 | 3 | 1.75 | 1 | 3.33 | 0.479 | 0.692 | (B) BRAF Class 2 Melanoma |
| ERBB3 | 12q13.2 | 21 | 2.60 | 4 | 3.01 | 0.48 | 0.692 | (B) BRAF Class 2 Melanoma |
| CYSLTR2 | 13q14.2 | 3 | 1.44 | 1 | 2.70 | 0.481 | 0.692 | (B) BRAF Class 2 Melanoma |
| ID3 | 1p36.12 | 4 | 0.57 | 1 | 1.02 | 0.482 | 0.692 | (B) BRAF Class 2 Melanoma |
| EGFL7 | 9q34.3 | 4 | 0.98 | 1 | 1.75 | 0.482 | 0.692 | (B) BRAF Class 2 Melanoma |
| IL10 | 1q32.1 | 4 | 0.98 | 1 | 1.75 | 0.482 | 0.692 | (B) BRAF Class 2 Melanoma |

| | | | | | | | | |
|---|---|---|---|---|---|---|---|---|
| HIST1H3E | 6p22.2 | 4 | 1.02 | 1 | 1.82 | 0.483 | 0.692 | (B) BRAF Class 2 Melanoma |
| NKX3-1 | 8p21.2 | 4 | 0.68 | 1 | 1.20 | 0.484 | 0.692 | (B) BRAF Class 2 Melanoma |
| KNSTRN | 15q15.1 | 15 | 6.20 | 3 | 7.50 | 0.484 | 0.692 | (B) BRAF Class 2 Melanoma |
| CDKN1B | 12p13.1 | 4 | 0.50 | 1 | 0.88 | 0.485 | 0.693 | (B) BRAF Class 2 Melanoma |
| SDHC | 1q23.3 | 4 | 0.53 | 1 | 0.93 | 0.487 | 0.694 | (B) BRAF Class 2 Melanoma |
| H3F3A | 1q42.12 | 4 | 0.50 | 1 | 0.88 | 0.489 | 0.695 | (B) BRAF Class 2 Melanoma |
| FGF23 | 12p13.32 | 6 | 2.08 | 1 | 3.45 | 0.491 | 0.697 | (B) BRAF Class 2 Melanoma |
| CD79B | 17q23.3 | 4 | 0.52 | 1 | 0.90 | 0.492 | 0.697 | (B) BRAF Class 2 Melanoma |
| CDKN2C | 1p32.3 | 4 | 0.52 | 1 | 0.90 | 0.492 | 0.697 | (B) BRAF Class 2 Melanoma |
| CCN6 | 6q21 | 6 | 2.16 | 1 | 3.57 | 0.493 | 0.697 | (B) BRAF Class 2 Melanoma |
| DOT1L | 19p13.3 | 25 | 4.66 | 4 | 5.26 | 0.495 | 0.698 | (B) BRAF Class 2 Melanoma |
| HLA-A | 6p22.1 | 5 | 1.12 | 0 | 0.00 | 0.495 | 0.698 | (A) BRAF Class 1 Melanoma |
| FANCG | 9p13.3 | 5 | 0.95 | 1 | 1.56 | 0.499 | 0.702 | (B) BRAF Class 2 Melanoma |
| ZNF703 | 8p11.23 | 7 | 2.42 | 0 | 0.00 | 0.509 | 0.715 | (A) BRAF Class 1 Melanoma |
| CEBPA | 19q13.11 | 5 | 0.66 | 0 | 0.00 | 0.511 | 0.716 | (A) BRAF Class 1 Melanoma |
| POLE | 12q24.33 | 61 | 8.05 | 8 | 7.48 | 0.511 | 0.716 | (A) BRAF Class 1 Melanoma |
| BCL2L1 | 20q11.21 | 5 | 0.66 | 0 | 0.00 | 0.513 | 0.718 | (A) BRAF Class 1 Melanoma |
| CDK4 | 12q14.1 | 28 | 3.48 | 4 | 3.01 | 0.516 | 0.72 | (A) BRAF Class 1 Melanoma |
| MAPK3 | 16p11.2 | 5 | 1.28 | 0 | 0.00 | 0.517 | 0.72 | (A) BRAF Class 1 Melanoma |
| XIAP | Xq25 | 5 | 1.23 | 0 | 0.00 | 0.518 | 0.72 | (A) BRAF Class 1 Melanoma |
| TRAF7 | 16p13.3 | 12 | 1.86 | 1 | 1.12 | 0.518 | 0.72 | (A) BRAF Class 1 Melanoma |
| TDG | 12q23.3 | 5 | 2.50 | 1 | 3.85 | 0.524 | 0.727 | (B) BRAF Class 2 Melanoma |
| CSF1R | 5q32 | 28 | 3.05 | 3 | 2.56 | 0.529 | 0.731 | (A) BRAF Class 1 Melanoma |
| ERCC2 | 19q13.32 | 26 | 3.29 | 4 | 3.54 | 0.529 | 0.731 | (B) BRAF Class 2 Melanoma |
| SLITRK6 | 13q31.1 | 6 | 8.11 | 0 | 0.00 | 0.529 | 0.731 | (A) BRAF Class 1 Melanoma |
| IKZF3 | 7q12-q21. | 5 | 4.13 | 1 | 6.25 | 0.532 | 0.734 | (B) BRAF Class 2 Melanoma |
| FANCM | 14q21.2 | 12 | 4.43 | 2 | 5.26 | 0.533 | 0.734 | (B) BRAF Class 2 Melanoma |
| PDCD1 | 2q37.3 | 5 | 1.14 | 1 | 1.69 | 0.534 | 0.735 | (B) BRAF Class 2 Melanoma |
| FGF4 | 11q13.3 | 12 | 2.23 | 2 | 2.63 | 0.535 | 0.735 | (B) BRAF Class 2 Melanoma |
| HIST1H1C | 6p22.2 | 5 | 1.18 | 1 | 1.72 | 0.539 | 0.737 | (B) BRAF Class 2 Melanoma |
| FAH | 15q25.1 | 5 | 2.50 | 0 | 0.00 | 0.54 | 0.737 | (A) BRAF Class 1 Melanoma |
| MBD4 | 3q21.3 | 5 | 2.50 | 0 | 0.00 | 0.54 | 0.737 | (A) BRAF Class 1 Melanoma |
| NAV3 | 12q21.2 | 6 | 11.11 | 1 | 16.67 | 0.541 | 0.738 | (B) BRAF Class 2 Melanoma |
| FANCC | 9q22.32 | 11 | 1.43 | 1 | 0.90 | 0.541 | 0.738 | (A) BRAF Class 1 Melanoma |
| UIMC1 | 5q35.2 | 5 | 2.48 | 0 | 0.00 | 0.543 | 0.738 | (A) BRAF Class 1 Melanoma |
| MYC | 8q24.21 | 23 | 2.85 | 4 | 3.01 | 0.545 | 0.738 | (B) BRAF Class 2 Melanoma |
| JUN | 1p32.1 | 5 | 1.00 | 1 | 1.43 | 0.545 | 0.738 | (B) BRAF Class 2 Melanoma |
| RAD51D | 17q12 | 5 | 0.67 | 1 | 0.95 | 0.546 | 0.738 | (B) BRAF Class 2 Melanoma |
| VTCN1 | lp13.1-p12 | 5 | 1.23 | 1 | 1.75 | 0.547 | 0.738 | (B) BRAF Class 2 Melanoma |
| PIK3R2 | 19p13.11 | 11 | 2.09 | 1 | 1.33 | 0.547 | 0.738 | (A) BRAF Class 1 Melanoma |
| RAB35 | 12q24.23 | 5 | 1.28 | 1 | 1.82 | 0.547 | 0.738 | (B) BRAF Class 2 Melanoma |
| RANBP2 | 2q13 | 20 | 8.77 | 2 | 10.00 | 0.549 | 0.739 | (B) BRAF Class 2 Melanoma |
| GATA1 | Xp11.23 | 5 | 0.94 | 1 | 1.33 | 0.55 | 0.739 | (B) BRAF Class 2 Melanoma |
| PAK1 | .q13.5-q14 | 5 | 1.13 | 1 | 1.59 | 0.552 | 0.74 | (B) BRAF Class 2 Melanoma |
| SOCS1 | 16p13.13 | 5 | 0.65 | 1 | 0.90 | 0.555 | 0.744 | (B) BRAF Class 2 Melanoma |

| | | | | | | | | |
|---|---|---|---|---|---|---|---|---|
| DROSHA | 5p13.3 | 9 | 4.66 | 1 | 3.03 | 0.556 | 0.745 | (A) BRAF Class 1 Melanoma |
| BCORL1 | Xq26.1 | 28 | 5.32 | 3 | 4.69 | 0.56 | 0.748 | (A) BRAF Class 1 Melanoma |
| CD274 | 9p24.1 | 5 | 0.66 | 1 | 0.90 | 0.561 | 0.748 | (B) BRAF Class 2 Melanoma |
| FGF3 | 11q13.3 | 13 | 2.27 | 2 | 2.53 | 0.561 | 0.748 | (B) BRAF Class 2 Melanoma |
| NSD2 | 4p16.3 | 12 | 2.81 | 2 | 3.13 | 0.565 | 0.751 | (B) BRAF Class 2 Melanoma |
| GNAQ | 9q21.2 | 12 | 1.07 | 2 | 1.18 | 0.567 | 0.753 | (B) BRAF Class 2 Melanoma |
| RELA | 11q13.1 | 6 | 2.37 | 1 | 3.13 | 0.57 | 0.754 | (B) BRAF Class 2 Melanoma |
| RPA1 | 17p13.3 | 5 | 1.97 | 1 | 2.63 | 0.57 | 0.754 | (B) BRAF Class 2 Melanoma |
| TLR4 | 9q33.1 | 7 | 7.07 | 0 | 0.00 | 0.571 | 0.754 | (A) BRAF Class 1 Melanoma |
| CCND1 | 11q13.3 | 9 | 1.13 | 1 | 0.75 | 0.571 | 0.754 | (A) BRAF Class 1 Melanoma |
| FAS | 10q23.31 | 5 | 1.03 | 0 | 0.00 | 0.573 | 0.754 | (A) BRAF Class 1 Melanoma |
| MTAP | 9p21.3 | 19 | 8.88 | 2 | 7.41 | 0.574 | 0.754 | (A) BRAF Class 1 Melanoma |
| ETV4 | 17q21.31 | 5 | 1.03 | 0 | 0.00 | 0.574 | 0.754 | (A) BRAF Class 1 Melanoma |
| CTCF | 16q22.1 | 13 | 1.81 | 2 | 1.96 | 0.575 | 0.754 | (B) BRAF Class 2 Melanoma |
| INSRR | 1q23.1 | 4 | 0.34 | 0 | 0.00 | 0.575 | 0.754 | (A) BRAF Class 1 Melanoma |
| IRS2 | 13q34 | 28 | 5.20 | 4 | 5.26 | 0.576 | 0.754 | (B) BRAF Class 2 Melanoma |
| BRD3 | 9q34.2 | 7 | 3.02 | 1 | 3.85 | 0.578 | 0.754 | (B) BRAF Class 2 Melanoma |
| FAN1 | 15q13.3 | 6 | 3.00 | 1 | 3.85 | 0.58 | 0.754 | (B) BRAF Class 2 Melanoma |
| HELQ | 4q21.23 | 6 | 3.00 | 1 | 3.85 | 0.58 | 0.754 | (B) BRAF Class 2 Melanoma |
| SOX2 | 3q26.33 | 4 | 0.52 | 0 | 0.00 | 0.58 | 0.754 | (A) BRAF Class 1 Melanoma |
| STAG2 | Xq25 | 23 | 2.99 | 3 | 2.70 | 0.58 | 0.754 | (A) BRAF Class 1 Melanoma |
| NKX2-1 | 14q13.3 | 10 | 1.30 | 1 | 0.89 | 0.583 | 0.757 | (A) BRAF Class 1 Melanoma |
| QKI | 6q26 | 5 | 1.08 | 0 | 0.00 | 0.586 | 0.759 | (A) BRAF Class 1 Melanoma |
| ARHGAP35 | 19q13.32 | 14 | 5.86 | 2 | 6.25 | 0.586 | 0.759 | (B) BRAF Class 2 Melanoma |
| PIK3R3 | 1p34.1 | 4 | 0.98 | 0 | 0.00 | 0.591 | 0.764 | (A) BRAF Class 1 Melanoma |
| CCND2 | 12p13.32 | 10 | 1.25 | 1 | 0.88 | 0.592 | 0.765 | (A) BRAF Class 1 Melanoma |
| FOXA1 | 14q21.1 | 17 | 2.61 | 2 | 2.25 | 0.595 | 0.766 | (A) BRAF Class 1 Melanoma |
| TGFBR2 | 3p24.1 | 10 | 1.86 | 1 | 1.32 | 0.596 | 0.767 | (A) BRAF Class 1 Melanoma |
| STAG1 | 3q22.3 | 6 | 2.80 | 0 | 0.00 | 0.597 | 0.767 | (A) BRAF Class 1 Melanoma |
| STK19 | 6p21.33 | 8 | 4.68 | 1 | 3.33 | 0.6 | 0.769 | (A) BRAF Class 1 Melanoma |
| MYBL1 | 8q13.1 | 4 | 1.32 | 0 | 0.00 | 0.6 | 0.769 | (A) BRAF Class 1 Melanoma |
| DPYD | 1p21.3 | 4 | 3.81 | 0 | 0.00 | 0.602 | 0.77 | (A) BRAF Class 1 Melanoma |
| YES1 | 18p11.32 | 6 | 1.47 | 1 | 1.75 | 0.603 | 0.77 | (B) BRAF Class 2 Melanoma |
| BCR | 22q11.23 | 10 | 3.92 | 1 | 4.55 | 0.605 | 0.771 | (B) BRAF Class 2 Melanoma |
| MAP3K9 | 14q24.2 | 4 | 6.45 | 0 | 0.00 | 0.608 | 0.775 | (A) BRAF Class 1 Melanoma |
| SHOC2 | 10q25.2 | 3 | 1.75 | 0 | 0.00 | 0.614 | 0.781 | (A) BRAF Class 1 Melanoma |
| TOP2A | 17q21.2 | 9 | 3.41 | 1 | 3.85 | 0.615 | 0.782 | (B) BRAF Class 2 Melanoma |
| SMAD4 | 18q21.2 | 10 | 0.89 | 1 | 0.66 | 0.619 | 0.786 | (A) BRAF Class 1 Melanoma |
| PAK3 | Xq23 | 9 | 3.23 | 1 | 3.57 | 0.621 | 0.787 | (B) BRAF Class 2 Melanoma |
| DUSP4 | 8p12 | 5 | 2.42 | 1 | 2.78 | 0.622 | 0.787 | (B) BRAF Class 2 Melanoma |
| DDB2 | 11p11.2 | 4 | 1.42 | 0 | 0.00 | 0.625 | 0.789 | (A) BRAF Class 1 Melanoma |
| GATA4 | 8p23.1 | 11 | 2.34 | 1 | 1.79 | 0.63 | 0.794 | (A) BRAF Class 1 Melanoma |
| PIK3CD | 1p36.22 | 7 | 1.62 | 1 | 1.75 | 0.632 | 0.796 | (B) BRAF Class 2 Melanoma |
| MALT1 | 18q21.32 | 7 | 1.67 | 1 | 1.79 | 0.636 | 0.801 | (B) BRAF Class 2 Melanoma |
| FANCB | Xp22.2 | 8 | 3.65 | 1 | 3.85 | 0.642 | 0.806 | (B) BRAF Class 2 Melanoma |

| | | | | | | | | |
|---|---|---|---|---|---|---|---|---|
| BIRC3 | 11q22.2 | 7 | 1.67 | 1 | 1.75 | 0.642 | 0.806 | (B) BRAF Class 2 Melanoma |
| FYN | 6q21 | 9 | 2.08 | 1 | 1.64 | 0.644 | 0.807 | (A) BRAF Class 1 Melanoma |
| TOP1 | 20q12 | 9 | 1.62 | 1 | 1.28 | 0.647 | 0.809 | (A) BRAF Class 1 Melanoma |
| SUZ12 | 17q11.2 | 9 | 1.31 | 1 | 1.04 | 0.648 | 0.809 | (A) BRAF Class 1 Melanoma |
| CLDN18 | 3q22.3 | 4 | 7.41 | 0 | 0.00 | 0.649 | 0.809 | (A) BRAF Class 1 Melanoma |
| CD276 | 15q24.1 | 7 | 1.72 | 1 | 1.75 | 0.653 | 0.81 | (B) BRAF Class 2 Melanoma |
| INPP4A | 2q11.2 | 7 | 1.72 | 1 | 1.75 | 0.653 | 0.81 | (B) BRAF Class 2 Melanoma |
| IRS1 | 2q36.3 | 7 | 1.72 | 1 | 1.75 | 0.653 | 0.81 | (B) BRAF Class 2 Melanoma |
| CTLA4 | 2q33.2 | 7 | 1.18 | 1 | 1.20 | 0.653 | 0.81 | (B) BRAF Class 2 Melanoma |
| HIST1H3C | 6p22.2 | 7 | 1.17 | 1 | 1.19 | 0.654 | 0.81 | (B) BRAF Class 2 Melanoma |
| YAP1 | 11q22.1 | 7 | 1.12 | 1 | 1.12 | 0.657 | 0.812 | (B) BRAF Class 2 Melanoma |
| SMARCD1 | 12q13.12 | 3 | 0.65 | 0 | 0.00 | 0.658 | 0.812 | (A) BRAF Class 1 Melanoma |
| RAD54L | 1p34.1 | 3 | 0.64 | 0 | 0.00 | 0.66 | 0.814 | (A) BRAF Class 1 Melanoma |
| MT-ND5 | - | 3 | 0.25 | 0 | 0.00 | 0.66 | 0.814 | (A) BRAF Class 1 Melanoma |
| WRN | 8p12 | 16 | 5.03 | 2 | 4.88 | 0.661 | 0.814 | (A) BRAF Class 1 Melanoma |
| CBFB | 16q22.1 | 3 | 0.42 | 0 | 0.00 | 0.67 | 0.823 | (A) BRAF Class 1 Melanoma |
| ABRAXAS1 | 4q21.23 | 3 | 0.45 | 0 | 0.00 | 0.674 | 0.826 | (A) BRAF Class 1 Melanoma |
| ETV6 | 12p13.2 | 14 | 1.83 | 2 | 1.82 | 0.674 | 0.826 | (A) BRAF Class 1 Melanoma |
| BCL2L11 | 2q13 | 3 | 0.73 | 0 | 0.00 | 0.675 | 0.826 | (A) BRAF Class 1 Melanoma |
| LMO2 | 11p13 | 3 | 1.19 | 0 | 0.00 | 0.676 | 0.826 | (A) BRAF Class 1 Melanoma |
| FAAP100 | 17q25.3 | 9 | 4.50 | 1 | 3.85 | 0.677 | 0.826 | (A) BRAF Class 1 Melanoma |
| KCNQ1 | .p15.5-p15 | 9 | 4.50 | 1 | 3.85 | 0.677 | 0.826 | (A) BRAF Class 1 Melanoma |
| IKZF2 | 2q34 | 4 | 5.63 | 0 | 0.00 | 0.681 | 0.829 | (A) BRAF Class 1 Melanoma |
| DKC1 | Xq28 | 3 | 1.50 | 0 | 0.00 | 0.683 | 0.83 | (A) BRAF Class 1 Melanoma |
| GBA | 1q22 | 3 | 1.50 | 0 | 0.00 | 0.692 | 0.838 | (A) BRAF Class 1 Melanoma |
| NEIL1 | 15q24.2 | 3 | 1.50 | 0 | 0.00 | 0.692 | 0.838 | (A) BRAF Class 1 Melanoma |
| SERPINA1 | 14q32.13 | 3 | 1.50 | 0 | 0.00 | 0.692 | 0.838 | (A) BRAF Class 1 Melanoma |
| XRCC5 | 2q35 | 3 | 1.49 | 0 | 0.00 | 0.694 | 0.839 | (A) BRAF Class 1 Melanoma |
| RAD51C | 17q22 | 8 | 1.07 | 1 | 0.95 | 0.694 | 0.839 | (A) BRAF Class 1 Melanoma |
| USP8 | 15q21.2 | 3 | 1.48 | 0 | 0.00 | 0.695 | 0.839 | (A) BRAF Class 1 Melanoma |
| SF1 | 11q13.1 | 3 | 3.16 | 0 | 0.00 | 0.697 | 0.84 | (A) BRAF Class 1 Melanoma |
| IGF2R | 6q25.3 | 3 | 2.34 | 0 | 0.00 | 0.7 | 0.841 | (A) BRAF Class 1 Melanoma |
| JAZF1 | p15.2-p15. | 3 | 1.18 | 0 | 0.00 | 0.7 | 0.841 | (A) BRAF Class 1 Melanoma |
| PAX7 | 1p36.13 | 5 | 2.25 | 0 | 0.00 | 0.704 | 0.844 | (A) BRAF Class 1 Melanoma |
| SH2D1A | Xq25 | 8 | 1.30 | 1 | 1.20 | 0.708 | 0.848 | (A) BRAF Class 1 Melanoma |
| XRCC1 | 19q13.31 | 3 | 1.38 | 0 | 0.00 | 0.711 | 0.851 | (A) BRAF Class 1 Melanoma |
| AGO2 | 8q24.3 | 6 | 3.51 | 1 | 3.33 | 0.719 | 0.854 | (A) BRAF Class 1 Melanoma |
| PRDM14 | 8q13.3 | 6 | 3.51 | 1 | 3.33 | 0.719 | 0.854 | (A) BRAF Class 1 Melanoma |
| SRC | 20q11.23 | 8 | 0.86 | 1 | 0.83 | 0.721 | 0.854 | (A) BRAF Class 1 Melanoma |
| RHOH | 4p14 | 3 | 1.32 | 0 | 0.00 | 0.721 | 0.854 | (A) BRAF Class 1 Melanoma |
| NUF2 | 1q23.3 | 2 | 1.17 | 0 | 0.00 | 0.723 | 0.854 | (A) BRAF Class 1 Melanoma |
| RAC2 | 22q13.1 | 2 | 1.17 | 0 | 0.00 | 0.723 | 0.854 | (A) BRAF Class 1 Melanoma |
| SESN1 | 6q21 | 2 | 1.17 | 0 | 0.00 | 0.723 | 0.854 | (A) BRAF Class 1 Melanoma |
| )RCS8-MEF. | 19p13.11 | 7 | 0.59 | 1 | 0.57 | 0.723 | 0.854 | (A) BRAF Class 1 Melanoma |
| BRINP3 | 1q31.1 | 5 | 10.64 | 0 | 0.00 | 0.724 | 0.854 | (A) BRAF Class 1 Melanoma |

| | | | | | | | | |
|---|---|---|---|---|---|---|---|---|
| GEN1 | 2p24.2 | 3 | 0.74 | 0 | 0.00 | 0.725 | 0.854 | (A) BRAF Class 1 Melanoma |
| ROBO1 | 3p12.3 | 3 | 5.56 | 0 | 0.00 | 0.725 | 0.854 | (A) BRAF Class 1 Melanoma |
| EZH1 | 17q21.2 | 2 | 0.97 | 0 | 0.00 | 0.725 | 0.854 | (A) BRAF Class 1 Melanoma |
| PML | 15q24.1 | 9 | 3.88 | 1 | 3.85 | 0.734 | 0.862 | (A) BRAF Class 1 Melanoma |
| VEGFA | 6p21.1 | 7 | 1.00 | 1 | 1.00 | 0.735 | 0.862 | (A) BRAF Class 1 Melanoma |
| RAD51B | 14q24.1 | 7 | 1.38 | 1 | 1.37 | 0.735 | 0.862 | (A) BRAF Class 1 Melanoma |
| BABAM1 | 19p13.11 | 2 | 0.57 | 0 | 0.00 | 0.745 | 0.872 | (A) BRAF Class 1 Melanoma |
| PNRC1 | 6q15 | 2 | 0.39 | 0 | 0.00 | 0.752 | 0.88 | (A) BRAF Class 1 Melanoma |
| XRCC3 | 14q32.33 | 2 | 0.79 | 0 | 0.00 | 0.756 | 0.884 | (A) BRAF Class 1 Melanoma |
| HDAC9 | 7p21.1 | 3 | 9.38 | 0 | 0.00 | 0.758 | 0.884 | (A) BRAF Class 1 Melanoma |
| H19 | 11p15.5 | 2 | 0.17 | 0 | 0.00 | 0.758 | 0.884 | (A) BRAF Class 1 Melanoma |
| BBC3 | 19q13.32 | 2 | 0.49 | 0 | 0.00 | 0.769 | 0.894 | (A) BRAF Class 1 Melanoma |
| STK40 | 1p34.3 | 2 | 0.49 | 0 | 0.00 | 0.769 | 0.894 | (A) BRAF Class 1 Melanoma |
| HSP90AA1 | 14q32.31 | 3 | 1.26 | 0 | 0.00 | 0.775 | 0.897 | (A) BRAF Class 1 Melanoma |
| CDK5 | 7q36.1 | 2 | 2.94 | 0 | 0.00 | 0.779 | 0.897 | (A) BRAF Class 1 Melanoma |
| FKBP9 | 7p14.3 | 2 | 2.94 | 0 | 0.00 | 0.779 | 0.897 | (A) BRAF Class 1 Melanoma |
| GSTM5 | 1p13.3 | 2 | 2.94 | 0 | 0.00 | 0.779 | 0.897 | (A) BRAF Class 1 Melanoma |
| RBL2 | 16q12.2 | 2 | 2.94 | 0 | 0.00 | 0.779 | 0.897 | (A) BRAF Class 1 Melanoma |
| ANKRD26 | 10p12.1 | 4 | 2.52 | 0 | 0.00 | 0.782 | 0.897 | (A) BRAF Class 1 Melanoma |
| DMC1 | 22q13.1 | 2 | 1.00 | 0 | 0.00 | 0.783 | 0.897 | (A) BRAF Class 1 Melanoma |
| NR0B1 | Xp21.2 | 2 | 1.00 | 0 | 0.00 | 0.783 | 0.897 | (A) BRAF Class 1 Melanoma |
| OGG1 | 3p25.3 | 2 | 1.00 | 0 | 0.00 | 0.783 | 0.897 | (A) BRAF Class 1 Melanoma |
| RHOT1 | 17q11.2 | 2 | 1.00 | 0 | 0.00 | 0.783 | 0.897 | (A) BRAF Class 1 Melanoma |
| CADM2 | 3p12.1 | 3 | 5.00 | 0 | 0.00 | 0.783 | 0.897 | (A) BRAF Class 1 Melanoma |
| DEPDC5 | !q12.2-q12 | 3 | 5.00 | 0 | 0.00 | 0.783 | 0.897 | (A) BRAF Class 1 Melanoma |
| TAL2 | 9q31.2 | 2 | 0.99 | 0 | 0.00 | 0.784 | 0.897 | (A) BRAF Class 1 Melanoma |
| XPA | 9q22.33 | 2 | 0.71 | 0 | 0.00 | 0.791 | 0.903 | (A) BRAF Class 1 Melanoma |
| MYH9 | 22q12.3 | 3 | 3.70 | 0 | 0.00 | 0.805 | 0.918 | (A) BRAF Class 1 Melanoma |
| CHD1 | ;q15-q21.1 | 2 | 3.70 | 0 | 0.00 | 0.808 | 0.918 | (A) BRAF Class 1 Melanoma |
| SOS2 | 14q21.3 | 2 | 3.70 | 0 | 0.00 | 0.808 | 0.918 | (A) BRAF Class 1 Melanoma |
| ZBTB20 | 3q13.31 | 2 | 3.70 | 0 | 0.00 | 0.808 | 0.918 | (A) BRAF Class 1 Melanoma |
| EPHB6 | 7q34 | 2 | 2.17 | 0 | 0.00 | 0.813 | 0.922 | (A) BRAF Class 1 Melanoma |
| LTK | 15q15.1 | 2 | 1.94 | 0 | 0.00 | 0.816 | 0.922 | (A) BRAF Class 1 Melanoma |
| PDK1 | 2q31.1 | 2 | 0.69 | 0 | 0.00 | 0.826 | 0.922 | (A) BRAF Class 1 Melanoma |
| PPP2R2A | 8p21.2 | 3 | 1.76 | 0 | 0.00 | 0.828 | 0.922 | (A) BRAF Class 1 Melanoma |
| TCF7L1 | 2p11.2 | 2 | 2.15 | 0 | 0.00 | 0.831 | 0.922 | (A) BRAF Class 1 Melanoma |
| DHX15 | 4p15.2 | 3 | 1.89 | 0 | 0.00 | 0.832 | 0.922 | (A) BRAF Class 1 Melanoma |
| FGF1 | 5q31.3 | 3 | 1.89 | 0 | 0.00 | 0.832 | 0.922 | (A) BRAF Class 1 Melanoma |
| FGF9 | 13q12.11 | 3 | 1.89 | 0 | 0.00 | 0.832 | 0.922 | (A) BRAF Class 1 Melanoma |
| TSHZ3 | 19q12 | 2 | 6.25 | 0 | 0.00 | 0.834 | 0.922 | (A) BRAF Class 1 Melanoma |
| PRSS8 | 16p11.2 | 2 | 0.86 | 0 | 0.00 | 0.835 | 0.922 | (A) BRAF Class 1 Melanoma |
| DDX3X | Xp11.4 | 2 | 2.56 | 0 | 0.00 | 0.841 | 0.922 | (A) BRAF Class 1 Melanoma |
| KCNIP1 | 5q35.1 | 2 | 3.33 | 0 | 0.00 | 0.851 | 0.922 | (A) BRAF Class 1 Melanoma |
| CDC42 | 1p36.12 | 1 | 0.48 | 0 | 0.00 | 0.852 | 0.922 | (A) BRAF Class 1 Melanoma |
| KMT5A | 12q24.31 | 1 | 0.58 | 0 | 0.00 | 0.852 | 0.922 | (A) BRAF Class 1 Melanoma |

| | | | | | | | | |
|---|---|---|---|---|---|---|---|---|
| PRKD1 | 14q12 | 1 | 0.53 | 0 | 0.00 | 0.862 | 0.922 | (A) BRAF Class 1 Melanoma |
| ABCA17P | 16p13.3 | 1 | 0.08 | 0 | 0.00 | 0.871 | 0.922 | (A) BRAF Class 1 Melanoma |
| ABR | 17p13.3 | 1 | 0.08 | 0 | 0.00 | 0.871 | 0.922 | (A) BRAF Class 1 Melanoma |
| ARHGAP4 | Xq28 | 1 | 0.08 | 0 | 0.00 | 0.871 | 0.922 | (A) BRAF Class 1 Melanoma |
| ARSB | 5q14.1 | 1 | 0.08 | 0 | 0.00 | 0.871 | 0.922 | (A) BRAF Class 1 Melanoma |
| CAMK1 | 3p25.3 | 1 | 0.08 | 0 | 0.00 | 0.871 | 0.922 | (A) BRAF Class 1 Melanoma |
| CASZ1 | 1p36.22 | 1 | 0.08 | 0 | 0.00 | 0.871 | 0.922 | (A) BRAF Class 1 Melanoma |
| CTDSP1 | 2q35 | 1 | 0.08 | 0 | 0.00 | 0.871 | 0.922 | (A) BRAF Class 1 Melanoma |
| DDAH1 | 1p22.3 | 1 | 0.08 | 0 | 0.00 | 0.871 | 0.922 | (A) BRAF Class 1 Melanoma |
| DEF8 | 16q24.3 | 1 | 0.08 | 0 | 0.00 | 0.871 | 0.922 | (A) BRAF Class 1 Melanoma |
| DENND4C | 9p22.1 | 1 | 0.08 | 0 | 0.00 | 0.871 | 0.922 | (A) BRAF Class 1 Melanoma |
| DHRSX | .33 and Yp | 1 | 0.08 | 0 | 0.00 | 0.871 | 0.922 | (A) BRAF Class 1 Melanoma |
| GALNT11 | 36.1\|7q36 | 1 | 0.08 | 0 | 0.00 | 0.871 | 0.922 | (A) BRAF Class 1 Melanoma |
| GRIA1 | 5q33.2 | 1 | 0.08 | 0 | 0.00 | 0.871 | 0.922 | (A) BRAF Class 1 Melanoma |
| GRIK1 | 21q21.3 | 1 | 0.08 | 0 | 0.00 | 0.871 | 0.922 | (A) BRAF Class 1 Melanoma |
| HECTD4 | 12q24.13 | 1 | 0.08 | 0 | 0.00 | 0.871 | 0.922 | (A) BRAF Class 1 Melanoma |
| HSD17B6 | 12q13.3 | 1 | 0.08 | 0 | 0.00 | 0.871 | 0.922 | (A) BRAF Class 1 Melanoma |
| C1053744 | 2p22.2 | 1 | 0.08 | 0 | 0.00 | 0.871 | 0.922 | (A) BRAF Class 1 Melanoma |
| MYO1D | 17q11.2 | 1 | 0.08 | 0 | 0.00 | 0.871 | 0.922 | (A) BRAF Class 1 Melanoma |
| MYSM1 | 1p32.1 | 1 | 0.08 | 0 | 0.00 | 0.871 | 0.922 | (A) BRAF Class 1 Melanoma |
| PADI2 | 1p36.13 | 1 | 0.08 | 0 | 0.00 | 0.871 | 0.922 | (A) BRAF Class 1 Melanoma |
| PAX9 | 14q13.3 | 1 | 0.08 | 0 | 0.00 | 0.871 | 0.922 | (A) BRAF Class 1 Melanoma |
| PFKFB4 | 3p21.31 | 1 | 0.08 | 0 | 0.00 | 0.871 | 0.922 | (A) BRAF Class 1 Melanoma |
| PLVAP | 19p13.11 | 1 | 0.08 | 0 | 0.00 | 0.871 | 0.922 | (A) BRAF Class 1 Melanoma |
| PNPLA6 | 19p13.2 | 1 | 0.08 | 0 | 0.00 | 0.871 | 0.922 | (A) BRAF Class 1 Melanoma |
| PRMT6 | 1p13.3 | 1 | 0.08 | 0 | 0.00 | 0.871 | 0.922 | (A) BRAF Class 1 Melanoma |
| RAB2A | q12.1-q12. | 1 | 0.08 | 0 | 0.00 | 0.871 | 0.922 | (A) BRAF Class 1 Melanoma |
| SAE1 | 19q13.32 | 1 | 0.08 | 0 | 0.00 | 0.871 | 0.922 | (A) BRAF Class 1 Melanoma |
| SCARA5 | 8p21.1 | 1 | 0.08 | 0 | 0.00 | 0.871 | 0.922 | (A) BRAF Class 1 Melanoma |
| SDCCAG8 | 1q43-q44 | 1 | 0.08 | 0 | 0.00 | 0.871 | 0.922 | (A) BRAF Class 1 Melanoma |
| SEMA6D | 15q21.1 | 1 | 0.08 | 0 | 0.00 | 0.871 | 0.922 | (A) BRAF Class 1 Melanoma |
| SIK3 | 11q23.3 | 1 | 0.08 | 0 | 0.00 | 0.871 | 0.922 | (A) BRAF Class 1 Melanoma |
| TRAJ42 | 14q11.2 | 1 | 0.08 | 0 | 0.00 | 0.871 | 0.922 | (A) BRAF Class 1 Melanoma |
| UGT1A6 | 2q37.1 | 1 | 0.08 | 0 | 0.00 | 0.871 | 0.922 | (A) BRAF Class 1 Melanoma |
| ZNF423 | 16q12.1 | 1 | 0.08 | 0 | 0.00 | 0.871 | 0.922 | (A) BRAF Class 1 Melanoma |
| RPS6KB1 | 17q23.1 | 2 | 1.26 | 0 | 0.00 | 0.874 | 0.922 | (A) BRAF Class 1 Melanoma |
| HIST2H3D | 1q21.2 | 1 | 0.26 | 0 | 0.00 | 0.876 | 0.922 | (A) BRAF Class 1 Melanoma |
| TYRO3 | 15q15.1 | 1 | 1.28 | 0 | 0.00 | 0.876 | 0.922 | (A) BRAF Class 1 Melanoma |
| HIST2H3C | 1q21.2 | 1 | 0.26 | 0 | 0.00 | 0.877 | 0.922 | (A) BRAF Class 1 Melanoma |
| PMAIP1 | 18q21.32 | 1 | 0.25 | 0 | 0.00 | 0.877 | 0.922 | (A) BRAF Class 1 Melanoma |
| DDR1 | 6p21.33 | 1 | 1.00 | 0 | 0.00 | 0.877 | 0.922 | (A) BRAF Class 1 Melanoma |
| PDPK1 | 16p13.3 | 1 | 0.23 | 0 | 0.00 | 0.877 | 0.922 | (A) BRAF Class 1 Melanoma |
| PHLPP2 | 16q22.2 | 1 | 1.35 | 0 | 0.00 | 0.881 | 0.922 | (A) BRAF Class 1 Melanoma |
| UGT1A1 | 2q37.1 | 1 | 0.95 | 0 | 0.00 | 0.882 | 0.922 | (A) BRAF Class 1 Melanoma |
| CDK1 | 10q21.2 | 1 | 1.47 | 0 | 0.00 | 0.883 | 0.922 | (A) BRAF Class 1 Melanoma |

| | | | | | | | | |
|---|---|---|---|---|---|---|---|---|
| CDK9 | 9q34.11 | 1 | 1.47 | 0 | 0.00 | 0.883 | 0.922 | (A) BRAF Class 1 Melanoma |
| CRTC2 | 1q21.3 | 1 | 1.47 | 0 | 0.00 | 0.883 | 0.922 | (A) BRAF Class 1 Melanoma |
| PRAME | 22q11.22 | 1 | 1.47 | 0 | 0.00 | 0.883 | 0.922 | (A) BRAF Class 1 Melanoma |
| PSMD13 | 11p15.5 | 1 | 1.47 | 0 | 0.00 | 0.883 | 0.922 | (A) BRAF Class 1 Melanoma |
| PTK2 | 8q24.3 | 1 | 1.47 | 0 | 0.00 | 0.883 | 0.922 | (A) BRAF Class 1 Melanoma |
| RHPN2 | 19q13.11 | 1 | 1.47 | 0 | 0.00 | 0.883 | 0.922 | (A) BRAF Class 1 Melanoma |
| SRP19 | 5q22.2 | 1 | 0.69 | 0 | 0.00 | 0.883 | 0.922 | (A) BRAF Class 1 Melanoma |
| FGF5 | 4q21.21 | 2 | 1.26 | 0 | 0.00 | 0.885 | 0.922 | (A) BRAF Class 1 Melanoma |
| ZBTB7A | 19p13.3 | 2 | 1.26 | 0 | 0.00 | 0.885 | 0.922 | (A) BRAF Class 1 Melanoma |
| ENG | 9q34.11 | 1 | 0.50 | 0 | 0.00 | 0.885 | 0.922 | (A) BRAF Class 1 Melanoma |
| FAAP24 | 19q13.11 | 1 | 0.50 | 0 | 0.00 | 0.885 | 0.922 | (A) BRAF Class 1 Melanoma |
| KLF2 | 19p13.11 | 1 | 0.50 | 0 | 0.00 | 0.885 | 0.922 | (A) BRAF Class 1 Melanoma |
| UBE2T | 1q32.1 | 1 | 0.50 | 0 | 0.00 | 0.885 | 0.922 | (A) BRAF Class 1 Melanoma |
| FANCF | 11p14.3 | 1 | 0.19 | 0 | 0.00 | 0.891 | 0.927 | (A) BRAF Class 1 Melanoma |
| PHF6 | Xq26.2 | 1 | 0.21 | 0 | 0.00 | 0.897 | 0.93 | (A) BRAF Class 1 Melanoma |
| ATP1B2 | 17p13.1 | 0 | 0.00 | 1 | 0.57 | 0.897 | 0.93 | (B) BRAF Class 2 Melanoma |
| CDH12 | 5p14.3 | 0 | 0.00 | 1 | 0.57 | 0.897 | 0.93 | (B) BRAF Class 2 Melanoma |
| KIF5B | 10p11.22 | 2 | 1.09 | 0 | 0.00 | 0.899 | 0.93 | (A) BRAF Class 1 Melanoma |
| MAP3K2 | 2q14.3 | 1 | 1.85 | 0 | 0.00 | 0.9 | 0.93 | (A) BRAF Class 1 Melanoma |
| MGMT | 10q26.3 | 1 | 1.85 | 0 | 0.00 | 0.9 | 0.93 | (A) BRAF Class 1 Melanoma |
| CDH20 | 18q21.33 | 1 | 1.09 | 0 | 0.00 | 0.902 | 0.931 | (A) BRAF Class 1 Melanoma |
| BTG1 | 12q21.33 | 1 | 0.36 | 0 | 0.00 | 0.906 | 0.934 | (A) BRAF Class 1 Melanoma |
| FRS2 | 12q15 | 1 | 0.38 | 0 | 0.00 | 0.907 | 0.934 | (A) BRAF Class 1 Melanoma |
| HDAC4 | 2q37.3 | 1 | 1.41 | 0 | 0.00 | 0.91 | 0.935 | (A) BRAF Class 1 Melanoma |
| GNA13 | 17q24.1 | 1 | 0.34 | 0 | 0.00 | 0.913 | 0.935 | (A) BRAF Class 1 Melanoma |
| SAMD9 | 7q21.2 | 2 | 3.08 | 0 | 0.00 | 0.913 | 0.935 | (A) BRAF Class 1 Melanoma |
| APOBEC3G | 22q13.1 | 1 | 3.13 | 0 | 0.00 | 0.914 | 0.935 | (A) BRAF Class 1 Melanoma |
| HSPA5 | 9q33.3 | 1 | 3.13 | 0 | 0.00 | 0.914 | 0.935 | (A) BRAF Class 1 Melanoma |
| USP7 | 16p13.2 | 1 | 3.13 | 0 | 0.00 | 0.914 | 0.935 | (A) BRAF Class 1 Melanoma |
| MAP3K7 | 6q15 | 1 | 1.27 | 0 | 0.00 | 0.919 | 0.938 | (A) BRAF Class 1 Melanoma |
| IRF2 | 4q35.1 | 1 | 0.42 | 0 | 0.00 | 0.919 | 0.938 | (A) BRAF Class 1 Melanoma |
| RACK1 | 5q35.3 | 1 | 1.67 | 0 | 0.00 | 0.923 | 0.94 | (A) BRAF Class 1 Melanoma |
| RPL10 | Xq28 | 1 | 2.70 | 0 | 0.00 | 0.925 | 0.94 | (A) BRAF Class 1 Melanoma |
| HSP90AB1 | 6p21.1 | 1 | 1.27 | 0 | 0.00 | 0.929 | 0.94 | (A) BRAF Class 1 Melanoma |
| DDX31 | 9q34.13 | 1 | 2.50 | 0 | 0.00 | 0.93 | 0.94 | (A) BRAF Class 1 Melanoma |
| FOSB | 19q13.32 | 1 | 2.50 | 0 | 0.00 | 0.93 | 0.94 | (A) BRAF Class 1 Melanoma |
| OGA | 10q24.32 | 1 | 2.50 | 0 | 0.00 | 0.93 | 0.94 | (A) BRAF Class 1 Melanoma |
| OTX2 | 14q22.3 | 1 | 2.50 | 0 | 0.00 | 0.93 | 0.94 | (A) BRAF Class 1 Melanoma |
| POLR2A | 17p13.1 | 1 | 2.50 | 0 | 0.00 | 0.93 | 0.94 | (A) BRAF Class 1 Melanoma |
| TBXT | 6q27 | 1 | 2.50 | 0 | 0.00 | 0.93 | 0.94 | (A) BRAF Class 1 Melanoma |
| PLAG1 | 8q12.1 | 1 | 1.16 | 0 | 0.00 | 0.935 | 0.943 | (A) BRAF Class 1 Melanoma |
| CSNK1A1 | 5q32 | 1 | 0.63 | 0 | 0.00 | 0.941 | 0.947 | (A) BRAF Class 1 Melanoma |
| NAB2 | 12q13.3 | 1 | 0.63 | 0 | 0.00 | 0.941 | 0.947 | (A) BRAF Class 1 Melanoma |
| LAMP1 | 13q34 | 1 | 0.54 | 0 | 0.00 | 0.948 | 0.954 | (A) BRAF Class 1 Melanoma |
| ETNK1 | 12p12.1 | 1 | 1.75 | 0 | 0.00 | 0.95 | 0.954 | (A) BRAF Class 1 Melanoma |

| | | | | | | | | |
|---|---|---|---|---|---|---|---|---|
| CD22 | 19q13.12 | 1 | 2.17 | 0 | 0.00 | 0.958 | 0.962 | (A) BRAF Class 1 Melanoma |
| ECT2L | 6q24.1 | 1 | 2.38 | 0 | 0.00 | 0.977 | 0.978 | (A) BRAF Class 1 Melanoma |
| TBL1XR1 | 3q26.32 | 1 | 2.38 | 0 | 0.00 | 0.977 | 0.978 | (A) BRAF Class 1 Melanoma |
| BRAF | 7q34 | 1180 | 100.00 | 175 | 100.00 | 1 | 1 | (B) BRAF Class 2 Melanoma |

#### Supplementary Table 8

| Gene | Cytoband | Class 1 Occurrence | Class 1 % | Class 2 occurrence | Class 2 % | SumClass | pvalue | qvalue | Enriched In |
|---|---|---|---|---|---|---|---|---|---|
| SYK | 9q22.2 | 16 | 2.72 | 9 | 10.59 | 25 | 0.002 | 0.0423 | (B) BRAF Class 2 Melanoma |
| CSF3R | 1p34.3 | 13 | 1.85 | 8 | 8.33 | 21 | 0.00177 | 0.039 | (B) BRAF Class 2 Melanoma |
| PPARG | 3p25.2 | 17 | 2.64 | 9 | 10.71 | 26 | 0.00144 | 0.036 | (B) BRAF Class 2 Melanoma |
| NOTCH4 | 6p21.32 | 32 | 6.54 | 13 | 18.84 | 45 | 0.00152 | 0.036 | (B) BRAF Class 2 Melanoma |
| PAK5 | 20p12.2 | 62 | 13.45 | 20 | 28.99 | 82 | 0.00156 | 0.036 | (B) BRAF Class 2 Melanoma |
| ATM | 11q22.3 | 66 | 5.97 | 20 | 13.61 | 86 | 0.00133 | 0.036 | (B) BRAF Class 2 Melanoma |
| TET2 | 4q24 | 70 | 9.00 | 22 | 19.30 | 92 | 0.00138 | 0.036 | (B) BRAF Class 2 Melanoma |
| TP53 | 17p13.1 | 175 | 14.83 | 39 | 24.84 | 214 | 0.00154 | 0.036 | (B) BRAF Class 2 Melanoma |
| JAK1 | 1p31.3 | 18 | 2.37 | 11 | 8.87 | 29 | 0.00101 | 0.03 | (B) BRAF Class 2 Melanoma |
| KDR | 4q12 | 87 | 7.87 | 25 | 17.01 | 112 | 0.00061 | 0.022 | (B) BRAF Class 2 Melanoma |
| SLIT2 | 4p15.31 | 28 | 10.61 | 10 | 38.46 | 38 | 0.00054 | 0.0201 | (B) BRAF Class 2 Melanoma |
| SF3B1 | 2q33.1 | 22 | 2.65 | 12 | 9.92 | 34 | 0.0005 | 0.0196 | (B) BRAF Class 2 Melanoma |
| FOXP1 | 3p13 | 9 | 1.74 | 8 | 10.81 | 17 | 0.0004 | 0.0185 | (B) BRAF Class 2 Melanoma |
| COL7A1 | 3p21.31 | 46 | 23.00 | 15 | 57.69 | 61 | 0.00043 | 0.0185 | (B) BRAF Class 2 Melanoma |
| FLT1 | 13q12.3 | 69 | 8.96 | 23 | 20.72 | 92 | 0.00041 | 0.0185 | (B) BRAF Class 2 Melanoma |
| ERBB4 | 2q34 | 107 | 9.54 | 32 | 18.93 | 139 | 0.00045 | 0.0185 | (B) BRAF Class 2 Melanoma |
| GRIN2A | 16p13.2 | 92 | 17.49 | 27 | 36.00 | 119 | 0.00033 | 0.0164 | (B) BRAF Class 2 Melanoma |
| MAGI2 | 7q21.11 | 18 | 7.89 | 8 | 40.00 | 26 | 0.00027 | 0.0158 | (B) BRAF Class 2 Melanoma |
| MET | 7q31.2 | 68 | 5.80 | 24 | 13.87 | 92 | 0.00028 | 0.0158 | (B) BRAF Class 2 Melanoma |
| BLM | 15q26.1 | 23 | 3.03 | 13 | 11.82 | 36 | 0.0002 | 0.0128 | (B) BRAF Class 2 Melanoma |
| RPTOR | 17q25.3 | 34 | 4.73 | 16 | 15.69 | 50 | 0.00014 | 0.0096 | (B) BRAF Class 2 Melanoma |

| | | | | | | | | | |
|---|---|---|---|---|---|---|---|---|---|
| EPHA7 | 6q16.1 | 57 | 10.27 | 23 | 28.40 | 80 | 3E-05 | 0.00267 | (B) BRAF Class 2 Melanoma |
| MPL | 1p34.2 | 19 | 1.77 | 13 | 9.03 | 32 | 2.5E-05 | 0.00247 | (B) BRAF Class 2 Melanoma |
| ATR | 3q23 | 27 | 3.58 | 16 | 15.24 | 43 | 1.3E-05 | 0.00146 | (B) BRAF Class 2 Melanoma |
| KMT2A | 11q23.3 | 67 | 8.61 | 27 | 23.89 | 94 | 7.8E-06 | 0.001 | (B) BRAF Class 2 Melanoma |
| KMT2D | 12q13.12 | 118 | 15.32 | 38 | 34.23 | 156 | 4.8E-06 | 0.00072 | (B) BRAF Class 2 Melanoma |
| CARD11 | 7p22.2 | 61 | 7.92 | 27 | 24.32 | 88 | 1.5E-06 | 0.00028 | (B) BRAF Class 2 Melanoma |
| NRAS | 1p13.2 | 10 | 0.85 | 14 | 8.00 | 24 | 1.4E-07 | 3E-05 | (B) BRAF Class 2 Melanoma |
| HNF1A | 12q24.31 | 18 | 1.63 | 17 | 11.64 | 35 | 4.3E-08 | 1.3E-05 | (B) BRAF Class 2 Melanoma |
| FLT3 | 13q12.2 | 41 | 3.70 | 28 | 19.05 | 69 | 2E-10 | 9.1E-08 | (B) BRAF Class 2 Melanoma |
| NF1 | 17q11.2 | 50 | 6.19 | 36 | 27.07 | 86 | 2.1E-11 | 1.9E-08 | (B) BRAF Class 2 Melanoma |

#### Supplementary Table 9

| Gene | Cytoband | Class 1 Occurrence | Class 1 % | Class 3 occurrence | Class 3 % | pvalue | qvalue | Enriched In |
|---|---|---|---|---|---|---|---|---|
| NF1 | 17q11.2 | 52 | 6.41 | 79 | 64.75 | 8.93E-48 | 8.13E-45 | (B) BRAF Class 3 Melanoma |
| NRAS | 1p13.2 | 10 | 0.85 | 54 | 31.58 | 4.27E-42 | 1.94E-39 | (B) BRAF Class 3 Melanoma |
| KRAS | 12p12.1 | 5 | 0.42 | 17 | 9.94 | 3.81E-12 | 1.16E-09 | (B) BRAF Class 3 Melanoma |
| FLT3 | 13q12.2 | 41 | 3.68 | 31 | 19.62 | 1.52E-11 | 3.46E-09 | (B) BRAF Class 3 Melanoma |
| KMT2A | 11q23.3 | 69 | 8.83 | 36 | 30.51 | 1.5E-09 | 2.74E-07 | (B) BRAF Class 3 Melanoma |
| NOTCH2 | 1p12 | 44 | 5.54 | 28 | 24.35 | 2.55E-09 | 3.73E-07 | (B) BRAF Class 3 Melanoma |
| SETD2 | 3p21.31 | 60 | 7.43 | 33 | 27.27 | 2.86E-09 | 3.73E-07 | (B) BRAF Class 3 Melanoma |
| ARID2 | 12q12 | 66 | 8.66 | 34 | 29.31 | 7.44E-09 | 8.47E-07 | (B) BRAF Class 3 Melanoma |
| FLT1 | 13q12.3 | 71 | 9.18 | 35 | 29.91 | 9.19E-09 | 8.81E-07 | (B) BRAF Class 3 Melanoma |
| KMT2D | 12q13.12 | 119 | 15.39 | 46 | 39.32 | 9.67E-09 | 8.81E-07 | (B) BRAF Class 3 Melanoma |
| ATM | 11q22.3 | 70 | 6.31 | 34 | 21.52 | 1.1E-08 | 9.08E-07 | (B) BRAF Class 3 Melanoma |
| KDR | 4q12 | 87 | 7.84 | 38 | 24.05 | 1.26E-08 | 9.56E-07 | (B) BRAF Class 3 Melanoma |
| FGFR4 | 5q35.2 | 26 | 3.19 | 21 | 17.07 | 3.92E-08 | 2.75E-06 | (B) BRAF Class 3 Melanoma |
| PTPRD | 9p24.1-p23 | 88 | 15.71 | 36 | 43.37 | 4.36E-08 | 2.84E-06 | (B) BRAF Class 3 Melanoma |
| ERBB4 | 2q34 | 109 | 9.69 | 42 | 25.77 | 6.04E-08 | 3.67E-06 | (B) BRAF Class 3 Melanoma |
| TET2 | 4q24 | 70 | 8.96 | 33 | 27.97 | 6.66E-08 | 3.79E-06 | (B) BRAF Class 3 Melanoma |
| HRAS | 11p15.5 | 4 | 0.34 | 11 | 6.43 | 8.47E-08 | 4.54E-06 | (B) BRAF Class 3 Melanoma |
| KIT | 4q12 | 24 | 2.03 | 19 | 11.11 | 1.72E-07 | 8.7E-06 | (B) BRAF Class 3 Melanoma |
| AXL | 19q13.2 | 30 | 3.71 | 21 | 17.36 | 2.01E-07 | 9.63E-06 | (B) BRAF Class 3 Melanoma |
| NTRK3 | 15q25.3 | 52 | 6.44 | 27 | 22.31 | 2.87E-07 | 1.31E-05 | (B) BRAF Class 3 Melanoma |
| RUNX1T1 | 8q21.3 | 35 | 7.58 | 21 | 31.34 | 3.29E-07 | 1.43E-05 | (B) BRAF Class 3 Melanoma |
| ROS1 | 6q22.1 | 150 | 17.38 | 49 | 37.69 | 3.66E-07 | 1.52E-05 | (B) BRAF Class 3 Melanoma |
| MET | 7q31.2 | 70 | 5.96 | 31 | 18.24 | 4.56E-07 | 1.79E-05 | (B) BRAF Class 3 Melanoma |
| CARD11 | 7p22.2 | 61 | 7.89 | 29 | 24.79 | 4.72E-07 | 1.79E-05 | (B) BRAF Class 3 Melanoma |
| BLM | 15q26.1 | 23 | 3.02 | 18 | 15.52 | 6.37E-07 | 2.32E-05 | (B) BRAF Class 3 Melanoma |
| IKZF1 | 7p12.2 | 62 | 8.07 | 29 | 24.79 | 7.02E-07 | 2.46E-05 | (B) BRAF Class 3 Melanoma |
| MECOM | 3q26.2 | 39 | 14.66 | 21 | 50.00 | 1.19E-06 | 4.02E-05 | (B) BRAF Class 3 Melanoma |
| SMARCA4 | 19p13.2 | 50 | 6.19 | 25 | 20.66 | 1.52E-06 | 4.9E-05 | (B) BRAF Class 3 Melanoma |
| FGFR2 | 10q26.13 | 47 | 3.97 | 24 | 14.04 | 1.56E-06 | 4.9E-05 | (B) BRAF Class 3 Melanoma |
| IL7R | 5p13.2 | 49 | 6.89 | 24 | 23.08 | 1.91E-06 | 5.81E-05 | (B) BRAF Class 3 Melanoma |
| NOTCH1 | 9q34.3 | 49 | 4.20 | 24 | 14.46 | 2.07E-06 | 6.07E-05 | (B) BRAF Class 3 Melanoma |
| CBL | 11q23.3 | 30 | 3.68 | 19 | 15.57 | 2.51E-06 | 6.97E-05 | (B) BRAF Class 3 Melanoma |
| GRIN2A | 16p13.2 | 92 | 17.46 | 32 | 42.67 | 2.52E-06 | 6.97E-05 | (B) BRAF Class 3 Melanoma |
| PREX2 | 8q13.2 | 75 | 17.69 | 27 | 45.76 | 4.48E-06 | 0.00012 | (B) BRAF Class 3 Melanoma |
| ARID1A | 1p36.11 | 63 | 7.80 | 27 | 22.31 | 5.67E-06 | 0.000148 | (B) BRAF Class 3 Melanoma |
| RET | 10q11.21 | 44 | 3.72 | 22 | 12.87 | 5.86E-06 | 0.000148 | (B) BRAF Class 3 Melanoma |
| ABCB11 | 2q31.1 | 11 | 4.93 | 11 | 34.38 | 6.08E-06 | 0.00015 | (B) BRAF Class 3 Melanoma |
| DDR2 | 1q23.3 | 19 | 2.16 | 15 | 11.19 | 6.41E-06 | 0.000154 | (B) BRAF Class 3 Melanoma |
| GLI1 | 12q13.3 | 31 | 4.23 | 18 | 16.51 | 1.04E-05 | 0.000243 | (B) BRAF Class 3 Melanoma |
| ALK | p23.2-p23. | 71 | 6.00 | 28 | 16.37 | 1.1E-05 | 0.000249 | (B) BRAF Class 3 Melanoma |
| TP53 | 17p13.1 | 175 | 14.79 | 49 | 28.82 | 1.17E-05 | 0.000261 | (B) BRAF Class 3 Melanoma |
| CSF3R | 1p34.3 | 13 | 1.85 | 12 | 11.65 | 1.27E-05 | 0.000274 | (B) BRAF Class 3 Melanoma |
| ASXL1 | 20q11.21 | 31 | 3.97 | 18 | 15.25 | 1.32E-05 | 0.000279 | (B) BRAF Class 3 Melanoma |
| EGFR | 7p11.2 | 51 | 4.31 | 23 | 13.45 | 1.35E-05 | 0.000279 | (B) BRAF Class 3 Melanoma |
| EPHA3 | 3p11.1 | 45 | 7.63 | 21 | 23.86 | 1.9E-05 | 0.000384 | (B) BRAF Class 3 Melanoma |

| | | | | | | | | |
|---|---|---|---|---|---|---|---|---|
| SYK | 9q22.2 | 17 | 2.88 | 13 | 14.77 | 2.37E-05 | 0.000469 | (B) BRAF Class 3 Melanoma |
| SETBP1 | 18q12.3 | 40 | 8.70 | 19 | 27.94 | 2.79E-05 | 0.000542 | (B) BRAF Class 3 Melanoma |
| PTPRT | 0q12-q13.1 | 90 | 19.19 | 28 | 43.75 | 2.93E-05 | 0.000555 | (B) BRAF Class 3 Melanoma |
| TET1 | 10q21.3 | 31 | 5.02 | 16 | 19.05 | 3.14E-05 | 0.000577 | (B) BRAF Class 3 Melanoma |
| PTPN11 | 12q24.13 | 12 | 1.07 | 11 | 6.79 | 3.17E-05 | 0.000577 | (B) BRAF Class 3 Melanoma |
| EPHA7 | 6q16.1 | 58 | 10.41 | 23 | 28.40 | 3.53E-05 | 0.000631 | (B) BRAF Class 3 Melanoma |
| COL7A1 | 3p21.31 | 46 | 22.89 | 18 | 62.07 | 3.67E-05 | 0.000643 | (B) BRAF Class 3 Melanoma |
| SYNE1 | 6q25.2 | 5 | 2.50 | 7 | 29.17 | 3.76E-05 | 0.000645 | (B) BRAF Class 3 Melanoma |
| EP300 | 22q13.2 | 42 | 5.43 | 20 | 17.09 | 4.01E-05 | 0.000676 | (B) BRAF Class 3 Melanoma |
| APC | 5q22.2 | 74 | 6.86 | 27 | 17.42 | 4.23E-05 | 0.000701 | (B) BRAF Class 3 Melanoma |
| DICER1 | 14q32.13 | 25 | 3.30 | 15 | 13.27 | 4.96E-05 | 0.000799 | (B) BRAF Class 3 Melanoma |
| JAK3 | 19p13.11 | 37 | 3.32 | 18 | 11.32 | 5E-05 | 0.000799 | (B) BRAF Class 3 Melanoma |
| PDGFRB | 5q32 | 32 | 3.96 | 17 | 14.05 | 5.21E-05 | 0.000818 | (B) BRAF Class 3 Melanoma |
| NOTCH3 | 19p13.12 | 87 | 11.65 | 29 | 26.85 | 5.86E-05 | 0.000905 | (B) BRAF Class 3 Melanoma |
| RPTOR | 17q25.3 | 34 | 4.72 | 17 | 16.19 | 6.26E-05 | 0.00095 | (B) BRAF Class 3 Melanoma |
| SOX17 | 8q11.23 | 5 | 1.23 | 7 | 13.73 | 7.22E-05 | 0.001078 | (B) BRAF Class 3 Melanoma |
| STAT5B | 17q21.2 | 3 | 0.72 | 6 | 11.76 | 8.24E-05 | 0.001196 | (B) BRAF Class 3 Melanoma |
| BARD1 | 2q35 | 6 | 0.83 | 8 | 7.62 | 8.27E-05 | 0.001196 | (B) BRAF Class 3 Melanoma |
| CIC | 19q13.2 | 40 | 5.54 | 18 | 17.14 | 0.000108 | 0.001531 | (B) BRAF Class 3 Melanoma |
| BRD4 | 19p13.12 | 39 | 5.13 | 18 | 15.93 | 0.000113 | 0.001584 | (B) BRAF Class 3 Melanoma |
| ATRX | Xq21.1 | 47 | 5.82 | 20 | 16.53 | 0.000127 | 0.001748 | (B) BRAF Class 3 Melanoma |
| DAXX | 6p21.32 | 16 | 2.26 | 11 | 11.11 | 0.000134 | 0.001802 | (B) BRAF Class 3 Melanoma |
| PBRM1 | 3p21.1 | 24 | 3.14 | 14 | 12.07 | 0.000137 | 0.001802 | (B) BRAF Class 3 Melanoma |
| NOTCH4 | 6p21.32 | 33 | 6.73 | 15 | 22.73 | 0.000137 | 0.001802 | (B) BRAF Class 3 Melanoma |
| PTK2B | 8p21.2 | 9 | 4.19 | 9 | 25.71 | 0.000149 | 0.001941 | (B) BRAF Class 3 Melanoma |
| PAK5 | 20p12.2 | 63 | 13.64 | 21 | 33.87 | 0.000164 | 0.002099 | (B) BRAF Class 3 Melanoma |
| BRCA2 | 13q13.1 | 57 | 7.06 | 22 | 18.18 | 0.000181 | 0.002287 | (B) BRAF Class 3 Melanoma |
| GNAS | 20q13.32 | 13 | 1.15 | 10 | 6.17 | 0.000198 | 0.002446 | (B) BRAF Class 3 Melanoma |
| CDH4 | 20q13.33 | 15 | 7.46 | 10 | 34.48 | 0.000199 | 0.002446 | (B) BRAF Class 3 Melanoma |
| FANCE | 6p21.31 | 7 | 1.35 | 8 | 10.13 | 0.000201 | 0.002446 | (B) BRAF Class 3 Melanoma |
| MAP3K5 | 6q23.3 | 0 | 0.00 | 4 | 44.44 | 0.000212 | 0.002536 | (B) BRAF Class 3 Melanoma |
| PRKDC | 8q11.21 | 54 | 10.53 | 21 | 26.58 | 0.000225 | 0.002649 | (B) BRAF Class 3 Melanoma |
| MTOR | 1p36.22 | 41 | 4.73 | 18 | 13.85 | 0.000227 | 0.002649 | (B) BRAF Class 3 Melanoma |
| FANCM | 14q21.2 | 13 | 4.78 | 10 | 23.26 | 0.000245 | 0.002791 | (B) BRAF Class 3 Melanoma |
| TSC2 | 16p13.3 | 30 | 3.70 | 15 | 12.40 | 0.000245 | 0.002791 | (B) BRAF Class 3 Melanoma |
| CCND2 | 12p13.32 | 10 | 1.25 | 9 | 7.44 | 0.00025 | 0.002813 | (B) BRAF Class 3 Melanoma |
| PTCH1 | 9q22.32 | 27 | 3.34 | 14 | 11.57 | 0.000312 | 0.00347 | (B) BRAF Class 3 Melanoma |
| HNF1A | 12q24.31 | 18 | 1.63 | 11 | 7.10 | 0.00033 | 0.003619 | (B) BRAF Class 3 Melanoma |
| MAP3K1 | 5q11.2 | 27 | 3.50 | 14 | 11.97 | 0.000345 | 0.003679 | (B) BRAF Class 3 Melanoma |
| GATA3 | 10p14 | 20 | 2.59 | 12 | 10.26 | 0.000347 | 0.003679 | (B) BRAF Class 3 Melanoma |
| IGF1 | 12q23.2 | 2 | 0.43 | 5 | 8.06 | 0.000347 | 0.003679 | (B) BRAF Class 3 Melanoma |
| FOXO1 | 13q14.11 | 7 | 1.52 | 7 | 11.48 | 0.000379 | 0.003917 | (B) BRAF Class 3 Melanoma |
| STAT4 | q32.2-q32. | 10 | 3.60 | 8 | 21.05 | 0.000381 | 0.003917 | (B) BRAF Class 3 Melanoma |
| BMPR1A | 10q23.2 | 6 | 0.81 | 7 | 6.54 | 0.000383 | 0.003917 | (B) BRAF Class 3 Melanoma |
| TP53BP1 | 15q15.3 | 18 | 4.41 | 11 | 18.03 | 0.000396 | 0.004003 | (B) BRAF Class 3 Melanoma |
| PDGFRA | 4q12 | 50 | 4.32 | 19 | 11.52 | 0.000409 | 0.004092 | (B) BRAF Class 3 Melanoma |
| GATA6 | 18q11.2 | 4 | 0.85 | 6 | 8.96 | 0.000415 | 0.004105 | (B) BRAF Class 3 Melanoma |
| KDM5C | Xp11.22 | 24 | 3.14 | 13 | 11.21 | 0.00042 | 0.004118 | (B) BRAF Class 3 Melanoma |

| | | | | | | | | |
|---|---|---|---|---|---|---|---|---|
| POLE | 12q24.33 | 60 | 7.91 | 21 | 19.27 | 0.000425 | 0.004121 | (B) BRAF Class 3 Melanoma |
| SLIT2 | 4p15.31 | 28 | 10.61 | 12 | 35.29 | 0.000439 | 0.004206 | (B) BRAF Class 3 Melanoma |
| MGA | 15q15.1 | 68 | 11.11 | 22 | 25.58 | 0.000461 | 0.004373 | (B) BRAF Class 3 Melanoma |
| TMPRSS2 | 21q22.3 | 15 | 2.09 | 10 | 9.52 | 0.000475 | 0.004461 | (B) BRAF Class 3 Melanoma |
| FANCD2 | 3p25.3 | 23 | 4.45 | 12 | 16.22 | 0.000482 | 0.004479 | (B) BRAF Class 3 Melanoma |
| TERT | 5p15.33 | 460 | 60.85 | 87 | 76.99 | 0.000488 | 0.004489 | (B) BRAF Class 3 Melanoma |
| KDM5A | 12p13.33 | 22 | 3.07 | 12 | 11.43 | 0.000498 | 0.004534 | (B) BRAF Class 3 Melanoma |
| CIITA | 16p13.13 | 18 | 6.52 | 11 | 25.00 | 0.000528 | 0.004728 | (B) BRAF Class 3 Melanoma |
| MPL | 1p34.2 | 19 | 1.76 | 11 | 7.14 | 0.000535 | 0.004728 | (B) BRAF Class 3 Melanoma |
| FBXW7 | 4q31.3 | 30 | 2.66 | 14 | 8.64 | 0.000537 | 0.004728 | (B) BRAF Class 3 Melanoma |
| CUX1 | 7q22.1 | 34 | 7.62 | 15 | 22.39 | 0.00054 | 0.004728 | (B) BRAF Class 3 Melanoma |
| PIK3C2B | 1q32.1 | 24 | 4.99 | 12 | 17.65 | 0.000562 | 0.004879 | (B) BRAF Class 3 Melanoma |
| PIK3CA | 3q26.32 | 34 | 2.87 | 15 | 8.77 | 0.000583 | 0.005011 | (B) BRAF Class 3 Melanoma |
| BRIP1 | 17q23.2 | 29 | 3.75 | 14 | 11.97 | 0.000599 | 0.005098 | (B) BRAF Class 3 Melanoma |
| ARID1B | 6q25.3 | 46 | 6.16 | 18 | 16.07 | 0.000641 | 0.005407 | (B) BRAF Class 3 Melanoma |
| CBLB | 3q13.11 | 8 | 2.76 | 8 | 16.00 | 0.00065 | 0.005436 | (B) BRAF Class 3 Melanoma |
| VEGFA | 6p21.1 | 7 | 1.00 | 7 | 6.93 | 0.000678 | 0.005588 | (B) BRAF Class 3 Melanoma |
| TP63 | 3q28 | 38 | 8.94 | 14 | 25.93 | 0.000683 | 0.005588 | (B) BRAF Class 3 Melanoma |
| KMT2B | 19q13.12 | 40 | 10.96 | 15 | 30.00 | 0.000687 | 0.005588 | (B) BRAF Class 3 Melanoma |
| BRCA1 | 17q21.31 | 34 | 4.21 | 15 | 12.30 | 0.000799 | 0.00644 | (B) BRAF Class 3 Melanoma |
| NTRK1 | 1q23.1 | 42 | 5.10 | 17 | 13.60 | 0.000815 | 0.006512 | (B) BRAF Class 3 Melanoma |
| JAK2 | 9p24.1 | 24 | 2.16 | 12 | 7.55 | 0.000834 | 0.006609 | (B) BRAF Class 3 Melanoma |
| ERG | 21q22.2 | 28 | 3.88 | 13 | 12.38 | 0.000875 | 0.006874 | (B) BRAF Class 3 Melanoma |
| ERBB2 | 17q12 | 24 | 2.03 | 12 | 7.02 | 0.000909 | 0.007081 | (B) BRAF Class 3 Melanoma |
| SPTA1 | 1q23.1 | 53 | 20.08 | 16 | 47.06 | 0.000952 | 0.007352 | (B) BRAF Class 3 Melanoma |
| SUFU | 10q24.32 | 13 | 1.68 | 9 | 7.69 | 0.000973 | 0.007446 | (B) BRAF Class 3 Melanoma |
| POLQ | 3q13.33 | 13 | 6.05 | 9 | 25.71 | 0.001002 | 0.007609 | (B) BRAF Class 3 Melanoma |
| PIK3CG | 7q22.3 | 30 | 5.68 | 13 | 17.33 | 0.001037 | 0.007804 | (B) BRAF Class 3 Melanoma |
| FLT4 | 5q35.3 | 61 | 8.01 | 21 | 18.10 | 0.001075 | 0.00803 | (B) BRAF Class 3 Melanoma |
| RUNX1 | 21q22.12 | 24 | 3.10 | 12 | 10.26 | 0.001202 | 0.008902 | (B) BRAF Class 3 Melanoma |
| FAT1 | 4q35.2 | 92 | 13.22 | 26 | 26.00 | 0.001224 | 0.008994 | (B) BRAF Class 3 Melanoma |
| MAGI2 | 7q21.11 | 18 | 7.89 | 8 | 32.00 | 0.001402 | 0.0102 | (B) BRAF Class 3 Melanoma |
| CCNE1 | 19q12 | 8 | 0.99 | 7 | 5.79 | 0.001409 | 0.0102 | (B) BRAF Class 3 Melanoma |
| MUTYH | 1p34.1 | 14 | 1.81 | 9 | 7.69 | 0.001423 | 0.0102 | (B) BRAF Class 3 Melanoma |
| NFKBIZ | 3q12.3 | 1 | 0.40 | 4 | 9.76 | 0.001511 | 0.0108 | (B) BRAF Class 3 Melanoma |
| TRAF7 | 16p13.3 | 12 | 1.86 | 8 | 8.70 | 0.001533 | 0.0108 | (B) BRAF Class 3 Melanoma |
| RECQL4 | 8q24.3 | 35 | 5.13 | 14 | 14.29 | 0.001534 | 0.0108 | (B) BRAF Class 3 Melanoma |
| TOP1 | 20q12 | 9 | 1.61 | 7 | 8.86 | 0.001576 | 0.011 | (B) BRAF Class 3 Melanoma |
| FGF14 | 13q33.1 | 4 | 1.38 | 5 | 12.82 | 0.001638 | 0.0113 | (B) BRAF Class 3 Melanoma |
| NTRK2 | 9q21.33 | 18 | 2.23 | 10 | 8.26 | 0.001647 | 0.0113 | (B) BRAF Class 3 Melanoma |
| UPF1 | 19p13.11 | 0 | 0.00 | 3 | 12.50 | 0.001663 | 0.0113 | (B) BRAF Class 3 Melanoma |
| ITK | 5q33.3 | 4 | 1.92 | 5 | 17.24 | 0.001746 | 0.0118 | (B) BRAF Class 3 Melanoma |
| GALNT12 | 9q22.33 | 0 | 0.00 | 3 | 7.50 | 0.001971 | 0.0132 | (B) BRAF Class 3 Melanoma |
| MEF2B | 19p13.11 | 6 | 0.78 | 6 | 5.13 | 0.002169 | 0.0144 | (B) BRAF Class 3 Melanoma |
| ALOX12B | 17p13.1 | 13 | 2.48 | 8 | 10.67 | 0.002201 | 0.0145 | (B) BRAF Class 3 Melanoma |
| EBF1 | 5q33.3 | 0 | 0.00 | 3 | 25.00 | 0.002227 | 0.0146 | (B) BRAF Class 3 Melanoma |
| CCNQ | Xq28 | 2 | 1.17 | 4 | 16.67 | 0.002291 | 0.0149 | (B) BRAF Class 3 Melanoma |
| FANCA | 16q24.3 | 27 | 3.34 | 12 | 9.92 | 0.002456 | 0.0159 | (B) BRAF Class 3 Melanoma |

| | | | | | | | | |
|---|---|---|---|---|---|---|---|---|
| NF2 | 22q12.2 | 16 | 1.97 | 9 | 7.44 | 0.002595 | 0.0165 | (B) BRAF Class 3 Melanoma |
| PIK3C2G | 12p12.3 | 47 | 9.94 | 15 | 23.81 | 0.002601 | 0.0165 | (B) BRAF Class 3 Melanoma |
| NECTIN4 | 1q23.3 | 2 | 1.00 | 4 | 13.79 | 0.002606 | 0.0165 | (B) BRAF Class 3 Melanoma |
| SNCAIP | 5q23.2 | 36 | 13.09 | 12 | 34.29 | 0.002689 | 0.0169 | (B) BRAF Class 3 Melanoma |
| CHEK1 | 11q24.2 | 2 | 0.26 | 4 | 3.67 | 0.002922 | 0.0182 | (B) BRAF Class 3 Melanoma |
| PTEN | 10q23.31 | 174 | 14.71 | 12 | 7.06 | 0.002968 | 0.0184 | (A) BRAF Class 1 Melanoma |
| TCF3 | 19p13.3 | 14 | 2.12 | 8 | 8.60 | 0.002987 | 0.0184 | (B) BRAF Class 3 Melanoma |
| LRP1B | q22.1-q22. | 86 | 30.82 | 21 | 55.26 | 0.003066 | 0.0186 | (B) BRAF Class 3 Melanoma |
| EZH2 | 7q36.1 | 33 | 3.19 | 13 | 8.67 | 0.003069 | 0.0186 | (B) BRAF Class 3 Melanoma |
| TFE3 | Xp11.23 | 5 | 1.22 | 5 | 8.93 | 0.003315 | 0.02 | (B) BRAF Class 3 Melanoma |
| PTPRB | 12q15 | 5 | 9.26 | 5 | 55.56 | 0.003333 | 0.02 | (B) BRAF Class 3 Melanoma |
| STAG2 | Xq25 | 24 | 3.11 | 11 | 9.40 | 0.003358 | 0.02 | (B) BRAF Class 3 Melanoma |
| XPO1 | 2p15 | 7 | 0.87 | 6 | 4.96 | 0.00342 | 0.0201 | (B) BRAF Class 3 Melanoma |
| COP1 | q25.1-q25. | 5 | 1.09 | 5 | 7.94 | 0.003427 | 0.0201 | (B) BRAF Class 3 Melanoma |
| HABP2 | 10q25.3 | 8 | 3.98 | 6 | 20.69 | 0.003529 | 0.0206 | (B) BRAF Class 3 Melanoma |
| MCM8 | 20p12.3 | 5 | 2.49 | 5 | 17.24 | 0.003657 | 0.0212 | (B) BRAF Class 3 Melanoma |
| SPRED1 | 15q14 | 2 | 0.97 | 4 | 12.12 | 0.003702 | 0.0213 | (B) BRAF Class 3 Melanoma |
| SRC | 20q11.23 | 8 | 0.86 | 6 | 4.65 | 0.003822 | 0.0219 | (B) BRAF Class 3 Melanoma |
| CXCR4 | 2q22.1 | 5 | 0.78 | 5 | 5.56 | 0.00392 | 0.0223 | (B) BRAF Class 3 Melanoma |
| IGF1R | 15q26.3 | 42 | 5.15 | 15 | 12.30 | 0.003961 | 0.0224 | (B) BRAF Class 3 Melanoma |
| MST1R | 3p21.31 | 13 | 2.77 | 7 | 11.67 | 0.003998 | 0.0225 | (B) BRAF Class 3 Melanoma |
| ERBB3 | 12q13.2 | 21 | 2.59 | 10 | 8.20 | 0.004023 | 0.0225 | (B) BRAF Class 3 Melanoma |
| CDK8 | 13q12.13 | 11 | 1.52 | 7 | 6.67 | 0.004225 | 0.0235 | (B) BRAF Class 3 Melanoma |
| ARHGAP35 | 19q13.32 | 14 | 5.83 | 8 | 21.05 | 0.004481 | 0.0247 | (B) BRAF Class 3 Melanoma |
| IDH2 | 15q26.1 | 8 | 0.72 | 6 | 3.68 | 0.005041 | 0.0277 | (B) BRAF Class 3 Melanoma |
| MCL1 | 1q21.2 | 5 | 0.65 | 5 | 4.27 | 0.005298 | 0.0289 | (B) BRAF Class 3 Melanoma |
| MTA1 | 14q32.33 | 3 | 1.49 | 4 | 13.79 | 0.005543 | 0.0301 | (B) BRAF Class 3 Melanoma |
| KEAP1 | 19p13.2 | 8 | 1.05 | 6 | 5.17 | 0.00567 | 0.0306 | (B) BRAF Class 3 Melanoma |
| RHOH | 4p14 | 3 | 1.32 | 4 | 12.12 | 0.005709 | 0.0306 | (B) BRAF Class 3 Melanoma |
| PPARG | 3p25.2 | 17 | 2.64 | 8 | 9.20 | 0.005741 | 0.0306 | (B) BRAF Class 3 Melanoma |
| CUL3 | 2q36.2 | 6 | 1.13 | 5 | 6.85 | 0.005817 | 0.0308 | (B) BRAF Class 3 Melanoma |
| MALT1 | 18q21.32 | 7 | 1.67 | 5 | 9.62 | 0.005844 | 0.0308 | (B) BRAF Class 3 Melanoma |
| RAF1 | 3p25.2 | 8 | 0.97 | 6 | 4.76 | 0.005915 | 0.031 | (B) BRAF Class 3 Melanoma |
| DOCK8 | 9p24.3 | 13 | 6.47 | 7 | 24.14 | 0.00597 | 0.031 | (B) BRAF Class 3 Melanoma |
| COL2A1 | 12q13.11 | 3 | 5.56 | 4 | 44.44 | 0.005983 | 0.031 | (B) BRAF Class 3 Melanoma |
| LZTR1 | 21\|22q11.: | 7 | 2.62 | 5 | 14.71 | 0.006094 | 0.0313 | (B) BRAF Class 3 Melanoma |
| FGFR3 | 4p16.3 | 32 | 2.70 | 12 | 7.02 | 0.006109 | 0.0313 | (B) BRAF Class 3 Melanoma |
| USP6 | 17p13.2 | 0 | 0.00 | 2 | 33.33 | 0.006211 | 0.0316 | (B) BRAF Class 3 Melanoma |
| AR | Xq12 | 31 | 3.84 | 12 | 9.84 | 0.006425 | 0.0325 | (B) BRAF Class 3 Melanoma |
| PIK3CD | 1p36.22 | 7 | 1.62 | 5 | 9.09 | 0.006499 | 0.0327 | (B) BRAF Class 3 Melanoma |
| DNMT3B | 20q11.21 | 11 | 2.70 | 6 | 11.76 | 0.006669 | 0.0334 | (B) BRAF Class 3 Melanoma |
| HOXB13 | 17q21.32 | 6 | 0.90 | 5 | 5.26 | 0.00676 | 0.0337 | (B) BRAF Class 3 Melanoma |
| LYN | 8q12.1 | 15 | 3.83 | 7 | 14.00 | 0.007009 | 0.0347 | (B) BRAF Class 3 Melanoma |
| TCF7L2 | )q25.2-q25 | 9 | 1.30 | 6 | 5.88 | 0.007453 | 0.0367 | (B) BRAF Class 3 Melanoma |
| NSD2 | 4p16.3 | 12 | 2.80 | 7 | 10.45 | 0.008096 | 0.0397 | (B) BRAF Class 3 Melanoma |
| PALB2 | 16p12.2 | 24 | 2.97 | 10 | 8.26 | 0.008157 | 0.0397 | (B) BRAF Class 3 Melanoma |
| TOPBP1 | 3q22.1 | 7 | 3.40 | 5 | 17.24 | 0.008681 | 0.0421 | (B) BRAF Class 3 Melanoma |
| GLI3 | 7p14.1 | 6 | 8.70 | 5 | 41.67 | 0.008746 | 0.0422 | (B) BRAF Class 3 Melanoma |

| | | | | | | | | |
|---|---|---|---|---|---|---|---|---|
| ZFHX4 | 8q21.13 | 11 | 20.37 | 6 | 66.67 | 0.008836 | 0.0422 | (B) BRAF Class 3 Melanoma |
| ARAF | Xp11.3 | 6 | 0.73 | 5 | 4.00 | 0.00884 | 0.0422 | (B) BRAF Class 3 Melanoma |
| XPC | 3p25.1 | 9 | 3.17 | 6 | 13.33 | 0.009062 | 0.043 | (B) BRAF Class 3 Melanoma |
| FAT3 | 11q14.3 | 8 | 7.41 | 6 | 30.00 | 0.009239 | 0.0436 | (B) BRAF Class 3 Melanoma |
| PRKCZ | 1p36.33 | 1 | 1.45 | 3 | 25.00 | 0.009422 | 0.0442 | (B) BRAF Class 3 Melanoma |
| PAX7 | 1p36.13 | 5 | 2.25 | 4 | 14.81 | 0.009574 | 0.0447 | (B) BRAF Class 3 Melanoma |
| SMAD4 | 18q21.2 | 10 | 0.89 | 6 | 3.70 | 0.009732 | 0.0452 | (B) BRAF Class 3 Melanoma |
| FANCI | 15q26.1 | 11 | 2.46 | 6 | 9.84 | 0.009962 | 0.0461 | (B) BRAF Class 3 Melanoma |
| XRCC4 | 5q14.2 | 4 | 1.99 | 4 | 13.79 | 0.0101 | 0.0465 | (B) BRAF Class 3 Melanoma |
| SLX4 | 16p13.3 | 29 | 5.47 | 10 | 14.08 | 0.0105 | 0.0481 | (B) BRAF Class 3 Melanoma |
| INHBA | 7p14.1 | 23 | 4.49 | 9 | 12.50 | 0.0106 | 0.0483 | (B) BRAF Class 3 Melanoma |
| EPCAM | 2p21 | 4 | 0.61 | 4 | 4.30 | 0.0107 | 0.0483 | (B) BRAF Class 3 Melanoma |
| CDH1 | 16q22.1 | 18 | 1.67 | 8 | 5.19 | 0.0107 | 0.0483 | (B) BRAF Class 3 Melanoma |
| TENT5C | 1p12 | 17 | 2.20 | 8 | 6.84 | 0.011 | 0.0493 | (B) BRAF Class 3 Melanoma |
| RAC1 | 7p22.1 | 36 | 4.38 | 12 | 10.17 | 0.0116 | 0.0519 | (B) BRAF Class 3 Melanoma |
| ATR | 3q23 | 27 | 3.57 | 10 | 9.17 | 0.012 | 0.053 | (B) BRAF Class 3 Melanoma |
| POLD1 | 19q13.33 | 27 | 3.82 | 10 | 9.80 | 0.012 | 0.053 | (B) BRAF Class 3 Melanoma |
| PGR | 11q22.1 | 18 | 4.58 | 7 | 14.58 | 0.0121 | 0.0534 | (B) BRAF Class 3 Melanoma |
| SPEN | :36.21-p36. | 53 | 9.85 | 15 | 19.74 | 0.0124 | 0.0543 | (B) BRAF Class 3 Melanoma |
| SOX9 | 17q24.3 | 7 | 0.92 | 5 | 4.42 | 0.0126 | 0.0548 | (B) BRAF Class 3 Melanoma |
| PHOX2B | 4p13 | 18 | 2.55 | 8 | 7.62 | 0.0128 | 0.0556 | (B) BRAF Class 3 Melanoma |
| DNMT3A | 2p23.3 | 26 | 3.33 | 10 | 8.47 | 0.0132 | 0.057 | (B) BRAF Class 3 Melanoma |
| MYCN | 2p24.3 | 14 | 1.73 | 7 | 5.74 | 0.0134 | 0.0577 | (B) BRAF Class 3 Melanoma |
| DIS3 | 13q21.33 | 14 | 1.87 | 7 | 6.14 | 0.0138 | 0.0589 | (B) BRAF Class 3 Melanoma |
| CREBBP | 16p13.3 | 40 | 4.95 | 13 | 10.74 | 0.0138 | 0.0589 | (B) BRAF Class 3 Melanoma |
| PPP2R1A | 19q13.41 | 15 | 1.98 | 7 | 6.42 | 0.014 | 0.0595 | (B) BRAF Class 3 Melanoma |
| MSI2 | 17q22 | 0 | 0.00 | 2 | 8.33 | 0.0146 | 0.0615 | (B) BRAF Class 3 Melanoma |
| RB1 | 13q14.2 | 33 | 2.96 | 11 | 6.96 | 0.015 | 0.0629 | (B) BRAF Class 3 Melanoma |
| POLB | 8p11.21 | 0 | 0.00 | 2 | 6.90 | 0.0154 | 0.0644 | (B) BRAF Class 3 Melanoma |
| GLI2 | 2q14.2 | 32 | 11.11 | 12 | 24.00 | 0.0156 | 0.065 | (B) BRAF Class 3 Melanoma |
| ETV1 | 7p21.2 | 15 | 2.07 | 7 | 6.54 | 0.0159 | 0.0658 | (B) BRAF Class 3 Melanoma |
| ARHGEF12 | 11q23.3 | 5 | 2.46 | 4 | 13.79 | 0.0161 | 0.0659 | (B) BRAF Class 3 Melanoma |
| UIMC1 | 5q35.2 | 5 | 2.46 | 4 | 13.79 | 0.0161 | 0.0659 | (B) BRAF Class 3 Melanoma |
| RICTOR | 5p13.1 | 24 | 3.17 | 9 | 8.26 | 0.0161 | 0.0659 | (B) BRAF Class 3 Melanoma |
| EML4 | 2p21 | 4 | 2.15 | 3 | 16.67 | 0.0163 | 0.0662 | (B) BRAF Class 3 Melanoma |
| PMS1 | 2q32.2 | 15 | 2.13 | 7 | 6.67 | 0.0163 | 0.0662 | (B) BRAF Class 3 Melanoma |
| RHBDF2 | 17q25.1 | 5 | 2.49 | 4 | 13.79 | 0.0166 | 0.0669 | (B) BRAF Class 3 Melanoma |
| DOT1L | 19p13.3 | 25 | 4.65 | 9 | 11.84 | 0.0167 | 0.0671 | (B) BRAF Class 3 Melanoma |
| DROSHA | 5p13.3 | 9 | 4.66 | 5 | 18.52 | 0.0176 | 0.0703 | (B) BRAF Class 3 Melanoma |
| MLLT3 | 9p21.3 | 1 | 0.60 | 2 | 13.33 | 0.018 | 0.0714 | (B) BRAF Class 3 Melanoma |
| SPOP | 17q21.33 | 5 | 0.66 | 4 | 3.67 | 0.0182 | 0.0714 | (B) BRAF Class 3 Melanoma |
| RRAGC | 1p34.3 | 0 | 0.00 | 2 | 6.06 | 0.0184 | 0.0714 | (B) BRAF Class 3 Melanoma |
| RRAS | 19q13.33 | 0 | 0.00 | 2 | 6.06 | 0.0184 | 0.0714 | (B) BRAF Class 3 Melanoma |
| RRAS2 | 11p15.2 | 0 | 0.00 | 2 | 6.06 | 0.0184 | 0.0714 | (B) BRAF Class 3 Melanoma |
| POT1 | 7q31.33 | 2 | 0.74 | 3 | 7.14 | 0.0184 | 0.0714 | (B) BRAF Class 3 Melanoma |
| PRKCA | 17q24.2 | 0 | 0.00 | 2 | 16.67 | 0.0185 | 0.0714 | (B) BRAF Class 3 Melanoma |
| TFEB | 6p21.1 | 0 | 0.00 | 2 | 16.67 | 0.0185 | 0.0714 | (B) BRAF Class 3 Melanoma |
| MYCL | 1p34.2 | 8 | 0.99 | 5 | 4.13 | 0.0187 | 0.0718 | (B) BRAF Class 3 Melanoma |

| | | | | | | | | |
|---|---|---|---|---|---|---|---|---|
| LIFR | 5p13.1 | 2 | 3.39 | 3 | 30.00 | 0.0194 | 0.0742 | (B) BRAF Class 3 Melanoma |
| RBBP8 | 18q11.2 | 9 | 4.48 | 5 | 17.24 | 0.02 | 0.0762 | (B) BRAF Class 3 Melanoma |
| CEBPA | 19q13.11 | 5 | 0.66 | 4 | 3.51 | 0.0203 | 0.0768 | (B) BRAF Class 3 Melanoma |
| E2F3 | 6p22.3 | 3 | 0.74 | 3 | 5.88 | 0.0203 | 0.0768 | (B) BRAF Class 3 Melanoma |
| ADGRA2 | 8p11.23 | 6 | 2.29 | 4 | 11.43 | 0.0204 | 0.0768 | (B) BRAF Class 3 Melanoma |
| NCKAP5 | 2q21.2 | 2 | 6.25 | 3 | 50.00 | 0.0207 | 0.0775 | (B) BRAF Class 3 Melanoma |
| GSK3B | 3q13.33 | 9 | 1.67 | 5 | 6.58 | 0.0208 | 0.0775 | (B) BRAF Class 3 Melanoma |
| MDM2 | 12q15 | 12 | 1.49 | 6 | 4.96 | 0.0212 | 0.079 | (B) BRAF Class 3 Melanoma |
| ZNRF3 | 22q12.1 | 9 | 3.56 | 5 | 13.51 | 0.0219 | 0.0809 | (B) BRAF Class 3 Melanoma |
| MSH2 | 2p21-p16.3 | 16 | 2.04 | 7 | 5.93 | 0.022 | 0.0811 | (B) BRAF Class 3 Melanoma |
| CDC73 | 1q31.2 | 12 | 1.55 | 6 | 5.13 | 0.0221 | 0.0813 | (B) BRAF Class 3 Melanoma |
| GRM3 | 21.11-q21. | 31 | 11.27 | 9 | 25.71 | 0.0224 | 0.0819 | (B) BRAF Class 3 Melanoma |
| PRDM14 | 8q13.3 | 6 | 3.51 | 4 | 16.67 | 0.0226 | 0.0823 | (B) BRAF Class 3 Melanoma |
| AMER1 | Xq11.2 | 22 | 4.08 | 8 | 10.53 | 0.0227 | 0.0823 | (B) BRAF Class 3 Melanoma |
| BCL6 | 3q27.3 | 16 | 2.07 | 7 | 5.98 | 0.0228 | 0.0823 | (B) BRAF Class 3 Melanoma |
| NSD3 | 8p11.23 | 9 | 2.45 | 5 | 9.26 | 0.0233 | 0.0837 | (B) BRAF Class 3 Melanoma |
| ABL1 | 9q34.12 | 26 | 2.41 | 9 | 5.84 | 0.0233 | 0.0837 | (B) BRAF Class 3 Melanoma |
| DCC | 18q21.2 | 13 | 16.46 | 6 | 46.15 | 0.0242 | 0.0864 | (B) BRAF Class 3 Melanoma |
| HGF | 7q21.11 | 37 | 6.88 | 11 | 14.47 | 0.0247 | 0.088 | (B) BRAF Class 3 Melanoma |
| NEIL3 | 4q34.3 | 6 | 2.99 | 4 | 13.79 | 0.0252 | 0.089 | (B) BRAF Class 3 Melanoma |
| PTPN14 | 1q32.3-q41 | 6 | 2.99 | 4 | 13.79 | 0.0252 | 0.089 | (B) BRAF Class 3 Melanoma |
| TSC1 | 9q34.13 | 30 | 3.70 | 10 | 8.26 | 0.0259 | 0.0911 | (B) BRAF Class 3 Melanoma |
| MEN1 | 11q13.1 | 9 | 1.16 | 5 | 4.27 | 0.027 | 0.0947 | (B) BRAF Class 3 Melanoma |
| ELOC | 8q21.11 | 3 | 0.49 | 3 | 3.49 | 0.0274 | 0.0957 | (B) BRAF Class 3 Melanoma |
| SF3B1 | 2q33.1 | 21 | 2.53 | 8 | 6.35 | 0.0278 | 0.096 | (B) BRAF Class 3 Melanoma |
| BABAM2 | 2p23.2 | 3 | 1.49 | 3 | 10.34 | 0.0279 | 0.096 | (B) BRAF Class 3 Melanoma |
| SERPINA1 | 14q32.13 | 3 | 1.49 | 3 | 10.34 | 0.0279 | 0.096 | (B) BRAF Class 3 Melanoma |
| UROD | 1p34.1 | 3 | 1.49 | 3 | 10.34 | 0.0279 | 0.096 | (B) BRAF Class 3 Melanoma |
| ACVR1 | 2q24.1 | 10 | 1.59 | 5 | 5.62 | 0.0286 | 0.0979 | (B) BRAF Class 3 Melanoma |
| PAX5 | 9p13.2 | 13 | 1.66 | 6 | 5.08 | 0.0288 | 0.0981 | (B) BRAF Class 3 Melanoma |
| RAD51B | 14q24.1 | 7 | 1.37 | 4 | 5.97 | 0.0292 | 0.0991 | (B) BRAF Class 3 Melanoma |
| XRCC1 | 19q13.31 | 3 | 1.38 | 3 | 9.38 | 0.0293 | 0.0991 | (B) BRAF Class 3 Melanoma |
| TBX3 | 12q24.21 | 28 | 5.29 | 9 | 12.00 | 0.0294 | 0.0993 | (B) BRAF Class 3 Melanoma |
| CDKN1A | 6p21.2 | 3 | 0.40 | 3 | 2.65 | 0.0317 | 0.107 | (B) BRAF Class 3 Melanoma |
| EIF4A2 | 3q27.3 | 1 | 0.25 | 2 | 4.17 | 0.0322 | 0.108 | (B) BRAF Class 3 Melanoma |
| SRSF2 | 17q25.1 | 10 | 1.44 | 5 | 4.90 | 0.0328 | 0.109 | (B) BRAF Class 3 Melanoma |
| COL1A1 | 17q21.33 | 3 | 3.70 | 3 | 23.08 | 0.0329 | 0.109 | (B) BRAF Class 3 Melanoma |
| NCOA2 | 8q13.3 | 3 | 3.70 | 3 | 23.08 | 0.0329 | 0.109 | (B) BRAF Class 3 Melanoma |
| FGF5 | 4q21.21 | 2 | 1.26 | 2 | 14.29 | 0.0333 | 0.11 | (B) BRAF Class 3 Melanoma |
| EED | 11q14.2 | 7 | 1.40 | 4 | 5.80 | 0.0338 | 0.111 | (B) BRAF Class 3 Melanoma |
| ESR1 | q25.1-q25. | 18 | 2.08 | 7 | 5.38 | 0.0346 | 0.114 | (B) BRAF Class 3 Melanoma |
| FRS2 | 12q15 | 1 | 0.38 | 2 | 5.88 | 0.0353 | 0.115 | (B) BRAF Class 3 Melanoma |
| TYK2 | 19p13.2 | 3 | 3.85 | 3 | 23.08 | 0.036 | 0.117 | (B) BRAF Class 3 Melanoma |
| MLH3 | 14q24.3 | 3 | 1.15 | 3 | 7.14 | 0.0368 | 0.119 | (B) BRAF Class 3 Melanoma |
| RIT1 | 1q22 | 7 | 1.08 | 4 | 4.35 | 0.0374 | 0.121 | (B) BRAF Class 3 Melanoma |
| CDK12 | 17q12 | 24 | 3.17 | 8 | 7.34 | 0.0379 | 0.122 | (B) BRAF Class 3 Melanoma |
| MDC1 | 6p21.33 | 29 | 7.11 | 8 | 15.69 | 0.0401 | 0.128 | (B) BRAF Class 3 Melanoma |
| KAT6A | 8p11.21 | 20 | 4.28 | 7 | 10.45 | 0.0404 | 0.128 | (B) BRAF Class 3 Melanoma |

| | | | | | | | | |
|---|---|---|---|---|---|---|---|---|
| EPAS1 | 2p21 | 1 | 0.58 | 2 | 8.33 | 0.0404 | 0.128 | (B) BRAF Class 3 Melanoma |
| SESN2 | 1p35.3 | 1 | 0.58 | 2 | 8.33 | 0.0404 | 0.128 | (B) BRAF Class 3 Melanoma |
| FLCN | 17p11.2 | 7 | 0.91 | 4 | 3.51 | 0.0424 | 0.134 | (B) BRAF Class 3 Melanoma |
| GPC3 | Xq26.2 | 6 | 1.92 | 4 | 7.55 | 0.0427 | 0.134 | (B) BRAF Class 3 Melanoma |
| PLCG2 | 16q23.3 | 18 | 3.84 | 6 | 10.00 | 0.0434 | 0.136 | (B) BRAF Class 3 Melanoma |
| NIPBL | 5p13.2 | 1 | 1.69 | 2 | 22.22 | 0.0441 | 0.138 | (B) BRAF Class 3 Melanoma |
| GNAQ | 9q21.2 | 11 | 0.98 | 5 | 3.01 | 0.0445 | 0.139 | (B) BRAF Class 3 Melanoma |
| BCL10 | 1p22.3 | 5 | 1.20 | 3 | 5.77 | 0.0479 | 0.149 | (B) BRAF Class 3 Melanoma |
| PDCD1 | 2q37.3 | 5 | 1.14 | 3 | 5.45 | 0.0491 | 0.152 | (B) BRAF Class 3 Melanoma |
| USP28 | 11q23.2 | 8 | 3.98 | 4 | 13.79 | 0.0494 | 0.153 | (B) BRAF Class 3 Melanoma |
| RAD21 | 8q24.11 | 16 | 2.27 | 6 | 5.83 | 0.0499 | 0.154 | (B) BRAF Class 3 Melanoma |
| ERCC3 | 2q14.3 | 12 | 1.79 | 5 | 5.21 | 0.0506 | 0.155 | (B) BRAF Class 3 Melanoma |
| CDKN1B | 12p13.1 | 4 | 0.50 | 3 | 2.48 | 0.051 | 0.156 | (B) BRAF Class 3 Melanoma |
| SMARCB1 | ı11.23\|22c | 8 | 0.72 | 4 | 2.53 | 0.0512 | 0.156 | (B) BRAF Class 3 Melanoma |
| CDKN2C | 1p32.3 | 4 | 0.52 | 3 | 2.56 | 0.0525 | 0.159 | (B) BRAF Class 3 Melanoma |
| PIK3CB | 3q22.3 | 14 | 2.64 | 5 | 7.35 | 0.0533 | 0.161 | (B) BRAF Class 3 Melanoma |
| RASA2 | 3q23 | 1 | 1.41 | 2 | 16.67 | 0.0534 | 0.161 | (B) BRAF Class 3 Melanoma |
| CHD5 | 1p36.31 | 4 | 7.41 | 3 | 33.33 | 0.054 | 0.162 | (B) BRAF Class 3 Melanoma |
| PTK2 | 8q24.3 | 1 | 1.45 | 2 | 16.67 | 0.056 | 0.167 | (B) BRAF Class 3 Melanoma |
| RHPN2 | 19q13.11 | 1 | 1.45 | 2 | 16.67 | 0.056 | 0.167 | (B) BRAF Class 3 Melanoma |
| BCORL1 | Xq26.1 | 30 | 5.68 | 9 | 11.25 | 0.0566 | 0.168 | (B) BRAF Class 3 Melanoma |
| FGF3 | 11q13.3 | 13 | 2.27 | 5 | 6.25 | 0.0575 | 0.17 | (B) BRAF Class 3 Melanoma |
| RSPO2 | 8q23.1 | 4 | 1.28 | 3 | 6.00 | 0.0577 | 0.17 | (B) BRAF Class 3 Melanoma |
| MRE11 | 11q21 | 17 | 2.25 | 6 | 5.50 | 0.0578 | 0.17 | (B) BRAF Class 3 Melanoma |
| IKZF2 | 2q34 | 4 | 5.63 | 3 | 25.00 | 0.0588 | 0.173 | (B) BRAF Class 3 Melanoma |
| MITF | 3p13 | 35 | 4.53 | 10 | 8.55 | 0.0591 | 0.173 | (B) BRAF Class 3 Melanoma |
| CSDE1 | 1p13.2 | 2 | 0.64 | 2 | 5.26 | 0.0599 | 0.175 | (B) BRAF Class 3 Melanoma |
| HIST3H3 | 1q42.13 | 2 | 0.51 | 2 | 4.17 | 0.0604 | 0.176 | (B) BRAF Class 3 Melanoma |
| REL | 2p16.1 | 8 | 1.04 | 4 | 3.42 | 0.0612 | 0.177 | (B) BRAF Class 3 Melanoma |
| CSF1R | 5q32 | 28 | 3.04 | 8 | 6.25 | 0.0615 | 0.178 | (B) BRAF Class 3 Melanoma |
| FAS | 10q23.31 | 5 | 1.02 | 3 | 4.41 | 0.0624 | 0.18 | (B) BRAF Class 3 Melanoma |
| STK40 | 1p34.3 | 2 | 0.49 | 2 | 3.92 | 0.0628 | 0.18 | (B) BRAF Class 3 Melanoma |
| NEGR1 | 1p31.1 | 14 | 3.22 | 5 | 8.47 | 0.0636 | 0.182 | (B) BRAF Class 3 Melanoma |
| IRF4 | 6p25.3 | 5 | 0.93 | 3 | 3.95 | 0.0642 | 0.183 | (B) BRAF Class 3 Melanoma |
| ZFHX3 | 5q22.2-q22 | 36 | 8.39 | 9 | 15.79 | 0.0651 | 0.185 | (B) BRAF Class 3 Melanoma |
| GATA2 | 3q21.3 | 13 | 1.74 | 5 | 4.63 | 0.0653 | 0.185 | (B) BRAF Class 3 Melanoma |
| ERCC2 | 19q13.32 | 26 | 3.27 | 8 | 6.67 | 0.0655 | 0.185 | (B) BRAF Class 3 Melanoma |
| PRF1 | 10q22.1 | 12 | 4.63 | 5 | 12.20 | 0.0658 | 0.185 | (B) BRAF Class 3 Melanoma |
| MTAP | 9p21.3 | 19 | 8.84 | 6 | 20.00 | 0.0659 | 0.185 | (B) BRAF Class 3 Melanoma |
| EXT1 | 8q24.11 | 8 | 2.82 | 4 | 8.89 | 0.0662 | 0.185 | (B) BRAF Class 3 Melanoma |
| TRAF2 | 9q34.3 | 6 | 1.46 | 3 | 5.88 | 0.0663 | 0.185 | (B) BRAF Class 3 Melanoma |
| YES1 | 18p11.32 | 6 | 1.47 | 3 | 5.88 | 0.0671 | 0.187 | (B) BRAF Class 3 Melanoma |
| NFKBIE | 6p21.1 | 8 | 3.38 | 4 | 10.53 | 0.0677 | 0.188 | (B) BRAF Class 3 Melanoma |
| SOS1 | 2p22.1 | 8 | 2.05 | 4 | 6.45 | 0.0678 | 0.188 | (B) BRAF Class 3 Melanoma |
| ERCC5 | 13q33.1 | 18 | 2.69 | 6 | 6.25 | 0.0682 | 0.188 | (B) BRAF Class 3 Melanoma |
| ABL2 | 1q25.2 | 6 | 2.19 | 3 | 8.57 | 0.0695 | 0.191 | (B) BRAF Class 3 Melanoma |
| STK11 | 19p13.3 | 18 | 1.60 | 6 | 3.70 | 0.0697 | 0.191 | (B) BRAF Class 3 Melanoma |
| SMARCAL1 | 2q35 | 0 | 0.00 | 1 | 33.33 | 0.0698 | 0.191 | (B) BRAF Class 3 Melanoma |

| | | | | | | | | |
|---|---|---|---|---|---|---|---|---|
| JAK1 | 1p31.3 | 18 | 2.37 | 6 | 5.45 | 0.0708 | 0.193 | (B) BRAF Class 3 Melanoma |
| ANKRD11 | 16q24.3 | 28 | 7.12 | 7 | 14.58 | 0.0714 | 0.194 | (B) BRAF Class 3 Melanoma |
| BCL2L12 | 19q13.33 | 4 | 1.50 | 3 | 6.38 | 0.0718 | 0.195 | (B) BRAF Class 3 Melanoma |
| PPM1D | 17q23.2 | 14 | 2.29 | 5 | 5.81 | 0.0724 | 0.196 | (B) BRAF Class 3 Melanoma |
| SMAD3 | 15q22.33 | 2 | 0.39 | 2 | 2.82 | 0.0743 | 0.2 | (B) BRAF Class 3 Melanoma |
| FANCG | 9p13.3 | 5 | 0.95 | 3 | 3.75 | 0.0749 | 0.201 | (B) BRAF Class 3 Melanoma |
| KDM6A | Xp11.3 | 13 | 1.68 | 5 | 4.27 | 0.0752 | 0.201 | (B) BRAF Class 3 Melanoma |
| TNFAIP3 | 6q23.3 | 13 | 1.68 | 5 | 4.27 | 0.0752 | 0.201 | (B) BRAF Class 3 Melanoma |
| MYB | 6q23.3 | 9 | 1.75 | 4 | 5.19 | 0.0756 | 0.201 | (B) BRAF Class 3 Melanoma |
| H3F3B | 17q25.1 | 2 | 0.32 | 2 | 2.25 | 0.0768 | 0.204 | (B) BRAF Class 3 Melanoma |
| PDCD1LG2 | 9p24.1 | 9 | 1.36 | 4 | 4.00 | 0.0775 | 0.205 | (B) BRAF Class 3 Melanoma |
| DMC1 | 22q13.1 | 2 | 1.00 | 2 | 6.90 | 0.0785 | 0.206 | (B) BRAF Class 3 Melanoma |
| NR0B1 | Xp21.2 | 2 | 1.00 | 2 | 6.90 | 0.0785 | 0.206 | (B) BRAF Class 3 Melanoma |
| RHOT1 | 17q11.2 | 2 | 1.00 | 2 | 6.90 | 0.0785 | 0.206 | (B) BRAF Class 3 Melanoma |
| FOXP1 | 3p13 | 10 | 1.93 | 4 | 5.56 | 0.0789 | 0.207 | (B) BRAF Class 3 Melanoma |
| EPHA5 | q13.1-q13. | 18 | 3.11 | 6 | 6.90 | 0.0802 | 0.209 | (B) BRAF Class 3 Melanoma |
| TGFBR2 | 3p24.1 | 10 | 1.86 | 4 | 5.26 | 0.0821 | 0.214 | (B) BRAF Class 3 Melanoma |
| MSH6 | 2p16.3 | 28 | 3.45 | 8 | 6.61 | 0.0827 | 0.215 | (B) BRAF Class 3 Melanoma |
| ZMYM2 | 13q12.11 | 0 | 0.00 | 1 | 25.00 | 0.0833 | 0.216 | (B) BRAF Class 3 Melanoma |
| BCL11B | 14q32.2 | 15 | 6.44 | 5 | 15.15 | 0.0845 | 0.218 | (B) BRAF Class 3 Melanoma |
| ADAMTS20 | 12q12 | 0 | 0.00 | 1 | 25.00 | 0.0851 | 0.218 | (B) BRAF Class 3 Melanoma |
| MAGEA1 | Xq28 | 0 | 0.00 | 1 | 25.00 | 0.0851 | 0.218 | (B) BRAF Class 3 Melanoma |
| THBS1 | 15q14 | 0 | 0.00 | 1 | 25.00 | 0.0851 | 0.218 | (B) BRAF Class 3 Melanoma |
| SLITRK6 | 13q31.1 | 6 | 8.11 | 4 | 23.53 | 0.0865 | 0.221 | (B) BRAF Class 3 Melanoma |
| WAS | Xp11.23 | 6 | 2.58 | 3 | 9.09 | 0.0868 | 0.221 | (B) BRAF Class 3 Melanoma |
| IRS1 | 2q36.3 | 7 | 1.72 | 3 | 5.88 | 0.0884 | 0.224 | (B) BRAF Class 3 Melanoma |
| NCOR1 | .7p12-p11.2 | 30 | 5.83 | 8 | 10.81 | 0.0894 | 0.226 | (B) BRAF Class 3 Melanoma |
| FLI1 | 11q24.3 | 4 | 2.17 | 2 | 11.11 | 0.0909 | 0.229 | (B) BRAF Class 3 Melanoma |
| EZH1 | 17q21.2 | 2 | 0.97 | 2 | 6.06 | 0.0922 | 0.232 | (B) BRAF Class 3 Melanoma |
| ETV6 | 12p13.2 | 14 | 1.82 | 5 | 4.27 | 0.0935 | 0.234 | (B) BRAF Class 3 Melanoma |
| ETS1 | 11q24.3 | 3 | 1.36 | 2 | 7.41 | 0.0937 | 0.234 | (B) BRAF Class 3 Melanoma |
| RSPO3 | 6q22.33 | 2 | 0.79 | 2 | 4.88 | 0.0943 | 0.235 | (B) BRAF Class 3 Melanoma |
| DDX3X | Xp11.4 | 2 | 2.56 | 2 | 15.38 | 0.0963 | 0.24 | (B) BRAF Class 3 Melanoma |
| H3F3C | 12p11.21 | 3 | 0.74 | 2 | 3.92 | 0.0973 | 0.241 | (B) BRAF Class 3 Melanoma |
| MAP2K2 | 19p13.3 | 15 | 1.98 | 5 | 4.55 | 0.0974 | 0.241 | (B) BRAF Class 3 Melanoma |
| GNA11 | 19p13.3 | 6 | 0.54 | 3 | 1.81 | 0.0987 | 0.244 | (B) BRAF Class 3 Melanoma |
| PRKAR1A | 17q24.2 | 10 | 1.29 | 4 | 3.42 | 0.0995 | 0.245 | (B) BRAF Class 3 Melanoma |
| TLR4 | 9q33.1 | 7 | 7.07 | 4 | 19.05 | 0.1 | 0.246 | (B) BRAF Class 3 Melanoma |
| RBL2 | 16q12.2 | 2 | 2.90 | 2 | 16.67 | 0.102 | 0.251 | (B) BRAF Class 3 Melanoma |
| GATA4 | 8p23.1 | 11 | 2.33 | 4 | 5.97 | 0.103 | 0.251 | (B) BRAF Class 3 Melanoma |
| RIF1 | 2q23.3 | 11 | 5.47 | 4 | 13.79 | 0.103 | 0.252 | (B) BRAF Class 3 Melanoma |
| PIK3R2 | 19p13.11 | 11 | 2.08 | 4 | 5.33 | 0.104 | 0.252 | (B) BRAF Class 3 Melanoma |
| BIRC2 | 11q22.2 | 0 | 0.00 | 1 | 20.00 | 0.104 | 0.252 | (B) BRAF Class 3 Melanoma |
| TEX12 | 11q23.1 | 0 | 0.00 | 1 | 5.88 | 0.105 | 0.254 | (B) BRAF Class 3 Melanoma |
| GEN1 | 2p24.2 | 3 | 0.74 | 2 | 3.70 | 0.106 | 0.256 | (B) BRAF Class 3 Melanoma |
| LATS2 | 13q12.11 | 13 | 3.17 | 4 | 7.84 | 0.107 | 0.256 | (B) BRAF Class 3 Melanoma |
| DNAH9 | 17p12 | 2 | 7.41 | 1 | 100.00 | 0.107 | 0.257 | (B) BRAF Class 3 Melanoma |
| NRG1 | 8p12 | 18 | 4.72 | 5 | 10.20 | 0.107 | 0.257 | (B) BRAF Class 3 Melanoma |

| | | | | | | | | |
|---|---|---|---|---|---|---|---|---|
| KAT6B | 10q22.2 | 16 | 7.02 | 5 | 15.15 | 0.108 | 0.257 | (B) BRAF Class 3 Melanoma |
| FOXF1 | 16q24.1 | 1 | 4.76 | 0 | 0.00 | 0.109 | 0.259 | (A) BRAF Class 1 Melanoma |
| SLFN11 | 17q12 | 1 | 4.76 | 0 | 0.00 | 0.109 | 0.259 | (A) BRAF Class 1 Melanoma |
| PLCB4 | [illegible]p12.3-p12 | 5 | 7.25 | 3 | 23.08 | 0.11 | 0.259 | (B) BRAF Class 3 Melanoma |
| TSHZ2 | 20q13.2 | 2 | 6.25 | 2 | 33.33 | 0.11 | 0.259 | (B) BRAF Class 3 Melanoma |
| ABRAXAS1 | 4q21.23 | 3 | 0.45 | 2 | 2.17 | 0.115 | 0.27 | (B) BRAF Class 3 Melanoma |
| SHOC2 | 10q25.2 | 3 | 1.75 | 2 | 8.33 | 0.115 | 0.27 | (B) BRAF Class 3 Melanoma |
| CDH23 | 10q22.1 | 2 | 8.00 | 1 | 100.00 | 0.115 | 0.27 | (B) BRAF Class 3 Melanoma |
| MED12 | Xq13.1 | 37 | 4.89 | 9 | 8.18 | 0.116 | 0.272 | (B) BRAF Class 3 Melanoma |
| FAN1 | 15q13.3 | 7 | 3.48 | 3 | 10.34 | 0.118 | 0.273 | (B) BRAF Class 3 Melanoma |
| XRCC6 | 22q13.2 | 7 | 3.48 | 3 | 10.34 | 0.118 | 0.273 | (B) BRAF Class 3 Melanoma |
| USP8 | 15q21.2 | 3 | 1.47 | 2 | 6.90 | 0.118 | 0.274 | (B) BRAF Class 3 Melanoma |
| MCM7 | 7q22.1 | 1 | 4.35 | 0 | 0.00 | 0.119 | 0.274 | (A) BRAF Class 1 Melanoma |
| TAL1 | 1p33 | 7 | 3.07 | 3 | 9.09 | 0.119 | 0.274 | (B) BRAF Class 3 Melanoma |
| DDB1 | 11q12.2 | 3 | 1.49 | 2 | 6.90 | 0.121 | 0.277 | (B) BRAF Class 3 Melanoma |
| RINT1 | 7q22.3 | 3 | 1.49 | 2 | 6.90 | 0.121 | 0.277 | (B) BRAF Class 3 Melanoma |
| NSD1 | 5q35.3 | 48 | 6.80 | 11 | 10.58 | 0.121 | 0.277 | (B) BRAF Class 3 Melanoma |
| FANCC | 9q22.32 | 11 | 1.42 | 4 | 3.42 | 0.122 | 0.279 | (B) BRAF Class 3 Melanoma |
| PGBD3 | 10q11.23 | 3 | 0.25 | 2 | 1.17 | 0.123 | 0.279 | (B) BRAF Class 3 Melanoma |
| RECQL | 12p12.1 | 0 | 0.00 | 1 | 4.17 | 0.123 | 0.28 | (B) BRAF Class 3 Melanoma |
| TSHR | 14q31.1 | 16 | 2.20 | 5 | 4.63 | 0.123 | 0.28 | (B) BRAF Class 3 Melanoma |
| MAF | 16q23.2 | 0 | 0.00 | 1 | 2.94 | 0.125 | 0.281 | (B) BRAF Class 3 Melanoma |
| SRY | Yp11.2 | 0 | 0.00 | 1 | 3.45 | 0.125 | 0.281 | (B) BRAF Class 3 Melanoma |
| RNF8 | 6p21.2 | 0 | 0.00 | 1 | 3.45 | 0.126 | 0.281 | (B) BRAF Class 3 Melanoma |
| INHBE | 12q13.3 | 0 | 0.00 | 1 | 0.58 | 0.126 | 0.281 | (B) BRAF Class 3 Melanoma |
| SNTG1 | 8q11.21 | 0 | 0.00 | 1 | 0.58 | 0.126 | 0.281 | (B) BRAF Class 3 Melanoma |
| TRBC2 | 7q34 | 0 | 0.00 | 1 | 0.58 | 0.126 | 0.281 | (B) BRAF Class 3 Melanoma |
| TAF1 | Xq13.1 | 10 | 4.24 | 3 | 11.54 | 0.127 | 0.281 | (B) BRAF Class 3 Melanoma |
| EPHB1 | 3q22.2 | 28 | 5.19 | 7 | 9.21 | 0.127 | 0.281 | (B) BRAF Class 3 Melanoma |
| IRS2 | 13q34 | 28 | 5.19 | 7 | 9.21 | 0.127 | 0.281 | (B) BRAF Class 3 Melanoma |
| NBN | 8q21.3 | 16 | 1.99 | 5 | 4.13 | 0.127 | 0.281 | (B) BRAF Class 3 Melanoma |
| LRIG3 | 12q14.1 | 1 | 4.00 | 0 | 0.00 | 0.128 | 0.281 | (A) BRAF Class 1 Melanoma |
| RBM10 | Xp11.3 | 17 | 2.40 | 5 | 4.95 | 0.128 | 0.283 | (B) BRAF Class 3 Melanoma |
| TRRAP | 7q22.1 | 0 | 0.00 | 1 | 6.67 | 0.132 | 0.289 | (B) BRAF Class 3 Melanoma |
| GNB1 | 1p36.33 | 1 | 3.85 | 0 | 0.00 | 0.132 | 0.289 | (A) BRAF Class 1 Melanoma |
| PTPN1 | 20q13.13 | 0 | 0.00 | 1 | 11.11 | 0.132 | 0.289 | (B) BRAF Class 3 Melanoma |
| BCL2 | 18q21.33 | 0 | 0.00 | 1 | 0.85 | 0.133 | 0.289 | (B) BRAF Class 3 Melanoma |
| CDK2 | 12q13.2 | 0 | 0.00 | 1 | 6.25 | 0.133 | 0.29 | (B) BRAF Class 3 Melanoma |
| AKT1 | 14q32.33 | 17 | 1.44 | 5 | 2.92 | 0.134 | 0.29 | (B) BRAF Class 3 Melanoma |
| FGFR1 | 8p11.23 | 17 | 1.44 | 5 | 2.92 | 0.134 | 0.29 | (B) BRAF Class 3 Melanoma |
| DNMT1 | 19p13.2 | 14 | 3.13 | 4 | 7.02 | 0.135 | 0.29 | (B) BRAF Class 3 Melanoma |
| EWSR1 | 22q12.2 | 7 | 1.34 | 3 | 3.75 | 0.135 | 0.29 | (B) BRAF Class 3 Melanoma |
| SH2D1A | Xq25 | 8 | 1.30 | 3 | 3.57 | 0.135 | 0.29 | (B) BRAF Class 3 Melanoma |
| EGFL7 | 9q34.3 | 4 | 0.98 | 2 | 3.92 | 0.136 | 0.291 | (B) BRAF Class 3 Melanoma |
| NRIP1 | 1q11.2-q21 | 1 | 2.50 | 1 | 33.33 | 0.136 | 0.291 | (B) BRAF Class 3 Melanoma |
| SMARCA2 | 9p24.3 | 0 | 0.00 | 1 | 11.11 | 0.136 | 0.291 | (B) BRAF Class 3 Melanoma |
| CDKN2A | 9p21.3 | 298 | 26.42 | 50 | 30.86 | 0.137 | 0.291 | (B) BRAF Class 3 Melanoma |
| DDIT3 | 12q13.3 | 0 | 0.00 | 1 | 7.69 | 0.138 | 0.292 | (B) BRAF Class 3 Melanoma |

| | | | | | | | | |
|---|---|---|---|---|---|---|---|---|
| MAML2 | 11q21 | 0 | 0.00 | 1 | 7.69 | 0.138 | 0.292 | (B) BRAF Class 3 Melanoma |
| PDGFB | 22q13.1 | 0 | 0.00 | 1 | 7.69 | 0.138 | 0.292 | (B) BRAF Class 3 Melanoma |
| RTEL1 | 20q13.33 | 8 | 4.68 | 3 | 12.50 | 0.139 | 0.294 | (B) BRAF Class 3 Melanoma |
| CRTC1 | 19p13.11 | 11 | 3.94 | 4 | 8.89 | 0.14 | 0.294 | (B) BRAF Class 3 Melanoma |
| EPHA2 | 1p36.13 | 0 | 0.00 | 1 | 5.88 | 0.14 | 0.295 | (B) BRAF Class 3 Melanoma |
| HIF1A | 14q23.2 | 0 | 0.00 | 1 | 7.69 | 0.141 | 0.296 | (B) BRAF Class 3 Melanoma |
| PRSS1 | 7q34 | 17 | 6.37 | 5 | 12.50 | 0.142 | 0.296 | (B) BRAF Class 3 Melanoma |
| NCOA3 | 20q13.12 | 19 | 4.26 | 5 | 8.33 | 0.142 | 0.297 | (B) BRAF Class 3 Melanoma |
| ASH2L | 8p11.23 | 0 | 0.00 | 1 | 11.11 | 0.143 | 0.297 | (B) BRAF Class 3 Melanoma |
| LMO2 | 11p13 | 3 | 1.18 | 2 | 4.88 | 0.144 | 0.298 | (B) BRAF Class 3 Melanoma |
| LEF1 | 4q25 | 0 | 0.00 | 1 | 8.33 | 0.145 | 0.299 | (B) BRAF Class 3 Melanoma |
| ROBO1 | 3p12.3 | 3 | 5.56 | 2 | 22.22 | 0.145 | 0.3 | (B) BRAF Class 3 Melanoma |
| PTCH2 | 1p34.1 | 3 | 2.91 | 2 | 11.76 | 0.146 | 0.301 | (B) BRAF Class 3 Melanoma |
| MLH1 | 3p22.2 | 18 | 1.62 | 5 | 3.16 | 0.147 | 0.303 | (B) BRAF Class 3 Melanoma |
| LMO3 | 12p12.3 | 0 | 0.00 | 1 | 8.33 | 0.148 | 0.303 | (B) BRAF Class 3 Melanoma |
| ZNF708 | 19p12 | 0 | 0.00 | 1 | 8.33 | 0.148 | 0.303 | (B) BRAF Class 3 Melanoma |
| RAD50 | 5q31.1 | 23 | 3.08 | 6 | 5.56 | 0.148 | 0.303 | (B) BRAF Class 3 Melanoma |
| INSR | 19p13.2 | 20 | 4.22 | 5 | 8.06 | 0.151 | 0.307 | (B) BRAF Class 3 Melanoma |
| BCOR | Xp11.4 | 38 | 4.92 | 9 | 7.69 | 0.152 | 0.309 | (B) BRAF Class 3 Melanoma |
| EGR1 | 5q31.2 | 0 | 0.00 | 1 | 14.29 | 0.152 | 0.309 | (B) BRAF Class 3 Melanoma |
| FYN | 6q21 | 9 | 2.07 | 3 | 5.26 | 0.153 | 0.31 | (B) BRAF Class 3 Melanoma |
| INSIG1 | 7q36.3 | 0 | 0.00 | 1 | 9.09 | 0.155 | 0.312 | (B) BRAF Class 3 Melanoma |
| NPRL3 | 16p13.3 | 0 | 0.00 | 1 | 9.09 | 0.155 | 0.312 | (B) BRAF Class 3 Melanoma |
| NAB2 | 12q13.3 | 1 | 0.63 | 1 | 7.14 | 0.156 | 0.313 | (B) BRAF Class 3 Melanoma |
| FUBP1 | 1p31.1 | 4 | 0.76 | 2 | 2.78 | 0.156 | 0.313 | (B) BRAF Class 3 Melanoma |
| CD79A | 19q13.2 | 4 | 0.76 | 2 | 2.74 | 0.158 | 0.313 | (B) BRAF Class 3 Melanoma |
| DGKH | 13q14.11 | 0 | 0.00 | 1 | 16.67 | 0.158 | 0.313 | (B) BRAF Class 3 Melanoma |
| ESPL1 | 12q13.13 | 0 | 0.00 | 1 | 16.67 | 0.158 | 0.313 | (B) BRAF Class 3 Melanoma |
| OR5L1 | 11q12.1 | 0 | 0.00 | 1 | 16.67 | 0.158 | 0.313 | (B) BRAF Class 3 Melanoma |
| TRIM28 | 19q13.43 | 0 | 0.00 | 1 | 16.67 | 0.158 | 0.313 | (B) BRAF Class 3 Melanoma |
| TRIM33 | 1p13.2 | 1 | 2.22 | 1 | 25.00 | 0.158 | 0.313 | (B) BRAF Class 3 Melanoma |
| SMYD3 | 1q44 | 4 | 2.34 | 2 | 8.33 | 0.16 | 0.316 | (B) BRAF Class 3 Melanoma |
| TEK | 9p21.2 | 19 | 7.51 | 5 | 13.89 | 0.163 | 0.321 | (B) BRAF Class 3 Melanoma |
| PAK3 | Xq23 | 9 | 3.23 | 3 | 7.89 | 0.163 | 0.321 | (B) BRAF Class 3 Melanoma |
| NUTM1 | 15q14 | 17 | 7.42 | 4 | 14.81 | 0.167 | 0.327 | (B) BRAF Class 3 Melanoma |
| EXO1 | 1q43 | 4 | 1.99 | 2 | 6.90 | 0.167 | 0.327 | (B) BRAF Class 3 Melanoma |
| INPP4B | 4q31.21 | 9 | 1.69 | 3 | 4.11 | 0.168 | 0.328 | (B) BRAF Class 3 Melanoma |
| CADM2 | 3p12.1 | 3 | 5.00 | 2 | 18.18 | 0.169 | 0.328 | (B) BRAF Class 3 Melanoma |
| HDAC9 | 7p21.1 | 3 | 9.38 | 2 | 33.33 | 0.169 | 0.328 | (B) BRAF Class 3 Melanoma |
| INSRR | 1q23.1 | 4 | 0.34 | 2 | 1.17 | 0.169 | 0.328 | (B) BRAF Class 3 Melanoma |
| KIF1B | 1p36.22 | 14 | 6.97 | 4 | 13.79 | 0.176 | 0.342 | (B) BRAF Class 3 Melanoma |
| PMS2 | 7p22.1 | 19 | 2.39 | 5 | 4.35 | 0.177 | 0.342 | (B) BRAF Class 3 Melanoma |
| CBFA2T3 | 16q24.3 | 9 | 4.43 | 3 | 10.34 | 0.178 | 0.343 | (B) BRAF Class 3 Melanoma |
| SOX2 | 3q26.33 | 4 | 0.52 | 2 | 1.71 | 0.18 | 0.348 | (B) BRAF Class 3 Melanoma |
| CD79B | 17q23.3 | 4 | 0.52 | 2 | 1.71 | 0.181 | 0.349 | (B) BRAF Class 3 Melanoma |
| SUZ12 | 17q11.2 | 9 | 1.30 | 3 | 3.03 | 0.182 | 0.35 | (B) BRAF Class 3 Melanoma |
| STAT3 | 17q21.2 | 14 | 1.77 | 4 | 3.48 | 0.185 | 0.354 | (B) BRAF Class 3 Melanoma |
| KLHL6 | 3q27.1 | 10 | 3.40 | 3 | 7.69 | 0.186 | 0.354 | (B) BRAF Class 3 Melanoma |

| | | | | | | | | |
|---|---|---|---|---|---|---|---|---|
| IRF2 | 4q35.1 | 1 | 0.42 | 1 | 3.85 | 0.187 | 0.356 | (B) BRAF Class 3 Melanoma |
| DDB2 | 11p11.2 | 4 | 1.41 | 2 | 4.44 | 0.192 | 0.365 | (B) BRAF Class 3 Melanoma |
| MAP3K9 | 14q24.2 | 4 | 6.45 | 2 | 20.00 | 0.192 | 0.365 | (B) BRAF Class 3 Melanoma |
| PIK3C3 | 18q12.3 | 5 | 1.07 | 2 | 3.17 | 0.196 | 0.372 | (B) BRAF Class 3 Melanoma |
| PAK1 | 1q13.5-q14 | 5 | 1.13 | 2 | 3.33 | 0.198 | 0.374 | (B) BRAF Class 3 Melanoma |
| AURKA | 20q13.2 | 9 | 1.16 | 3 | 2.56 | 0.201 | 0.38 | (B) BRAF Class 3 Melanoma |
| CCND1 | 11q13.3 | 9 | 1.12 | 3 | 2.46 | 0.203 | 0.382 | (B) BRAF Class 3 Melanoma |
| PAX8 | 2q14.1 | 6 | 2.64 | 2 | 7.41 | 0.204 | 0.384 | (B) BRAF Class 3 Melanoma |
| ARID5B | 10q21.2 | 22 | 4.95 | 5 | 8.33 | 0.208 | 0.39 | (B) BRAF Class 3 Melanoma |
| STAG1 | 3q22.3 | 6 | 2.79 | 2 | 7.69 | 0.209 | 0.391 | (B) BRAF Class 3 Melanoma |
| GATA1 | Xp11.23 | 5 | 0.94 | 2 | 2.67 | 0.211 | 0.393 | (B) BRAF Class 3 Melanoma |
| QKI | 6q26 | 5 | 1.08 | 2 | 3.03 | 0.213 | 0.396 | (B) BRAF Class 3 Melanoma |
| KCNQ1 | 1p15.5-p15 | 10 | 4.98 | 3 | 10.34 | 0.216 | 0.401 | (B) BRAF Class 3 Melanoma |
| RAD51 | 15q15.1 | 5 | 0.66 | 2 | 1.83 | 0.218 | 0.404 | (B) BRAF Class 3 Melanoma |
| MYBL1 | 8q13.1 | 4 | 1.31 | 2 | 3.77 | 0.218 | 0.404 | (B) BRAF Class 3 Melanoma |
| ETV4 | 17q21.31 | 5 | 1.02 | 2 | 2.82 | 0.219 | 0.404 | (B) BRAF Class 3 Melanoma |
| RAD51D | 17q12 | 5 | 0.67 | 2 | 1.83 | 0.219 | 0.404 | (B) BRAF Class 3 Melanoma |
| SH2B3 | 12q24.12 | 20 | 2.67 | 5 | 4.42 | 0.222 | 0.408 | (B) BRAF Class 3 Melanoma |
| DNAJB1 | 19p13.12 | 1 | 0.22 | 1 | 1.67 | 0.223 | 0.41 | (B) BRAF Class 3 Melanoma |
| CD274 | 9p24.1 | 5 | 0.66 | 2 | 1.77 | 0.226 | 0.414 | (B) BRAF Class 3 Melanoma |
| SDHA | 5p15.33 | 27 | 3.63 | 6 | 5.61 | 0.226 | 0.414 | (B) BRAF Class 3 Melanoma |
| SMO | 7q32.1 | 39 | 3.51 | 8 | 5.03 | 0.227 | 0.414 | (B) BRAF Class 3 Melanoma |
| WWTR1 | 3q25.1 | 1 | 0.51 | 1 | 3.70 | 0.229 | 0.417 | (B) BRAF Class 3 Melanoma |
| RAD52 | 12p13.33 | 1 | 0.15 | 1 | 1.09 | 0.231 | 0.419 | (B) BRAF Class 3 Melanoma |
| ERF | 19q13.2 | 1 | 0.58 | 1 | 4.17 | 0.232 | 0.419 | (B) BRAF Class 3 Melanoma |
| SESN3 | 11q21 | 1 | 0.58 | 1 | 4.17 | 0.232 | 0.419 | (B) BRAF Class 3 Melanoma |
| AXIN1 | 16p13.3 | 11 | 2.09 | 3 | 4.17 | 0.232 | 0.419 | (B) BRAF Class 3 Melanoma |
| MYD88 | 3p22.2 | 10 | 1.23 | 3 | 2.46 | 0.234 | 0.422 | (B) BRAF Class 3 Melanoma |
| SDHAF2 | 11q12.2 | 1 | 0.15 | 1 | 1.02 | 0.235 | 0.424 | (B) BRAF Class 3 Melanoma |
| CAMK1 | 3p25.3 | 1 | 0.08 | 1 | 0.58 | 0.237 | 0.424 | (B) BRAF Class 3 Melanoma |
| ENG | 9q34.11 | 1 | 0.50 | 1 | 3.45 | 0.237 | 0.424 | (B) BRAF Class 3 Melanoma |
| KLF2 | 19p13.11 | 1 | 0.50 | 1 | 3.45 | 0.237 | 0.424 | (B) BRAF Class 3 Melanoma |
| NT5C2 | q24.32-q24 | 5 | 1.84 | 2 | 4.76 | 0.238 | 0.424 | (B) BRAF Class 3 Melanoma |
| CRKL | 22q11.21 | 10 | 1.29 | 3 | 2.56 | 0.238 | 0.424 | (B) BRAF Class 3 Melanoma |
| RARA | 17q21.2 | 10 | 1.29 | 3 | 2.56 | 0.238 | 0.424 | (B) BRAF Class 3 Melanoma |
| PHF6 | Xq26.2 | 1 | 0.21 | 1 | 1.45 | 0.239 | 0.425 | (B) BRAF Class 3 Melanoma |
| HNRNPK | 9q21.32 | 2 | 1.06 | 1 | 5.56 | 0.241 | 0.427 | (B) BRAF Class 3 Melanoma |
| PRKCI | 3q26.2 | 16 | 2.56 | 4 | 4.40 | 0.241 | 0.427 | (B) BRAF Class 3 Melanoma |
| FGF23 | 12p13.32 | 6 | 2.08 | 2 | 5.13 | 0.244 | 0.43 | (B) BRAF Class 3 Melanoma |
| PLAG1 | 8q12.1 | 1 | 1.16 | 1 | 7.69 | 0.247 | 0.434 | (B) BRAF Class 3 Melanoma |
| SOX10 | 22q13.1 | 6 | 2.16 | 2 | 5.26 | 0.248 | 0.436 | (B) BRAF Class 3 Melanoma |
| PRPF8 | 17p13.3 | 1 | 1.04 | 1 | 6.67 | 0.253 | 0.444 | (B) BRAF Class 3 Melanoma |
| MAP3K13 | 3q27.2 | 12 | 2.54 | 3 | 4.76 | 0.254 | 0.444 | (B) BRAF Class 3 Melanoma |
| BUB1B | 15q15.1 | 10 | 3.52 | 3 | 6.67 | 0.256 | 0.448 | (B) BRAF Class 3 Melanoma |
| AGO2 | 8q24.3 | 6 | 3.51 | 2 | 8.33 | 0.256 | 0.448 | (B) BRAF Class 3 Melanoma |
| NUP93 | 16q13 | 6 | 1.17 | 2 | 2.78 | 0.258 | 0.448 | (B) BRAF Class 3 Melanoma |
| CDKN1C | 11p15.4 | 1 | 0.36 | 1 | 2.27 | 0.258 | 0.448 | (B) BRAF Class 3 Melanoma |
| CASR | q13.33-q21 | 3 | 7.50 | 1 | 33.33 | 0.259 | 0.45 | (B) BRAF Class 3 Melanoma |

| | | | | | | | | |
|---|---|---|---|---|---|---|---|---|
| PAXIP1 | 7q36.2 | 6 | 2.99 | 2 | 6.90 | 0.266 | 0.461 | (B) BRAF Class 3 Melanoma |
| FH | 1q43 | 11 | 1.43 | 3 | 2.63 | 0.267 | 0.462 | (B) BRAF Class 3 Melanoma |
| HDAC4 | 2q37.3 | 1 | 1.41 | 1 | 8.33 | 0.27 | 0.465 | (B) BRAF Class 3 Melanoma |
| CDKN2B | 9p21.3 | 113 | 14.04 | 20 | 16.53 | 0.272 | 0.469 | (B) BRAF Class 3 Melanoma |
| PRKN | 6q26 | 34 | 4.49 | 7 | 6.19 | 0.275 | 0.472 | (B) BRAF Class 3 Melanoma |
| BTK | Xq22.1 | 12 | 2.09 | 3 | 3.75 | 0.275 | 0.472 | (B) BRAF Class 3 Melanoma |
| CDK1 | 10q21.2 | 1 | 1.45 | 1 | 8.33 | 0.276 | 0.472 | (B) BRAF Class 3 Melanoma |
| PRAME | 22q11.22 | 1 | 1.45 | 1 | 8.33 | 0.276 | 0.472 | (B) BRAF Class 3 Melanoma |
| NFKBIA | 14q13.2 | 9 | 1.19 | 0 | 0.00 | 0.281 | 0.48 | (A) BRAF Class 1 Melanoma |
| SOCS1 | 16p13.13 | 6 | 0.78 | 2 | 1.71 | 0.284 | 0.483 | (B) BRAF Class 3 Melanoma |
| DHX15 | 4p15.2 | 3 | 1.89 | 1 | 7.14 | 0.289 | 0.49 | (B) BRAF Class 3 Melanoma |
| ESR2 | 4q23.2-q23 | 1 | 0.93 | 1 | 5.00 | 0.289 | 0.49 | (B) BRAF Class 3 Melanoma |
| PRSS8 | 16p11.2 | 2 | 0.86 | 1 | 3.57 | 0.29 | 0.49 | (B) BRAF Class 3 Melanoma |
| RNF43 | 17q22 | 12 | 1.59 | 3 | 2.75 | 0.291 | 0.491 | (B) BRAF Class 3 Melanoma |
| CTLA4 | 2q33.2 | 7 | 1.18 | 2 | 2.50 | 0.291 | 0.491 | (B) BRAF Class 3 Melanoma |
| HIST1H3G | 6p22.2 | 2 | 0.51 | 1 | 2.08 | 0.293 | 0.492 | (B) BRAF Class 3 Melanoma |
| HIST1H3J | 6p22.1 | 2 | 0.51 | 1 | 2.08 | 0.293 | 0.492 | (B) BRAF Class 3 Melanoma |
| KMT2C | 7q36.1 | 76 | 14.93 | 12 | 18.18 | 0.297 | 0.497 | (B) BRAF Class 3 Melanoma |
| PTPRS | 19p13.3 | 21 | 5.15 | 4 | 7.84 | 0.298 | 0.497 | (B) BRAF Class 3 Melanoma |
| PML | 15q24.1 | 9 | 3.86 | 0 | 0.00 | 0.298 | 0.497 | (A) BRAF Class 1 Melanoma |
| INHA | 2q35 | 8 | 2.04 | 2 | 4.17 | 0.298 | 0.497 | (B) BRAF Class 3 Melanoma |
| ICOSLG | 21q22.3 | 2 | 0.49 | 1 | 1.96 | 0.298 | 0.497 | (B) BRAF Class 3 Melanoma |
| MYC | 8q24.21 | 23 | 2.84 | 5 | 4.10 | 0.299 | 0.497 | (B) BRAF Class 3 Melanoma |
| CHD2 | 15q26.1 | 14 | 5.30 | 3 | 8.82 | 0.304 | 0.505 | (B) BRAF Class 3 Melanoma |
| INPPL1 | 11q13.4 | 7 | 4.09 | 2 | 8.33 | 0.306 | 0.507 | (B) BRAF Class 3 Melanoma |
| SHQ1 | 3p13 | 8 | 1.96 | 2 | 3.92 | 0.307 | 0.507 | (B) BRAF Class 3 Melanoma |
| PRDM1 | 6q21 | 53 | 6.86 | 10 | 8.55 | 0.308 | 0.507 | (B) BRAF Class 3 Melanoma |
| IKBKE | 1q32.1 | 7 | 1.30 | 2 | 2.63 | 0.308 | 0.507 | (B) BRAF Class 3 Melanoma |
| BRD3 | 9q34.2 | 7 | 3.00 | 2 | 6.06 | 0.309 | 0.509 | (B) BRAF Class 3 Melanoma |
| EPHB6 | 7q34 | 2 | 2.17 | 1 | 8.33 | 0.31 | 0.509 | (B) BRAF Class 3 Melanoma |
| HIST1H3B | 6p22.2 | 7 | 1.01 | 2 | 2.00 | 0.315 | 0.516 | (B) BRAF Class 3 Melanoma |
| CTNNA1 | 5q31.2 | 7 | 1.48 | 2 | 2.94 | 0.315 | 0.516 | (B) BRAF Class 3 Melanoma |
| PNKP | 19q13.33 | 7 | 3.48 | 2 | 6.90 | 0.317 | 0.516 | (B) BRAF Class 3 Melanoma |
| PDK1 | 2q31.1 | 2 | 0.69 | 1 | 2.56 | 0.317 | 0.516 | (B) BRAF Class 3 Melanoma |
| DRCS8-MEF2 | 19p13.11 | 7 | 0.59 | 2 | 1.17 | 0.317 | 0.516 | (B) BRAF Class 3 Melanoma |
| CHEK2 | 22q12.1 | 18 | 2.23 | 4 | 3.31 | 0.319 | 0.518 | (B) BRAF Class 3 Melanoma |
| NAV3 | 12q21.2 | 6 | 11.11 | 2 | 22.22 | 0.32 | 0.518 | (B) BRAF Class 3 Melanoma |
| FOXA1 | 14q21.1 | 17 | 2.60 | 1 | 1.08 | 0.321 | 0.519 | (A) BRAF Class 1 Melanoma |
| IGF2 | 11p15.5 | 7 | 0.99 | 2 | 1.92 | 0.323 | 0.522 | (B) BRAF Class 3 Melanoma |
| MAPK1 | 22q11.22 | 7 | 0.93 | 2 | 1.82 | 0.324 | 0.522 | (B) BRAF Class 3 Melanoma |
| RAC2 | 22q13.1 | 2 | 1.17 | 1 | 4.17 | 0.327 | 0.525 | (B) BRAF Class 3 Melanoma |
| RXRA | 9q34.2 | 2 | 1.17 | 1 | 4.17 | 0.327 | 0.525 | (B) BRAF Class 3 Melanoma |
| TAP1 | 6p21.32 | 2 | 1.17 | 1 | 4.17 | 0.327 | 0.525 | (B) BRAF Class 3 Melanoma |
| SMC1A | Xp11.22 | 2 | 0.73 | 1 | 2.56 | 0.329 | 0.526 | (B) BRAF Class 3 Melanoma |
| WT1 | 11p13 | 24 | 3.07 | 5 | 4.24 | 0.329 | 0.526 | (B) BRAF Class 3 Melanoma |
| TAL2 | 9q31.2 | 2 | 0.99 | 1 | 3.45 | 0.331 | 0.527 | (B) BRAF Class 3 Melanoma |
| TLX3 | 5q35.1 | 2 | 0.99 | 1 | 3.45 | 0.331 | 0.527 | (B) BRAF Class 3 Melanoma |
| CDK6 | 7q21.2 | 7 | 0.87 | 2 | 1.65 | 0.331 | 0.527 | (B) BRAF Class 3 Melanoma |

| | | | | | | | | |
|---|---|---|---|---|---|---|---|---|
| ZRSR2 | Xp22.2 | 8 | 1.15 | 0 | 0.00 | 0.332 | 0.527 | (A) BRAF Class 1 Melanoma |
| PNRC1 | 6q15 | 2 | 0.39 | 1 | 1.35 | 0.333 | 0.528 | (B) BRAF Class 3 Melanoma |
| FAAP20 | 1p36.33 | 2 | 1.00 | 1 | 3.45 | 0.334 | 0.528 | (B) BRAF Class 3 Melanoma |
| SMAD2 | 18q21.1 | 7 | 0.91 | 2 | 1.71 | 0.336 | 0.53 | (B) BRAF Class 3 Melanoma |
| PARP1 | 1q42.12 | 8 | 1.61 | 2 | 2.99 | 0.336 | 0.53 | (B) BRAF Class 3 Melanoma |
| NTHL1 | 16p13.3 | 2 | 0.53 | 1 | 1.79 | 0.341 | 0.537 | (B) BRAF Class 3 Melanoma |
| CYSLTR2 | 13q14.2 | 2 | 0.96 | 1 | 3.23 | 0.342 | 0.537 | (B) BRAF Class 3 Melanoma |
| BABAM1 | 19p13.11 | 2 | 0.56 | 1 | 1.89 | 0.343 | 0.537 | (B) BRAF Class 3 Melanoma |
| CCND3 | 6p21.1 | 2 | 0.25 | 1 | 0.83 | 0.343 | 0.537 | (B) BRAF Class 3 Melanoma |
| KNSTRN | 15q15.1 | 15 | 6.20 | 1 | 2.70 | 0.346 | 0.54 | (A) BRAF Class 1 Melanoma |
| PIK3R1 | 5q13.1 | 19 | 2.34 | 4 | 3.31 | 0.349 | 0.544 | (B) BRAF Class 3 Melanoma |
| AXIN2 | 17q24.1 | 20 | 3.11 | 4 | 4.35 | 0.352 | 0.548 | (B) BRAF Class 3 Melanoma |
| STK19 | 6p21.33 | 8 | 4.68 | 2 | 8.33 | 0.355 | 0.551 | (B) BRAF Class 3 Melanoma |
| SS18 | 18q11.2 | 2 | 0.77 | 1 | 2.44 | 0.356 | 0.551 | (B) BRAF Class 3 Melanoma |
| RASA1 | 5q14.3 | 14 | 2.23 | 3 | 3.37 | 0.356 | 0.551 | (B) BRAF Class 3 Melanoma |
| BAP1 | 3p21.1 | 30 | 3.71 | 3 | 2.48 | 0.357 | 0.552 | (A) BRAF Class 1 Melanoma |
| SBDS | 7q11.21 | 2 | 0.70 | 1 | 2.22 | 0.358 | 0.552 | (B) BRAF Class 3 Melanoma |
| CHD4 | 12p13.31 | 9 | 3.41 | 2 | 5.88 | 0.363 | 0.56 | (B) BRAF Class 3 Melanoma |
| BCR | 22q11.23 | 10 | 3.92 | 2 | 6.67 | 0.366 | 0.564 | (B) BRAF Class 3 Melanoma |
| HIST1H3I | 6p22.1 | 3 | 0.76 | 1 | 2.08 | 0.37 | 0.569 | (B) BRAF Class 3 Melanoma |
| GRM8 | 7q31.33 | 4 | 9.30 | 1 | 25.00 | 0.372 | 0.571 | (B) BRAF Class 3 Melanoma |
| U2AF1 | 21q22.3 | 8 | 1.00 | 2 | 1.69 | 0.374 | 0.572 | (B) BRAF Class 3 Melanoma |
| STAT5A | 17q21.2 | 3 | 0.73 | 1 | 1.96 | 0.375 | 0.572 | (B) BRAF Class 3 Melanoma |
| SOS2 | 14q21.3 | 2 | 3.70 | 1 | 11.11 | 0.375 | 0.572 | (B) BRAF Class 3 Melanoma |
| CRLF2 | 2.33 and Yp | 14 | 1.87 | 3 | 2.73 | 0.376 | 0.572 | (B) BRAF Class 3 Melanoma |
| TCF7L1 | 2p11.2 | 2 | 2.13 | 1 | 6.25 | 0.379 | 0.576 | (B) BRAF Class 3 Melanoma |
| CDK5 | 7q36.1 | 2 | 2.90 | 1 | 8.33 | 0.386 | 0.584 | (B) BRAF Class 3 Melanoma |
| FKBP9 | 7p14.3 | 2 | 2.90 | 1 | 8.33 | 0.386 | 0.584 | (B) BRAF Class 3 Melanoma |
| GSTM5 | 1p13.3 | 2 | 2.90 | 1 | 8.33 | 0.386 | 0.584 | (B) BRAF Class 3 Melanoma |
| FUS | 16p11.2 | 7 | 2.26 | 2 | 3.77 | 0.387 | 0.585 | (B) BRAF Class 3 Melanoma |
| WRN | 8p12 | 18 | 5.63 | 4 | 7.41 | 0.395 | 0.596 | (B) BRAF Class 3 Melanoma |
| B2M | 15q21.1 | 22 | 3.30 | 2 | 2.08 | 0.399 | 0.601 | (A) BRAF Class 1 Melanoma |
| CALR | 19p13.13 | 7 | 1.07 | 0 | 0.00 | 0.4 | 0.602 | (A) BRAF Class 1 Melanoma |
| KCNIP1 | 5q35.1 | 2 | 3.33 | 1 | 9.09 | 0.401 | 0.602 | (B) BRAF Class 3 Melanoma |
| CDK4 | 12q14.1 | 28 | 3.47 | 3 | 2.46 | 0.403 | 0.604 | (A) BRAF Class 1 Melanoma |
| EXT2 | 11p11.2 | 8 | 2.82 | 2 | 4.44 | 0.408 | 0.61 | (B) BRAF Class 3 Melanoma |
| MSI1 | 12q24.31 | 3 | 1.75 | 1 | 4.17 | 0.411 | 0.614 | (B) BRAF Class 3 Melanoma |
| TSHZ3 | 19q12 | 2 | 6.25 | 1 | 16.67 | 0.412 | 0.614 | (B) BRAF Class 3 Melanoma |
| MT-ND5 | - | 3 | 0.25 | 1 | 0.58 | 0.418 | 0.619 | (B) BRAF Class 3 Melanoma |
| DCLRE1C | 10p13 | 3 | 1.49 | 1 | 3.45 | 0.419 | 0.619 | (B) BRAF Class 3 Melanoma |
| GBA | 1q22 | 3 | 1.49 | 1 | 3.45 | 0.419 | 0.619 | (B) BRAF Class 3 Melanoma |
| HMBS | 11q23.3 | 3 | 1.49 | 1 | 3.45 | 0.419 | 0.619 | (B) BRAF Class 3 Melanoma |
| MUS81 | 11q13.1 | 3 | 1.49 | 1 | 3.45 | 0.419 | 0.619 | (B) BRAF Class 3 Melanoma |
| NEIL1 | 15q24.2 | 3 | 1.49 | 1 | 3.45 | 0.419 | 0.619 | (B) BRAF Class 3 Melanoma |
| POLH | 6p21.1 | 3 | 1.38 | 1 | 3.13 | 0.424 | 0.625 | (B) BRAF Class 3 Melanoma |
| MSH3 | 5q14.1 | 10 | 2.66 | 2 | 3.92 | 0.431 | 0.634 | (B) BRAF Class 3 Melanoma |
| AURKB | 17p13.1 | 9 | 1.16 | 2 | 1.71 | 0.435 | 0.638 | (B) BRAF Class 3 Melanoma |
| CD276 | 15q24.1 | 7 | 1.72 | 0 | 0.00 | 0.436 | 0.638 | (A) BRAF Class 1 Melanoma |

| | | | | | | | | |
|---|---|---|---|---|---|---|---|---|
| LATS1 | 6q25.1 | 7 | 1.72 | 0 | 0.00 | 0.436 | 0.638 | (A) BRAF Class 1 Melanoma |
| LIG4 | 13q33.3 | 6 | 2.99 | 0 | 0.00 | 0.441 | 0.645 | (A) BRAF Class 1 Melanoma |
| ELANE | 19p13.3 | 6 | 2.69 | 0 | 0.00 | 0.443 | 0.647 | (A) BRAF Class 1 Melanoma |
| FANCL | 2p16.1 | 6 | 1.27 | 0 | 0.00 | 0.444 | 0.647 | (A) BRAF Class 1 Melanoma |
| FOXL2 | 3q22.3 | 6 | 0.79 | 0 | 0.00 | 0.445 | 0.647 | (A) BRAF Class 1 Melanoma |
| RPS6KB2 | 11q13.2 | 4 | 0.98 | 1 | 1.96 | 0.447 | 0.647 | (B) BRAF Class 3 Melanoma |
| RYBP | 3p13 | 4 | 0.98 | 1 | 1.96 | 0.447 | 0.647 | (B) BRAF Class 3 Melanoma |
| CASP8 | 2q33.1 | 10 | 1.42 | 2 | 2.02 | 0.447 | 0.647 | (B) BRAF Class 3 Melanoma |
| MAP3K4 | 6q26 | 6 | 3.77 | 1 | 7.14 | 0.452 | 0.654 | (B) BRAF Class 3 Melanoma |
| PIM1 | 6p21.2 | 10 | 1.48 | 2 | 2.06 | 0.457 | 0.66 | (B) BRAF Class 3 Melanoma |
| IGF2R | 6q25.3 | 3 | 2.34 | 1 | 4.76 | 0.459 | 0.662 | (B) BRAF Class 3 Melanoma |
| ASXL2 | 2p23.3 | 25 | 5.63 | 4 | 6.67 | 0.461 | 0.663 | (B) BRAF Class 3 Melanoma |
| RAD54B | 8q22.1 | 10 | 4.98 | 2 | 6.90 | 0.462 | 0.663 | (B) BRAF Class 3 Melanoma |
| KLF4 | 9q31.2 | 17 | 2.63 | 3 | 3.26 | 0.464 | 0.666 | (B) BRAF Class 3 Melanoma |
| FGF10 | 5p12 | 6 | 2.08 | 0 | 0.00 | 0.465 | 0.666 | (A) BRAF Class 1 Melanoma |
| IDH1 | 2q34 | 41 | 3.47 | 5 | 2.92 | 0.466 | 0.666 | (A) BRAF Class 1 Melanoma |
| SIN3A | 15q24.2 | 3 | 5.56 | 1 | 11.11 | 0.469 | 0.67 | (B) BRAF Class 3 Melanoma |
| DMD | :p21.2-p21. | 3 | 4.35 | 1 | 8.33 | 0.48 | 0.684 | (B) BRAF Class 3 Melanoma |
| AKT2 | 19q13.2 | 12 | 1.48 | 1 | 0.83 | 0.481 | 0.684 | (A) BRAF Class 1 Melanoma |
| GABRA6 | 5q34 | 7 | 2.93 | 0 | 0.00 | 0.481 | 0.684 | (A) BRAF Class 1 Melanoma |
| NKX2-1 | 14q13.3 | 10 | 1.29 | 2 | 1.71 | 0.482 | 0.684 | (B) BRAF Class 3 Melanoma |
| XRCC5 | 2q35 | 4 | 1.97 | 1 | 3.45 | 0.49 | 0.695 | (B) BRAF Class 3 Melanoma |
| SLC25A13 | 7q21.3 | 4 | 1.99 | 1 | 3.45 | 0.494 | 0.698 | (B) BRAF Class 3 Melanoma |
| STAT6 | 12q13.3 | 9 | 2.90 | 2 | 3.77 | 0.495 | 0.699 | (B) BRAF Class 3 Melanoma |
| ID3 | 1p36.12 | 4 | 0.57 | 1 | 0.97 | 0.496 | 0.7 | (B) BRAF Class 3 Melanoma |
| RAB35 | 12q24.23 | 5 | 1.27 | 1 | 2.08 | 0.501 | 0.706 | (B) BRAF Class 3 Melanoma |
| SLC34A2 | 4p15.2 | 11 | 4.78 | 2 | 6.06 | 0.503 | 0.707 | (B) BRAF Class 3 Melanoma |
| ERCC4 | 16p13.12 | 18 | 2.47 | 3 | 2.86 | 0.505 | 0.709 | (B) BRAF Class 3 Melanoma |
| MBD4 | 3q21.3 | 5 | 2.49 | 0 | 0.00 | 0.506 | 0.71 | (A) BRAF Class 1 Melanoma |
| VTCN1 | 1p13.1-p12 | 5 | 1.23 | 1 | 1.96 | 0.509 | 0.711 | (B) BRAF Class 3 Melanoma |
| XIAP | Xq25 | 5 | 1.23 | 1 | 1.96 | 0.509 | 0.711 | (B) BRAF Class 3 Melanoma |
| MAP2K1 | 15q22.31 | 50 | 5.68 | 7 | 5.22 | 0.513 | 0.714 | (A) BRAF Class 1 Melanoma |
| DPYD | 1p21.3 | 4 | 3.81 | 1 | 6.25 | 0.514 | 0.714 | (B) BRAF Class 3 Melanoma |
| TNFRSF14 | 1p36.32 | 5 | 0.93 | 0 | 0.00 | 0.515 | 0.714 | (A) BRAF Class 1 Melanoma |
| ACVR1B | 12q13.13 | 5 | 1.82 | 1 | 2.86 | 0.516 | 0.714 | (B) BRAF Class 3 Melanoma |
| CTNNB1 | 3p22.1 | 67 | 5.66 | 10 | 5.85 | 0.516 | 0.714 | (B) BRAF Class 3 Melanoma |
| XRCC2 | 7q36.1 | 12 | 1.82 | 2 | 2.22 | 0.516 | 0.714 | (B) BRAF Class 3 Melanoma |
| ZMYM3 | Xq13.1 | 4 | 5.63 | 0 | 0.00 | 0.529 | 0.731 | (A) BRAF Class 1 Melanoma |
| CLDN18 | 3q22.3 | 4 | 7.41 | 0 | 0.00 | 0.531 | 0.732 | (A) BRAF Class 1 Melanoma |
| CYLD | 16q12.1 | 18 | 2.70 | 3 | 3.00 | 0.531 | 0.732 | (B) BRAF Class 3 Melanoma |
| JUN | 1p32.1 | 5 | 0.99 | 0 | 0.00 | 0.534 | 0.733 | (A) BRAF Class 1 Melanoma |
| FGF4 | 11q13.3 | 12 | 2.23 | 2 | 2.63 | 0.534 | 0.733 | (B) BRAF Class 3 Melanoma |
| EMSY | 11q13.5 | 5 | 1.73 | 1 | 2.56 | 0.535 | 0.733 | (B) BRAF Class 3 Melanoma |
| FGF6 | 12p13.32 | 5 | 1.73 | 1 | 2.56 | 0.535 | 0.733 | (B) BRAF Class 3 Melanoma |
| HLA-A | 6p22.1 | 5 | 1.12 | 0 | 0.00 | 0.537 | 0.734 | (A) BRAF Class 1 Melanoma |
| KDM6B | 17p13.1 | 4 | 5.41 | 1 | 8.33 | 0.537 | 0.734 | (B) BRAF Class 3 Melanoma |
| GREM1 | 15q13.3 | 5 | 0.74 | 1 | 1.06 | 0.542 | 0.739 | (B) BRAF Class 3 Melanoma |
| HIST1H1C | 6p22.2 | 5 | 1.18 | 0 | 0.00 | 0.548 | 0.746 | (A) BRAF Class 1 Melanoma |

| | | | | | | | | |
|---|---|---|---|---|---|---|---|---|
| TRAF3 | 14q32.32 | 4 | 1.53 | 0 | 0.00 | 0.549 | 0.746 | (A) BRAF Class 1 Melanoma |
| ETV5 | 3q27.2 | 12 | 2.46 | 2 | 2.82 | 0.549 | 0.746 | (B) BRAF Class 3 Melanoma |
| BRCC3 | Xq28 | 5 | 2.43 | 1 | 3.45 | 0.55 | 0.746 | (B) BRAF Class 3 Melanoma |
| FGF19 | 11q13.3 | 11 | 1.92 | 1 | 1.25 | 0.556 | 0.753 | (A) BRAF Class 1 Melanoma |
| FAH | 15q25.1 | 5 | 2.49 | 1 | 3.45 | 0.559 | 0.755 | (B) BRAF Class 3 Melanoma |
| MAPK3 | 16p11.2 | 5 | 1.27 | 0 | 0.00 | 0.56 | 0.756 | (A) BRAF Class 1 Melanoma |
| PLK2 | 5q11.2 | 6 | 1.47 | 1 | 1.96 | 0.564 | 0.76 | (B) BRAF Class 3 Melanoma |
| PPP6C | 9q33.3 | 37 | 8.49 | 5 | 8.62 | 0.565 | 0.76 | (B) BRAF Class 3 Melanoma |
| BCL2L1 | 20q11.21 | 5 | 0.66 | 1 | 0.88 | 0.568 | 0.763 | (B) BRAF Class 3 Melanoma |
| MAP2K4 | 17p12 | 10 | 1.24 | 1 | 0.83 | 0.569 | 0.763 | (A) BRAF Class 1 Melanoma |
| KEL | 7q34 | 22 | 9.21 | 2 | 7.69 | 0.574 | 0.768 | (A) BRAF Class 1 Melanoma |
| HFE | 6p22.2 | 4 | 1.99 | 0 | 0.00 | 0.581 | 0.777 | (A) BRAF Class 1 Melanoma |
| SDHC | 1q23.3 | 4 | 0.53 | 0 | 0.00 | 0.585 | 0.781 | (A) BRAF Class 1 Melanoma |
| RPA1 | 17p13.3 | 5 | 1.96 | 1 | 2.50 | 0.586 | 0.782 | (B) BRAF Class 3 Melanoma |
| CTCF | 16q22.1 | 13 | 1.80 | 2 | 1.90 | 0.588 | 0.783 | (B) BRAF Class 3 Melanoma |
| DUSP4 | 8p12 | 5 | 2.42 | 1 | 3.03 | 0.592 | 0.787 | (B) BRAF Class 3 Melanoma |
| ELF3 | 1q32.1 | 5 | 2.42 | 1 | 3.03 | 0.592 | 0.787 | (B) BRAF Class 3 Melanoma |
| HSD3B1 | 1p12 | 5 | 2.09 | 0 | 0.00 | 0.594 | 0.788 | (A) BRAF Class 1 Melanoma |
| CCN6 | 6q21 | 6 | 2.16 | 1 | 2.63 | 0.596 | 0.789 | (B) BRAF Class 3 Melanoma |
| DEPDC5 | 2q12.2-q12 | 3 | 5.00 | 0 | 0.00 | 0.599 | 0.791 | (A) BRAF Class 1 Melanoma |
| ARFRP1 | 20q13.33 | 4 | 1.39 | 0 | 0.00 | 0.599 | 0.791 | (A) BRAF Class 1 Melanoma |
| NKX3-1 | 8p21.2 | 4 | 0.68 | 0 | 0.00 | 0.601 | 0.792 | (A) BRAF Class 1 Melanoma |
| BIRC3 | 11q22.2 | 7 | 1.67 | 1 | 1.92 | 0.61 | 0.802 | (B) BRAF Class 3 Melanoma |
| NPM1 | 5q35.1 | 6 | 0.56 | 1 | 0.65 | 0.61 | 0.802 | (B) BRAF Class 3 Melanoma |
| INPP4A | 2q11.2 | 7 | 1.72 | 1 | 1.96 | 0.613 | 0.804 | (B) BRAF Class 3 Melanoma |
| DIS3L2 | 2q37.1 | 6 | 2.69 | 1 | 3.13 | 0.614 | 0.804 | (B) BRAF Class 3 Melanoma |
| MDM4 | 1q32.1 | 15 | 1.86 | 2 | 1.65 | 0.615 | 0.804 | (A) BRAF Class 1 Melanoma |
| HELQ | 4q21.23 | 6 | 2.99 | 1 | 3.45 | 0.616 | 0.804 | (B) BRAF Class 3 Melanoma |
| TDG | 12q23.3 | 6 | 2.99 | 1 | 3.45 | 0.616 | 0.804 | (B) BRAF Class 3 Melanoma |
| RHEB | 7q36.1 | 6 | 0.87 | 1 | 0.99 | 0.618 | 0.805 | (B) BRAF Class 3 Melanoma |
| PTPRK | 6q22.33 | 5 | 9.26 | 1 | 11.11 | 0.62 | 0.807 | (B) BRAF Class 3 Melanoma |
| SMC3 | 10q25.2 | 9 | 1.97 | 1 | 1.47 | 0.622 | 0.808 | (A) BRAF Class 1 Melanoma |
| IL10 | 1q32.1 | 4 | 0.98 | 0 | 0.00 | 0.623 | 0.808 | (A) BRAF Class 1 Melanoma |
| PIK3R3 | 1p34.1 | 4 | 0.98 | 0 | 0.00 | 0.623 | 0.808 | (A) BRAF Class 1 Melanoma |
| RANBP2 | 2q13 | 20 | 8.77 | 2 | 8.00 | 0.627 | 0.811 | (A) BRAF Class 1 Melanoma |
| GPS2 | 17p13.1 | 4 | 1.00 | 0 | 0.00 | 0.629 | 0.812 | (A) BRAF Class 1 Melanoma |
| HIST1H3E | 6p22.2 | 4 | 1.02 | 0 | 0.00 | 0.63 | 0.812 | (A) BRAF Class 1 Melanoma |
| ERCC6 | 10q11.23 | 9 | 4.48 | 1 | 3.45 | 0.634 | 0.815 | (A) BRAF Class 1 Melanoma |
| FAAP100 | 17q25.3 | 9 | 4.48 | 1 | 3.45 | 0.634 | 0.815 | (A) BRAF Class 1 Melanoma |
| HIST1H3C | 6p22.2 | 7 | 1.17 | 1 | 1.25 | 0.636 | 0.817 | (B) BRAF Class 3 Melanoma |
| MYH9 | 22q12.3 | 3 | 3.70 | 0 | 0.00 | 0.637 | 0.817 | (A) BRAF Class 1 Melanoma |
| JAZF1 | p15.2-p15. | 3 | 1.17 | 0 | 0.00 | 0.639 | 0.817 | (A) BRAF Class 1 Melanoma |
| FGF7 | 15q21.2 | 4 | 1.88 | 0 | 0.00 | 0.64 | 0.817 | (A) BRAF Class 1 Melanoma |
| RNF213 | 17q25.3 | 5 | 11.11 | 0 | 0.00 | 0.641 | 0.817 | (A) BRAF Class 1 Melanoma |
| ZNF703 | 8p11.23 | 7 | 2.42 | 1 | 2.56 | 0.641 | 0.817 | (B) BRAF Class 3 Melanoma |
| ZNF217 | 20q13.2 | 14 | 2.68 | 2 | 2.50 | 0.642 | 0.817 | (A) BRAF Class 1 Melanoma |
| SF1 | 11q13.1 | 3 | 3.13 | 0 | 0.00 | 0.644 | 0.82 | (A) BRAF Class 1 Melanoma |
| RPS6KA4 | 11q13.1 | 10 | 2.45 | 1 | 1.96 | 0.65 | 0.825 | (A) BRAF Class 1 Melanoma |

| | | | | | | | | |
|---|---|---|---|---|---|---|---|---|
| EIF1AX | Xp22.12 | 7 | 1.53 | 1 | 1.56 | 0.651 | 0.826 | (B) BRAF Class 3 Melanoma |
| BRINP3 | 1q31.1 | 5 | 10.64 | 0 | 0.00 | 0.653 | 0.827 | (A) BRAF Class 1 Melanoma |
| RELA | 11q13.1 | 6 | 2.36 | 1 | 2.44 | 0.653 | 0.827 | (B) BRAF Class 3 Melanoma |
| AKT3 | 1q43-q44 | 14 | 1.73 | 2 | 1.65 | 0.655 | 0.827 | (A) BRAF Class 1 Melanoma |
| YAP1 | 11q22.1 | 7 | 1.12 | 1 | 1.12 | 0.656 | 0.828 | (B) BRAF Class 3 Melanoma |
| VHL | 3p25.3 | 7 | 0.65 | 1 | 0.65 | 0.658 | 0.828 | (B) BRAF Class 3 Melanoma |
| ERRFI1 | 1p36.23 | 9 | 1.77 | 1 | 1.45 | 0.66 | 0.83 | (A) BRAF Class 1 Melanoma |
| H3F3A | 1q42.12 | 3 | 0.38 | 0 | 0.00 | 0.661 | 0.83 | (A) BRAF Class 1 Melanoma |
| CBFB | 16q22.1 | 3 | 0.42 | 0 | 0.00 | 0.663 | 0.832 | (A) BRAF Class 1 Melanoma |
| DKC1 | Xq28 | 3 | 1.49 | 0 | 0.00 | 0.666 | 0.834 | (A) BRAF Class 1 Melanoma |
| IKZF3 | [illegible]7q12-q21.[illegible] | 5 | 4.10 | 1 | 4.17 | 0.667 | 0.834 | (B) BRAF Class 3 Melanoma |
| PPP4R2 | 3p13 | 3 | 1.75 | 0 | 0.00 | 0.673 | 0.841 | (A) BRAF Class 1 Melanoma |
| FANCB | Xp22.2 | 8 | 3.64 | 1 | 3.13 | 0.68 | 0.848 | (A) BRAF Class 1 Melanoma |
| RAD51C | 17q22 | 8 | 1.07 | 1 | 0.92 | 0.681 | 0.848 | (A) BRAF Class 1 Melanoma |
| TOP2A | 17q21.2 | 9 | 3.41 | 1 | 2.94 | 0.681 | 0.848 | (A) BRAF Class 1 Melanoma |
| TMEM127 | 2q11.2 | 3 | 0.49 | 0 | 0.00 | 0.683 | 0.848 | (A) BRAF Class 1 Melanoma |
| KLLN | 10q23.31 | 8 | 3.98 | 1 | 3.45 | 0.683 | 0.848 | (A) BRAF Class 1 Melanoma |
| SMARCD1 | 12q13.12 | 3 | 0.65 | 0 | 0.00 | 0.685 | 0.849 | (A) BRAF Class 1 Melanoma |
| RAD54L | 1p34.1 | 3 | 0.63 | 0 | 0.00 | 0.687 | 0.85 | (A) BRAF Class 1 Melanoma |
| RHOA | 3p21.31 | 8 | 1.18 | 1 | 1.04 | 0.691 | 0.854 | (A) BRAF Class 1 Melanoma |
| MYOD1 | 11p15.1 | 3 | 0.70 | 0 | 0.00 | 0.701 | 0.865 | (A) BRAF Class 1 Melanoma |
| BCL2L11 | 2q13 | 3 | 0.73 | 0 | 0.00 | 0.703 | 0.867 | (A) BRAF Class 1 Melanoma |
| CENPA | 2p23.3 | 3 | 0.76 | 0 | 0.00 | 0.707 | 0.868 | (A) BRAF Class 1 Melanoma |
| EIF4E | 4q23 | 3 | 0.76 | 0 | 0.00 | 0.707 | 0.868 | (A) BRAF Class 1 Melanoma |
| HIST1H3D | 6p22.2 | 3 | 0.76 | 0 | 0.00 | 0.707 | 0.868 | (A) BRAF Class 1 Melanoma |
| ANKRD26 | 10p12.1 | 4 | 2.52 | 0 | 0.00 | 0.711 | 0.872 | (A) BRAF Class 1 Melanoma |
| NFE2L2 | 2q31.2 | 7 | 0.87 | 1 | 0.83 | 0.72 | 0.882 | (A) BRAF Class 1 Melanoma |
| LMO1 | 11p15.4 | 7 | 0.97 | 1 | 0.94 | 0.725 | 0.887 | (A) BRAF Class 1 Melanoma |
| HSP90AA1 | 14q32.31 | 3 | 1.26 | 0 | 0.00 | 0.732 | 0.891 | (A) BRAF Class 1 Melanoma |
| CHD1 | 5q15-q21.1 | 2 | 3.70 | 0 | 0.00 | 0.733 | 0.891 | (A) BRAF Class 1 Melanoma |
| TTYH1 | 19q13.42 | 2 | 3.70 | 0 | 0.00 | 0.733 | 0.891 | (A) BRAF Class 1 Melanoma |
| ZBTB20 | 3q13.31 | 2 | 3.70 | 0 | 0.00 | 0.733 | 0.891 | (A) BRAF Class 1 Melanoma |
| TRIM37 | 17q22 | 7 | 3.48 | 1 | 3.45 | 0.734 | 0.891 | (A) BRAF Class 1 Melanoma |
| XPA | 9q22.33 | 2 | 0.70 | 0 | 0.00 | 0.745 | 0.903 | (A) BRAF Class 1 Melanoma |
| XRCC3 | 14q32.33 | 2 | 0.78 | 0 | 0.00 | 0.747 | 0.905 | (A) BRAF Class 1 Melanoma |
| SDHD | 11q23.1 | 2 | 0.26 | 0 | 0.00 | 0.758 | 0.917 | (A) BRAF Class 1 Melanoma |
| EP400 | 12q24.33 | 3 | 6.98 | 0 | 0.00 | 0.761 | 0.918 | (A) BRAF Class 1 Melanoma |
| RPS6KA2 | 6q27 | 3 | 6.98 | 0 | 0.00 | 0.761 | 0.918 | (A) BRAF Class 1 Melanoma |
| OGG1 | 3p25.3 | 2 | 1.00 | 0 | 0.00 | 0.763 | 0.919 | (A) BRAF Class 1 Melanoma |
| H19 | 11p15.5 | 2 | 0.17 | 0 | 0.00 | 0.763 | 0.919 | (A) BRAF Class 1 Melanoma |
| CARM1 | 19p13.2 | 2 | 1.17 | 0 | 0.00 | 0.768 | 0.921 | (A) BRAF Class 1 Melanoma |
| NUF2 | 1q23.3 | 2 | 1.17 | 0 | 0.00 | 0.768 | 0.921 | (A) BRAF Class 1 Melanoma |
| SESN1 | 6q21 | 2 | 1.17 | 0 | 0.00 | 0.768 | 0.921 | (A) BRAF Class 1 Melanoma |
| PPP2R2A | 8p21.2 | 3 | 1.76 | 0 | 0.00 | 0.775 | 0.926 | (A) BRAF Class 1 Melanoma |
| FGF1 | 5q31.3 | 3 | 1.89 | 0 | 0.00 | 0.775 | 0.926 | (A) BRAF Class 1 Melanoma |
| FGF9 | 13q12.11 | 3 | 1.89 | 0 | 0.00 | 0.775 | 0.926 | (A) BRAF Class 1 Melanoma |
| MAX | 14q23.3 | 2 | 0.31 | 0 | 0.00 | 0.778 | 0.927 | (A) BRAF Class 1 Melanoma |
| LTK | 15q15.1 | 2 | 1.94 | 0 | 0.00 | 0.788 | 0.937 | (A) BRAF Class 1 Melanoma |

| | | | | | | | | |
|---|---|---|---|---|---|---|---|---|
| BBC3 | 19q13.32 | 2 | 0.49 | 0 | 0.00 | 0.79 | 0.937 | (A) BRAF Class 1 Melanoma |
| HIST1H2BD | 6p22.2 | 2 | 0.49 | 0 | 0.00 | 0.79 | 0.937 | (A) BRAF Class 1 Melanoma |
| TGFBR1 | 9q22.33 | 2 | 0.49 | 0 | 0.00 | 0.79 | 0.937 | (A) BRAF Class 1 Melanoma |
| HIST1H3A | 6p22.2 | 2 | 0.51 | 0 | 0.00 | 0.794 | 0.938 | (A) BRAF Class 1 Melanoma |
| HIST1H3F | 6p22.2 | 2 | 0.51 | 0 | 0.00 | 0.794 | 0.938 | (A) BRAF Class 1 Melanoma |
| DDX41 | 5q35.3 | 2 | 1.03 | 0 | 0.00 | 0.8 | 0.938 | (A) BRAF Class 1 Melanoma |
| SAMD9 | 7q21.2 | 2 | 3.08 | 0 | 0.00 | 0.814 | 0.938 | (A) BRAF Class 1 Melanoma |
| NLRP1 | 17p13.2 | 2 | 4.65 | 0 | 0.00 | 0.835 | 0.938 | (A) BRAF Class 1 Melanoma |
| KIF5B | 10p11.22 | 2 | 1.09 | 0 | 0.00 | 0.837 | 0.938 | (A) BRAF Class 1 Melanoma |
| CDH11 | 16q21 | 2 | 4.44 | 0 | 0.00 | 0.842 | 0.938 | (A) BRAF Class 1 Melanoma |
| NIN | 14q22.1 | 2 | 4.44 | 0 | 0.00 | 0.842 | 0.938 | (A) BRAF Class 1 Melanoma |
| APOBEC3G | 22q13.1 | 1 | 3.13 | 0 | 0.00 | 0.842 | 0.938 | (A) BRAF Class 1 Melanoma |
| HSPA5 | 9q33.3 | 1 | 3.13 | 0 | 0.00 | 0.842 | 0.938 | (A) BRAF Class 1 Melanoma |
| USP7 | 16p13.2 | 1 | 3.13 | 0 | 0.00 | 0.842 | 0.938 | (A) BRAF Class 1 Melanoma |
| RPS6KB1 | 17q23.1 | 2 | 1.26 | 0 | 0.00 | 0.844 | 0.938 | (A) BRAF Class 1 Melanoma |
| ZBTB7A | 19p13.3 | 2 | 1.26 | 0 | 0.00 | 0.844 | 0.938 | (A) BRAF Class 1 Melanoma |
| RACK1 | 5q35.3 | 1 | 1.67 | 0 | 0.00 | 0.845 | 0.938 | (A) BRAF Class 1 Melanoma |
| CDK9 | 9q34.11 | 1 | 1.45 | 0 | 0.00 | 0.852 | 0.938 | (A) BRAF Class 1 Melanoma |
| CRTC2 | 1q21.3 | 1 | 1.45 | 0 | 0.00 | 0.852 | 0.938 | (A) BRAF Class 1 Melanoma |
| PSMD13 | 11p15.5 | 1 | 1.45 | 0 | 0.00 | 0.852 | 0.938 | (A) BRAF Class 1 Melanoma |
| MAP3K2 | 2q14.3 | 1 | 1.85 | 0 | 0.00 | 0.857 | 0.938 | (A) BRAF Class 1 Melanoma |
| MGMT | 10q26.3 | 1 | 1.85 | 0 | 0.00 | 0.857 | 0.938 | (A) BRAF Class 1 Melanoma |
| HSP90AB1 | 6p21.1 | 1 | 1.27 | 0 | 0.00 | 0.859 | 0.938 | (A) BRAF Class 1 Melanoma |
| MAP3K7 | 6q15 | 1 | 1.27 | 0 | 0.00 | 0.859 | 0.938 | (A) BRAF Class 1 Melanoma |
| RPL10 | Xq28 | 1 | 2.70 | 0 | 0.00 | 0.86 | 0.938 | (A) BRAF Class 1 Melanoma |
| CDC42 | 1p36.12 | 1 | 0.48 | 0 | 0.00 | 0.863 | 0.938 | (A) BRAF Class 1 Melanoma |
| SAMD9L | 7q21.2 | 2 | 5.00 | 0 | 0.00 | 0.864 | 0.938 | (A) BRAF Class 1 Melanoma |
| FANCF | 11p14.3 | 1 | 0.19 | 0 | 0.00 | 0.868 | 0.938 | (A) BRAF Class 1 Melanoma |
| UGT1A1 | 2q37.1 | 1 | 0.95 | 0 | 0.00 | 0.868 | 0.938 | (A) BRAF Class 1 Melanoma |
| SDHB | 1p36.13 | 1 | 0.13 | 0 | 0.00 | 0.871 | 0.938 | (A) BRAF Class 1 Melanoma |
| CNOT3 | 19q13.42 | 1 | 1.64 | 0 | 0.00 | 0.871 | 0.938 | (A) BRAF Class 1 Melanoma |
| ABCA17P | 16p13.3 | 1 | 0.08 | 0 | 0.00 | 0.874 | 0.938 | (A) BRAF Class 1 Melanoma |
| ABR | 17p13.3 | 1 | 0.08 | 0 | 0.00 | 0.874 | 0.938 | (A) BRAF Class 1 Melanoma |
| ARHGAP4 | Xq28 | 1 | 0.08 | 0 | 0.00 | 0.874 | 0.938 | (A) BRAF Class 1 Melanoma |
| ARSB | 5q14.1 | 1 | 0.08 | 0 | 0.00 | 0.874 | 0.938 | (A) BRAF Class 1 Melanoma |
| C2ORF83 | 2q36.3 | 1 | 0.08 | 0 | 0.00 | 0.874 | 0.938 | (A) BRAF Class 1 Melanoma |
| CASZ1 | 1p36.22 | 1 | 0.08 | 0 | 0.00 | 0.874 | 0.938 | (A) BRAF Class 1 Melanoma |
| CTDSP1 | 2q35 | 1 | 0.08 | 0 | 0.00 | 0.874 | 0.938 | (A) BRAF Class 1 Melanoma |
| DDAH1 | 1p22.3 | 1 | 0.08 | 0 | 0.00 | 0.874 | 0.938 | (A) BRAF Class 1 Melanoma |
| DEF8 | 16q24.3 | 1 | 0.08 | 0 | 0.00 | 0.874 | 0.938 | (A) BRAF Class 1 Melanoma |
| DENND4C | 9p22.1 | 1 | 0.08 | 0 | 0.00 | 0.874 | 0.938 | (A) BRAF Class 1 Melanoma |
| DHRSX | 2.33 and Yp | 1 | 0.08 | 0 | 0.00 | 0.874 | 0.938 | (A) BRAF Class 1 Melanoma |
| GALNT11 | 136.1\|7q36 | 1 | 0.08 | 0 | 0.00 | 0.874 | 0.938 | (A) BRAF Class 1 Melanoma |
| GRIA1 | 5q33.2 | 1 | 0.08 | 0 | 0.00 | 0.874 | 0.938 | (A) BRAF Class 1 Melanoma |
| GRIK1 | 21q21.3 | 1 | 0.08 | 0 | 0.00 | 0.874 | 0.938 | (A) BRAF Class 1 Melanoma |
| HECTD4 | 12q24.13 | 1 | 0.08 | 0 | 0.00 | 0.874 | 0.938 | (A) BRAF Class 1 Melanoma |
| HSD17B6 | 12q13.3 | 1 | 0.08 | 0 | 0.00 | 0.874 | 0.938 | (A) BRAF Class 1 Melanoma |
| )C1053744( | 2p22.2 | 1 | 0.08 | 0 | 0.00 | 0.874 | 0.938 | (A) BRAF Class 1 Melanoma |

| | | | | | | | | |
|---|---|---|---|---|---|---|---|---|
| MYO1D | 17q11.2 | 1 | 0.08 | 0 | 0.00 | 0.874 | 0.938 | (A) BRAF Class 1 Melanoma |
| MYSM1 | 1p32.1 | 1 | 0.08 | 0 | 0.00 | 0.874 | 0.938 | (A) BRAF Class 1 Melanoma |
| PADI2 | 1p36.13 | 1 | 0.08 | 0 | 0.00 | 0.874 | 0.938 | (A) BRAF Class 1 Melanoma |
| PAX9 | 14q13.3 | 1 | 0.08 | 0 | 0.00 | 0.874 | 0.938 | (A) BRAF Class 1 Melanoma |
| PFKFB4 | 3p21.31 | 1 | 0.08 | 0 | 0.00 | 0.874 | 0.938 | (A) BRAF Class 1 Melanoma |
| PLVAP | 19p13.11 | 1 | 0.08 | 0 | 0.00 | 0.874 | 0.938 | (A) BRAF Class 1 Melanoma |
| PNPLA6 | 19p13.2 | 1 | 0.08 | 0 | 0.00 | 0.874 | 0.938 | (A) BRAF Class 1 Melanoma |
| PRMT6 | 1p13.3 | 1 | 0.08 | 0 | 0.00 | 0.874 | 0.938 | (A) BRAF Class 1 Melanoma |
| RAB2A | q12.1-q12. | 1 | 0.08 | 0 | 0.00 | 0.874 | 0.938 | (A) BRAF Class 1 Melanoma |
| SAE1 | 19q13.32 | 1 | 0.08 | 0 | 0.00 | 0.874 | 0.938 | (A) BRAF Class 1 Melanoma |
| SCARA5 | 8p21.1 | 1 | 0.08 | 0 | 0.00 | 0.874 | 0.938 | (A) BRAF Class 1 Melanoma |
| SDCCAG8 | 1q43-q44 | 1 | 0.08 | 0 | 0.00 | 0.874 | 0.938 | (A) BRAF Class 1 Melanoma |
| SEMA6D | 15q21.1 | 1 | 0.08 | 0 | 0.00 | 0.874 | 0.938 | (A) BRAF Class 1 Melanoma |
| SIK3 | 11q23.3 | 1 | 0.08 | 0 | 0.00 | 0.874 | 0.938 | (A) BRAF Class 1 Melanoma |
| TRAJ42 | 14q11.2 | 1 | 0.08 | 0 | 0.00 | 0.874 | 0.938 | (A) BRAF Class 1 Melanoma |
| UGT1A6 | 2q37.1 | 1 | 0.08 | 0 | 0.00 | 0.874 | 0.938 | (A) BRAF Class 1 Melanoma |
| ZNF423 | 16q12.1 | 1 | 0.08 | 0 | 0.00 | 0.874 | 0.938 | (A) BRAF Class 1 Melanoma |
| EME1 | 17q21.33 | 1 | 0.50 | 0 | 0.00 | 0.874 | 0.938 | (A) BRAF Class 1 Melanoma |
| FAAP24 | 19q13.11 | 1 | 0.50 | 0 | 0.00 | 0.874 | 0.938 | (A) BRAF Class 1 Melanoma |
| ID4 | 6p22.3 | 1 | 0.50 | 0 | 0.00 | 0.874 | 0.938 | (A) BRAF Class 1 Melanoma |
| UBE2T | 1q32.1 | 1 | 0.50 | 0 | 0.00 | 0.874 | 0.938 | (A) BRAF Class 1 Melanoma |
| PRKD1 | 14q12 | 1 | 0.53 | 0 | 0.00 | 0.874 | 0.938 | (A) BRAF Class 1 Melanoma |
| PTP4A1 | 6q12 | 1 | 0.58 | 0 | 0.00 | 0.877 | 0.938 | (A) BRAF Class 1 Melanoma |
| KMT5A | 12q24.31 | 1 | 0.58 | 0 | 0.00 | 0.878 | 0.938 | (A) BRAF Class 1 Melanoma |
| ERCC1 | 19q13.32 | 1 | 0.26 | 0 | 0.00 | 0.878 | 0.938 | (A) BRAF Class 1 Melanoma |
| SMARCE1 | 17q21.2 | 1 | 0.43 | 0 | 0.00 | 0.879 | 0.938 | (A) BRAF Class 1 Melanoma |
| BTG1 | 12q21.33 | 1 | 0.36 | 0 | 0.00 | 0.88 | 0.938 | (A) BRAF Class 1 Melanoma |
| GNA13 | 17q24.1 | 1 | 0.34 | 0 | 0.00 | 0.883 | 0.938 | (A) BRAF Class 1 Melanoma |
| CDH20 | 18q21.33 | 1 | 1.09 | 0 | 0.00 | 0.885 | 0.938 | (A) BRAF Class 1 Melanoma |
| EPHB4 | 7q22.1 | 1 | 0.80 | 0 | 0.00 | 0.887 | 0.938 | (A) BRAF Class 1 Melanoma |
| DCUN1D1 | 3q26.33 | 1 | 0.25 | 0 | 0.00 | 0.887 | 0.938 | (A) BRAF Class 1 Melanoma |
| PDPK1 | 16p13.3 | 1 | 0.23 | 0 | 0.00 | 0.888 | 0.938 | (A) BRAF Class 1 Melanoma |
| PMAIP1 | 18q21.32 | 1 | 0.25 | 0 | 0.00 | 0.889 | 0.938 | (A) BRAF Class 1 Melanoma |
| HIST2H3D | 1q21.2 | 1 | 0.26 | 0 | 0.00 | 0.89 | 0.938 | (A) BRAF Class 1 Melanoma |
| HIST1H3H | 6p22.1 | 1 | 0.25 | 0 | 0.00 | 0.891 | 0.938 | (A) BRAF Class 1 Melanoma |
| HIST2H3C | 1q21.2 | 1 | 0.25 | 0 | 0.00 | 0.891 | 0.938 | (A) BRAF Class 1 Melanoma |
| PHLPP2 | 16q22.2 | 1 | 1.35 | 0 | 0.00 | 0.892 | 0.938 | (A) BRAF Class 1 Melanoma |
| PAX3 | 2q36.1 | 1 | 0.45 | 0 | 0.00 | 0.892 | 0.938 | (A) BRAF Class 1 Melanoma |
| DDR1 | 6p21.33 | 1 | 1.00 | 0 | 0.00 | 0.893 | 0.938 | (A) BRAF Class 1 Melanoma |
| SRP19 | 5q22.2 | 1 | 0.69 | 0 | 0.00 | 0.895 | 0.938 | (A) BRAF Class 1 Melanoma |
| TYRO3 | 15q15.1 | 1 | 1.28 | 0 | 0.00 | 0.897 | 0.938 | (A) BRAF Class 1 Melanoma |
| ZBTB2 | 6q25.1 | 1 | 0.44 | 0 | 0.00 | 0.901 | 0.938 | (A) BRAF Class 1 Melanoma |
| ETNK1 | 12p12.1 | 1 | 1.75 | 0 | 0.00 | 0.905 | 0.938 | (A) BRAF Class 1 Melanoma |
| CAMTA1 | 36.31-p36. | 1 | 1.69 | 0 | 0.00 | 0.908 | 0.938 | (A) BRAF Class 1 Melanoma |
| NR4A3 | 9q31.1 | 1 | 1.69 | 0 | 0.00 | 0.908 | 0.938 | (A) BRAF Class 1 Melanoma |
| LAMP1 | 13q34 | 1 | 0.54 | 0 | 0.00 | 0.911 | 0.938 | (A) BRAF Class 1 Melanoma |
| TCF12 | 15q21.3 | 1 | 1.39 | 0 | 0.00 | 0.911 | 0.938 | (A) BRAF Class 1 Melanoma |
| NCOR2 | 12q24.31 | 1 | 2.38 | 0 | 0.00 | 0.913 | 0.938 | (A) BRAF Class 1 Melanoma |

| | | | | | | | | |
|---|---|---|---|---|---|---|---|---|
| AURKC | 19q13.43 | 1 | 2.33 | 0 | 0.00 | 0.915 | 0.938 | (A) BRAF Class 1 Melanoma |
| CSMD3 | 8q23.3 | 1 | 2.33 | 0 | 0.00 | 0.915 | 0.938 | (A) BRAF Class 1 Melanoma |
| CYP2C19 | 10q23.33 | 1 | 2.33 | 0 | 0.00 | 0.915 | 0.938 | (A) BRAF Class 1 Melanoma |
| DST | 6p12.1 | 1 | 2.33 | 0 | 0.00 | 0.915 | 0.938 | (A) BRAF Class 1 Melanoma |
| FN1 | 2q35 | 1 | 2.33 | 0 | 0.00 | 0.915 | 0.938 | (A) BRAF Class 1 Melanoma |
| FOXP4 | 6p21.1 | 1 | 2.33 | 0 | 0.00 | 0.915 | 0.938 | (A) BRAF Class 1 Melanoma |
| GDNF | 5p13.2 | 1 | 2.33 | 0 | 0.00 | 0.915 | 0.938 | (A) BRAF Class 1 Melanoma |
| GUCY1A2 | 11q22.3 | 1 | 2.33 | 0 | 0.00 | 0.915 | 0.938 | (A) BRAF Class 1 Melanoma |
| IKBKB | 8p11.21 | 1 | 2.33 | 0 | 0.00 | 0.915 | 0.938 | (A) BRAF Class 1 Melanoma |
| ITGA10 | 1q21.1 | 1 | 2.33 | 0 | 0.00 | 0.915 | 0.938 | (A) BRAF Class 1 Melanoma |
| STK36 | 2q35 | 1 | 2.33 | 0 | 0.00 | 0.915 | 0.938 | (A) BRAF Class 1 Melanoma |
| TNK2 | 3q29 | 1 | 2.33 | 0 | 0.00 | 0.915 | 0.938 | (A) BRAF Class 1 Melanoma |
| TRIM24 | 7q33-q34 | 1 | 2.33 | 0 | 0.00 | 0.915 | 0.938 | (A) BRAF Class 1 Melanoma |
| ASPSCR1 | 17q25.3 | 1 | 2.27 | 0 | 0.00 | 0.917 | 0.938 | (A) BRAF Class 1 Melanoma |
| PRDM16 | 1p36.32 | 1 | 2.27 | 0 | 0.00 | 0.917 | 0.938 | (A) BRAF Class 1 Melanoma |
| RBM15 | 1p13.3 | 1 | 2.27 | 0 | 0.00 | 0.917 | 0.938 | (A) BRAF Class 1 Melanoma |
| ARNT | 1q21.3 | 1 | 2.22 | 0 | 0.00 | 0.918 | 0.938 | (A) BRAF Class 1 Melanoma |
| KNL1 | 15q15.1 | 1 | 2.22 | 0 | 0.00 | 0.918 | 0.938 | (A) BRAF Class 1 Melanoma |
| NFKB2 | 10q24.32 | 1 | 2.22 | 0 | 0.00 | 0.918 | 0.938 | (A) BRAF Class 1 Melanoma |
| PDE4DIP | 1q21.2 | 1 | 2.22 | 0 | 0.00 | 0.918 | 0.938 | (A) BRAF Class 1 Melanoma |
| PER1 | 17p13.1 | 1 | 2.22 | 0 | 0.00 | 0.918 | 0.938 | (A) BRAF Class 1 Melanoma |
| TPR | 1q31.1 | 1 | 2.22 | 0 | 0.00 | 0.918 | 0.938 | (A) BRAF Class 1 Melanoma |
| ZNF384 | 12p13.31 | 1 | 2.22 | 0 | 0.00 | 0.918 | 0.938 | (A) BRAF Class 1 Melanoma |
| CSNK1A1 | 5q32 | 1 | 0.63 | 0 | 0.00 | 0.919 | 0.938 | (A) BRAF Class 1 Melanoma |
| FGF8 | 10q24.32 | 1 | 0.63 | 0 | 0.00 | 0.919 | 0.938 | (A) BRAF Class 1 Melanoma |
| CD22 | 19q13.12 | 1 | 2.17 | 0 | 0.00 | 0.92 | 0.938 | (A) BRAF Class 1 Melanoma |
| PDCD11 | 10q24.33 | 1 | 2.86 | 0 | 0.00 | 0.921 | 0.938 | (A) BRAF Class 1 Melanoma |
| PCDHAC2 | 5q31.3 | 2 | 8.00 | 0 | 0.00 | 0.923 | 0.938 | (A) BRAF Class 1 Melanoma |
| TGFB1 | 19q13.2 | 2 | 8.00 | 0 | 0.00 | 0.923 | 0.938 | (A) BRAF Class 1 Melanoma |
| ADGRB3 | 6q12-q13 | 2 | 7.69 | 0 | 0.00 | 0.926 | 0.938 | (A) BRAF Class 1 Melanoma |
| NUP214 | 9q34.13 | 1 | 2.00 | 0 | 0.00 | 0.926 | 0.938 | (A) BRAF Class 1 Melanoma |
| UBR5 | 8q22.3 | 1 | 2.00 | 0 | 0.00 | 0.926 | 0.938 | (A) BRAF Class 1 Melanoma |
| BEND2 | Xp22.13 | 1 | 2.50 | 0 | 0.00 | 0.93 | 0.938 | (A) BRAF Class 1 Melanoma |
| DDX31 | 9q34.13 | 1 | 2.50 | 0 | 0.00 | 0.93 | 0.938 | (A) BRAF Class 1 Melanoma |
| FOSB | 19q13.32 | 1 | 2.50 | 0 | 0.00 | 0.93 | 0.938 | (A) BRAF Class 1 Melanoma |
| FRK | 6q22.1 | 1 | 2.50 | 0 | 0.00 | 0.93 | 0.938 | (A) BRAF Class 1 Melanoma |
| GLMN | 1p22.1 | 1 | 2.50 | 0 | 0.00 | 0.93 | 0.938 | (A) BRAF Class 1 Melanoma |
| OGA | 10q24.32 | 1 | 2.50 | 0 | 0.00 | 0.93 | 0.938 | (A) BRAF Class 1 Melanoma |
| OTX2 | 14q22.3 | 1 | 2.50 | 0 | 0.00 | 0.93 | 0.938 | (A) BRAF Class 1 Melanoma |
| POLR2A | 17p13.1 | 1 | 2.50 | 0 | 0.00 | 0.93 | 0.938 | (A) BRAF Class 1 Melanoma |
| TBXT | 6q27 | 1 | 2.50 | 0 | 0.00 | 0.93 | 0.938 | (A) BRAF Class 1 Melanoma |
| ZFPM1 | 16q24.2 | 1 | 2.50 | 0 | 0.00 | 0.93 | 0.938 | (A) BRAF Class 1 Melanoma |
| ECT2L | 6q24.1 | 1 | 2.38 | 0 | 0.00 | 0.933 | 0.94 | (A) BRAF Class 1 Melanoma |
| TBL1XR1 | 3q26.32 | 1 | 2.38 | 0 | 0.00 | 0.933 | 0.94 | (A) BRAF Class 1 Melanoma |
| CACNA1E | 1q25.3 | 1 | 4.00 | 0 | 0.00 | 0.962 | 0.964 | (A) BRAF Class 1 Melanoma |
| GNAI3 | 1p13.3 | 1 | 4.00 | 0 | 0.00 | 0.962 | 0.964 | (A) BRAF Class 1 Melanoma |
| SP140 | 2q37.1 | 1 | 4.00 | 0 | 0.00 | 0.962 | 0.964 | (A) BRAF Class 1 Melanoma |
| UMODL1 | 21q22.3 | 1 | 4.00 | 0 | 0.00 | 0.962 | 0.964 | (A) BRAF Class 1 Melanoma |

| | | | | | | | | |
|---|---|---|---|---|---|---|---|---|
| RPL22 | 1p36.31 | 1 | 3.70 | 0 | 0.00 | 0.964 | 0.965 | (A) BRAF Class 1 Melanoma |
| BRAF | 7q34 | 1183 | 100.00 | 171 | 100.00 | 1 | 1 | (B) BRAF Class 3 Melanoma |

#### Supplementary Table 10

| Gene | Cytoband | Class 1 Occurrence | Class 1 % | Class 3 occurrence | Class 3 % | SumClass | pvalue | qvalue | Enriched In |
|---|---|---|---|---|---|---|---|---|---|
| TENT5C | 1p12 | 17 | 2.20 | 8 | 6.84 | 25 | 0.011 | 0.0493 | (B) BRAF Class 3 Melanoma |
| CDH1 | 16q22.1 | 18 | 1.67 | 8 | 5.19 | 26 | 0.0107 | 0.0483 | (B) BRAF Class 3 Melanoma |
| INHBA | 7p14.1 | 23 | 4.49 | 9 | 12.50 | 32 | 0.0106 | 0.0483 | (B) BRAF Class 3 Melanoma |
| SLX4 | 16p13.3 | 29 | 5.47 | 10 | 14.08 | 39 | 0.0105 | 0.0481 | (B) BRAF Class 3 Melanoma |
| FANCI | 15q26.1 | 11 | 2.46 | 6 | 9.84 | 17 | 0.009962 | 0.0461 | (B) BRAF Class 3 Melanoma |
| SMAD4 | 18q21.2 | 10 | 0.89 | 6 | 3.70 | 16 | 0.009732 | 0.0452 | (B) BRAF Class 3 Melanoma |
| XPC | 3p25.1 | 9 | 3.17 | 6 | 13.33 | 15 | 0.009062 | 0.043 | (B) BRAF Class 3 Melanoma |
| ZFHX4 | 8q21.13 | 11 | 20.37 | 6 | 66.67 | 17 | 0.008836 | 0.0422 | (B) BRAF Class 3 Melanoma |
| NSD2 | 4p16.3 | 12 | 2.80 | 7 | 10.45 | 19 | 0.008096 | 0.0397 | (B) BRAF Class 3 Melanoma |
| PALB2 | 16p12.2 | 24 | 2.97 | 10 | 8.26 | 34 | 0.008157 | 0.0397 | (B) BRAF Class 3 Melanoma |
| TCF7L2 | 0q25.2-q25.[illegible] | 9 | 1.30 | 6 | 5.88 | 15 | 0.007453 | 0.0367 | (B) BRAF Class 3 Melanoma |
| LYN | 8q12.1 | 15 | 3.83 | 7 | 14.00 | 22 | 0.007009 | 0.0347 | (B) BRAF Class 3 Melanoma |
| DNMT3B | 20q11.21 | 11 | 2.70 | 6 | 11.76 | 17 | 0.006669 | 0.0334 | (B) BRAF Class 3 Melanoma |
| AR | Xq12 | 31 | 3.84 | 12 | 9.84 | 43 | 0.006425 | 0.0325 | (B) BRAF Class 3 Melanoma |

| | | | | | | | | | |
|---|---|---|---|---|---|---|---|---|---|
| FGFR3 | 4p16.3 | 32 | 2.70 | 12 | 7.02 | 44 | 0.006109 | 0.0313 | (B) BRAF Class 3 Melanoma |
| DOCK8 | 9p24.3 | 13 | 6.47 | 7 | 24.14 | 20 | 0.00597 | 0.031 | (B) BRAF Class 3 Melanoma |
| PPARG | 3p25.2 | 17 | 2.64 | 8 | 9.20 | 25 | 0.005741 | 0.0306 | (B) BRAF Class 3 Melanoma |
| ARHGAP35 | 19q13.32 | 14 | 5.83 | 8 | 21.05 | 22 | 0.004481 | 0.0247 | (B) BRAF Class 3 Melanoma |
| CDK8 | 13q12.13 | 11 | 1.52 | 7 | 6.67 | 18 | 0.004225 | 0.0235 | (B) BRAF Class 3 Melanoma |
| MST1R | 3p21.31 | 13 | 2.77 | 7 | 11.67 | 20 | 0.003998 | 0.0225 | (B) BRAF Class 3 Melanoma |
| ERBB3 | 12q13.2 | 21 | 2.59 | 10 | 8.20 | 31 | 0.004023 | 0.0225 | (B) BRAF Class 3 Melanoma |
| IGF1R | 15q26.3 | 42 | 5.15 | 15 | 12.30 | 57 | 0.003961 | 0.0224 | (B) BRAF Class 3 Melanoma |
| STAG2 | Xq25 | 24 | 3.11 | 11 | 9.40 | 35 | 0.003358 | 0.02 | (B) BRAF Class 3 Melanoma |
| EZH2 | 7q36.1 | 33 | 3.19 | 13 | 8.67 | 46 | 0.003069 | 0.0186 | (B) BRAF Class 3 Melanoma |
| LRP1B | 2q22.1-q22.2 | 86 | 30.82 | 21 | 55.26 | 107 | 0.003066 | 0.0186 | (B) BRAF Class 3 Melanoma |
| TCF3 | 19p13.3 | 14 | 2.12 | 8 | 8.60 | 22 | 0.002987 | 0.0184 | (B) BRAF Class 3 Melanoma |
| PTEN | 10q23.31 | 174 | 14.71 | 12 | 7.06 | 186 | 0.002968 | 0.0184 | (A) BRAF Class 1 Melanoma |
| SNCAIP | 5q23.2 | 36 | 13.09 | 12 | 34.29 | 48 | 0.002689 | 0.0169 | (B) BRAF Class 3 Melanoma |
| NF2 | 22q12.2 | 16 | 1.97 | 9 | 7.44 | 25 | 0.002595 | 0.0165 | (B) BRAF Class 3 Melanoma |

| | | | | | | | | | |
|---|---|---|---|---|---|---|---|---|---|
| PIK3C2G | 12p12.3 | 47 | 9.94 | 15 | 23.81 | 62 | 0.002601 | 0.0165 | (B) BRAF Class 3 Melanoma |
| FANCA | 16q24.3 | 27 | 3.34 | 12 | 9.92 | 39 | 0.002456 | 0.0159 | (B) BRAF Class 3 Melanoma |
| ALOX12B | 17p13.1 | 13 | 2.48 | 8 | 10.67 | 21 | 0.002201 | 0.0145 | (B) BRAF Class 3 Melanoma |
| NTRK2 | 9q21.33 | 18 | 2.23 | 10 | 8.26 | 28 | 0.001647 | 0.0113 | (B) BRAF Class 3 Melanoma |
| TOP1 | 20q12 | 9 | 1.61 | 7 | 8.86 | 16 | 0.001576 | 0.011 | (B) BRAF Class 3 Melanoma |
| TRAF7 | 16p13.3 | 12 | 1.86 | 8 | 8.70 | 20 | 0.001533 | 0.0108 | (B) BRAF Class 3 Melanoma |
| RECQL4 | 8q24.3 | 35 | 5.13 | 14 | 14.29 | 49 | 0.001534 | 0.0108 | (B) BRAF Class 3 Melanoma |
| CCNE1 | 19q12 | 8 | 0.99 | 7 | 5.79 | 15 | 0.001409 | 0.0102 | (B) BRAF Class 3 Melanoma |
| MUTYH | 1p34.1 | 14 | 1.81 | 9 | 7.69 | 23 | 0.001423 | 0.0102 | (B) BRAF Class 3 Melanoma |
| MAGI2 | 7q21.11 | 18 | 7.89 | 8 | 32.00 | 26 | 0.001402 | 0.0102 | (B) BRAF Class 3 Melanoma |
| FAT1 | 4q35.2 | 92 | 13.22 | 26 | 26.00 | 118 | 0.001224 | 0.008994 | (B) BRAF Class 3 Melanoma |
| RUNX1 | 21q22.12 | 24 | 3.10 | 12 | 10.26 | 36 | 0.001202 | 0.008902 | (B) BRAF Class 3 Melanoma |
| FLT4 | 5q35.3 | 61 | 8.01 | 21 | 18.10 | 82 | 0.001075 | 0.00803 | (B) BRAF Class 3 Melanoma |
| PIK3CG | 7q22.3 | 30 | 5.68 | 13 | 17.33 | 43 | 0.001037 | 0.007804 | (B) BRAF Class 3 Melanoma |
| POLQ | 3q13.33 | 13 | 6.05 | 9 | 25.71 | 22 | 0.001002 | 0.007609 | (B) BRAF Class 3 Melanoma |

| | | | | | | | | | |
|---|---|---|---|---|---|---|---|---|---|
| SUFU | 10q24.32 | 13 | 1.68 | 9 | 7.69 | 22 | 0.0009726 | 0.007446 | (B) BRAF Class 3 Melanoma |
| SPTA1 | 1q23.1 | 53 | 20.08 | 16 | 47.06 | 69 | 0.0009523 | 0.007352 | (B) BRAF Class 3 Melanoma |
| ERBB2 | 17q12 | 24 | 2.03 | 12 | 7.02 | 36 | 0.0009094 | 0.007081 | (B) BRAF Class 3 Melanoma |
| ERG | 21q22.2 | 28 | 3.88 | 13 | 12.38 | 41 | 0.0008753 | 0.006874 | (B) BRAF Class 3 Melanoma |
| JAK2 | 9p24.1 | 24 | 2.16 | 12 | 7.55 | 36 | 0.0008343 | 0.006609 | (B) BRAF Class 3 Melanoma |
| NTRK1 | 1q23.1 | 42 | 5.10 | 17 | 13.60 | 59 | 0.0008148 | 0.006512 | (B) BRAF Class 3 Melanoma |
| BRCA1 | 17q21.31 | 34 | 4.21 | 15 | 12.30 | 49 | 0.0007988 | 0.00644 | (B) BRAF Class 3 Melanoma |
| TP63 | 3q28 | 38 | 8.94 | 14 | 25.93 | 52 | 0.0006826 | 0.005588 | (B) BRAF Class 3 Melanoma |
| KMT2B | 19q13.12 | 40 | 10.96 | 15 | 30.00 | 55 | 0.0006869 | 0.005588 | (B) BRAF Class 3 Melanoma |
| CBLB | 3q13.11 | 8 | 2.76 | 8 | 16.00 | 16 | 0.0006504 | 0.005436 | (B) BRAF Class 3 Melanoma |
| ARID1B | 6q25.3 | 46 | 6.16 | 18 | 16.07 | 64 | 0.000641 | 0.005407 | (B) BRAF Class 3 Melanoma |
| BRIP1 | 17q23.2 | 29 | 3.75 | 14 | 11.97 | 43 | 0.0005988 | 0.005098 | (B) BRAF Class 3 Melanoma |
| PIK3CA | 3q26.32 | 34 | 2.87 | 15 | 8.77 | 49 | 0.0005831 | 0.005011 | (B) BRAF Class 3 Melanoma |
| PIK3C2B | 1q32.1 | 24 | 4.99 | 12 | 17.65 | 36 | 0.0005623 | 0.004879 | (B) BRAF Class 3 Melanoma |
| CIITA | 16p13.13 | 18 | 6.52 | 11 | 25.00 | 29 | 0.0005282 | 0.004728 | (B) BRAF Class 3 Melanoma |

| | | | | | | | | | |
|---|---|---|---|---|---|---|---|---|---|
| MPL | 1p34.2 | 19 | 1.76 | 11 | 7.14 | 30 | 0.0005354 | 0.004728 | (B) BRAF Class 3 Melanoma |
| FBXW7 | 4q31.3 | 30 | 2.66 | 14 | 8.64 | 44 | 0.0005373 | 0.004728 | (B) BRAF Class 3 Melanoma |
| CUX1 | 7q22.1 | 34 | 7.62 | 15 | 22.39 | 49 | 0.0005398 | 0.004728 | (B) BRAF Class 3 Melanoma |
| KDM5A | 12p13.33 | 22 | 3.07 | 12 | 11.43 | 34 | 0.0004977 | 0.004534 | (B) BRAF Class 3 Melanoma |
| TERT | 5p15.33 | 460 | 60.85 | 87 | 76.99 | 547 | 0.0004878 | 0.004489 | (B) BRAF Class 3 Melanoma |
| FANCD2 | 3p25.3 | 23 | 4.45 | 12 | 16.22 | 35 | 0.0004818 | 0.004479 | (B) BRAF Class 3 Melanoma |
| TMPRSS2 | 21q22.3 | 15 | 2.09 | 10 | 9.52 | 25 | 0.000475 | 0.004461 | (B) BRAF Class 3 Melanoma |
| MGA | 15q15.1 | 68 | 11.11 | 22 | 25.58 | 90 | 0.0004609 | 0.004373 | (B) BRAF Class 3 Melanoma |
| SLIT2 | 4p15.31 | 28 | 10.61 | 12 | 35.29 | 40 | 0.0004386 | 0.004206 | (B) BRAF Class 3 Melanoma |
| POLE | 12q24.33 | 60 | 7.91 | 21 | 19.27 | 81 | 0.0004253 | 0.004121 | (B) BRAF Class 3 Melanoma |
| KDM5C | Xp11.22 | 24 | 3.14 | 13 | 11.21 | 37 | 0.0004204 | 0.004118 | (B) BRAF Class 3 Melanoma |
| PDGFRA | 4q12 | 50 | 4.32 | 19 | 11.52 | 69 | 0.0004088 | 0.004092 | (B) BRAF Class 3 Melanoma |
| TP53BP1 | 15q15.3 | 18 | 4.41 | 11 | 18.03 | 29 | 0.0003955 | 0.004003 | (B) BRAF Class 3 Melanoma |
| STAT4 | 2q32.2-q32.3 | 10 | 3.60 | 8 | 21.05 | 18 | 0.0003809 | 0.003917 | (B) BRAF Class 3 Melanoma |
| GATA3 | 10p14 | 20 | 2.59 | 12 | 10.26 | 32 | 0.0003467 | 0.003679 | (B) BRAF Class 3 Melanoma |

| | | | | | | | | | |
|---|---|---|---|---|---|---|---|---|---|
| MAP3K1 | 5q11.2 | 27 | 3.50 | 14 | 11.97 | 41 | 0.0003446 | 0.003679 | (B) BRAF Class 3 Melanoma |
| HNF1A | 12q24.31 | 18 | 1.63 | 11 | 7.10 | 29 | 0.0003298 | 0.003619 | (B) BRAF Class 3 Melanoma |
| PTCH1 | 9q22.32 | 27 | 3.34 | 14 | 11.57 | 41 | 0.0003124 | 0.00347 | (B) BRAF Class 3 Melanoma |
| CCND2 | 12p13.32 | 10 | 1.25 | 9 | 7.44 | 19 | 0.0002501 | 0.002813 | (B) BRAF Class 3 Melanoma |
| FANCM | 14q21.2 | 13 | 4.78 | 10 | 23.26 | 23 | 0.0002446 | 0.002791 | (B) BRAF Class 3 Melanoma |
| TSC2 | 16p13.3 | 30 | 3.70 | 15 | 12.40 | 45 | 0.0002451 | 0.002791 | (B) BRAF Class 3 Melanoma |
| MTOR | 1p36.22 | 41 | 4.73 | 18 | 13.85 | 59 | 0.0002268 | 0.002649 | (B) BRAF Class 3 Melanoma |
| PRKDC | 8q11.21 | 54 | 10.53 | 21 | 26.58 | 75 | 0.0002246 | 0.002649 | (B) BRAF Class 3 Melanoma |
| FANCE | 6p21.31 | 7 | 1.35 | 8 | 10.13 | 15 | 0.0002013 | 0.002446 | (B) BRAF Class 3 Melanoma |
| GNAS | 20q13.32 | 13 | 1.15 | 10 | 6.17 | 23 | 0.0001979 | 0.002446 | (B) BRAF Class 3 Melanoma |
| CDH4 | 20q13.33 | 15 | 7.46 | 10 | 34.48 | 25 | 0.0001989 | 0.002446 | (B) BRAF Class 3 Melanoma |
| BRCA2 | 13q13.1 | 57 | 7.06 | 22 | 18.18 | 79 | 0.0001807 | 0.002287 | (B) BRAF Class 3 Melanoma |
| PAK5 | 20p12.2 | 63 | 13.64 | 21 | 33.87 | 84 | 0.0001636 | 0.002099 | (B) BRAF Class 3 Melanoma |
| PTK2B | 8p21.2 | 9 | 4.19 | 9 | 25.71 | 18 | 0.0001491 | 0.001941 | (B) BRAF Class 3 Melanoma |
| DAXX | 6p21.32 | 16 | 2.26 | 11 | 11.11 | 27 | 0.0001339 | 0.001802 | (B) BRAF Class 3 Melanoma |

| | | | | | | | | | |
|---|---|---|---|---|---|---|---|---|---|
| PBRM1 | 3p21.1 | 24 | 3.14 | 14 | 12.07 | 38 | 0.0001365 | 0.001802 | (B) BRAF Class 3 Melanoma |
| NOTCH4 | 6p21.32 | 33 | 6.73 | 15 | 22.73 | 48 | 0.0001365 | 0.001802 | (B) BRAF Class 3 Melanoma |
| ATRX | Xq21.1 | 47 | 5.82 | 20 | 16.53 | 67 | 0.0001266 | 0.001748 | (B) BRAF Class 3 Melanoma |
| BRD4 | 19p13.12 | 39 | 5.13 | 18 | 15.93 | 57 | 0.000113 | 0.001584 | (B) BRAF Class 3 Melanoma |
| CIC | 19q13.2 | 40 | 5.54 | 18 | 17.14 | 58 | 0.0001075 | 0.001531 | (B) BRAF Class 3 Melanoma |
| RPTOR | 17q25.3 | 34 | 4.72 | 17 | 16.19 | 51 | 6.259E-05 | 0.0009504 | (B) BRAF Class 3 Melanoma |
| NOTCH3 | 19p13.12 | 87 | 11.65 | 29 | 26.85 | 116 | 5.859E-05 | 0.0009046 | (B) BRAF Class 3 Melanoma |
| PDGFRB | 5q32 | 32 | 3.96 | 17 | 14.05 | 49 | 5.207E-05 | 0.0008178 | (B) BRAF Class 3 Melanoma |
| DICER1 | 14q32.13 | 25 | 3.30 | 15 | 13.27 | 40 | 0.0000496 | 0.0007988 | (B) BRAF Class 3 Melanoma |
| JAK3 | 19p13.11 | 37 | 3.32 | 18 | 11.32 | 55 | 4.998E-05 | 0.0007988 | (B) BRAF Class 3 Melanoma |
| APC | 5q22.2 | 74 | 6.86 | 27 | 17.42 | 101 | 4.231E-05 | 0.0007008 | (B) BRAF Class 3 Melanoma |
| EP300 | 22q13.2 | 42 | 5.43 | 20 | 17.09 | 62 | 4.008E-05 | 0.0006762 | (B) BRAF Class 3 Melanoma |
| COL7A1 | 3p21.31 | 46 | 22.89 | 18 | 62.07 | 64 | 3.672E-05 | 0.0006434 | (B) BRAF Class 3 Melanoma |
| EPHA7 | 6q16.1 | 58 | 10.41 | 23 | 28.40 | 81 | 3.533E-05 | 0.000631 | (B) BRAF Class 3 Melanoma |
| PTPN11 | 12q24.13 | 12 | 1.07 | 11 | 6.79 | 23 | 3.165E-05 | 0.0005767 | (B) BRAF Class 3 Melanoma |

| | | | | | | | | | |
|---|---|---|---|---|---|---|---|---|---|
| TET1 | 10q21.3 | 31 | 5.02 | 16 | 19.05 | 47 | 3.144E-05 | 0.0005767 | (B) BRAF Class 3 Melanoma |
| PTPRT | 20q12-q13.1 | 90 | 19.19 | 28 | 43.75 | 118 | 2.925E-05 | 0.0005551 | (B) BRAF Class 3 Melanoma |
| SETBP1 | 18q12.3 | 40 | 8.70 | 19 | 27.94 | 59 | 2.794E-05 | 0.0005416 | (B) BRAF Class 3 Melanoma |
| SYK | 9q22.2 | 17 | 2.88 | 13 | 14.77 | 30 | 2.368E-05 | 0.000469 | (B) BRAF Class 3 Melanoma |
| EPHA3 | 3p11.1 | 45 | 7.63 | 21 | 23.86 | 66 | 1.896E-05 | 0.0003839 | (B) BRAF Class 3 Melanoma |
| ASXL1 | 20q11.21 | 31 | 3.97 | 18 | 15.25 | 49 | 1.323E-05 | 0.0002793 | (B) BRAF Class 3 Melanoma |
| EGFR | 7p11.2 | 51 | 4.31 | 23 | 13.45 | 74 | 1.349E-05 | 0.0002793 | (B) BRAF Class 3 Melanoma |
| CSF3R | 1p34.3 | 13 | 1.85 | 12 | 11.65 | 25 | 1.265E-05 | 0.0002743 | (B) BRAF Class 3 Melanoma |
| TP53 | 17p13.1 | 175 | 14.79 | 49 | 28.82 | 224 | 1.172E-05 | 0.0002605 | (B) BRAF Class 3 Melanoma |
| ALK | 2p23.2-p23.1 | 71 | 6.00 | 28 | 16.37 | 99 | 1.095E-05 | 0.0002494 | (B) BRAF Class 3 Melanoma |
| GLI1 | 12q13.3 | 31 | 4.23 | 18 | 16.51 | 49 | 1.038E-05 | 0.0002425 | (B) BRAF Class 3 Melanoma |
| DDR2 | 1q23.3 | 19 | 2.16 | 15 | 11.19 | 34 | 6.405E-06 | 0.0001535 | (B) BRAF Class 3 Melanoma |
| ABCB11 | 2q31.1 | 11 | 4.93 | 11 | 34.38 | 22 | 6.084E-06 | 0.0001498 | (B) BRAF Class 3 Melanoma |
| RET | 10q11.21 | 44 | 3.72 | 22 | 12.87 | 66 | 5.855E-06 | 0.0001482 | (B) BRAF Class 3 Melanoma |
| ARID1A | 1p36.11 | 63 | 7.80 | 27 | 22.31 | 90 | 5.67E-06 | 0.0001476 | (B) BRAF Class 3 Melanoma |

| | | | | | | | | | |
|---|---|---|---|---|---|---|---|---|---|
| PREX2 | 8q13.2 | 75 | 17.69 | 27 | 45.76 | 102 | 4.483E-06 | 0.0001201 | (B) BRAF Class 3 Melanoma |
| CBL | 11q23.3 | 30 | 3.68 | 19 | 15.57 | 49 | 2.509E-06 | 6.969E-05 | (B) BRAF Class 3 Melanoma |
| GRIN2A | 16p13.2 | 92 | 17.46 | 32 | 42.67 | 124 | 2.524E-06 | 6.969E-05 | (B) BRAF Class 3 Melanoma |
| NOTCH1 | 9q34.3 | 49 | 4.20 | 24 | 14.46 | 73 | 2.065E-06 | 6.068E-05 | (B) BRAF Class 3 Melanoma |
| IL7R | 5p13.2 | 49 | 6.89 | 24 | 23.08 | 73 | 1.913E-06 | 0.0000581 | (B) BRAF Class 3 Melanoma |
| FGFR2 | 10q26.13 | 47 | 3.97 | 24 | 14.04 | 71 | 1.561E-06 | 4.903E-05 | (B) BRAF Class 3 Melanoma |
| SMARCA4 | 19p13.2 | 50 | 6.19 | 25 | 20.66 | 75 | 1.519E-06 | 4.903E-05 | (B) BRAF Class 3 Melanoma |
| MECOM | 3q26.2 | 39 | 14.66 | 21 | 50.00 | 60 | 1.191E-06 | 4.018E-05 | (B) BRAF Class 3 Melanoma |
| IKZF1 | 7p12.2 | 62 | 8.07 | 29 | 24.79 | 91 | 7.02E-07 | 2.461E-05 | (B) BRAF Class 3 Melanoma |
| BLM | 15q26.1 | 23 | 3.02 | 18 | 15.52 | 41 | 6.37E-07 | 2.322E-05 | (B) BRAF Class 3 Melanoma |
| CARD11 | 7p22.2 | 61 | 7.89 | 29 | 24.79 | 90 | 4.72E-07 | 1.793E-05 | (B) BRAF Class 3 Melanoma |
| MET | 7q31.2 | 70 | 5.96 | 31 | 18.24 | 101 | 4.56E-07 | 1.793E-05 | (B) BRAF Class 3 Melanoma |
| ROS1 | 6q22.1 | 150 | 17.38 | 49 | 37.69 | 199 | 3.66E-07 | 1.516E-05 | (B) BRAF Class 3 Melanoma |
| RUNX1T1 | 8q21.3 | 35 | 7.58 | 21 | 31.34 | 56 | 3.29E-07 | 1.425E-05 | (B) BRAF Class 3 Melanoma |
| NTRK3 | 15q25.3 | 52 | 6.44 | 27 | 22.31 | 79 | 2.87E-07 | 1.307E-05 | (B) BRAF Class 3 Melanoma |

| | | | | | | | | | |
|---|---|---|---|---|---|---|---|---|---|
| AXL | 19q13.2 | 30 | 3.71 | 21 | 17.36 | 51 | 2.01E-07 | 9.628E-06 | (B) BRAF Class 3 Melanoma |
| KIT | 4q12 | 24 | 2.03 | 19 | 11.11 | 43 | 1.72E-07 | 8.697E-06 | (B) BRAF Class 3 Melanoma |
| HRAS | 11p15.5 | 4 | 0.34 | 11 | 6.43 | 15 | 8.47E-08 | 4.539E-06 | (B) BRAF Class 3 Melanoma |
| TET2 | 4q24 | 70 | 8.96 | 33 | 27.97 | 103 | 6.66E-08 | 3.791E-06 | (B) BRAF Class 3 Melanoma |
| ERBB4 | 2q34 | 109 | 9.69 | 42 | 25.77 | 151 | 6.04E-08 | 3.671E-06 | (B) BRAF Class 3 Melanoma |
| PTPRD | 9p24.1-p23 | 88 | 15.71 | 36 | 43.37 | 124 | 4.36E-08 | 2.836E-06 | (B) BRAF Class 3 Melanoma |
| FGFR4 | 5q35.2 | 26 | 3.19 | 21 | 17.07 | 47 | 3.92E-08 | 2.746E-06 | (B) BRAF Class 3 Melanoma |
| KDR | 4q12 | 87 | 7.84 | 38 | 24.05 | 125 | 1.26E-08 | 9.56E-07 | (B) BRAF Class 3 Melanoma |
| ATM | 11q22.3 | 70 | 6.31 | 34 | 21.52 | 104 | 1.1E-08 | 9.08E-07 | (B) BRAF Class 3 Melanoma |
| FLT1 | 13q12.3 | 71 | 9.18 | 35 | 29.91 | 106 | 9.19E-09 | 8.81E-07 | (B) BRAF Class 3 Melanoma |
| KMT2D | 12q13.12 | 119 | 15.39 | 46 | 39.32 | 165 | 9.67E-09 | 8.81E-07 | (B) BRAF Class 3 Melanoma |
| ARID2 | 12q12 | 66 | 8.66 | 34 | 29.31 | 100 | 7.44E-09 | 8.47E-07 | (B) BRAF Class 3 Melanoma |
| NOTCH2 | 1p12 | 44 | 5.54 | 28 | 24.35 | 72 | 2.55E-09 | 3.73E-07 | (B) BRAF Class 3 Melanoma |
| SETD2 | 3p21.31 | 60 | 7.43 | 33 | 27.27 | 93 | 2.86E-09 | 3.73E-07 | (B) BRAF Class 3 Melanoma |
| KMT2A | 11q23.3 | 69 | 8.83 | 36 | 30.51 | 105 | 1.5E-09 | 2.74E-07 | (B) BRAF Class 3 Melanoma |

| | | | | | | | | | |
|---|---|---|---|---|---|---|---|---|---|
| FLT3 | 13q12.2 | 41 | 3.68 | 31 | 19.62 | 72 | 1.52E-11 | 3.46E-09 | (B) BRAF Class 3 Melanoma |
| KRAS | 12p12.1 | 5 | 0.42 | 17 | 9.94 | 22 | 3.81E-12 | 1.16E-09 | (B) BRAF Class 3 Melanoma |
| NRAS | 1p13.2 | 10 | 0.85 | 54 | 31.58 | 64 | 4.27E-42 | 1.94E-39 | (B) BRAF Class 3 Melanoma |
| NF1 | 17q11.2 | 52 | 6.41 | 79 | 64.75 | 131 | 8.93E-48 | 8.13E-45 | (B) BRAF Class 3 Melanoma |

#### Supplementary Table 11

| Gene | Cytoband | Class 1 Occurrence | Class 1 % | Class 2 Occurrence | Class 2 % | pval | qval | Most Enriched In |
|---|---|---|---|---|---|---|---|---|
| KRAS | 12p12.1 | 15 | 1.79 | 12 | 17.91 | 7.19E-08 | 5.83E-05 | (B) BRAF Class 2 Colorectal |
| RNF43 | 17q22 | 227 | 41.20 | 2 | 4.76 | 2.18E-07 | 8.83E-05 | (A) BRAF Class 1 Colorectal |
| NRAS | 1p13.2 | 14 | 1.67 | 7 | 10.45 | 0.000446 | 0.0987 | (B) BRAF Class 2 Colorectal |
| TGFBR2 | 3p24.1 | 92 | 25.56 | 0 | 0.00 | 0.000487 | 0.0987 | (A) BRAF Class 1 Colorectal |
| KMT2D | 12q13.12 | 178 | 29.82 | 4 | 8.70 | 0.000822 | 0.127 | (A) BRAF Class 1 Colorectal |
| MSH6 | 2p16.3 | 88 | 14.04 | 0 | 0.00 | 0.000941 | 0.127 | (A) BRAF Class 1 Colorectal |
| MTOR | 1p36.22 | 72 | 11.78 | 0 | 0.00 | 0.001239 | 0.143 | (A) BRAF Class 1 Colorectal |
| KMT2A | 11q23.3 | 106 | 17.52 | 1 | 2.13 | 0.001723 | 0.175 | (A) BRAF Class 1 Colorectal |
| POLD1 | 19q13.33 | 80 | 15.24 | 0 | 0.00 | 0.002041 | 0.184 | (A) BRAF Class 1 Colorectal |
| MSH3 | 5q14.1 | 76 | 29.69 | 0 | 0.00 | 0.003188 | 0.259 | (A) BRAF Class 1 Colorectal |
| APC | 5q22.2 | 234 | 29.36 | 30 | 45.45 | 0.00582 | 0.405 | (B) BRAF Class 2 Colorectal |
| NOTCH3 | 19p13.12 | 112 | 20.40 | 2 | 4.76 | 0.005995 | 0.405 | (A) BRAF Class 1 Colorectal |
| CREBBP | 16p13.3 | 111 | 18.32 | 2 | 4.35 | 0.007224 | 0.407 | (A) BRAF Class 1 Colorectal |
| GLI1 | 12q13.3 | 66 | 11.74 | 0 | 0.00 | 0.007239 | 0.407 | (A) BRAF Class 1 Colorectal |
| CIC | 19q13.2 | 87 | 16.70 | 1 | 2.50 | 0.007534 | 0.407 | (A) BRAF Class 1 Colorectal |
| JAK1 | 1p31.3 | 50 | 9.07 | 0 | 0.00 | 0.008981 | 0.455 | (A) BRAF Class 1 Colorectal |
| BRD4 | 19p13.12 | 59 | 9.98 | 0 | 0.00 | 0.009603 | 0.458 | (A) BRAF Class 1 Colorectal |
| ZFHX3 | 6q22.2-q22. | 85 | 30.80 | 1 | 5.26 | 0.0106 | 0.467 | (A) BRAF Class 1 Colorectal |
| FBXW7 | 4q31.3 | 140 | 16.77 | 3 | 5.26 | 0.0109 | 0.467 | (A) BRAF Class 1 Colorectal |
| CUX1 | 7q22.1 | 58 | 16.29 | 0 | 0.00 | 0.0139 | 0.494 | (A) BRAF Class 1 Colorectal |
| ADGRB3 | 6q12-q13 | 1 | 100.00 | 0 | 0.00 | 0.0147 | 0.494 | (A) BRAF Class 1 Colorectal |
| ETAA1 | 2p14 | 1 | 100.00 | 0 | 0.00 | 0.0147 | 0.494 | (A) BRAF Class 1 Colorectal |
| OLIG2 | 21q22.11 | 1 | 100.00 | 0 | 0.00 | 0.0147 | 0.494 | (A) BRAF Class 1 Colorectal |
| RET | 10q11.21 | 53 | 6.31 | 0 | 0.00 | 0.0151 | 0.494 | (A) BRAF Class 1 Colorectal |
| BRCA2 | 13q13.1 | 100 | 16.50 | 2 | 4.35 | 0.0152 | 0.494 | (A) BRAF Class 1 Colorectal |
| BCORL1 | Xq26.1 | 76 | 19.49 | 1 | 3.45 | 0.0179 | 0.557 | (A) BRAF Class 1 Colorectal |
| AXL | 19q13.2 | 51 | 8.42 | 0 | 0.00 | 0.0205 | 0.604 | (A) BRAF Class 1 Colorectal |
| PIK3C2B | 1q32.1 | 51 | 13.46 | 0 | 0.00 | 0.0234 | 0.604 | (A) BRAF Class 1 Colorectal |
| FAT1 | 4q35.2 | 114 | 22.62 | 3 | 8.11 | 0.0236 | 0.604 | (A) BRAF Class 1 Colorectal |
| PTPRD | 9p24.1-p23 | 71 | 17.93 | 1 | 3.33 | 0.0238 | 0.604 | (A) BRAF Class 1 Colorectal |
| SMO | 7q32.1 | 46 | 5.86 | 0 | 0.00 | 0.0245 | 0.604 | (A) BRAF Class 1 Colorectal |
| IKZF1 | 7p12.2 | 48 | 8.04 | 0 | 0.00 | 0.0246 | 0.604 | (A) BRAF Class 1 Colorectal |
| FGFR3 | 4p16.3 | 47 | 5.60 | 0 | 0.00 | 0.0246 | 0.604 | (A) BRAF Class 1 Colorectal |
| GRIN2A | 16p13.2 | 51 | 14.57 | 0 | 0.00 | 0.0262 | 0.622 | (A) BRAF Class 1 Colorectal |
| SETD2 | 3p21.31 | 71 | 11.72 | 1 | 2.17 | 0.0269 | 0.622 | (A) BRAF Class 1 Colorectal |
| MLF1 | 3q25.32 | 1 | 50.00 | 0 | 0.00 | 0.029 | 0.622 | (A) BRAF Class 1 Colorectal |
| RPL22 | 1p36.31 | 1 | 50.00 | 0 | 0.00 | 0.029 | 0.622 | (A) BRAF Class 1 Colorectal |
| NOTCH2 | 1p12 | 46 | 7.36 | 0 | 0.00 | 0.0294 | 0.622 | (A) BRAF Class 1 Colorectal |
| KMT2C | 7q36.1 | 72 | 20.99 | 1 | 4.17 | 0.0299 | 0.622 | (A) BRAF Class 1 Colorectal |
| ARAF | Xp11.3 | 47 | 7.40 | 0 | 0.00 | 0.0308 | 0.625 | (A) BRAF Class 1 Colorectal |
| ARID1A | 1p36.11 | 141 | 23.27 | 5 | 10.87 | 0.0326 | 0.644 | (A) BRAF Class 1 Colorectal |
| ATRX | Xq21.1 | 87 | 14.36 | 2 | 4.35 | 0.0351 | 0.665 | (A) BRAF Class 1 Colorectal |
| ARID1B | 6q25.3 | 107 | 18.45 | 3 | 6.98 | 0.0359 | 0.665 | (A) BRAF Class 1 Colorectal |
| ERBB3 | 12q13.2 | 39 | 6.22 | 8 | 13.79 | 0.0365 | 0.665 | (B) BRAF Class 2 Colorectal |
| FGFR4 | 5q35.2 | 36 | 5.86 | 0 | 0.00 | 0.0374 | 0.665 | (A) BRAF Class 1 Colorectal |

| | | | | | | | | |
|---|---|---|---|---|---|---|---|---|
| BLM | 15q26.1 | 68 | 11.58 | 1 | 2.33 | 0.0381 | 0.665 | (A) BRAF Class 1 Colorectal |
| GNAS | 20q13.32 | 73 | 8.74 | 1 | 1.75 | 0.0396 | 0.665 | (A) BRAF Class 1 Colorectal |
| SPEN | ›36.21-p36.: | 64 | 17.78 | 1 | 3.70 | 0.0399 | 0.665 | (A) BRAF Class 1 Colorectal |
| PDGFRB | 5q32 | 42 | 6.93 | 0 | 0.00 | 0.0417 | 0.665 | (A) BRAF Class 1 Colorectal |
| AR | Xq12 | 70 | 11.55 | 2 | 3.57 | 0.0421 | 0.665 | (A) BRAF Class 1 Colorectal |
| EGR1 | 5q31.2 | 1 | 33.33 | 0 | 0.00 | 0.0429 | 0.665 | (A) BRAF Class 1 Colorectal |
| MLLT1 | 19p13.3 | 1 | 33.33 | 0 | 0.00 | 0.0429 | 0.665 | (A) BRAF Class 1 Colorectal |
| CADM2 | 3p12.1 | 2 | 3.57 | 2 | 33.33 | 0.0434 | 0.665 | (B) BRAF Class 2 Colorectal |
| PTCH1 | 9q22.32 | 83 | 13.70 | 2 | 4.35 | 0.0448 | 0.67 | (A) BRAF Class 1 Colorectal |
| CSDE1 | 1p13.2 | 6 | 2.87 | 2 | 20.00 | 0.0455 | 0.67 | (B) BRAF Class 2 Colorectal |
| MGA | 15q15.1 | 83 | 18.99 | 2 | 6.25 | 0.0482 | 0.679 | (A) BRAF Class 1 Colorectal |
| IRS1 | 2q36.3 | 44 | 15.22 | 0 | 0.00 | 0.0485 | 0.679 | (A) BRAF Class 1 Colorectal |
| FAN1 | 15q13.3 | 9 | 5.59 | 3 | 23.08 | 0.0486 | 0.679 | (B) BRAF Class 2 Colorectal |
| DOT1L | 19p13.3 | 39 | 10.83 | 0 | 0.00 | 0.0511 | 0.702 | (A) BRAF Class 1 Colorectal |
| KDM5A | 12p13.33 | 79 | 15.16 | 2 | 5.00 | 0.0523 | 0.707 | (A) BRAF Class 1 Colorectal |
| CARD11 | 7p22.2 | 79 | 13.23 | 2 | 4.35 | 0.0532 | 0.707 | (A) BRAF Class 1 Colorectal |
| PGR | 11q22.1 | 44 | 16.24 | 0 | 0.00 | 0.0546 | 0.714 | (A) BRAF Class 1 Colorectal |
| FLT4 | 5q35.3 | 81 | 13.80 | 2 | 4.65 | 0.0588 | 0.757 | (A) BRAF Class 1 Colorectal |
| EPHA3 | 3p11.1 | 37 | 8.49 | 0 | 0.00 | 0.06 | 0.76 | (A) BRAF Class 1 Colorectal |
| PREX2 | 8q13.2 | 46 | 17.42 | 0 | 0.00 | 0.062 | 0.774 | (A) BRAF Class 1 Colorectal |
| CASP8 | 2q33.1 | 74 | 14.37 | 2 | 5.00 | 0.0674 | 0.801 | (A) BRAF Class 1 Colorectal |
| WT1 | 11p13 | 35 | 5.79 | 0 | 0.00 | 0.0677 | 0.801 | (A) BRAF Class 1 Colorectal |
| MYC | 8q24.21 | 47 | 7.50 | 1 | 1.72 | 0.0707 | 0.801 | (A) BRAF Class 1 Colorectal |
| IDH2 | 15q26.1 | 33 | 4.15 | 0 | 0.00 | 0.0712 | 0.801 | (A) BRAF Class 1 Colorectal |
| B2M | 15q21.1 | 59 | 10.55 | 1 | 2.50 | 0.0731 | 0.801 | (A) BRAF Class 1 Colorectal |
| CDK5 | 7q36.1 | 0 | 0.00 | 1 | 16.67 | 0.0732 | 0.801 | (B) BRAF Class 2 Colorectal |
| CAMKMT | 2p21 | 0 | 0.00 | 1 | 1.49 | 0.0739 | 0.801 | (B) BRAF Class 2 Colorectal |
| PAXIP1-AS2 | 7q36.2 | 0 | 0.00 | 1 | 1.49 | 0.0739 | 0.801 | (B) BRAF Class 2 Colorectal |
| KEAP1 | 19p13.2 | 33 | 5.72 | 0 | 0.00 | 0.0742 | 0.801 | (A) BRAF Class 1 Colorectal |
| ERCC2 | 19q13.32 | 36 | 6.08 | 0 | 0.00 | 0.0743 | 0.801 | (A) BRAF Class 1 Colorectal |
| CDH1 | 16q22.1 | 37 | 4.64 | 0 | 0.00 | 0.0766 | 0.801 | (A) BRAF Class 1 Colorectal |
| BCOR | Xp11.4 | 90 | 15.08 | 3 | 6.52 | 0.0769 | 0.801 | (A) BRAF Class 1 Colorectal |
| TSC1 | 9q34.13 | 34 | 5.61 | 0 | 0.00 | 0.0777 | 0.801 | (A) BRAF Class 1 Colorectal |
| LATS1 | 6q25.1 | 14 | 4.84 | 3 | 15.79 | 0.078 | 0.801 | (B) BRAF Class 2 Colorectal |
| PTPRT | 0q12-q13.1 | 57 | 17.43 | 1 | 4.35 | 0.0792 | 0.803 | (A) BRAF Class 1 Colorectal |
| TNFAIP3 | 6q23.3 | 33 | 5.53 | 0 | 0.00 | 0.0809 | 0.806 | (A) BRAF Class 1 Colorectal |
| FGFR2 | 10q26.13 | 32 | 3.81 | 0 | 0.00 | 0.0821 | 0.806 | (A) BRAF Class 1 Colorectal |
| DDR1 | 6p21.33 | 1 | 3.85 | 2 | 33.33 | 0.0827 | 0.806 | (B) BRAF Class 2 Colorectal |
| ATM | 11q22.3 | 118 | 15.03 | 4 | 7.41 | 0.0835 | 0.806 | (A) BRAF Class 1 Colorectal |
| JAK3 | 19p13.11 | 50 | 6.20 | 1 | 1.52 | 0.0869 | 0.817 | (A) BRAF Class 1 Colorectal |
| PTPRS | 19p13.3 | 78 | 26.99 | 2 | 10.53 | 0.0877 | 0.817 | (A) BRAF Class 1 Colorectal |
| RAD50 | 5q31.1 | 56 | 10.77 | 1 | 2.70 | 0.0885 | 0.817 | (A) BRAF Class 1 Colorectal |
| CTNNB1 | 3p22.1 | 28 | 3.33 | 5 | 7.46 | 0.089 | 0.817 | (B) BRAF Class 2 Colorectal |
| XPO1 | 2p15 | 32 | 5.28 | 0 | 0.00 | 0.0906 | 0.817 | (A) BRAF Class 1 Colorectal |
| FOXL2 | 3q22.3 | 32 | 6.04 | 0 | 0.00 | 0.091 | 0.817 | (A) BRAF Class 1 Colorectal |
| MALT1 | 18q21.32 | 7 | 2.30 | 2 | 10.53 | 0.0917 | 0.817 | (B) BRAF Class 2 Colorectal |
| IDH1 | 2q34 | 30 | 3.57 | 0 | 0.00 | 0.0962 | 0.848 | (A) BRAF Class 1 Colorectal |
| AMER1 | Xq11.2 | 32 | 8.89 | 5 | 18.52 | 0.102 | 0.886 | (B) BRAF Class 2 Colorectal |

| | | | | | | | | |
|---|---|---|---|---|---|---|---|---|
| COL7A1 | 3p21.31 | 31 | 19.25 | 5 | 38.46 | 0.103 | 0.886 | (B) BRAF Class 2 Colorectal |
| GATA2 | 3q21.3 | 32 | 6.15 | 0 | 0.00 | 0.104 | 0.886 | (A) BRAF Class 1 Colorectal |
| MECOM | 3q26.2 | 27 | 11.30 | 0 | 0.00 | 0.113 | 0.942 | (A) BRAF Class 1 Colorectal |
| PIK3CG | 7q22.3 | 31 | 8.86 | 0 | 0.00 | 0.117 | 0.942 | (A) BRAF Class 1 Colorectal |
| PAX5 | 9p13.2 | 28 | 4.63 | 0 | 0.00 | 0.118 | 0.942 | (A) BRAF Class 1 Colorectal |
| INSR | 19p13.2 | 32 | 9.58 | 0 | 0.00 | 0.118 | 0.942 | (A) BRAF Class 1 Colorectal |
| LATS2 | 13q12.11 | 31 | 10.47 | 0 | 0.00 | 0.118 | 0.942 | (A) BRAF Class 1 Colorectal |
| SOS1 | 2p22.1 | 27 | 9.22 | 0 | 0.00 | 0.118 | 0.942 | (A) BRAF Class 1 Colorectal |
| ARID2 | 12q12 | 68 | 11.58 | 2 | 4.65 | 0.121 | 0.942 | (A) BRAF Class 1 Colorectal |
| TET2 | 4q24 | 47 | 7.77 | 1 | 2.13 | 0.121 | 0.942 | (A) BRAF Class 1 Colorectal |
| POLE | 12q24.33 | 84 | 15.19 | 3 | 7.32 | 0.121 | 0.942 | (A) BRAF Class 1 Colorectal |
| DICER1 | 14q32.13 | 49 | 8.42 | 1 | 2.33 | 0.122 | 0.942 | (A) BRAF Class 1 Colorectal |
| NBN | 8q21.3 | 47 | 7.81 | 1 | 2.17 | 0.126 | 0.942 | (A) BRAF Class 1 Colorectal |
| PHOX2B | 4p13 | 28 | 4.96 | 0 | 0.00 | 0.128 | 0.942 | (A) BRAF Class 1 Colorectal |
| CDK2 | 12q13.2 | 1 | 1.18 | 1 | 16.67 | 0.128 | 0.942 | (B) BRAF Class 2 Colorectal |
| HNF1A | 12q24.31 | 71 | 9.10 | 2 | 3.70 | 0.129 | 0.942 | (A) BRAF Class 1 Colorectal |
| PIK3C2G | 12p12.3 | 28 | 8.83 | 0 | 0.00 | 0.129 | 0.942 | (A) BRAF Class 1 Colorectal |
| CDKN1A | 6p21.2 | 7 | 1.19 | 2 | 4.35 | 0.133 | 0.942 | (B) BRAF Class 2 Colorectal |
| ELF3 | 1q32.1 | 1 | 0.76 | 1 | 10.00 | 0.136 | 0.942 | (B) BRAF Class 2 Colorectal |
| FLCN | 17p11.2 | 45 | 7.60 | 1 | 2.17 | 0.136 | 0.942 | (A) BRAF Class 1 Colorectal |
| ALK | :p23.2-p23.: | 77 | 9.17 | 3 | 4.48 | 0.137 | 0.942 | (A) BRAF Class 1 Colorectal |
| NF1 | 17q11.2 | 66 | 10.89 | 3 | 5.36 | 0.14 | 0.942 | (A) BRAF Class 1 Colorectal |
| EP300 | 22q13.2 | 94 | 15.75 | 4 | 8.70 | 0.14 | 0.942 | (A) BRAF Class 1 Colorectal |
| TSHR | 14q31.1 | 28 | 5.19 | 0 | 0.00 | 0.142 | 0.942 | (A) BRAF Class 1 Colorectal |
| MST1R | 3p21.31 | 27 | 8.28 | 0 | 0.00 | 0.147 | 0.942 | (A) BRAF Class 1 Colorectal |
| SF3B1 | 2q33.1 | 26 | 4.04 | 0 | 0.00 | 0.148 | 0.942 | (A) BRAF Class 1 Colorectal |
| GSK3B | 3q13.33 | 8 | 2.22 | 2 | 7.41 | 0.149 | 0.942 | (B) BRAF Class 2 Colorectal |
| RAD21 | 8q24.11 | 25 | 4.49 | 0 | 0.00 | 0.15 | 0.942 | (A) BRAF Class 1 Colorectal |
| FANCF | 11p14.3 | 9 | 2.37 | 2 | 7.69 | 0.152 | 0.942 | (B) BRAF Class 2 Colorectal |
| SMAD4 | 18q21.2 | 188 | 22.51 | 9 | 15.79 | 0.154 | 0.942 | (A) BRAF Class 1 Colorectal |
| TLR4 | 9q33.1 | 1 | 1.10 | 1 | 12.50 | 0.156 | 0.942 | (B) BRAF Class 2 Colorectal |
| FANCM | 14q21.2 | 24 | 11.37 | 0 | 0.00 | 0.157 | 0.942 | (A) BRAF Class 1 Colorectal |
| TET1 | 10q21.3 | 65 | 13.43 | 2 | 5.88 | 0.157 | 0.942 | (A) BRAF Class 1 Colorectal |
| RPTOR | 17q25.3 | 43 | 8.25 | 1 | 2.50 | 0.158 | 0.942 | (A) BRAF Class 1 Colorectal |
| PIK3CD | 1p36.22 | 28 | 8.70 | 0 | 0.00 | 0.158 | 0.942 | (A) BRAF Class 1 Colorectal |
| CD274 | 9p24.1 | 8 | 1.35 | 2 | 4.35 | 0.159 | 0.942 | (B) BRAF Class 2 Colorectal |
| PIK3CA | 3q26.32 | 199 | 23.69 | 20 | 29.85 | 0.162 | 0.942 | (B) BRAF Class 2 Colorectal |
| PRDM1 | 6q21 | 24 | 4.02 | 0 | 0.00 | 0.163 | 0.942 | (A) BRAF Class 1 Colorectal |
| SOX2 | 3q26.33 | 24 | 4.02 | 0 | 0.00 | 0.163 | 0.942 | (A) BRAF Class 1 Colorectal |
| KDM5C | Xp11.22 | 41 | 7.11 | 1 | 2.17 | 0.164 | 0.942 | (A) BRAF Class 1 Colorectal |
| RB1 | 13q14.2 | 47 | 5.83 | 1 | 1.79 | 0.164 | 0.942 | (A) BRAF Class 1 Colorectal |
| EXO1 | 1q43 | 8 | 4.97 | 2 | 15.38 | 0.165 | 0.942 | (B) BRAF Class 2 Colorectal |
| MYCN | 2p24.3 | 20 | 3.30 | 0 | 0.00 | 0.166 | 0.942 | (A) BRAF Class 1 Colorectal |
| RECQL4 | 8q24.3 | 58 | 10.68 | 2 | 4.76 | 0.171 | 0.942 | (A) BRAF Class 1 Colorectal |
| REL | 2p16.1 | 23 | 3.88 | 0 | 0.00 | 0.174 | 0.942 | (A) BRAF Class 1 Colorectal |
| CYLD | 16q12.1 | 26 | 5.18 | 0 | 0.00 | 0.174 | 0.942 | (A) BRAF Class 1 Colorectal |
| AKT1 | 14q32.33 | 72 | 8.57 | 3 | 4.48 | 0.175 | 0.942 | (A) BRAF Class 1 Colorectal |
| CDC73 | 1q31.2 | 23 | 3.85 | 0 | 0.00 | 0.176 | 0.942 | (A) BRAF Class 1 Colorectal |

| | | | | | | | | |
|---|---|---|---|---|---|---|---|---|
| ASXL1 | 20q11.21 | 73 | 12.07 | 3 | 6.38 | 0.177 | 0.942 | (A) BRAF Class 1 Colorectal |
| MPL | 1p34.2 | 25 | 3.14 | 0 | 0.00 | 0.179 | 0.942 | (A) BRAF Class 1 Colorectal |
| QKI | 6q26 | 41 | 11.45 | 1 | 3.70 | 0.18 | 0.942 | (A) BRAF Class 1 Colorectal |
| ROS1 | 6q22.1 | 62 | 10.15 | 3 | 5.36 | 0.18 | 0.942 | (A) BRAF Class 1 Colorectal |
| CRTC1 | 19p13.11 | 22 | 8.12 | 0 | 0.00 | 0.181 | 0.942 | (A) BRAF Class 1 Colorectal |
| PLCG2 | 16q23.3 | 27 | 8.41 | 0 | 0.00 | 0.183 | 0.942 | (A) BRAF Class 1 Colorectal |
| SLX4 | 16p13.3 | 50 | 13.19 | 1 | 4.35 | 0.183 | 0.942 | (A) BRAF Class 1 Colorectal |
| SQSTM1 | 5q35.3 | 1 | 1.79 | 1 | 16.67 | 0.186 | 0.942 | (B) BRAF Class 2 Colorectal |
| PIK3CB | 3q22.3 | 24 | 6.70 | 0 | 0.00 | 0.188 | 0.942 | (A) BRAF Class 1 Colorectal |
| ANKRD11 | 16q24.3 | 47 | 17.34 | 1 | 5.88 | 0.19 | 0.942 | (A) BRAF Class 1 Colorectal |
| CDK8 | 13q12.13 | 22 | 4.22 | 0 | 0.00 | 0.19 | 0.942 | (A) BRAF Class 1 Colorectal |
| NCOR1 | 17p12-p11.2 | 43 | 12.50 | 1 | 4.17 | 0.191 | 0.942 | (A) BRAF Class 1 Colorectal |
| PRKDC | 8q11.21 | 79 | 20.79 | 3 | 11.54 | 0.191 | 0.942 | (A) BRAF Class 1 Colorectal |
| KAT6B | 10q22.2 | 21 | 10.77 | 0 | 0.00 | 0.194 | 0.942 | (A) BRAF Class 1 Colorectal |
| TSC2 | 16p13.3 | 56 | 9.24 | 2 | 4.35 | 0.201 | 0.942 | (A) BRAF Class 1 Colorectal |
| DAXX | 6p21.32 | 21 | 4.05 | 0 | 0.00 | 0.204 | 0.942 | (A) BRAF Class 1 Colorectal |
| NRG1 | 8p12 | 23 | 8.27 | 0 | 0.00 | 0.206 | 0.942 | (A) BRAF Class 1 Colorectal |
| PRAME | 22q11.22 | 2 | 2.63 | 1 | 16.67 | 0.206 | 0.942 | (B) BRAF Class 2 Colorectal |
| OGG1 | 3p25.3 | 2 | 1.24 | 1 | 7.69 | 0.209 | 0.942 | (B) BRAF Class 2 Colorectal |
| BAP1 | 3p21.1 | 21 | 3.47 | 0 | 0.00 | 0.21 | 0.942 | (A) BRAF Class 1 Colorectal |
| MTA1 | 14q32.33 | 19 | 11.80 | 0 | 0.00 | 0.21 | 0.942 | (A) BRAF Class 1 Colorectal |
| TFE3 | Xp11.23 | 23 | 8.16 | 0 | 0.00 | 0.21 | 0.942 | (A) BRAF Class 1 Colorectal |
| BRD3 | 9q34.2 | 20 | 10.26 | 0 | 0.00 | 0.211 | 0.942 | (A) BRAF Class 1 Colorectal |
| GATA3 | 10p14 | 38 | 6.37 | 1 | 2.17 | 0.212 | 0.942 | (A) BRAF Class 1 Colorectal |
| STK11 | 19p13.3 | 23 | 2.75 | 0 | 0.00 | 0.215 | 0.942 | (A) BRAF Class 1 Colorectal |
| CCND1 | 11q13.3 | 17 | 2.81 | 0 | 0.00 | 0.218 | 0.942 | (A) BRAF Class 1 Colorectal |
| TMPRSS2 | 21q22.3 | 20 | 3.87 | 0 | 0.00 | 0.219 | 0.942 | (A) BRAF Class 1 Colorectal |
| HRAS | 11p15.5 | 10 | 1.19 | 2 | 2.99 | 0.22 | 0.942 | (B) BRAF Class 2 Colorectal |
| MEF2B | 19p13.11 | 20 | 3.35 | 0 | 0.00 | 0.221 | 0.942 | (A) BRAF Class 1 Colorectal |
| CDKN1C | 11p15.4 | 2 | 0.83 | 1 | 4.76 | 0.223 | 0.942 | (B) BRAF Class 2 Colorectal |
| ERBB4 | 2q34 | 81 | 9.95 | 4 | 6.15 | 0.224 | 0.942 | (A) BRAF Class 1 Colorectal |
| KDM6A | Xp11.3 | 37 | 6.20 | 1 | 2.17 | 0.225 | 0.942 | (A) BRAF Class 1 Colorectal |
| ERCC4 | 16p13.12 | 20 | 3.43 | 0 | 0.00 | 0.228 | 0.942 | (A) BRAF Class 1 Colorectal |
| SMARCE1 | 17q21.2 | 2 | 1.20 | 1 | 6.67 | 0.23 | 0.942 | (B) BRAF Class 2 Colorectal |
| CDKN2B | 9p21.3 | 11 | 1.82 | 2 | 4.35 | 0.232 | 0.942 | (B) BRAF Class 2 Colorectal |
| CTNNA1 | 5q31.2 | 20 | 6.37 | 0 | 0.00 | 0.233 | 0.942 | (A) BRAF Class 1 Colorectal |
| RPS6KB2 | 11q13.2 | 13 | 4.50 | 2 | 10.53 | 0.234 | 0.942 | (B) BRAF Class 2 Colorectal |
| SHQ1 | 3p13 | 13 | 4.50 | 2 | 10.53 | 0.234 | 0.942 | (B) BRAF Class 2 Colorectal |
| PIK3R2 | 19p13.11 | 21 | 6.00 | 0 | 0.00 | 0.239 | 0.942 | (A) BRAF Class 1 Colorectal |
| PIK3C3 | 18q12.3 | 22 | 7.14 | 0 | 0.00 | 0.239 | 0.942 | (A) BRAF Class 1 Colorectal |
| IRF2 | 4q35.1 | 4 | 2.84 | 1 | 12.50 | 0.244 | 0.942 | (B) BRAF Class 2 Colorectal |
| HLA-B | 6p21.33 | 21 | 15.00 | 0 | 0.00 | 0.244 | 0.942 | (A) BRAF Class 1 Colorectal |
| AXIN2 | 17q24.1 | 53 | 10.88 | 2 | 5.56 | 0.245 | 0.942 | (A) BRAF Class 1 Colorectal |
| RICTOR | 5p13.1 | 36 | 6.79 | 1 | 2.50 | 0.247 | 0.942 | (A) BRAF Class 1 Colorectal |
| DDB1 | 11q12.2 | 17 | 10.56 | 0 | 0.00 | 0.25 | 0.942 | (A) BRAF Class 1 Colorectal |
| RBBP8 | 18q11.2 | 17 | 10.56 | 0 | 0.00 | 0.25 | 0.942 | (A) BRAF Class 1 Colorectal |
| DNMT3B | 20q11.21 | 21 | 7.27 | 0 | 0.00 | 0.251 | 0.942 | (A) BRAF Class 1 Colorectal |
| CD58 | 1p13.1 | 3 | 2.78 | 1 | 12.50 | 0.252 | 0.942 | (B) BRAF Class 2 Colorectal |

| | | | | | | | | |
|---|---|---|---|---|---|---|---|---|
| MLH3 | 14q24.3 | 16 | 8.04 | 0 | 0.00 | 0.256 | 0.942 | (A) BRAF Class 1 Colorectal |
| SMARCA4 | 19p13.2 | 81 | 13.37 | 4 | 8.70 | 0.257 | 0.942 | (A) BRAF Class 1 Colorectal |
| SDHA | 5p15.33 | 33 | 5.80 | 1 | 2.17 | 0.258 | 0.942 | (A) BRAF Class 1 Colorectal |
| CBL | 11q23.3 | 18 | 2.93 | 0 | 0.00 | 0.26 | 0.942 | (A) BRAF Class 1 Colorectal |
| MED12 | Xq13.1 | 53 | 10.00 | 3 | 6.00 | 0.264 | 0.942 | (A) BRAF Class 1 Colorectal |
| CRTC2 | 1q21.3 | 3 | 3.95 | 1 | 16.67 | 0.266 | 0.942 | (B) BRAF Class 2 Colorectal |
| BRCA1 | 17q21.31 | 54 | 8.91 | 3 | 5.36 | 0.267 | 0.942 | (A) BRAF Class 1 Colorectal |
| MEN1 | 11q13.1 | 34 | 5.70 | 1 | 2.17 | 0.267 | 0.942 | (A) BRAF Class 1 Colorectal |
| NKX2-1 | 14q13.3 | 34 | 5.70 | 1 | 2.17 | 0.267 | 0.942 | (A) BRAF Class 1 Colorectal |
| RPS6KA4 | 11q13.1 | 20 | 6.92 | 0 | 0.00 | 0.268 | 0.942 | (A) BRAF Class 1 Colorectal |
| FOXP1 | 3p13 | 19 | 5.51 | 0 | 0.00 | 0.269 | 0.942 | (A) BRAF Class 1 Colorectal |
| KDM6B | 17p13.1 | 12 | 15.79 | 2 | 33.33 | 0.271 | 0.942 | (B) BRAF Class 2 Colorectal |
| PDK1 | 2q31.1 | 4 | 2.61 | 1 | 10.00 | 0.274 | 0.942 | (B) BRAF Class 2 Colorectal |
| ETV1 | 7p21.2 | 36 | 6.23 | 1 | 2.44 | 0.278 | 0.942 | (A) BRAF Class 1 Colorectal |
| RSPO3 | 6q22.33 | 15 | 7.58 | 0 | 0.00 | 0.278 | 0.942 | (A) BRAF Class 1 Colorectal |
| SOCS1 | 16p13.13 | 17 | 2.85 | 0 | 0.00 | 0.278 | 0.942 | (A) BRAF Class 1 Colorectal |
| GRM3 | ׀21.11-q21.: | 18 | 12.33 | 0 | 0.00 | 0.282 | 0.942 | (A) BRAF Class 1 Colorectal |
| CTLA4 | 2q33.2 | 18 | 3.99 | 0 | 0.00 | 0.285 | 0.942 | (A) BRAF Class 1 Colorectal |
| DNMT1 | 19p13.2 | 35 | 10.80 | 1 | 4.35 | 0.285 | 0.942 | (A) BRAF Class 1 Colorectal |
| RAD52 | 12p13.33 | 17 | 3.56 | 0 | 0.00 | 0.285 | 0.942 | (A) BRAF Class 1 Colorectal |
| FANCA | 16q24.3 | 33 | 5.45 | 1 | 2.17 | 0.29 | 0.942 | (A) BRAF Class 1 Colorectal |
| SMARCD1 | 12q13.12 | 19 | 6.19 | 0 | 0.00 | 0.291 | 0.942 | (A) BRAF Class 1 Colorectal |
| BCL11B | 14q32.2 | 13 | 6.67 | 2 | 13.33 | 0.291 | 0.942 | (B) BRAF Class 2 Colorectal |
| PBRM1 | 3p21.1 | 60 | 10.40 | 3 | 6.52 | 0.293 | 0.942 | (A) BRAF Class 1 Colorectal |
| ABL1 | 9q34.12 | 52 | 6.52 | 2 | 3.57 | 0.294 | 0.942 | (A) BRAF Class 1 Colorectal |
| EGFR | 7p11.2 | 46 | 5.48 | 2 | 2.99 | 0.294 | 0.942 | (A) BRAF Class 1 Colorectal |
| NSD1 | 5q35.3 | 50 | 9.78 | 2 | 5.41 | 0.296 | 0.942 | (A) BRAF Class 1 Colorectal |
| CDH4 | 20q13.33 | 15 | 9.32 | 0 | 0.00 | 0.296 | 0.942 | (A) BRAF Class 1 Colorectal |
| NEIL3 | 4q34.3 | 15 | 9.32 | 0 | 0.00 | 0.296 | 0.942 | (A) BRAF Class 1 Colorectal |
| AXIN1 | 16p13.3 | 32 | 9.04 | 1 | 3.70 | 0.298 | 0.942 | (A) BRAF Class 1 Colorectal |
| FGFR1 | 8p11.23 | 31 | 3.69 | 1 | 1.49 | 0.3 | 0.942 | (A) BRAF Class 1 Colorectal |
| FUBP1 | 1p31.1 | 16 | 4.52 | 0 | 0.00 | 0.301 | 0.942 | (A) BRAF Class 1 Colorectal |
| GATA4 | 8p23.1 | 17 | 4.50 | 0 | 0.00 | 0.302 | 0.942 | (A) BRAF Class 1 Colorectal |
| GREM1 | 15q13.3 | 4 | 0.80 | 1 | 2.70 | 0.302 | 0.942 | (B) BRAF Class 2 Colorectal |
| U2AF1 | 21q22.3 | 16 | 2.66 | 0 | 0.00 | 0.303 | 0.942 | (A) BRAF Class 1 Colorectal |
| MUTYH | 1p34.1 | 24 | 4.02 | 3 | 6.52 | 0.303 | 0.942 | (B) BRAF Class 2 Colorectal |
| ERCC3 | 2q14.3 | 17 | 3.04 | 0 | 0.00 | 0.304 | 0.942 | (A) BRAF Class 1 Colorectal |
| MYCL | 1p34.2 | 16 | 2.64 | 0 | 0.00 | 0.306 | 0.942 | (A) BRAF Class 1 Colorectal |
| STAG1 | 3q22.3 | 21 | 13.29 | 2 | 25.00 | 0.306 | 0.942 | (B) BRAF Class 2 Colorectal |
| PPARG | 3p25.2 | 17 | 3.87 | 0 | 0.00 | 0.307 | 0.942 | (A) BRAF Class 1 Colorectal |
| PPP2R1A | 19q13.41 | 16 | 3.02 | 0 | 0.00 | 0.307 | 0.942 | (A) BRAF Class 1 Colorectal |
| KAT6A | 8p11.21 | 36 | 11.84 | 1 | 5.00 | 0.307 | 0.942 | (A) BRAF Class 1 Colorectal |
| RUNX1T1 | 8q21.3 | 18 | 5.92 | 0 | 0.00 | 0.308 | 0.942 | (A) BRAF Class 1 Colorectal |
| IL7R | 5p13.2 | 33 | 6.46 | 1 | 2.70 | 0.312 | 0.942 | (A) BRAF Class 1 Colorectal |
| FOXO1 | 13q14.11 | 17 | 5.48 | 0 | 0.00 | 0.319 | 0.942 | (A) BRAF Class 1 Colorectal |
| HLA-A | 6p22.1 | 37 | 12.80 | 1 | 5.56 | 0.32 | 0.942 | (A) BRAF Class 1 Colorectal |
| CHD4 | 12p13.31 | 21 | 15.44 | 0 | 0.00 | 0.32 | 0.942 | (A) BRAF Class 1 Colorectal |
| MAP2K1 | 15q22.31 | 12 | 1.82 | 2 | 3.39 | 0.321 | 0.942 | (B) BRAF Class 2 Colorectal |

| | | | | | | | | |
|---|---|---|---|---|---|---|---|---|
| SLC34A2 | 4p15.2 | 13 | 6.37 | 0 | 0.00 | 0.323 | 0.942 | (A) BRAF Class 1 Colorectal |
| KIT | 4q12 | 44 | 5.24 | 2 | 2.99 | 0.323 | 0.942 | (A) BRAF Class 1 Colorectal |
| RANBP2 | 2q13 | 27 | 20.61 | 0 | 0.00 | 0.325 | 0.942 | (A) BRAF Class 1 Colorectal |
| TOP1 | 20q12 | 17 | 4.74 | 0 | 0.00 | 0.325 | 0.942 | (A) BRAF Class 1 Colorectal |
| INPP4B | 4q31.21 | 15 | 4.24 | 0 | 0.00 | 0.325 | 0.942 | (A) BRAF Class 1 Colorectal |
| POLB | 8p11.21 | 4 | 2.48 | 1 | 7.69 | 0.325 | 0.942 | (B) BRAF Class 2 Colorectal |
| NSD2 | 4p16.3 | 28 | 8.31 | 1 | 3.57 | 0.325 | 0.942 | (A) BRAF Class 1 Colorectal |
| ZNRF3 | 22q12.1 | 28 | 14.29 | 1 | 6.25 | 0.325 | 0.942 | (A) BRAF Class 1 Colorectal |
| PTPN11 | 12q24.13 | 17 | 2.09 | 0 | 0.00 | 0.326 | 0.942 | (A) BRAF Class 1 Colorectal |
| PLK2 | 5q11.2 | 17 | 5.88 | 0 | 0.00 | 0.329 | 0.942 | (A) BRAF Class 1 Colorectal |
| FANCE | 6p21.31 | 5 | 1.32 | 1 | 3.85 | 0.329 | 0.942 | (B) BRAF Class 2 Colorectal |
| EWSR1 | 22q12.2 | 15 | 3.89 | 0 | 0.00 | 0.331 | 0.942 | (A) BRAF Class 1 Colorectal |
| FGF4 | 11q13.3 | 15 | 4.17 | 0 | 0.00 | 0.331 | 0.942 | (A) BRAF Class 1 Colorectal |
| PRF1 | 10q22.1 | 14 | 5.88 | 0 | 0.00 | 0.331 | 0.942 | (A) BRAF Class 1 Colorectal |
| BCL6 | 3q27.3 | 30 | 5.03 | 1 | 2.17 | 0.332 | 0.942 | (A) BRAF Class 1 Colorectal |
| ERCC5 | 13q33.1 | 48 | 8.57 | 2 | 5.00 | 0.332 | 0.942 | (A) BRAF Class 1 Colorectal |
| CTCF | 16q22.1 | 30 | 5.76 | 1 | 2.50 | 0.333 | 0.942 | (A) BRAF Class 1 Colorectal |
| CYSLTR2 | 13q14.2 | 5 | 3.14 | 1 | 9.09 | 0.335 | 0.942 | (B) BRAF Class 2 Colorectal |
| ABCB11 | 2q31.1 | 12 | 7.32 | 0 | 0.00 | 0.338 | 0.942 | (A) BRAF Class 1 Colorectal |
| FOXA1 | 14q21.1 | 15 | 3.07 | 0 | 0.00 | 0.339 | 0.942 | (A) BRAF Class 1 Colorectal |
| TOP2A | 17q21.2 | 20 | 14.71 | 0 | 0.00 | 0.34 | 0.942 | (A) BRAF Class 1 Colorectal |
| NOTCH1 | 9q34.3 | 96 | 11.84 | 5 | 8.93 | 0.345 | 0.942 | (A) BRAF Class 1 Colorectal |
| MAP3K13 | 3q27.2 | 5 | 1.58 | 1 | 4.35 | 0.345 | 0.942 | (B) BRAF Class 2 Colorectal |
| DOCK8 | 9p24.3 | 27 | 16.77 | 1 | 7.69 | 0.347 | 0.942 | (A) BRAF Class 1 Colorectal |
| RBM10 | Xp11.3 | 43 | 8.37 | 2 | 5.00 | 0.35 | 0.942 | (A) BRAF Class 1 Colorectal |
| INHBA | 7p14.1 | 16 | 4.82 | 0 | 0.00 | 0.35 | 0.942 | (A) BRAF Class 1 Colorectal |
| KCNQ1 | 1p15.5-p15. | 13 | 8.07 | 0 | 0.00 | 0.351 | 0.942 | (A) BRAF Class 1 Colorectal |
| AKT3 | 1q43-q44 | 14 | 2.23 | 0 | 0.00 | 0.352 | 0.942 | (A) BRAF Class 1 Colorectal |
| NF2 | 22q12.2 | 14 | 2.23 | 0 | 0.00 | 0.352 | 0.942 | (A) BRAF Class 1 Colorectal |
| FANCI | 15q26.1 | 20 | 6.62 | 2 | 11.11 | 0.356 | 0.942 | (B) BRAF Class 2 Colorectal |
| NTRK2 | 9q21.33 | 27 | 4.46 | 3 | 6.52 | 0.356 | 0.942 | (B) BRAF Class 2 Colorectal |
| IRF4 | 6p25.3 | 14 | 3.89 | 0 | 0.00 | 0.357 | 0.942 | (A) BRAF Class 1 Colorectal |
| MAP2K4 | 17p12 | 27 | 4.46 | 3 | 6.52 | 0.357 | 0.942 | (B) BRAF Class 2 Colorectal |
| FLT1 | 13q12.3 | 43 | 7.20 | 2 | 4.35 | 0.358 | 0.942 | (A) BRAF Class 1 Colorectal |
| VHL | 3p25.3 | 15 | 1.88 | 0 | 0.00 | 0.358 | 0.942 | (A) BRAF Class 1 Colorectal |
| YAP1 | 11q22.1 | 14 | 3.08 | 0 | 0.00 | 0.359 | 0.942 | (A) BRAF Class 1 Colorectal |
| MTAP | 9p21.3 | 4 | 2.33 | 1 | 6.25 | 0.362 | 0.942 | (B) BRAF Class 2 Colorectal |
| MSH2 | 2p21-p16.3 | 45 | 7.46 | 2 | 4.55 | 0.363 | 0.942 | (A) BRAF Class 1 Colorectal |
| RARA | 17q21.2 | 27 | 4.52 | 3 | 6.52 | 0.365 | 0.942 | (B) BRAF Class 2 Colorectal |
| NSD3 | 8p11.23 | 15 | 5.02 | 2 | 8.33 | 0.365 | 0.942 | (B) BRAF Class 2 Colorectal |
| NTRK3 | 15q25.3 | 43 | 7.10 | 2 | 4.35 | 0.367 | 0.942 | (A) BRAF Class 1 Colorectal |
| GNA11 | 19p13.3 | 13 | 1.58 | 0 | 0.00 | 0.37 | 0.942 | (A) BRAF Class 1 Colorectal |
| GNAQ | 9q21.2 | 13 | 1.57 | 0 | 0.00 | 0.371 | 0.942 | (A) BRAF Class 1 Colorectal |
| GLI3 | 7p14.1 | 12 | 15.79 | 0 | 0.00 | 0.374 | 0.942 | (A) BRAF Class 1 Colorectal |
| INPP4A | 2q11.2 | 15 | 5.19 | 0 | 0.00 | 0.376 | 0.942 | (A) BRAF Class 1 Colorectal |
| FANCD2 | 3p25.3 | 20 | 5.21 | 2 | 8.33 | 0.377 | 0.942 | (B) BRAF Class 2 Colorectal |
| HIST1H1C | 6p22.2 | 15 | 4.67 | 0 | 0.00 | 0.379 | 0.942 | (A) BRAF Class 1 Colorectal |
| ADGRA2 | 8p11.23 | 20 | 18.02 | 0 | 0.00 | 0.381 | 0.942 | (A) BRAF Class 1 Colorectal |

| | | | | | | | | |
|---|---|---|---|---|---|---|---|---|
| RHBDF2 | 17q25.1 | 12 | 7.45 | 0 | 0.00 | 0.381 | 0.942 | (A) BRAF Class 1 Colorectal |
| CCND2 | 12p13.32 | 13 | 2.15 | 0 | 0.00 | 0.383 | 0.942 | (A) BRAF Class 1 Colorectal |
| CBFA2T3 | 16q24.3 | 12 | 7.41 | 0 | 0.00 | 0.384 | 0.942 | (A) BRAF Class 1 Colorectal |
| TLX3 | 5q35.1 | 12 | 7.41 | 0 | 0.00 | 0.384 | 0.942 | (A) BRAF Class 1 Colorectal |
| ZNF217 | 20q13.2 | 29 | 7.44 | 3 | 10.34 | 0.385 | 0.942 | (B) BRAF Class 2 Colorectal |
| SMAD2 | 18q21.1 | 28 | 4.69 | 3 | 6.52 | 0.385 | 0.942 | (B) BRAF Class 2 Colorectal |
| ERG | 21q22.2 | 27 | 5.18 | 1 | 2.50 | 0.39 | 0.942 | (A) BRAF Class 1 Colorectal |
| TP53BP1 | 15q15.3 | 28 | 9.30 | 1 | 4.55 | 0.391 | 0.942 | (A) BRAF Class 1 Colorectal |
| CRLF2 | 2.33 and Yp1 | 13 | 2.22 | 0 | 0.00 | 0.394 | 0.942 | (A) BRAF Class 1 Colorectal |
| PIK3R1 | 5q13.1 | 41 | 6.54 | 2 | 4.17 | 0.394 | 0.942 | (A) BRAF Class 1 Colorectal |
| ANKRD26 | 10p12.1 | 31 | 37.80 | 0 | 0.00 | 0.395 | 0.942 | (A) BRAF Class 1 Colorectal |
| DROSHA | 5p13.3 | 12 | 9.23 | 0 | 0.00 | 0.395 | 0.942 | (A) BRAF Class 1 Colorectal |
| CHEK2 | 22q12.1 | 29 | 4.79 | 3 | 6.52 | 0.396 | 0.942 | (B) BRAF Class 2 Colorectal |
| IGF1R | 15q26.3 | 53 | 8.63 | 5 | 10.64 | 0.397 | 0.942 | (B) BRAF Class 2 Colorectal |
| TBX3 | 12q24.21 | 27 | 7.54 | 1 | 3.70 | 0.397 | 0.942 | (A) BRAF Class 1 Colorectal |
| SLITRK6 | 13q31.1 | 4 | 6.90 | 1 | 16.67 | 0.399 | 0.942 | (B) BRAF Class 2 Colorectal |
| PML | 15q24.1 | 12 | 6.15 | 0 | 0.00 | 0.401 | 0.942 | (A) BRAF Class 1 Colorectal |
| FGF14 | 13q33.1 | 7 | 4.58 | 1 | 10.00 | 0.404 | 0.942 | (B) BRAF Class 2 Colorectal |
| TP63 | 3q28 | 14 | 4.36 | 0 | 0.00 | 0.405 | 0.942 | (A) BRAF Class 1 Colorectal |
| CALR | 19p13.13 | 12 | 2.49 | 0 | 0.00 | 0.407 | 0.942 | (A) BRAF Class 1 Colorectal |
| DIS3L2 | 2q37.1 | 10 | 6.10 | 0 | 0.00 | 0.407 | 0.942 | (A) BRAF Class 1 Colorectal |
| SUFU | 10q24.32 | 26 | 4.36 | 1 | 2.17 | 0.41 | 0.942 | (A) BRAF Class 1 Colorectal |
| EXT1 | 8q24.11 | 6 | 2.21 | 1 | 4.76 | 0.41 | 0.942 | (B) BRAF Class 2 Colorectal |
| CCND3 | 6p21.1 | 12 | 1.98 | 0 | 0.00 | 0.412 | 0.942 | (A) BRAF Class 1 Colorectal |
| SPTA1 | 1q23.1 | 26 | 19.12 | 2 | 28.57 | 0.413 | 0.942 | (B) BRAF Class 2 Colorectal |
| ARHGAP35 | 19q13.32 | 10 | 5.99 | 0 | 0.00 | 0.413 | 0.942 | (A) BRAF Class 1 Colorectal |
| FAAP100 | 17q25.3 | 11 | 6.83 | 0 | 0.00 | 0.414 | 0.942 | (A) BRAF Class 1 Colorectal |
| LIG4 | 13q33.3 | 11 | 6.83 | 0 | 0.00 | 0.414 | 0.942 | (A) BRAF Class 1 Colorectal |
| PTPN14 | 1q32.3-q41 | 11 | 6.83 | 0 | 0.00 | 0.414 | 0.942 | (A) BRAF Class 1 Colorectal |
| PTK2B | 8p21.2 | 11 | 6.75 | 0 | 0.00 | 0.419 | 0.942 | (A) BRAF Class 1 Colorectal |
| KLF4 | 9q31.2 | 12 | 2.46 | 0 | 0.00 | 0.422 | 0.942 | (A) BRAF Class 1 Colorectal |
| ASXL2 | 2p23.3 | 27 | 9.18 | 1 | 4.76 | 0.423 | 0.942 | (A) BRAF Class 1 Colorectal |
| XRCC3 | 14q32.33 | 11 | 6.15 | 0 | 0.00 | 0.427 | 0.942 | (A) BRAF Class 1 Colorectal |
| PMS1 | 2q32.2 | 26 | 4.60 | 1 | 2.38 | 0.428 | 0.942 | (A) BRAF Class 1 Colorectal |
| ICOSLG | 21q22.3 | 13 | 4.50 | 0 | 0.00 | 0.43 | 0.942 | (A) BRAF Class 1 Colorectal |
| IFNGR1 | 6q23.3 | 13 | 4.50 | 0 | 0.00 | 0.43 | 0.942 | (A) BRAF Class 1 Colorectal |
| TGFBR1 | 9q22.33 | 13 | 4.50 | 0 | 0.00 | 0.43 | 0.942 | (A) BRAF Class 1 Colorectal |
| TCF7L2 | 0q25.2-q25. | 58 | 10.60 | 5 | 12.50 | 0.433 | 0.942 | (B) BRAF Class 2 Colorectal |
| RTEL1 | 20q13.33 | 8 | 6.30 | 1 | 12.50 | 0.433 | 0.942 | (B) BRAF Class 2 Colorectal |
| TP53 | 17p13.1 | 498 | 59.29 | 35 | 61.40 | 0.433 | 0.942 | (B) BRAF Class 2 Colorectal |
| RAC1 | 7p22.1 | 7 | 1.25 | 1 | 2.44 | 0.434 | 0.942 | (B) BRAF Class 2 Colorectal |
| KDR | 4q12 | 48 | 6.11 | 4 | 7.41 | 0.435 | 0.942 | (B) BRAF Class 2 Colorectal |
| NOTCH4 | 6p21.32 | 44 | 12.19 | 2 | 8.33 | 0.435 | 0.942 | (A) BRAF Class 1 Colorectal |
| RECQL | 12p12.1 | 13 | 10.24 | 0 | 0.00 | 0.435 | 0.942 | (A) BRAF Class 1 Colorectal |
| NEGR1 | 1p31.1 | 12 | 3.67 | 0 | 0.00 | 0.436 | 0.942 | (A) BRAF Class 1 Colorectal |
| ABRAXAS1 | 4q21.23 | 12 | 2.42 | 0 | 0.00 | 0.437 | 0.942 | (A) BRAF Class 1 Colorectal |
| GATA1 | Xp11.23 | 12 | 3.23 | 0 | 0.00 | 0.438 | 0.942 | (A) BRAF Class 1 Colorectal |
| MAP2K2 | 19p13.3 | 9 | 1.70 | 0 | 0.00 | 0.442 | 0.942 | (A) BRAF Class 1 Colorectal |

| | | | | | | | | |
|---|---|---|---|---|---|---|---|---|
| RAD51D | 17q12 | 11 | 2.10 | 0 | 0.00 | 0.443 | 0.942 | (A) BRAF Class 1 Colorectal |
| PRKCI | 3q26.2 | 27 | 5.35 | 1 | 2.86 | 0.444 | 0.942 | (A) BRAF Class 1 Colorectal |
| CDK6 | 7q21.2 | 11 | 1.82 | 0 | 0.00 | 0.444 | 0.942 | (A) BRAF Class 1 Colorectal |
| RAF1 | 3p25.2 | 21 | 3.31 | 1 | 1.75 | 0.445 | 0.942 | (A) BRAF Class 1 Colorectal |
| MAPK3 | 16p11.2 | 13 | 4.80 | 0 | 0.00 | 0.446 | 0.942 | (A) BRAF Class 1 Colorectal |
| PMS2 | 7p22.1 | 19 | 3.15 | 2 | 4.35 | 0.446 | 0.942 | (B) BRAF Class 2 Colorectal |
| ELANE | 19p13.3 | 9 | 5.49 | 0 | 0.00 | 0.446 | 0.942 | (A) BRAF Class 1 Colorectal |
| FH | 1q43 | 24 | 4.05 | 1 | 2.17 | 0.449 | 0.942 | (A) BRAF Class 1 Colorectal |
| CDK4 | 12q14.1 | 9 | 1.49 | 0 | 0.00 | 0.449 | 0.942 | (A) BRAF Class 1 Colorectal |
| PRKAR1A | 17q24.2 | 7 | 1.17 | 1 | 2.17 | 0.45 | 0.942 | (B) BRAF Class 2 Colorectal |
| GLI2 | 2q14.2 | 33 | 13.64 | 2 | 9.52 | 0.45 | 0.942 | (A) BRAF Class 1 Colorectal |
| KIF1B | 1p36.22 | 10 | 6.21 | 0 | 0.00 | 0.45 | 0.942 | (A) BRAF Class 1 Colorectal |
| SERPINA1 | 14q32.13 | 10 | 6.21 | 0 | 0.00 | 0.45 | 0.942 | (A) BRAF Class 1 Colorectal |
| SETBP1 | 18q12.3 | 41 | 11.45 | 2 | 8.00 | 0.45 | 0.942 | (A) BRAF Class 1 Colorectal |
| SLIT2 | 4p15.31 | 15 | 11.03 | 0 | 0.00 | 0.452 | 0.942 | (A) BRAF Class 1 Colorectal |
| PTEN | 10q23.31 | 136 | 16.19 | 10 | 17.54 | 0.453 | 0.942 | (B) BRAF Class 2 Colorectal |
| CDK12 | 17q12 | 38 | 7.17 | 2 | 5.00 | 0.454 | 0.942 | (A) BRAF Class 1 Colorectal |
| BARD1 | 2q35 | 24 | 4.61 | 1 | 2.50 | 0.454 | 0.942 | (A) BRAF Class 1 Colorectal |
| BABAM1 | 19p13.11 | 11 | 3.82 | 0 | 0.00 | 0.455 | 0.942 | (A) BRAF Class 1 Colorectal |
| SOX17 | 8q11.23 | 43 | 14.88 | 2 | 10.53 | 0.455 | 0.942 | (A) BRAF Class 1 Colorectal |
| RAD54L | 1p34.1 | 11 | 3.47 | 0 | 0.00 | 0.457 | 0.942 | (A) BRAF Class 1 Colorectal |
| EPHA7 | 6q16.1 | 35 | 8.71 | 3 | 10.71 | 0.459 | 0.942 | (B) BRAF Class 2 Colorectal |
| PGBD3 | 10q11.23 | 7 | 0.83 | 1 | 1.49 | 0.46 | 0.942 | (B) BRAF Class 2 Colorectal |
| ESR1 | ;q25.1-q25.2 | 31 | 4.91 | 2 | 3.45 | 0.463 | 0.942 | (A) BRAF Class 1 Colorectal |
| KMT2B | 19q13.12 | 66 | 26.83 | 3 | 21.43 | 0.465 | 0.942 | (A) BRAF Class 1 Colorectal |
| RXRA | 9q34.2 | 12 | 9.45 | 0 | 0.00 | 0.465 | 0.942 | (A) BRAF Class 1 Colorectal |
| PARP1 | 1q42.12 | 25 | 7.14 | 1 | 4.00 | 0.468 | 0.942 | (A) BRAF Class 1 Colorectal |
| NCOA3 | 20q13.12 | 26 | 8.44 | 1 | 4.76 | 0.47 | 0.942 | (A) BRAF Class 1 Colorectal |
| AKT2 | 19q13.2 | 20 | 3.19 | 2 | 4.17 | 0.473 | 0.942 | (B) BRAF Class 2 Colorectal |
| HOXB13 | 17q21.32 | 8 | 1.77 | 1 | 3.03 | 0.473 | 0.942 | (B) BRAF Class 2 Colorectal |
| AURKB | 17p13.1 | 10 | 1.68 | 0 | 0.00 | 0.473 | 0.942 | (A) BRAF Class 1 Colorectal |
| FANCL | 2p16.1 | 8 | 2.55 | 1 | 4.35 | 0.475 | 0.942 | (B) BRAF Class 2 Colorectal |
| MYBL1 | 8q13.1 | 21 | 7.66 | 1 | 4.35 | 0.475 | 0.942 | (A) BRAF Class 1 Colorectal |
| ROBO1 | 3p12.3 | 2 | 40.00 | 0 | 0.00 | 0.476 | 0.942 | (A) BRAF Class 1 Colorectal |
| ZFHX4 | 8q21.13 | 2 | 40.00 | 0 | 0.00 | 0.476 | 0.942 | (A) BRAF Class 1 Colorectal |
| BCL10 | 1p22.3 | 12 | 3.95 | 0 | 0.00 | 0.477 | 0.942 | (A) BRAF Class 1 Colorectal |
| RAD51C | 17q22 | 10 | 1.90 | 0 | 0.00 | 0.477 | 0.942 | (A) BRAF Class 1 Colorectal |
| SMAD3 | 15q22.33 | 22 | 6.41 | 2 | 8.33 | 0.477 | 0.942 | (B) BRAF Class 2 Colorectal |
| LYN | 8q12.1 | 10 | 3.73 | 1 | 6.25 | 0.478 | 0.942 | (B) BRAF Class 2 Colorectal |
| DNAJB1 | 19p13.12 | 11 | 3.57 | 0 | 0.00 | 0.478 | 0.942 | (A) BRAF Class 1 Colorectal |
| DIS3 | 13q21.33 | 35 | 6.25 | 2 | 4.55 | 0.483 | 0.942 | (A) BRAF Class 1 Colorectal |
| SYK | 9q22.2 | 23 | 5.28 | 1 | 3.03 | 0.484 | 0.942 | (A) BRAF Class 1 Colorectal |
| TCF3 | 19p13.3 | 39 | 7.20 | 2 | 5.26 | 0.485 | 0.942 | (A) BRAF Class 1 Colorectal |
| TEK | 9p21.2 | 8 | 5.23 | 0 | 0.00 | 0.489 | 0.942 | (A) BRAF Class 1 Colorectal |
| ENG | 9q34.11 | 9 | 5.59 | 0 | 0.00 | 0.489 | 0.942 | (A) BRAF Class 1 Colorectal |
| BTK | Xq22.1 | 10 | 2.71 | 0 | 0.00 | 0.489 | 0.942 | (A) BRAF Class 1 Colorectal |
| FGF3 | 11q13.3 | 10 | 2.71 | 0 | 0.00 | 0.489 | 0.942 | (A) BRAF Class 1 Colorectal |
| CIITA | 16p13.13 | 22 | 8.15 | 1 | 4.76 | 0.49 | 0.942 | (A) BRAF Class 1 Colorectal |

| | | | | | | | | |
|---|---|---|---|---|---|---|---|---|
| PALB2 | 16p12.2 | 49 | 8.09 | 3 | 6.52 | 0.49 | 0.942 | (A) BRAF Class 1 Colorectal |
| ERCC6 | 10q11.23 | 21 | 13.04 | 1 | 7.69 | 0.49 | 0.942 | (A) BRAF Class 1 Colorectal |
| YES1 | 18p11.32 | 11 | 3.81 | 0 | 0.00 | 0.49 | 0.942 | (A) BRAF Class 1 Colorectal |
| CHEK1 | 11q24.2 | 21 | 3.96 | 2 | 5.00 | 0.491 | 0.942 | (B) BRAF Class 2 Colorectal |
| ACVR1 | 2q24.1 | 10 | 2.13 | 0 | 0.00 | 0.493 | 0.942 | (A) BRAF Class 1 Colorectal |
| GALNT12 | 9q22.33 | 8 | 4.52 | 0 | 0.00 | 0.494 | 0.942 | (A) BRAF Class 1 Colorectal |
| NFKBIZ | 3q12.3 | 9 | 3.80 | 0 | 0.00 | 0.494 | 0.942 | (A) BRAF Class 1 Colorectal |
| BUB1B | 15q15.1 | 8 | 2.95 | 1 | 4.76 | 0.494 | 0.942 | (B) BRAF Class 2 Colorectal |
| EZH2 | 7q36.1 | 9 | 1.21 | 1 | 1.92 | 0.495 | 0.942 | (B) BRAF Class 2 Colorectal |
| EPAS1 | 2p21 | 11 | 8.66 | 0 | 0.00 | 0.497 | 0.942 | (A) BRAF Class 1 Colorectal |
| MET | 7q31.2 | 34 | 4.09 | 2 | 3.03 | 0.498 | 0.942 | (A) BRAF Class 1 Colorectal |
| BMPR1A | 10q23.2 | 36 | 6.25 | 2 | 4.65 | 0.498 | 0.942 | (A) BRAF Class 1 Colorectal |
| RUNX1 | 21q22.12 | 36 | 5.84 | 2 | 4.35 | 0.498 | 0.942 | (A) BRAF Class 1 Colorectal |
| FAS | 10q23.31 | 10 | 2.63 | 0 | 0.00 | 0.499 | 0.942 | (A) BRAF Class 1 Colorectal |
| ETV5 | 3q27.2 | 10 | 2.62 | 0 | 0.00 | 0.5 | 0.942 | (A) BRAF Class 1 Colorectal |
| ORCS8-MEF2 | 19p13.11 | 8 | 0.95 | 1 | 1.49 | 0.5 | 0.942 | (B) BRAF Class 2 Colorectal |
| NTHL1 | 16p13.3 | 8 | 2.75 | 1 | 4.35 | 0.5 | 0.942 | (B) BRAF Class 2 Colorectal |
| LRP1B | :q22.1-q22.: | 33 | 23.08 | 1 | 14.29 | 0.502 | 0.942 | (A) BRAF Class 1 Colorectal |
| NTRK1 | 1q23.1 | 36 | 5.67 | 2 | 4.26 | 0.504 | 0.942 | (A) BRAF Class 1 Colorectal |
| DDB2 | 11p11.2 | 9 | 3.32 | 0 | 0.00 | 0.506 | 0.942 | (A) BRAF Class 1 Colorectal |
| CDKN1B | 12p13.1 | 22 | 3.63 | 1 | 2.17 | 0.507 | 0.942 | (A) BRAF Class 1 Colorectal |
| GEN1 | 2p24.2 | 11 | 3.82 | 0 | 0.00 | 0.507 | 0.942 | (A) BRAF Class 1 Colorectal |
| MLH1 | 3p22.2 | 24 | 2.98 | 1 | 1.79 | 0.508 | 0.942 | (A) BRAF Class 1 Colorectal |
| PRKN | 6q26 | 34 | 5.76 | 2 | 4.35 | 0.508 | 0.942 | (A) BRAF Class 1 Colorectal |
| LZTR1 | 21\|22q11.1 | 13 | 9.42 | 0 | 0.00 | 0.511 | 0.942 | (A) BRAF Class 1 Colorectal |
| ARID5B | 10q21.2 | 23 | 7.82 | 1 | 4.76 | 0.511 | 0.942 | (A) BRAF Class 1 Colorectal |
| MCM8 | 20p12.3 | 8 | 4.97 | 1 | 7.69 | 0.511 | 0.942 | (B) BRAF Class 2 Colorectal |
| CUL3 | 2q36.2 | 9 | 2.54 | 0 | 0.00 | 0.512 | 0.942 | (A) BRAF Class 1 Colorectal |
| PNRC1 | 6q15 | 10 | 2.61 | 0 | 0.00 | 0.515 | 0.942 | (A) BRAF Class 1 Colorectal |
| NFKBIA | 14q13.2 | 9 | 1.55 | 0 | 0.00 | 0.515 | 0.942 | (A) BRAF Class 1 Colorectal |
| MDM4 | 1q32.1 | 9 | 1.49 | 0 | 0.00 | 0.515 | 0.942 | (A) BRAF Class 1 Colorectal |
| VEGFA | 6p21.1 | 9 | 1.79 | 0 | 0.00 | 0.516 | 0.942 | (A) BRAF Class 1 Colorectal |
| RAD51 | 15q15.1 | 9 | 1.70 | 0 | 0.00 | 0.516 | 0.942 | (A) BRAF Class 1 Colorectal |
| TNFRSF14 | 1p36.32 | 9 | 2.50 | 0 | 0.00 | 0.518 | 0.942 | (A) BRAF Class 1 Colorectal |
| ERRFI1 | 1p36.23 | 9 | 2.69 | 0 | 0.00 | 0.519 | 0.942 | (A) BRAF Class 1 Colorectal |
| RIT1 | 1q22 | 9 | 1.82 | 0 | 0.00 | 0.519 | 0.942 | (A) BRAF Class 1 Colorectal |
| SDHAF2 | 11q12.2 | 9 | 1.60 | 0 | 0.00 | 0.52 | 0.942 | (A) BRAF Class 1 Colorectal |
| MITF | 3p13 | 21 | 3.52 | 1 | 2.17 | 0.523 | 0.942 | (A) BRAF Class 1 Colorectal |
| CD276 | 15q24.1 | 10 | 3.46 | 0 | 0.00 | 0.524 | 0.942 | (A) BRAF Class 1 Colorectal |
| ATR | 3q23 | 73 | 13.77 | 5 | 12.50 | 0.525 | 0.942 | (A) BRAF Class 1 Colorectal |
| NPM1 | 5q35.1 | 10 | 1.29 | 1 | 1.85 | 0.526 | 0.942 | (B) BRAF Class 2 Colorectal |
| CBFB | 16q22.1 | 9 | 1.73 | 1 | 2.50 | 0.526 | 0.942 | (B) BRAF Class 2 Colorectal |
| DEPDC5 | 2q12.2-q12. | 6 | 10.71 | 1 | 16.67 | 0.528 | 0.942 | (B) BRAF Class 2 Colorectal |
| TRAF3 | 14q32.32 | 8 | 4.02 | 1 | 5.88 | 0.529 | 0.942 | (B) BRAF Class 2 Colorectal |
| BABAM2 | 2p23.2 | 8 | 4.97 | 0 | 0.00 | 0.53 | 0.942 | (A) BRAF Class 1 Colorectal |
| HELQ | 4q21.23 | 8 | 4.97 | 0 | 0.00 | 0.53 | 0.942 | (A) BRAF Class 1 Colorectal |
| NR0B1 | Xp21.2 | 8 | 4.97 | 0 | 0.00 | 0.53 | 0.942 | (A) BRAF Class 1 Colorectal |
| CEBPA | 19q13.11 | 21 | 3.63 | 1 | 2.27 | 0.53 | 0.942 | (A) BRAF Class 1 Colorectal |

| | | | | | | | | |
|---|---|---|---|---|---|---|---|---|
| XPC | 3p25.1 | 9 | 3.32 | 1 | 4.76 | 0.532 | 0.942 | (B) BRAF Class 2 Colorectal |
| PRKD1 | 14q12 | 10 | 6.29 | 0 | 0.00 | 0.534 | 0.942 | (A) BRAF Class 1 Colorectal |
| FANCB | Xp22.2 | 8 | 4.00 | 0 | 0.00 | 0.535 | 0.942 | (A) BRAF Class 1 Colorectal |
| RIF1 | 2q23.3 | 21 | 13.04 | 2 | 15.38 | 0.536 | 0.942 | (B) BRAF Class 2 Colorectal |
| LMO1 | 11p15.4 | 9 | 1.55 | 0 | 0.00 | 0.539 | 0.942 | (A) BRAF Class 1 Colorectal |
| SUZ12 | 17q11.2 | 21 | 3.73 | 1 | 2.38 | 0.54 | 0.942 | (A) BRAF Class 1 Colorectal |
| ZRSR2 | Xp22.2 | 24 | 4.39 | 2 | 5.00 | 0.542 | 0.942 | (B) BRAF Class 2 Colorectal |
| SRSF2 | 17q25.1 | 10 | 1.83 | 1 | 2.50 | 0.543 | 0.942 | (B) BRAF Class 2 Colorectal |
| TERT | 5p15.33 | 48 | 7.88 | 3 | 6.82 | 0.544 | 0.942 | (A) BRAF Class 1 Colorectal |
| EXT2 | 11p11.2 | 8 | 2.95 | 0 | 0.00 | 0.546 | 0.942 | (A) BRAF Class 1 Colorectal |
| STAG2 | Xq25 | 23 | 3.85 | 2 | 4.35 | 0.547 | 0.942 | (B) BRAF Class 2 Colorectal |
| MCL1 | 1q21.2 | 8 | 1.34 | 0 | 0.00 | 0.55 | 0.942 | (A) BRAF Class 1 Colorectal |
| SH2B3 | 12q24.12 | 34 | 6.02 | 2 | 4.88 | 0.553 | 0.942 | (A) BRAF Class 1 Colorectal |
| PRPF40B | 12q13.12 | 7 | 8.86 | 1 | 12.50 | 0.554 | 0.942 | (B) BRAF Class 2 Colorectal |
| IKBKE | 1q32.1 | 10 | 2.78 | 1 | 3.70 | 0.554 | 0.942 | (B) BRAF Class 2 Colorectal |
| TAF1 | Xq13.1 | 15 | 11.36 | 0 | 0.00 | 0.555 | 0.942 | (A) BRAF Class 1 Colorectal |
| MYOD1 | 11p15.1 | 21 | 7.19 | 1 | 4.76 | 0.555 | 0.942 | (A) BRAF Class 1 Colorectal |
| NFE2L2 | 2q31.2 | 10 | 1.65 | 1 | 2.17 | 0.556 | 0.942 | (B) BRAF Class 2 Colorectal |
| CSF1R | 5q32 | 22 | 3.52 | 1 | 2.33 | 0.556 | 0.942 | (A) BRAF Class 1 Colorectal |
| NUP93 | 16q13 | 9 | 2.71 | 0 | 0.00 | 0.557 | 0.942 | (A) BRAF Class 1 Colorectal |
| MAP3K4 | 6q26 | 21 | 25.61 | 0 | 0.00 | 0.56 | 0.942 | (A) BRAF Class 1 Colorectal |
| SOX9 | 17q24.3 | 93 | 15.76 | 7 | 15.22 | 0.561 | 0.942 | (A) BRAF Class 1 Colorectal |
| ERBB2 | 17q12 | 49 | 5.83 | 4 | 5.97 | 0.562 | 0.942 | (B) BRAF Class 2 Colorectal |
| IRS2 | 13q34 | 38 | 10.56 | 3 | 11.11 | 0.563 | 0.942 | (B) BRAF Class 2 Colorectal |
| CDKN2A | 9p21.3 | 27 | 3.23 | 2 | 3.51 | 0.565 | 0.942 | (B) BRAF Class 2 Colorectal |
| MAX | 14q23.3 | 8 | 1.73 | 0 | 0.00 | 0.565 | 0.942 | (A) BRAF Class 1 Colorectal |
| UPF1 | 19p13.11 | 12 | 9.45 | 1 | 12.50 | 0.565 | 0.942 | (B) BRAF Class 2 Colorectal |
| CSF3R | 1p34.3 | 30 | 5.50 | 2 | 4.55 | 0.566 | 0.942 | (A) BRAF Class 1 Colorectal |
| TAP1 | 6p21.32 | 9 | 7.09 | 0 | 0.00 | 0.567 | 0.942 | (A) BRAF Class 1 Colorectal |
| MRE11 | 11q21 | 32 | 6.04 | 2 | 5.00 | 0.567 | 0.942 | (A) BRAF Class 1 Colorectal |
| ALOX12B | 17p13.1 | 20 | 5.09 | 1 | 3.45 | 0.568 | 0.942 | (A) BRAF Class 1 Colorectal |
| PDCD1LG2 | 9p24.1 | 8 | 1.57 | 0 | 0.00 | 0.57 | 0.942 | (A) BRAF Class 1 Colorectal |
| SRC | 20q11.23 | 12 | 1.86 | 1 | 2.33 | 0.571 | 0.942 | (B) BRAF Class 2 Colorectal |
| PPM1D | 17q23.2 | 20 | 4.58 | 1 | 3.13 | 0.572 | 0.942 | (A) BRAF Class 1 Colorectal |
| FANCC | 9q22.32 | 19 | 3.18 | 1 | 2.17 | 0.574 | 0.942 | (A) BRAF Class 1 Colorectal |
| NKX3-1 | 8p21.2 | 8 | 1.78 | 0 | 0.00 | 0.575 | 0.942 | (A) BRAF Class 1 Colorectal |
| DMC1 | 22q13.1 | 7 | 4.35 | 0 | 0.00 | 0.575 | 0.942 | (A) BRAF Class 1 Colorectal |
| MBD4 | 3q21.3 | 7 | 4.35 | 0 | 0.00 | 0.575 | 0.942 | (A) BRAF Class 1 Colorectal |
| USP28 | 11q23.2 | 7 | 4.35 | 0 | 0.00 | 0.575 | 0.942 | (A) BRAF Class 1 Colorectal |
| ZNF703 | 8p11.23 | 12 | 7.84 | 1 | 10.00 | 0.575 | 0.942 | (B) BRAF Class 2 Colorectal |
| EPHA5 | 4q13.1-q13.2 | 56 | 13.15 | 4 | 13.33 | 0.575 | 0.942 | (B) BRAF Class 2 Colorectal |
| DDR2 | 1q23.3 | 21 | 3.28 | 2 | 3.51 | 0.576 | 0.942 | (B) BRAF Class 2 Colorectal |
| MDC1 | 6p21.33 | 22 | 7.61 | 1 | 5.26 | 0.576 | 0.942 | (A) BRAF Class 1 Colorectal |
| RSPO2 | 8q23.1 | 6 | 2.71 | 0 | 0.00 | 0.577 | 0.942 | (A) BRAF Class 1 Colorectal |
| USP8 | 15q21.2 | 7 | 4.32 | 0 | 0.00 | 0.577 | 0.942 | (A) BRAF Class 1 Colorectal |
| MAP3K1 | 5q11.2 | 43 | 7.20 | 3 | 6.52 | 0.578 | 0.942 | (A) BRAF Class 1 Colorectal |
| PDGFRA | 4q12 | 40 | 5.02 | 3 | 4.55 | 0.579 | 0.942 | (A) BRAF Class 1 Colorectal |
| COP1 | .q25.1-q25.[illegible] | 21 | 5.75 | 1 | 4.00 | 0.58 | 0.942 | (A) BRAF Class 1 Colorectal |

| | | | | | | | | |
|---|---|---|---|---|---|---|---|---|
| TRAF2 | 9q34.3 | 13 | 4.29 | 1 | 5.26 | 0.581 | 0.942 | (B) BRAF Class 2 Colorectal |
| PDCD1 | 2q37.3 | 26 | 7.83 | 2 | 8.33 | 0.582 | 0.942 | (B) BRAF Class 2 Colorectal |
| STAT5B | 17q21.2 | 30 | 9.87 | 2 | 10.53 | 0.582 | 0.942 | (B) BRAF Class 2 Colorectal |
| INHA | 2q35 | 13 | 4.80 | 1 | 5.88 | 0.582 | 0.942 | (B) BRAF Class 2 Colorectal |
| TOPBP1 | 3q22.1 | 10 | 6.21 | 1 | 7.69 | 0.586 | 0.942 | (B) BRAF Class 2 Colorectal |
| EIF1AX | Xp22.12 | 8 | 2.48 | 0 | 0.00 | 0.587 | 0.942 | (A) BRAF Class 1 Colorectal |
| XRCC1 | 19q13.31 | 7 | 3.63 | 0 | 0.00 | 0.587 | 0.942 | (A) BRAF Class 1 Colorectal |
| TRAF7 | 16p13.3 | 26 | 5.33 | 2 | 5.56 | 0.589 | 0.942 | (B) BRAF Class 2 Colorectal |
| SDHC | 1q23.3 | 7 | 1.19 | 0 | 0.00 | 0.59 | 0.942 | (A) BRAF Class 1 Colorectal |
| XPA | 9q22.33 | 7 | 2.58 | 0 | 0.00 | 0.59 | 0.942 | (A) BRAF Class 1 Colorectal |
| CBLB | 3q13.11 | 7 | 2.57 | 0 | 0.00 | 0.591 | 0.942 | (A) BRAF Class 1 Colorectal |
| SMC3 | 10q25.2 | 12 | 3.36 | 1 | 4.00 | 0.591 | 0.942 | (B) BRAF Class 2 Colorectal |
| BCL2 | 18q21.33 | 7 | 1.18 | 0 | 0.00 | 0.591 | 0.942 | (A) BRAF Class 1 Colorectal |
| DNMT3A | 2p23.3 | 54 | 8.93 | 4 | 8.51 | 0.591 | 0.942 | (A) BRAF Class 1 Colorectal |
| CXCR4 | 2q22.1 | 7 | 1.46 | 0 | 0.00 | 0.592 | 0.942 | (A) BRAF Class 1 Colorectal |
| ETV4 | 17q21.31 | 12 | 3.15 | 1 | 3.70 | 0.595 | 0.942 | (B) BRAF Class 2 Colorectal |
| PIK3R3 | 1p34.1 | 8 | 2.77 | 0 | 0.00 | 0.597 | 0.942 | (A) BRAF Class 1 Colorectal |
| STK40 | 1p34.3 | 8 | 2.77 | 0 | 0.00 | 0.597 | 0.942 | (A) BRAF Class 1 Colorectal |
| XIAP | Xq25 | 8 | 2.77 | 0 | 0.00 | 0.597 | 0.942 | (A) BRAF Class 1 Colorectal |
| PRPF8 | 17p13.3 | 8 | 10.13 | 1 | 12.50 | 0.599 | 0.942 | (B) BRAF Class 2 Colorectal |
| ABL2 | 1q25.2 | 10 | 7.25 | 0 | 0.00 | 0.6 | 0.942 | (A) BRAF Class 1 Colorectal |
| STAT3 | 17q21.2 | 26 | 4.46 | 2 | 4.55 | 0.6 | 0.942 | (B) BRAF Class 2 Colorectal |
| ABCB4 | 7q21.12 | 1 | 33.33 | 0 | 0.00 | 0.6 | 0.942 | (A) BRAF Class 1 Colorectal |
| SAMD9L | 7q21.2 | 1 | 33.33 | 0 | 0.00 | 0.6 | 0.942 | (A) BRAF Class 1 Colorectal |
| SMARCAL1 | 2q35 | 1 | 33.33 | 0 | 0.00 | 0.6 | 0.942 | (A) BRAF Class 1 Colorectal |
| MAGI2 | 7q21.11 | 13 | 9.92 | 0 | 0.00 | 0.6 | 0.942 | (A) BRAF Class 1 Colorectal |
| TMEM127 | 2q11.2 | 7 | 1.55 | 0 | 0.00 | 0.601 | 0.942 | (A) BRAF Class 1 Colorectal |
| RHOA | 3p21.31 | 19 | 3.83 | 1 | 2.78 | 0.602 | 0.942 | (A) BRAF Class 1 Colorectal |
| FLT3 | 13q12.2 | 46 | 5.71 | 3 | 5.36 | 0.604 | 0.942 | (A) BRAF Class 1 Colorectal |
| PRDM14 | 8q13.3 | 8 | 6.30 | 0 | 0.00 | 0.605 | 0.942 | (A) BRAF Class 1 Colorectal |
| XRCC2 | 7q36.1 | 7 | 1.43 | 0 | 0.00 | 0.606 | 0.942 | (A) BRAF Class 1 Colorectal |
| CCNE1 | 19q12 | 18 | 2.97 | 1 | 2.17 | 0.607 | 0.942 | (A) BRAF Class 1 Colorectal |
| MDM2 | 12q15 | 18 | 2.97 | 1 | 2.17 | 0.607 | 0.942 | (A) BRAF Class 1 Colorectal |
| STAT5A | 17q21.2 | 14 | 4.62 | 1 | 5.26 | 0.607 | 0.942 | (B) BRAF Class 2 Colorectal |
| FGF19 | 11q13.3 | 7 | 1.90 | 0 | 0.00 | 0.608 | 0.942 | (A) BRAF Class 1 Colorectal |
| SF1 | 11q13.1 | 5 | 6.33 | 0 | 0.00 | 0.61 | 0.942 | (A) BRAF Class 1 Colorectal |
| EPHB1 | 3q22.2 | 30 | 8.33 | 2 | 7.41 | 0.61 | 0.942 | (A) BRAF Class 1 Colorectal |
| STAT4 | q32.2-q32. | 10 | 6.99 | 0 | 0.00 | 0.611 | 0.942 | (A) BRAF Class 1 Colorectal |
| POT1 | 7q31.33 | 6 | 2.99 | 0 | 0.00 | 0.611 | 0.942 | (A) BRAF Class 1 Colorectal |
| ETV6 | 12p13.2 | 29 | 4.89 | 2 | 4.35 | 0.611 | 0.942 | (A) BRAF Class 1 Colorectal |
| EIF4A2 | 3q27.3 | 8 | 2.94 | 0 | 0.00 | 0.612 | 0.942 | (A) BRAF Class 1 Colorectal |
| GPS2 | 17p13.1 | 8 | 2.94 | 0 | 0.00 | 0.612 | 0.942 | (A) BRAF Class 1 Colorectal |
| RASA1 | 5q14.3 | 18 | 3.96 | 1 | 2.94 | 0.614 | 0.942 | (A) BRAF Class 1 Colorectal |
| STAT6 | 12q13.3 | 6 | 2.19 | 0 | 0.00 | 0.614 | 0.942 | (A) BRAF Class 1 Colorectal |
| UIMC1 | 5q35.2 | 6 | 3.57 | 0 | 0.00 | 0.614 | 0.942 | (A) BRAF Class 1 Colorectal |
| XRCC5 | 2q35 | 6 | 3.57 | 0 | 0.00 | 0.614 | 0.942 | (A) BRAF Class 1 Colorectal |
| GPC3 | Xq26.2 | 6 | 2.18 | 0 | 0.00 | 0.615 | 0.942 | (A) BRAF Class 1 Colorectal |
| WRN | 8p12 | 24 | 8.70 | 2 | 8.70 | 0.617 | 0.942 | (B) BRAF Class 2 Colorectal |

| | | | | | | | | |
|---|---|---|---|---|---|---|---|---|
| DCUN1D1 | 3q26.33 | 5 | 1.73 | 0 | 0.00 | 0.618 | 0.942 | (A) BRAF Class 1 Colorectal |
| AURKA | 20q13.2 | 12 | 2.01 | 1 | 2.17 | 0.623 | 0.942 | (B) BRAF Class 2 Colorectal |
| CD79A | 19q13.2 | 18 | 5.26 | 1 | 4.00 | 0.623 | 0.942 | (A) BRAF Class 1 Colorectal |
| MUS81 | 11q13.1 | 6 | 3.73 | 0 | 0.00 | 0.623 | 0.942 | (A) BRAF Class 1 Colorectal |
| NEIL1 | 15q24.2 | 6 | 3.73 | 0 | 0.00 | 0.623 | 0.942 | (A) BRAF Class 1 Colorectal |
| RINT1 | 7q22.3 | 6 | 3.73 | 0 | 0.00 | 0.623 | 0.942 | (A) BRAF Class 1 Colorectal |
| TRIM37 | 17q22 | 6 | 3.73 | 0 | 0.00 | 0.623 | 0.942 | (A) BRAF Class 1 Colorectal |
| DMD | (p21.2-p21.: | 6 | 7.89 | 0 | 0.00 | 0.624 | 0.942 | (A) BRAF Class 1 Colorectal |
| PTK2 | 8q24.3 | 6 | 7.89 | 0 | 0.00 | 0.624 | 0.942 | (A) BRAF Class 1 Colorectal |
| TENT5C | 1p12 | 17 | 2.85 | 1 | 2.17 | 0.626 | 0.942 | (A) BRAF Class 1 Colorectal |
| MYD88 | 3p22.2 | 17 | 2.77 | 1 | 2.13 | 0.629 | 0.942 | (A) BRAF Class 1 Colorectal |
| NT5C2 | q24.32-q24. | 5 | 2.39 | 0 | 0.00 | 0.631 | 0.942 | (A) BRAF Class 1 Colorectal |
| RPA1 | 17p13.3 | 6 | 3.35 | 0 | 0.00 | 0.633 | 0.942 | (A) BRAF Class 1 Colorectal |
| BRIP1 | 17q23.2 | 28 | 4.69 | 2 | 4.35 | 0.635 | 0.942 | (A) BRAF Class 1 Colorectal |
| EMSY | 11q13.5 | 7 | 4.58 | 0 | 0.00 | 0.636 | 0.942 | (A) BRAF Class 1 Colorectal |
| SDHB | 1p36.13 | 6 | 1.01 | 0 | 0.00 | 0.637 | 0.942 | (A) BRAF Class 1 Colorectal |
| PAX7 | 1p36.13 | 9 | 7.44 | 0 | 0.00 | 0.637 | 0.942 | (A) BRAF Class 1 Colorectal |
| EGFL7 | 9q34.3 | 7 | 2.42 | 0 | 0.00 | 0.637 | 0.942 | (A) BRAF Class 1 Colorectal |
| CD79B | 17q23.3 | 6 | 1.01 | 0 | 0.00 | 0.638 | 0.942 | (A) BRAF Class 1 Colorectal |
| HGF | 7q21.11 | 17 | 4.72 | 1 | 3.70 | 0.638 | 0.942 | (A) BRAF Class 1 Colorectal |
| INPPL1 | 11q13.4 | 20 | 15.75 | 1 | 12.50 | 0.639 | 0.942 | (A) BRAF Class 1 Colorectal |
| MAPK1 | 22q11.22 | 6 | 1.03 | 0 | 0.00 | 0.639 | 0.942 | (A) BRAF Class 1 Colorectal |
| CHD2 | 15q26.1 | 18 | 13.24 | 1 | 14.29 | 0.64 | 0.942 | (B) BRAF Class 2 Colorectal |
| JUN | 1p32.1 | 18 | 5.07 | 1 | 4.00 | 0.64 | 0.942 | (A) BRAF Class 1 Colorectal |
| SPRED1 | 15q14 | 6 | 4.55 | 0 | 0.00 | 0.64 | 0.942 | (A) BRAF Class 1 Colorectal |
| BCR | 22q11.23 | 8 | 5.56 | 0 | 0.00 | 0.642 | 0.942 | (A) BRAF Class 1 Colorectal |
| PAK5 | 20p12.2 | 15 | 4.89 | 1 | 5.00 | 0.645 | 0.942 | (B) BRAF Class 2 Colorectal |
| ERF | 19q13.2 | 7 | 5.51 | 0 | 0.00 | 0.645 | 0.942 | (A) BRAF Class 1 Colorectal |
| JAK2 | 9p24.1 | 26 | 3.31 | 2 | 3.13 | 0.646 | 0.942 | (A) BRAF Class 1 Colorectal |
| NFKBIE | 6p21.1 | 5 | 3.01 | 0 | 0.00 | 0.646 | 0.942 | (A) BRAF Class 1 Colorectal |
| SMARCB1 | q11.23\|22q | 18 | 2.23 | 1 | 1.79 | 0.647 | 0.942 | (A) BRAF Class 1 Colorectal |
| EPCAM | 2p21 | 6 | 1.21 | 0 | 0.00 | 0.647 | 0.942 | (A) BRAF Class 1 Colorectal |
| FUS | 16p11.2 | 5 | 1.95 | 0 | 0.00 | 0.648 | 0.942 | (A) BRAF Class 1 Colorectal |
| MAP3K9 | 14q24.2 | 2 | 15.38 | 0 | 0.00 | 0.65 | 0.943 | (A) BRAF Class 1 Colorectal |
| POLQ | 3q13.33 | 26 | 15.95 | 2 | 15.38 | 0.658 | 0.95 | (A) BRAF Class 1 Colorectal |
| PAK1 | 1q13.5-q14. | 6 | 2.04 | 0 | 0.00 | 0.659 | 0.95 | (A) BRAF Class 1 Colorectal |
| GATA6 | 18q11.2 | 17 | 4.50 | 1 | 3.70 | 0.659 | 0.95 | (A) BRAF Class 1 Colorectal |
| RELA | 11q13.1 | 5 | 2.53 | 0 | 0.00 | 0.66 | 0.95 | (A) BRAF Class 1 Colorectal |
| FANCG | 9p13.3 | 16 | 4.10 | 1 | 3.45 | 0.668 | 0.95 | (A) BRAF Class 1 Colorectal |
| PPP6C | 9q33.3 | 6 | 2.17 | 0 | 0.00 | 0.669 | 0.95 | (A) BRAF Class 1 Colorectal |
| PAX3 | 2q36.1 | 8 | 6.61 | 0 | 0.00 | 0.671 | 0.95 | (A) BRAF Class 1 Colorectal |
| HSD3B1 | 1p12 | 7 | 4.96 | 0 | 0.00 | 0.674 | 0.95 | (A) BRAF Class 1 Colorectal |
| FYN | 6q21 | 17 | 6.16 | 1 | 5.26 | 0.674 | 0.95 | (A) BRAF Class 1 Colorectal |
| BRCC3 | Xq28 | 5 | 3.11 | 0 | 0.00 | 0.675 | 0.95 | (A) BRAF Class 1 Colorectal |
| HABP2 | 10q25.3 | 5 | 3.11 | 0 | 0.00 | 0.675 | 0.95 | (A) BRAF Class 1 Colorectal |
| PNKP | 19q13.33 | 5 | 3.11 | 0 | 0.00 | 0.675 | 0.95 | (A) BRAF Class 1 Colorectal |
| UROD | 1p34.1 | 5 | 3.11 | 0 | 0.00 | 0.675 | 0.95 | (A) BRAF Class 1 Colorectal |
| CCN6 | 6q21 | 8 | 5.59 | 0 | 0.00 | 0.676 | 0.95 | (A) BRAF Class 1 Colorectal |

| | | | | | | | | |
|---|---|---|---|---|---|---|---|---|
| ITK | 5q33.3 | 5 | 3.09 | 0 | 0.00 | 0.677 | 0.95 | (A) BRAF Class 1 Colorectal |
| KLHL6 | 3q27.1 | 6 | 3.92 | 0 | 0.00 | 0.68 | 0.95 | (A) BRAF Class 1 Colorectal |
| BCL2L12 | 19q13.33 | 5 | 2.09 | 0 | 0.00 | 0.68 | 0.95 | (A) BRAF Class 1 Colorectal |
| VTCN1 | 1p13.1-p12 | 6 | 2.08 | 0 | 0.00 | 0.68 | 0.95 | (A) BRAF Class 1 Colorectal |
| PHF1 | 6p21.32 | 5 | 0.60 | 0 | 0.00 | 0.681 | 0.95 | (A) BRAF Class 1 Colorectal |
| FAT3 | 11q14.3 | 3 | 13.04 | 0 | 0.00 | 0.681 | 0.95 | (A) BRAF Class 1 Colorectal |
| BCL2L1 | 20q11.21 | 5 | 0.85 | 0 | 0.00 | 0.685 | 0.95 | (A) BRAF Class 1 Colorectal |
| PAXIP1 | 7q36.2 | 14 | 8.70 | 1 | 7.69 | 0.689 | 0.95 | (A) BRAF Class 1 Colorectal |
| EZH1 | 17q21.2 | 5 | 3.79 | 0 | 0.00 | 0.69 | 0.95 | (A) BRAF Class 1 Colorectal |
| MYB | 6q23.3 | 5 | 1.31 | 0 | 0.00 | 0.692 | 0.95 | (A) BRAF Class 1 Colorectal |
| ARHGEF12 | 11q23.3 | 14 | 8.64 | 1 | 7.69 | 0.692 | 0.95 | (A) BRAF Class 1 Colorectal |
| HIST1H3J | 6p22.1 | 6 | 2.21 | 0 | 0.00 | 0.692 | 0.95 | (A) BRAF Class 1 Colorectal |
| MST1 | 3p21.31 | 6 | 2.21 | 0 | 0.00 | 0.692 | 0.95 | (A) BRAF Class 1 Colorectal |
| RAB35 | 12q24.23 | 6 | 2.21 | 0 | 0.00 | 0.692 | 0.95 | (A) BRAF Class 1 Colorectal |
| BIRC3 | 11q22.2 | 6 | 1.97 | 0 | 0.00 | 0.694 | 0.95 | (A) BRAF Class 1 Colorectal |
| SPOP | 17q21.33 | 5 | 0.94 | 0 | 0.00 | 0.694 | 0.95 | (A) BRAF Class 1 Colorectal |
| PIM1 | 6p21.2 | 5 | 0.88 | 0 | 0.00 | 0.694 | 0.95 | (A) BRAF Class 1 Colorectal |
| H3F3A | 1q42.12 | 5 | 0.82 | 0 | 0.00 | 0.7 | 0.95 | (A) BRAF Class 1 Colorectal |
| PAK3 | Xq23 | 7 | 4.90 | 0 | 0.00 | 0.711 | 0.95 | (A) BRAF Class 1 Colorectal |
| AGO2 | 8q24.3 | 17 | 13.39 | 1 | 12.50 | 0.711 | 0.95 | (A) BRAF Class 1 Colorectal |
| HIST1H3B | 6p22.2 | 5 | 0.95 | 0 | 0.00 | 0.711 | 0.95 | (A) BRAF Class 1 Colorectal |
| NUTM1 | 15q14 | 5 | 3.85 | 0 | 0.00 | 0.712 | 0.95 | (A) BRAF Class 1 Colorectal |
| PHF6 | Xq26.2 | 15 | 4.09 | 1 | 3.85 | 0.714 | 0.95 | (A) BRAF Class 1 Colorectal |
| AJUBA | 14q11.2 | 1 | 20.00 | 0 | 0.00 | 0.714 | 0.95 | (A) BRAF Class 1 Colorectal |
| CHD5 | 1p36.31 | 1 | 20.00 | 0 | 0.00 | 0.714 | 0.95 | (A) BRAF Class 1 Colorectal |
| ZBTB20 | 3q13.31 | 1 | 20.00 | 0 | 0.00 | 0.714 | 0.95 | (A) BRAF Class 1 Colorectal |
| BBC3 | 19q13.32 | 5 | 1.73 | 0 | 0.00 | 0.726 | 0.95 | (A) BRAF Class 1 Colorectal |
| E2F3 | 6p22.3 | 5 | 1.73 | 0 | 0.00 | 0.726 | 0.95 | (A) BRAF Class 1 Colorectal |
| HIST1H2BD | 6p22.2 | 5 | 1.73 | 0 | 0.00 | 0.726 | 0.95 | (A) BRAF Class 1 Colorectal |
| BCL2L11 | 2q13 | 5 | 1.72 | 0 | 0.00 | 0.727 | 0.95 | (A) BRAF Class 1 Colorectal |
| EED | 11q14.2 | 4 | 1.03 | 0 | 0.00 | 0.729 | 0.95 | (A) BRAF Class 1 Colorectal |
| IGF2 | 11p15.5 | 14 | 2.75 | 1 | 2.70 | 0.731 | 0.95 | (A) BRAF Class 1 Colorectal |
| GBA | 1q22 | 4 | 2.48 | 0 | 0.00 | 0.731 | 0.95 | (A) BRAF Class 1 Colorectal |
| KLF2 | 19p13.11 | 4 | 2.48 | 0 | 0.00 | 0.731 | 0.95 | (A) BRAF Class 1 Colorectal |
| RAD54B | 8q22.1 | 4 | 2.48 | 0 | 0.00 | 0.731 | 0.95 | (A) BRAF Class 1 Colorectal |
| RNF8 | 6p21.2 | 4 | 2.48 | 0 | 0.00 | 0.731 | 0.95 | (A) BRAF Class 1 Colorectal |
| XRCC6 | 22q13.2 | 4 | 2.48 | 0 | 0.00 | 0.731 | 0.95 | (A) BRAF Class 1 Colorectal |
| KCNIP1 | 5q35.1 | 3 | 5.36 | 0 | 0.00 | 0.733 | 0.95 | (A) BRAF Class 1 Colorectal |
| CARM1 | 19p13.2 | 5 | 3.94 | 0 | 0.00 | 0.733 | 0.95 | (A) BRAF Class 1 Colorectal |
| MSI1 | 12q24.31 | 5 | 3.94 | 0 | 0.00 | 0.733 | 0.95 | (A) BRAF Class 1 Colorectal |
| PRKCZ | 1p36.33 | 4 | 5.26 | 0 | 0.00 | 0.734 | 0.95 | (A) BRAF Class 1 Colorectal |
| BRAF | 7q34 | 836 | 99.52 | 67 | 100.00 | 0.735 | 0.95 | (B) BRAF Class 2 Colorectal |
| SBDS | 7q11.21 | 4 | 1.48 | 0 | 0.00 | 0.741 | 0.95 | (A) BRAF Class 1 Colorectal |
| SMC1A | Xp11.22 | 5 | 2.55 | 0 | 0.00 | 0.741 | 0.95 | (A) BRAF Class 1 Colorectal |
| SDHD | 11q23.1 | 4 | 0.68 | 0 | 0.00 | 0.741 | 0.95 | (A) BRAF Class 1 Colorectal |
| CDKN2C | 1p32.3 | 4 | 0.67 | 0 | 0.00 | 0.741 | 0.95 | (A) BRAF Class 1 Colorectal |
| WAS | Xp11.23 | 4 | 2.05 | 0 | 0.00 | 0.742 | 0.95 | (A) BRAF Class 1 Colorectal |
| ERCC1 | 19q13.32 | 5 | 1.79 | 0 | 0.00 | 0.742 | 0.95 | (A) BRAF Class 1 Colorectal |

| | | | | | | | | |
|---|---|---|---|---|---|---|---|---|
| SOX10 | 22q13.1 | 6 | 4.20 | 0 | 0.00 | 0.747 | 0.95 | (A) BRAF Class 1 Colorectal |
| SMARCA2 | 9p24.3 | 1 | 16.67 | 0 | 0.00 | 0.75 | 0.95 | (A) BRAF Class 1 Colorectal |
| TCF7L1 | 2p11.2 | 4 | 3.67 | 0 | 0.00 | 0.75 | 0.95 | (A) BRAF Class 1 Colorectal |
| GABRA6 | 5q34 | 5 | 3.55 | 0 | 0.00 | 0.756 | 0.95 | (A) BRAF Class 1 Colorectal |
| ACVR1B | 12q13.13 | 4 | 2.74 | 0 | 0.00 | 0.765 | 0.95 | (A) BRAF Class 1 Colorectal |
| SNCAIP | 5q23.2 | 4 | 2.74 | 0 | 0.00 | 0.765 | 0.95 | (A) BRAF Class 1 Colorectal |
| EPHB4 | 7q22.1 | 2 | 3.39 | 0 | 0.00 | 0.774 | 0.95 | (A) BRAF Class 1 Colorectal |
| H3F3C | 12p11.21 | 4 | 1.38 | 0 | 0.00 | 0.774 | 0.95 | (A) BRAF Class 1 Colorectal |
| DHX15 | 4p15.2 | 10 | 12.20 | 0 | 0.00 | 0.775 | 0.95 | (A) BRAF Class 1 Colorectal |
| MAF | 16q23.2 | 3 | 1.46 | 0 | 0.00 | 0.776 | 0.95 | (A) BRAF Class 1 Colorectal |
| IGF1 | 12q23.2 | 4 | 1.30 | 0 | 0.00 | 0.776 | 0.95 | (A) BRAF Class 1 Colorectal |
| MAPKAP1 | 9q33.3 | 4 | 3.15 | 0 | 0.00 | 0.781 | 0.95 | (A) BRAF Class 1 Colorectal |
| NUF2 | 1q23.3 | 4 | 3.15 | 0 | 0.00 | 0.781 | 0.95 | (A) BRAF Class 1 Colorectal |
| SESN2 | 1p35.3 | 4 | 3.15 | 0 | 0.00 | 0.781 | 0.95 | (A) BRAF Class 1 Colorectal |
| SESN3 | 11q21 | 4 | 3.15 | 0 | 0.00 | 0.781 | 0.95 | (A) BRAF Class 1 Colorectal |
| SHOC2 | 10q25.2 | 4 | 3.15 | 0 | 0.00 | 0.781 | 0.95 | (A) BRAF Class 1 Colorectal |
| STK19 | 6p21.33 | 4 | 3.15 | 0 | 0.00 | 0.781 | 0.95 | (A) BRAF Class 1 Colorectal |
| SS18 | 18q11.2 | 3 | 1.51 | 0 | 0.00 | 0.781 | 0.95 | (A) BRAF Class 1 Colorectal |
| HIST1H3E | 6p22.2 | 4 | 1.48 | 0 | 0.00 | 0.783 | 0.95 | (A) BRAF Class 1 Colorectal |
| DKC1 | Xq28 | 3 | 1.86 | 0 | 0.00 | 0.791 | 0.95 | (A) BRAF Class 1 Colorectal |
| EGLN1 | 1q42.2 | 3 | 1.86 | 0 | 0.00 | 0.791 | 0.95 | (A) BRAF Class 1 Colorectal |
| FAH | 15q25.1 | 3 | 1.86 | 0 | 0.00 | 0.791 | 0.95 | (A) BRAF Class 1 Colorectal |
| ID4 | 6p22.3 | 3 | 1.86 | 0 | 0.00 | 0.791 | 0.95 | (A) BRAF Class 1 Colorectal |
| NECTIN4 | 1q23.3 | 3 | 1.86 | 0 | 0.00 | 0.791 | 0.95 | (A) BRAF Class 1 Colorectal |
| RHOT1 | 17q11.2 | 3 | 1.86 | 0 | 0.00 | 0.791 | 0.95 | (A) BRAF Class 1 Colorectal |
| SLC25A13 | 7q21.3 | 3 | 1.86 | 0 | 0.00 | 0.791 | 0.95 | (A) BRAF Class 1 Colorectal |
| TDG | 12q23.3 | 3 | 1.86 | 0 | 0.00 | 0.791 | 0.95 | (A) BRAF Class 1 Colorectal |
| RBL2 | 16q12.2 | 3 | 3.95 | 0 | 0.00 | 0.794 | 0.95 | (A) BRAF Class 1 Colorectal |
| NAB2 | 12q13.3 | 9 | 10.98 | 0 | 0.00 | 0.796 | 0.95 | (A) BRAF Class 1 Colorectal |
| ZBTB7A | 19p13.3 | 9 | 10.98 | 0 | 0.00 | 0.796 | 0.95 | (A) BRAF Class 1 Colorectal |
| POLH | 6p21.1 | 3 | 1.55 | 0 | 0.00 | 0.798 | 0.95 | (A) BRAF Class 1 Colorectal |
| MAFB | 20q12 | 3 | 1.54 | 0 | 0.00 | 0.8 | 0.95 | (A) BRAF Class 1 Colorectal |
| FIP1L1 | 4q12 | 1 | 8.33 | 0 | 0.00 | 0.8 | 0.95 | (A) BRAF Class 1 Colorectal |
| KEL | 7q34 | 4 | 2.84 | 0 | 0.00 | 0.8 | 0.95 | (A) BRAF Class 1 Colorectal |
| CRKL | 22q11.21 | 3 | 0.50 | 0 | 0.00 | 0.8 | 0.95 | (A) BRAF Class 1 Colorectal |
| RRAGC | 1p34.3 | 3 | 2.27 | 0 | 0.00 | 0.802 | 0.95 | (A) BRAF Class 1 Colorectal |
| RRAS | 19q13.33 | 3 | 2.27 | 0 | 0.00 | 0.802 | 0.95 | (A) BRAF Class 1 Colorectal |
| KIF5B | 10p11.22 | 6 | 5.22 | 0 | 0.00 | 0.811 | 0.95 | (A) BRAF Class 1 Colorectal |
| PDPK1 | 16p13.3 | 3 | 1.03 | 0 | 0.00 | 0.811 | 0.95 | (A) BRAF Class 1 Colorectal |
| KNSTRN | 15q15.1 | 3 | 2.13 | 0 | 0.00 | 0.813 | 0.95 | (A) BRAF Class 1 Colorectal |
| RACK1 | 5q35.3 | 2 | 3.57 | 0 | 0.00 | 0.814 | 0.95 | (A) BRAF Class 1 Colorectal |
| FGF5 | 4q21.21 | 8 | 9.76 | 0 | 0.00 | 0.818 | 0.95 | (A) BRAF Class 1 Colorectal |
| NCOA2 | 8q13.3 | 2 | 5.13 | 0 | 0.00 | 0.821 | 0.95 | (A) BRAF Class 1 Colorectal |
| MLLT3 | 9p21.3 | 8 | 9.52 | 0 | 0.00 | 0.822 | 0.95 | (A) BRAF Class 1 Colorectal |
| PMAIP1 | 18q21.32 | 3 | 1.04 | 0 | 0.00 | 0.826 | 0.95 | (A) BRAF Class 1 Colorectal |
| RYBP | 3p13 | 3 | 1.04 | 0 | 0.00 | 0.826 | 0.95 | (A) BRAF Class 1 Colorectal |
| FGF23 | 12p13.32 | 3 | 1.96 | 0 | 0.00 | 0.826 | 0.95 | (A) BRAF Class 1 Colorectal |
| FGF6 | 12p13.32 | 3 | 1.96 | 0 | 0.00 | 0.826 | 0.95 | (A) BRAF Class 1 Colorectal |

| | | | | | | | | |
|---|---|---|---|---|---|---|---|---|
| IGF2R | 6q25.3 | 2 | 3.92 | 0 | 0.00 | 0.828 | 0.95 | (A) BRAF Class 1 Colorectal |
| CCNQ | Xq28 | 3 | 2.36 | 0 | 0.00 | 0.831 | 0.95 | (A) BRAF Class 1 Colorectal |
| TAP2 | 6p21.32 | 3 | 2.36 | 0 | 0.00 | 0.831 | 0.95 | (A) BRAF Class 1 Colorectal |
| HIST3H3 | 1q42.13 | 3 | 1.11 | 0 | 0.00 | 0.833 | 0.95 | (A) BRAF Class 1 Colorectal |
| EML4 | 2p21 | 5 | 4.31 | 0 | 0.00 | 0.842 | 0.95 | (A) BRAF Class 1 Colorectal |
| JAZF1 | 'p15.2-p15.: | 2 | 1.01 | 0 | 0.00 | 0.848 | 0.95 | (A) BRAF Class 1 Colorectal |
| CD74 | 5q33.1 | 3 | 2.38 | 0 | 0.00 | 0.849 | 0.95 | (A) BRAF Class 1 Colorectal |
| PPP2R2A | 8p21.2 | 3 | 3.26 | 0 | 0.00 | 0.852 | 0.95 | (A) BRAF Class 1 Colorectal |
| TYRO3 | 15q15.1 | 1 | 4.35 | 0 | 0.00 | 0.852 | 0.95 | (A) BRAF Class 1 Colorectal |
| DCLRE1C | 10p13 | 2 | 1.24 | 0 | 0.00 | 0.856 | 0.95 | (A) BRAF Class 1 Colorectal |
| EME1 | 17q21.33 | 2 | 1.24 | 0 | 0.00 | 0.856 | 0.95 | (A) BRAF Class 1 Colorectal |
| HFE | 6p22.2 | 2 | 1.24 | 0 | 0.00 | 0.856 | 0.95 | (A) BRAF Class 1 Colorectal |
| HMBS | 11q23.3 | 2 | 1.24 | 0 | 0.00 | 0.856 | 0.95 | (A) BRAF Class 1 Colorectal |
| UBE2T | 1q32.1 | 2 | 1.24 | 0 | 0.00 | 0.856 | 0.95 | (A) BRAF Class 1 Colorectal |
| XRCC4 | 5q14.2 | 2 | 1.24 | 0 | 0.00 | 0.856 | 0.95 | (A) BRAF Class 1 Colorectal |
| PTCH2 | 1p34.1 | 1 | 5.56 | 0 | 0.00 | 0.857 | 0.95 | (A) BRAF Class 1 Colorectal |
| EPB41L4A | ;q22.1-q22.: | 2 | 0.24 | 0 | 0.00 | 0.858 | 0.95 | (A) BRAF Class 1 Colorectal |
| MT-ND5 | - | 2 | 0.24 | 0 | 0.00 | 0.858 | 0.95 | (A) BRAF Class 1 Colorectal |
| TRBC2 | 7q34 | 2 | 0.24 | 0 | 0.00 | 0.858 | 0.95 | (A) BRAF Class 1 Colorectal |
| FKBP9 | 7p14.3 | 2 | 2.63 | 0 | 0.00 | 0.858 | 0.95 | (A) BRAF Class 1 Colorectal |
| PSMD13 | 11p15.5 | 2 | 2.63 | 0 | 0.00 | 0.858 | 0.95 | (A) BRAF Class 1 Colorectal |
| RHPN2 | 19q13.11 | 2 | 2.63 | 0 | 0.00 | 0.858 | 0.95 | (A) BRAF Class 1 Colorectal |
| ZNF708 | 19p12 | 2 | 2.63 | 0 | 0.00 | 0.858 | 0.95 | (A) BRAF Class 1 Colorectal |
| HSP90AA1 | 14q32.31 | 4 | 3.01 | 0 | 0.00 | 0.861 | 0.95 | (A) BRAF Class 1 Colorectal |
| TAL1 | 1p33 | 2 | 1.03 | 0 | 0.00 | 0.862 | 0.95 | (A) BRAF Class 1 Colorectal |
| TFRC | 3q29 | 6 | 7.23 | 0 | 0.00 | 0.863 | 0.95 | (A) BRAF Class 1 Colorectal |
| CDC42 | 1p36.12 | 2 | 1.52 | 0 | 0.00 | 0.864 | 0.95 | (A) BRAF Class 1 Colorectal |
| DUSP4 | 8p12 | 2 | 1.52 | 0 | 0.00 | 0.864 | 0.95 | (A) BRAF Class 1 Colorectal |
| RHEB | 7q36.1 | 2 | 0.40 | 0 | 0.00 | 0.864 | 0.95 | (A) BRAF Class 1 Colorectal |
| HIST1H3C | 6p22.2 | 2 | 0.46 | 0 | 0.00 | 0.868 | 0.95 | (A) BRAF Class 1 Colorectal |
| ELOC | 8q21.11 | 2 | 0.46 | 0 | 0.00 | 0.868 | 0.95 | (A) BRAF Class 1 Colorectal |
| H3F3B | 17q25.1 | 2 | 0.43 | 0 | 0.00 | 0.87 | 0.95 | (A) BRAF Class 1 Colorectal |
| LAMP1 | 13q34 | 4 | 3.48 | 0 | 0.00 | 0.871 | 0.95 | (A) BRAF Class 1 Colorectal |
| SH2D1A | Xq25 | 2 | 0.41 | 0 | 0.00 | 0.873 | 0.95 | (A) BRAF Class 1 Colorectal |
| RAD51B | 14q24.1 | 2 | 0.61 | 0 | 0.00 | 0.873 | 0.95 | (A) BRAF Class 1 Colorectal |
| KMT5A | 12q24.31 | 2 | 1.49 | 0 | 0.00 | 0.878 | 0.95 | (A) BRAF Class 1 Colorectal |
| IL10 | 1q32.1 | 2 | 0.69 | 0 | 0.00 | 0.88 | 0.95 | (A) BRAF Class 1 Colorectal |
| FGF10 | 5p12 | 2 | 1.31 | 0 | 0.00 | 0.881 | 0.95 | (A) BRAF Class 1 Colorectal |
| GNA13 | 17q24.1 | 2 | 1.31 | 0 | 0.00 | 0.881 | 0.95 | (A) BRAF Class 1 Colorectal |
| MSI2 | 17q22 | 2 | 1.57 | 0 | 0.00 | 0.885 | 0.95 | (A) BRAF Class 1 Colorectal |
| PPP4R2 | 3p13 | 2 | 1.57 | 0 | 0.00 | 0.885 | 0.95 | (A) BRAF Class 1 Colorectal |
| RAC2 | 22q13.1 | 2 | 1.57 | 0 | 0.00 | 0.885 | 0.95 | (A) BRAF Class 1 Colorectal |
| SESN1 | 6q21 | 2 | 1.57 | 0 | 0.00 | 0.885 | 0.95 | (A) BRAF Class 1 Colorectal |
| SMYD3 | 1q44 | 2 | 1.57 | 0 | 0.00 | 0.885 | 0.95 | (A) BRAF Class 1 Colorectal |
| CENPA | 2p23.3 | 2 | 0.74 | 0 | 0.00 | 0.885 | 0.95 | (A) BRAF Class 1 Colorectal |
| EIF4E | 4q23 | 2 | 0.74 | 0 | 0.00 | 0.885 | 0.95 | (A) BRAF Class 1 Colorectal |
| HIST1H3G | 6p22.2 | 2 | 0.74 | 0 | 0.00 | 0.885 | 0.95 | (A) BRAF Class 1 Colorectal |
| HIST1H3H | 6p22.1 | 2 | 0.74 | 0 | 0.00 | 0.885 | 0.95 | (A) BRAF Class 1 Colorectal |

| | | | | | | | | |
|---|---|---|---|---|---|---|---|---|
| HIST1H3I | 6p22.1 | 2 | 0.74 | 0 | 0.00 | 0.885 | 0.95 | (A) BRAF Class 1 Colorectal |
| SYNE1 | 6q25.2 | 3 | 2.42 | 0 | 0.00 | 0.887 | 0.95 | (A) BRAF Class 1 Colorectal |
| DNAH9 | 17p12 | 1 | 12.50 | 0 | 0.00 | 0.889 | 0.95 | (A) BRAF Class 1 Colorectal |
| CD22 | 19q13.12 | 1 | 2.38 | 0 | 0.00 | 0.894 | 0.95 | (A) BRAF Class 1 Colorectal |
| PRSS8 | 16p11.2 | 3 | 2.21 | 0 | 0.00 | 0.897 | 0.95 | (A) BRAF Class 1 Colorectal |
| THADA | 2p21 | 1 | 2.86 | 0 | 0.00 | 0.897 | 0.95 | (A) BRAF Class 1 Colorectal |
| BIRC5 | 17q25.3 | 1 | 2.78 | 0 | 0.00 | 0.9 | 0.95 | (A) BRAF Class 1 Colorectal |
| SAMD9 | 7q21.2 | 1 | 2.78 | 0 | 0.00 | 0.9 | 0.95 | (A) BRAF Class 1 Colorectal |
| XBP1 | 2q12.1\|22q1 | 1 | 2.70 | 0 | 0.00 | 0.902 | 0.95 | (A) BRAF Class 1 Colorectal |
| ZMYM3 | Xq13.1 | 1 | 2.70 | 0 | 0.00 | 0.902 | 0.95 | (A) BRAF Class 1 Colorectal |
| LTK | 15q15.1 | 1 | 1.79 | 0 | 0.00 | 0.903 | 0.95 | (A) BRAF Class 1 Colorectal |
| NPRL2 | 3p21.31 | 1 | 1.79 | 0 | 0.00 | 0.903 | 0.95 | (A) BRAF Class 1 Colorectal |
| NPRL3 | 16p13.3 | 1 | 1.79 | 0 | 0.00 | 0.903 | 0.95 | (A) BRAF Class 1 Colorectal |
| FRS2 | 12q15 | 2 | 1.47 | 0 | 0.00 | 0.904 | 0.95 | (A) BRAF Class 1 Colorectal |
| TFEB | 6p21.1 | 1 | 2.63 | 0 | 0.00 | 0.905 | 0.95 | (A) BRAF Class 1 Colorectal |
| ETS1 | 11q24.3 | 2 | 1.67 | 0 | 0.00 | 0.907 | 0.95 | (A) BRAF Class 1 Colorectal |
| MYH9 | 22q12.3 | 1 | 2.56 | 0 | 0.00 | 0.907 | 0.95 | (A) BRAF Class 1 Colorectal |
| PDGFB | 22q13.1 | 1 | 2.56 | 0 | 0.00 | 0.907 | 0.95 | (A) BRAF Class 1 Colorectal |
| PAX8 | 2q14.1 | 2 | 1.65 | 0 | 0.00 | 0.907 | 0.95 | (A) BRAF Class 1 Colorectal |
| HNRNPK | 9q21.32 | 2 | 2.33 | 0 | 0.00 | 0.913 | 0.95 | (A) BRAF Class 1 Colorectal |
| FAAP20 | 1p36.33 | 1 | 0.62 | 0 | 0.00 | 0.925 | 0.95 | (A) BRAF Class 1 Colorectal |
| FAAP24 | 19q13.11 | 1 | 0.62 | 0 | 0.00 | 0.925 | 0.95 | (A) BRAF Class 1 Colorectal |
| KLLN | 10q23.31 | 1 | 0.62 | 0 | 0.00 | 0.925 | 0.95 | (A) BRAF Class 1 Colorectal |
| NEIL2 | 8p23.1 | 1 | 0.62 | 0 | 0.00 | 0.925 | 0.95 | (A) BRAF Class 1 Colorectal |
| SLX1B | 16p11.2 | 1 | 0.62 | 0 | 0.00 | 0.925 | 0.95 | (A) BRAF Class 1 Colorectal |
| TAZ | Xq28 | 1 | 0.62 | 0 | 0.00 | 0.925 | 0.95 | (A) BRAF Class 1 Colorectal |
| TAL2 | 9q31.2 | 1 | 0.62 | 0 | 0.00 | 0.926 | 0.95 | (A) BRAF Class 1 Colorectal |
| LMO2 | 11p13 | 1 | 0.42 | 0 | 0.00 | 0.926 | 0.95 | (A) BRAF Class 1 Colorectal |
| ANKRD30A | 10p11.21 | 1 | 0.12 | 0 | 0.00 | 0.926 | 0.95 | (A) BRAF Class 1 Colorectal |
| AP1G1 | 16q22.2 | 1 | 0.12 | 0 | 0.00 | 0.926 | 0.95 | (A) BRAF Class 1 Colorectal |
| CAMK1 | 3p25.3 | 1 | 0.12 | 0 | 0.00 | 0.926 | 0.95 | (A) BRAF Class 1 Colorectal |
| CUL1 | 7q36.1 | 1 | 0.12 | 0 | 0.00 | 0.926 | 0.95 | (A) BRAF Class 1 Colorectal |
| EIF3E | 8q23.1 | 1 | 0.12 | 0 | 0.00 | 0.926 | 0.95 | (A) BRAF Class 1 Colorectal |
| EMID1 | 22q12.2 | 1 | 0.12 | 0 | 0.00 | 0.926 | 0.95 | (A) BRAF Class 1 Colorectal |
| F11-AS1 | 4q35.2 | 1 | 0.12 | 0 | 0.00 | 0.926 | 0.95 | (A) BRAF Class 1 Colorectal |
| GOLT1A | 1q32.1 | 1 | 0.12 | 0 | 0.00 | 0.926 | 0.95 | (A) BRAF Class 1 Colorectal |
| GRAPL | 17p11.2 | 1 | 0.12 | 0 | 0.00 | 0.926 | 0.95 | (A) BRAF Class 1 Colorectal |
| HLA-DOB | 6p21.32 | 1 | 0.12 | 0 | 0.00 | 0.926 | 0.95 | (A) BRAF Class 1 Colorectal |
| HOOK2 | 19p13.13 | 1 | 0.12 | 0 | 0.00 | 0.926 | 0.95 | (A) BRAF Class 1 Colorectal |
| NDUFB2 | 7q34 | 1 | 0.12 | 0 | 0.00 | 0.926 | 0.95 | (A) BRAF Class 1 Colorectal |
| PLEKHA7 | 1p15.2-p15. | 1 | 0.12 | 0 | 0.00 | 0.926 | 0.95 | (A) BRAF Class 1 Colorectal |
| SCAP | 3p21.31 | 1 | 0.12 | 0 | 0.00 | 0.926 | 0.95 | (A) BRAF Class 1 Colorectal |
| TRAJ32 | 14q11.2 | 1 | 0.12 | 0 | 0.00 | 0.926 | 0.95 | (A) BRAF Class 1 Colorectal |
| TRBJ2-2 | 7q34 | 1 | 0.12 | 0 | 0.00 | 0.926 | 0.95 | (A) BRAF Class 1 Colorectal |
| TRGJP | 7p14.1 | 1 | 0.12 | 0 | 0.00 | 0.926 | 0.95 | (A) BRAF Class 1 Colorectal |
| TSPOAP1 | 17q22 | 1 | 0.12 | 0 | 0.00 | 0.926 | 0.95 | (A) BRAF Class 1 Colorectal |
| CDK1 | 10q21.2 | 1 | 1.32 | 0 | 0.00 | 0.927 | 0.95 | (A) BRAF Class 1 Colorectal |
| CDK9 | 9q34.11 | 1 | 1.32 | 0 | 0.00 | 0.927 | 0.95 | (A) BRAF Class 1 Colorectal |

| | | | | | | | | |
|---|---|---|---|---|---|---|---|---|
| GSTM5 | 1p13.3 | 1 | 1.32 | 0 | 0.00 | 0.927 | 0.95 | (A) BRAF Class 1 Colorectal |
| ZBTB2 | 6q25.1 | 2 | 1.53 | 0 | 0.00 | 0.928 | 0.95 | (A) BRAF Class 1 Colorectal |
| PRSS1 | 7q34 | 1 | 0.49 | 0 | 0.00 | 0.928 | 0.95 | (A) BRAF Class 1 Colorectal |
| ID3 | 1p36.12 | 1 | 0.18 | 0 | 0.00 | 0.928 | 0.95 | (A) BRAF Class 1 Colorectal |
| WWTR1 | 3q25.1 | 1 | 0.76 | 0 | 0.00 | 0.929 | 0.95 | (A) BRAF Class 1 Colorectal |
| CSNK1A1 | 5q32 | 3 | 3.66 | 0 | 0.00 | 0.929 | 0.95 | (A) BRAF Class 1 Colorectal |
| FGF2 | 4q28.1 | 3 | 3.66 | 0 | 0.00 | 0.929 | 0.95 | (A) BRAF Class 1 Colorectal |
| RPS6KB1 | 17q23.1 | 3 | 3.66 | 0 | 0.00 | 0.929 | 0.95 | (A) BRAF Class 1 Colorectal |
| RRAS2 | 11p15.2 | 1 | 0.76 | 0 | 0.00 | 0.93 | 0.95 | (A) BRAF Class 1 Colorectal |
| FLI1 | 11q24.3 | 2 | 1.74 | 0 | 0.00 | 0.934 | 0.95 | (A) BRAF Class 1 Colorectal |
| ARFRP1 | 20q13.33 | 1 | 0.66 | 0 | 0.00 | 0.938 | 0.95 | (A) BRAF Class 1 Colorectal |
| CDH20 | 18q21.33 | 1 | 2.17 | 0 | 0.00 | 0.939 | 0.95 | (A) BRAF Class 1 Colorectal |
| CDH5 | 16q21 | 1 | 2.17 | 0 | 0.00 | 0.939 | 0.95 | (A) BRAF Class 1 Colorectal |
| HIST2H3D | 1q21.2 | 1 | 0.37 | 0 | 0.00 | 0.941 | 0.95 | (A) BRAF Class 1 Colorectal |
| HIST1H3D | 6p22.2 | 1 | 0.37 | 0 | 0.00 | 0.941 | 0.95 | (A) BRAF Class 1 Colorectal |
| HIST1H3F | 6p22.2 | 1 | 0.37 | 0 | 0.00 | 0.941 | 0.95 | (A) BRAF Class 1 Colorectal |
| MAP3K14 | 17q21.31 | 1 | 0.33 | 0 | 0.00 | 0.941 | 0.95 | (A) BRAF Class 1 Colorectal |
| ACVR2A | !q22.3-q23.: | 1 | 3.03 | 0 | 0.00 | 0.943 | 0.95 | (A) BRAF Class 1 Colorectal |
| ADAMTS20 | 12q12 | 1 | 3.03 | 0 | 0.00 | 0.943 | 0.95 | (A) BRAF Class 1 Colorectal |
| BLNK | 10q24.1 | 1 | 3.03 | 0 | 0.00 | 0.943 | 0.95 | (A) BRAF Class 1 Colorectal |
| CSMD3 | 8q23.3 | 1 | 3.03 | 0 | 0.00 | 0.943 | 0.95 | (A) BRAF Class 1 Colorectal |
| FOXP4 | 6p21.1 | 1 | 3.03 | 0 | 0.00 | 0.943 | 0.95 | (A) BRAF Class 1 Colorectal |
| IRF8 | 16q24.1 | 1 | 3.03 | 0 | 0.00 | 0.943 | 0.95 | (A) BRAF Class 1 Colorectal |
| ITGA9 | 3p22.2 | 1 | 3.03 | 0 | 0.00 | 0.943 | 0.95 | (A) BRAF Class 1 Colorectal |
| RRM1 | 11p15.4 | 1 | 3.03 | 0 | 0.00 | 0.943 | 0.95 | (A) BRAF Class 1 Colorectal |
| TAF1L | 9p21.1 | 1 | 3.03 | 0 | 0.00 | 0.943 | 0.95 | (A) BRAF Class 1 Colorectal |
| THBS1 | 15q14 | 1 | 3.03 | 0 | 0.00 | 0.943 | 0.95 | (A) BRAF Class 1 Colorectal |
| TRIM24 | 7q33-q34 | 1 | 3.03 | 0 | 0.00 | 0.943 | 0.95 | (A) BRAF Class 1 Colorectal |
| AFDN | 6q27 | 1 | 2.94 | 0 | 0.00 | 0.944 | 0.95 | (A) BRAF Class 1 Colorectal |
| AFF1 | ‹q21.3-q22.: | 1 | 2.94 | 0 | 0.00 | 0.944 | 0.95 | (A) BRAF Class 1 Colorectal |
| BCL9 | 1q21.2 | 1 | 2.94 | 0 | 0.00 | 0.944 | 0.95 | (A) BRAF Class 1 Colorectal |
| FOXO3 | 6q21 | 1 | 2.94 | 0 | 0.00 | 0.944 | 0.95 | (A) BRAF Class 1 Colorectal |
| KNL1 | 15q15.1 | 1 | 2.94 | 0 | 0.00 | 0.944 | 0.95 | (A) BRAF Class 1 Colorectal |
| LCK | 1p35.2 | 1 | 2.94 | 0 | 0.00 | 0.944 | 0.95 | (A) BRAF Class 1 Colorectal |
| MUC1 | 1q22 | 1 | 2.94 | 0 | 0.00 | 0.944 | 0.95 | (A) BRAF Class 1 Colorectal |
| NIN | 14q22.1 | 1 | 2.94 | 0 | 0.00 | 0.944 | 0.95 | (A) BRAF Class 1 Colorectal |
| NUP214 | 9q34.13 | 1 | 2.94 | 0 | 0.00 | 0.944 | 0.95 | (A) BRAF Class 1 Colorectal |
| PDE4DIP | 1q21.2 | 1 | 2.94 | 0 | 0.00 | 0.944 | 0.95 | (A) BRAF Class 1 Colorectal |
| RNF213 | 17q25.3 | 1 | 2.94 | 0 | 0.00 | 0.944 | 0.95 | (A) BRAF Class 1 Colorectal |
| TLX1 | 10q24.31 | 1 | 2.94 | 0 | 0.00 | 0.944 | 0.95 | (A) BRAF Class 1 Colorectal |
| UBR5 | 8q22.3 | 1 | 2.94 | 0 | 0.00 | 0.944 | 0.95 | (A) BRAF Class 1 Colorectal |
| GID4 | ɔ11.2\|17p1: | 1 | 0.68 | 0 | 0.00 | 0.949 | 0.953 | (A) BRAF Class 1 Colorectal |
| FGF9 | 13q12.11 | 2 | 2.44 | 0 | 0.00 | 0.953 | 0.956 | (A) BRAF Class 1 Colorectal |
| DDX41 | 5q35.3 | 1 | 1.15 | 0 | 0.00 | 0.956 | 0.958 | (A) BRAF Class 1 Colorectal |
| FGF8 | 10q24.32 | 1 | 1.22 | 0 | 0.00 | 0.976 | 0.976 | (A) BRAF Class 1 Colorectal |
| TSPAN31 | 12q14.1 | 1 | 1.22 | 0 | 0.00 | 0.976 | 0.976 | (A) BRAF Class 1 Colorectal |

#### Supplementary Table 12

| Gene | Cytoband | Class 1 Occurrence | Class 1 % | Class 2 Occurrence | Class 2 % | SumClass | pval | qval | Most Enriched In |
|---|---|---|---|---|---|---|---|---|---|
| NRAS | 1p13.2 | 14 | 1.67 | 7 | 10.45 | 21 | 0.00045 | 0.0987 | (B) BRAF Class 2 Colorectal |
| TGFBR2 | 3p24.1 | 92 | 25.56 | 0 | 0.00 | 92 | 0.00049 | 0.0987 | (A) BRAF Class 1 Colorectal |
| RNF43 | 17q22 | 227 | 41.20 | 2 | 4.76 | 229 | 2.2E-07 | 8.8E-05 | (A) BRAF Class 1 Colorectal |
| KRAS | 12p12.1 | 15 | 1.79 | 12 | 17.91 | 27 | 7.2E-08 | 5.8E-05 | (B) BRAF Class 2 Colorectal |

#### Supplementary Table 13

| Gene | Cytoband | Class 1 Occurrence | Class 1 % | Class 3 Occurrence | Class 3 % | pval | qval | Enriched In |
|---|---|---|---|---|---|---|---|---|
| APC | 5q22.2 | 234 | 29.40 | 90 | 69.23 | 7.89E-18 | 6.58E-15 | (B) BRAF Class 3 Colorectal |
| RNF43 | 17q22 | 226 | 41.09 | 4 | 4.40 | 3.33E-14 | 1.39E-11 | (A) BRAF Class 1 Colorectal |
| KRAS | 12p12.1 | 15 | 1.79 | 23 | 17.16 | 7.27E-12 | 2.02E-09 | (B) BRAF Class 3 Colorectal |
| FAT1 | 4q35.2 | 113 | 22.47 | 1 | 1.20 | 8.36E-08 | 1.74E-05 | (A) BRAF Class 1 Colorectal |
| BCOR | Xp11.4 | 89 | 14.93 | 0 | 0.00 | 3.11E-07 | 5.19E-05 | (A) BRAF Class 1 Colorectal |
| MSH6 | 2p16.3 | 88 | 14.06 | 0 | 0.00 | 5.15E-07 | 7.15E-05 | (A) BRAF Class 1 Colorectal |
| KMT2D | 12q13.12 | 177 | 29.70 | 10 | 9.90 | 6.87E-06 | 0.000818 | (A) BRAF Class 1 Colorectal |
| MSH3 | 5q14.1 | 76 | 29.69 | 0 | 0.00 | 8.35E-06 | 0.00087 | (A) BRAF Class 1 Colorectal |
| KMT2A | 11q23.3 | 106 | 17.55 | 3 | 2.88 | 1.29E-05 | 0.001196 | (A) BRAF Class 1 Colorectal |
| ARID1B | 6q25.3 | 106 | 18.31 | 3 | 3.13 | 2.1E-05 | 0.001667 | (A) BRAF Class 1 Colorectal |
| ZFHX3 | 5q22.2-q22. | 85 | 30.80 | 2 | 4.26 | 2.39E-05 | 0.001667 | (A) BRAF Class 1 Colorectal |
| PTPRD | 9p24.1-p23 | 71 | 17.93 | 1 | 1.35 | 2.43E-05 | 0.001667 | (A) BRAF Class 1 Colorectal |
| PTPRS | 19p13.3 | 78 | 26.99 | 1 | 2.17 | 2.66E-05 | 0.001667 | (A) BRAF Class 1 Colorectal |
| POLE | 12q24.33 | 84 | 15.25 | 1 | 1.16 | 2.8E-05 | 0.001667 | (A) BRAF Class 1 Colorectal |
| CREBBP | 16p13.3 | 110 | 18.18 | 4 | 3.92 | 4.69E-05 | 0.002608 | (A) BRAF Class 1 Colorectal |
| CIC | 19q13.2 | 86 | 16.54 | 2 | 2.27 | 5.62E-05 | 0.002932 | (A) BRAF Class 1 Colorectal |
| AR | Xq12 | 70 | 11.57 | 1 | 0.96 | 9.93E-05 | 0.004505 | (A) BRAF Class 1 Colorectal |
| EP300 | 22q13.2 | 94 | 15.77 | 3 | 2.97 | 9.96E-05 | 0.004505 | (A) BRAF Class 1 Colorectal |
| SETD2 | 3p21.31 | 71 | 11.74 | 1 | 0.98 | 0.000103 | 0.004505 | (A) BRAF Class 1 Colorectal |
| NRAS | 1p13.2 | 13 | 1.55 | 11 | 8.21 | 0.000117 | 0.00488 | (B) BRAF Class 3 Colorectal |
| PTCH1 | 9q22.32 | 82 | 13.55 | 2 | 1.96 | 0.000124 | 0.004933 | (A) BRAF Class 1 Colorectal |
| FLT4 | 5q35.3 | 81 | 13.82 | 2 | 2.02 | 0.000138 | 0.005243 | (A) BRAF Class 1 Colorectal |
| BLM | 15q26.1 | 68 | 11.60 | 1 | 1.01 | 0.00016 | 0.005624 | (A) BRAF Class 1 Colorectal |
| HNF1A | 12q24.31 | 71 | 9.11 | 1 | 0.79 | 0.000162 | 0.005624 | (A) BRAF Class 1 Colorectal |
| CASP8 | 2q33.1 | 74 | 14.40 | 2 | 2.35 | 0.000452 | 0.0151 | (A) BRAF Class 1 Colorectal |
| NOTCH1 | 9q34.3 | 96 | 11.85 | 4 | 3.03 | 0.000616 | 0.0195 | (A) BRAF Class 1 Colorectal |
| TET1 | 10q21.3 | 64 | 13.25 | 1 | 1.35 | 0.000632 | 0.0195 | (A) BRAF Class 1 Colorectal |
| SPEN | 36.21-p36. | 64 | 17.78 | 2 | 3.08 | 0.000736 | 0.0218 | (A) BRAF Class 1 Colorectal |
| BRD4 | 19p13.12 | 59 | 10.00 | 1 | 1.02 | 0.000758 | 0.0218 | (A) BRAF Class 1 Colorectal |
| NOTCH3 | 19p13.12 | 111 | 20.26 | 6 | 6.82 | 0.000923 | 0.0257 | (A) BRAF Class 1 Colorectal |
| AKT1 | 14q32.33 | 72 | 8.59 | 2 | 1.49 | 0.001026 | 0.0273 | (A) BRAF Class 1 Colorectal |
| PGR | 11q22.1 | 44 | 16.24 | 0 | 0.00 | 0.001048 | 0.0273 | (A) BRAF Class 1 Colorectal |
| DOT1L | 19p13.3 | 39 | 10.83 | 0 | 0.00 | 0.001102 | 0.0279 | (A) BRAF Class 1 Colorectal |
| ATRX | Xq21.1 | 87 | 14.38 | 4 | 3.92 | 0.001165 | 0.0286 | (A) BRAF Class 1 Colorectal |
| KIT | 4q12 | 44 | 5.25 | 0 | 0.00 | 0.001246 | 0.0293 | (A) BRAF Class 1 Colorectal |
| BRCA1 | 17q21.31 | 54 | 8.93 | 1 | 0.96 | 0.001263 | 0.0293 | (A) BRAF Class 1 Colorectal |
| HIST1H3C | 6p22.2 | 1 | 0.23 | 4 | 5.97 | 0.001328 | 0.0299 | (B) BRAF Class 3 Colorectal |
| DNMT3A | 2p23.3 | 53 | 8.77 | 1 | 0.96 | 0.001456 | 0.032 | (A) BRAF Class 1 Colorectal |
| IGF1R | 15q26.3 | 53 | 8.65 | 1 | 0.95 | 0.001519 | 0.0325 | (A) BRAF Class 1 Colorectal |
| GLI1 | 12q13.3 | 66 | 11.76 | 2 | 2.17 | 0.001618 | 0.0337 | (A) BRAF Class 1 Colorectal |
| TGFBR2 | 3p24.1 | 92 | 25.56 | 6 | 9.23 | 0.001847 | 0.0372 | (A) BRAF Class 1 Colorectal |
| POLD1 | 19q13.33 | 79 | 15.11 | 3 | 3.70 | 0.001876 | 0.0372 | (A) BRAF Class 1 Colorectal |
| HLA-A | 6p22.1 | 37 | 12.80 | 0 | 0.00 | 0.002117 | 0.0411 | (A) BRAF Class 1 Colorectal |
| BRCA2 | 13q13.1 | 100 | 16.53 | 6 | 5.88 | 0.002187 | 0.0415 | (A) BRAF Class 1 Colorectal |
| BCORL1 | Xq26.1 | 76 | 19.54 | 3 | 5.00 | 0.002362 | 0.0438 | (A) BRAF Class 1 Colorectal |
| FGFR4 | 5q35.2 | 35 | 5.71 | 0 | 0.00 | 0.003087 | 0.056 | (A) BRAF Class 1 Colorectal |

| | | | | | | | | |
|---|---|---|---|---|---|---|---|---|
| TCF3 | 19p13.3 | 39 | 7.21 | 0 | 0.00 | 0.00316 | 0.0561 | (A) BRAF Class 1 Colorectal |
| WT1 | 11p13 | 35 | 5.79 | 0 | 0.00 | 0.003311 | 0.0575 | (A) BRAF Class 1 Colorectal |
| ATM | 11q22.3 | 117 | 14.92 | 8 | 6.30 | 0.003992 | 0.0679 | (A) BRAF Class 1 Colorectal |
| KMT2C | 7q36.1 | 72 | 20.99 | 4 | 6.67 | 0.004257 | 0.071 | (A) BRAF Class 1 Colorectal |
| AXIN1 | 16p13.3 | 32 | 9.04 | 0 | 0.00 | 0.004583 | 0.0744 | (A) BRAF Class 1 Colorectal |
| KEAP1 | 19p13.2 | 33 | 5.73 | 0 | 0.00 | 0.004639 | 0.0744 | (A) BRAF Class 1 Colorectal |
| TNFAIP3 | 6q23.3 | 33 | 5.54 | 0 | 0.00 | 0.004997 | 0.0762 | (A) BRAF Class 1 Colorectal |
| IL7R | 5p13.2 | 33 | 6.47 | 0 | 0.00 | 0.005002 | 0.0762 | (A) BRAF Class 1 Colorectal |
| SDHA | 5p15.33 | 33 | 5.81 | 0 | 0.00 | 0.005028 | 0.0762 | (A) BRAF Class 1 Colorectal |
| GNAS | 20q13.32 | 72 | 8.64 | 3 | 2.33 | 0.005291 | 0.0774 | (A) BRAF Class 1 Colorectal |
| ANKRD11 | 16q24.3 | 47 | 17.34 | 1 | 2.38 | 0.005292 | 0.0774 | (A) BRAF Class 1 Colorectal |
| RECQL4 | 8q24.3 | 58 | 10.70 | 2 | 2.30 | 0.005568 | 0.0783 | (A) BRAF Class 1 Colorectal |
| KDR | 4q12 | 47 | 5.99 | 1 | 0.79 | 0.005583 | 0.0783 | (A) BRAF Class 1 Colorectal |
| MTOR | 1p36.22 | 72 | 11.80 | 4 | 3.74 | 0.005635 | 0.0783 | (A) BRAF Class 1 Colorectal |
| GATA2 | 3q21.3 | 32 | 6.17 | 0 | 0.00 | 0.006086 | 0.0807 | (A) BRAF Class 1 Colorectal |
| INSR | 19p13.2 | 32 | 9.58 | 0 | 0.00 | 0.00609 | 0.0807 | (A) BRAF Class 1 Colorectal |
| MSH2 | 2p21-p16.3 | 45 | 7.48 | 1 | 1.00 | 0.006093 | 0.0807 | (A) BRAF Class 1 Colorectal |
| EGFR | 7p11.2 | 46 | 5.49 | 1 | 0.75 | 0.006984 | 0.0904 | (A) BRAF Class 1 Colorectal |
| RPTOR | 17q25.3 | 43 | 8.27 | 1 | 1.14 | 0.007106 | 0.0904 | (A) BRAF Class 1 Colorectal |
| ETAA1 | 2p14 | 1 | 100.00 | 0 | 0.00 | 0.007407 | 0.0904 | (A) BRAF Class 1 Colorectal |
| OLIG2 | 21q22.11 | 1 | 100.00 | 0 | 0.00 | 0.007407 | 0.0904 | (A) BRAF Class 1 Colorectal |
| NTRK3 | 15q25.3 | 43 | 7.11 | 1 | 0.98 | 0.007478 | 0.0904 | (A) BRAF Class 1 Colorectal |
| ATR | 3q23 | 72 | 13.61 | 4 | 4.49 | 0.007479 | 0.0904 | (A) BRAF Class 1 Colorectal |
| ALK | p23.2-p23. | 76 | 9.07 | 4 | 2.99 | 0.008118 | 0.0967 | (A) BRAF Class 1 Colorectal |
| KDM5A | 12p13.33 | 79 | 15.19 | 5 | 5.68 | 0.008561 | 0.101 | (A) BRAF Class 1 Colorectal |
| CHEK2 | 22q12.1 | 29 | 4.79 | 0 | 0.00 | 0.009873 | 0.114 | (A) BRAF Class 1 Colorectal |
| ARID1A | 1p36.11 | 140 | 23.14 | 13 | 12.75 | 0.0102 | 0.115 | (A) BRAF Class 1 Colorectal |
| SLX4 | 16p13.3 | 49 | 12.96 | 1 | 1.96 | 0.0103 | 0.115 | (A) BRAF Class 1 Colorectal |
| KDM5C | Xp11.22 | 40 | 6.94 | 1 | 1.01 | 0.0103 | 0.115 | (A) BRAF Class 1 Colorectal |
| ASXL2 | 2p23.3 | 27 | 9.18 | 0 | 0.00 | 0.011 | 0.121 | (A) BRAF Class 1 Colorectal |
| EPHA5 | q13.1-q13. | 56 | 13.15 | 3 | 3.95 | 0.0114 | 0.123 | (A) BRAF Class 1 Colorectal |
| NSD1 | 5q35.3 | 50 | 9.80 | 2 | 2.33 | 0.0115 | 0.123 | (A) BRAF Class 1 Colorectal |
| AXIN2 | 17q24.1 | 53 | 10.91 | 2 | 2.63 | 0.0121 | 0.128 | (A) BRAF Class 1 Colorectal |
| ABL1 | 9q34.12 | 52 | 6.53 | 2 | 1.56 | 0.0129 | 0.134 | (A) BRAF Class 1 Colorectal |
| PHOX2B | 4p13 | 28 | 4.96 | 0 | 0.00 | 0.0132 | 0.134 | (A) BRAF Class 1 Colorectal |
| ERG | 21q22.2 | 27 | 5.19 | 0 | 0.00 | 0.0133 | 0.134 | (A) BRAF Class 1 Colorectal |
| ARID2 | 12q12 | 67 | 11.43 | 4 | 4.04 | 0.0136 | 0.134 | (A) BRAF Class 1 Colorectal |
| NF1 | 17q11.2 | 66 | 10.91 | 4 | 3.85 | 0.0136 | 0.134 | (A) BRAF Class 1 Colorectal |
| NTRK2 | 9q21.33 | 27 | 4.46 | 0 | 0.00 | 0.0137 | 0.134 | (A) BRAF Class 1 Colorectal |
| EP400 | 12q24.33 | 0 | 0.00 | 2 | 40.00 | 0.0142 | 0.134 | (B) BRAF Class 3 Colorectal |
| STAT5B | 17q21.2 | 30 | 9.87 | 0 | 0.00 | 0.0143 | 0.134 | (A) BRAF Class 1 Colorectal |
| CUX1 | 7q22.1 | 58 | 16.34 | 2 | 4.17 | 0.0143 | 0.134 | (A) BRAF Class 1 Colorectal |
| HLA-B | 6p21.33 | 21 | 15.00 | 0 | 0.00 | 0.0145 | 0.134 | (A) BRAF Class 1 Colorectal |
| DICER1 | 14q32.13 | 48 | 8.26 | 2 | 2.02 | 0.0145 | 0.134 | (A) BRAF Class 1 Colorectal |
| MST1R | 3p21.31 | 27 | 8.28 | 0 | 0.00 | 0.0146 | 0.134 | (A) BRAF Class 1 Colorectal |
| EPHA7 | 6q16.1 | 35 | 8.71 | 1 | 1.39 | 0.0166 | 0.151 | (A) BRAF Class 1 Colorectal |
| NBN | 8q21.3 | 47 | 7.82 | 2 | 1.96 | 0.017 | 0.152 | (A) BRAF Class 1 Colorectal |
| CDK12 | 17q12 | 37 | 6.99 | 1 | 1.12 | 0.0175 | 0.155 | (A) BRAF Class 1 Colorectal |

| | | | | | | | | |
|---|---|---|---|---|---|---|---|---|
| TET2 | 4q24 | 46 | 7.62 | 2 | 1.92 | 0.0179 | 0.157 | (A) BRAF Class 1 Colorectal |
| PRKDC | 8q11.21 | 78 | 20.58 | 5 | 8.62 | 0.0182 | 0.158 | (A) BRAF Class 1 Colorectal |
| ROS1 | 6q22.1 | 62 | 10.16 | 4 | 3.74 | 0.019 | 0.163 | (A) BRAF Class 1 Colorectal |
| ERCC2 | 19q13.32 | 36 | 6.09 | 1 | 1.00 | 0.0199 | 0.169 | (A) BRAF Class 1 Colorectal |
| RUNX1 | 21q22.12 | 36 | 5.85 | 1 | 0.96 | 0.0201 | 0.169 | (A) BRAF Class 1 Colorectal |
| TSC2 | 16p13.3 | 55 | 9.09 | 3 | 2.94 | 0.0204 | 0.169 | (A) BRAF Class 1 Colorectal |
| SYK | 9q22.2 | 23 | 5.28 | 0 | 0.00 | 0.0207 | 0.169 | (A) BRAF Class 1 Colorectal |
| TRAF7 | 16p13.3 | 26 | 5.34 | 0 | 0.00 | 0.021 | 0.169 | (A) BRAF Class 1 Colorectal |
| FLCN | 17p11.2 | 45 | 7.61 | 2 | 1.98 | 0.0211 | 0.169 | (A) BRAF Class 1 Colorectal |
| BMPR1A | 10q23.2 | 36 | 6.26 | 1 | 1.04 | 0.0211 | 0.169 | (A) BRAF Class 1 Colorectal |
| MUTYH | 1p34.1 | 24 | 4.03 | 0 | 0.00 | 0.0218 | 0.169 | (A) BRAF Class 1 Colorectal |
| PRDM1 | 6q21 | 24 | 4.03 | 0 | 0.00 | 0.0218 | 0.169 | (A) BRAF Class 1 Colorectal |
| ARID5B | 10q21.2 | 23 | 7.82 | 0 | 0.00 | 0.0221 | 0.169 | (A) BRAF Class 1 Colorectal |
| NCOR1 | 17p12-p11.[illegible] | 43 | 12.50 | 2 | 3.33 | 0.0221 | 0.169 | (A) BRAF Class 1 Colorectal |
| MED12 | Xq13.1 | 53 | 10.02 | 3 | 3.30 | 0.0224 | 0.169 | (A) BRAF Class 1 Colorectal |
| PIK3C3 | 18q12.3 | 22 | 7.14 | 0 | 0.00 | 0.0225 | 0.169 | (A) BRAF Class 1 Colorectal |
| QKI | 6q26 | 41 | 11.48 | 1 | 2.00 | 0.0226 | 0.169 | (A) BRAF Class 1 Colorectal |
| ARAF | Xp11.3 | 47 | 7.42 | 2 | 1.96 | 0.0227 | 0.169 | (A) BRAF Class 1 Colorectal |
| ETV1 | 7p21.2 | 36 | 6.24 | 1 | 1.06 | 0.0236 | 0.173 | (A) BRAF Class 1 Colorectal |
| SMO | 7q32.1 | 46 | 5.87 | 2 | 1.55 | 0.0236 | 0.173 | (A) BRAF Class 1 Colorectal |
| NOTCH4 | 6p21.32 | 44 | 12.19 | 2 | 3.33 | 0.0252 | 0.183 | (A) BRAF Class 1 Colorectal |
| ZRSR2 | Xp22.2 | 24 | 4.40 | 0 | 0.00 | 0.0258 | 0.185 | (A) BRAF Class 1 Colorectal |
| MEN1 | 11q13.1 | 34 | 5.70 | 1 | 0.99 | 0.0262 | 0.185 | (A) BRAF Class 1 Colorectal |
| NKX2-1 | 14q13.3 | 34 | 5.70 | 1 | 0.99 | 0.0262 | 0.185 | (A) BRAF Class 1 Colorectal |
| RB1 | 13q14.2 | 46 | 5.71 | 2 | 1.55 | 0.0271 | 0.19 | (A) BRAF Class 1 Colorectal |
| MAP3K1 | 5q11.2 | 43 | 7.21 | 2 | 1.98 | 0.0285 | 0.198 | (A) BRAF Class 1 Colorectal |
| PIK3CG | 7q22.3 | 31 | 8.86 | 1 | 1.59 | 0.0292 | 0.201 | (A) BRAF Class 1 Colorectal |
| IRS1 | 2q36.3 | 44 | 15.22 | 2 | 4.35 | 0.0295 | 0.202 | (A) BRAF Class 1 Colorectal |
| CDKN1B | 12p13.1 | 22 | 3.64 | 0 | 0.00 | 0.0307 | 0.206 | (A) BRAF Class 1 Colorectal |
| SH2B3 | 12q24.12 | 33 | 5.85 | 1 | 1.05 | 0.0307 | 0.206 | (A) BRAF Class 1 Colorectal |
| DNMT1 | 19p13.2 | 35 | 10.80 | 1 | 2.00 | 0.0312 | 0.208 | (A) BRAF Class 1 Colorectal |
| INPPL1 | 11q13.4 | 20 | 15.75 | 0 | 0.00 | 0.0321 | 0.212 | (A) BRAF Class 1 Colorectal |
| AXL | 19q13.2 | 51 | 8.43 | 3 | 2.94 | 0.0325 | 0.214 | (A) BRAF Class 1 Colorectal |
| ERBB4 | 2q34 | 80 | 9.85 | 6 | 4.65 | 0.0338 | 0.22 | (A) BRAF Class 1 Colorectal |
| MDC1 | 6p21.33 | 22 | 7.61 | 0 | 0.00 | 0.0346 | 0.222 | (A) BRAF Class 1 Colorectal |
| PTEN | 10q23.31 | 136 | 16.23 | 13 | 9.85 | 0.0347 | 0.222 | (A) BRAF Class 1 Colorectal |
| MRE11 | 11q21 | 32 | 6.05 | 1 | 1.12 | 0.0352 | 0.224 | (A) BRAF Class 1 Colorectal |
| JAK1 | 1p31.3 | 50 | 9.09 | 3 | 3.23 | 0.0354 | 0.224 | (A) BRAF Class 1 Colorectal |
| RET | 10q11.21 | 53 | 6.32 | 3 | 2.24 | 0.0361 | 0.226 | (A) BRAF Class 1 Colorectal |
| IDH2 | 15q26.1 | 33 | 4.16 | 1 | 0.78 | 0.0366 | 0.226 | (A) BRAF Class 1 Colorectal |
| DDR2 | 1q23.3 | 21 | 3.29 | 0 | 0.00 | 0.0367 | 0.226 | (A) BRAF Class 1 Colorectal |
| MYOD1 | 11p15.1 | 21 | 7.19 | 0 | 0.00 | 0.0368 | 0.226 | (A) BRAF Class 1 Colorectal |
| PDGFRB | 5q32 | 41 | 6.78 | 2 | 1.96 | 0.0374 | 0.228 | (A) BRAF Class 1 Colorectal |
| FOXP1 | 3p13 | 19 | 5.51 | 0 | 0.00 | 0.0382 | 0.231 | (A) BRAF Class 1 Colorectal |
| ASXL1 | 20q11.21 | 72 | 11.92 | 6 | 5.77 | 0.0396 | 0.237 | (A) BRAF Class 1 Colorectal |
| MET | 7q31.2 | 33 | 3.98 | 1 | 0.76 | 0.0411 | 0.245 | (A) BRAF Class 1 Colorectal |
| MYBL1 | 8q13.1 | 21 | 7.69 | 0 | 0.00 | 0.0415 | 0.245 | (A) BRAF Class 1 Colorectal |
| MEF2B | 19p13.11 | 20 | 3.36 | 0 | 0.00 | 0.0417 | 0.245 | (A) BRAF Class 1 Colorectal |

| | | | | | | | | |
|---|---|---|---|---|---|---|---|---|
| CEBPA | 19q13.11 | 20 | 3.46 | 0 | 0.00 | 0.0428 | 0.25 | (A) BRAF Class 1 Colorectal |
| SUZ12 | 17q11.2 | 21 | 3.74 | 0 | 0.00 | 0.0433 | 0.251 | (A) BRAF Class 1 Colorectal |
| SMARCA4 | 19p13.2 | 80 | 13.22 | 7 | 6.86 | 0.0436 | 0.251 | (A) BRAF Class 1 Colorectal |
| ERCC4 | 16p13.12 | 20 | 3.44 | 0 | 0.00 | 0.045 | 0.256 | (A) BRAF Class 1 Colorectal |
| JAK3 | 19p13.11 | 50 | 6.21 | 3 | 2.29 | 0.0453 | 0.256 | (A) BRAF Class 1 Colorectal |
| EPHA3 | 3p11.1 | 37 | 8.49 | 2 | 2.56 | 0.0455 | 0.256 | (A) BRAF Class 1 Colorectal |
| MGA | 15q15.1 | 82 | 18.81 | 7 | 10.00 | 0.0461 | 0.256 | (A) BRAF Class 1 Colorectal |
| PDGFRA | 4q12 | 40 | 5.03 | 2 | 1.52 | 0.0467 | 0.256 | (A) BRAF Class 1 Colorectal |
| RAD50 | 5q31.1 | 56 | 10.79 | 4 | 4.60 | 0.0471 | 0.256 | (A) BRAF Class 1 Colorectal |
| CSF3R | 1p34.3 | 30 | 5.51 | 1 | 1.10 | 0.0473 | 0.256 | (A) BRAF Class 1 Colorectal |
| RPS6KA4 | 11q13.1 | 20 | 6.92 | 0 | 0.00 | 0.0474 | 0.256 | (A) BRAF Class 1 Colorectal |
| PPM1D | 17q23.2 | 20 | 4.59 | 0 | 0.00 | 0.0478 | 0.256 | (A) BRAF Class 1 Colorectal |
| RANBP2 | 2q13 | 27 | 20.61 | 0 | 0.00 | 0.0479 | 0.256 | (A) BRAF Class 1 Colorectal |
| CTNNB1 | 3p22.1 | 27 | 3.22 | 9 | 6.72 | 0.0482 | 0.256 | (B) BRAF Class 3 Colorectal |
| SMC3 | 10q25.2 | 12 | 3.37 | 5 | 9.80 | 0.0485 | 0.256 | (B) BRAF Class 3 Colorectal |
| CD79A | 19q13.2 | 18 | 5.26 | 0 | 0.00 | 0.0486 | 0.256 | (A) BRAF Class 1 Colorectal |
| CIITA | 16p13.13 | 22 | 8.18 | 0 | 0.00 | 0.0489 | 0.256 | (A) BRAF Class 1 Colorectal |
| FGFR1 | 8p11.23 | 31 | 3.70 | 1 | 0.75 | 0.0504 | 0.262 | (A) BRAF Class 1 Colorectal |
| ETV6 | 12p13.2 | 29 | 4.90 | 1 | 0.99 | 0.0507 | 0.262 | (A) BRAF Class 1 Colorectal |
| PIK3C2G | 12p12.3 | 28 | 8.83 | 1 | 1.82 | 0.0515 | 0.265 | (A) BRAF Class 1 Colorectal |
| B2M | 15q21.1 | 59 | 10.57 | 4 | 4.60 | 0.052 | 0.266 | (A) BRAF Class 1 Colorectal |
| GATA3 | 10p14 | 38 | 6.38 | 2 | 1.98 | 0.0526 | 0.267 | (A) BRAF Class 1 Colorectal |
| CBL | 11q23.3 | 18 | 2.94 | 0 | 0.00 | 0.056 | 0.282 | (A) BRAF Class 1 Colorectal |
| NSD2 | 4p16.3 | 28 | 8.33 | 1 | 1.75 | 0.0562 | 0.282 | (A) BRAF Class 1 Colorectal |
| JUN | 1p32.1 | 18 | 5.07 | 0 | 0.00 | 0.0564 | 0.282 | (A) BRAF Class 1 Colorectal |
| IDH1 | 2q34 | 30 | 3.58 | 1 | 0.75 | 0.0572 | 0.284 | (A) BRAF Class 1 Colorectal |
| KDM6A | Xp11.3 | 37 | 6.21 | 2 | 1.98 | 0.0593 | 0.293 | (A) BRAF Class 1 Colorectal |
| SOX9 | 17q24.3 | 92 | 15.62 | 9 | 9.18 | 0.0601 | 0.294 | (A) BRAF Class 1 Colorectal |
| TBX3 | 12q24.21 | 27 | 7.54 | 1 | 1.61 | 0.0606 | 0.294 | (A) BRAF Class 1 Colorectal |
| TOP2A | 17q21.2 | 20 | 14.71 | 0 | 0.00 | 0.0606 | 0.294 | (A) BRAF Class 1 Colorectal |
| DIS3 | 13q21.33 | 35 | 6.26 | 2 | 2.02 | 0.0628 | 0.302 | (A) BRAF Class 1 Colorectal |
| FYN | 6q21 | 17 | 6.16 | 0 | 0.00 | 0.0641 | 0.302 | (A) BRAF Class 1 Colorectal |
| TP53 | 17p13.1 | 497 | 59.31 | 88 | 66.67 | 0.0645 | 0.302 | (B) BRAF Class 3 Colorectal |
| CCND1 | 11q13.3 | 17 | 2.81 | 0 | 0.00 | 0.0652 | 0.302 | (A) BRAF Class 1 Colorectal |
| PLCG2 | 16q23.3 | 27 | 8.41 | 1 | 1.85 | 0.066 | 0.302 | (A) BRAF Class 1 Colorectal |
| ERCC5 | 13q33.1 | 48 | 8.59 | 3 | 3.45 | 0.0661 | 0.302 | (A) BRAF Class 1 Colorectal |
| MAPKAP1 | 9q33.3 | 4 | 3.15 | 3 | 13.64 | 0.0662 | 0.302 | (B) BRAF Class 3 Colorectal |
| ZNRF3 | 22q12.1 | 27 | 13.85 | 1 | 3.13 | 0.0666 | 0.302 | (A) BRAF Class 1 Colorectal |
| CDKN2B | 9p21.3 | 11 | 1.82 | 5 | 4.90 | 0.067 | 0.302 | (B) BRAF Class 3 Colorectal |
| SMARCB1 | q11.23\|22c | 18 | 2.24 | 0 | 0.00 | 0.0671 | 0.302 | (A) BRAF Class 1 Colorectal |
| FGFR3 | 4p16.3 | 47 | 5.61 | 3 | 2.24 | 0.0673 | 0.302 | (A) BRAF Class 1 Colorectal |
| MTA1 | 14q32.33 | 19 | 11.88 | 0 | 0.00 | 0.0674 | 0.302 | (A) BRAF Class 1 Colorectal |
| SOCS1 | 16p13.13 | 17 | 2.85 | 0 | 0.00 | 0.0675 | 0.302 | (A) BRAF Class 1 Colorectal |
| TENT5C | 1p12 | 17 | 2.85 | 0 | 0.00 | 0.0675 | 0.302 | (A) BRAF Class 1 Colorectal |
| RARA | 17q21.2 | 27 | 4.53 | 1 | 0.99 | 0.0678 | 0.302 | (A) BRAF Class 1 Colorectal |
| MDM2 | 12q15 | 17 | 2.81 | 0 | 0.00 | 0.0685 | 0.304 | (A) BRAF Class 1 Colorectal |
| NOTCH2 | 1p12 | 46 | 7.37 | 3 | 2.97 | 0.0688 | 0.304 | (A) BRAF Class 1 Colorectal |
| PIK3CD | 1p36.22 | 28 | 8.70 | 1 | 1.96 | 0.0707 | 0.308 | (A) BRAF Class 1 Colorectal |

| | | | | | | | | |
|---|---|---|---|---|---|---|---|---|
| SETBP1 | 18q12.3 | 41 | 11.48 | 2 | 3.92 | 0.0709 | 0.308 | (A) BRAF Class 1 Colorectal |
| RHOA | 3p21.31 | 18 | 3.64 | 0 | 0.00 | 0.071 | 0.308 | (A) BRAF Class 1 Colorectal |
| FUBP1 | 1p31.1 | 16 | 4.52 | 0 | 0.00 | 0.0718 | 0.31 | (A) BRAF Class 1 Colorectal |
| CTLA4 | 2q33.2 | 18 | 4.00 | 0 | 0.00 | 0.0732 | 0.315 | (A) BRAF Class 1 Colorectal |
| RASA1 | 5q14.3 | 17 | 3.74 | 0 | 0.00 | 0.0735 | 0.315 | (A) BRAF Class 1 Colorectal |
| CDH1 | 16q22.1 | 37 | 4.65 | 2 | 1.56 | 0.0744 | 0.317 | (A) BRAF Class 1 Colorectal |
| PTPRT | 0q12-q13.1 | 57 | 17.43 | 5 | 8.93 | 0.075 | 0.317 | (A) BRAF Class 1 Colorectal |
| TERT | 5p15.33 | 47 | 7.74 | 3 | 3.19 | 0.0753 | 0.317 | (A) BRAF Class 1 Colorectal |
| PLK2 | 5q11.2 | 17 | 5.88 | 0 | 0.00 | 0.0759 | 0.318 | (A) BRAF Class 1 Colorectal |
| GRM3 | 21.11-q21. | 18 | 12.33 | 0 | 0.00 | 0.0772 | 0.319 | (A) BRAF Class 1 Colorectal |
| SUFU | 10q24.32 | 26 | 4.36 | 1 | 0.99 | 0.0774 | 0.319 | (A) BRAF Class 1 Colorectal |
| SLC25A13 | 7q21.3 | 2 | 1.25 | 2 | 8.70 | 0.0776 | 0.319 | (B) BRAF Class 3 Colorectal |
| XRCC4 | 5q14.2 | 2 | 1.25 | 2 | 8.70 | 0.0776 | 0.319 | (B) BRAF Class 3 Colorectal |
| ZNF217 | 20q13.2 | 28 | 7.20 | 1 | 1.67 | 0.0785 | 0.321 | (A) BRAF Class 1 Colorectal |
| STAT3 | 17q21.2 | 26 | 4.47 | 1 | 1.02 | 0.0788 | 0.321 | (A) BRAF Class 1 Colorectal |
| FGF4 | 11q13.3 | 15 | 4.17 | 0 | 0.00 | 0.0792 | 0.321 | (A) BRAF Class 1 Colorectal |
| SYNE1 | 6q25.2 | 3 | 2.42 | 2 | 14.29 | 0.0803 | 0.324 | (B) BRAF Class 3 Colorectal |
| AMER1 | Xq11.2 | 32 | 8.89 | 2 | 3.08 | 0.0811 | 0.324 | (A) BRAF Class 1 Colorectal |
| NTRK1 | 1q23.1 | 36 | 5.69 | 2 | 1.96 | 0.0813 | 0.324 | (A) BRAF Class 1 Colorectal |
| PARP1 | 1q42.12 | 25 | 7.14 | 1 | 1.67 | 0.0822 | 0.327 | (A) BRAF Class 1 Colorectal |
| PTPN11 | 12q24.13 | 17 | 2.09 | 0 | 0.00 | 0.0826 | 0.327 | (A) BRAF Class 1 Colorectal |
| CARD11 | 7p22.2 | 79 | 13.26 | 8 | 7.92 | 0.086 | 0.337 | (A) BRAF Class 1 Colorectal |
| FANCI | 15q26.1 | 20 | 6.64 | 0 | 0.00 | 0.0862 | 0.337 | (A) BRAF Class 1 Colorectal |
| PAK5 | 20p12.2 | 15 | 4.89 | 0 | 0.00 | 0.0871 | 0.339 | (A) BRAF Class 1 Colorectal |
| STAG1 | 3q22.3 | 21 | 13.29 | 0 | 0.00 | 0.0895 | 0.346 | (A) BRAF Class 1 Colorectal |
| PRKN | 6q26 | 34 | 5.77 | 2 | 2.04 | 0.0899 | 0.346 | (A) BRAF Class 1 Colorectal |
| SOX17 | 8q11.23 | 43 | 14.88 | 3 | 6.52 | 0.0905 | 0.346 | (A) BRAF Class 1 Colorectal |
| KMT2B | 19q13.12 | 66 | 26.83 | 5 | 14.71 | 0.0905 | 0.346 | (A) BRAF Class 1 Colorectal |
| POLQ | 3q13.33 | 25 | 15.43 | 1 | 3.85 | 0.091 | 0.346 | (A) BRAF Class 1 Colorectal |
| CYLD | 16q12.1 | 26 | 5.19 | 1 | 1.25 | 0.092 | 0.349 | (A) BRAF Class 1 Colorectal |
| GLI2 | 2q14.2 | 32 | 13.28 | 2 | 4.88 | 0.0954 | 0.36 | (A) BRAF Class 1 Colorectal |
| IRS2 | 13q34 | 38 | 10.56 | 3 | 4.62 | 0.0967 | 0.363 | (A) BRAF Class 1 Colorectal |
| JAK2 | 9p24.1 | 25 | 3.19 | 1 | 0.78 | 0.0973 | 0.364 | (A) BRAF Class 1 Colorectal |
| FH | 1q43 | 24 | 4.06 | 1 | 0.99 | 0.0979 | 0.364 | (A) BRAF Class 1 Colorectal |
| PPARG | 3p25.2 | 16 | 3.65 | 0 | 0.00 | 0.0988 | 0.366 | (A) BRAF Class 1 Colorectal |
| BARD1 | 2q35 | 24 | 4.62 | 1 | 1.14 | 0.1 | 0.369 | (A) BRAF Class 1 Colorectal |
| RAD21 | 8q24.11 | 25 | 4.50 | 1 | 1.11 | 0.101 | 0.369 | (A) BRAF Class 1 Colorectal |
| FOXA1 | 14q21.1 | 15 | 3.07 | 0 | 0.00 | 0.102 | 0.374 | (A) BRAF Class 1 Colorectal |
| ANKRD26 | 10p12.1 | 31 | 37.80 | 0 | 0.00 | 0.103 | 0.376 | (A) BRAF Class 1 Colorectal |
| EPHB1 | 3q22.2 | 30 | 8.33 | 2 | 3.08 | 0.104 | 0.377 | (A) BRAF Class 1 Colorectal |
| PRKCI | 3q26.2 | 27 | 5.36 | 1 | 1.35 | 0.104 | 0.377 | (A) BRAF Class 1 Colorectal |
| PDCD1 | 2q37.3 | 26 | 7.83 | 1 | 2.00 | 0.106 | 0.381 | (A) BRAF Class 1 Colorectal |
| FOXL2 | 3q22.3 | 32 | 6.05 | 2 | 2.25 | 0.108 | 0.385 | (A) BRAF Class 1 Colorectal |
| IGF2 | 11p15.5 | 14 | 2.75 | 0 | 0.00 | 0.109 | 0.39 | (A) BRAF Class 1 Colorectal |
| RSPO3 | 6q22.33 | 15 | 7.61 | 0 | 0.00 | 0.111 | 0.392 | (A) BRAF Class 1 Colorectal |
| IKZF1 | 7p12.2 | 47 | 7.89 | 4 | 3.96 | 0.111 | 0.392 | (A) BRAF Class 1 Colorectal |
| REL | 2p16.1 | 23 | 3.89 | 1 | 0.99 | 0.112 | 0.392 | (A) BRAF Class 1 Colorectal |
| FANCG | 9p13.3 | 15 | 3.86 | 0 | 0.00 | 0.112 | 0.392 | (A) BRAF Class 1 Colorectal |

| | | | | | | | | |
|---|---|---|---|---|---|---|---|---|
| MYC | 8q24.21 | 47 | 7.51 | 4 | 3.77 | 0.112 | 0.392 | (A) BRAF Class 1 Colorectal |
| RECQL | 12p12.1 | 13 | 10.24 | 0 | 0.00 | 0.114 | 0.394 | (A) BRAF Class 1 Colorectal |
| AKT3 | 1q43-q44 | 14 | 2.24 | 0 | 0.00 | 0.114 | 0.394 | (A) BRAF Class 1 Colorectal |
| PREX2 | 8q13.2 | 46 | 17.42 | 4 | 9.09 | 0.118 | 0.406 | (A) BRAF Class 1 Colorectal |
| HIST1H1C | 6p22.2 | 15 | 4.67 | 0 | 0.00 | 0.118 | 0.406 | (A) BRAF Class 1 Colorectal |
| DROSHA | 5p13.3 | 12 | 9.23 | 0 | 0.00 | 0.12 | 0.412 | (A) BRAF Class 1 Colorectal |
| SMAD3 | 15q22.33 | 22 | 6.41 | 7 | 11.67 | 0.121 | 0.413 | (B) BRAF Class 3 Colorectal |
| ADGRA2 | 8p11.23 | 20 | 18.02 | 1 | 5.00 | 0.125 | 0.418 | (A) BRAF Class 1 Colorectal |
| SLIT2 | 4p15.31 | 15 | 11.03 | 0 | 0.00 | 0.127 | 0.418 | (A) BRAF Class 1 Colorectal |
| CDH11 | 16q21 | 0 | 0.00 | 1 | 20.00 | 0.128 | 0.418 | (B) BRAF Class 3 Colorectal |
| HLF | 17q22 | 0 | 0.00 | 1 | 20.00 | 0.128 | 0.418 | (B) BRAF Class 3 Colorectal |
| HOOK3 | 8p11.21 | 0 | 0.00 | 1 | 20.00 | 0.128 | 0.418 | (B) BRAF Class 3 Colorectal |
| PBX1 | 1q23.3 | 0 | 0.00 | 1 | 20.00 | 0.128 | 0.418 | (B) BRAF Class 3 Colorectal |
| RBM15 | 1p13.3 | 0 | 0.00 | 1 | 20.00 | 0.128 | 0.418 | (B) BRAF Class 3 Colorectal |
| PBRM1 | 3p21.1 | 59 | 10.24 | 6 | 6.06 | 0.129 | 0.418 | (A) BRAF Class 1 Colorectal |
| CDC73 | 1q31.2 | 22 | 3.69 | 1 | 0.99 | 0.129 | 0.418 | (A) BRAF Class 1 Colorectal |
| KDM6B | 17p13.1 | 12 | 15.79 | 0 | 0.00 | 0.131 | 0.418 | (A) BRAF Class 1 Colorectal |
| FN1 | 2q35 | 0 | 0.00 | 1 | 20.00 | 0.132 | 0.418 | (B) BRAF Class 3 Colorectal |
| G6PD | Xq28 | 0 | 0.00 | 1 | 20.00 | 0.132 | 0.418 | (B) BRAF Class 3 Colorectal |
| GRM8 | 7q31.33 | 0 | 0.00 | 1 | 20.00 | 0.132 | 0.418 | (B) BRAF Class 3 Colorectal |
| KLF6 | 10p15.2 | 0 | 0.00 | 1 | 20.00 | 0.132 | 0.418 | (B) BRAF Class 3 Colorectal |
| MAGI1 | 3p14.1 | 0 | 0.00 | 1 | 20.00 | 0.132 | 0.418 | (B) BRAF Class 3 Colorectal |
| MMP2 | 16q12.2 | 0 | 0.00 | 1 | 20.00 | 0.132 | 0.418 | (B) BRAF Class 3 Colorectal |
| USP9X | Xp11.4 | 0 | 0.00 | 1 | 20.00 | 0.132 | 0.418 | (B) BRAF Class 3 Colorectal |
| BCL6 | 3q27.3 | 30 | 5.03 | 2 | 1.98 | 0.132 | 0.418 | (A) BRAF Class 1 Colorectal |
| FBXW7 | 4q31.3 | 139 | 16.69 | 16 | 12.40 | 0.134 | 0.423 | (A) BRAF Class 1 Colorectal |
| RXRA | 9q34.2 | 12 | 9.45 | 0 | 0.00 | 0.136 | 0.423 | (A) BRAF Class 1 Colorectal |
| UPF1 | 19p13.11 | 12 | 9.45 | 0 | 0.00 | 0.136 | 0.423 | (A) BRAF Class 1 Colorectal |
| CSF1R | 5q32 | 22 | 3.52 | 1 | 0.96 | 0.136 | 0.423 | (A) BRAF Class 1 Colorectal |
| TP63 | 3q28 | 14 | 4.36 | 0 | 0.00 | 0.137 | 0.423 | (A) BRAF Class 1 Colorectal |
| FNDC3B | 3q26.31 | 0 | 0.00 | 1 | 0.75 | 0.138 | 0.423 | (B) BRAF Class 3 Colorectal |
| MIR99AHG | 21q21.1 | 0 | 0.00 | 1 | 0.75 | 0.138 | 0.423 | (B) BRAF Class 3 Colorectal |
| TMIE | 3p21.31 | 0 | 0.00 | 1 | 0.75 | 0.138 | 0.423 | (B) BRAF Class 3 Colorectal |
| ZNF284 | 19q13.31 | 0 | 0.00 | 1 | 0.75 | 0.138 | 0.423 | (B) BRAF Class 3 Colorectal |
| IFNGR1 | 6q23.3 | 13 | 4.50 | 0 | 0.00 | 0.141 | 0.426 | (A) BRAF Class 1 Colorectal |
| RPS6KB2 | 11q13.2 | 13 | 4.50 | 0 | 0.00 | 0.141 | 0.426 | (A) BRAF Class 1 Colorectal |
| SHQ1 | 3p13 | 13 | 4.50 | 0 | 0.00 | 0.141 | 0.426 | (A) BRAF Class 1 Colorectal |
| TGFBR1 | 9q22.33 | 13 | 4.50 | 0 | 0.00 | 0.141 | 0.426 | (A) BRAF Class 1 Colorectal |
| GNAQ | 9q21.2 | 13 | 1.58 | 0 | 0.00 | 0.141 | 0.426 | (A) BRAF Class 1 Colorectal |
| ARHGEF12 | 11q23.3 | 14 | 8.70 | 0 | 0.00 | 0.143 | 0.429 | (A) BRAF Class 1 Colorectal |
| STAT5A | 17q21.2 | 14 | 4.62 | 0 | 0.00 | 0.144 | 0.431 | (A) BRAF Class 1 Colorectal |
| INHA | 2q35 | 13 | 4.80 | 0 | 0.00 | 0.148 | 0.438 | (A) BRAF Class 1 Colorectal |
| MAPK3 | 16p11.2 | 13 | 4.80 | 0 | 0.00 | 0.148 | 0.438 | (A) BRAF Class 1 Colorectal |
| BAP1 | 3p21.1 | 21 | 3.47 | 1 | 0.98 | 0.149 | 0.438 | (A) BRAF Class 1 Colorectal |
| CHEK1 | 11q24.2 | 21 | 3.97 | 1 | 1.12 | 0.149 | 0.438 | (A) BRAF Class 1 Colorectal |
| CCND3 | 6p21.1 | 12 | 1.98 | 0 | 0.00 | 0.152 | 0.445 | (A) BRAF Class 1 Colorectal |
| PRF1 | 10q22.1 | 13 | 5.49 | 0 | 0.00 | 0.152 | 0.445 | (A) BRAF Class 1 Colorectal |
| FLT3 | 13q12.2 | 46 | 5.71 | 4 | 3.10 | 0.154 | 0.45 | (A) BRAF Class 1 Colorectal |

| | | | | | | | | |
|---|---|---|---|---|---|---|---|---|
| CRKL | 22q11.21 | 3 | 0.50 | 2 | 1.98 | 0.155 | 0.45 | (B) BRAF Class 3 Colorectal |
| DAXX | 6p21.32 | 21 | 4.05 | 1 | 1.18 | 0.158 | 0.457 | (A) BRAF Class 1 Colorectal |
| RICTOR | 5p13.1 | 36 | 6.81 | 3 | 3.37 | 0.159 | 0.458 | (A) BRAF Class 1 Colorectal |
| ABRAXAS1 | 4q21.23 | 12 | 2.42 | 0 | 0.00 | 0.159 | 0.458 | (A) BRAF Class 1 Colorectal |
| MYCN | 2p24.3 | 20 | 3.31 | 1 | 0.96 | 0.161 | 0.459 | (A) BRAF Class 1 Colorectal |
| COP1 | q25.1-q25.[illegible] | 21 | 5.75 | 1 | 1.69 | 0.161 | 0.459 | (A) BRAF Class 1 Colorectal |
| EPAS1 | 2p21 | 11 | 8.66 | 0 | 0.00 | 0.161 | 0.459 | (A) BRAF Class 1 Colorectal |
| TMPRSS2 | 21q22.3 | 20 | 3.88 | 1 | 1.14 | 0.163 | 0.461 | (A) BRAF Class 1 Colorectal |
| BRIP1 | 17q23.2 | 28 | 4.70 | 2 | 1.98 | 0.164 | 0.461 | (A) BRAF Class 1 Colorectal |
| BCL11B | 14q32.2 | 13 | 6.70 | 0 | 0.00 | 0.164 | 0.461 | (A) BRAF Class 1 Colorectal |
| CALR | 19p13.13 | 12 | 2.50 | 0 | 0.00 | 0.165 | 0.461 | (A) BRAF Class 1 Colorectal |
| TP53BP1 | 15q15.3 | 27 | 9.00 | 2 | 3.85 | 0.165 | 0.461 | (A) BRAF Class 1 Colorectal |
| PRKAR1A | 17q24.2 | 7 | 1.17 | 3 | 2.97 | 0.165 | 0.461 | (B) BRAF Class 3 Colorectal |
| TRAF2 | 9q34.3 | 13 | 4.29 | 0 | 0.00 | 0.166 | 0.461 | (A) BRAF Class 1 Colorectal |
| PIK3CA | 3q26.32 | 198 | 23.63 | 26 | 19.40 | 0.167 | 0.462 | (A) BRAF Class 1 Colorectal |
| RAD54L | 1p34.1 | 11 | 3.47 | 0 | 0.00 | 0.168 | 0.463 | (A) BRAF Class 1 Colorectal |
| MTAP | 9p21.3 | 4 | 2.34 | 2 | 8.00 | 0.17 | 0.468 | (B) BRAF Class 3 Colorectal |
| SOS1 | 2p22.1 | 27 | 9.25 | 2 | 4.00 | 0.17 | 0.468 | (A) BRAF Class 1 Colorectal |
| XRCC3 | 14q32.33 | 11 | 6.18 | 0 | 0.00 | 0.172 | 0.471 | (A) BRAF Class 1 Colorectal |
| BCL10 | 1p22.3 | 12 | 3.95 | 0 | 0.00 | 0.173 | 0.471 | (A) BRAF Class 1 Colorectal |
| AKT2 | 19q13.2 | 20 | 3.19 | 1 | 0.96 | 0.174 | 0.472 | (A) BRAF Class 1 Colorectal |
| PALB2 | 16p12.2 | 49 | 8.10 | 5 | 4.90 | 0.179 | 0.486 | (A) BRAF Class 1 Colorectal |
| SMAD4 | 18q21.2 | 188 | 22.57 | 24 | 18.60 | 0.186 | 0.501 | (A) BRAF Class 1 Colorectal |
| IKBKE | 1q32.1 | 10 | 2.78 | 0 | 0.00 | 0.187 | 0.501 | (A) BRAF Class 1 Colorectal |
| RBM10 | Xp11.3 | 42 | 8.19 | 4 | 4.71 | 0.187 | 0.501 | (A) BRAF Class 1 Colorectal |
| FANCC | 9q22.32 | 19 | 3.19 | 1 | 0.99 | 0.188 | 0.501 | (A) BRAF Class 1 Colorectal |
| FGF3 | 11q13.3 | 10 | 2.71 | 0 | 0.00 | 0.189 | 0.501 | (A) BRAF Class 1 Colorectal |
| PML | 15q24.1 | 12 | 6.19 | 0 | 0.00 | 0.19 | 0.501 | (A) BRAF Class 1 Colorectal |
| TSHR | 14q31.1 | 28 | 5.20 | 2 | 2.33 | 0.191 | 0.501 | (A) BRAF Class 1 Colorectal |
| CBFA2T3 | 16q24.3 | 12 | 7.45 | 0 | 0.00 | 0.191 | 0.501 | (A) BRAF Class 1 Colorectal |
| TLX3 | 5q35.1 | 12 | 7.45 | 0 | 0.00 | 0.191 | 0.501 | (A) BRAF Class 1 Colorectal |
| TFE3 | Xp11.23 | 23 | 8.19 | 1 | 2.63 | 0.191 | 0.501 | (A) BRAF Class 1 Colorectal |
| TSC1 | 9q34.13 | 34 | 5.62 | 3 | 2.94 | 0.191 | 0.501 | (A) BRAF Class 1 Colorectal |
| DNAJB1 | 19p13.12 | 11 | 3.57 | 0 | 0.00 | 0.192 | 0.501 | (A) BRAF Class 1 Colorectal |
| GRIN2A | 16p13.2 | 51 | 14.57 | 6 | 9.52 | 0.194 | 0.504 | (A) BRAF Class 1 Colorectal |
| BABAM1 | 19p13.11 | 11 | 3.83 | 0 | 0.00 | 0.196 | 0.508 | (A) BRAF Class 1 Colorectal |
| ERCC6 | 10q11.23 | 21 | 13.13 | 1 | 4.35 | 0.198 | 0.512 | (A) BRAF Class 1 Colorectal |
| KLF4 | 9q31.2 | 11 | 2.26 | 0 | 0.00 | 0.2 | 0.513 | (A) BRAF Class 1 Colorectal |
| KAT6B | 10q22.2 | 21 | 10.82 | 1 | 3.57 | 0.2 | 0.513 | (A) BRAF Class 1 Colorectal |
| PNRC1 | 6q15 | 10 | 2.61 | 0 | 0.00 | 0.2 | 0.513 | (A) BRAF Class 1 Colorectal |
| NFKBIZ | 3q12.3 | 9 | 3.81 | 3 | 8.33 | 0.202 | 0.515 | (B) BRAF Class 3 Colorectal |
| TEK | 9p21.2 | 8 | 5.23 | 0 | 0.00 | 0.203 | 0.515 | (A) BRAF Class 1 Colorectal |
| CREB1 | 2q33.3 | 0 | 0.00 | 1 | 10.00 | 0.204 | 0.516 | (B) BRAF Class 3 Colorectal |
| MAML2 | 11q21 | 0 | 0.00 | 1 | 10.00 | 0.204 | 0.516 | (B) BRAF Class 3 Colorectal |
| RAD51C | 17q22 | 10 | 1.91 | 0 | 0.00 | 0.206 | 0.518 | (A) BRAF Class 1 Colorectal |
| AURKB | 17p13.1 | 10 | 1.68 | 0 | 0.00 | 0.207 | 0.519 | (A) BRAF Class 1 Colorectal |
| CHD4 | 12p13.31 | 21 | 15.44 | 1 | 5.26 | 0.207 | 0.519 | (A) BRAF Class 1 Colorectal |
| HGF | 7q21.11 | 17 | 4.72 | 1 | 1.54 | 0.208 | 0.519 | (A) BRAF Class 1 Colorectal |

| | | | | | | | | |
|---|---|---|---|---|---|---|---|---|
| NFE2L2 | 2q31.2 | 10 | 1.65 | 0 | 0.00 | 0.208 | 0.519 | (A) BRAF Class 1 Colorectal |
| FANCA | 16q24.3 | 33 | 5.45 | 3 | 2.94 | 0.21 | 0.521 | (A) BRAF Class 1 Colorectal |
| CUL3 | 2q36.2 | 9 | 2.54 | 0 | 0.00 | 0.216 | 0.533 | (A) BRAF Class 1 Colorectal |
| SESN3 | 11q21 | 4 | 3.15 | 2 | 9.09 | 0.216 | 0.533 | (B) BRAF Class 3 Colorectal |
| ABL2 | 1q25.2 | 10 | 7.25 | 0 | 0.00 | 0.217 | 0.535 | (A) BRAF Class 1 Colorectal |
| PTPN14 | 1q32.3-q41 | 11 | 6.88 | 0 | 0.00 | 0.218 | 0.535 | (A) BRAF Class 1 Colorectal |
| NPM1 | 5q35.1 | 10 | 1.29 | 0 | 0.00 | 0.219 | 0.536 | (A) BRAF Class 1 Colorectal |
| TNFRSF14 | 1p36.32 | 9 | 2.50 | 0 | 0.00 | 0.221 | 0.536 | (A) BRAF Class 1 Colorectal |
| PIK3R1 | 5q13.1 | 40 | 6.39 | 4 | 3.85 | 0.221 | 0.536 | (A) BRAF Class 1 Colorectal |
| SRSF2 | 17q25.1 | 10 | 1.83 | 0 | 0.00 | 0.222 | 0.536 | (A) BRAF Class 1 Colorectal |
| ETV4 | 17q21.31 | 11 | 2.89 | 0 | 0.00 | 0.222 | 0.536 | (A) BRAF Class 1 Colorectal |
| ROBO1 | 3p12.3 | 2 | 40.00 | 0 | 0.00 | 0.222 | 0.536 | (A) BRAF Class 1 Colorectal |
| TAP1 | 6p21.32 | 9 | 7.09 | 0 | 0.00 | 0.227 | 0.546 | (A) BRAF Class 1 Colorectal |
| STAT4 | q32.2-q32.: | 10 | 6.99 | 0 | 0.00 | 0.229 | 0.548 | (A) BRAF Class 1 Colorectal |
| XPO1 | 2p15 | 32 | 5.29 | 3 | 2.94 | 0.229 | 0.548 | (A) BRAF Class 1 Colorectal |
| MYD88 | 3p22.2 | 17 | 2.78 | 1 | 0.95 | 0.233 | 0.549 | (A) BRAF Class 1 Colorectal |
| RELA | 11q13.1 | 5 | 2.54 | 2 | 6.67 | 0.233 | 0.549 | (B) BRAF Class 3 Colorectal |
| ERRFI1 | 1p36.23 | 9 | 2.69 | 0 | 0.00 | 0.234 | 0.549 | (A) BRAF Class 1 Colorectal |
| ESR1 | ·q25.1-q25.: | 31 | 4.91 | 3 | 2.75 | 0.235 | 0.549 | (A) BRAF Class 1 Colorectal |
| PRKD1 | 14q12 | 10 | 6.29 | 0 | 0.00 | 0.236 | 0.549 | (A) BRAF Class 1 Colorectal |
| ACVR1 | 2q24.1 | 10 | 2.14 | 0 | 0.00 | 0.236 | 0.549 | (A) BRAF Class 1 Colorectal |
| CDK4 | 12q14.1 | 9 | 1.49 | 0 | 0.00 | 0.238 | 0.549 | (A) BRAF Class 1 Colorectal |
| FOXO1 | 13q14.11 | 17 | 5.48 | 1 | 1.92 | 0.239 | 0.549 | (A) BRAF Class 1 Colorectal |
| PIK3CB | 3q22.3 | 24 | 6.70 | 2 | 3.28 | 0.24 | 0.549 | (A) BRAF Class 1 Colorectal |
| FANCD2 | 3p25.3 | 20 | 5.22 | 1 | 1.85 | 0.241 | 0.549 | (A) BRAF Class 1 Colorectal |
| SF3B1 | 2q33.1 | 25 | 3.90 | 2 | 1.90 | 0.242 | 0.549 | (A) BRAF Class 1 Colorectal |
| MAP3K4 | 6q26 | 21 | 25.61 | 0 | 0.00 | 0.242 | 0.549 | (A) BRAF Class 1 Colorectal |
| CBFB | 16q22.1 | 9 | 1.73 | 0 | 0.00 | 0.242 | 0.549 | (A) BRAF Class 1 Colorectal |
| AFDN | 6q27 | 1 | 2.94 | 1 | 20.00 | 0.243 | 0.549 | (B) BRAF Class 3 Colorectal |
| AFF1 | ·q21.3-q22. | 1 | 2.94 | 1 | 20.00 | 0.243 | 0.549 | (B) BRAF Class 3 Colorectal |
| BCL9 | 1q21.2 | 1 | 2.94 | 1 | 20.00 | 0.243 | 0.549 | (B) BRAF Class 3 Colorectal |
| NIN | 14q22.1 | 1 | 2.94 | 1 | 20.00 | 0.243 | 0.549 | (B) BRAF Class 3 Colorectal |
| NUP214 | 9q34.13 | 1 | 2.94 | 1 | 20.00 | 0.243 | 0.549 | (B) BRAF Class 3 Colorectal |
| RNF213 | 17q25.3 | 1 | 2.94 | 1 | 20.00 | 0.243 | 0.549 | (B) BRAF Class 3 Colorectal |
| UBR5 | 8q22.3 | 1 | 2.94 | 1 | 20.00 | 0.243 | 0.549 | (B) BRAF Class 3 Colorectal |
| MDM4 | 1q32.1 | 9 | 1.49 | 0 | 0.00 | 0.244 | 0.55 | (A) BRAF Class 1 Colorectal |
| ACVR2A | q22.3-q23. | 1 | 3.03 | 1 | 20.00 | 0.249 | 0.551 | (B) BRAF Class 3 Colorectal |
| ADAMTS20 | 12q12 | 1 | 3.03 | 1 | 20.00 | 0.249 | 0.551 | (B) BRAF Class 3 Colorectal |
| CSMD3 | 8q23.3 | 1 | 3.03 | 1 | 20.00 | 0.249 | 0.551 | (B) BRAF Class 3 Colorectal |
| FOXP4 | 6p21.1 | 1 | 3.03 | 1 | 20.00 | 0.249 | 0.551 | (B) BRAF Class 3 Colorectal |
| TAF1L | 9p21.1 | 1 | 3.03 | 1 | 20.00 | 0.249 | 0.551 | (B) BRAF Class 3 Colorectal |
| THBS1 | 15q14 | 1 | 3.03 | 1 | 20.00 | 0.249 | 0.551 | (B) BRAF Class 3 Colorectal |
| PIK3C2B | 1q32.1 | 51 | 13.49 | 5 | 9.09 | 0.251 | 0.551 | (A) BRAF Class 1 Colorectal |
| MAGI2 | 7q21.11 | 13 | 9.92 | 0 | 0.00 | 0.251 | 0.551 | (A) BRAF Class 1 Colorectal |
| SERPINA1 | 14q32.13 | 10 | 6.25 | 0 | 0.00 | 0.252 | 0.551 | (A) BRAF Class 1 Colorectal |
| TOPBP1 | 3q22.1 | 10 | 6.25 | 0 | 0.00 | 0.252 | 0.551 | (A) BRAF Class 1 Colorectal |
| DOCK8 | 9p24.3 | 27 | 16.88 | 2 | 8.70 | 0.252 | 0.551 | (A) BRAF Class 1 Colorectal |
| PRSS1 | 7q34 | 1 | 0.49 | 1 | 3.13 | 0.252 | 0.551 | (B) BRAF Class 3 Colorectal |

| | | | | | | | | |
|---|---|---|---|---|---|---|---|---|
| RAD52 | 12p13.33 | 17 | 3.56 | 1 | 1.28 | 0.253 | 0.552 | (A) BRAF Class 1 Colorectal |
| FAS | 10q23.31 | 10 | 2.64 | 0 | 0.00 | 0.254 | 0.552 | (A) BRAF Class 1 Colorectal |
| ETV5 | 3q27.2 | 10 | 2.63 | 0 | 0.00 | 0.255 | 0.552 | (A) BRAF Class 1 Colorectal |
| GALNT12 | 9q22.33 | 8 | 4.55 | 0 | 0.00 | 0.256 | 0.554 | (A) BRAF Class 1 Colorectal |
| GEN1 | 2p24.2 | 11 | 3.83 | 0 | 0.00 | 0.258 | 0.555 | (A) BRAF Class 1 Colorectal |
| IGF2R | 6q25.3 | 2 | 3.92 | 2 | 11.76 | 0.258 | 0.555 | (B) BRAF Class 3 Colorectal |
| ID3 | 1p36.12 | 1 | 0.18 | 1 | 1.11 | 0.26 | 0.556 | (B) BRAF Class 3 Colorectal |
| LYN | 8q12.1 | 10 | 3.73 | 0 | 0.00 | 0.26 | 0.556 | (A) BRAF Class 1 Colorectal |
| GSK3B | 3q13.33 | 8 | 2.22 | 0 | 0.00 | 0.262 | 0.559 | (A) BRAF Class 1 Colorectal |
| LMO1 | 11p15.4 | 9 | 1.55 | 0 | 0.00 | 0.267 | 0.568 | (A) BRAF Class 1 Colorectal |
| HOXB13 | 17q21.32 | 8 | 1.78 | 0 | 0.00 | 0.268 | 0.568 | (A) BRAF Class 1 Colorectal |
| RIT1 | 1q22 | 9 | 1.83 | 0 | 0.00 | 0.268 | 0.568 | (A) BRAF Class 1 Colorectal |
| PRDM14 | 8q13.3 | 8 | 6.30 | 0 | 0.00 | 0.269 | 0.568 | (A) BRAF Class 1 Colorectal |
| MYCL | 1p34.2 | 16 | 2.64 | 1 | 0.98 | 0.27 | 0.568 | (A) BRAF Class 1 Colorectal |
| FAT3 | 11q14.3 | 3 | 13.04 | 0 | 0.00 | 0.271 | 0.568 | (A) BRAF Class 1 Colorectal |
| STAG2 | Xq25 | 23 | 3.86 | 2 | 1.98 | 0.272 | 0.569 | (A) BRAF Class 1 Colorectal |
| SDHAF2 | 11q12.2 | 9 | 1.60 | 0 | 0.00 | 0.275 | 0.572 | (A) BRAF Class 1 Colorectal |
| BCL2L1 | 20q11.21 | 5 | 0.85 | 2 | 1.98 | 0.275 | 0.572 | (B) BRAF Class 3 Colorectal |
| XPC | 3p25.1 | 9 | 3.33 | 0 | 0.00 | 0.275 | 0.572 | (A) BRAF Class 1 Colorectal |
| MLH1 | 3p22.2 | 24 | 2.98 | 2 | 1.55 | 0.28 | 0.58 | (A) BRAF Class 1 Colorectal |
| PRPF40B | 12q13.12 | 7 | 8.86 | 0 | 0.00 | 0.283 | 0.586 | (A) BRAF Class 1 Colorectal |
| RAD51 | 15q15.1 | 8 | 1.52 | 0 | 0.00 | 0.285 | 0.589 | (A) BRAF Class 1 Colorectal |
| CHD2 | 15q26.1 | 18 | 13.24 | 1 | 5.26 | 0.286 | 0.589 | (A) BRAF Class 1 Colorectal |
| PIK3R2 | 19p13.11 | 21 | 6.00 | 2 | 3.17 | 0.289 | 0.59 | (A) BRAF Class 1 Colorectal |
| GPS2 | 17p13.1 | 8 | 2.94 | 0 | 0.00 | 0.289 | 0.59 | (A) BRAF Class 1 Colorectal |
| SPTA1 | 1q23.1 | 26 | 19.12 | 2 | 10.53 | 0.29 | 0.59 | (A) BRAF Class 1 Colorectal |
| FAN1 | 15q13.3 | 9 | 5.63 | 0 | 0.00 | 0.29 | 0.59 | (A) BRAF Class 1 Colorectal |
| KIF1B | 1p36.22 | 9 | 5.63 | 0 | 0.00 | 0.29 | 0.59 | (A) BRAF Class 1 Colorectal |
| TCF7L2 | )q25.2-q25. | 58 | 10.62 | 7 | 7.95 | 0.291 | 0.59 | (A) BRAF Class 1 Colorectal |
| CTCF | 16q22.1 | 29 | 5.58 | 3 | 3.41 | 0.293 | 0.593 | (A) BRAF Class 1 Colorectal |
| XBP1 | 2q12.1\|22q | 1 | 2.70 | 1 | 14.29 | 0.296 | 0.598 | (B) BRAF Class 3 Colorectal |
| PAX8 | 2q14.1 | 2 | 1.65 | 1 | 6.67 | 0.298 | 0.6 | (B) BRAF Class 3 Colorectal |
| SAMD9 | 7q21.2 | 1 | 2.78 | 1 | 14.29 | 0.302 | 0.604 | (B) BRAF Class 3 Colorectal |
| STK40 | 1p34.3 | 8 | 2.77 | 0 | 0.00 | 0.303 | 0.604 | (A) BRAF Class 1 Colorectal |
| XIAP | Xq25 | 8 | 2.77 | 0 | 0.00 | 0.303 | 0.604 | (A) BRAF Class 1 Colorectal |
| NPRL3 | 16p13.3 | 1 | 1.79 | 1 | 9.09 | 0.303 | 0.604 | (B) BRAF Class 3 Colorectal |
| ORCS8-MEF2 | 19p13.11 | 8 | 0.95 | 0 | 0.00 | 0.304 | 0.604 | (A) BRAF Class 1 Colorectal |
| EIF1AX | Xp22.12 | 8 | 2.48 | 0 | 0.00 | 0.305 | 0.604 | (A) BRAF Class 1 Colorectal |
| RBBP8 | 18q11.2 | 17 | 10.63 | 1 | 4.35 | 0.305 | 0.604 | (A) BRAF Class 1 Colorectal |
| PAX5 | 9p13.2 | 28 | 4.64 | 3 | 2.88 | 0.307 | 0.607 | (A) BRAF Class 1 Colorectal |
| ELANE | 19p13.3 | 8 | 4.91 | 0 | 0.00 | 0.312 | 0.615 | (A) BRAF Class 1 Colorectal |
| EIF4A2 | 3q27.3 | 8 | 2.94 | 0 | 0.00 | 0.313 | 0.615 | (A) BRAF Class 1 Colorectal |
| FGF19 | 11q13.3 | 7 | 1.90 | 0 | 0.00 | 0.313 | 0.615 | (A) BRAF Class 1 Colorectal |
| DDB2 | 11p11.2 | 8 | 2.96 | 0 | 0.00 | 0.318 | 0.621 | (A) BRAF Class 1 Colorectal |
| EXT2 | 11p11.2 | 8 | 2.96 | 0 | 0.00 | 0.318 | 0.621 | (A) BRAF Class 1 Colorectal |
| ERF | 19q13.2 | 7 | 5.51 | 0 | 0.00 | 0.319 | 0.621 | (A) BRAF Class 1 Colorectal |
| SPRED1 | 15q14 | 6 | 4.55 | 0 | 0.00 | 0.321 | 0.624 | (A) BRAF Class 1 Colorectal |
| FANCL | 2p16.1 | 8 | 2.56 | 0 | 0.00 | 0.323 | 0.626 | (A) BRAF Class 1 Colorectal |

| | | | | | | | | |
|---|---|---|---|---|---|---|---|---|
| DEPDC5 | 2q12.2-q12. | 6 | 10.71 | 0 | 0.00 | 0.325 | 0.629 | (A) BRAF Class 1 Colorectal |
| FANCM | 14q21.2 | 23 | 10.95 | 2 | 6.25 | 0.326 | 0.629 | (A) BRAF Class 1 Colorectal |
| INPP4A | 2q11.2 | 15 | 5.19 | 1 | 2.17 | 0.327 | 0.629 | (A) BRAF Class 1 Colorectal |
| PRSS8 | 16p11.2 | 3 | 2.21 | 1 | 7.14 | 0.327 | 0.629 | (B) BRAF Class 3 Colorectal |
| MITF | 3p13 | 21 | 3.52 | 2 | 1.98 | 0.328 | 0.629 | (A) BRAF Class 1 Colorectal |
| NSD3 | 8p11.23 | 15 | 5.03 | 1 | 2.13 | 0.332 | 0.634 | (A) BRAF Class 1 Colorectal |
| EGR1 | 5q31.2 | 1 | 33.33 | 0 | 0.00 | 0.333 | 0.634 | (A) BRAF Class 1 Colorectal |
| BABAM2 | 2p23.2 | 8 | 5.00 | 0 | 0.00 | 0.334 | 0.634 | (A) BRAF Class 1 Colorectal |
| MCM8 | 20p12.3 | 8 | 5.00 | 0 | 0.00 | 0.334 | 0.634 | (A) BRAF Class 1 Colorectal |
| GATA4 | 8p23.1 | 17 | 4.51 | 1 | 1.92 | 0.334 | 0.634 | (A) BRAF Class 1 Colorectal |
| PAX7 | 1p36.13 | 9 | 7.44 | 0 | 0.00 | 0.338 | 0.636 | (A) BRAF Class 1 Colorectal |
| DDB1 | 11q12.2 | 16 | 10.00 | 1 | 4.35 | 0.338 | 0.636 | (A) BRAF Class 1 Colorectal |
| CDKN1A | 6p21.2 | 7 | 1.19 | 0 | 0.00 | 0.339 | 0.636 | (A) BRAF Class 1 Colorectal |
| SDHC | 1q23.3 | 7 | 1.19 | 0 | 0.00 | 0.339 | 0.636 | (A) BRAF Class 1 Colorectal |
| EWSR1 | 22q12.2 | 15 | 3.90 | 1 | 1.67 | 0.339 | 0.636 | (A) BRAF Class 1 Colorectal |
| COL7A1 | 3p21.31 | 31 | 19.38 | 3 | 13.04 | 0.343 | 0.642 | (A) BRAF Class 1 Colorectal |
| ALOX12B | 17p13.1 | 20 | 5.09 | 2 | 2.94 | 0.344 | 0.642 | (A) BRAF Class 1 Colorectal |
| NF2 | 22q12.2 | 14 | 2.24 | 1 | 0.96 | 0.345 | 0.642 | (A) BRAF Class 1 Colorectal |
| NTHL1 | 16p13.3 | 7 | 2.41 | 0 | 0.00 | 0.346 | 0.642 | (A) BRAF Class 1 Colorectal |
| XRCC2 | 7q36.1 | 7 | 1.44 | 0 | 0.00 | 0.347 | 0.643 | (A) BRAF Class 1 Colorectal |
| EGFL7 | 9q34.3 | 7 | 2.42 | 0 | 0.00 | 0.352 | 0.649 | (A) BRAF Class 1 Colorectal |
| PGBD3 | 10q11.23 | 7 | 0.84 | 0 | 0.00 | 0.353 | 0.649 | (A) BRAF Class 1 Colorectal |
| EMSY | 11q13.5 | 7 | 4.58 | 0 | 0.00 | 0.354 | 0.649 | (A) BRAF Class 1 Colorectal |
| FGF14 | 13q33.1 | 7 | 4.58 | 0 | 0.00 | 0.354 | 0.649 | (A) BRAF Class 1 Colorectal |
| YAP1 | 11q22.1 | 14 | 3.08 | 1 | 1.35 | 0.354 | 0.649 | (A) BRAF Class 1 Colorectal |
| SRC | 20q11.23 | 12 | 1.86 | 3 | 2.83 | 0.357 | 0.652 | (B) BRAF Class 3 Colorectal |
| CDKN1C | 11p15.4 | 2 | 0.84 | 1 | 2.63 | 0.359 | 0.652 | (B) BRAF Class 3 Colorectal |
| MAP2K1 | 15q22.31 | 12 | 1.82 | 3 | 2.75 | 0.36 | 0.652 | (B) BRAF Class 3 Colorectal |
| MECOM | 3q26.2 | 27 | 11.34 | 3 | 7.69 | 0.361 | 0.652 | (A) BRAF Class 1 Colorectal |
| DNMT3B | 20q11.21 | 21 | 7.27 | 2 | 4.35 | 0.362 | 0.652 | (A) BRAF Class 1 Colorectal |
| VHL | 3p25.3 | 14 | 1.76 | 1 | 0.78 | 0.362 | 0.652 | (A) BRAF Class 1 Colorectal |
| BCR | 22q11.23 | 8 | 5.56 | 0 | 0.00 | 0.362 | 0.652 | (A) BRAF Class 1 Colorectal |
| CXCR4 | 2q22.1 | 7 | 1.46 | 0 | 0.00 | 0.363 | 0.652 | (A) BRAF Class 1 Colorectal |
| MALT1 | 18q21.32 | 7 | 2.30 | 0 | 0.00 | 0.363 | 0.652 | (A) BRAF Class 1 Colorectal |
| LATS1 | 6q25.1 | 14 | 4.84 | 1 | 2.17 | 0.364 | 0.652 | (A) BRAF Class 1 Colorectal |
| EXO1 | 1q43 | 8 | 5.00 | 2 | 8.70 | 0.365 | 0.653 | (B) BRAF Class 3 Colorectal |
| PPP6C | 9q33.3 | 6 | 2.17 | 0 | 0.00 | 0.366 | 0.654 | (A) BRAF Class 1 Colorectal |
| ERBB3 | 12q13.2 | 39 | 6.23 | 8 | 7.55 | 0.368 | 0.656 | (B) BRAF Class 3 Colorectal |
| SMAD2 | 18q21.1 | 28 | 4.70 | 6 | 5.94 | 0.369 | 0.656 | (B) BRAF Class 3 Colorectal |
| MAP3K9 | 14q24.2 | 2 | 15.38 | 0 | 0.00 | 0.371 | 0.659 | (A) BRAF Class 1 Colorectal |
| NEIL3 | 4q34.3 | 15 | 9.38 | 1 | 4.35 | 0.373 | 0.661 | (A) BRAF Class 1 Colorectal |
| CDH20 | 18q21.33 | 1 | 2.17 | 1 | 8.33 | 0.374 | 0.661 | (B) BRAF Class 3 Colorectal |
| PTK2 | 8q24.3 | 6 | 7.89 | 0 | 0.00 | 0.376 | 0.663 | (A) BRAF Class 1 Colorectal |
| CCND2 | 12p13.32 | 13 | 2.15 | 1 | 0.98 | 0.377 | 0.663 | (A) BRAF Class 1 Colorectal |
| SMARCD1 | 12q13.12 | 19 | 6.19 | 2 | 3.77 | 0.377 | 0.663 | (A) BRAF Class 1 Colorectal |
| PAK1 | 1q13.5-q14. | 6 | 2.04 | 0 | 0.00 | 0.38 | 0.664 | (A) BRAF Class 1 Colorectal |
| ZNF708 | 19p12 | 2 | 2.63 | 1 | 7.69 | 0.381 | 0.664 | (B) BRAF Class 3 Colorectal |
| PAX3 | 2q36.1 | 8 | 6.61 | 0 | 0.00 | 0.382 | 0.664 | (A) BRAF Class 1 Colorectal |

| | | | | | | | | |
|---|---|---|---|---|---|---|---|---|
| DMC1 | 22q13.1 | 7 | 4.38 | 0 | 0.00 | 0.384 | 0.664 | (A) BRAF Class 1 Colorectal |
| HELQ | 4q21.23 | 7 | 4.38 | 0 | 0.00 | 0.384 | 0.664 | (A) BRAF Class 1 Colorectal |
| MBD4 | 3q21.3 | 7 | 4.38 | 0 | 0.00 | 0.384 | 0.664 | (A) BRAF Class 1 Colorectal |
| USP28 | 11q23.2 | 7 | 4.38 | 0 | 0.00 | 0.384 | 0.664 | (A) BRAF Class 1 Colorectal |
| SDHB | 1p36.13 | 6 | 1.02 | 0 | 0.00 | 0.387 | 0.667 | (A) BRAF Class 1 Colorectal |
| CD79B | 17q23.3 | 6 | 1.01 | 0 | 0.00 | 0.387 | 0.667 | (A) BRAF Class 1 Colorectal |
| EZH1 | 17q21.2 | 5 | 3.79 | 0 | 0.00 | 0.389 | 0.668 | (A) BRAF Class 1 Colorectal |
| CRLF2 | 2.33 and Yp | 13 | 2.23 | 1 | 1.04 | 0.389 | 0.668 | (A) BRAF Class 1 Colorectal |
| GLI3 | 7p14.1 | 12 | 15.79 | 1 | 7.69 | 0.395 | 0.677 | (A) BRAF Class 1 Colorectal |
| GATA1 | Xp11.23 | 12 | 3.23 | 1 | 1.54 | 0.398 | 0.678 | (A) BRAF Class 1 Colorectal |
| CASR | [illegible]13.33-q21[illegible] | 0 | 0.00 | 1 | 50.00 | 0.4 | 0.678 | (B) BRAF Class 3 Colorectal |
| DDX31 | 9q34.13 | 0 | 0.00 | 1 | 50.00 | 0.4 | 0.678 | (B) BRAF Class 3 Colorectal |
| MLF1 | 3q25.32 | 1 | 50.00 | 0 | 0.00 | 0.4 | 0.678 | (A) BRAF Class 1 Colorectal |
| RPL22 | 1p36.31 | 1 | 50.00 | 0 | 0.00 | 0.4 | 0.678 | (A) BRAF Class 1 Colorectal |
| PMS1 | 2q32.2 | 26 | 4.61 | 3 | 3.26 | 0.401 | 0.678 | (A) BRAF Class 1 Colorectal |
| MAPK1 | 22q11.22 | 6 | 1.03 | 0 | 0.00 | 0.402 | 0.679 | (A) BRAF Class 1 Colorectal |
| ICOSLG | 21q22.3 | 13 | 4.50 | 1 | 2.17 | 0.403 | 0.679 | (A) BRAF Class 1 Colorectal |
| WRN | 8p12 | 24 | 8.73 | 5 | 10.87 | 0.404 | 0.679 | (B) BRAF Class 3 Colorectal |
| EPCAM | 2p21 | 6 | 1.21 | 0 | 0.00 | 0.405 | 0.679 | (A) BRAF Class 1 Colorectal |
| VTCN1 | 1p13.1-p12 | 6 | 2.08 | 0 | 0.00 | 0.409 | 0.679 | (A) BRAF Class 1 Colorectal |
| SF1 | 11q13.1 | 5 | 6.33 | 0 | 0.00 | 0.411 | 0.679 | (A) BRAF Class 1 Colorectal |
| CDH4 | 20q13.33 | 14 | 8.75 | 1 | 4.35 | 0.411 | 0.679 | (A) BRAF Class 1 Colorectal |
| PAXIP1 | 7q36.2 | 14 | 8.75 | 1 | 4.35 | 0.411 | 0.679 | (A) BRAF Class 1 Colorectal |
| KLHL6 | 3q27.1 | 6 | 3.92 | 0 | 0.00 | 0.412 | 0.679 | (A) BRAF Class 1 Colorectal |
| CDKN2A | 9p21.3 | 27 | 3.24 | 3 | 2.33 | 0.412 | 0.679 | (A) BRAF Class 1 Colorectal |
| PMS2 | 7p22.1 | 19 | 3.15 | 2 | 2.02 | 0.412 | 0.679 | (A) BRAF Class 1 Colorectal |
| SLITRK6 | 13q31.1 | 4 | 6.90 | 0 | 0.00 | 0.412 | 0.679 | (A) BRAF Class 1 Colorectal |
| STAT6 | 12q13.3 | 6 | 2.20 | 0 | 0.00 | 0.413 | 0.679 | (A) BRAF Class 1 Colorectal |
| GPC3 | Xq26.2 | 6 | 2.19 | 0 | 0.00 | 0.414 | 0.679 | (A) BRAF Class 1 Colorectal |
| PHF6 | Xq26.2 | 15 | 4.10 | 3 | 5.56 | 0.414 | 0.679 | (B) BRAF Class 3 Colorectal |
| FLT1 | 13q12.3 | 43 | 7.21 | 6 | 5.94 | 0.417 | 0.679 | (A) BRAF Class 1 Colorectal |
| SOX10 | 22q13.1 | 6 | 4.20 | 0 | 0.00 | 0.418 | 0.679 | (A) BRAF Class 1 Colorectal |
| HIST1H3J | 6p22.1 | 6 | 2.21 | 0 | 0.00 | 0.418 | 0.679 | (A) BRAF Class 1 Colorectal |
| MST1 | 3p21.31 | 6 | 2.21 | 0 | 0.00 | 0.418 | 0.679 | (A) BRAF Class 1 Colorectal |
| RAB35 | 12q24.23 | 6 | 2.21 | 0 | 0.00 | 0.418 | 0.679 | (A) BRAF Class 1 Colorectal |
| DKC1 | Xq28 | 3 | 1.88 | 1 | 4.35 | 0.418 | 0.679 | (B) BRAF Class 3 Colorectal |
| KEL | 7q34 | 4 | 2.84 | 1 | 6.25 | 0.42 | 0.68 | (B) BRAF Class 3 Colorectal |
| LRP1B | q22.1-q22.[illegible] | 33 | 23.08 | 4 | 18.18 | 0.421 | 0.68 | (A) BRAF Class 1 Colorectal |
| EXT1 | 8q24.11 | 6 | 2.22 | 0 | 0.00 | 0.425 | 0.685 | (A) BRAF Class 1 Colorectal |
| NCOA3 | 20q13.12 | 26 | 8.44 | 5 | 10.20 | 0.425 | 0.685 | (B) BRAF Class 3 Colorectal |
| TAF1 | Xq13.1 | 15 | 11.36 | 1 | 5.88 | 0.428 | 0.688 | (A) BRAF Class 1 Colorectal |
| CCNE1 | 19q12 | 18 | 2.98 | 2 | 1.96 | 0.431 | 0.689 | (A) BRAF Class 1 Colorectal |
| TOP1 | 20q12 | 17 | 4.74 | 2 | 3.13 | 0.431 | 0.689 | (A) BRAF Class 1 Colorectal |
| FGFR2 | 10q26.13 | 32 | 3.82 | 4 | 2.99 | 0.431 | 0.689 | (A) BRAF Class 1 Colorectal |
| CTNNA1 | 5q31.2 | 20 | 6.39 | 2 | 4.26 | 0.432 | 0.689 | (A) BRAF Class 1 Colorectal |
| SOX2 | 3q26.33 | 24 | 4.03 | 3 | 2.97 | 0.433 | 0.689 | (A) BRAF Class 1 Colorectal |
| MCL1 | 1q21.2 | 8 | 1.34 | 2 | 1.98 | 0.438 | 0.695 | (B) BRAF Class 3 Colorectal |
| AGO2 | 8q24.3 | 17 | 13.39 | 2 | 9.09 | 0.441 | 0.695 | (A) BRAF Class 1 Colorectal |

| | | | | | | | | |
|---|---|---|---|---|---|---|---|---|
| FANCF | 11p14.3 | 9 | 2.37 | 2 | 3.45 | 0.441 | 0.695 | (B) BRAF Class 3 Colorectal |
| NEIL1 | 15q24.2 | 6 | 3.75 | 0 | 0.00 | 0.441 | 0.695 | (A) BRAF Class 1 Colorectal |
| TRIM37 | 17q22 | 6 | 3.75 | 0 | 0.00 | 0.441 | 0.695 | (A) BRAF Class 1 Colorectal |
| UIMC1 | 5q35.2 | 6 | 3.59 | 0 | 0.00 | 0.442 | 0.695 | (A) BRAF Class 1 Colorectal |
| CARM1 | 19p13.2 | 5 | 3.94 | 0 | 0.00 | 0.445 | 0.697 | (A) BRAF Class 1 Colorectal |
| MSI1 | 12q24.31 | 5 | 3.94 | 0 | 0.00 | 0.445 | 0.697 | (A) BRAF Class 1 Colorectal |
| MAP3K13 | 3q27.2 | 5 | 1.58 | 0 | 0.00 | 0.447 | 0.7 | (A) BRAF Class 1 Colorectal |
| FUS | 16p11.2 | 5 | 1.96 | 0 | 0.00 | 0.449 | 0.701 | (A) BRAF Class 1 Colorectal |
| RTEL1 | 20q13.33 | 8 | 6.30 | 2 | 9.09 | 0.451 | 0.703 | (B) BRAF Class 3 Colorectal |
| BRD3 | 9q34.2 | 20 | 10.31 | 2 | 7.14 | 0.454 | 0.707 | (A) BRAF Class 1 Colorectal |
| HSP90AA1 | 14q32.31 | 4 | 3.01 | 1 | 5.88 | 0.457 | 0.709 | (B) BRAF Class 3 Colorectal |
| CRTC1 | 19p13.11 | 22 | 8.15 | 4 | 9.76 | 0.457 | 0.709 | (B) BRAF Class 3 Colorectal |
| SPOP | 17q21.33 | 5 | 0.95 | 0 | 0.00 | 0.458 | 0.709 | (A) BRAF Class 1 Colorectal |
| RAD51D | 17q12 | 11 | 2.10 | 1 | 1.12 | 0.46 | 0.71 | (A) BRAF Class 1 Colorectal |
| ABCB11 | 2q31.1 | 12 | 7.36 | 1 | 4.00 | 0.461 | 0.71 | (A) BRAF Class 1 Colorectal |
| DCUN1D1 | 3q26.33 | 5 | 1.73 | 0 | 0.00 | 0.462 | 0.71 | (A) BRAF Class 1 Colorectal |
| HSD3B1 | 1p12 | 7 | 4.96 | 0 | 0.00 | 0.464 | 0.711 | (A) BRAF Class 1 Colorectal |
| CDK6 | 7q21.2 | 11 | 1.82 | 1 | 0.98 | 0.464 | 0.711 | (A) BRAF Class 1 Colorectal |
| ZBTB7A | 19p13.3 | 9 | 10.98 | 1 | 20.00 | 0.465 | 0.711 | (B) BRAF Class 3 Colorectal |
| BCL2L12 | 19q13.33 | 5 | 2.10 | 0 | 0.00 | 0.465 | 0.711 | (A) BRAF Class 1 Colorectal |
| MLLT3 | 9p21.3 | 8 | 9.52 | 0 | 0.00 | 0.468 | 0.714 | (A) BRAF Class 1 Colorectal |
| INHBA | 7p14.1 | 16 | 4.82 | 2 | 3.39 | 0.472 | 0.716 | (A) BRAF Class 1 Colorectal |
| MPL | 1p34.2 | 24 | 3.02 | 3 | 2.34 | 0.472 | 0.716 | (A) BRAF Class 1 Colorectal |
| XRCC1 | 19q13.31 | 6 | 3.13 | 0 | 0.00 | 0.475 | 0.716 | (A) BRAF Class 1 Colorectal |
| PHF1 | 6p21.32 | 5 | 0.60 | 0 | 0.00 | 0.476 | 0.716 | (A) BRAF Class 1 Colorectal |
| BBC3 | 19q13.32 | 5 | 1.73 | 0 | 0.00 | 0.476 | 0.716 | (A) BRAF Class 1 Colorectal |
| E2F3 | 6p22.3 | 5 | 1.73 | 0 | 0.00 | 0.476 | 0.716 | (A) BRAF Class 1 Colorectal |
| HIST1H2BD | 6p22.2 | 5 | 1.73 | 0 | 0.00 | 0.476 | 0.716 | (A) BRAF Class 1 Colorectal |
| BCL2L11 | 2q13 | 5 | 1.72 | 0 | 0.00 | 0.477 | 0.716 | (A) BRAF Class 1 Colorectal |
| RAF1 | 3p25.2 | 21 | 3.32 | 4 | 3.85 | 0.479 | 0.719 | (B) BRAF Class 3 Colorectal |
| PIM1 | 6p21.2 | 5 | 0.88 | 0 | 0.00 | 0.482 | 0.721 | (A) BRAF Class 1 Colorectal |
| H3F3A | 1q42.12 | 5 | 0.82 | 0 | 0.00 | 0.483 | 0.721 | (A) BRAF Class 1 Colorectal |
| BTK | Xq22.1 | 10 | 2.71 | 1 | 1.52 | 0.483 | 0.721 | (A) BRAF Class 1 Colorectal |
| NFKBIA | 14q13.2 | 9 | 1.56 | 2 | 2.06 | 0.485 | 0.722 | (B) BRAF Class 3 Colorectal |
| CYSLTR2 | 13q14.2 | 5 | 3.16 | 0 | 0.00 | 0.489 | 0.723 | (A) BRAF Class 1 Colorectal |
| FANCE | 6p21.31 | 5 | 1.32 | 0 | 0.00 | 0.489 | 0.723 | (A) BRAF Class 1 Colorectal |
| MLH3 | 14q24.3 | 15 | 7.58 | 3 | 9.09 | 0.49 | 0.723 | (B) BRAF Class 3 Colorectal |
| INPP4B | 4q31.21 | 15 | 4.24 | 2 | 3.08 | 0.493 | 0.723 | (A) BRAF Class 1 Colorectal |
| RAD54B | 8q22.1 | 4 | 2.50 | 1 | 4.35 | 0.493 | 0.723 | (B) BRAF Class 3 Colorectal |
| XRCC6 | 22q13.2 | 4 | 2.50 | 1 | 4.35 | 0.493 | 0.723 | (B) BRAF Class 3 Colorectal |
| KCNQ1 | 1p15.5-p15. | 12 | 7.50 | 1 | 4.35 | 0.494 | 0.723 | (A) BRAF Class 1 Colorectal |
| U2AF1 | 21q22.3 | 15 | 2.50 | 3 | 2.94 | 0.499 | 0.723 | (B) BRAF Class 3 Colorectal |
| ZFHX4 | 8q21.13 | 2 | 40.00 | 1 | 20.00 | 0.5 | 0.723 | (A) BRAF Class 1 Colorectal |
| AJUBA | 14q11.2 | 1 | 20.00 | 0 | 0.00 | 0.5 | 0.723 | (A) BRAF Class 1 Colorectal |
| CHD1 | 5q15-q21.1 | 0 | 0.00 | 1 | 20.00 | 0.5 | 0.723 | (B) BRAF Class 3 Colorectal |
| CHD5 | 1p36.31 | 1 | 20.00 | 0 | 0.00 | 0.5 | 0.723 | (A) BRAF Class 1 Colorectal |
| CLDN18 | 3q22.3 | 0 | 0.00 | 1 | 20.00 | 0.5 | 0.723 | (B) BRAF Class 3 Colorectal |
| ZBTB20 | 3q13.31 | 1 | 20.00 | 0 | 0.00 | 0.5 | 0.723 | (A) BRAF Class 1 Colorectal |

| | | | | | | | | |
|---|---|---|---|---|---|---|---|---|
| MLLT1 | 19p13.3 | 1 | 33.33 | 0 | 0.00 | 0.5 | 0.723 | (A) BRAF Class 1 Colorectal |
| ADGRB3 | 6q12-q13 | 1 | 100.00 | 1 | 33.33 | 0.5 | 0.723 | (A) BRAF Class 1 Colorectal |
| STK11 | 19p13.3 | 23 | 2.76 | 4 | 3.10 | 0.5 | 0.723 | (B) BRAF Class 3 Colorectal |
| MAP2K4 | 17p12 | 27 | 4.47 | 5 | 4.90 | 0.502 | 0.723 | (B) BRAF Class 3 Colorectal |
| HIST1H3B | 6p22.2 | 5 | 0.95 | 0 | 0.00 | 0.503 | 0.723 | (A) BRAF Class 1 Colorectal |
| BRCC3 | Xq28 | 5 | 3.13 | 0 | 0.00 | 0.507 | 0.723 | (A) BRAF Class 1 Colorectal |
| HABP2 | 10q25.3 | 5 | 3.13 | 0 | 0.00 | 0.507 | 0.723 | (A) BRAF Class 1 Colorectal |
| UROD | 1p34.1 | 5 | 3.13 | 0 | 0.00 | 0.507 | 0.723 | (A) BRAF Class 1 Colorectal |
| NRG1 | 8p12 | 23 | 8.30 | 2 | 6.25 | 0.508 | 0.723 | (A) BRAF Class 1 Colorectal |
| RUNX1T1 | 8q21.3 | 18 | 5.94 | 2 | 4.44 | 0.508 | 0.723 | (A) BRAF Class 1 Colorectal |
| ITK | 5q33.3 | 5 | 3.11 | 0 | 0.00 | 0.509 | 0.723 | (A) BRAF Class 1 Colorectal |
| HRAS | 11p15.5 | 10 | 1.19 | 2 | 1.49 | 0.509 | 0.723 | (B) BRAF Class 3 Colorectal |
| ARHGAP35 | 19q13.32 | 10 | 6.02 | 1 | 3.57 | 0.51 | 0.723 | (A) BRAF Class 1 Colorectal |
| SMC1A | Xp11.22 | 5 | 2.55 | 0 | 0.00 | 0.51 | 0.723 | (A) BRAF Class 1 Colorectal |
| NUP93 | 16q13 | 9 | 2.71 | 2 | 3.39 | 0.514 | 0.727 | (B) BRAF Class 3 Colorectal |
| EPHB4 | 7q22.1 | 2 | 3.39 | 1 | 6.25 | 0.519 | 0.732 | (B) BRAF Class 3 Colorectal |
| ERCC3 | 2q14.3 | 17 | 3.04 | 3 | 3.45 | 0.519 | 0.732 | (B) BRAF Class 3 Colorectal |
| KCNIP1 | 5q35.1 | 3 | 5.36 | 1 | 9.09 | 0.521 | 0.732 | (B) BRAF Class 3 Colorectal |
| ERCC1 | 19q13.32 | 5 | 1.80 | 1 | 2.78 | 0.521 | 0.732 | (B) BRAF Class 3 Colorectal |
| NUF2 | 1q23.3 | 4 | 3.15 | 0 | 0.00 | 0.524 | 0.732 | (A) BRAF Class 1 Colorectal |
| SESN2 | 1p35.3 | 4 | 3.15 | 0 | 0.00 | 0.524 | 0.732 | (A) BRAF Class 1 Colorectal |
| SHOC2 | 10q25.2 | 4 | 3.15 | 0 | 0.00 | 0.524 | 0.732 | (A) BRAF Class 1 Colorectal |
| STK19 | 6p21.33 | 4 | 3.15 | 0 | 0.00 | 0.524 | 0.732 | (A) BRAF Class 1 Colorectal |
| PRKCZ | 1p36.33 | 4 | 5.26 | 0 | 0.00 | 0.525 | 0.733 | (A) BRAF Class 1 Colorectal |
| IGF1 | 12q23.2 | 4 | 1.30 | 0 | 0.00 | 0.527 | 0.734 | (A) BRAF Class 1 Colorectal |
| NFKBIE | 6p21.1 | 4 | 2.42 | 0 | 0.00 | 0.531 | 0.736 | (A) BRAF Class 1 Colorectal |
| CDKN2C | 1p32.3 | 4 | 0.68 | 0 | 0.00 | 0.532 | 0.736 | (A) BRAF Class 1 Colorectal |
| PTK2B | 8p21.2 | 10 | 6.17 | 1 | 3.85 | 0.534 | 0.736 | (A) BRAF Class 1 Colorectal |
| SLC34A2 | 4p15.2 | 12 | 5.91 | 1 | 3.70 | 0.534 | 0.736 | (A) BRAF Class 1 Colorectal |
| CDK8 | 13q12.13 | 22 | 4.23 | 4 | 4.55 | 0.534 | 0.736 | (B) BRAF Class 3 Colorectal |
| DHX15 | 4p15.2 | 10 | 12.20 | 0 | 0.00 | 0.535 | 0.736 | (A) BRAF Class 1 Colorectal |
| TCF7L1 | 2p11.2 | 4 | 3.67 | 0 | 0.00 | 0.538 | 0.736 | (A) BRAF Class 1 Colorectal |
| FAAP100 | 17q25.3 | 11 | 6.88 | 1 | 4.35 | 0.539 | 0.736 | (A) BRAF Class 1 Colorectal |
| LIG4 | 13q33.3 | 11 | 6.88 | 1 | 4.35 | 0.539 | 0.736 | (A) BRAF Class 1 Colorectal |
| RHBDF2 | 17q25.1 | 11 | 6.88 | 1 | 4.35 | 0.539 | 0.736 | (A) BRAF Class 1 Colorectal |
| ERBB2 | 17q12 | 49 | 5.85 | 8 | 5.97 | 0.539 | 0.736 | (B) BRAF Class 3 Colorectal |
| CD276 | 15q24.1 | 10 | 3.46 | 1 | 2.17 | 0.54 | 0.736 | (A) BRAF Class 1 Colorectal |
| IRF4 | 6p25.3 | 14 | 3.89 | 2 | 3.08 | 0.546 | 0.743 | (A) BRAF Class 1 Colorectal |
| SDHD | 11q23.1 | 4 | 0.68 | 1 | 0.99 | 0.547 | 0.743 | (B) BRAF Class 3 Colorectal |
| PPP2R1A | 19q13.41 | 15 | 2.84 | 2 | 2.25 | 0.547 | 0.743 | (A) BRAF Class 1 Colorectal |
| EED | 11q14.2 | 4 | 1.03 | 0 | 0.00 | 0.548 | 0.743 | (A) BRAF Class 1 Colorectal |
| GREM1 | 15q13.3 | 4 | 0.80 | 0 | 0.00 | 0.55 | 0.744 | (A) BRAF Class 1 Colorectal |
| BRAF | 7q34 | 834 | 99.52 | 134 | 100.00 | 0.552 | 0.745 | (B) BRAF Class 3 Colorectal |
| H3F3C | 12p11.21 | 4 | 1.38 | 0 | 0.00 | 0.552 | 0.745 | (A) BRAF Class 1 Colorectal |
| YES1 | 18p11.32 | 11 | 3.81 | 2 | 4.35 | 0.554 | 0.745 | (B) BRAF Class 3 Colorectal |
| MAP2K2 | 19p13.3 | 9 | 1.70 | 1 | 1.10 | 0.555 | 0.745 | (A) BRAF Class 1 Colorectal |
| PDK1 | 2q31.1 | 4 | 2.61 | 0 | 0.00 | 0.555 | 0.745 | (A) BRAF Class 1 Colorectal |
| DIS3L2 | 2q37.1 | 10 | 6.13 | 1 | 4.00 | 0.556 | 0.745 | (A) BRAF Class 1 Colorectal |

| | | | | | | | | |
|---|---|---|---|---|---|---|---|---|
| KAT6A | 8p11.21 | 36 | 11.88 | 5 | 11.11 | 0.558 | 0.747 | (A) BRAF Class 1 Colorectal |
| HIST1H3E | 6p22.2 | 4 | 1.48 | 0 | 0.00 | 0.56 | 0.749 | (A) BRAF Class 1 Colorectal |
| PRPF8 | 17p13.3 | 8 | 10.13 | 1 | 6.67 | 0.561 | 0.749 | (A) BRAF Class 1 Colorectal |
| SBDS | 7q11.21 | 4 | 1.48 | 0 | 0.00 | 0.566 | 0.753 | (A) BRAF Class 1 Colorectal |
| LZTR1 | 21\|22q11.1 | 13 | 9.42 | 2 | 10.53 | 0.568 | 0.753 | (B) BRAF Class 3 Colorectal |
| LATS2 | 13q12.11 | 31 | 10.47 | 5 | 10.64 | 0.568 | 0.753 | (B) BRAF Class 3 Colorectal |
| RRAGC | 1p34.3 | 3 | 2.27 | 0 | 0.00 | 0.57 | 0.753 | (A) BRAF Class 1 Colorectal |
| RRAS | 19q13.33 | 3 | 2.27 | 0 | 0.00 | 0.57 | 0.753 | (A) BRAF Class 1 Colorectal |
| NAB2 | 12q13.3 | 9 | 10.98 | 0 | 0.00 | 0.571 | 0.753 | (A) BRAF Class 1 Colorectal |
| EZH2 | 7q36.1 | 9 | 1.22 | 1 | 0.82 | 0.575 | 0.753 | (A) BRAF Class 1 Colorectal |
| DNAH9 | 17p12 | 1 | 12.50 | 1 | 25.00 | 0.576 | 0.753 | (B) BRAF Class 3 Colorectal |
| CSDE1 | 1p13.2 | 6 | 2.87 | 1 | 3.70 | 0.578 | 0.753 | (B) BRAF Class 3 Colorectal |
| KNSTRN | 15q15.1 | 3 | 2.13 | 0 | 0.00 | 0.579 | 0.753 | (A) BRAF Class 1 Colorectal |
| GABRA6 | 5q34 | 5 | 3.55 | 0 | 0.00 | 0.58 | 0.753 | (A) BRAF Class 1 Colorectal |
| NT5C2 | q24.32-q24. | 5 | 2.40 | 1 | 3.13 | 0.58 | 0.753 | (B) BRAF Class 3 Colorectal |
| WAS | Xp11.23 | 4 | 2.06 | 0 | 0.00 | 0.581 | 0.753 | (A) BRAF Class 1 Colorectal |
| ACVR1B | 12q13.13 | 4 | 2.74 | 0 | 0.00 | 0.581 | 0.753 | (A) BRAF Class 1 Colorectal |
| SNCAIP | 5q23.2 | 4 | 2.74 | 0 | 0.00 | 0.581 | 0.753 | (A) BRAF Class 1 Colorectal |
| KLF2 | 19p13.11 | 4 | 2.50 | 0 | 0.00 | 0.582 | 0.753 | (A) BRAF Class 1 Colorectal |
| PNKP | 19q13.33 | 4 | 2.50 | 0 | 0.00 | 0.582 | 0.753 | (A) BRAF Class 1 Colorectal |
| POLB | 8p11.21 | 4 | 2.50 | 0 | 0.00 | 0.582 | 0.753 | (A) BRAF Class 1 Colorectal |
| RNF8 | 6p21.2 | 4 | 2.50 | 0 | 0.00 | 0.582 | 0.753 | (A) BRAF Class 1 Colorectal |
| VEGFA | 6p21.1 | 9 | 1.80 | 1 | 1.23 | 0.584 | 0.755 | (A) BRAF Class 1 Colorectal |
| MYB | 6q23.3 | 5 | 1.32 | 1 | 1.67 | 0.587 | 0.758 | (B) BRAF Class 3 Colorectal |
| GATA6 | 18q11.2 | 17 | 4.51 | 2 | 3.85 | 0.589 | 0.759 | (A) BRAF Class 1 Colorectal |
| ZNF703 | 8p11.23 | 12 | 7.84 | 2 | 8.33 | 0.594 | 0.765 | (B) BRAF Class 3 Colorectal |
| NUTM1 | 15q14 | 5 | 3.85 | 0 | 0.00 | 0.595 | 0.765 | (A) BRAF Class 1 Colorectal |
| PTCH2 | 1p34.1 | 1 | 5.56 | 0 | 0.00 | 0.6 | 0.766 | (A) BRAF Class 1 Colorectal |
| ABCB4 | 7q21.12 | 1 | 33.33 | 0 | 0.00 | 0.6 | 0.766 | (A) BRAF Class 1 Colorectal |
| DYNC1I1 | 7q21.3 | 0 | 0.00 | 1 | 33.33 | 0.6 | 0.766 | (B) BRAF Class 3 Colorectal |
| SMARCAL1 | 2q35 | 1 | 33.33 | 0 | 0.00 | 0.6 | 0.766 | (A) BRAF Class 1 Colorectal |
| GNA11 | 19p13.3 | 12 | 1.46 | 2 | 1.50 | 0.602 | 0.768 | (B) BRAF Class 3 Colorectal |
| NEGR1 | 1p31.1 | 12 | 3.67 | 2 | 3.77 | 0.605 | 0.771 | (B) BRAF Class 3 Colorectal |
| FGF5 | 4q21.21 | 8 | 9.76 | 0 | 0.00 | 0.61 | 0.775 | (A) BRAF Class 1 Colorectal |
| RPA1 | 17p13.3 | 5 | 2.81 | 1 | 3.33 | 0.612 | 0.777 | (B) BRAF Class 3 Colorectal |
| XRCC5 | 2q35 | 6 | 3.59 | 1 | 4.17 | 0.616 | 0.777 | (B) BRAF Class 3 Colorectal |
| MUS81 | 11q13.1 | 6 | 3.75 | 1 | 4.35 | 0.616 | 0.777 | (B) BRAF Class 3 Colorectal |
| RINT1 | 7q22.3 | 6 | 3.75 | 1 | 4.35 | 0.616 | 0.777 | (B) BRAF Class 3 Colorectal |
| CCNQ | Xq28 | 3 | 2.36 | 0 | 0.00 | 0.617 | 0.777 | (A) BRAF Class 1 Colorectal |
| TAP2 | 6p21.32 | 3 | 2.36 | 0 | 0.00 | 0.617 | 0.777 | (A) BRAF Class 1 Colorectal |
| CRTC2 | 1q21.3 | 3 | 3.95 | 0 | 0.00 | 0.619 | 0.778 | (A) BRAF Class 1 Colorectal |
| RBL2 | 16q12.2 | 3 | 3.95 | 0 | 0.00 | 0.619 | 0.778 | (A) BRAF Class 1 Colorectal |
| CD274 | 9p24.1 | 8 | 1.36 | 1 | 1.02 | 0.625 | 0.784 | (A) BRAF Class 1 Colorectal |
| NCOA2 | 8q13.3 | 2 | 5.13 | 0 | 0.00 | 0.63 | 0.789 | (A) BRAF Class 1 Colorectal |
| XPA | 9q22.33 | 6 | 2.22 | 1 | 2.44 | 0.632 | 0.79 | (B) BRAF Class 3 Colorectal |
| PDPK1 | 16p13.3 | 3 | 1.03 | 0 | 0.00 | 0.633 | 0.79 | (A) BRAF Class 1 Colorectal |
| ENG | 9q34.11 | 9 | 5.63 | 1 | 4.35 | 0.635 | 0.791 | (A) BRAF Class 1 Colorectal |
| BIRC3 | 11q22.2 | 6 | 1.97 | 1 | 2.13 | 0.637 | 0.793 | (B) BRAF Class 3 Colorectal |

| | | | | | | | | |
|---|---|---|---|---|---|---|---|---|
| PMAIP1 | 18q21.32 | 3 | 1.04 | 0 | 0.00 | 0.641 | 0.796 | (A) BRAF Class 1 Colorectal |
| RYBP | 3p13 | 3 | 1.04 | 0 | 0.00 | 0.641 | 0.796 | (A) BRAF Class 1 Colorectal |
| PIK3R3 | 1p34.1 | 8 | 2.77 | 1 | 2.17 | 0.643 | 0.796 | (A) BRAF Class 1 Colorectal |
| FGF23 | 12p13.32 | 3 | 1.96 | 0 | 0.00 | 0.644 | 0.796 | (A) BRAF Class 1 Colorectal |
| FGF6 | 12p13.32 | 3 | 1.96 | 0 | 0.00 | 0.644 | 0.796 | (A) BRAF Class 1 Colorectal |
| RIF1 | 2q23.3 | 21 | 13.13 | 3 | 13.04 | 0.646 | 0.797 | (A) BRAF Class 1 Colorectal |
| IRF2 | 4q35.1 | 4 | 2.84 | 0 | 0.00 | 0.648 | 0.797 | (A) BRAF Class 1 Colorectal |
| HIST3H3 | 1q42.13 | 3 | 1.11 | 0 | 0.00 | 0.648 | 0.797 | (A) BRAF Class 1 Colorectal |
| MAX | 14q23.3 | 8 | 1.73 | 1 | 1.39 | 0.652 | 0.798 | (A) BRAF Class 1 Colorectal |
| SS18 | 18q11.2 | 3 | 1.52 | 0 | 0.00 | 0.654 | 0.798 | (A) BRAF Class 1 Colorectal |
| PDCD1LG2 | 9p24.1 | 8 | 1.57 | 1 | 1.27 | 0.655 | 0.798 | (A) BRAF Class 1 Colorectal |
| EML4 | 2p21 | 5 | 4.31 | 0 | 0.00 | 0.657 | 0.798 | (A) BRAF Class 1 Colorectal |
| CCN6 | 6q21 | 8 | 5.59 | 1 | 4.55 | 0.657 | 0.798 | (A) BRAF Class 1 Colorectal |
| NKX3-1 | 8p21.2 | 8 | 1.78 | 1 | 1.45 | 0.658 | 0.798 | (A) BRAF Class 1 Colorectal |
| MAF | 16q23.2 | 3 | 1.47 | 0 | 0.00 | 0.661 | 0.798 | (A) BRAF Class 1 Colorectal |
| POT1 | 7q31.33 | 6 | 3.00 | 1 | 3.03 | 0.662 | 0.798 | (B) BRAF Class 3 Colorectal |
| BUB1B | 15q15.1 | 8 | 2.96 | 1 | 2.44 | 0.663 | 0.798 | (A) BRAF Class 1 Colorectal |
| USP8 | 15q21.2 | 7 | 4.35 | 1 | 4.35 | 0.664 | 0.798 | (B) BRAF Class 3 Colorectal |
| MAFB | 20q12 | 3 | 1.55 | 0 | 0.00 | 0.666 | 0.798 | (A) BRAF Class 1 Colorectal |
| FIP1L1 | 4q12 | 1 | 8.33 | 0 | 0.00 | 0.667 | 0.798 | (A) BRAF Class 1 Colorectal |
| EGLN1 | 1q42.2 | 3 | 1.88 | 0 | 0.00 | 0.667 | 0.798 | (A) BRAF Class 1 Colorectal |
| FAH | 15q25.1 | 3 | 1.88 | 0 | 0.00 | 0.667 | 0.798 | (A) BRAF Class 1 Colorectal |
| GBA | 1q22 | 3 | 1.88 | 0 | 0.00 | 0.667 | 0.798 | (A) BRAF Class 1 Colorectal |
| ID4 | 6p22.3 | 3 | 1.88 | 0 | 0.00 | 0.667 | 0.798 | (A) BRAF Class 1 Colorectal |
| NECTIN4 | 1q23.3 | 3 | 1.88 | 0 | 0.00 | 0.667 | 0.798 | (A) BRAF Class 1 Colorectal |
| RHOT1 | 17q11.2 | 3 | 1.88 | 0 | 0.00 | 0.667 | 0.798 | (A) BRAF Class 1 Colorectal |
| TDG | 12q23.3 | 3 | 1.88 | 0 | 0.00 | 0.667 | 0.798 | (A) BRAF Class 1 Colorectal |
| AURKA | 20q13.2 | 12 | 2.01 | 2 | 1.98 | 0.669 | 0.8 | (A) BRAF Class 1 Colorectal |
| BCL2 | 18q21.33 | 7 | 1.18 | 1 | 0.99 | 0.671 | 0.8 | (A) BRAF Class 1 Colorectal |
| CD58 | 1p13.1 | 3 | 2.78 | 0 | 0.00 | 0.675 | 0.804 | (A) BRAF Class 1 Colorectal |
| TRAF3 | 14q32.32 | 7 | 3.54 | 1 | 3.03 | 0.68 | 0.809 | (A) BRAF Class 1 Colorectal |
| RAC1 | 7p22.1 | 7 | 1.26 | 1 | 1.09 | 0.683 | 0.812 | (A) BRAF Class 1 Colorectal |
| NR0B1 | Xp21.2 | 8 | 5.00 | 1 | 4.35 | 0.685 | 0.812 | (A) BRAF Class 1 Colorectal |
| CDC42 | 1p36.12 | 2 | 1.52 | 0 | 0.00 | 0.688 | 0.814 | (A) BRAF Class 1 Colorectal |
| DUSP4 | 8p12 | 2 | 1.52 | 0 | 0.00 | 0.688 | 0.814 | (A) BRAF Class 1 Colorectal |
| POLH | 6p21.1 | 3 | 1.56 | 0 | 0.00 | 0.691 | 0.817 | (A) BRAF Class 1 Colorectal |
| KIF5B | 10p11.22 | 6 | 5.22 | 0 | 0.00 | 0.696 | 0.818 | (A) BRAF Class 1 Colorectal |
| TFRC | 3q29 | 6 | 7.23 | 0 | 0.00 | 0.696 | 0.818 | (A) BRAF Class 1 Colorectal |
| CADM2 | 3p12.1 | 2 | 3.57 | 0 | 0.00 | 0.697 | 0.818 | (A) BRAF Class 1 Colorectal |
| RACK1 | 5q35.3 | 2 | 3.57 | 0 | 0.00 | 0.697 | 0.818 | (A) BRAF Class 1 Colorectal |
| SAMD9L | 7q21.2 | 1 | 33.33 | 1 | 50.00 | 0.7 | 0.821 | (B) BRAF Class 3 Colorectal |
| TMEM127 | 2q11.2 | 7 | 1.55 | 1 | 1.41 | 0.703 | 0.822 | (A) BRAF Class 1 Colorectal |
| DDR1 | 6p21.33 | 1 | 3.85 | 0 | 0.00 | 0.703 | 0.822 | (A) BRAF Class 1 Colorectal |
| PAK3 | Xq23 | 7 | 4.90 | 1 | 4.55 | 0.711 | 0.83 | (A) BRAF Class 1 Colorectal |
| LAMP1 | 13q34 | 4 | 3.48 | 0 | 0.00 | 0.713 | 0.831 | (A) BRAF Class 1 Colorectal |
| RSPO2 | 8q23.1 | 6 | 2.73 | 1 | 2.56 | 0.716 | 0.831 | (A) BRAF Class 1 Colorectal |
| CBLB | 3q13.11 | 7 | 2.58 | 1 | 2.44 | 0.717 | 0.831 | (A) BRAF Class 1 Colorectal |
| TYRO3 | 15q15.1 | 1 | 4.35 | 0 | 0.00 | 0.719 | 0.831 | (A) BRAF Class 1 Colorectal |

| | | | | | | | | |
|---|---|---|---|---|---|---|---|---|
| FANCB | Xp22.2 | 8 | 4.02 | 1 | 3.85 | 0.721 | 0.831 | (A) BRAF Class 1 Colorectal |
| MSI2 | 17q22 | 2 | 1.57 | 0 | 0.00 | 0.726 | 0.831 | (A) BRAF Class 1 Colorectal |
| PPP4R2 | 3p13 | 2 | 1.57 | 0 | 0.00 | 0.726 | 0.831 | (A) BRAF Class 1 Colorectal |
| RAC2 | 22q13.1 | 2 | 1.57 | 0 | 0.00 | 0.726 | 0.831 | (A) BRAF Class 1 Colorectal |
| SESN1 | 6q21 | 2 | 1.57 | 0 | 0.00 | 0.726 | 0.831 | (A) BRAF Class 1 Colorectal |
| SMYD3 | 1q44 | 2 | 1.57 | 0 | 0.00 | 0.726 | 0.831 | (A) BRAF Class 1 Colorectal |
| SMARCA2 | 9p24.3 | 1 | 16.67 | 1 | 20.00 | 0.727 | 0.831 | (B) BRAF Class 3 Colorectal |
| KMT5A | 12q24.31 | 2 | 1.49 | 0 | 0.00 | 0.728 | 0.831 | (A) BRAF Class 1 Colorectal |
| FKBP9 | 7p14.3 | 2 | 2.63 | 0 | 0.00 | 0.728 | 0.831 | (A) BRAF Class 1 Colorectal |
| PRAME | 22q11.22 | 2 | 2.63 | 0 | 0.00 | 0.728 | 0.831 | (A) BRAF Class 1 Colorectal |
| PSMD13 | 11p15.5 | 2 | 2.63 | 0 | 0.00 | 0.728 | 0.831 | (A) BRAF Class 1 Colorectal |
| RHPN2 | 19q13.11 | 2 | 2.63 | 0 | 0.00 | 0.728 | 0.831 | (A) BRAF Class 1 Colorectal |
| RAD51B | 14q24.1 | 2 | 0.61 | 0 | 0.00 | 0.729 | 0.831 | (A) BRAF Class 1 Colorectal |
| DMD | :p21.2-p21.: | 6 | 7.89 | 1 | 7.69 | 0.73 | 0.831 | (A) BRAF Class 1 Colorectal |
| RHEB | 7q36.1 | 2 | 0.40 | 0 | 0.00 | 0.738 | 0.837 | (A) BRAF Class 1 Colorectal |
| ELOC | 8q21.11 | 2 | 0.46 | 0 | 0.00 | 0.742 | 0.837 | (A) BRAF Class 1 Colorectal |
| EPB41L4A | q22.1-q22.: | 2 | 0.24 | 0 | 0.00 | 0.743 | 0.837 | (A) BRAF Class 1 Colorectal |
| MT-ND5 | - | 2 | 0.24 | 0 | 0.00 | 0.743 | 0.837 | (A) BRAF Class 1 Colorectal |
| TRBC2 | 7q34 | 2 | 0.24 | 0 | 0.00 | 0.743 | 0.837 | (A) BRAF Class 1 Colorectal |
| IL10 | 1q32.1 | 2 | 0.69 | 0 | 0.00 | 0.744 | 0.837 | (A) BRAF Class 1 Colorectal |
| FGF10 | 5p12 | 2 | 1.31 | 0 | 0.00 | 0.747 | 0.837 | (A) BRAF Class 1 Colorectal |
| GNA13 | 17q24.1 | 2 | 1.31 | 0 | 0.00 | 0.747 | 0.837 | (A) BRAF Class 1 Colorectal |
| CENPA | 2p23.3 | 2 | 0.74 | 0 | 0.00 | 0.749 | 0.837 | (A) BRAF Class 1 Colorectal |
| EIF4E | 4q23 | 2 | 0.74 | 0 | 0.00 | 0.749 | 0.837 | (A) BRAF Class 1 Colorectal |
| HIST1H3G | 6p22.2 | 2 | 0.74 | 0 | 0.00 | 0.749 | 0.837 | (A) BRAF Class 1 Colorectal |
| HIST1H3H | 6p22.1 | 2 | 0.74 | 0 | 0.00 | 0.749 | 0.837 | (A) BRAF Class 1 Colorectal |
| HIST1H3I | 6p22.1 | 2 | 0.74 | 0 | 0.00 | 0.749 | 0.837 | (A) BRAF Class 1 Colorectal |
| BOD1L1 | 4p15.33 | 0 | 0.00 | 1 | 33.33 | 0.75 | 0.837 | (B) BRAF Class 3 Colorectal |
| FGA | 4q31.3 | 0 | 0.00 | 1 | 33.33 | 0.75 | 0.837 | (B) BRAF Class 3 Colorectal |
| SH2D1A | Xq25 | 2 | 0.41 | 0 | 0.00 | 0.751 | 0.837 | (A) BRAF Class 1 Colorectal |
| H3F3B | 17q25.1 | 2 | 0.43 | 0 | 0.00 | 0.751 | 0.837 | (A) BRAF Class 1 Colorectal |
| SMARCE1 | 17q21.2 | 2 | 1.21 | 0 | 0.00 | 0.754 | 0.837 | (A) BRAF Class 1 Colorectal |
| JAZF1 | p15.2-p15. | 2 | 1.01 | 0 | 0.00 | 0.754 | 0.837 | (A) BRAF Class 1 Colorectal |
| CD74 | 5q33.1 | 3 | 2.38 | 0 | 0.00 | 0.76 | 0.839 | (A) BRAF Class 1 Colorectal |
| TAL1 | 1p33 | 2 | 1.03 | 0 | 0.00 | 0.763 | 0.839 | (A) BRAF Class 1 Colorectal |
| DCLRE1C | 10p13 | 2 | 1.25 | 0 | 0.00 | 0.764 | 0.839 | (A) BRAF Class 1 Colorectal |
| EME1 | 17q21.33 | 2 | 1.25 | 0 | 0.00 | 0.764 | 0.839 | (A) BRAF Class 1 Colorectal |
| HFE | 6p22.2 | 2 | 1.25 | 0 | 0.00 | 0.764 | 0.839 | (A) BRAF Class 1 Colorectal |
| HMBS | 11q23.3 | 2 | 1.25 | 0 | 0.00 | 0.764 | 0.839 | (A) BRAF Class 1 Colorectal |
| OGG1 | 3p25.3 | 2 | 1.25 | 0 | 0.00 | 0.764 | 0.839 | (A) BRAF Class 1 Colorectal |
| UBE2T | 1q32.1 | 2 | 1.25 | 0 | 0.00 | 0.764 | 0.839 | (A) BRAF Class 1 Colorectal |
| FRS2 | 12q15 | 2 | 1.47 | 0 | 0.00 | 0.769 | 0.844 | (A) BRAF Class 1 Colorectal |
| ETS1 | 11q24.3 | 2 | 1.67 | 0 | 0.00 | 0.789 | 0.865 | (A) BRAF Class 1 Colorectal |
| CDH5 | 16q21 | 1 | 2.17 | 0 | 0.00 | 0.793 | 0.868 | (A) BRAF Class 1 Colorectal |
| MYH9 | 22q12.3 | 1 | 2.56 | 0 | 0.00 | 0.796 | 0.869 | (A) BRAF Class 1 Colorectal |
| PDGFB | 22q13.1 | 1 | 2.56 | 0 | 0.00 | 0.796 | 0.869 | (A) BRAF Class 1 Colorectal |
| LTK | 15q15.1 | 1 | 1.79 | 0 | 0.00 | 0.8 | 0.872 | (A) BRAF Class 1 Colorectal |
| PPP2R2A | 8p21.2 | 3 | 3.26 | 0 | 0.00 | 0.801 | 0.872 | (A) BRAF Class 1 Colorectal |

| | | | | | | | | |
|---|---|---|---|---|---|---|---|---|
| HNRNPK | 9q21.32 | 2 | 2.33 | 0 | 0.00 | 0.802 | 0.872 | (A) BRAF Class 1 Colorectal |
| ZBTB2 | 6q25.1 | 2 | 1.53 | 0 | 0.00 | 0.816 | 0.882 | (A) BRAF Class 1 Colorectal |
| TLR4 | 9q33.1 | 1 | 1.10 | 0 | 0.00 | 0.827 | 0.882 | (A) BRAF Class 1 Colorectal |
| ELF3 | 1q32.1 | 1 | 0.76 | 0 | 0.00 | 0.83 | 0.882 | (A) BRAF Class 1 Colorectal |
| RRAS2 | 11p15.2 | 1 | 0.76 | 0 | 0.00 | 0.83 | 0.882 | (A) BRAF Class 1 Colorectal |
| CSNK1A1 | 5q32 | 3 | 3.66 | 0 | 0.00 | 0.836 | 0.882 | (A) BRAF Class 1 Colorectal |
| FGF2 | 4q28.1 | 3 | 3.66 | 0 | 0.00 | 0.836 | 0.882 | (A) BRAF Class 1 Colorectal |
| RPS6KB1 | 17q23.1 | 3 | 3.66 | 0 | 0.00 | 0.836 | 0.882 | (A) BRAF Class 1 Colorectal |
| NPRL2 | 3p21.31 | 1 | 1.79 | 0 | 0.00 | 0.836 | 0.882 | (A) BRAF Class 1 Colorectal |
| SQSTM1 | 5q35.3 | 1 | 1.79 | 0 | 0.00 | 0.836 | 0.882 | (A) BRAF Class 1 Colorectal |
| BIRC5 | 17q25.3 | 1 | 2.78 | 0 | 0.00 | 0.837 | 0.882 | (A) BRAF Class 1 Colorectal |
| ZMYM3 | Xq13.1 | 1 | 2.70 | 0 | 0.00 | 0.841 | 0.882 | (A) BRAF Class 1 Colorectal |
| TFEB | 6p21.1 | 1 | 2.63 | 0 | 0.00 | 0.844 | 0.882 | (A) BRAF Class 1 Colorectal |
| WWTR1 | 3q25.1 | 1 | 0.76 | 0 | 0.00 | 0.845 | 0.882 | (A) BRAF Class 1 Colorectal |
| FLI1 | 11q24.3 | 2 | 1.74 | 0 | 0.00 | 0.846 | 0.882 | (A) BRAF Class 1 Colorectal |
| CDK1 | 10q21.2 | 1 | 1.32 | 0 | 0.00 | 0.854 | 0.882 | (A) BRAF Class 1 Colorectal |
| CDK9 | 9q34.11 | 1 | 1.32 | 0 | 0.00 | 0.854 | 0.882 | (A) BRAF Class 1 Colorectal |
| GSTM5 | 1p13.3 | 1 | 1.32 | 0 | 0.00 | 0.854 | 0.882 | (A) BRAF Class 1 Colorectal |
| CDK2 | 12q13.2 | 1 | 1.18 | 0 | 0.00 | 0.859 | 0.882 | (A) BRAF Class 1 Colorectal |
| ANKRD30A | 10p11.21 | 1 | 0.12 | 0 | 0.00 | 0.862 | 0.882 | (A) BRAF Class 1 Colorectal |
| AP1G1 | 16q22.2 | 1 | 0.12 | 0 | 0.00 | 0.862 | 0.882 | (A) BRAF Class 1 Colorectal |
| CAMK1 | 3p25.3 | 1 | 0.12 | 0 | 0.00 | 0.862 | 0.882 | (A) BRAF Class 1 Colorectal |
| CUL1 | 7q36.1 | 1 | 0.12 | 0 | 0.00 | 0.862 | 0.882 | (A) BRAF Class 1 Colorectal |
| EIF3E | 8q23.1 | 1 | 0.12 | 0 | 0.00 | 0.862 | 0.882 | (A) BRAF Class 1 Colorectal |
| EMID1 | 22q12.2 | 1 | 0.12 | 0 | 0.00 | 0.862 | 0.882 | (A) BRAF Class 1 Colorectal |
| F11-AS1 | 4q35.2 | 1 | 0.12 | 0 | 0.00 | 0.862 | 0.882 | (A) BRAF Class 1 Colorectal |
| GOLT1A | 1q32.1 | 1 | 0.12 | 0 | 0.00 | 0.862 | 0.882 | (A) BRAF Class 1 Colorectal |
| GRAPL | 17p11.2 | 1 | 0.12 | 0 | 0.00 | 0.862 | 0.882 | (A) BRAF Class 1 Colorectal |
| HLA-DOB | 6p21.32 | 1 | 0.12 | 0 | 0.00 | 0.862 | 0.882 | (A) BRAF Class 1 Colorectal |
| HOOK2 | 19p13.13 | 1 | 0.12 | 0 | 0.00 | 0.862 | 0.882 | (A) BRAF Class 1 Colorectal |
| NDUFB2 | 7q34 | 1 | 0.12 | 0 | 0.00 | 0.862 | 0.882 | (A) BRAF Class 1 Colorectal |
| PLEKHA7 | 1p15.2-p15. | 1 | 0.12 | 0 | 0.00 | 0.862 | 0.882 | (A) BRAF Class 1 Colorectal |
| SCAP | 3p21.31 | 1 | 0.12 | 0 | 0.00 | 0.862 | 0.882 | (A) BRAF Class 1 Colorectal |
| TRAJ32 | 14q11.2 | 1 | 0.12 | 0 | 0.00 | 0.862 | 0.882 | (A) BRAF Class 1 Colorectal |
| TRBJ2-2 | 7q34 | 1 | 0.12 | 0 | 0.00 | 0.862 | 0.882 | (A) BRAF Class 1 Colorectal |
| TRGJP | 7p14.1 | 1 | 0.12 | 0 | 0.00 | 0.862 | 0.882 | (A) BRAF Class 1 Colorectal |
| TSPOAP1 | 17q22 | 1 | 0.12 | 0 | 0.00 | 0.862 | 0.882 | (A) BRAF Class 1 Colorectal |
| ARFRP1 | 20q13.33 | 1 | 0.66 | 0 | 0.00 | 0.864 | 0.882 | (A) BRAF Class 1 Colorectal |
| HIST2H3D | 1q21.2 | 1 | 0.37 | 0 | 0.00 | 0.865 | 0.882 | (A) BRAF Class 1 Colorectal |
| HIST1H3D | 6p22.2 | 1 | 0.37 | 0 | 0.00 | 0.866 | 0.882 | (A) BRAF Class 1 Colorectal |
| HIST1H3F | 6p22.2 | 1 | 0.37 | 0 | 0.00 | 0.866 | 0.882 | (A) BRAF Class 1 Colorectal |
| LMO2 | 11p13 | 1 | 0.42 | 0 | 0.00 | 0.868 | 0.882 | (A) BRAF Class 1 Colorectal |
| BLNK | 10q24.1 | 1 | 3.03 | 0 | 0.00 | 0.868 | 0.882 | (A) BRAF Class 1 Colorectal |
| IRF8 | 16q24.1 | 1 | 3.03 | 0 | 0.00 | 0.868 | 0.882 | (A) BRAF Class 1 Colorectal |
| ITGA9 | 3p22.2 | 1 | 3.03 | 0 | 0.00 | 0.868 | 0.882 | (A) BRAF Class 1 Colorectal |
| RRM1 | 11p15.4 | 1 | 3.03 | 0 | 0.00 | 0.868 | 0.882 | (A) BRAF Class 1 Colorectal |
| TRIM24 | 7q33-q34 | 1 | 3.03 | 0 | 0.00 | 0.868 | 0.882 | (A) BRAF Class 1 Colorectal |
| FOXO3 | 6q21 | 1 | 2.94 | 0 | 0.00 | 0.872 | 0.882 | (A) BRAF Class 1 Colorectal |

| | | | | | | | | |
|---|---|---|---|---|---|---|---|---|
| KNL1 | 15q15.1 | 1 | 2.94 | 0 | 0.00 | 0.872 | 0.882 | (A) BRAF Class 1 Colorectal |
| LCK | 1p35.2 | 1 | 2.94 | 0 | 0.00 | 0.872 | 0.882 | (A) BRAF Class 1 Colorectal |
| MUC1 | 1q22 | 1 | 2.94 | 0 | 0.00 | 0.872 | 0.882 | (A) BRAF Class 1 Colorectal |
| PDE4DIP | 1q21.2 | 1 | 2.94 | 0 | 0.00 | 0.872 | 0.882 | (A) BRAF Class 1 Colorectal |
| TLX1 | 10q24.31 | 1 | 2.94 | 0 | 0.00 | 0.872 | 0.882 | (A) BRAF Class 1 Colorectal |
| MAP3K14 | 17q21.31 | 1 | 0.33 | 0 | 0.00 | 0.873 | 0.882 | (A) BRAF Class 1 Colorectal |
| FAAP20 | 1p36.33 | 1 | 0.63 | 0 | 0.00 | 0.874 | 0.882 | (A) BRAF Class 1 Colorectal |
| FAAP24 | 19q13.11 | 1 | 0.63 | 0 | 0.00 | 0.874 | 0.882 | (A) BRAF Class 1 Colorectal |
| KLLN | 10q23.31 | 1 | 0.63 | 0 | 0.00 | 0.874 | 0.882 | (A) BRAF Class 1 Colorectal |
| NEIL2 | 8p23.1 | 1 | 0.63 | 0 | 0.00 | 0.874 | 0.882 | (A) BRAF Class 1 Colorectal |
| SLX1B | 16p11.2 | 1 | 0.63 | 0 | 0.00 | 0.874 | 0.882 | (A) BRAF Class 1 Colorectal |
| TAZ | Xq28 | 1 | 0.63 | 0 | 0.00 | 0.874 | 0.882 | (A) BRAF Class 1 Colorectal |
| TAL2 | 9q31.2 | 1 | 0.62 | 0 | 0.00 | 0.875 | 0.882 | (A) BRAF Class 1 Colorectal |
| GID4 | ›11.2\|17p1 | 1 | 0.68 | 0 | 0.00 | 0.886 | 0.893 | (A) BRAF Class 1 Colorectal |
| FGF9 | 13q12.11 | 2 | 2.44 | 0 | 0.00 | 0.888 | 0.893 | (A) BRAF Class 1 Colorectal |
| DDX41 | 5q35.3 | 1 | 1.15 | 0 | 0.00 | 0.897 | 0.901 | (A) BRAF Class 1 Colorectal |
| THADA | 2p21 | 1 | 2.86 | 0 | 0.00 | 0.897 | 0.901 | (A) BRAF Class 1 Colorectal |
| CD22 | 19q13.12 | 1 | 2.38 | 0 | 0.00 | 0.913 | 0.915 | (A) BRAF Class 1 Colorectal |
| FGF8 | 10q24.32 | 1 | 1.22 | 0 | 0.00 | 0.943 | 0.943 | (A) BRAF Class 1 Colorectal |
| TSPAN31 | 12q14.1 | 1 | 1.22 | 0 | 0.00 | 0.943 | 0.943 | (A) BRAF Class 1 Colorectal |

#### Supplementary Table 14

| Gene | Cytoband | Class 1 Occurrence | Class 1 % | Class 3 Occurrence | Class 3 % | SumClass | pval | qval | Enriched In |
|---|---|---|---|---|---|---|---|---|---|
| BCORL1 | Xq26.1 | 76 | 19.54 | 3 | 5.00 | 79 | 0.00236 | 0.0438 | (A) BRAF Class 1 Colorectal |
| BRCA2 | 13q13.1 | 100 | 16.53 | 6 | 5.88 | 106 | 0.00219 | 0.0415 | (A) BRAF Class 1 Colorectal |
| HLA-A | 6p22.1 | 37 | 12.80 | 0 | 0.00 | 37 | 0.00212 | 0.0411 | (A) BRAF Class 1 Colorectal |
| POLD1 | 19q13.33 | 79 | 15.11 | 3 | 3.70 | 82 | 0.00188 | 0.0372 | (A) BRAF Class 1 Colorectal |
| TGFBR2 | 3p24.1 | 92 | 25.56 | 6 | 9.23 | 98 | 0.00185 | 0.0372 | (A) BRAF Class 1 Colorectal |
| GLI1 | 12q13.3 | 66 | 11.76 | 2 | 2.17 | 68 | 0.00162 | 0.0337 | (A) BRAF Class 1 Colorectal |
| IGF1R | 15q26.3 | 53 | 8.65 | 1 | 0.95 | 54 | 0.00152 | 0.0325 | (A) BRAF Class 1 Colorectal |
| DNMT3A | 2p23.3 | 53 | 8.77 | 1 | 0.96 | 54 | 0.00146 | 0.032 | (A) BRAF Class 1 Colorectal |
| KIT | 4q12 | 44 | 5.25 | 0 | 0.00 | 44 | 0.00125 | 0.0293 | (A) BRAF Class 1 Colorectal |
| BRCA1 | 17q21.31 | 54 | 8.93 | 1 | 0.96 | 55 | 0.00126 | 0.0293 | (A) BRAF Class 1 Colorectal |
| ATRX | Xq21.1 | 87 | 14.38 | 4 | 3.92 | 91 | 0.00117 | 0.0286 | (A) BRAF Class 1 Colorectal |
| DOT1L | 19p13.3 | 39 | 10.83 | 0 | 0.00 | 39 | 0.0011 | 0.0279 | (A) BRAF Class 1 Colorectal |
| PGR | 11q22.1 | 44 | 16.24 | 0 | 0.00 | 44 | 0.00105 | 0.0273 | (A) BRAF Class 1 Colorectal |
| AKT1 | 14q32.33 | 72 | 8.59 | 2 | 1.49 | 74 | 0.00103 | 0.0273 | (A) BRAF Class 1 Colorectal |
| NOTCH3 | 19p13.12 | 111 | 20.26 | 6 | 6.82 | 117 | 0.00092 | 0.0257 | (A) BRAF Class 1 Colorectal |
| BRD4 | 19p13.12 | 59 | 10.00 | 1 | 1.02 | 60 | 0.00076 | 0.0218 | (A) BRAF Class 1 Colorectal |
| SPEN | ɔ36.21-p36.1 | 64 | 17.78 | 2 | 3.08 | 66 | 0.00074 | 0.0218 | (A) BRAF Class 1 Colorectal |
| TET1 | 10q21.3 | 64 | 13.25 | 1 | 1.35 | 65 | 0.00063 | 0.0195 | (A) BRAF Class 1 Colorectal |
| NOTCH1 | 9q34.3 | 96 | 11.85 | 4 | 3.03 | 100 | 0.00062 | 0.0195 | (A) BRAF Class 1 Colorectal |
| CASP8 | 2q33.1 | 74 | 14.40 | 2 | 2.35 | 76 | 0.00045 | 0.0151 | (A) BRAF Class 1 Colorectal |
| BLM | 15q26.1 | 68 | 11.60 | 1 | 1.01 | 69 | 0.00016 | 0.00562 | (A) BRAF Class 1 Colorectal |

| | | | | | | | | | |
|---|---|---|---|---|---|---|---|---|---|
| HNF1A | 12q24.31 | 71 | 9.11 | 1 | 0.79 | 72 | 0.00016 | 0.00562 | (A) BRAF Class 1 Colorectal |
| FLT4 | 5q35.3 | 81 | 13.82 | 2 | 2.02 | 83 | 0.00014 | 0.00524 | (A) BRAF Class 1 Colorectal |
| PTCH1 | 9q22.32 | 82 | 13.55 | 2 | 1.96 | 84 | 0.00012 | 0.00493 | (A) BRAF Class 1 Colorectal |
| NRAS | 1p13.2 | 13 | 1.55 | 11 | 8.21 | 24 | 0.00012 | 0.00488 | (B) BRAF Class 3 Colorectal |
| AR | Xq12 | 70 | 11.57 | 1 | 0.96 | 71 | 9.9E-05 | 0.00451 | (A) BRAF Class 1 Colorectal |
| SETD2 | 3p21.31 | 71 | 11.74 | 1 | 0.98 | 72 | 0.0001 | 0.00451 | (A) BRAF Class 1 Colorectal |
| EP300 | 22q13.2 | 94 | 15.77 | 3 | 2.97 | 97 | 1E-04 | 0.00451 | (A) BRAF Class 1 Colorectal |
| CIC | 19q13.2 | 86 | 16.54 | 2 | 2.27 | 88 | 5.6E-05 | 0.00293 | (A) BRAF Class 1 Colorectal |
| CREBBP | 16p13.3 | 110 | 18.18 | 4 | 3.92 | 114 | 4.7E-05 | 0.00261 | (A) BRAF Class 1 Colorectal |
| PTPRD | 9p24.1-p23 | 71 | 17.93 | 1 | 1.35 | 72 | 2.4E-05 | 0.00167 | (A) BRAF Class 1 Colorectal |
| PTPRS | 19p13.3 | 78 | 26.99 | 1 | 2.17 | 79 | 2.7E-05 | 0.00167 | (A) BRAF Class 1 Colorectal |
| POLE | 12q24.33 | 84 | 15.25 | 1 | 1.16 | 85 | 2.8E-05 | 0.00167 | (A) BRAF Class 1 Colorectal |
| ZFHX3 | 6q22.2-q22. | 85 | 30.80 | 2 | 4.26 | 87 | 2.4E-05 | 0.00167 | (A) BRAF Class 1 Colorectal |
| ARID1B | 6q25.3 | 106 | 18.31 | 3 | 3.13 | 109 | 2.1E-05 | 0.00167 | (A) BRAF Class 1 Colorectal |
| KMT2A | 11q23.3 | 106 | 17.55 | 3 | 2.88 | 109 | 1.3E-05 | 0.0012 | (A) BRAF Class 1 Colorectal |
| MSH3 | 5q14.1 | 76 | 29.69 | 0 | 0.00 | 76 | 8.3E-06 | 0.00087 | (A) BRAF Class 1 Colorectal |
| KMT2D | 12q13.12 | 177 | 29.70 | 10 | 9.90 | 187 | 6.9E-06 | 0.00082 | (A) BRAF Class 1 Colorectal |
| MSH6 | 2p16.3 | 88 | 14.06 | 0 | 0.00 | 88 | 5.2E-07 | 7.2E-05 | (A) BRAF Class 1 Colorectal |
| BCOR | Xp11.4 | 89 | 14.93 | 0 | 0.00 | 89 | 3.1E-07 | 5.2E-05 | (A) BRAF Class 1 Colorectal |
| FAT1 | 4q35.2 | 113 | 22.47 | 1 | 1.20 | 114 | 8.4E-08 | 1.7E-05 | (A) BRAF Class 1 Colorectal |
| KRAS | 12p12.1 | 15 | 1.79 | 23 | 17.16 | 38 | 7.3E-12 | 2E-09 | (B) BRAF Class 3 Colorectal |
| RNF43 | 17q22 | 226 | 41.09 | 4 | 4.40 | 230 | 3.3E-14 | 1.4E-11 | (A) BRAF Class 1 Colorectal |

| | | | | | | | | | |
|---|---|---|---|---|---|---|---|---|---|
| APC | 5q22.2 | 234 | 29.40 | 90 | 69.23 | 324 | 7.9E-18 | 6.6E-15 | (B) BRAF Class 3 Colorectal |

#### Supplementary Table 15

| Gene | Cytoband | Class 1 Occurrence | Class 1 % | Class 2 Occurrence | Class 2 % | pval | qval | Enriched in |
|---|---|---|---|---|---|---|---|---|
| STK11 | 19p13.3 | 6 | 2.59 | 60 | 27.52 | 4.2E-15 | 2.7E-12 | (B) BRAF Class 2 Lung |
| SETD2 | 3p21.31 | 71 | 39.66 | 11 | 6.67 | 8.6E-14 | 2.8E-11 | (A) BRAF Class 1 Lung |
| KEAP1 | 19p13.2 | 3 | 1.80 | 33 | 21.29 | 6.7E-09 | 1.4E-06 | (B) BRAF Class 2 Lung |
| TP53 | 17p13.1 | 74 | 31.36 | 114 | 50.22 | 2.6E-05 | 0.00414 | (B) BRAF Class 2 Lung |
| EPHA3 | 3p11.1 | 3 | 2.38 | 20 | 15.15 | 0.00021 | 0.0269 | (B) BRAF Class 2 Lung |
| ERBB4 | 2q34 | 2 | 0.88 | 17 | 7.46 | 0.00029 | 0.0306 | (B) BRAF Class 2 Lung |
| ATR | 3q23 | 6 | 3.97 | 20 | 14.71 | 0.00136 | 0.125 | (B) BRAF Class 2 Lung |
| NF1 | 17q11.2 | 5 | 2.78 | 19 | 10.73 | 0.00218 | 0.16 | (B) BRAF Class 2 Lung |
| KRAS | 12p12.1 | 9 | 3.81 | 26 | 10.92 | 0.00235 | 0.16 | (B) BRAF Class 2 Lung |
| PTPRT | 0q12-q13.1 | 2 | 2.41 | 14 | 15.38 | 0.00248 | 0.16 | (B) BRAF Class 2 Lung |
| KAT6B | 10q22.2 | 0 | 0.00 | 6 | 15.38 | 0.00375 | 0.219 | (B) BRAF Class 2 Lung |
| BRIP1 | 17q23.2 | 0 | 0.00 | 7 | 4.27 | 0.00545 | 0.28 | (B) BRAF Class 2 Lung |
| GLI2 | 2q14.2 | 2 | 2.38 | 10 | 14.71 | 0.00565 | 0.28 | (B) BRAF Class 2 Lung |
| RAF1 | 3p25.2 | 0 | 0.00 | 7 | 3.85 | 0.00695 | 0.319 | (B) BRAF Class 2 Lung |
| DIS3L2 | 2q37.1 | 0 | 0.00 | 5 | 13.89 | 0.00858 | 0.368 | (B) BRAF Class 2 Lung |
| EPHA5 | q13.1-q13.[illegible] | 3 | 2.46 | 13 | 10.24 | 0.0108 | 0.435 | (B) BRAF Class 2 Lung |
| SMARCA4 | 19p13.2 | 11 | 6.15 | 23 | 13.94 | 0.0122 | 0.444 | (B) BRAF Class 2 Lung |
| ETV6 | 12p13.2 | 0 | 0.00 | 6 | 3.66 | 0.0124 | 0.444 | (B) BRAF Class 2 Lung |
| TERT | 5p15.33 | 10 | 5.75 | 22 | 13.33 | 0.0134 | 0.452 | (B) BRAF Class 2 Lung |
| PGR | 11q22.1 | 0 | 0.00 | 6 | 8.00 | 0.0147 | 0.461 | (B) BRAF Class 2 Lung |
| FGFR2 | 10q26.13 | 0 | 0.00 | 6 | 2.52 | 0.0155 | 0.461 | (B) BRAF Class 2 Lung |
| NRAS | 1p13.2 | 3 | 1.27 | 12 | 5.04 | 0.0169 | 0.461 | (B) BRAF Class 2 Lung |
| HGF | 7q21.11 | 0 | 0.00 | 6 | 5.83 | 0.0174 | 0.461 | (B) BRAF Class 2 Lung |
| NOTCH2 | 1p12 | 5 | 2.82 | 14 | 8.64 | 0.0175 | 0.461 | (B) BRAF Class 2 Lung |
| PDGFRB | 5q32 | 3 | 1.68 | 11 | 6.67 | 0.0181 | 0.461 | (B) BRAF Class 2 Lung |
| ERCC6 | 10q11.23 | 0 | 0.00 | 4 | 12.50 | 0.0186 | 0.461 | (B) BRAF Class 2 Lung |
| WT1 | 11p13 | 2 | 1.11 | 9 | 5.45 | 0.0218 | 0.519 | (B) BRAF Class 2 Lung |
| AURKB | 17p13.1 | 0 | 0.00 | 5 | 3.05 | 0.0246 | 0.564 | (B) BRAF Class 2 Lung |
| FOXA1 | 14q21.1 | 1 | 0.74 | 7 | 5.60 | 0.0257 | 0.57 | (B) BRAF Class 2 Lung |
| NSD2 | 4p16.3 | 1 | 1.04 | 7 | 7.61 | 0.0281 | 0.591 | (B) BRAF Class 2 Lung |
| CDKN2A | 9p21.3 | 17 | 7.42 | 29 | 13.30 | 0.0291 | 0.591 | (B) BRAF Class 2 Lung |
| TSC1 | 9q34.13 | 3 | 1.68 | 10 | 6.06 | 0.031 | 0.591 | (B) BRAF Class 2 Lung |
| JAK3 | 19p13.11 | 2 | 0.90 | 9 | 4.04 | 0.0311 | 0.591 | (B) BRAF Class 2 Lung |
| INHBA | 7p14.1 | 0 | 0.00 | 5 | 5.32 | 0.0312 | 0.591 | (B) BRAF Class 2 Lung |
| EGFR | 7p11.2 | 18 | 7.63 | 8 | 3.36 | 0.0322 | 0.591 | (A) BRAF Class 1 Lung |
| AKT1 | 14q32.33 | 7 | 2.97 | 1 | 0.42 | 0.0331 | 0.591 | (A) BRAF Class 1 Lung |
| PDGFRA | 4q12 | 4 | 1.75 | 12 | 5.29 | 0.0351 | 0.602 | (B) BRAF Class 2 Lung |
| GATA6 | 18q11.2 | 1 | 0.91 | 6 | 6.52 | 0.0356 | 0.602 | (B) BRAF Class 2 Lung |
| BCORL1 | Xq26.1 | 1 | 0.85 | 6 | 5.88 | 0.0395 | 0.636 | (B) BRAF Class 2 Lung |
| FANCA | 16q24.3 | 2 | 1.12 | 8 | 4.85 | 0.0396 | 0.636 | (B) BRAF Class 2 Lung |
| PTPRD | 9p24.1-p23 | 5 | 4.72 | 14 | 12.07 | 0.0414 | 0.641 | (B) BRAF Class 2 Lung |
| KDR | 4q12 | 10 | 4.44 | 19 | 9.00 | 0.0425 | 0.641 | (B) BRAF Class 2 Lung |
| ASXL1 | 20q11.21 | 14 | 7.78 | 5 | 3.03 | 0.0433 | 0.641 | (A) BRAF Class 1 Lung |
| RASA1 | 5q14.3 | 1 | 0.76 | 6 | 5.08 | 0.0438 | 0.641 | (B) BRAF Class 2 Lung |
| CEBPA | 19q13.11 | 1 | 0.57 | 6 | 3.75 | 0.0466 | 0.666 | (B) BRAF Class 2 Lung |

| | | | | | | | | |
|---|---|---|---|---|---|---|---|---|
| PRKN | 6q26 | 1 | 0.58 | 6 | 3.73 | 0.0497 | 0.681 | (B) BRAF Class 2 Lung |
| FLCN | 17p11.2 | 1 | 0.57 | 6 | 3.66 | 0.05 | 0.681 | (B) BRAF Class 2 Lung |
| AKT3 | 1q43-q44 | 0 | 0.00 | 4 | 2.41 | 0.052 | 0.681 | (B) BRAF Class 2 Lung |
| RAD54B | 8q22.1 | 0 | 0.00 | 3 | 9.38 | 0.0521 | 0.681 | (B) BRAF Class 2 Lung |
| SUZ12 | 17q11.2 | 0 | 0.00 | 4 | 2.65 | 0.0537 | 0.681 | (B) BRAF Class 2 Lung |
| VHL | 3p25.3 | 0 | 0.00 | 4 | 1.90 | 0.054 | 0.681 | (B) BRAF Class 2 Lung |
| KEL | 7q34 | 0 | 0.00 | 4 | 12.90 | 0.0603 | 0.685 | (B) BRAF Class 2 Lung |
| RUNX1 | 21q22.12 | 2 | 1.12 | 7 | 4.35 | 0.065 | 0.685 | (B) BRAF Class 2 Lung |
| EPHB1 | 3q22.2 | 0 | 0.00 | 4 | 3.88 | 0.067 | 0.685 | (B) BRAF Class 2 Lung |
| PREX2 | 8q13.2 | 3 | 4.62 | 10 | 13.33 | 0.0672 | 0.685 | (B) BRAF Class 2 Lung |
| RBM10 | Xp11.3 | 5 | 3.45 | 11 | 8.33 | 0.0686 | 0.685 | (B) BRAF Class 2 Lung |
| RAD51 | 15q15.1 | 4 | 2.68 | 0 | 0.00 | 0.0754 | 0.685 | (A) BRAF Class 1 Lung |
| MYC | 8q24.21 | 4 | 2.22 | 10 | 5.65 | 0.0805 | 0.685 | (B) BRAF Class 2 Lung |
| RICTOR | 5p13.1 | 3 | 2.00 | 8 | 5.88 | 0.0806 | 0.685 | (B) BRAF Class 2 Lung |
| POT1 | 7q31.33 | 3 | 4.92 | 7 | 14.58 | 0.0809 | 0.685 | (B) BRAF Class 2 Lung |
| TSHR | 14q31.1 | 1 | 0.65 | 5 | 3.68 | 0.0819 | 0.685 | (B) BRAF Class 2 Lung |
| NTRK3 | 15q25.3 | 6 | 3.35 | 12 | 7.27 | 0.082 | 0.685 | (B) BRAF Class 2 Lung |
| SPTA1 | 1q23.1 | 4 | 12.90 | 10 | 30.30 | 0.0829 | 0.685 | (B) BRAF Class 2 Lung |
| RHBDF2 | 17q25.1 | 5 | 9.62 | 0 | 0.00 | 0.0842 | 0.685 | (A) BRAF Class 1 Lung |
| FANCL | 2p16.1 | 0 | 0.00 | 3 | 4.11 | 0.0878 | 0.685 | (B) BRAF Class 2 Lung |
| ERCC5 | 13q33.1 | 1 | 0.63 | 5 | 3.40 | 0.0886 | 0.685 | (B) BRAF Class 2 Lung |
| ROS1 | 6q22.1 | 4 | 2.19 | 10 | 5.41 | 0.0889 | 0.685 | (B) BRAF Class 2 Lung |
| BAP1 | 3p21.1 | 1 | 0.56 | 5 | 3.03 | 0.0895 | 0.685 | (B) BRAF Class 2 Lung |
| KIT | 4q12 | 2 | 0.85 | 7 | 2.94 | 0.0898 | 0.685 | (B) BRAF Class 2 Lung |
| CBLB | 3q13.11 | 0 | 0.00 | 3 | 4.29 | 0.0917 | 0.685 | (B) BRAF Class 2 Lung |
| ERBB2 | 17q12 | 2 | 0.86 | 7 | 2.94 | 0.093 | 0.685 | (B) BRAF Class 2 Lung |
| STAT6 | 12q13.3 | 0 | 0.00 | 3 | 4.17 | 0.0943 | 0.685 | (B) BRAF Class 2 Lung |
| EZH2 | 7q36.1 | 1 | 0.46 | 5 | 2.45 | 0.0949 | 0.685 | (B) BRAF Class 2 Lung |
| NKX2-1 | 14q13.3 | 4 | 2.25 | 9 | 5.49 | 0.0995 | 0.685 | (B) BRAF Class 2 Lung |
| SLX4 | 16p13.3 | 2 | 1.94 | 6 | 6.59 | 0.103 | 0.685 | (B) BRAF Class 2 Lung |
| MYB | 6q23.3 | 0 | 0.00 | 3 | 2.94 | 0.104 | 0.685 | (B) BRAF Class 2 Lung |
| SMAD4 | 18q21.2 | 17 | 7.33 | 9 | 4.13 | 0.105 | 0.685 | (A) BRAF Class 1 Lung |
| CDK12 | 17q12 | 0 | 0.00 | 3 | 2.21 | 0.105 | 0.685 | (B) BRAF Class 2 Lung |
| TET1 | 10q21.3 | 6 | 4.55 | 11 | 9.32 | 0.106 | 0.685 | (B) BRAF Class 2 Lung |
| LATS2 | 13q12.11 | 1 | 1.22 | 5 | 6.02 | 0.108 | 0.685 | (B) BRAF Class 2 Lung |
| CXCR4 | 2q22.1 | 0 | 0.00 | 3 | 2.44 | 0.108 | 0.685 | (B) BRAF Class 2 Lung |
| MDM4 | 1q32.1 | 0 | 0.00 | 3 | 1.82 | 0.109 | 0.685 | (B) BRAF Class 2 Lung |
| PIK3C2G | 12p12.3 | 1 | 1.15 | 5 | 5.62 | 0.111 | 0.685 | (B) BRAF Class 2 Lung |
| MAGI2 | 7q21.11 | 0 | 0.00 | 3 | 11.54 | 0.111 | 0.685 | (B) BRAF Class 2 Lung |
| RUNX1T1 | 8q21.3 | 1 | 1.14 | 4 | 5.88 | 0.114 | 0.685 | (B) BRAF Class 2 Lung |
| PPARG | 3p25.2 | 0 | 0.00 | 3 | 2.68 | 0.114 | 0.685 | (B) BRAF Class 2 Lung |
| TET2 | 4q24 | 10 | 5.56 | 4 | 2.42 | 0.114 | 0.685 | (A) BRAF Class 1 Lung |
| MET | 7q31.2 | 3 | 1.28 | 8 | 3.38 | 0.115 | 0.685 | (B) BRAF Class 2 Lung |
| GATA3 | 10p14 | 2 | 1.12 | 6 | 3.66 | 0.117 | 0.685 | (B) BRAF Class 2 Lung |
| RAD21 | 8q24.11 | 5 | 3.05 | 1 | 0.65 | 0.121 | 0.685 | (A) BRAF Class 1 Lung |
| ATRX | Xq21.1 | 6 | 3.35 | 11 | 6.67 | 0.121 | 0.685 | (B) BRAF Class 2 Lung |
| BRAF | 7q34 | 233 | 98.73 | 238 | 100.00 | 0.123 | 0.685 | (B) BRAF Class 2 Lung |
| PAK5 | 20p12.2 | 0 | 0.00 | 3 | 3.57 | 0.123 | 0.685 | (B) BRAF Class 2 Lung |

| | | | | | | | | |
|---|---|---|---|---|---|---|---|---|
| ASXL2 | 2p23.3 | 1 | 1.25 | 5 | 5.81 | 0.123 | 0.685 | (B) BRAF Class 2 Lung |
| TBX3 | 12q24.21 | 3 | 3.16 | 8 | 8.00 | 0.124 | 0.685 | (B) BRAF Class 2 Lung |
| FH | 1q43 | 5 | 2.84 | 1 | 0.61 | 0.125 | 0.685 | (A) BRAF Class 1 Lung |
| E2F3 | 6p22.3 | 0 | 0.00 | 3 | 3.80 | 0.13 | 0.685 | (B) BRAF Class 2 Lung |
| SHQ1 | 3p13 | 0 | 0.00 | 3 | 3.80 | 0.13 | 0.685 | (B) BRAF Class 2 Lung |
| VTCN1 | 1p13.1-p12 | 0 | 0.00 | 3 | 3.80 | 0.13 | 0.685 | (B) BRAF Class 2 Lung |
| IRF4 | 6p25.3 | 0 | 0.00 | 3 | 2.91 | 0.133 | 0.685 | (B) BRAF Class 2 Lung |
| LRP1B | q22.1-q22.: | 6 | 16.67 | 11 | 30.56 | 0.133 | 0.685 | (B) BRAF Class 2 Lung |
| FGF4 | 11q13.3 | 0 | 0.00 | 3 | 2.91 | 0.135 | 0.685 | (B) BRAF Class 2 Lung |
| FGF23 | 12p13.32 | 0 | 0.00 | 3 | 7.32 | 0.135 | 0.685 | (B) BRAF Class 2 Lung |
| ARID1B | 6q25.3 | 3 | 1.76 | 7 | 4.49 | 0.135 | 0.685 | (B) BRAF Class 2 Lung |
| AXL | 19q13.2 | 3 | 1.68 | 7 | 4.24 | 0.137 | 0.685 | (B) BRAF Class 2 Lung |
| PRPF40B | 12q13.12 | 0 | 0.00 | 3 | 9.09 | 0.137 | 0.685 | (B) BRAF Class 2 Lung |
| ERCC2 | 19q13.32 | 3 | 1.73 | 7 | 4.38 | 0.138 | 0.685 | (B) BRAF Class 2 Lung |
| SDHA | 5p15.33 | 3 | 1.85 | 7 | 4.67 | 0.138 | 0.685 | (B) BRAF Class 2 Lung |
| FAT1 | 4q35.2 | 14 | 9.86 | 19 | 14.96 | 0.139 | 0.685 | (B) BRAF Class 2 Lung |
| MEN1 | 11q13.1 | 3 | 1.69 | 0 | 0.00 | 0.14 | 0.685 | (A) BRAF Class 1 Lung |
| XRCC3 | 14q32.33 | 4 | 6.67 | 0 | 0.00 | 0.141 | 0.685 | (A) BRAF Class 1 Lung |
| DCLRE1C | 10p13 | 0 | 0.00 | 2 | 6.25 | 0.142 | 0.685 | (B) BRAF Class 2 Lung |
| EXO1 | 1q43 | 0 | 0.00 | 2 | 6.25 | 0.142 | 0.685 | (B) BRAF Class 2 Lung |
| GBA | 1q22 | 0 | 0.00 | 2 | 6.25 | 0.142 | 0.685 | (B) BRAF Class 2 Lung |
| TDG | 12q23.3 | 0 | 0.00 | 2 | 6.25 | 0.142 | 0.685 | (B) BRAF Class 2 Lung |
| U2AF1 | 21q22.3 | 3 | 1.70 | 7 | 4.24 | 0.143 | 0.685 | (B) BRAF Class 2 Lung |
| SMC3 | 10q25.2 | 1 | 0.97 | 4 | 4.44 | 0.145 | 0.685 | (B) BRAF Class 2 Lung |
| SMC1A | Xp11.22 | 0 | 0.00 | 3 | 5.17 | 0.147 | 0.685 | (B) BRAF Class 2 Lung |
| ZNF217 | 20q13.2 | 3 | 2.54 | 0 | 0.00 | 0.152 | 0.685 | (A) BRAF Class 1 Lung |
| GRIN2A | 16p13.2 | 2 | 2.13 | 6 | 6.12 | 0.153 | 0.685 | (B) BRAF Class 2 Lung |
| NTRK1 | 1q23.1 | 4 | 2.16 | 8 | 4.68 | 0.154 | 0.685 | (B) BRAF Class 2 Lung |
| CHEK1 | 11q24.2 | 1 | 0.66 | 4 | 2.94 | 0.154 | 0.685 | (B) BRAF Class 2 Lung |
| CTCF | 16q22.1 | 1 | 0.67 | 4 | 2.96 | 0.156 | 0.685 | (B) BRAF Class 2 Lung |
| FANCE | 6p21.31 | 3 | 2.61 | 0 | 0.00 | 0.158 | 0.685 | (A) BRAF Class 1 Lung |
| CIITA | 16p13.13 | 3 | 3.66 | 6 | 8.96 | 0.158 | 0.685 | (B) BRAF Class 2 Lung |
| SETBP1 | 18q12.3 | 8 | 7.69 | 12 | 13.04 | 0.159 | 0.685 | (B) BRAF Class 2 Lung |
| TNFAIP3 | 6q23.3 | 1 | 0.56 | 4 | 2.44 | 0.161 | 0.685 | (B) BRAF Class 2 Lung |
| KLF4 | 9q31.2 | 1 | 0.75 | 4 | 3.23 | 0.162 | 0.685 | (B) BRAF Class 2 Lung |
| CBFA2T3 | 16q24.3 | 1 | 1.89 | 3 | 8.82 | 0.163 | 0.685 | (B) BRAF Class 2 Lung |
| AGO2 | 8q24.3 | 0 | 0.00 | 3 | 7.32 | 0.164 | 0.685 | (B) BRAF Class 2 Lung |
| NF2 | 22q12.2 | 5 | 2.78 | 9 | 5.42 | 0.165 | 0.685 | (B) BRAF Class 2 Lung |
| BRD3 | 9q34.2 | 0 | 0.00 | 2 | 5.13 | 0.166 | 0.685 | (B) BRAF Class 2 Lung |
| CYLD | 16q12.1 | 1 | 0.68 | 4 | 2.90 | 0.166 | 0.685 | (B) BRAF Class 2 Lung |
| ARID2 | 12q12 | 7 | 4.00 | 11 | 6.92 | 0.174 | 0.685 | (B) BRAF Class 2 Lung |
| CDKN2B | 9p21.3 | 7 | 3.93 | 11 | 6.75 | 0.179 | 0.685 | (B) BRAF Class 2 Lung |
| GALNT12 | 9q22.33 | 1 | 1.67 | 3 | 7.32 | 0.181 | 0.685 | (B) BRAF Class 2 Lung |
| RPTOR | 17q25.3 | 2 | 1.34 | 5 | 3.70 | 0.185 | 0.685 | (B) BRAF Class 2 Lung |
| ATM | 11q22.3 | 16 | 7.11 | 21 | 9.95 | 0.186 | 0.685 | (B) BRAF Class 2 Lung |
| BCL11B | 14q32.2 | 2 | 3.57 | 4 | 10.26 | 0.187 | 0.685 | (B) BRAF Class 2 Lung |
| LZTR1 | 21\|22q11.1 | 1 | 2.94 | 4 | 11.43 | 0.187 | 0.685 | (B) BRAF Class 2 Lung |
| RANBP2 | 2q13 | 2 | 7.41 | 5 | 19.23 | 0.194 | 0.685 | (B) BRAF Class 2 Lung |

| | | | | | | | | |
|---|---|---|---|---|---|---|---|---|
| IRS1 | 2q36.3 | 1 | 1.32 | 4 | 5.06 | 0.196 | 0.685 | (B) BRAF Class 2 Lung |
| RSPO2 | 8q23.1 | 0 | 0.00 | 2 | 3.77 | 0.196 | 0.685 | (B) BRAF Class 2 Lung |
| ARHGAP35 | 19q13.32 | 5 | 8.77 | 1 | 2.44 | 0.198 | 0.685 | (A) BRAF Class 1 Lung |
| DDB2 | 11p11.2 | 0 | 0.00 | 2 | 2.94 | 0.199 | 0.685 | (B) BRAF Class 2 Lung |
| DDX31 | 9q34.13 | 0 | 0.00 | 3 | 75.00 | 0.2 | 0.685 | (B) BRAF Class 2 Lung |
| IKBKE | 1q32.1 | 1 | 1.02 | 4 | 3.88 | 0.2 | 0.685 | (B) BRAF Class 2 Lung |
| FGF19 | 11q13.3 | 1 | 1.02 | 4 | 3.85 | 0.203 | 0.685 | (B) BRAF Class 2 Lung |
| ETV5 | 3q27.2 | 0 | 0.00 | 2 | 2.11 | 0.211 | 0.685 | (B) BRAF Class 2 Lung |
| BRCA2 | 13q13.1 | 11 | 6.08 | 6 | 3.64 | 0.213 | 0.685 | (A) BRAF Class 1 Lung |
| FUS | 16p11.2 | 0 | 0.00 | 2 | 3.03 | 0.214 | 0.685 | (B) BRAF Class 2 Lung |
| MLH3 | 14q24.3 | 2 | 3.45 | 4 | 9.09 | 0.219 | 0.685 | (B) BRAF Class 2 Lung |
| TMEM127 | 2q11.2 | 0 | 0.00 | 2 | 1.74 | 0.219 | 0.685 | (B) BRAF Class 2 Lung |
| GRM3 | 21.11-q21. | 2 | 5.88 | 0 | 0.00 | 0.219 | 0.685 | (A) BRAF Class 1 Lung |
| PHF6 | Xq26.2 | 0 | 0.00 | 2 | 2.11 | 0.22 | 0.685 | (B) BRAF Class 2 Lung |
| MAX | 14q23.3 | 4 | 3.03 | 1 | 0.85 | 0.222 | 0.685 | (A) BRAF Class 1 Lung |
| AR | Xq12 | 7 | 3.91 | 11 | 6.25 | 0.223 | 0.685 | (B) BRAF Class 2 Lung |
| ERG | 21q22.2 | 0 | 0.00 | 2 | 1.48 | 0.223 | 0.685 | (B) BRAF Class 2 Lung |
| JAK2 | 9p24.1 | 2 | 0.90 | 5 | 2.25 | 0.225 | 0.685 | (B) BRAF Class 2 Lung |
| POLE | 12q24.33 | 4 | 2.61 | 7 | 4.96 | 0.226 | 0.685 | (B) BRAF Class 2 Lung |
| PALB2 | 16p12.2 | 4 | 2.23 | 7 | 4.24 | 0.227 | 0.685 | (B) BRAF Class 2 Lung |
| SH2B3 | 12q24.12 | 0 | 0.00 | 2 | 1.29 | 0.228 | 0.685 | (B) BRAF Class 2 Lung |
| CCND2 | 12p13.32 | 0 | 0.00 | 2 | 1.21 | 0.229 | 0.685 | (B) BRAF Class 2 Lung |
| CDK6 | 7q21.2 | 0 | 0.00 | 2 | 1.21 | 0.229 | 0.685 | (B) BRAF Class 2 Lung |
| XPO1 | 2p15 | 0 | 0.00 | 2 | 1.21 | 0.229 | 0.685 | (B) BRAF Class 2 Lung |
| MEF2B | 19p13.11 | 0 | 0.00 | 2 | 1.22 | 0.231 | 0.685 | (B) BRAF Class 2 Lung |
| ARAF | Xp11.3 | 4 | 2.17 | 7 | 4.09 | 0.231 | 0.685 | (B) BRAF Class 2 Lung |
| HABP2 | 10q25.3 | 3 | 5.77 | 0 | 0.00 | 0.232 | 0.685 | (A) BRAF Class 1 Lung |
| KCNQ1 | 1p15.5-p15. | 3 | 5.77 | 0 | 0.00 | 0.232 | 0.685 | (A) BRAF Class 1 Lung |
| MTA1 | 14q32.33 | 3 | 5.77 | 0 | 0.00 | 0.232 | 0.685 | (A) BRAF Class 1 Lung |
| BCL2 | 18q21.33 | 0 | 0.00 | 2 | 1.22 | 0.233 | 0.685 | (B) BRAF Class 2 Lung |
| CD79B | 17q23.3 | 0 | 0.00 | 2 | 1.22 | 0.233 | 0.685 | (B) BRAF Class 2 Lung |
| BCL2L1 | 20q11.21 | 0 | 0.00 | 2 | 1.23 | 0.235 | 0.685 | (B) BRAF Class 2 Lung |
| DIS3 | 13q21.33 | 5 | 2.91 | 8 | 5.06 | 0.235 | 0.685 | (B) BRAF Class 2 Lung |
| ERRFI1 | 1p36.23 | 2 | 2.20 | 0 | 0.00 | 0.235 | 0.685 | (A) BRAF Class 1 Lung |
| NPM1 | 5q35.1 | 0 | 0.00 | 2 | 0.96 | 0.236 | 0.685 | (B) BRAF Class 2 Lung |
| FLT3 | 13q12.2 | 4 | 1.77 | 7 | 3.30 | 0.237 | 0.685 | (B) BRAF Class 2 Lung |
| BRCA1 | 17q21.31 | 3 | 1.66 | 6 | 3.41 | 0.237 | 0.685 | (B) BRAF Class 2 Lung |
| FGFR4 | 5q35.2 | 3 | 1.65 | 6 | 3.37 | 0.24 | 0.685 | (B) BRAF Class 2 Lung |
| HIST1H3G | 6p22.2 | 2 | 2.70 | 0 | 0.00 | 0.245 | 0.685 | (A) BRAF Class 1 Lung |
| PAK3 | Xq23 | 0 | 0.00 | 2 | 5.56 | 0.246 | 0.685 | (B) BRAF Class 2 Lung |
| PBRM1 | 3p21.1 | 6 | 3.59 | 9 | 5.81 | 0.249 | 0.685 | (B) BRAF Class 2 Lung |
| PRKDC | 8q11.21 | 10 | 8.62 | 12 | 12.37 | 0.251 | 0.685 | (B) BRAF Class 2 Lung |
| RAB35 | 12q24.23 | 0 | 0.00 | 2 | 2.67 | 0.252 | 0.685 | (B) BRAF Class 2 Lung |
| SOX10 | 22q13.1 | 0 | 0.00 | 2 | 5.56 | 0.254 | 0.685 | (B) BRAF Class 2 Lung |
| NUP93 | 16q13 | 0 | 0.00 | 2 | 2.13 | 0.254 | 0.685 | (B) BRAF Class 2 Lung |
| HSD3B1 | 1p12 | 0 | 0.00 | 2 | 6.45 | 0.254 | 0.685 | (B) BRAF Class 2 Lung |
| PRAME | 22q11.22 | 0 | 0.00 | 2 | 6.90 | 0.254 | 0.685 | (B) BRAF Class 2 Lung |
| PRKCZ | 1p36.33 | 0 | 0.00 | 2 | 6.90 | 0.254 | 0.685 | (B) BRAF Class 2 Lung |

| | | | | | | | | |
|---|---|---|---|---|---|---|---|---|
| BCOR | Xp11.4 | 7 | 3.95 | 10 | 6.10 | 0.255 | 0.685 | (B) BRAF Class 2 Lung |
| FANCG | 9p13.3 | 1 | 0.84 | 3 | 2.94 | 0.255 | 0.685 | (B) BRAF Class 2 Lung |
| PIK3R2 | 19p13.11 | 0 | 0.00 | 2 | 2.04 | 0.257 | 0.685 | (B) BRAF Class 2 Lung |
| ABCB11 | 2q31.1 | 4 | 7.41 | 5 | 13.89 | 0.257 | 0.685 | (B) BRAF Class 2 Lung |
| DNMT3B | 20q11.21 | 0 | 0.00 | 2 | 2.53 | 0.258 | 0.685 | (B) BRAF Class 2 Lung |
| STK40 | 1p34.3 | 0 | 0.00 | 2 | 2.53 | 0.258 | 0.685 | (B) BRAF Class 2 Lung |
| TGFBR1 | 9q22.33 | 0 | 0.00 | 2 | 2.53 | 0.258 | 0.685 | (B) BRAF Class 2 Lung |
| YES1 | 18p11.32 | 0 | 0.00 | 2 | 2.53 | 0.258 | 0.685 | (B) BRAF Class 2 Lung |
| GNA11 | 19p13.3 | 0 | 0.00 | 2 | 0.84 | 0.259 | 0.685 | (B) BRAF Class 2 Lung |
| KDM6A | Xp11.3 | 5 | 2.81 | 2 | 1.22 | 0.259 | 0.685 | (A) BRAF Class 1 Lung |
| ARFRP1 | 20q13.33 | 0 | 0.00 | 2 | 5.13 | 0.26 | 0.685 | (B) BRAF Class 2 Lung |
| TNFRSF14 | 1p36.32 | 0 | 0.00 | 2 | 1.94 | 0.261 | 0.685 | (B) BRAF Class 2 Lung |
| NRG1 | 8p12 | 4 | 5.56 | 1 | 1.75 | 0.264 | 0.685 | (A) BRAF Class 1 Lung |
| GSK3B | 3q13.33 | 0 | 0.00 | 2 | 1.94 | 0.264 | 0.685 | (B) BRAF Class 2 Lung |
| NCOR1 | l7p12-p11.: | 0 | 0.00 | 2 | 2.11 | 0.265 | 0.685 | (B) BRAF Class 2 Lung |
| GATA2 | 3q21.3 | 1 | 0.68 | 3 | 2.29 | 0.267 | 0.685 | (B) BRAF Class 2 Lung |
| ARID5B | 10q21.2 | 0 | 0.00 | 2 | 2.33 | 0.267 | 0.685 | (B) BRAF Class 2 Lung |
| FLI1 | 11q24.3 | 0 | 0.00 | 2 | 9.09 | 0.268 | 0.685 | (B) BRAF Class 2 Lung |
| TCF7L1 | 2p11.2 | 0 | 0.00 | 2 | 5.88 | 0.27 | 0.685 | (B) BRAF Class 2 Lung |
| LMO1 | 11p15.4 | 1 | 0.59 | 3 | 1.99 | 0.27 | 0.685 | (B) BRAF Class 2 Lung |
| PRPF8 | 17p13.3 | 3 | 10.00 | 1 | 3.03 | 0.271 | 0.685 | (A) BRAF Class 1 Lung |
| MRE11 | 11q21 | 1 | 0.66 | 3 | 2.21 | 0.273 | 0.685 | (B) BRAF Class 2 Lung |
| MYD88 | 3p22.2 | 2 | 1.11 | 0 | 0.00 | 0.273 | 0.685 | (A) BRAF Class 1 Lung |
| RIT1 | 1q22 | 2 | 1.44 | 0 | 0.00 | 0.274 | 0.685 | (A) BRAF Class 1 Lung |
| TCF3 | 19p13.3 | 1 | 0.63 | 3 | 2.10 | 0.275 | 0.685 | (B) BRAF Class 2 Lung |
| ACVR1B | 12q13.13 | 0 | 0.00 | 2 | 5.26 | 0.275 | 0.685 | (B) BRAF Class 2 Lung |
| CRLF2 | 2.33 and Yp | 1 | 0.58 | 3 | 1.92 | 0.276 | 0.685 | (B) BRAF Class 2 Lung |
| DDR2 | 1q23.3 | 4 | 2.12 | 7 | 3.66 | 0.277 | 0.685 | (B) BRAF Class 2 Lung |
| VEGFA | 6p21.1 | 2 | 1.37 | 0 | 0.00 | 0.279 | 0.685 | (A) BRAF Class 1 Lung |
| TRAF7 | 16p13.3 | 1 | 0.75 | 3 | 2.42 | 0.282 | 0.685 | (B) BRAF Class 2 Lung |
| TENT5C | 1p12 | 1 | 0.56 | 3 | 1.83 | 0.283 | 0.685 | (B) BRAF Class 2 Lung |
| CDKN1B | 12p13.1 | 1 | 0.56 | 3 | 1.82 | 0.283 | 0.685 | (B) BRAF Class 2 Lung |
| CUL3 | 2q36.2 | 3 | 3.19 | 1 | 0.99 | 0.284 | 0.685 | (A) BRAF Class 1 Lung |
| DEPDC5 | 2q12.2-q12. | 0 | 0.00 | 2 | 10.00 | 0.285 | 0.685 | (B) BRAF Class 2 Lung |
| GPC3 | Xq26.2 | 2 | 2.33 | 0 | 0.00 | 0.287 | 0.685 | (A) BRAF Class 1 Lung |
| TP53BP1 | 15q15.3 | 2 | 2.20 | 0 | 0.00 | 0.288 | 0.685 | (A) BRAF Class 1 Lung |
| MPL | 1p34.2 | 1 | 0.45 | 3 | 1.42 | 0.293 | 0.685 | (B) BRAF Class 2 Lung |
| GATA4 | 8p23.1 | 2 | 1.82 | 0 | 0.00 | 0.295 | 0.685 | (A) BRAF Class 1 Lung |
| JAK1 | 1p31.3 | 1 | 0.66 | 3 | 2.03 | 0.301 | 0.685 | (B) BRAF Class 2 Lung |
| PPP4R2 | 3p13 | 0 | 0.00 | 2 | 4.88 | 0.304 | 0.685 | (B) BRAF Class 2 Lung |
| NOTCH3 | 19p13.12 | 3 | 2.03 | 5 | 3.76 | 0.304 | 0.685 | (B) BRAF Class 2 Lung |
| TEK | 9p21.2 | 0 | 0.00 | 2 | 3.64 | 0.306 | 0.685 | (B) BRAF Class 2 Lung |
| MSH2 | 2p21-p16.3 | 3 | 1.69 | 5 | 3.11 | 0.311 | 0.685 | (B) BRAF Class 2 Lung |
| PMS1 | 2q32.2 | 2 | 1.22 | 4 | 2.60 | 0.313 | 0.685 | (B) BRAF Class 2 Lung |
| CTNNB1 | 3p22.1 | 11 | 4.66 | 8 | 3.36 | 0.314 | 0.685 | (A) BRAF Class 1 Lung |
| DUSP4 | 8p12 | 0 | 0.00 | 2 | 4.17 | 0.316 | 0.685 | (B) BRAF Class 2 Lung |
| CDC73 | 1q31.2 | 3 | 1.69 | 5 | 3.05 | 0.318 | 0.685 | (B) BRAF Class 2 Lung |
| KDM5A | 12p13.33 | 5 | 3.36 | 7 | 5.19 | 0.319 | 0.685 | (B) BRAF Class 2 Lung |

| | | | | | | | | |
|---|---|---|---|---|---|---|---|---|
| NFKBIZ | 3q12.3 | 2 | 2.50 | 0 | 0.00 | 0.32 | 0.685 | (A) BRAF Class 1 Lung |
| TSC2 | 16p13.3 | 13 | 7.26 | 9 | 5.45 | 0.322 | 0.685 | (A) BRAF Class 1 Lung |
| NECTIN4 | 1q23.3 | 1 | 1.92 | 2 | 6.25 | 0.323 | 0.685 | (B) BRAF Class 2 Lung |
| NR0B1 | Xp21.2 | 1 | 1.92 | 2 | 6.25 | 0.323 | 0.685 | (B) BRAF Class 2 Lung |
| MDC1 | 6p21.33 | 1 | 1.32 | 3 | 3.80 | 0.325 | 0.685 | (B) BRAF Class 2 Lung |
| FLT1 | 13q12.3 | 4 | 2.25 | 6 | 3.66 | 0.325 | 0.685 | (B) BRAF Class 2 Lung |
| IKZF1 | 7p12.2 | 4 | 2.25 | 6 | 3.66 | 0.325 | 0.685 | (B) BRAF Class 2 Lung |
| DICER1 | 14q32.13 | 5 | 2.89 | 7 | 4.40 | 0.329 | 0.685 | (B) BRAF Class 2 Lung |
| STAG2 | Xq25 | 4 | 2.26 | 6 | 3.66 | 0.329 | 0.685 | (B) BRAF Class 2 Lung |
| MTAP | 9p21.3 | 2 | 3.57 | 3 | 7.69 | 0.332 | 0.685 | (B) BRAF Class 2 Lung |
| MST1R | 3p21.31 | 1 | 1.19 | 3 | 3.37 | 0.332 | 0.685 | (B) BRAF Class 2 Lung |
| FGF3 | 11q13.3 | 1 | 1.02 | 3 | 2.88 | 0.333 | 0.685 | (B) BRAF Class 2 Lung |
| IDH2 | 15q26.1 | 4 | 1.76 | 2 | 0.87 | 0.335 | 0.685 | (A) BRAF Class 1 Lung |
| ITK | 5q33.3 | 1 | 1.89 | 2 | 5.88 | 0.337 | 0.685 | (B) BRAF Class 2 Lung |
| INPP4B | 4q31.21 | 1 | 1.06 | 3 | 2.97 | 0.338 | 0.685 | (B) BRAF Class 2 Lung |
| SRP19 | 5q22.2 | 2 | 5.00 | 0 | 0.00 | 0.342 | 0.685 | (A) BRAF Class 1 Lung |
| MYCL | 1p34.2 | 3 | 1.68 | 1 | 0.61 | 0.343 | 0.685 | (A) BRAF Class 1 Lung |
| SPEN | 36.21-p36. | 7 | 7.22 | 5 | 4.85 | 0.343 | 0.685 | (A) BRAF Class 1 Lung |
| SOX2 | 3q26.33 | 3 | 1.69 | 1 | 0.61 | 0.343 | 0.685 | (A) BRAF Class 1 Lung |
| MTOR | 1p36.22 | 10 | 5.46 | 13 | 7.03 | 0.344 | 0.685 | (B) BRAF Class 2 Lung |
| RHEB | 7q36.1 | 3 | 2.03 | 1 | 0.74 | 0.347 | 0.685 | (A) BRAF Class 1 Lung |
| ANKRD11 | 16q24.3 | 2 | 2.70 | 4 | 5.33 | 0.347 | 0.685 | (B) BRAF Class 2 Lung |
| BARD1 | 2q35 | 3 | 2.01 | 1 | 0.74 | 0.349 | 0.685 | (A) BRAF Class 1 Lung |
| FGFR3 | 4p16.3 | 5 | 2.15 | 3 | 1.26 | 0.35 | 0.685 | (A) BRAF Class 1 Lung |
| FOXL2 | 3q22.3 | 3 | 1.99 | 1 | 0.74 | 0.352 | 0.685 | (A) BRAF Class 1 Lung |
| ALOX12B | 17p13.1 | 2 | 1.74 | 4 | 3.39 | 0.354 | 0.685 | (B) BRAF Class 2 Lung |
| EML4 | 2p21 | 1 | 4.76 | 3 | 12.50 | 0.357 | 0.685 | (B) BRAF Class 2 Lung |
| POLH | 6p21.1 | 1 | 1.89 | 2 | 5.56 | 0.357 | 0.685 | (B) BRAF Class 2 Lung |
| NFE2L2 | 2q31.2 | 9 | 5.03 | 6 | 3.64 | 0.359 | 0.685 | (A) BRAF Class 1 Lung |
| NOTCH1 | 9q34.3 | 5 | 2.17 | 7 | 3.17 | 0.359 | 0.685 | (B) BRAF Class 2 Lung |
| COP1 | q25.1-q25. | 2 | 1.92 | 4 | 3.70 | 0.36 | 0.685 | (B) BRAF Class 2 Lung |
| SHH | 7q36.3 | 1 | 25.00 | 0 | 0.00 | 0.364 | 0.685 | (A) BRAF Class 1 Lung |
| TOPBP1 | 3q22.1 | 4 | 7.69 | 1 | 3.13 | 0.365 | 0.685 | (A) BRAF Class 1 Lung |
| ARHGEF12 | 11q23.3 | 2 | 3.77 | 0 | 0.00 | 0.368 | 0.685 | (A) BRAF Class 1 Lung |
| FN1 | 2q35 | 1 | 33.33 | 0 | 0.00 | 0.375 | 0.685 | (A) BRAF Class 1 Lung |
| SMO | 7q32.1 | 5 | 2.22 | 7 | 3.15 | 0.377 | 0.685 | (B) BRAF Class 2 Lung |
| TP63 | 3q28 | 2 | 2.60 | 4 | 4.82 | 0.377 | 0.685 | (B) BRAF Class 2 Lung |
| RET | 10q11.21 | 7 | 2.97 | 5 | 2.10 | 0.38 | 0.685 | (A) BRAF Class 1 Lung |
| DDB1 | 11q12.2 | 0 | 0.00 | 1 | 3.13 | 0.381 | 0.685 | (B) BRAF Class 2 Lung |
| EGLN1 | 1q42.2 | 0 | 0.00 | 1 | 3.13 | 0.381 | 0.685 | (B) BRAF Class 2 Lung |
| FAN1 | 15q13.3 | 0 | 0.00 | 1 | 3.13 | 0.381 | 0.685 | (B) BRAF Class 2 Lung |
| MCM8 | 20p12.3 | 0 | 0.00 | 1 | 3.13 | 0.381 | 0.685 | (B) BRAF Class 2 Lung |
| POLB | 8p11.21 | 0 | 0.00 | 1 | 3.13 | 0.381 | 0.685 | (B) BRAF Class 2 Lung |
| UBE2T | 1q32.1 | 0 | 0.00 | 1 | 3.13 | 0.381 | 0.685 | (B) BRAF Class 2 Lung |
| CRKL | 22q11.21 | 4 | 2.25 | 2 | 1.22 | 0.382 | 0.685 | (A) BRAF Class 1 Lung |
| ZFHX4 | 8q21.13 | 0 | 0.00 | 2 | 28.57 | 0.382 | 0.685 | (B) BRAF Class 2 Lung |
| NFKBIA | 14q13.2 | 4 | 2.29 | 2 | 1.25 | 0.385 | 0.685 | (A) BRAF Class 1 Lung |
| POLQ | 3q13.33 | 4 | 7.41 | 4 | 11.43 | 0.387 | 0.685 | (B) BRAF Class 2 Lung |

| | | | | | | | | |
|---|---|---|---|---|---|---|---|---|
| MED12 | Xq13.1 | 6 | 4.00 | 4 | 2.72 | 0.387 | 0.685 | (A) BRAF Class 1 Lung |
| FGFR1 | 8p11.23 | 4 | 1.72 | 6 | 2.52 | 0.389 | 0.685 | (B) BRAF Class 2 Lung |
| EPHA7 | 6q16.1 | 4 | 3.39 | 6 | 4.96 | 0.39 | 0.685 | (B) BRAF Class 2 Lung |
| TLX3 | 5q35.1 | 0 | 0.00 | 1 | 2.94 | 0.391 | 0.685 | (B) BRAF Class 2 Lung |
| BUB1B | 15q15.1 | 3 | 3.57 | 1 | 1.47 | 0.393 | 0.685 | (A) BRAF Class 1 Lung |
| TRAF3 | 14q32.32 | 1 | 1.72 | 2 | 4.55 | 0.397 | 0.685 | (B) BRAF Class 2 Lung |
| GNAS | 20q13.32 | 5 | 2.16 | 3 | 1.38 | 0.397 | 0.685 | (A) BRAF Class 1 Lung |
| PTEN | 10q23.31 | 6 | 2.54 | 4 | 1.76 | 0.4 | 0.685 | (A) BRAF Class 1 Lung |
| GNA14 | 9q21.2 | 0 | 0.00 | 2 | 50.00 | 0.4 | 0.685 | (B) BRAF Class 2 Lung |
| PARP1 | 1q42.12 | 4 | 4.44 | 6 | 6.38 | 0.401 | 0.685 | (B) BRAF Class 2 Lung |
| ARID1A | 1p36.11 | 21 | 11.73 | 17 | 10.30 | 0.402 | 0.685 | (A) BRAF Class 1 Lung |
| SPRED1 | 15q14 | 2 | 5.41 | 1 | 2.08 | 0.402 | 0.685 | (A) BRAF Class 1 Lung |
| RAD50 | 5q31.1 | 5 | 3.38 | 6 | 4.58 | 0.417 | 0.685 | (B) BRAF Class 2 Lung |
| DNAH9 | 17p12 | 0 | 0.00 | 1 | 20.00 | 0.417 | 0.685 | (B) BRAF Class 2 Lung |
| EXT2 | 11p11.2 | 1 | 1.19 | 2 | 2.94 | 0.421 | 0.685 | (B) BRAF Class 2 Lung |
| DNMT3A | 2p23.3 | 9 | 5.00 | 10 | 6.06 | 0.422 | 0.685 | (B) BRAF Class 2 Lung |
| NUTM1 | 15q14 | 1 | 3.85 | 0 | 0.00 | 0.426 | 0.685 | (A) BRAF Class 1 Lung |
| RSPO3 | 6q22.33 | 0 | 0.00 | 1 | 2.33 | 0.43 | 0.685 | (B) BRAF Class 2 Lung |
| BRD4 | 19p13.12 | 7 | 4.02 | 5 | 3.07 | 0.431 | 0.685 | (A) BRAF Class 1 Lung |
| FAS | 10q23.31 | 1 | 0.88 | 2 | 2.11 | 0.435 | 0.685 | (B) BRAF Class 2 Lung |
| RRAS2 | 11p15.2 | 1 | 2.70 | 0 | 0.00 | 0.435 | 0.685 | (A) BRAF Class 1 Lung |
| BCL2L12 | 19q13.33 | 0 | 0.00 | 1 | 1.56 | 0.438 | 0.685 | (B) BRAF Class 2 Lung |
| MECOM | 3q26.2 | 0 | 0.00 | 1 | 1.56 | 0.438 | 0.685 | (B) BRAF Class 2 Lung |
| CDKN1C | 11p15.4 | 0 | 0.00 | 1 | 1.54 | 0.442 | 0.685 | (B) BRAF Class 2 Lung |
| POLD1 | 19q13.33 | 6 | 4.03 | 4 | 2.99 | 0.442 | 0.685 | (A) BRAF Class 1 Lung |
| PTCH1 | 9q22.32 | 5 | 2.79 | 6 | 3.64 | 0.444 | 0.685 | (B) BRAF Class 2 Lung |
| CARM1 | 19p13.2 | 1 | 3.03 | 0 | 0.00 | 0.446 | 0.685 | (A) BRAF Class 1 Lung |
| UPF1 | 19p13.11 | 1 | 3.03 | 0 | 0.00 | 0.446 | 0.685 | (A) BRAF Class 1 Lung |
| CRTC1 | 19p13.11 | 0 | 0.00 | 1 | 1.47 | 0.447 | 0.685 | (B) BRAF Class 2 Lung |
| EXT1 | 8q24.11 | 0 | 0.00 | 1 | 1.47 | 0.447 | 0.685 | (B) BRAF Class 2 Lung |
| MAP3K1 | 5q11.2 | 5 | 2.82 | 6 | 3.66 | 0.448 | 0.685 | (B) BRAF Class 2 Lung |
| NTRK2 | 9q21.33 | 5 | 2.81 | 6 | 3.64 | 0.448 | 0.685 | (B) BRAF Class 2 Lung |
| TFE3 | Xp11.23 | 0 | 0.00 | 1 | 1.59 | 0.45 | 0.685 | (B) BRAF Class 2 Lung |
| NT5C2 | q24.32-q24. | 0 | 0.00 | 1 | 2.00 | 0.45 | 0.685 | (B) BRAF Class 2 Lung |
| SLITRK6 | 13q31.1 | 1 | 5.26 | 0 | 0.00 | 0.452 | 0.685 | (A) BRAF Class 1 Lung |
| KMT2B | 19q13.12 | 2 | 3.64 | 4 | 5.80 | 0.453 | 0.685 | (B) BRAF Class 2 Lung |
| IL7R | 5p13.2 | 6 | 4.08 | 4 | 3.08 | 0.453 | 0.685 | (A) BRAF Class 1 Lung |
| EPCAM | 2p21 | 2 | 1.43 | 3 | 2.36 | 0.454 | 0.685 | (B) BRAF Class 2 Lung |
| BLM | 15q26.1 | 2 | 1.14 | 3 | 1.89 | 0.455 | 0.685 | (B) BRAF Class 2 Lung |
| MCL1 | 1q21.2 | 3 | 1.69 | 4 | 2.44 | 0.455 | 0.685 | (B) BRAF Class 2 Lung |
| RARA | 17q21.2 | 3 | 1.69 | 4 | 2.44 | 0.455 | 0.685 | (B) BRAF Class 2 Lung |
| FANCF | 11p14.3 | 0 | 0.00 | 1 | 1.03 | 0.455 | 0.685 | (B) BRAF Class 2 Lung |
| ADGRA2 | 8p11.23 | 2 | 5.71 | 3 | 9.38 | 0.457 | 0.685 | (B) BRAF Class 2 Lung |
| FBXW7 | 4q31.3 | 5 | 2.16 | 6 | 2.75 | 0.457 | 0.685 | (B) BRAF Class 2 Lung |
| YAP1 | 11q22.1 | 1 | 0.76 | 2 | 1.69 | 0.458 | 0.685 | (B) BRAF Class 2 Lung |
| MYBL1 | 8q13.1 | 0 | 0.00 | 1 | 1.39 | 0.459 | 0.685 | (B) BRAF Class 2 Lung |
| AXIN2 | 17q24.1 | 2 | 1.50 | 3 | 2.46 | 0.459 | 0.685 | (B) BRAF Class 2 Lung |
| IGF1R | 15q26.3 | 2 | 1.10 | 3 | 1.81 | 0.459 | 0.685 | (B) BRAF Class 2 Lung |

| | | | | | | | | |
|---|---|---|---|---|---|---|---|---|
| NSD3 | 8p11.23 | 1 | 1.12 | 2 | 2.50 | 0.46 | 0.685 | (B) BRAF Class 2 Lung |
| ZFHX3 | 5q22.2-q22. | 7 | 8.97 | 6 | 7.32 | 0.462 | 0.685 | (A) BRAF Class 1 Lung |
| ETV1 | 7p21.2 | 1 | 0.60 | 2 | 1.32 | 0.462 | 0.685 | (B) BRAF Class 2 Lung |
| HIST1H3C | 6p22.2 | 0 | 0.00 | 1 | 0.90 | 0.464 | 0.685 | (B) BRAF Class 2 Lung |
| NKX3-1 | 8p21.2 | 0 | 0.00 | 1 | 0.90 | 0.464 | 0.685 | (B) BRAF Class 2 Lung |
| CREBBP | 16p13.3 | 9 | 5.03 | 7 | 4.24 | 0.466 | 0.685 | (A) BRAF Class 1 Lung |
| LYN | 8q12.1 | 1 | 1.59 | 0 | 0.00 | 0.467 | 0.685 | (A) BRAF Class 1 Lung |
| MAP2K4 | 17p12 | 1 | 0.56 | 2 | 1.23 | 0.467 | 0.685 | (B) BRAF Class 2 Lung |
| CTLA4 | 2q33.2 | 0 | 0.00 | 1 | 0.88 | 0.467 | 0.685 | (B) BRAF Class 2 Lung |
| IGF2 | 11p15.5 | 0 | 0.00 | 1 | 0.78 | 0.467 | 0.685 | (B) BRAF Class 2 Lung |
| KMT2C | 7q36.1 | 7 | 7.69 | 6 | 6.32 | 0.467 | 0.685 | (A) BRAF Class 1 Lung |
| B2M | 15q21.1 | 1 | 0.63 | 2 | 1.38 | 0.468 | 0.685 | (B) BRAF Class 2 Lung |
| NCOA3 | 20q13.12 | 2 | 2.53 | 1 | 1.16 | 0.468 | 0.685 | (A) BRAF Class 1 Lung |
| H3F3A | 1q42.12 | 1 | 0.56 | 2 | 1.22 | 0.469 | 0.685 | (B) BRAF Class 2 Lung |
| PRDM1 | 6q21 | 1 | 0.56 | 2 | 1.22 | 0.469 | 0.685 | (B) BRAF Class 2 Lung |
| SOCS1 | 16p13.13 | 1 | 0.56 | 2 | 1.22 | 0.469 | 0.685 | (B) BRAF Class 2 Lung |
| XRCC2 | 7q36.1 | 0 | 0.00 | 1 | 0.82 | 0.469 | 0.685 | (B) BRAF Class 2 Lung |
| AKT2 | 19q13.2 | 1 | 0.56 | 2 | 1.20 | 0.47 | 0.685 | (B) BRAF Class 2 Lung |
| SH2D1A | Xq25 | 0 | 0.00 | 1 | 0.86 | 0.47 | 0.685 | (B) BRAF Class 2 Lung |
| KMT2A | 11q23.3 | 9 | 5.00 | 7 | 4.24 | 0.47 | 0.685 | (A) BRAF Class 1 Lung |
| TMPRSS2 | 21q22.3 | 1 | 0.68 | 2 | 1.48 | 0.471 | 0.685 | (B) BRAF Class 2 Lung |
| DAXX | 6p21.32 | 0 | 0.00 | 1 | 0.76 | 0.471 | 0.685 | (B) BRAF Class 2 Lung |
| ZRSR2 | Xp22.2 | 1 | 0.62 | 2 | 1.33 | 0.473 | 0.685 | (B) BRAF Class 2 Lung |
| GEN1 | 2p24.2 | 4 | 5.26 | 2 | 3.45 | 0.475 | 0.685 | (A) BRAF Class 1 Lung |
| EP300 | 22q13.2 | 10 | 5.62 | 8 | 4.88 | 0.476 | 0.685 | (A) BRAF Class 1 Lung |
| PPP2R2A | 8p21.2 | 1 | 5.00 | 0 | 0.00 | 0.476 | 0.685 | (A) BRAF Class 1 Lung |
| KMT2D | 12q13.12 | 16 | 8.99 | 16 | 9.76 | 0.476 | 0.685 | (B) BRAF Class 2 Lung |
| PTPN11 | 12q24.13 | 1 | 0.43 | 2 | 0.92 | 0.477 | 0.685 | (B) BRAF Class 2 Lung |
| MYOD1 | 11p15.1 | 2 | 2.56 | 1 | 1.20 | 0.477 | 0.685 | (A) BRAF Class 1 Lung |
| CASP8 | 2q33.1 | 0 | 0.00 | 1 | 0.76 | 0.478 | 0.685 | (B) BRAF Class 2 Lung |
| FANCC | 9q22.32 | 0 | 0.00 | 1 | 0.61 | 0.48 | 0.685 | (B) BRAF Class 2 Lung |
| TRAF2 | 9q34.3 | 2 | 2.67 | 1 | 1.27 | 0.48 | 0.685 | (A) BRAF Class 1 Lung |
| SDHC | 1q23.3 | 0 | 0.00 | 1 | 0.62 | 0.481 | 0.685 | (B) BRAF Class 2 Lung |
| SRSF2 | 17q25.1 | 0 | 0.00 | 1 | 0.67 | 0.482 | 0.685 | (B) BRAF Class 2 Lung |
| SDHD | 11q23.1 | 0 | 0.00 | 1 | 0.61 | 0.482 | 0.685 | (B) BRAF Class 2 Lung |
| CDKN2C | 1p32.3 | 0 | 0.00 | 1 | 0.61 | 0.484 | 0.685 | (B) BRAF Class 2 Lung |
| SDHAF2 | 11q12.2 | 0 | 0.00 | 1 | 0.67 | 0.484 | 0.685 | (B) BRAF Class 2 Lung |
| PHOX2B | 4p13 | 0 | 0.00 | 1 | 0.65 | 0.484 | 0.685 | (B) BRAF Class 2 Lung |
| CHD4 | 12p13.31 | 1 | 3.23 | 0 | 0.00 | 0.484 | 0.685 | (A) BRAF Class 1 Lung |
| TOP2A | 17q21.2 | 1 | 3.23 | 0 | 0.00 | 0.484 | 0.685 | (A) BRAF Class 1 Lung |
| APC | 5q22.2 | 15 | 6.67 | 16 | 7.21 | 0.484 | 0.685 | (B) BRAF Class 2 Lung |
| FANCD2 | 3p25.3 | 5 | 4.42 | 3 | 3.30 | 0.485 | 0.685 | (A) BRAF Class 1 Lung |
| NOTCH4 | 6p21.32 | 5 | 5.68 | 4 | 4.44 | 0.486 | 0.685 | (A) BRAF Class 1 Lung |
| TFRC | 3q29 | 1 | 5.56 | 0 | 0.00 | 0.486 | 0.685 | (A) BRAF Class 1 Lung |
| ERBB3 | 12q13.2 | 6 | 3.33 | 7 | 3.95 | 0.487 | 0.685 | (B) BRAF Class 2 Lung |
| PMAIP1 | 18q21.32 | 1 | 1.32 | 0 | 0.00 | 0.49 | 0.685 | (A) BRAF Class 1 Lung |
| CCND1 | 11q13.3 | 3 | 1.68 | 4 | 2.27 | 0.491 | 0.685 | (B) BRAF Class 2 Lung |
| CDK9 | 9q34.11 | 1 | 3.57 | 0 | 0.00 | 0.491 | 0.685 | (A) BRAF Class 1 Lung |

| | | | | | | | | |
|---|---|---|---|---|---|---|---|---|
| RPL26 | 17p13.1 | 1 | 3.57 | 0 | 0.00 | 0.491 | 0.685 | (A) BRAF Class 1 Lung |
| CCN6 | 6q21 | 1 | 2.86 | 0 | 0.00 | 0.493 | 0.685 | (A) BRAF Class 1 Lung |
| STAT4 | q32.2-q32.: | 1 | 2.86 | 0 | 0.00 | 0.493 | 0.685 | (A) BRAF Class 1 Lung |
| CDH4 | 20q13.33 | 2 | 3.85 | 2 | 6.25 | 0.493 | 0.685 | (B) BRAF Class 2 Lung |
| PAXIP1 | 7q36.2 | 2 | 3.85 | 2 | 6.25 | 0.493 | 0.685 | (B) BRAF Class 2 Lung |
| RIF1 | 2q23.3 | 2 | 3.85 | 2 | 6.25 | 0.493 | 0.685 | (B) BRAF Class 2 Lung |
| RINT1 | 7q22.3 | 2 | 3.85 | 2 | 6.25 | 0.493 | 0.685 | (B) BRAF Class 2 Lung |
| EIF4A2 | 3q27.3 | 1 | 1.33 | 0 | 0.00 | 0.493 | 0.685 | (A) BRAF Class 1 Lung |
| MAP3K13 | 3q27.2 | 1 | 1.15 | 0 | 0.00 | 0.494 | 0.685 | (A) BRAF Class 1 Lung |
| STAG1 | 3q22.3 | 1 | 2.22 | 0 | 0.00 | 0.495 | 0.685 | (A) BRAF Class 1 Lung |
| HIST1H3D | 6p22.2 | 1 | 1.35 | 0 | 0.00 | 0.497 | 0.685 | (A) BRAF Class 1 Lung |
| HIST1H3E | 6p22.2 | 1 | 1.35 | 0 | 0.00 | 0.497 | 0.685 | (A) BRAF Class 1 Lung |
| HIST1H3H | 6p22.1 | 1 | 1.35 | 0 | 0.00 | 0.497 | 0.685 | (A) BRAF Class 1 Lung |
| HIST1H3I | 6p22.1 | 1 | 1.35 | 0 | 0.00 | 0.497 | 0.685 | (A) BRAF Class 1 Lung |
| HIST1H3J | 6p22.1 | 1 | 1.35 | 0 | 0.00 | 0.497 | 0.685 | (A) BRAF Class 1 Lung |
| HIST2H3C | 1q21.2 | 1 | 1.35 | 0 | 0.00 | 0.497 | 0.685 | (A) BRAF Class 1 Lung |
| CLIP1 | 12q24.31 | 1 | 0.42 | 0 | 0.00 | 0.498 | 0.685 | (A) BRAF Class 1 Lung |
| MT-ND6 | - | 1 | 0.42 | 0 | 0.00 | 0.498 | 0.685 | (A) BRAF Class 1 Lung |
| PIK3C3 | 18q12.3 | 2 | 2.35 | 1 | 1.18 | 0.5 | 0.685 | (A) BRAF Class 1 Lung |
| FAT3 | 11q14.3 | 1 | 8.33 | 2 | 16.67 | 0.5 | 0.685 | (B) BRAF Class 2 Lung |
| FGF2 | 4q28.1 | 0 | 0.00 | 1 | 5.88 | 0.5 | 0.685 | (B) BRAF Class 2 Lung |
| NKX2-1-AS1 | 14q13.3 | 0 | 0.00 | 1 | 5.88 | 0.5 | 0.685 | (B) BRAF Class 2 Lung |
| PCDHAC2 | 5q31.3 | 0 | 0.00 | 1 | 100.00 | 0.5 | 0.685 | (B) BRAF Class 2 Lung |
| HIST2H3D | 1q21.2 | 1 | 1.37 | 0 | 0.00 | 0.5 | 0.685 | (A) BRAF Class 1 Lung |
| NSD1 | 5q35.3 | 9 | 6.12 | 7 | 5.38 | 0.5 | 0.685 | (A) BRAF Class 1 Lung |
| ADGRV1 | 5q14.3 | 0 | 0.00 | 1 | 0.42 | 0.502 | 0.685 | (B) BRAF Class 2 Lung |
| HRAS | 11p15.5 | 0 | 0.00 | 1 | 0.42 | 0.502 | 0.685 | (B) BRAF Class 2 Lung |
| LGALS17A | 19q13.2 | 0 | 0.00 | 1 | 0.42 | 0.502 | 0.685 | (B) BRAF Class 2 Lung |
| MTMR3 | 22q12.2 | 0 | 0.00 | 1 | 0.42 | 0.502 | 0.685 | (B) BRAF Class 2 Lung |
| NBPF1 | 1p36.13 | 0 | 0.00 | 1 | 0.42 | 0.502 | 0.685 | (B) BRAF Class 2 Lung |
| PGBD3 | 10q11.23 | 0 | 0.00 | 1 | 0.42 | 0.502 | 0.685 | (B) BRAF Class 2 Lung |
| SELENOI | 2p23.3 | 0 | 0.00 | 1 | 0.42 | 0.502 | 0.685 | (B) BRAF Class 2 Lung |
| TMEM132C | q24.32-q24. | 0 | 0.00 | 1 | 0.42 | 0.502 | 0.685 | (B) BRAF Class 2 Lung |
| ESR1 | q25.1-q25.: | 0 | 0.00 | 1 | 0.54 | 0.503 | 0.685 | (B) BRAF Class 2 Lung |
| MAPK3 | 16p11.2 | 0 | 0.00 | 1 | 1.33 | 0.503 | 0.685 | (B) BRAF Class 2 Lung |
| GNAQ | 9q21.2 | 1 | 0.43 | 2 | 0.84 | 0.505 | 0.685 | (B) BRAF Class 2 Lung |
| HIST3H3 | 1q42.13 | 1 | 1.35 | 2 | 2.67 | 0.505 | 0.685 | (B) BRAF Class 2 Lung |
| MAP2K1 | 15q22.31 | 3 | 1.58 | 4 | 2.08 | 0.506 | 0.685 | (B) BRAF Class 2 Lung |
| EME1 | 17q21.33 | 3 | 5.77 | 1 | 3.13 | 0.507 | 0.685 | (A) BRAF Class 1 Lung |
| CD79A | 19q13.2 | 0 | 0.00 | 1 | 1.01 | 0.508 | 0.685 | (B) BRAF Class 2 Lung |
| GATA1 | Xp11.23 | 0 | 0.00 | 1 | 1.01 | 0.508 | 0.685 | (B) BRAF Class 2 Lung |
| TOP1 | 20q12 | 0 | 0.00 | 1 | 1.01 | 0.508 | 0.685 | (B) BRAF Class 2 Lung |
| MYCN | 2p24.3 | 3 | 1.68 | 2 | 1.14 | 0.508 | 0.685 | (A) BRAF Class 1 Lung |
| GABRA6 | 5q34 | 0 | 0.00 | 1 | 3.23 | 0.508 | 0.685 | (B) BRAF Class 2 Lung |
| SYNE1 | 6q25.2 | 0 | 0.00 | 1 | 3.45 | 0.509 | 0.685 | (B) BRAF Class 2 Lung |
| BCL10 | 1p22.3 | 0 | 0.00 | 1 | 1.25 | 0.51 | 0.685 | (B) BRAF Class 2 Lung |
| IFNGR1 | 6q23.3 | 0 | 0.00 | 1 | 1.27 | 0.51 | 0.685 | (B) BRAF Class 2 Lung |
| INPP4A | 2q11.2 | 0 | 0.00 | 1 | 1.27 | 0.51 | 0.685 | (B) BRAF Class 2 Lung |

| | | | | | | | | |
|---|---|---|---|---|---|---|---|---|
| RYBP | 3p13 | 0 | 0.00 | 1 | 1.27 | 0.51 | 0.685 | (B) BRAF Class 2 Lung |
| XIAP | Xq25 | 0 | 0.00 | 1 | 1.27 | 0.51 | 0.685 | (B) BRAF Class 2 Lung |
| PDCD1LG2 | 9p24.1 | 1 | 0.68 | 0 | 0.00 | 0.51 | 0.685 | (A) BRAF Class 1 Lung |
| RAC1 | 7p22.1 | 1 | 0.63 | 0 | 0.00 | 0.511 | 0.685 | (A) BRAF Class 1 Lung |
| IDH1 | 2q34 | 3 | 1.29 | 4 | 1.68 | 0.512 | 0.685 | (B) BRAF Class 2 Lung |
| PPP6C | 9q33.3 | 0 | 0.00 | 1 | 1.20 | 0.512 | 0.685 | (B) BRAF Class 2 Lung |
| TGFBR2 | 3p24.1 | 0 | 0.00 | 1 | 0.97 | 0.512 | 0.685 | (B) BRAF Class 2 Lung |
| FYN | 6q21 | 0 | 0.00 | 1 | 1.22 | 0.512 | 0.685 | (B) BRAF Class 2 Lung |
| SRC | 20q11.23 | 0 | 0.00 | 1 | 0.56 | 0.513 | 0.685 | (B) BRAF Class 2 Lung |
| FOXP1 | 3p13 | 0 | 0.00 | 1 | 1.02 | 0.513 | 0.685 | (B) BRAF Class 2 Lung |
| CRTC2 | 1q21.3 | 1 | 3.57 | 2 | 6.90 | 0.513 | 0.685 | (B) BRAF Class 2 Lung |
| LATS1 | 6q25.1 | 1 | 1.32 | 2 | 2.53 | 0.515 | 0.685 | (B) BRAF Class 2 Lung |
| PTPRS | 19p13.3 | 1 | 1.32 | 2 | 2.53 | 0.515 | 0.685 | (B) BRAF Class 2 Lung |
| BCR | 22q11.23 | 0 | 0.00 | 1 | 2.94 | 0.515 | 0.685 | (B) BRAF Class 2 Lung |
| PIK3CD | 1p36.22 | 0 | 0.00 | 1 | 1.19 | 0.515 | 0.685 | (B) BRAF Class 2 Lung |
| NEGR1 | 1p31.1 | 1 | 1.10 | 2 | 2.11 | 0.516 | 0.685 | (B) BRAF Class 2 Lung |
| DMD | p21.2-p21. | 2 | 7.14 | 3 | 10.34 | 0.517 | 0.685 | (B) BRAF Class 2 Lung |
| GLI3 | 7p14.1 | 2 | 7.14 | 3 | 10.34 | 0.517 | 0.685 | (B) BRAF Class 2 Lung |
| PIM1 | 6p21.2 | 1 | 0.61 | 0 | 0.00 | 0.517 | 0.685 | (A) BRAF Class 1 Lung |
| SDHB | 1p36.13 | 1 | 0.57 | 0 | 0.00 | 0.518 | 0.685 | (A) BRAF Class 1 Lung |
| CDKN1A | 6p21.2 | 1 | 0.58 | 0 | 0.00 | 0.518 | 0.685 | (A) BRAF Class 1 Lung |
| ALK | p23.2-p23. | 8 | 3.43 | 9 | 3.78 | 0.518 | 0.685 | (B) BRAF Class 2 Lung |
| PRKCI | 3q26.2 | 1 | 0.70 | 0 | 0.00 | 0.518 | 0.685 | (A) BRAF Class 1 Lung |
| BMPR1A | 10q23.2 | 1 | 0.60 | 0 | 0.00 | 0.519 | 0.685 | (A) BRAF Class 1 Lung |
| BTK | Xq22.1 | 1 | 1.01 | 2 | 1.92 | 0.519 | 0.685 | (B) BRAF Class 2 Lung |
| AMER1 | Xq11.2 | 1 | 1.02 | 2 | 1.94 | 0.519 | 0.685 | (B) BRAF Class 2 Lung |
| IRS2 | 13q34 | 1 | 1.02 | 2 | 1.94 | 0.519 | 0.685 | (B) BRAF Class 2 Lung |
| BIRC3 | 11q22.2 | 1 | 1.28 | 2 | 2.44 | 0.519 | 0.685 | (B) BRAF Class 2 Lung |
| EMSY | 11q13.5 | 0 | 0.00 | 1 | 2.44 | 0.519 | 0.685 | (B) BRAF Class 2 Lung |
| FGF14 | 13q33.1 | 0 | 0.00 | 1 | 2.44 | 0.519 | 0.685 | (B) BRAF Class 2 Lung |
| PDK1 | 2q31.1 | 0 | 0.00 | 1 | 2.44 | 0.519 | 0.685 | (B) BRAF Class 2 Lung |
| CALR | 19p13.13 | 1 | 0.74 | 0 | 0.00 | 0.519 | 0.685 | (A) BRAF Class 1 Lung |
| PIK3CG | 7q22.3 | 4 | 4.21 | 5 | 5.10 | 0.52 | 0.685 | (B) BRAF Class 2 Lung |
| HNF1A | 12q24.31 | 3 | 1.36 | 2 | 0.95 | 0.52 | 0.685 | (A) BRAF Class 1 Lung |
| FUBP1 | 1p31.1 | 0 | 0.00 | 1 | 0.98 | 0.52 | 0.685 | (B) BRAF Class 2 Lung |
| SMAD2 | 18q21.1 | 1 | 0.56 | 0 | 0.00 | 0.52 | 0.685 | (A) BRAF Class 1 Lung |
| MAPK1 | 22q11.22 | 2 | 1.20 | 1 | 0.63 | 0.521 | 0.685 | (A) BRAF Class 1 Lung |
| RHOA | 3p21.31 | 1 | 0.75 | 0 | 0.00 | 0.521 | 0.685 | (A) BRAF Class 1 Lung |
| MLH1 | 3p22.2 | 3 | 1.35 | 2 | 0.94 | 0.524 | 0.685 | (A) BRAF Class 1 Lung |
| RNF43 | 17q22 | 1 | 0.67 | 0 | 0.00 | 0.524 | 0.685 | (A) BRAF Class 1 Lung |
| GREM1 | 15q13.3 | 1 | 0.73 | 0 | 0.00 | 0.525 | 0.685 | (A) BRAF Class 1 Lung |
| EPHB6 | 7q34 | 0 | 0.00 | 1 | 10.00 | 0.526 | 0.685 | (B) BRAF Class 2 Lung |
| PDCD1 | 2q37.3 | 0 | 0.00 | 1 | 1.11 | 0.526 | 0.685 | (B) BRAF Class 2 Lung |
| INSR | 19p13.2 | 2 | 2.41 | 3 | 3.41 | 0.528 | 0.685 | (B) BRAF Class 2 Lung |
| HIST1H1C | 6p22.2 | 1 | 1.30 | 2 | 2.41 | 0.528 | 0.685 | (B) BRAF Class 2 Lung |
| FGF6 | 12p13.32 | 1 | 2.63 | 2 | 4.88 | 0.529 | 0.685 | (B) BRAF Class 2 Lung |
| ZNF703 | 8p11.23 | 1 | 2.63 | 2 | 4.88 | 0.529 | 0.685 | (B) BRAF Class 2 Lung |
| STAT3 | 17q21.2 | 2 | 1.15 | 1 | 0.62 | 0.529 | 0.685 | (A) BRAF Class 1 Lung |

| | | | | | | | | |
|---|---|---|---|---|---|---|---|---|
| FANCI | 15q26.1 | 5 | 6.33 | 3 | 5.08 | 0.53 | 0.685 | (A) BRAF Class 1 Lung |
| EWSR1 | 22q12.2 | 1 | 0.87 | 0 | 0.00 | 0.53 | 0.685 | (A) BRAF Class 1 Lung |
| COL7A1 | 3p21.31 | 6 | 11.54 | 3 | 9.38 | 0.53 | 0.685 | (A) BRAF Class 1 Lung |
| FANCM | 14q21.2 | 2 | 3.28 | 2 | 4.88 | 0.53 | 0.685 | (B) BRAF Class 2 Lung |
| SUFU | 10q24.32 | 2 | 1.12 | 1 | 0.61 | 0.531 | 0.685 | (A) BRAF Class 1 Lung |
| RAD51D | 17q12 | 2 | 1.37 | 1 | 0.75 | 0.532 | 0.685 | (A) BRAF Class 1 Lung |
| PIK3C2B | 1q32.1 | 4 | 3.57 | 4 | 4.30 | 0.533 | 0.685 | (B) BRAF Class 2 Lung |
| PMS2 | 7p22.1 | 2 | 1.13 | 1 | 0.62 | 0.533 | 0.685 | (A) BRAF Class 1 Lung |
| HIST1H3B | 6p22.2 | 3 | 2.13 | 2 | 1.53 | 0.535 | 0.685 | (A) BRAF Class 1 Lung |
| TCF7L2 | )q25.2-q25 | 3 | 1.85 | 2 | 1.33 | 0.536 | 0.685 | (A) BRAF Class 1 Lung |
| DNMT1 | 19p13.2 | 1 | 1.27 | 2 | 2.30 | 0.536 | 0.685 | (B) BRAF Class 2 Lung |
| CSF1R | 5q32 | 3 | 1.88 | 4 | 2.34 | 0.537 | 0.685 | (B) BRAF Class 2 Lung |
| CSDE1 | 1p13.2 | 0 | 0.00 | 1 | 1.72 | 0.537 | 0.685 | (B) BRAF Class 2 Lung |
| DOT1L | 19p13.3 | 4 | 4.12 | 5 | 4.85 | 0.538 | 0.685 | (B) BRAF Class 2 Lung |
| BABAM1 | 19p13.11 | 1 | 1.18 | 0 | 0.00 | 0.538 | 0.685 | (A) BRAF Class 1 Lung |
| BCL6 | 3q27.3 | 3 | 1.69 | 2 | 1.22 | 0.539 | 0.685 | (A) BRAF Class 1 Lung |
| ACVR1 | 2q24.1 | 2 | 1.53 | 1 | 0.85 | 0.539 | 0.685 | (A) BRAF Class 1 Lung |
| NBN | 8q21.3 | 5 | 2.84 | 4 | 2.42 | 0.54 | 0.685 | (A) BRAF Class 1 Lung |
| BTG1 | 12q21.33 | 0 | 0.00 | 1 | 2.50 | 0.541 | 0.685 | (B) BRAF Class 2 Lung |
| RACK1 | 5q35.3 | 0 | 0.00 | 1 | 5.00 | 0.541 | 0.685 | (B) BRAF Class 2 Lung |
| HLA-B | 6p21.33 | 0 | 0.00 | 1 | 2.17 | 0.541 | 0.685 | (B) BRAF Class 2 Lung |
| MUTYH | 1p34.1 | 4 | 2.25 | 3 | 1.83 | 0.545 | 0.685 | (A) BRAF Class 1 Lung |
| SPOP | 17q21.33 | 3 | 2.00 | 2 | 1.47 | 0.546 | 0.685 | (A) BRAF Class 1 Lung |
| EED | 11q14.2 | 1 | 1.01 | 2 | 1.79 | 0.546 | 0.685 | (B) BRAF Class 2 Lung |
| ERCC3 | 2q14.3 | 4 | 2.50 | 3 | 2.04 | 0.547 | 0.685 | (A) BRAF Class 1 Lung |
| IKZF3 | l7q12-q21. | 0 | 0.00 | 1 | 2.50 | 0.548 | 0.685 | (B) BRAF Class 2 Lung |
| PPP2R1A | 19q13.41 | 3 | 1.99 | 2 | 1.47 | 0.549 | 0.685 | (A) BRAF Class 1 Lung |
| HOXB13 | 17q21.32 | 2 | 1.46 | 1 | 0.83 | 0.55 | 0.685 | (A) BRAF Class 1 Lung |
| CCNE1 | 19q12 | 5 | 2.79 | 4 | 2.42 | 0.551 | 0.685 | (A) BRAF Class 1 Lung |
| CARD11 | 7p22.2 | 5 | 2.81 | 4 | 2.44 | 0.551 | 0.685 | (A) BRAF Class 1 Lung |
| SBDS | 7q11.21 | 1 | 1.19 | 0 | 0.00 | 0.553 | 0.685 | (A) BRAF Class 1 Lung |
| XPA | 9q22.33 | 1 | 1.19 | 0 | 0.00 | 0.553 | 0.685 | (A) BRAF Class 1 Lung |
| PAX7 | 1p36.13 | 0 | 0.00 | 1 | 3.23 | 0.554 | 0.685 | (B) BRAF Class 2 Lung |
| CCNQ | Xq28 | 0 | 0.00 | 1 | 2.44 | 0.554 | 0.685 | (B) BRAF Class 2 Lung |
| EPAS1 | 2p21 | 0 | 0.00 | 1 | 2.44 | 0.554 | 0.685 | (B) BRAF Class 2 Lung |
| ERF | 19q13.2 | 0 | 0.00 | 1 | 2.44 | 0.554 | 0.685 | (B) BRAF Class 2 Lung |
| INPPL1 | 11q13.4 | 0 | 0.00 | 1 | 2.44 | 0.554 | 0.685 | (B) BRAF Class 2 Lung |
| MSI1 | 12q24.31 | 0 | 0.00 | 1 | 2.44 | 0.554 | 0.685 | (B) BRAF Class 2 Lung |
| PRDM14 | 8q13.3 | 0 | 0.00 | 1 | 2.44 | 0.554 | 0.685 | (B) BRAF Class 2 Lung |
| RECQL | 12p12.1 | 0 | 0.00 | 1 | 2.44 | 0.554 | 0.685 | (B) BRAF Class 2 Lung |
| SMYD3 | 1q44 | 0 | 0.00 | 1 | 2.44 | 0.554 | 0.685 | (B) BRAF Class 2 Lung |
| TAP1 | 6p21.32 | 0 | 0.00 | 1 | 2.44 | 0.554 | 0.685 | (B) BRAF Class 2 Lung |
| GLI1 | 12q13.3 | 4 | 2.38 | 3 | 1.97 | 0.555 | 0.685 | (A) BRAF Class 1 Lung |
| MSH6 | 2p16.3 | 6 | 3.33 | 5 | 3.01 | 0.555 | 0.685 | (A) BRAF Class 1 Lung |
| MDM2 | 12q15 | 6 | 3.35 | 5 | 3.03 | 0.556 | 0.685 | (A) BRAF Class 1 Lung |
| CIC | 19q13.2 | 5 | 3.33 | 5 | 3.70 | 0.558 | 0.685 | (B) BRAF Class 2 Lung |
| RB1 | 13q14.2 | 7 | 3.10 | 7 | 3.30 | 0.558 | 0.685 | (B) BRAF Class 2 Lung |
| ERCC1 | 19q13.32 | 1 | 1.33 | 0 | 0.00 | 0.56 | 0.685 | (A) BRAF Class 1 Lung |

| | | | | | | | | |
|---|---|---|---|---|---|---|---|---|
| ETV4 | 17q21.31 | 2 | 1.79 | 1 | 1.05 | 0.562 | 0.685 | (A) BRAF Class 1 Lung |
| JAZF1 | p15.2-p15. | 1 | 1.72 | 0 | 0.00 | 0.563 | 0.685 | (A) BRAF Class 1 Lung |
| SS18 | 18q11.2 | 1 | 1.72 | 0 | 0.00 | 0.563 | 0.685 | (A) BRAF Class 1 Lung |
| KNSTRN | 15q15.1 | 0 | 0.00 | 1 | 2.04 | 0.563 | 0.685 | (B) BRAF Class 2 Lung |
| ELF3 | 1q32.1 | 0 | 0.00 | 1 | 2.08 | 0.565 | 0.685 | (B) BRAF Class 2 Lung |
| FLT4 | 5q35.3 | 6 | 3.43 | 5 | 3.14 | 0.566 | 0.685 | (A) BRAF Class 1 Lung |
| PIK3CA | 3q26.32 | 14 | 5.93 | 14 | 5.88 | 0.568 | 0.686 | (A) BRAF Class 1 Lung |
| KAT6A | 8p11.21 | 6 | 6.82 | 5 | 7.35 | 0.569 | 0.686 | (B) BRAF Class 2 Lung |
| CUX1 | 7q22.1 | 4 | 3.92 | 3 | 3.37 | 0.575 | 0.692 | (A) BRAF Class 1 Lung |
| WRN | 8p12 | 3 | 3.41 | 2 | 2.67 | 0.575 | 0.692 | (A) BRAF Class 1 Lung |
| KDM5C | Xp11.22 | 5 | 2.99 | 5 | 3.23 | 0.578 | 0.693 | (B) BRAF Class 2 Lung |
| TRIM37 | 17q22 | 4 | 7.69 | 2 | 6.25 | 0.585 | 0.695 | (A) BRAF Class 1 Lung |
| RECQL4 | 8q24.3 | 5 | 3.31 | 5 | 3.52 | 0.586 | 0.695 | (B) BRAF Class 2 Lung |
| PAX5 | 9p13.2 | 4 | 2.22 | 4 | 2.42 | 0.589 | 0.695 | (B) BRAF Class 2 Lung |
| PML | 15q24.1 | 1 | 1.79 | 0 | 0.00 | 0.589 | 0.695 | (A) BRAF Class 1 Lung |
| MSH3 | 5q14.1 | 1 | 1.72 | 2 | 2.70 | 0.591 | 0.695 | (B) BRAF Class 2 Lung |
| TLR4 | 9q33.1 | 1 | 4.55 | 2 | 7.14 | 0.591 | 0.695 | (B) BRAF Class 2 Lung |
| FANCB | Xp22.2 | 1 | 1.69 | 0 | 0.00 | 0.596 | 0.695 | (A) BRAF Class 1 Lung |
| ERCC4 | 16p13.12 | 4 | 2.37 | 4 | 2.55 | 0.597 | 0.695 | (B) BRAF Class 2 Lung |
| IL2RB | 22q12.3 | 0 | 0.00 | 1 | 33.33 | 0.6 | 0.695 | (B) BRAF Class 2 Lung |
| NCKAP5 | 2q21.2 | 0 | 0.00 | 1 | 33.33 | 0.6 | 0.695 | (B) BRAF Class 2 Lung |
| OR5L1 | 11q12.1 | 0 | 0.00 | 1 | 33.33 | 0.6 | 0.695 | (B) BRAF Class 2 Lung |
| ABL1 | 9q34.12 | 4 | 1.78 | 4 | 1.90 | 0.602 | 0.695 | (B) BRAF Class 2 Lung |
| MGA | 15q15.1 | 8 | 6.15 | 7 | 6.14 | 0.605 | 0.695 | (A) BRAF Class 1 Lung |
| SLC34A2 | 4p15.2 | 2 | 3.51 | 1 | 2.33 | 0.605 | 0.695 | (A) BRAF Class 1 Lung |
| RTEL1 | 20q13.33 | 2 | 6.06 | 2 | 4.88 | 0.606 | 0.695 | (A) BRAF Class 1 Lung |
| CBL | 11q23.3 | 3 | 1.66 | 3 | 1.81 | 0.616 | 0.695 | (B) BRAF Class 2 Lung |
| IPMK | 10q21.1 | 1 | 25.00 | 1 | 14.29 | 0.618 | 0.695 | (A) BRAF Class 1 Lung |
| PRKAR1A | 17q24.2 | 3 | 1.69 | 3 | 1.83 | 0.618 | 0.695 | (B) BRAF Class 2 Lung |
| DKC1 | Xq28 | 1 | 1.92 | 0 | 0.00 | 0.619 | 0.695 | (A) BRAF Class 1 Lung |
| FAAP100 | 17q25.3 | 1 | 1.92 | 0 | 0.00 | 0.619 | 0.695 | (A) BRAF Class 1 Lung |
| FAH | 15q25.1 | 1 | 1.92 | 0 | 0.00 | 0.619 | 0.695 | (A) BRAF Class 1 Lung |
| HMBS | 11q23.3 | 1 | 1.92 | 0 | 0.00 | 0.619 | 0.695 | (A) BRAF Class 1 Lung |
| LIG4 | 13q33.3 | 1 | 1.92 | 0 | 0.00 | 0.619 | 0.695 | (A) BRAF Class 1 Lung |
| MBD4 | 3q21.3 | 1 | 1.92 | 0 | 0.00 | 0.619 | 0.695 | (A) BRAF Class 1 Lung |
| NEIL2 | 8p23.1 | 1 | 1.92 | 0 | 0.00 | 0.619 | 0.695 | (A) BRAF Class 1 Lung |
| TAZ | Xq28 | 1 | 1.92 | 0 | 0.00 | 0.619 | 0.695 | (A) BRAF Class 1 Lung |
| UROD | 1p34.1 | 1 | 1.92 | 0 | 0.00 | 0.619 | 0.695 | (A) BRAF Class 1 Lung |
| BRCC3 | Xq28 | 1 | 1.92 | 1 | 3.13 | 0.62 | 0.695 | (B) BRAF Class 2 Lung |
| ENG | 9q34.11 | 1 | 1.92 | 1 | 3.13 | 0.62 | 0.695 | (B) BRAF Class 2 Lung |
| HELQ | 4q21.23 | 1 | 1.92 | 1 | 3.13 | 0.62 | 0.695 | (B) BRAF Class 2 Lung |
| KIF1B | 1p36.22 | 1 | 1.92 | 1 | 3.13 | 0.62 | 0.695 | (B) BRAF Class 2 Lung |
| MUS81 | 11q13.1 | 1 | 1.92 | 1 | 3.13 | 0.62 | 0.695 | (B) BRAF Class 2 Lung |
| RHOT1 | 17q11.2 | 1 | 1.92 | 1 | 3.13 | 0.62 | 0.695 | (B) BRAF Class 2 Lung |
| USP28 | 11q23.2 | 1 | 1.92 | 1 | 3.13 | 0.62 | 0.695 | (B) BRAF Class 2 Lung |
| XRCC4 | 5q14.2 | 1 | 1.92 | 1 | 3.13 | 0.62 | 0.695 | (B) BRAF Class 2 Lung |
| TPMT | 6p22.3 | 1 | 10.00 | 0 | 0.00 | 0.625 | 0.695 | (A) BRAF Class 1 Lung |
| BRINP3 | 1q31.1 | 0 | 0.00 | 1 | 20.00 | 0.625 | 0.695 | (B) BRAF Class 2 Lung |

| | | | | | | | | |
|---|---|---|---|---|---|---|---|---|
| CSMD3 | 8q23.3 | 0 | 0.00 | 1 | 20.00 | 0.625 | 0.695 | (B) BRAF Class 2 Lung |
| FZR1 | 19p13.3 | 0 | 0.00 | 1 | 20.00 | 0.625 | 0.695 | (B) BRAF Class 2 Lung |
| LTF | 3p21.31 | 0 | 0.00 | 1 | 20.00 | 0.625 | 0.695 | (B) BRAF Class 2 Lung |
| MTR | 1q43 | 0 | 0.00 | 1 | 20.00 | 0.625 | 0.695 | (B) BRAF Class 2 Lung |
| TGM7 | 5q15.2-q15. | 0 | 0.00 | 1 | 20.00 | 0.625 | 0.695 | (B) BRAF Class 2 Lung |
| USP8 | 15q21.2 | 1 | 1.89 | 1 | 2.94 | 0.632 | 0.697 | (B) BRAF Class 2 Lung |
| PTK2B | 8p21.2 | 1 | 1.85 | 1 | 2.86 | 0.635 | 0.697 | (B) BRAF Class 2 Lung |
| DOCK8 | 9p24.3 | 3 | 5.77 | 2 | 6.25 | 0.635 | 0.697 | (B) BRAF Class 2 Lung |
| RBBP8 | 18q11.2 | 3 | 5.77 | 2 | 6.25 | 0.635 | 0.697 | (B) BRAF Class 2 Lung |
| NAV3 | 12q21.2 | 0 | 0.00 | 1 | 14.29 | 0.636 | 0.697 | (B) BRAF Class 2 Lung |
| NIN | 14q22.1 | 0 | 0.00 | 1 | 14.29 | 0.636 | 0.697 | (B) BRAF Class 2 Lung |
| PTPRB | 12q15 | 0 | 0.00 | 1 | 14.29 | 0.636 | 0.697 | (B) BRAF Class 2 Lung |
| PTPRK | 6q22.33 | 0 | 0.00 | 1 | 14.29 | 0.636 | 0.697 | (B) BRAF Class 2 Lung |
| ROBO1 | 3p12.3 | 0 | 0.00 | 1 | 14.29 | 0.636 | 0.697 | (B) BRAF Class 2 Lung |
| ZNRF3 | 22q12.1 | 1 | 1.69 | 1 | 2.56 | 0.64 | 0.7 | (B) BRAF Class 2 Lung |
| SMARCA2 | 9p24.3 | 0 | 0.00 | 1 | 11.11 | 0.643 | 0.702 | (B) BRAF Class 2 Lung |
| MAP2K2 | 19p13.3 | 3 | 1.99 | 3 | 2.04 | 0.645 | 0.703 | (B) BRAF Class 2 Lung |
| XRCC1 | 19q13.31 | 1 | 1.89 | 1 | 2.78 | 0.648 | 0.705 | (B) BRAF Class 2 Lung |
| PRSS1 | 7q34 | 1 | 1.69 | 1 | 2.44 | 0.654 | 0.707 | (B) BRAF Class 2 Lung |
| TAL1 | 1p33 | 1 | 1.79 | 1 | 2.56 | 0.655 | 0.707 | (B) BRAF Class 2 Lung |
| SF3B1 | 2q33.1 | 2 | 1.07 | 2 | 1.16 | 0.656 | 0.707 | (B) BRAF Class 2 Lung |
| AURKA | 20q13.2 | 2 | 1.12 | 2 | 1.22 | 0.657 | 0.707 | (B) BRAF Class 2 Lung |
| MITF | 3p13 | 2 | 1.12 | 2 | 1.22 | 0.657 | 0.707 | (B) BRAF Class 2 Lung |
| CHEK2 | 22q12.1 | 2 | 1.12 | 2 | 1.21 | 0.657 | 0.707 | (B) BRAF Class 2 Lung |
| PIK3R1 | 5q13.1 | 2 | 1.11 | 2 | 1.20 | 0.658 | 0.707 | (B) BRAF Class 2 Lung |
| REL | 2p16.1 | 2 | 1.14 | 2 | 1.22 | 0.664 | 0.711 | (B) BRAF Class 2 Lung |
| PCM1 | 8p22 | 0 | 0.00 | 1 | 50.00 | 0.667 | 0.711 | (B) BRAF Class 2 Lung |
| SERPINB4 | 18q21.33 | 0 | 0.00 | 1 | 50.00 | 0.667 | 0.711 | (B) BRAF Class 2 Lung |
| TYK2 | 19p13.2 | 0 | 0.00 | 1 | 8.33 | 0.667 | 0.711 | (B) BRAF Class 2 Lung |
| TAF1 | Xq13.1 | 2 | 6.90 | 2 | 7.41 | 0.667 | 0.711 | (B) BRAF Class 2 Lung |
| ABL2 | 1q25.2 | 2 | 5.88 | 2 | 5.56 | 0.671 | 0.714 | (A) BRAF Class 1 Lung |
| RHPN2 | 19q13.11 | 2 | 7.14 | 2 | 6.90 | 0.681 | 0.724 | (A) BRAF Class 1 Lung |
| CDK4 | 12q14.1 | 2 | 1.12 | 2 | 1.14 | 0.682 | 0.724 | (B) BRAF Class 2 Lung |
| PRF1 | 10q22.1 | 1 | 1.23 | 1 | 1.59 | 0.685 | 0.726 | (B) BRAF Class 2 Lung |
| XPC | 3p25.1 | 1 | 1.19 | 1 | 1.47 | 0.696 | 0.735 | (B) BRAF Class 2 Lung |
| MSI2 | 17q22 | 1 | 3.03 | 1 | 2.44 | 0.696 | 0.735 | (A) BRAF Class 1 Lung |
| PAX3 | 2q36.1 | 1 | 4.00 | 1 | 3.23 | 0.698 | 0.736 | (A) BRAF Class 1 Lung |
| RBM15 | 1p13.3 | 0 | 0.00 | 1 | 14.29 | 0.7 | 0.737 | (B) BRAF Class 2 Lung |
| USP6 | 17p13.2 | 0 | 0.00 | 1 | 10.00 | 0.714 | 0.75 | (B) BRAF Class 2 Lung |
| RAD52 | 12p13.33 | 1 | 0.72 | 1 | 0.83 | 0.715 | 0.75 | (B) BRAF Class 2 Lung |
| PPM1D | 17q23.2 | 1 | 0.77 | 1 | 0.88 | 0.717 | 0.751 | (B) BRAF Class 2 Lung |
| CDK8 | 13q12.13 | 1 | 0.67 | 1 | 0.74 | 0.724 | 0.751 | (B) BRAF Class 2 Lung |
| SOS1 | 2p22.1 | 1 | 1.12 | 1 | 1.25 | 0.724 | 0.751 | (B) BRAF Class 2 Lung |
| CBFB | 16q22.1 | 1 | 0.67 | 1 | 0.74 | 0.726 | 0.751 | (B) BRAF Class 2 Lung |
| AXIN1 | 16p13.3 | 1 | 1.06 | 1 | 0.98 | 0.73 | 0.751 | (A) BRAF Class 1 Lung |
| FOXO1 | 13q14.11 | 1 | 1.22 | 1 | 1.12 | 0.731 | 0.751 | (A) BRAF Class 1 Lung |
| SOX9 | 17q24.3 | 1 | 0.58 | 1 | 0.62 | 0.732 | 0.751 | (B) BRAF Class 2 Lung |
| ID3 | 1p36.12 | 1 | 0.61 | 1 | 0.65 | 0.735 | 0.751 | (B) BRAF Class 2 Lung |

| | | | | | | | | |
|---|---|---|---|---|---|---|---|---|
| CSF3R | 1p34.3 | 1 | 0.65 | 1 | 0.68 | 0.737 | 0.751 | (B) BRAF Class 2 Lung |
| SMAD3 | 15q22.33 | 1 | 1.11 | 1 | 1.05 | 0.738 | 0.751 | (A) BRAF Class 1 Lung |
| SMARCB1 | q11.23\|22c | 1 | 0.45 | 1 | 0.47 | 0.738 | 0.751 | (B) BRAF Class 2 Lung |
| CHD2 | 15q26.1 | 1 | 3.23 | 1 | 3.03 | 0.738 | 0.751 | (A) BRAF Class 1 Lung |
| SLIT2 | 4p15.31 | 1 | 3.23 | 1 | 3.03 | 0.738 | 0.751 | (A) BRAF Class 1 Lung |
| SYK | 9q22.2 | 1 | 0.79 | 1 | 0.76 | 0.739 | 0.751 | (A) BRAF Class 1 Lung |
| BBC3 | 19q13.32 | 1 | 1.32 | 1 | 1.27 | 0.742 | 0.751 | (A) BRAF Class 1 Lung |
| H3F3C | 12p11.21 | 1 | 1.32 | 1 | 1.27 | 0.742 | 0.751 | (A) BRAF Class 1 Lung |
| PIK3R3 | 1p34.1 | 1 | 1.32 | 1 | 1.27 | 0.742 | 0.751 | (A) BRAF Class 1 Lung |
| RPS6KA4 | 11q13.1 | 1 | 1.32 | 1 | 1.27 | 0.742 | 0.751 | (A) BRAF Class 1 Lung |
| SOX17 | 8q11.23 | 1 | 1.32 | 1 | 1.27 | 0.742 | 0.751 | (A) BRAF Class 1 Lung |
| PIK3CB | 3q22.3 | 1 | 1.09 | 1 | 1.05 | 0.743 | 0.751 | (A) BRAF Class 1 Lung |
| HLA-A | 6p22.1 | 1 | 1.22 | 1 | 1.25 | 0.745 | 0.751 | (B) BRAF Class 2 Lung |
| ZNF708 | 19p12 | 1 | 3.57 | 1 | 3.45 | 0.746 | 0.751 | (A) BRAF Class 1 Lung |
| RAD54L | 1p34.1 | 1 | 1.15 | 1 | 1.12 | 0.746 | 0.751 | (A) BRAF Class 1 Lung |
| HSP90AA1 | 14q32.31 | 1 | 3.33 | 1 | 3.45 | 0.746 | 0.751 | (B) BRAF Class 2 Lung |
| HIST1H3F | 6p22.2 | 1 | 1.35 | 1 | 1.33 | 0.748 | 0.751 | (A) BRAF Class 1 Lung |
| GPS2 | 17p13.1 | 1 | 1.33 | 1 | 1.32 | 0.748 | 0.751 | (A) BRAF Class 1 Lung |
| ORCS8-MEF2 | 19p13.11 | 1 | 0.42 | 1 | 0.42 | 0.748 | 0.751 | (A) BRAF Class 1 Lung |
| PNRC1 | 6q15 | 1 | 0.89 | 1 | 0.88 | 0.749 | 0.751 | (A) BRAF Class 1 Lung |
| PLCG2 | 16q23.3 | 1 | 1.18 | 1 | 1.18 | 0.751 | 0.753 | (B) BRAF Class 2 Lung |
| ANKRD26 | 10p12.1 | 1 | 5.88 | 1 | 5.88 | 0.758 | 0.758 | (B) BRAF Class 2 Lung |

#### Supplementary Table 16

| Gene | Cytoband | Class 1 Occurrence | Class 1 % | Class 2 Occurrence | Class 2 % | SumClass | pval | qval | Enriched in |
|---|---|---|---|---|---|---|---|---|---|
| ERBB4 | 2q34 | 2 | 0.88 | 17 | 7.46 | 19 | 0.00029 | 0.0306 | (B) BRAF Class 2 Lung |
| EPHA3 | 3p11.1 | 3 | 2.38 | 20 | 15.15 | 23 | 0.00021 | 0.0269 | (B) BRAF Class 2 Lung |
| TP53 | 17p13.1 | 74 | 31.36 | 114 | 50.22 | 188 | 2.6E-05 | 0.00414 | (B) BRAF Class 2 Lung |
| KEAP1 | 19p13.2 | 3 | 1.80 | 33 | 21.29 | 36 | 6.7E-09 | 1.4E-06 | (B) BRAF Class 2 Lung |
| SETD2 | 3p21.31 | 71 | 39.66 | 11 | 6.67 | 82 | 8.6E-14 | 2.8E-11 | (A) BRAF Class 1 Lung |
| STK11 | 19p13.3 | 6 | 2.59 | 60 | 27.52 | 66 | 4.2E-15 | 2.7E-12 | (B) BRAF Class 2 Lung |

#### Supplementary Table 17

| Gene | Cytoband | Class 1 Occurrence | Class 1 % | Class 3 Occurrence | Class 3 % | pval | qval | Most Enriched In |
|---|---|---|---|---|---|---|---|---|
| STK11 | 19p13.3 | 6 | 2.58 | 57 | 29.53 | 5.6E-16 | 3.5E-13 | (B) BRAF Class 3 Lung |
| KEAP1 | 19p13.2 | 3 | 1.79 | 40 | 27.40 | 5.4E-12 | 1.7E-09 | (B) BRAF Class 3 Lung |
| SETD2 | 3p21.31 | 71 | 39.44 | 15 | 9.80 | 2.1E-10 | 4.4E-08 | (A) BRAF Class 1 Lung |
| KRAS | 12p12.1 | 9 | 3.80 | 41 | 20.40 | 3.1E-08 | 4.8E-06 | (B) BRAF Class 3 Lung |
| TP53 | 17p13.1 | 74 | 31.22 | 101 | 50.75 | 2.5E-05 | 0.00317 | (B) BRAF Class 3 Lung |
| ERBB4 | 2q34 | 2 | 0.87 | 17 | 8.85 | 6.4E-05 | 0.00662 | (B) BRAF Class 3 Lung |
| PTPRD | )p24.1-p2[illegible] | 5 | 4.72 | 22 | 21.57 | 0.00024 | 0.0211 | (B) BRAF Class 3 Lung |
| RBM10 | Xp11.3 | 5 | 3.42 | 20 | 15.50 | 0.00044 | 0.0342 | (B) BRAF Class 3 Lung |
| NF1 | 17q11.2 | 5 | 2.76 | 19 | 12.18 | 0.00072 | 0.0496 | (B) BRAF Class 3 Lung |
| MET | 7q31.2 | 3 | 1.28 | 15 | 7.50 | 0.00107 | 0.0666 | (B) BRAF Class 3 Lung |
| NOTCH3 | 19p13.12 | 3 | 2.01 | 14 | 10.77 | 0.00216 | 0.122 | (B) BRAF Class 3 Lung |
| PTPRT | )q12-q13.[illegible] | 2 | 2.41 | 13 | 15.85 | 0.00236 | 0.122 | (B) BRAF Class 3 Lung |
| PDGFRA | 4q12 | 4 | 1.75 | 15 | 7.73 | 0.00288 | 0.138 | (B) BRAF Class 3 Lung |
| ATRX | Xq21.1 | 6 | 3.33 | 17 | 11.11 | 0.00479 | 0.213 | (B) BRAF Class 3 Lung |
| BCORL1 | Xq26.1 | 1 | 0.84 | 8 | 8.51 | 0.00683 | 0.266 | (B) BRAF Class 3 Lung |
| PIK3C2G | 12p12.3 | 1 | 1.15 | 9 | 10.98 | 0.00692 | 0.266 | (B) BRAF Class 3 Lung |
| BRCA1 | 17q21.31 | 3 | 1.65 | 12 | 7.64 | 0.00725 | 0.266 | (B) BRAF Class 3 Lung |
| RAF1 | 3p25.2 | 0 | 0.00 | 6 | 3.73 | 0.00948 | 0.328 | (B) BRAF Class 3 Lung |
| PAK5 | 20p12.2 | 0 | 0.00 | 6 | 7.89 | 0.0103 | 0.335 | (B) BRAF Class 3 Lung |
| ROS1 | 6q22.1 | 4 | 2.17 | 13 | 8.07 | 0.0107 | 0.335 | (B) BRAF Class 3 Lung |
| ERBB2 | 17q12 | 2 | 0.85 | 9 | 4.52 | 0.0163 | 0.466 | (B) BRAF Class 3 Lung |
| EPHA3 | 3p11.1 | 3 | 2.38 | 11 | 9.57 | 0.0165 | 0.466 | (B) BRAF Class 3 Lung |
| CDH1 | 16q22.1 | 0 | 0.00 | 5 | 2.72 | 0.0183 | 0.467 | (B) BRAF Class 3 Lung |
| BRIP1 | 17q23.2 | 0 | 0.00 | 5 | 3.29 | 0.0197 | 0.467 | (B) BRAF Class 3 Lung |
| SUZ12 | 17q11.2 | 0 | 0.00 | 5 | 3.60 | 0.0202 | 0.467 | (B) BRAF Class 3 Lung |
| CDK12 | 17q12 | 0 | 0.00 | 5 | 3.82 | 0.0204 | 0.467 | (B) BRAF Class 3 Lung |
| GRIN2A | 16p13.2 | 2 | 2.13 | 9 | 10.34 | 0.0211 | 0.467 | (B) BRAF Class 3 Lung |
| SMAD4 | 18q21.2 | 17 | 7.30 | 5 | 2.59 | 0.0225 | 0.467 | (A) BRAF Class 1 Lung |
| PPP2R1A | 19q13.41 | 3 | 1.97 | 10 | 7.63 | 0.0228 | 0.467 | (B) BRAF Class 3 Lung |
| SMARCA4 | 19p13.2 | 11 | 6.11 | 20 | 13.07 | 0.0233 | 0.467 | (B) BRAF Class 3 Lung |
| KEL | 7q34 | 0 | 0.00 | 5 | 17.24 | 0.0237 | 0.467 | (B) BRAF Class 3 Lung |
| MDM2 | 12q15 | 6 | 3.33 | 0 | 0.00 | 0.024 | 0.467 | (A) BRAF Class 1 Lung |
| ERCC6 | 10q11.23 | 0 | 0.00 | 4 | 10.81 | 0.0258 | 0.472 | (B) BRAF Class 3 Lung |
| EPHB1 | 3q22.2 | 0 | 0.00 | 5 | 5.38 | 0.0259 | 0.472 | (B) BRAF Class 3 Lung |
| FAT1 | 4q35.2 | 14 | 9.79 | 23 | 18.70 | 0.0277 | 0.472 | (B) BRAF Class 3 Lung |
| JAK3 | 19p13.11 | 2 | 0.89 | 8 | 4.28 | 0.028 | 0.472 | (B) BRAF Class 3 Lung |
| TP63 | 3q28 | 2 | 2.60 | 9 | 11.69 | 0.028 | 0.472 | (B) BRAF Class 3 Lung |
| PMS1 | 2q32.2 | 2 | 1.21 | 8 | 5.63 | 0.0307 | 0.494 | (B) BRAF Class 3 Lung |
| NOTCH2 | 1p12 | 5 | 2.81 | 12 | 8.00 | 0.031 | 0.494 | (B) BRAF Class 3 Lung |
| TNFAIP3 | 6q23.3 | 1 | 0.56 | 6 | 3.95 | 0.0382 | 0.561 | (B) BRAF Class 3 Lung |
| ERCC5 | 13q33.1 | 1 | 0.62 | 6 | 4.38 | 0.0383 | 0.561 | (B) BRAF Class 3 Lung |
| ARID2 | 12q12 | 8 | 4.55 | 15 | 10.27 | 0.0385 | 0.561 | (B) BRAF Class 3 Lung |
| NRAS | 1p13.2 | 3 | 1.27 | 9 | 4.48 | 0.0387 | 0.561 | (B) BRAF Class 3 Lung |
| FGFR2 | 10q26.13 | 0 | 0.00 | 4 | 1.99 | 0.0436 | 0.568 | (B) BRAF Class 3 Lung |
| ERG | 21q22.2 | 0 | 0.00 | 4 | 3.08 | 0.0447 | 0.568 | (B) BRAF Class 3 Lung |

| | | | | | | | | |
|---|---|---|---|---|---|---|---|---|
| ETV6 | 12p13.2 | 0 | 0.00 | 4 | 2.63 | 0.0457 | 0.568 | (B) BRAF Class 3 Lung |
| ESR1 | q25.1-q25. | 0 | 0.00 | 4 | 2.47 | 0.0466 | 0.568 | (B) BRAF Class 3 Lung |
| FGFR1 | 8p11.23 | 4 | 1.71 | 10 | 5.03 | 0.0469 | 0.568 | (B) BRAF Class 3 Lung |
| INHBA | 7p14.1 | 0 | 0.00 | 4 | 4.82 | 0.0487 | 0.568 | (B) BRAF Class 3 Lung |
| MYC | 8q24.21 | 4 | 2.21 | 10 | 6.41 | 0.0487 | 0.568 | (B) BRAF Class 3 Lung |
| NOTCH1 | 9q34.3 | 5 | 2.16 | 11 | 5.70 | 0.0497 | 0.568 | (B) BRAF Class 3 Lung |
| MAP3K13 | 3q27.2 | 1 | 1.15 | 6 | 7.32 | 0.0497 | 0.568 | (B) BRAF Class 3 Lung |
| IRS2 | 13q34 | 1 | 1.02 | 6 | 6.45 | 0.0511 | 0.568 | (B) BRAF Class 3 Lung |
| TGFBR1 | 9q22.33 | 0 | 0.00 | 4 | 5.56 | 0.0536 | 0.568 | (B) BRAF Class 3 Lung |
| PRSS1 | 7q34 | 1 | 1.67 | 5 | 10.87 | 0.0537 | 0.568 | (B) BRAF Class 3 Lung |
| SF3B1 | 2q33.1 | 2 | 1.06 | 7 | 4.38 | 0.0539 | 0.568 | (B) BRAF Class 3 Lung |
| POLE | 12q24.33 | 4 | 2.60 | 10 | 7.30 | 0.0546 | 0.568 | (B) BRAF Class 3 Lung |
| HGF | 7q21.11 | 0 | 0.00 | 4 | 4.30 | 0.0555 | 0.568 | (B) BRAF Class 3 Lung |
| ARID5B | 10q21.2 | 0 | 0.00 | 4 | 5.19 | 0.0556 | 0.568 | (B) BRAF Class 3 Lung |
| INPP4B | 4q31.21 | 1 | 1.06 | 6 | 6.52 | 0.0556 | 0.568 | (B) BRAF Class 3 Lung |
| FANCI | 15q26.1 | 5 | 6.25 | 0 | 0.00 | 0.0556 | 0.568 | (A) BRAF Class 1 Lung |
| RET | 10q11.21 | 7 | 2.95 | 13 | 6.47 | 0.0635 | 0.634 | (B) BRAF Class 3 Lung |
| RHBDF2 | 17q25.1 | 5 | 9.43 | 0 | 0.00 | 0.0653 | 0.634 | (A) BRAF Class 1 Lung |
| MCL1 | 1q21.2 | 3 | 1.68 | 8 | 5.26 | 0.0656 | 0.634 | (B) BRAF Class 3 Lung |
| PTPN14 | 1q32.3-q41 | 0 | 0.00 | 3 | 8.11 | 0.0661 | 0.634 | (B) BRAF Class 3 Lung |
| DROSHA | 5p13.3 | 0 | 0.00 | 4 | 10.53 | 0.0678 | 0.64 | (B) BRAF Class 3 Lung |
| GLI1 | 12q13.3 | 4 | 2.37 | 9 | 6.38 | 0.0705 | 0.651 | (B) BRAF Class 3 Lung |
| SDHA | 5p15.33 | 3 | 1.84 | 8 | 5.63 | 0.0711 | 0.651 | (B) BRAF Class 3 Lung |
| TSHR | 14q31.1 | 1 | 0.65 | 5 | 3.85 | 0.0724 | 0.654 | (B) BRAF Class 3 Lung |
| EPHA5 | q13.1-q13. | 3 | 2.46 | 8 | 7.34 | 0.0759 | 0.667 | (B) BRAF Class 3 Lung |
| NTRK1 | 1q23.1 | 4 | 2.15 | 9 | 5.66 | 0.0772 | 0.667 | (B) BRAF Class 3 Lung |
| PRKN | 6q26 | 1 | 0.57 | 5 | 3.31 | 0.0777 | 0.667 | (B) BRAF Class 3 Lung |
| DDB2 | 11p11.2 | 0 | 0.00 | 3 | 4.62 | 0.0792 | 0.667 | (B) BRAF Class 3 Lung |
| EXT1 | 8q24.11 | 0 | 0.00 | 3 | 4.62 | 0.0792 | 0.667 | (B) BRAF Class 3 Lung |
| CBLB | 3q13.11 | 0 | 0.00 | 3 | 4.41 | 0.0856 | 0.691 | (B) BRAF Class 3 Lung |
| FUS | 16p11.2 | 0 | 0.00 | 3 | 4.69 | 0.0911 | 0.691 | (B) BRAF Class 3 Lung |
| IRS1 | 2q36.3 | 1 | 1.32 | 5 | 6.94 | 0.0926 | 0.691 | (B) BRAF Class 3 Lung |
| ATR | 3q23 | 6 | 3.95 | 11 | 8.40 | 0.0935 | 0.691 | (B) BRAF Class 3 Lung |
| ELOC | 8q21.11 | 0 | 0.00 | 3 | 2.73 | 0.0937 | 0.691 | (B) BRAF Class 3 Lung |
| CHEK2 | 22q12.1 | 2 | 1.11 | 6 | 3.92 | 0.0949 | 0.691 | (B) BRAF Class 3 Lung |
| DAXX | 6p21.32 | 0 | 0.00 | 3 | 2.36 | 0.0962 | 0.691 | (B) BRAF Class 3 Lung |
| GEN1 | 2p24.2 | 4 | 5.19 | 0 | 0.00 | 0.0964 | 0.691 | (A) BRAF Class 1 Lung |
| SRC | 20q11.23 | 0 | 0.00 | 3 | 2.03 | 0.0988 | 0.691 | (B) BRAF Class 3 Lung |
| GNA11 | 19p13.3 | 0 | 0.00 | 3 | 1.51 | 0.099 | 0.691 | (B) BRAF Class 3 Lung |
| KMT2A | 11q23.3 | 9 | 4.97 | 14 | 9.15 | 0.0995 | 0.691 | (B) BRAF Class 3 Lung |
| ARID1B | 6q25.3 | 3 | 1.75 | 7 | 4.83 | 0.109 | 0.691 | (B) BRAF Class 3 Lung |
| PIK3CA | 3q26.32 | 14 | 5.91 | 19 | 9.45 | 0.112 | 0.691 | (B) BRAF Class 3 Lung |
| GABRA6 | 5q34 | 0 | 0.00 | 3 | 10.34 | 0.112 | 0.691 | (B) BRAF Class 3 Lung |
| RPS6KB2 | 11q13.2 | 0 | 0.00 | 3 | 4.17 | 0.113 | 0.691 | (B) BRAF Class 3 Lung |
| TGFBR2 | 3p24.1 | 0 | 0.00 | 3 | 3.23 | 0.114 | 0.691 | (B) BRAF Class 3 Lung |
| TOPBP1 | 3q22.1 | 4 | 7.55 | 0 | 0.00 | 0.115 | 0.691 | (A) BRAF Class 1 Lung |
| TERT | 5p15.33 | 11 | 6.29 | 16 | 10.53 | 0.117 | 0.691 | (B) BRAF Class 3 Lung |
| SNCAIP | 5q23.2 | 0 | 0.00 | 3 | 8.82 | 0.119 | 0.691 | (B) BRAF Class 3 Lung |

| | | | | | | | | |
|---|---|---|---|---|---|---|---|---|
| XRCC3 | 14q32.33 | 4 | 6.56 | 0 | 0.00 | 0.123 | 0.691 | (A) BRAF Class 1 Lung |
| DICER1 | 14q32.13 | 5 | 2.87 | 9 | 6.16 | 0.123 | 0.691 | (B) BRAF Class 3 Lung |
| CARD11 | 7p22.2 | 5 | 2.79 | 9 | 5.92 | 0.128 | 0.691 | (B) BRAF Class 3 Lung |
| MAP3K1 | 5q11.2 | 5 | 2.81 | 9 | 5.92 | 0.131 | 0.691 | (B) BRAF Class 3 Lung |
| PTPN11 | 12q24.13 | 1 | 0.43 | 4 | 2.08 | 0.133 | 0.691 | (B) BRAF Class 3 Lung |
| AKT1 | 14q32.33 | 7 | 2.95 | 2 | 1.00 | 0.135 | 0.691 | (A) BRAF Class 1 Lung |
| CEBPA | 19q13.11 | 1 | 0.56 | 4 | 2.70 | 0.135 | 0.691 | (B) BRAF Class 3 Lung |
| AGO2 | 8q24.3 | 0 | 0.00 | 3 | 8.33 | 0.136 | 0.691 | (B) BRAF Class 3 Lung |
| PRDM14 | 8q13.3 | 0 | 0.00 | 3 | 8.33 | 0.136 | 0.691 | (B) BRAF Class 3 Lung |
| TENT5C | 1p12 | 1 | 0.56 | 4 | 2.63 | 0.14 | 0.691 | (B) BRAF Class 3 Lung |
| DNMT3A | 2p23.3 | 9 | 4.97 | 13 | 8.50 | 0.142 | 0.691 | (B) BRAF Class 3 Lung |
| CDKN2B | 9p21.3 | 8 | 4.47 | 12 | 7.89 | 0.142 | 0.691 | (B) BRAF Class 3 Lung |
| SOS1 | 2p22.1 | 1 | 1.11 | 4 | 5.13 | 0.142 | 0.691 | (B) BRAF Class 3 Lung |
| HIST1H3B | 6p22.2 | 3 | 2.11 | 0 | 0.00 | 0.146 | 0.691 | (A) BRAF Class 1 Lung |
| KMT2D | 12q13.12 | 16 | 8.94 | 20 | 13.16 | 0.147 | 0.691 | (B) BRAF Class 3 Lung |
| JAK1 | 1p31.3 | 1 | 0.65 | 4 | 2.99 | 0.147 | 0.691 | (B) BRAF Class 3 Lung |
| NSD3 | 8p11.23 | 1 | 1.11 | 4 | 5.00 | 0.149 | 0.691 | (B) BRAF Class 3 Lung |
| FGFR3 | 4p16.3 | 5 | 2.14 | 1 | 0.50 | 0.15 | 0.691 | (A) BRAF Class 1 Lung |
| MED12 | Xq13.1 | 6 | 3.97 | 10 | 7.52 | 0.15 | 0.691 | (B) BRAF Class 3 Lung |
| TRAF7 | 16p13.3 | 1 | 0.74 | 4 | 3.33 | 0.15 | 0.691 | (B) BRAF Class 3 Lung |
| SPOP | 17q21.33 | 3 | 1.99 | 0 | 0.00 | 0.152 | 0.691 | (A) BRAF Class 1 Lung |
| CCND1 | 11q13.3 | 3 | 1.67 | 0 | 0.00 | 0.154 | 0.691 | (A) BRAF Class 1 Lung |
| PRKAR1A | 17q24.2 | 3 | 1.68 | 0 | 0.00 | 0.157 | 0.691 | (A) BRAF Class 1 Lung |
| RARA | 17q21.2 | 3 | 1.68 | 0 | 0.00 | 0.157 | 0.691 | (A) BRAF Class 1 Lung |
| BRAF | 7q34 | 234 | 98.73 | 201 | 100.00 | 0.158 | 0.691 | (B) BRAF Class 3 Lung |
| CUX1 | 7q22.1 | 4 | 3.88 | 7 | 8.43 | 0.16 | 0.691 | (B) BRAF Class 3 Lung |
| PMS2 | 7p22.1 | 2 | 1.12 | 5 | 3.33 | 0.16 | 0.691 | (B) BRAF Class 3 Lung |
| KIT | 4q12 | 2 | 0.84 | 5 | 2.49 | 0.163 | 0.691 | (B) BRAF Class 3 Lung |
| FANCA | 16q24.3 | 2 | 1.11 | 5 | 3.27 | 0.163 | 0.691 | (B) BRAF Class 3 Lung |
| PRPF8 | 17p13.3 | 3 | 10.00 | 0 | 0.00 | 0.164 | 0.691 | (A) BRAF Class 1 Lung |
| EXO1 | 1q43 | 0 | 0.00 | 2 | 5.41 | 0.166 | 0.691 | (B) BRAF Class 3 Lung |
| GBA | 1q22 | 0 | 0.00 | 2 | 5.41 | 0.166 | 0.691 | (B) BRAF Class 3 Lung |
| SLC25A13 | 7q21.3 | 0 | 0.00 | 2 | 5.41 | 0.166 | 0.691 | (B) BRAF Class 3 Lung |
| SOX17 | 8q11.23 | 1 | 1.32 | 4 | 5.56 | 0.167 | 0.691 | (B) BRAF Class 3 Lung |
| AMER1 | Xq11.2 | 1 | 1.02 | 4 | 4.30 | 0.168 | 0.691 | (B) BRAF Class 3 Lung |
| BTK | Xq22.1 | 1 | 1.01 | 4 | 4.26 | 0.168 | 0.691 | (B) BRAF Class 3 Lung |
| ATM | 11q22.3 | 16 | 7.08 | 19 | 10.22 | 0.169 | 0.691 | (B) BRAF Class 3 Lung |
| ELANE | 19p13.3 | 0 | 0.00 | 2 | 5.13 | 0.17 | 0.691 | (B) BRAF Class 3 Lung |
| CIITA | 16p13.13 | 3 | 3.61 | 0 | 0.00 | 0.174 | 0.691 | (A) BRAF Class 1 Lung |
| FLT3 | 13q12.2 | 4 | 1.76 | 7 | 3.74 | 0.174 | 0.691 | (B) BRAF Class 3 Lung |
| MECOM | 3q26.2 | 0 | 0.00 | 2 | 3.33 | 0.174 | 0.691 | (B) BRAF Class 3 Lung |
| MSH2 | 2p21-p16.3 | 3 | 1.69 | 6 | 3.97 | 0.177 | 0.691 | (B) BRAF Class 3 Lung |
| KAT6B | 10q22.2 | 0 | 0.00 | 2 | 4.65 | 0.182 | 0.691 | (B) BRAF Class 3 Lung |
| SPEN | 36.21-p36. | 7 | 7.22 | 3 | 3.23 | 0.183 | 0.691 | (A) BRAF Class 1 Lung |
| FANCF | 11p14.3 | 0 | 0.00 | 2 | 2.27 | 0.183 | 0.691 | (B) BRAF Class 3 Lung |
| ZFHX3 | 5q22.2-q22 | 7 | 8.97 | 11 | 15.07 | 0.183 | 0.691 | (B) BRAF Class 3 Lung |
| FLT4 | 5q35.3 | 6 | 3.41 | 9 | 6.16 | 0.183 | 0.691 | (B) BRAF Class 3 Lung |
| ITK | 5q33.3 | 1 | 1.85 | 3 | 7.89 | 0.189 | 0.691 | (B) BRAF Class 3 Lung |

| | | | | | | | | |
|---|---|---|---|---|---|---|---|---|
| POLD1 | 19q13.33 | 6 | 4.00 | 2 | 1.53 | 0.19 | 0.691 | (A) BRAF Class 1 Lung |
| PTEN | 10q23.31 | 6 | 2.53 | 9 | 4.52 | 0.191 | 0.691 | (B) BRAF Class 3 Lung |
| TFE3 | Xp11.23 | 0 | 0.00 | 2 | 3.23 | 0.194 | 0.691 | (B) BRAF Class 3 Lung |
| PARP1 | 1q42.12 | 4 | 4.44 | 1 | 1.15 | 0.195 | 0.691 | (A) BRAF Class 1 Lung |
| ARHGAP35 | 19q13.32 | 5 | 8.62 | 1 | 2.38 | 0.195 | 0.691 | (A) BRAF Class 1 Lung |
| STAT6 | 12q13.3 | 0 | 0.00 | 2 | 2.90 | 0.197 | 0.691 | (B) BRAF Class 3 Lung |
| TET2 | 4q24 | 10 | 5.52 | 13 | 8.50 | 0.197 | 0.691 | (B) BRAF Class 3 Lung |
| EME1 | 17q21.33 | 3 | 5.66 | 0 | 0.00 | 0.199 | 0.691 | (A) BRAF Class 1 Lung |
| HABP2 | 10q25.3 | 3 | 5.66 | 0 | 0.00 | 0.199 | 0.691 | (A) BRAF Class 1 Lung |
| CREBBP | 16p13.3 | 9 | 5.00 | 12 | 7.84 | 0.201 | 0.691 | (B) BRAF Class 3 Lung |
| TLR4 | 9q33.1 | 1 | 4.55 | 4 | 16.67 | 0.202 | 0.691 | (B) BRAF Class 3 Lung |
| EPHA7 | 6q16.1 | 4 | 3.39 | 7 | 6.73 | 0.202 | 0.691 | (B) BRAF Class 3 Lung |
| VHL | 3p25.3 | 0 | 0.00 | 2 | 1.08 | 0.203 | 0.691 | (B) BRAF Class 3 Lung |
| MAGI2 | 7q21.11 | 0 | 0.00 | 2 | 8.70 | 0.207 | 0.691 | (B) BRAF Class 3 Lung |
| IGF2 | 11p15.5 | 0 | 0.00 | 2 | 1.63 | 0.207 | 0.691 | (B) BRAF Class 3 Lung |
| RUNX1T1 | 8q21.3 | 1 | 1.12 | 3 | 4.55 | 0.207 | 0.691 | (B) BRAF Class 3 Lung |
| RECQL4 | 8q24.3 | 5 | 3.29 | 8 | 6.02 | 0.207 | 0.691 | (B) BRAF Class 3 Lung |
| MT-ND5 | - | 0 | 0.00 | 2 | 1.00 | 0.21 | 0.691 | (B) BRAF Class 3 Lung |
| AKT3 | 1q43-q44 | 0 | 0.00 | 2 | 1.30 | 0.211 | 0.691 | (B) BRAF Class 3 Lung |
| SDHC | 1q23.3 | 0 | 0.00 | 2 | 1.34 | 0.211 | 0.691 | (B) BRAF Class 3 Lung |
| SH2D1A | Xq25 | 0 | 0.00 | 2 | 1.75 | 0.214 | 0.691 | (B) BRAF Class 3 Lung |
| H3F3B | 17q25.1 | 0 | 0.00 | 2 | 1.72 | 0.216 | 0.691 | (B) BRAF Class 3 Lung |
| MTAP | 9p21.3 | 2 | 3.51 | 4 | 9.30 | 0.216 | 0.691 | (B) BRAF Class 3 Lung |
| XPC | 3p25.1 | 1 | 1.18 | 3 | 4.62 | 0.217 | 0.691 | (B) BRAF Class 3 Lung |
| CASP8 | 2q33.1 | 0 | 0.00 | 2 | 1.57 | 0.217 | 0.691 | (B) BRAF Class 3 Lung |
| GLI2 | 2q14.2 | 2 | 2.35 | 4 | 6.25 | 0.218 | 0.691 | (B) BRAF Class 3 Lung |
| SPRED1 | 15q14 | 2 | 5.41 | 0 | 0.00 | 0.222 | 0.691 | (A) BRAF Class 1 Lung |
| LRP1B | q22.1-q22. | 6 | 16.67 | 8 | 27.59 | 0.223 | 0.691 | (B) BRAF Class 3 Lung |
| IGF1 | 12q23.2 | 0 | 0.00 | 2 | 2.63 | 0.224 | 0.691 | (B) BRAF Class 3 Lung |
| RTEL1 | 20q13.33 | 2 | 6.06 | 0 | 0.00 | 0.225 | 0.691 | (A) BRAF Class 1 Lung |
| RHPN2 | 19q13.11 | 2 | 7.14 | 4 | 18.18 | 0.225 | 0.691 | (B) BRAF Class 3 Lung |
| GATA1 | Xp11.23 | 0 | 0.00 | 2 | 2.27 | 0.227 | 0.691 | (B) BRAF Class 3 Lung |
| PGR | 11q22.1 | 0 | 0.00 | 2 | 2.94 | 0.228 | 0.691 | (B) BRAF Class 3 Lung |
| CD79A | 19q13.2 | 0 | 0.00 | 2 | 2.25 | 0.23 | 0.691 | (B) BRAF Class 3 Lung |
| BRD4 | 19p13.12 | 7 | 4.00 | 3 | 1.97 | 0.232 | 0.691 | (A) BRAF Class 1 Lung |
| FYN | 6q21 | 0 | 0.00 | 2 | 2.74 | 0.232 | 0.691 | (B) BRAF Class 3 Lung |
| FOXP1 | 3p13 | 0 | 0.00 | 2 | 2.30 | 0.232 | 0.691 | (B) BRAF Class 3 Lung |
| TBX3 | 12q24.21 | 3 | 3.16 | 6 | 6.52 | 0.233 | 0.691 | (B) BRAF Class 3 Lung |
| RAD51 | 15q15.1 | 4 | 2.67 | 1 | 0.77 | 0.233 | 0.691 | (A) BRAF Class 1 Lung |
| ERCC4 | 16p13.12 | 4 | 2.35 | 1 | 0.68 | 0.234 | 0.691 | (A) BRAF Class 1 Lung |
| DNMT3B | 20q11.21 | 0 | 0.00 | 2 | 2.78 | 0.235 | 0.691 | (B) BRAF Class 3 Lung |
| ICOSLG | 21q22.3 | 0 | 0.00 | 2 | 2.78 | 0.235 | 0.691 | (B) BRAF Class 3 Lung |
| INPP4A | 2q11.2 | 0 | 0.00 | 2 | 2.78 | 0.235 | 0.691 | (B) BRAF Class 3 Lung |
| PLK2 | 5q11.2 | 0 | 0.00 | 2 | 2.78 | 0.235 | 0.691 | (B) BRAF Class 3 Lung |
| SHQ1 | 3p13 | 0 | 0.00 | 2 | 2.78 | 0.235 | 0.691 | (B) BRAF Class 3 Lung |
| YES1 | 18p11.32 | 0 | 0.00 | 2 | 2.78 | 0.235 | 0.691 | (B) BRAF Class 3 Lung |
| IDH2 | 15q26.1 | 4 | 1.75 | 1 | 0.51 | 0.237 | 0.691 | (A) BRAF Class 1 Lung |
| NRG1 | 8p12 | 4 | 5.48 | 6 | 10.34 | 0.238 | 0.691 | (B) BRAF Class 3 Lung |

| | | | | | | | | |
|---|---|---|---|---|---|---|---|---|
| ERCC3 | 2q14.3 | 4 | 2.48 | 1 | 0.73 | 0.24 | 0.691 | (A) BRAF Class 1 Lung |
| MAX | 14q23.3 | 4 | 3.01 | 1 | 0.88 | 0.24 | 0.691 | (A) BRAF Class 1 Lung |
| CRKL | 22q11.21 | 4 | 2.23 | 1 | 0.66 | 0.241 | 0.691 | (A) BRAF Class 1 Lung |
| CRLF2 | 2.33 and Yp | 1 | 0.58 | 3 | 2.10 | 0.243 | 0.691 | (B) BRAF Class 3 Lung |
| NFKBIA | 14q13.2 | 4 | 2.27 | 1 | 0.68 | 0.244 | 0.691 | (A) BRAF Class 1 Lung |
| PPM1D | 17q23.2 | 1 | 0.76 | 3 | 2.73 | 0.248 | 0.691 | (B) BRAF Class 3 Lung |
| PRDM1 | 6q21 | 1 | 0.56 | 3 | 1.97 | 0.253 | 0.691 | (B) BRAF Class 3 Lung |
| SMAD2 | 18q21.1 | 1 | 0.56 | 3 | 1.97 | 0.253 | 0.691 | (B) BRAF Class 3 Lung |
| AKT2 | 19q13.2 | 1 | 0.55 | 3 | 1.95 | 0.253 | 0.691 | (B) BRAF Class 3 Lung |
| RASA1 | 5q14.3 | 1 | 0.75 | 3 | 2.63 | 0.255 | 0.691 | (B) BRAF Class 3 Lung |
| PRKCI | 3q26.2 | 1 | 0.69 | 3 | 2.42 | 0.257 | 0.691 | (B) BRAF Class 3 Lung |
| MRE11 | 11q21 | 1 | 0.66 | 3 | 2.29 | 0.258 | 0.691 | (B) BRAF Class 3 Lung |
| RNF43 | 17q22 | 1 | 0.66 | 3 | 2.29 | 0.26 | 0.691 | (B) BRAF Class 3 Lung |
| SOX9 | 17q24.3 | 1 | 0.57 | 3 | 1.99 | 0.26 | 0.691 | (B) BRAF Class 3 Lung |
| EGFR | 7p11.2 | 19 | 8.02 | 12 | 5.97 | 0.261 | 0.691 | (A) BRAF Class 1 Lung |
| TP53BP1 | 15q15.3 | 2 | 2.17 | 4 | 5.19 | 0.261 | 0.691 | (B) BRAF Class 3 Lung |
| JAK2 | 9p24.1 | 2 | 0.90 | 4 | 2.15 | 0.262 | 0.691 | (B) BRAF Class 3 Lung |
| FOXA1 | 14q21.1 | 1 | 0.74 | 3 | 2.50 | 0.266 | 0.691 | (B) BRAF Class 3 Lung |
| SMO | 7q32.1 | 5 | 2.21 | 7 | 3.72 | 0.267 | 0.691 | (B) BRAF Class 3 Lung |
| EPAS1 | 2p21 | 0 | 0.00 | 2 | 5.56 | 0.269 | 0.691 | (B) BRAF Class 3 Lung |
| INPPL1 | 11q13.4 | 0 | 0.00 | 2 | 5.56 | 0.269 | 0.691 | (B) BRAF Class 3 Lung |
| HIST1H3G | 6p22.2 | 2 | 2.70 | 0 | 0.00 | 0.27 | 0.691 | (A) BRAF Class 1 Lung |
| ELF3 | 1q32.1 | 0 | 0.00 | 2 | 4.88 | 0.273 | 0.691 | (B) BRAF Class 3 Lung |
| EZH1 | 17q21.2 | 0 | 0.00 | 2 | 4.88 | 0.273 | 0.691 | (B) BRAF Class 3 Lung |
| DIS3 | 13q21.33 | 5 | 2.89 | 7 | 4.79 | 0.275 | 0.691 | (B) BRAF Class 3 Lung |
| ALK | p23.2-p23. | 8 | 3.42 | 10 | 5.03 | 0.276 | 0.691 | (B) BRAF Class 3 Lung |
| CDKN2A | 9p21.3 | 18 | 7.83 | 19 | 9.95 | 0.276 | 0.691 | (B) BRAF Class 3 Lung |
| AXL | 19q13.2 | 3 | 1.67 | 5 | 3.27 | 0.276 | 0.691 | (B) BRAF Class 3 Lung |
| ROBO1 | 3p12.3 | 0 | 0.00 | 2 | 40.00 | 0.278 | 0.691 | (B) BRAF Class 3 Lung |
| NTRK3 | 15q25.3 | 6 | 3.33 | 8 | 5.23 | 0.279 | 0.691 | (B) BRAF Class 3 Lung |
| FLT1 | 13q12.3 | 4 | 2.23 | 6 | 3.95 | 0.279 | 0.691 | (B) BRAF Class 3 Lung |
| IKZF1 | 7p12.2 | 4 | 2.23 | 6 | 3.95 | 0.279 | 0.691 | (B) BRAF Class 3 Lung |
| NKX2-1 | 14q13.3 | 4 | 2.23 | 6 | 3.95 | 0.279 | 0.691 | (B) BRAF Class 3 Lung |
| PALB2 | 16p12.2 | 4 | 2.22 | 6 | 3.92 | 0.279 | 0.691 | (B) BRAF Class 3 Lung |
| RIT1 | 1q22 | 2 | 1.43 | 4 | 3.28 | 0.279 | 0.691 | (B) BRAF Class 3 Lung |
| NEGR1 | 1p31.1 | 1 | 1.10 | 3 | 3.57 | 0.281 | 0.691 | (B) BRAF Class 3 Lung |
| STAG2 | Xq25 | 4 | 2.25 | 6 | 3.95 | 0.282 | 0.691 | (B) BRAF Class 3 Lung |
| NSD2 | 4p16.3 | 1 | 1.03 | 3 | 3.33 | 0.283 | 0.691 | (B) BRAF Class 3 Lung |
| RAD21 | 8q24.11 | 5 | 3.03 | 2 | 1.39 | 0.284 | 0.691 | (A) BRAF Class 1 Lung |
| ACVR1 | 2q24.1 | 2 | 1.52 | 0 | 0.00 | 0.285 | 0.691 | (A) BRAF Class 1 Lung |
| ARAF | Xp11.3 | 4 | 2.16 | 6 | 3.77 | 0.285 | 0.691 | (B) BRAF Class 3 Lung |
| BARD1 | 2q35 | 3 | 2.00 | 5 | 3.85 | 0.286 | 0.691 | (B) BRAF Class 3 Lung |
| EPCAM | 2p21 | 2 | 1.42 | 0 | 0.00 | 0.286 | 0.691 | (A) BRAF Class 1 Lung |
| ABL2 | 1q25.2 | 2 | 5.88 | 0 | 0.00 | 0.287 | 0.691 | (A) BRAF Class 1 Lung |
| VEGFA | 6p21.1 | 2 | 1.36 | 0 | 0.00 | 0.289 | 0.691 | (A) BRAF Class 1 Lung |
| H3F3C | 12p11.21 | 1 | 1.32 | 3 | 4.17 | 0.29 | 0.691 | (B) BRAF Class 3 Lung |
| PTPRS | 19p13.3 | 1 | 1.32 | 3 | 4.17 | 0.29 | 0.691 | (B) BRAF Class 3 Lung |
| RPS6KA4 | 11q13.1 | 1 | 1.32 | 3 | 4.17 | 0.29 | 0.691 | (B) BRAF Class 3 Lung |

| | | | | | | | | |
|---|---|---|---|---|---|---|---|---|
| STAT3 | 17q21.2 | 2 | 1.14 | 0 | 0.00 | 0.291 | 0.691 | (A) BRAF Class 1 Lung |
| SLC34A2 | 4p15.2 | 2 | 3.45 | 0 | 0.00 | 0.291 | 0.691 | (A) BRAF Class 1 Lung |
| AURKA | 20q13.2 | 2 | 1.12 | 0 | 0.00 | 0.292 | 0.691 | (A) BRAF Class 1 Lung |
| SUFU | 10q24.32 | 2 | 1.12 | 0 | 0.00 | 0.292 | 0.691 | (A) BRAF Class 1 Lung |
| MAP2K1 | 15q22.31 | 3 | 1.57 | 5 | 2.98 | 0.294 | 0.691 | (B) BRAF Class 3 Lung |
| DDR2 | 1q23.3 | 4 | 2.11 | 6 | 3.59 | 0.298 | 0.691 | (B) BRAF Class 3 Lung |
| OR5L1 | 11q12.1 | 0 | 0.00 | 2 | 66.67 | 0.3 | 0.691 | (B) BRAF Class 3 Lung |
| RAD50 | 5q31.1 | 5 | 3.36 | 2 | 1.60 | 0.302 | 0.691 | (A) BRAF Class 1 Lung |
| DOCK8 | 9p24.3 | 3 | 5.66 | 4 | 10.81 | 0.305 | 0.691 | (B) BRAF Class 3 Lung |
| TAF1 | Xq13.1 | 2 | 6.90 | 0 | 0.00 | 0.306 | 0.691 | (A) BRAF Class 1 Lung |
| AXIN1 | 16p13.3 | 1 | 1.06 | 3 | 3.23 | 0.306 | 0.691 | (B) BRAF Class 3 Lung |
| DMD | p21.2-p21. | 2 | 7.14 | 0 | 0.00 | 0.309 | 0.691 | (A) BRAF Class 1 Lung |
| FBXW7 | 4q31.3 | 5 | 2.15 | 2 | 1.04 | 0.309 | 0.691 | (A) BRAF Class 1 Lung |
| GNAS | 20q13.32 | 5 | 2.15 | 2 | 1.04 | 0.309 | 0.691 | (A) BRAF Class 1 Lung |
| ETV4 | 17q21.31 | 2 | 1.77 | 0 | 0.00 | 0.312 | 0.691 | (A) BRAF Class 1 Lung |
| TRIM37 | 17q22 | 4 | 7.55 | 1 | 2.70 | 0.312 | 0.691 | (A) BRAF Class 1 Lung |
| EED | 11q14.2 | 1 | 1.01 | 3 | 2.97 | 0.318 | 0.691 | (B) BRAF Class 3 Lung |
| CUL3 | 2q36.2 | 3 | 3.19 | 1 | 1.09 | 0.319 | 0.691 | (A) BRAF Class 1 Lung |
| PIK3CG | 7q22.3 | 4 | 4.21 | 6 | 6.90 | 0.32 | 0.691 | (B) BRAF Class 3 Lung |
| ERBB3 | 12q13.2 | 6 | 3.31 | 3 | 1.92 | 0.33 | 0.691 | (A) BRAF Class 1 Lung |
| EFNB3 | 17p13.1 | 0 | 0.00 | 1 | 33.33 | 0.333 | 0.691 | (B) BRAF Class 3 Lung |
| RINT1 | 7q22.3 | 2 | 3.77 | 0 | 0.00 | 0.344 | 0.691 | (A) BRAF Class 1 Lung |
| DOT1L | 19p13.3 | 4 | 4.12 | 6 | 6.45 | 0.347 | 0.691 | (B) BRAF Class 3 Lung |
| RB1 | 13q14.2 | 7 | 3.08 | 8 | 4.28 | 0.349 | 0.691 | (B) BRAF Class 3 Lung |
| MGA | 15q15.1 | 8 | 6.11 | 9 | 8.18 | 0.353 | 0.691 | (B) BRAF Class 3 Lung |
| KDR | 4q12 | 10 | 4.42 | 6 | 3.23 | 0.359 | 0.691 | (A) BRAF Class 1 Lung |
| POT1 | 7q31.33 | 3 | 4.84 | 4 | 8.33 | 0.359 | 0.691 | (B) BRAF Class 3 Lung |
| MSH3 | 5q14.1 | 1 | 1.72 | 3 | 4.55 | 0.36 | 0.691 | (B) BRAF Class 3 Lung |
| LIG4 | 13q33.3 | 1 | 1.89 | 2 | 5.41 | 0.367 | 0.691 | (B) BRAF Class 3 Lung |
| APC | 5q22.2 | 15 | 6.64 | 15 | 7.98 | 0.368 | 0.691 | (B) BRAF Class 3 Lung |
| FOXL2 | 3q22.3 | 3 | 1.97 | 1 | 0.76 | 0.369 | 0.691 | (A) BRAF Class 1 Lung |
| CBFA2T3 | 16q24.3 | 1 | 1.85 | 2 | 5.26 | 0.369 | 0.691 | (B) BRAF Class 3 Lung |
| FGFR4 | 5q35.2 | 3 | 1.64 | 1 | 0.64 | 0.371 | 0.691 | (A) BRAF Class 1 Lung |
| RHEB | 7q36.1 | 3 | 2.01 | 1 | 0.79 | 0.373 | 0.691 | (A) BRAF Class 1 Lung |
| FN1 | 2q35 | 1 | 33.33 | 0 | 0.00 | 0.375 | 0.691 | (A) BRAF Class 1 Lung |
| MEN1 | 11q13.1 | 3 | 1.68 | 1 | 0.66 | 0.375 | 0.691 | (A) BRAF Class 1 Lung |
| ABL1 | 9q34.12 | 4 | 1.77 | 5 | 2.69 | 0.381 | 0.691 | (B) BRAF Class 3 Lung |
| AR | Xq12 | 7 | 3.89 | 8 | 5.16 | 0.382 | 0.691 | (B) BRAF Class 3 Lung |
| TET1 | 10q21.3 | 7 | 5.26 | 8 | 6.96 | 0.384 | 0.691 | (B) BRAF Class 3 Lung |
| FANCB | Xp22.2 | 1 | 1.67 | 2 | 4.44 | 0.393 | 0.691 | (B) BRAF Class 3 Lung |
| SRP19 | 5q22.2 | 2 | 4.88 | 0 | 0.00 | 0.394 | 0.691 | (A) BRAF Class 1 Lung |
| PAX5 | 9p13.2 | 4 | 2.21 | 5 | 3.27 | 0.397 | 0.691 | (B) BRAF Class 3 Lung |
| CIC | 19q13.2 | 5 | 3.31 | 6 | 4.62 | 0.398 | 0.691 | (B) BRAF Class 3 Lung |
| ZNF217 | 20q13.2 | 3 | 2.52 | 1 | 1.06 | 0.404 | 0.691 | (A) BRAF Class 1 Lung |
| TSC2 | 16p13.3 | 13 | 7.22 | 13 | 8.50 | 0.409 | 0.691 | (B) BRAF Class 3 Lung |
| CBL | 11q23.3 | 3 | 1.65 | 4 | 2.60 | 0.409 | 0.691 | (B) BRAF Class 3 Lung |
| TSC1 | 9q34.13 | 3 | 1.67 | 4 | 2.61 | 0.411 | 0.691 | (B) BRAF Class 3 Lung |
| BABAM2 | 2p23.2 | 0 | 0.00 | 1 | 2.70 | 0.411 | 0.691 | (B) BRAF Class 3 Lung |

| | | | | | | | | |
|---|---|---|---|---|---|---|---|---|
| MCM8 | 20p12.3 | 0 | 0.00 | 1 | 2.70 | 0.411 | 0.691 | (B) BRAF Class 3 Lung |
| RNF8 | 6p21.2 | 0 | 0.00 | 1 | 2.70 | 0.411 | 0.691 | (B) BRAF Class 3 Lung |
| RUNX1 | 21q22.12 | 2 | 1.12 | 3 | 2.03 | 0.411 | 0.691 | (B) BRAF Class 3 Lung |
| SMC3 | 10q25.2 | 1 | 0.96 | 2 | 2.41 | 0.416 | 0.691 | (B) BRAF Class 3 Lung |
| MTOR | 1p36.22 | 10 | 5.43 | 7 | 4.35 | 0.417 | 0.691 | (A) BRAF Class 1 Lung |
| ASXL1 | 20q11.21 | 14 | 7.73 | 10 | 6.54 | 0.419 | 0.691 | (A) BRAF Class 1 Lung |
| U2AF1 | 21q22.3 | 3 | 1.69 | 4 | 2.61 | 0.42 | 0.691 | (B) BRAF Class 3 Lung |
| FANCE | 6p21.31 | 3 | 2.59 | 1 | 1.14 | 0.421 | 0.691 | (A) BRAF Class 1 Lung |
| WT1 | 11p13 | 2 | 1.10 | 3 | 1.96 | 0.421 | 0.691 | (B) BRAF Class 3 Lung |
| EWSR1 | 22q12.2 | 1 | 0.86 | 2 | 2.13 | 0.421 | 0.691 | (B) BRAF Class 3 Lung |
| RICTOR | 5p13.1 | 3 | 1.99 | 4 | 3.05 | 0.422 | 0.691 | (B) BRAF Class 3 Lung |
| RANBP2 | 2q13 | 2 | 7.41 | 3 | 13.04 | 0.422 | 0.691 | (B) BRAF Class 3 Lung |
| GATA3 | 10p14 | 2 | 1.12 | 3 | 1.97 | 0.423 | 0.691 | (B) BRAF Class 3 Lung |
| MITF | 3p13 | 2 | 1.12 | 3 | 1.97 | 0.423 | 0.691 | (B) BRAF Class 3 Lung |
| PIK3R1 | 5q13.1 | 2 | 1.10 | 3 | 1.95 | 0.424 | 0.691 | (B) BRAF Class 3 Lung |
| LZTR1 | 21\|22q11. | 1 | 2.94 | 2 | 7.14 | 0.426 | 0.691 | (B) BRAF Class 3 Lung |
| MAP2K2 | 19p13.3 | 3 | 1.97 | 4 | 3.01 | 0.427 | 0.691 | (B) BRAF Class 3 Lung |
| BCL2L12 | 19q13.33 | 0 | 0.00 | 1 | 1.61 | 0.428 | 0.691 | (B) BRAF Class 3 Lung |
| RHOH | 4p14 | 0 | 0.00 | 1 | 2.33 | 0.43 | 0.691 | (B) BRAF Class 3 Lung |
| WAS | Xp11.23 | 0 | 0.00 | 1 | 2.33 | 0.43 | 0.691 | (B) BRAF Class 3 Lung |
| PRKDC | 8q11.21 | 10 | 8.55 | 9 | 10.23 | 0.43 | 0.691 | (B) BRAF Class 3 Lung |
| SPTA1 | 1q23.1 | 4 | 12.90 | 5 | 17.86 | 0.433 | 0.691 | (B) BRAF Class 3 Lung |
| TCF3 | 19p13.3 | 1 | 0.63 | 2 | 1.50 | 0.433 | 0.691 | (B) BRAF Class 3 Lung |
| CRTC1 | 19p13.11 | 0 | 0.00 | 1 | 1.54 | 0.433 | 0.691 | (B) BRAF Class 3 Lung |
| CYLD | 16q12.1 | 1 | 0.68 | 2 | 1.61 | 0.434 | 0.691 | (B) BRAF Class 3 Lung |
| GATA2 | 3q21.3 | 1 | 0.67 | 2 | 1.60 | 0.434 | 0.691 | (B) BRAF Class 3 Lung |
| CSF1R | 5q32 | 3 | 1.88 | 4 | 2.82 | 0.434 | 0.691 | (B) BRAF Class 3 Lung |
| BABAM1 | 19p13.11 | 1 | 1.16 | 2 | 2.74 | 0.438 | 0.691 | (B) BRAF Class 3 Lung |
| BAP1 | 3p21.1 | 1 | 0.56 | 2 | 1.31 | 0.439 | 0.691 | (B) BRAF Class 3 Lung |
| ABCB11 | 2q31.1 | 4 | 7.27 | 4 | 10.26 | 0.439 | 0.691 | (B) BRAF Class 3 Lung |
| KDM6B | 17p13.1 | 0 | 0.00 | 1 | 4.55 | 0.44 | 0.691 | (B) BRAF Class 3 Lung |
| PRAME | 22q11.22 | 0 | 0.00 | 1 | 4.55 | 0.44 | 0.691 | (B) BRAF Class 3 Lung |
| PTK2 | 8q24.3 | 0 | 0.00 | 1 | 4.55 | 0.44 | 0.691 | (B) BRAF Class 3 Lung |
| MLLT3 | 9p21.3 | 0 | 0.00 | 1 | 6.67 | 0.441 | 0.691 | (B) BRAF Class 3 Lung |
| B2M | 15q21.1 | 1 | 0.63 | 2 | 1.47 | 0.441 | 0.691 | (B) BRAF Class 3 Lung |
| FANCL | 2p16.1 | 0 | 0.00 | 1 | 1.39 | 0.442 | 0.691 | (B) BRAF Class 3 Lung |
| EP300 | 22q13.2 | 10 | 5.59 | 7 | 4.61 | 0.442 | 0.691 | (A) BRAF Class 1 Lung |
| CDK8 | 13q12.13 | 1 | 0.66 | 2 | 1.54 | 0.444 | 0.691 | (B) BRAF Class 3 Lung |
| AXIN2 | 17q24.1 | 2 | 1.49 | 3 | 2.52 | 0.444 | 0.691 | (B) BRAF Class 3 Lung |
| CHEK1 | 11q24.2 | 1 | 0.66 | 2 | 1.53 | 0.444 | 0.691 | (B) BRAF Class 3 Lung |
| IPMK | 10q21.1 | 1 | 25.00 | 0 | 0.00 | 0.444 | 0.691 | (A) BRAF Class 1 Lung |
| SHH | 7q36.3 | 1 | 25.00 | 0 | 0.00 | 0.444 | 0.691 | (A) BRAF Class 1 Lung |
| PRPF40B | 12q13.12 | 0 | 0.00 | 1 | 4.17 | 0.444 | 0.691 | (B) BRAF Class 3 Lung |
| SF1 | 11q13.1 | 0 | 0.00 | 1 | 4.17 | 0.444 | 0.691 | (B) BRAF Class 3 Lung |
| PIM1 | 6p21.2 | 1 | 0.61 | 2 | 1.40 | 0.449 | 0.691 | (B) BRAF Class 3 Lung |
| MYB | 6q23.3 | 0 | 0.00 | 1 | 1.06 | 0.45 | 0.691 | (B) BRAF Class 3 Lung |
| NPM1 | 5q35.1 | 0 | 0.00 | 1 | 0.55 | 0.452 | 0.691 | (B) BRAF Class 3 Lung |
| DHX15 | 4p15.2 | 0 | 0.00 | 1 | 7.14 | 0.452 | 0.691 | (B) BRAF Class 3 Lung |

| | | | | | | | | |
|---|---|---|---|---|---|---|---|---|
| NAB2 | 12q13.3 | 0 | 0.00 | 1 | 7.14 | 0.452 | 0.691 | (B) BRAF Class 3 Lung |
| SOX10 | 22q13.1 | 0 | 0.00 | 1 | 3.45 | 0.453 | 0.691 | (B) BRAF Class 3 Lung |
| NFE2L2 | 2q31.2 | 9 | 5.00 | 9 | 5.88 | 0.453 | 0.691 | (B) BRAF Class 3 Lung |
| RBBP8 | 18q11.2 | 3 | 5.66 | 1 | 2.70 | 0.454 | 0.691 | (A) BRAF Class 1 Lung |
| CCNE1 | 19q12 | 5 | 2.78 | 3 | 1.96 | 0.454 | 0.691 | (A) BRAF Class 1 Lung |
| SH2B3 | 12q24.12 | 0 | 0.00 | 1 | 0.70 | 0.455 | 0.691 | (B) BRAF Class 3 Lung |
| KLF4 | 9q31.2 | 1 | 0.74 | 2 | 1.67 | 0.456 | 0.691 | (B) BRAF Class 3 Lung |
| NKX3-1 | 8p21.2 | 0 | 0.00 | 1 | 0.92 | 0.458 | 0.691 | (B) BRAF Class 3 Lung |
| CTLA4 | 2q33.2 | 0 | 0.00 | 1 | 0.91 | 0.458 | 0.691 | (B) BRAF Class 3 Lung |
| TMEM127 | 2q11.2 | 0 | 0.00 | 1 | 0.90 | 0.459 | 0.691 | (B) BRAF Class 3 Lung |
| ACP2 | 1.2\|11p12 | 0 | 0.00 | 1 | 0.50 | 0.459 | 0.691 | (B) BRAF Class 3 Lung |
| GON4L | 1q22 | 0 | 0.00 | 1 | 0.50 | 0.459 | 0.691 | (B) BRAF Class 3 Lung |
| HRAS | 11p15.5 | 0 | 0.00 | 1 | 0.50 | 0.459 | 0.691 | (B) BRAF Class 3 Lung |
| LINC00996 | 7q36.1 | 0 | 0.00 | 1 | 0.50 | 0.459 | 0.691 | (B) BRAF Class 3 Lung |
| LINC01514 | 10q24.31 | 0 | 0.00 | 1 | 0.50 | 0.459 | 0.691 | (B) BRAF Class 3 Lung |
| PAX8-AS1 | 2q14.1 | 0 | 0.00 | 1 | 0.50 | 0.459 | 0.691 | (B) BRAF Class 3 Lung |
| PTTG1IP | 21q22.3 | 0 | 0.00 | 1 | 0.50 | 0.459 | 0.691 | (B) BRAF Class 3 Lung |
| RLF | 1p34.2 | 0 | 0.00 | 1 | 0.50 | 0.459 | 0.691 | (B) BRAF Class 3 Lung |
| TRAJ22 | 14q11.2 | 0 | 0.00 | 1 | 0.50 | 0.459 | 0.691 | (B) BRAF Class 3 Lung |
| TTF1 | 9q34.13 | 0 | 0.00 | 1 | 0.50 | 0.459 | 0.691 | (B) BRAF Class 3 Lung |
| ZFP30 | 19q13.12 | 0 | 0.00 | 1 | 0.50 | 0.459 | 0.691 | (B) BRAF Class 3 Lung |
| FANCC | 9q22.32 | 0 | 0.00 | 1 | 0.66 | 0.459 | 0.691 | (B) BRAF Class 3 Lung |
| CDK6 | 7q21.2 | 0 | 0.00 | 1 | 0.65 | 0.459 | 0.691 | (B) BRAF Class 3 Lung |
| XPO1 | 2p15 | 0 | 0.00 | 1 | 0.65 | 0.459 | 0.691 | (B) BRAF Class 3 Lung |
| RSPO2 | 8q23.1 | 0 | 0.00 | 1 | 1.75 | 0.46 | 0.691 | (B) BRAF Class 3 Lung |
| NTHL1 | 16p13.3 | 0 | 0.00 | 1 | 1.33 | 0.46 | 0.691 | (B) BRAF Class 3 Lung |
| MEF2B | 19p13.11 | 0 | 0.00 | 1 | 0.66 | 0.461 | 0.691 | (B) BRAF Class 3 Lung |
| CHD2 | 15q26.1 | 1 | 3.23 | 2 | 7.14 | 0.461 | 0.691 | (B) BRAF Class 3 Lung |
| CHD4 | 12p13.31 | 1 | 3.23 | 2 | 7.14 | 0.461 | 0.691 | (B) BRAF Class 3 Lung |
| SDHD | 11q23.1 | 0 | 0.00 | 1 | 0.66 | 0.462 | 0.691 | (B) BRAF Class 3 Lung |
| PHOX2B | 4p13 | 0 | 0.00 | 1 | 0.70 | 0.463 | 0.691 | (B) BRAF Class 3 Lung |
| XRCC2 | 7q36.1 | 0 | 0.00 | 1 | 0.83 | 0.463 | 0.691 | (B) BRAF Class 3 Lung |
| CDKN2C | 1p32.3 | 0 | 0.00 | 1 | 0.66 | 0.463 | 0.691 | (B) BRAF Class 3 Lung |
| PBRM1 | 3p21.1 | 6 | 3.57 | 4 | 2.74 | 0.465 | 0.691 | (A) BRAF Class 1 Lung |
| CD274 | 9p24.1 | 0 | 0.00 | 1 | 0.66 | 0.465 | 0.691 | (B) BRAF Class 3 Lung |
| NT5C2 | ɿ24.32-q24 | 0 | 0.00 | 1 | 1.85 | 0.466 | 0.691 | (B) BRAF Class 3 Lung |
| BCL2L1 | 20q11.21 | 0 | 0.00 | 1 | 0.66 | 0.466 | 0.691 | (B) BRAF Class 3 Lung |
| LATS2 | 13q12.11 | 1 | 1.22 | 2 | 2.67 | 0.466 | 0.691 | (B) BRAF Class 3 Lung |
| KMT2B | 19q13.12 | 2 | 3.64 | 1 | 1.67 | 0.467 | 0.691 | (A) BRAF Class 1 Lung |
| RAD51C | 17q22 | 0 | 0.00 | 1 | 0.77 | 0.469 | 0.691 | (B) BRAF Class 3 Lung |
| PPARG | 3p25.2 | 0 | 0.00 | 1 | 0.94 | 0.471 | 0.691 | (B) BRAF Class 3 Lung |
| CXCR4 | 2q22.1 | 0 | 0.00 | 1 | 0.83 | 0.473 | 0.691 | (B) BRAF Class 3 Lung |
| NUP93 | 16q13 | 0 | 0.00 | 1 | 1.20 | 0.474 | 0.691 | (B) BRAF Class 3 Lung |
| RRAS2 | 11p15.2 | 1 | 2.70 | 0 | 0.00 | 0.474 | 0.691 | (A) BRAF Class 1 Lung |
| DDX41 | 5q35.3 | 0 | 0.00 | 1 | 5.26 | 0.475 | 0.691 | (B) BRAF Class 3 Lung |
| FGF12 | 3q28-q29 | 0 | 0.00 | 1 | 10.00 | 0.476 | 0.691 | (B) BRAF Class 3 Lung |
| CTNNB1 | 3p22.1 | 12 | 5.06 | 9 | 4.48 | 0.478 | 0.691 | (A) BRAF Class 1 Lung |
| PIK3R2 | 19p13.11 | 0 | 0.00 | 1 | 1.15 | 0.478 | 0.691 | (B) BRAF Class 3 Lung |

| | | | | | | | | |
|---|---|---|---|---|---|---|---|---|
| MSI2 | 17q22 | 1 | 3.03 | 0 | 0.00 | 0.478 | 0.691 | (A) BRAF Class 1 Lung |
| EIF4E | 4q23 | 0 | 0.00 | 1 | 1.47 | 0.479 | 0.691 | (B) BRAF Class 3 Lung |
| EMSY | 11q13.5 | 0 | 0.00 | 1 | 2.86 | 0.479 | 0.691 | (B) BRAF Class 3 Lung |
| FGF14 | 13q33.1 | 0 | 0.00 | 1 | 2.86 | 0.479 | 0.691 | (B) BRAF Class 3 Lung |
| FGF23 | 12p13.32 | 0 | 0.00 | 1 | 2.86 | 0.479 | 0.691 | (B) BRAF Class 3 Lung |
| PDK1 | 2q31.1 | 0 | 0.00 | 1 | 2.86 | 0.479 | 0.691 | (B) BRAF Class 3 Lung |
| LATS1 | 6q25.1 | 1 | 1.32 | 2 | 2.78 | 0.48 | 0.691 | (B) BRAF Class 3 Lung |
| TERC | 3q26.2 | 0 | 0.00 | 1 | 2.38 | 0.483 | 0.691 | (B) BRAF Class 3 Lung |
| JUN | 1p32.1 | 0 | 0.00 | 1 | 1.14 | 0.484 | 0.691 | (B) BRAF Class 3 Lung |
| FOXO1 | 13q14.11 | 1 | 1.22 | 2 | 2.53 | 0.486 | 0.691 | (B) BRAF Class 3 Lung |
| CD276 | 15q24.1 | 0 | 0.00 | 1 | 1.39 | 0.486 | 0.691 | (B) BRAF Class 3 Lung |
| RYBP | 3p13 | 0 | 0.00 | 1 | 1.39 | 0.486 | 0.691 | (B) BRAF Class 3 Lung |
| XIAP | Xq25 | 0 | 0.00 | 1 | 1.39 | 0.486 | 0.691 | (B) BRAF Class 3 Lung |
| IRF4 | 6p25.3 | 0 | 0.00 | 1 | 1.08 | 0.487 | 0.691 | (B) BRAF Class 3 Lung |
| TNFRSF14 | 1p36.32 | 0 | 0.00 | 1 | 1.08 | 0.487 | 0.691 | (B) BRAF Class 3 Lung |
| FLI1 | 11q24.3 | 0 | 0.00 | 1 | 5.26 | 0.487 | 0.691 | (B) BRAF Class 3 Lung |
| FGF4 | 11q13.3 | 0 | 0.00 | 1 | 1.08 | 0.489 | 0.691 | (B) BRAF Class 3 Lung |
| IL7R | 5p13.2 | 6 | 4.05 | 6 | 4.84 | 0.49 | 0.691 | (B) BRAF Class 3 Lung |
| PREX2 | 8q13.2 | 3 | 4.62 | 4 | 6.25 | 0.491 | 0.691 | (B) BRAF Class 3 Lung |
| NCOR1 | 7p12-p11. | 0 | 0.00 | 1 | 1.16 | 0.491 | 0.691 | (B) BRAF Class 3 Lung |
| HSD3B1 | 1p12 | 0 | 0.00 | 1 | 3.45 | 0.492 | 0.691 | (B) BRAF Class 3 Lung |
| DCUN1D1 | 3q26.33 | 0 | 0.00 | 1 | 1.35 | 0.493 | 0.691 | (B) BRAF Class 3 Lung |
| STAT5B | 17q21.2 | 0 | 0.00 | 1 | 1.35 | 0.493 | 0.691 | (B) BRAF Class 3 Lung |
| PIK3CD | 1p36.22 | 0 | 0.00 | 1 | 1.30 | 0.494 | 0.691 | (B) BRAF Class 3 Lung |
| PIK3C2B | 1q32.1 | 4 | 3.54 | 4 | 4.55 | 0.495 | 0.691 | (B) BRAF Class 3 Lung |
| MST1R | 3p21.31 | 1 | 1.19 | 2 | 2.41 | 0.495 | 0.691 | (B) BRAF Class 3 Lung |
| BCOR | Xp11.4 | 7 | 3.93 | 5 | 3.29 | 0.497 | 0.691 | (A) BRAF Class 1 Lung |
| DNAJB1 | 19p13.12 | 0 | 0.00 | 1 | 1.28 | 0.497 | 0.691 | (B) BRAF Class 3 Lung |
| COL7A1 | 3p21.31 | 6 | 11.32 | 5 | 13.51 | 0.5 | 0.691 | (B) BRAF Class 3 Lung |
| GRM3 | 21.11-q21. | 2 | 5.88 | 1 | 2.94 | 0.5 | 0.691 | (A) BRAF Class 1 Lung |
| ACVR1B | 12q13.13 | 0 | 0.00 | 1 | 2.94 | 0.5 | 0.691 | (B) BRAF Class 3 Lung |
| ETS1 | 11q24.3 | 0 | 0.00 | 1 | 4.17 | 0.5 | 0.691 | (B) BRAF Class 3 Lung |
| GAB2 | 11q14.1 | 0 | 0.00 | 1 | 20.00 | 0.5 | 0.691 | (B) BRAF Class 3 Lung |
| PPP2R2A | 8p21.2 | 1 | 5.00 | 0 | 0.00 | 0.5 | 0.691 | (A) BRAF Class 1 Lung |
| CASR | ı13.33-q21 | 0 | 0.00 | 1 | 50.00 | 0.5 | 0.691 | (B) BRAF Class 3 Lung |
| FRK | 6q22.1 | 0 | 0.00 | 1 | 50.00 | 0.5 | 0.691 | (B) BRAF Class 3 Lung |
| KRIT1 | 7q21.2 | 0 | 0.00 | 1 | 50.00 | 0.5 | 0.691 | (B) BRAF Class 3 Lung |
| POLR2A | 17p13.1 | 0 | 0.00 | 1 | 50.00 | 0.5 | 0.691 | (B) BRAF Class 3 Lung |
| PAX3 | 2q36.1 | 1 | 4.00 | 0 | 0.00 | 0.5 | 0.691 | (A) BRAF Class 1 Lung |
| PAX7 | 1p36.13 | 0 | 0.00 | 1 | 4.00 | 0.5 | 0.691 | (B) BRAF Class 3 Lung |
| DNMT1 | 19p13.2 | 1 | 1.27 | 2 | 2.53 | 0.5 | 0.691 | (B) BRAF Class 3 Lung |
| ARID1A | 1p36.11 | 21 | 11.67 | 17 | 11.11 | 0.507 | 0.691 | (A) BRAF Class 1 Lung |
| PDCD1 | 2q37.3 | 0 | 0.00 | 1 | 1.19 | 0.509 | 0.691 | (B) BRAF Class 3 Lung |
| INSR | 19p13.2 | 2 | 2.41 | 1 | 1.23 | 0.509 | 0.691 | (A) BRAF Class 1 Lung |
| TRAF2 | 9q34.3 | 2 | 2.67 | 1 | 1.37 | 0.51 | 0.691 | (A) BRAF Class 1 Lung |
| SMAD3 | 15q22.33 | 1 | 1.11 | 0 | 0.00 | 0.511 | 0.691 | (A) BRAF Class 1 Lung |
| LYN | 8q12.1 | 1 | 1.59 | 2 | 3.08 | 0.512 | 0.691 | (B) BRAF Class 3 Lung |
| EML4 | 2p21 | 1 | 4.76 | 0 | 0.00 | 0.512 | 0.691 | (A) BRAF Class 1 Lung |

| | | | | | | | | |
|---|---|---|---|---|---|---|---|---|
| KIF5B | 10p11.22 | 0 | 0.00 | 1 | 5.00 | 0.513 | 0.691 | (B) BRAF Class 3 Lung |
| PMAIP1 | 18q21.32 | 1 | 1.32 | 0 | 0.00 | 0.514 | 0.691 | (A) BRAF Class 1 Lung |
| RAD54L | 1p34.1 | 1 | 1.15 | 0 | 0.00 | 0.515 | 0.691 | (A) BRAF Class 1 Lung |
| CYSLTR2 | 13q14.2 | 0 | 0.00 | 1 | 2.27 | 0.518 | 0.691 | (B) BRAF Class 3 Lung |
| FGF6 | 12p13.32 | 1 | 2.63 | 0 | 0.00 | 0.521 | 0.691 | (A) BRAF Class 1 Lung |
| PTCH1 | 9q22.32 | 5 | 2.78 | 5 | 3.27 | 0.521 | 0.691 | (B) BRAF Class 3 Lung |
| EIF4A2 | 3q27.3 | 1 | 1.33 | 0 | 0.00 | 0.521 | 0.691 | (A) BRAF Class 1 Lung |
| HIST1H3D | 6p22.2 | 1 | 1.35 | 0 | 0.00 | 0.521 | 0.691 | (A) BRAF Class 1 Lung |
| HIST1H3E | 6p22.2 | 1 | 1.35 | 0 | 0.00 | 0.521 | 0.691 | (A) BRAF Class 1 Lung |
| HIST1H3H | 6p22.1 | 1 | 1.35 | 0 | 0.00 | 0.521 | 0.691 | (A) BRAF Class 1 Lung |
| HIST3H3 | 1q42.13 | 1 | 1.35 | 0 | 0.00 | 0.521 | 0.691 | (A) BRAF Class 1 Lung |
| DDR1 | 6p21.33 | 0 | 0.00 | 1 | 8.33 | 0.522 | 0.691 | (B) BRAF Class 3 Lung |
| ERF | 19q13.2 | 0 | 0.00 | 1 | 2.78 | 0.522 | 0.691 | (B) BRAF Class 3 Lung |
| SESN2 | 1p35.3 | 0 | 0.00 | 1 | 2.78 | 0.522 | 0.691 | (B) BRAF Class 3 Lung |
| TAP1 | 6p21.32 | 0 | 0.00 | 1 | 2.78 | 0.522 | 0.691 | (B) BRAF Class 3 Lung |
| PLCB4 | )p12.3-p12 | 0 | 0.00 | 1 | 9.09 | 0.524 | 0.691 | (B) BRAF Class 3 Lung |
| GPS2 | 17p13.1 | 1 | 1.33 | 0 | 0.00 | 0.524 | 0.691 | (A) BRAF Class 1 Lung |
| CDC42 | 1p36.12 | 0 | 0.00 | 1 | 2.44 | 0.526 | 0.691 | (B) BRAF Class 3 Lung |
| RRAS | 19q13.33 | 0 | 0.00 | 1 | 2.44 | 0.526 | 0.691 | (B) BRAF Class 3 Lung |
| RAC1 | 7p22.1 | 1 | 0.63 | 0 | 0.00 | 0.526 | 0.691 | (A) BRAF Class 1 Lung |
| CSF3R | 1p34.3 | 1 | 0.64 | 0 | 0.00 | 0.529 | 0.691 | (A) BRAF Class 1 Lung |
| ANKRD11 | 16q24.3 | 2 | 2.70 | 1 | 1.47 | 0.532 | 0.691 | (A) BRAF Class 1 Lung |
| SETBP1 | 18q12.3 | 8 | 7.62 | 7 | 8.33 | 0.532 | 0.691 | (B) BRAF Class 3 Lung |
| HLA-A | 6p22.1 | 1 | 1.22 | 0 | 0.00 | 0.532 | 0.691 | (A) BRAF Class 1 Lung |
| PNRC1 | 6q15 | 1 | 0.89 | 0 | 0.00 | 0.533 | 0.691 | (A) BRAF Class 1 Lung |
| KDM5C | Xp11.22 | 5 | 2.98 | 5 | 3.42 | 0.535 | 0.691 | (B) BRAF Class 3 Lung |
| H3F3A | 1q42.12 | 1 | 0.56 | 0 | 0.00 | 0.538 | 0.691 | (A) BRAF Class 1 Lung |
| FANCD2 | 3p25.3 | 5 | 4.39 | 3 | 3.57 | 0.538 | 0.691 | (A) BRAF Class 1 Lung |
| SDHB | 1p36.13 | 1 | 0.56 | 0 | 0.00 | 0.538 | 0.691 | (A) BRAF Class 1 Lung |
| CDKN1B | 12p13.1 | 1 | 0.56 | 0 | 0.00 | 0.539 | 0.691 | (A) BRAF Class 1 Lung |
| ZRSR2 | Xp22.2 | 1 | 0.62 | 0 | 0.00 | 0.54 | 0.691 | (A) BRAF Class 1 Lung |
| WRN | 8p12 | 3 | 3.37 | 3 | 4.29 | 0.54 | 0.691 | (B) BRAF Class 3 Lung |
| MAP2K4 | 17p12 | 1 | 0.56 | 0 | 0.00 | 0.541 | 0.691 | (A) BRAF Class 1 Lung |
| CLIP1 | 12q24.31 | 1 | 0.42 | 0 | 0.00 | 0.541 | 0.691 | (A) BRAF Class 1 Lung |
| BMPR1A | 10q23.2 | 1 | 0.59 | 0 | 0.00 | 0.542 | 0.691 | (A) BRAF Class 1 Lung |
| PIK3C3 | 18q12.3 | 2 | 2.35 | 1 | 1.32 | 0.542 | 0.691 | (A) BRAF Class 1 Lung |
| NSD1 | 5q35.3 | 9 | 6.08 | 7 | 5.65 | 0.545 | 0.691 | (A) BRAF Class 1 Lung |
| EZH2 | 7q36.1 | 1 | 0.46 | 0 | 0.00 | 0.545 | 0.691 | (A) BRAF Class 1 Lung |
| MUTYH | 1p34.1 | 4 | 2.23 | 4 | 2.63 | 0.545 | 0.691 | (B) BRAF Class 3 Lung |
| TFRC | 3q29 | 1 | 5.56 | 0 | 0.00 | 0.545 | 0.691 | (A) BRAF Class 1 Lung |
| RAD51D | 17q12 | 2 | 1.36 | 1 | 0.77 | 0.546 | 0.691 | (A) BRAF Class 1 Lung |
| PAXIP1 | 7q36.2 | 2 | 3.77 | 2 | 5.41 | 0.546 | 0.691 | (B) BRAF Class 3 Lung |
| STAT4 | q32.2-q32. | 1 | 2.86 | 0 | 0.00 | 0.547 | 0.691 | (A) BRAF Class 1 Lung |
| MAPK1 | 22q11.22 | 2 | 1.19 | 1 | 0.68 | 0.548 | 0.691 | (A) BRAF Class 1 Lung |
| ANKRD26 | 10p12.1 | 1 | 5.88 | 0 | 0.00 | 0.548 | 0.691 | (A) BRAF Class 1 Lung |
| ARHGEF12 | 11q23.3 | 2 | 3.70 | 2 | 5.26 | 0.549 | 0.691 | (B) BRAF Class 3 Lung |
| JAZF1 | p15.2-p15. | 1 | 1.69 | 0 | 0.00 | 0.551 | 0.691 | (A) BRAF Class 1 Lung |
| SS18 | 18q11.2 | 1 | 1.69 | 0 | 0.00 | 0.551 | 0.691 | (A) BRAF Class 1 Lung |

| | | | | | | | | |
|---|---|---|---|---|---|---|---|---|
| STAG1 | 3q22.3 | 1 | 2.22 | 0 | 0.00 | 0.556 | 0.691 | (A) BRAF Class 1 Lung |
| NIPBL | 5p13.2 | 0 | 0.00 | 1 | 20.00 | 0.556 | 0.691 | (B) BRAF Class 3 Lung |
| TTYH1 | 19q13.42 | 0 | 0.00 | 1 | 20.00 | 0.556 | 0.691 | (B) BRAF Class 3 Lung |
| ZFHX4 | 8q21.13 | 0 | 0.00 | 1 | 20.00 | 0.556 | 0.691 | (B) BRAF Class 3 Lung |
| CDK4 | 12q14.1 | 2 | 1.11 | 1 | 0.65 | 0.556 | 0.691 | (A) BRAF Class 1 Lung |
| BRCA2 | 13q13.1 | 11 | 6.04 | 9 | 5.81 | 0.557 | 0.691 | (A) BRAF Class 1 Lung |
| ERCC1 | 19q13.32 | 1 | 1.32 | 0 | 0.00 | 0.559 | 0.691 | (A) BRAF Class 1 Lung |
| CDK9 | 9q34.11 | 1 | 3.57 | 0 | 0.00 | 0.56 | 0.691 | (A) BRAF Class 1 Lung |
| CRTC2 | 1q21.3 | 1 | 3.57 | 0 | 0.00 | 0.56 | 0.691 | (A) BRAF Class 1 Lung |
| RPL26 | 17p13.1 | 1 | 3.57 | 0 | 0.00 | 0.56 | 0.691 | (A) BRAF Class 1 Lung |
| ZNF708 | 19p12 | 1 | 3.57 | 0 | 0.00 | 0.56 | 0.691 | (A) BRAF Class 1 Lung |
| KMT2C | 7q36.1 | 7 | 7.69 | 6 | 7.14 | 0.561 | 0.691 | (A) BRAF Class 1 Lung |
| FAS | 10q23.31 | 1 | 0.88 | 0 | 0.00 | 0.562 | 0.691 | (A) BRAF Class 1 Lung |
| POLH | 6p21.1 | 1 | 1.85 | 0 | 0.00 | 0.562 | 0.691 | (A) BRAF Class 1 Lung |
| XRCC1 | 19q13.31 | 1 | 1.85 | 0 | 0.00 | 0.562 | 0.691 | (A) BRAF Class 1 Lung |
| IGF1R | 15q26.3 | 2 | 1.10 | 1 | 0.65 | 0.563 | 0.691 | (A) BRAF Class 1 Lung |
| SLX4 | 16p13.3 | 2 | 1.92 | 1 | 1.14 | 0.563 | 0.691 | (A) BRAF Class 1 Lung |
| MYD88 | 3p22.2 | 2 | 1.10 | 1 | 0.65 | 0.563 | 0.691 | (A) BRAF Class 1 Lung |
| MLH1 | 3p22.2 | 3 | 1.34 | 3 | 1.62 | 0.564 | 0.691 | (B) BRAF Class 3 Lung |
| XPA | 9q22.33 | 1 | 1.18 | 0 | 0.00 | 0.567 | 0.691 | (A) BRAF Class 1 Lung |
| HNF1A | 12q24.31 | 3 | 1.36 | 3 | 1.63 | 0.568 | 0.691 | (B) BRAF Class 3 Lung |
| KAT6A | 8p11.21 | 6 | 6.74 | 4 | 6.06 | 0.569 | 0.691 | (A) BRAF Class 1 Lung |
| PML | 15q24.1 | 1 | 1.75 | 0 | 0.00 | 0.57 | 0.691 | (A) BRAF Class 1 Lung |
| MYCN | 2p24.3 | 3 | 1.67 | 2 | 1.29 | 0.57 | 0.691 | (A) BRAF Class 1 Lung |
| FANCM | 14q21.2 | 2 | 3.23 | 2 | 4.35 | 0.571 | 0.691 | (B) BRAF Class 3 Lung |
| FAT3 | 11q14.3 | 1 | 8.33 | 0 | 0.00 | 0.571 | 0.691 | (A) BRAF Class 1 Lung |
| ERCC2 | 19q13.32 | 3 | 1.72 | 3 | 2.04 | 0.575 | 0.691 | (B) BRAF Class 3 Lung |
| MYCL | 1p34.2 | 3 | 1.67 | 2 | 1.31 | 0.576 | 0.691 | (A) BRAF Class 1 Lung |
| CDC73 | 1q31.2 | 3 | 1.68 | 2 | 1.32 | 0.577 | 0.691 | (A) BRAF Class 1 Lung |
| PRF1 | 10q22.1 | 1 | 1.22 | 0 | 0.00 | 0.577 | 0.691 | (A) BRAF Class 1 Lung |
| BCL6 | 3q27.3 | 3 | 1.68 | 3 | 1.97 | 0.578 | 0.691 | (B) BRAF Class 3 Lung |
| SOX2 | 3q26.33 | 3 | 1.68 | 3 | 1.97 | 0.578 | 0.691 | (B) BRAF Class 3 Lung |
| TCF7L2 | )q25.2-q25 | 3 | 1.84 | 2 | 1.45 | 0.578 | 0.691 | (A) BRAF Class 1 Lung |
| PDGFRB | 5q32 | 3 | 1.67 | 3 | 1.96 | 0.578 | 0.691 | (B) BRAF Class 3 Lung |
| IDH1 | 2q34 | 3 | 1.28 | 3 | 1.51 | 0.578 | 0.691 | (B) BRAF Class 3 Lung |
| PTK2B | 8p21.2 | 1 | 1.82 | 0 | 0.00 | 0.579 | 0.691 | (A) BRAF Class 1 Lung |
| ATF1 | 12q13.12 | 0 | 0.00 | 1 | 9.09 | 0.579 | 0.691 | (B) BRAF Class 3 Lung |
| MYH9 | 22q12.3 | 0 | 0.00 | 1 | 9.09 | 0.579 | 0.691 | (B) BRAF Class 3 Lung |
| GPC3 | Xq26.2 | 2 | 2.30 | 1 | 1.43 | 0.581 | 0.691 | (A) BRAF Class 1 Lung |
| NOTCH4 | 6p21.32 | 5 | 5.68 | 5 | 6.10 | 0.582 | 0.691 | (B) BRAF Class 3 Lung |
| MLH3 | 14q24.3 | 2 | 3.39 | 1 | 2.13 | 0.585 | 0.691 | (A) BRAF Class 1 Lung |
| GALNT12 | 9q22.33 | 1 | 1.64 | 0 | 0.00 | 0.587 | 0.691 | (A) BRAF Class 1 Lung |
| GATA4 | 8p23.1 | 2 | 1.80 | 1 | 1.14 | 0.587 | 0.691 | (A) BRAF Class 1 Lung |
| HIF1A | 14q23.2 | 0 | 0.00 | 1 | 10.00 | 0.588 | 0.691 | (B) BRAF Class 3 Lung |
| KDM5A | 12p13.33 | 5 | 3.33 | 4 | 3.08 | 0.589 | 0.691 | (A) BRAF Class 1 Lung |
| DKC1 | Xq28 | 1 | 1.89 | 0 | 0.00 | 0.589 | 0.691 | (A) BRAF Class 1 Lung |
| FAAP100 | 17q25.3 | 1 | 1.89 | 0 | 0.00 | 0.589 | 0.691 | (A) BRAF Class 1 Lung |
| NECTIN4 | 1q23.3 | 1 | 1.89 | 0 | 0.00 | 0.589 | 0.691 | (A) BRAF Class 1 Lung |

| | | | | | | | | |
|---|---|---|---|---|---|---|---|---|
| NEIL2 | 8p23.1 | 1 | 1.89 | 0 | 0.00 | 0.589 | 0.691 | (A) BRAF Class 1 Lung |
| NR0B1 | Xp21.2 | 1 | 1.89 | 0 | 0.00 | 0.589 | 0.691 | (A) BRAF Class 1 Lung |
| RHOT1 | 17q11.2 | 1 | 1.89 | 0 | 0.00 | 0.589 | 0.691 | (A) BRAF Class 1 Lung |
| TAZ | Xq28 | 1 | 1.89 | 0 | 0.00 | 0.589 | 0.691 | (A) BRAF Class 1 Lung |
| USP28 | 11q23.2 | 1 | 1.89 | 0 | 0.00 | 0.589 | 0.691 | (A) BRAF Class 1 Lung |
| XRCC4 | 5q14.2 | 1 | 1.89 | 0 | 0.00 | 0.589 | 0.691 | (A) BRAF Class 1 Lung |
| NBN | 8q21.3 | 5 | 2.82 | 4 | 2.61 | 0.59 | 0.691 | (A) BRAF Class 1 Lung |
| GLI3 | 7p14.1 | 2 | 7.14 | 1 | 4.55 | 0.591 | 0.691 | (A) BRAF Class 1 Lung |
| FH | 1q43 | 5 | 2.82 | 4 | 2.63 | 0.594 | 0.692 | (A) BRAF Class 1 Lung |
| ZNRF3 | 22q12.1 | 1 | 1.67 | 0 | 0.00 | 0.594 | 0.692 | (A) BRAF Class 1 Lung |
| NTRK2 | 9q21.33 | 5 | 2.79 | 4 | 2.61 | 0.597 | 0.694 | (A) BRAF Class 1 Lung |
| NF2 | 22q12.2 | 5 | 2.76 | 4 | 2.60 | 0.599 | 0.694 | (A) BRAF Class 1 Lung |
| ZNF521 | 18q11.2 | 0 | 0.00 | 1 | 16.67 | 0.6 | 0.694 | (B) BRAF Class 3 Lung |
| KDM6A | Xp11.3 | 5 | 2.79 | 4 | 2.63 | 0.601 | 0.694 | (A) BRAF Class 1 Lung |
| BCL11B | 14q32.2 | 2 | 3.51 | 1 | 2.33 | 0.605 | 0.698 | (A) BRAF Class 1 Lung |
| MSH6 | 2p16.3 | 6 | 3.31 | 5 | 3.25 | 0.61 | 0.702 | (A) BRAF Class 1 Lung |
| HOXB13 | 17q21.32 | 2 | 1.45 | 2 | 1.75 | 0.614 | 0.706 | (B) BRAF Class 3 Lung |
| BLM | 15q26.1 | 2 | 1.14 | 2 | 1.37 | 0.616 | 0.706 | (B) BRAF Class 3 Lung |
| NFKBIZ | 3q12.3 | 2 | 2.47 | 1 | 1.69 | 0.618 | 0.707 | (A) BRAF Class 1 Lung |
| BUB1B | 15q15.1 | 3 | 3.53 | 2 | 3.08 | 0.625 | 0.715 | (A) BRAF Class 1 Lung |
| POLQ | 3q13.33 | 4 | 7.27 | 3 | 7.50 | 0.631 | 0.717 | (B) BRAF Class 3 Lung |
| REL | 2p16.1 | 2 | 1.14 | 2 | 1.32 | 0.632 | 0.717 | (B) BRAF Class 3 Lung |
| RPTOR | 17q25.3 | 2 | 1.33 | 2 | 1.54 | 0.633 | 0.717 | (B) BRAF Class 3 Lung |
| CDH4 | 20q13.33 | 2 | 3.77 | 1 | 2.70 | 0.633 | 0.717 | (A) BRAF Class 1 Lung |
| RIF1 | 2q23.3 | 2 | 3.77 | 1 | 2.70 | 0.633 | 0.717 | (A) BRAF Class 1 Lung |
| ADGRA2 | 8p11.23 | 2 | 5.71 | 1 | 4.17 | 0.641 | 0.724 | (A) BRAF Class 1 Lung |
| COP1 | q25.1-q25. | 2 | 1.92 | 2 | 2.13 | 0.65 | 0.726 | (B) BRAF Class 3 Lung |
| ALOX12B | 17p13.1 | 2 | 1.74 | 2 | 1.92 | 0.65 | 0.726 | (B) BRAF Class 3 Lung |
| USP8 | 15q21.2 | 1 | 1.85 | 1 | 2.70 | 0.651 | 0.726 | (B) BRAF Class 3 Lung |
| BRCC3 | Xq28 | 1 | 1.89 | 1 | 2.70 | 0.656 | 0.726 | (B) BRAF Class 3 Lung |
| ENG | 9q34.11 | 1 | 1.89 | 1 | 2.70 | 0.656 | 0.726 | (B) BRAF Class 3 Lung |
| FAH | 15q25.1 | 1 | 1.89 | 1 | 2.70 | 0.656 | 0.726 | (B) BRAF Class 3 Lung |
| HELQ | 4q21.23 | 1 | 1.89 | 1 | 2.70 | 0.656 | 0.726 | (B) BRAF Class 3 Lung |
| HMBS | 11q23.3 | 1 | 1.89 | 1 | 2.70 | 0.656 | 0.726 | (B) BRAF Class 3 Lung |
| KIF1B | 1p36.22 | 1 | 1.89 | 1 | 2.70 | 0.656 | 0.726 | (B) BRAF Class 3 Lung |
| MBD4 | 3q21.3 | 1 | 1.89 | 1 | 2.70 | 0.656 | 0.726 | (B) BRAF Class 3 Lung |
| MUS81 | 11q13.1 | 1 | 1.89 | 1 | 2.70 | 0.656 | 0.726 | (B) BRAF Class 3 Lung |
| UROD | 1p34.1 | 1 | 1.89 | 1 | 2.70 | 0.656 | 0.726 | (B) BRAF Class 3 Lung |
| RASA2 | 3q23 | 0 | 0.00 | 1 | 10.00 | 0.667 | 0.733 | (B) BRAF Class 3 Lung |
| TPMT | 6p22.3 | 1 | 10.00 | 0 | 0.00 | 0.667 | 0.733 | (A) BRAF Class 1 Lung |
| KCNQ1 | .p15.5-p15 | 3 | 5.66 | 2 | 5.41 | 0.667 | 0.733 | (A) BRAF Class 1 Lung |
| MTA1 | 14q32.33 | 3 | 5.66 | 2 | 5.41 | 0.667 | 0.733 | (A) BRAF Class 1 Lung |
| MYOD1 | 11p15.1 | 2 | 2.56 | 2 | 2.70 | 0.67 | 0.735 | (B) BRAF Class 3 Lung |
| ERRFI1 | 1p36.23 | 2 | 2.20 | 2 | 2.27 | 0.677 | 0.741 | (B) BRAF Class 3 Lung |
| TAL1 | 1p33 | 1 | 1.75 | 1 | 2.33 | 0.678 | 0.741 | (B) BRAF Class 3 Lung |
| EXT2 | 11p11.2 | 1 | 1.18 | 1 | 1.54 | 0.681 | 0.741 | (B) BRAF Class 3 Lung |
| SBDS | 7q11.21 | 1 | 1.18 | 1 | 1.54 | 0.681 | 0.741 | (B) BRAF Class 3 Lung |
| NCOA3 | 20q13.12 | 2 | 2.53 | 2 | 2.56 | 0.685 | 0.744 | (B) BRAF Class 3 Lung |

| | | | | | | | | |
|---|---|---|---|---|---|---|---|---|
| FANCG | 9p13.3 | 1 | 0.83 | 1 | 1.06 | 0.687 | 0.744 | (B) BRAF Class 3 Lung |
| GATA6 | 18q11.2 | 1 | 0.90 | 1 | 1.14 | 0.69 | 0.744 | (B) BRAF Class 3 Lung |
| TRAF3 | 14q32.32 | 1 | 1.69 | 1 | 2.13 | 0.693 | 0.744 | (B) BRAF Class 3 Lung |
| HSP90AA1 | 14q32.31 | 1 | 3.33 | 1 | 4.17 | 0.696 | 0.744 | (B) BRAF Class 3 Lung |
| MPL | 1p34.2 | 1 | 0.45 | 1 | 0.54 | 0.7 | 0.744 | (B) BRAF Class 3 Lung |
| SMARCB1 | ;11.23\|22( | 1 | 0.45 | 1 | 0.54 | 0.701 | 0.744 | (B) BRAF Class 3 Lung |
| LMO1 | 11p15.4 | 1 | 0.59 | 1 | 0.71 | 0.702 | 0.744 | (B) BRAF Class 3 Lung |
| ETV1 | 7p21.2 | 1 | 0.59 | 1 | 0.71 | 0.704 | 0.744 | (B) BRAF Class 3 Lung |
| CCN6 | 6q21 | 1 | 2.86 | 1 | 3.45 | 0.705 | 0.744 | (B) BRAF Class 3 Lung |
| )RCS8-MEF: | 19p13.11 | 1 | 0.42 | 1 | 0.50 | 0.708 | 0.744 | (B) BRAF Class 3 Lung |
| MT-ND6 | - | 1 | 0.42 | 1 | 0.50 | 0.708 | 0.744 | (B) BRAF Class 3 Lung |
| SOCS1 | 16p13.13 | 1 | 0.56 | 1 | 0.66 | 0.708 | 0.744 | (B) BRAF Class 3 Lung |
| NUTM1 | 15q14 | 1 | 3.85 | 1 | 3.23 | 0.709 | 0.744 | (A) BRAF Class 1 Lung |
| RAD52 | 12p13.33 | 1 | 0.71 | 1 | 0.84 | 0.709 | 0.744 | (B) BRAF Class 3 Lung |
| GNAQ | 9q21.2 | 1 | 0.42 | 1 | 0.50 | 0.709 | 0.744 | (B) BRAF Class 3 Lung |
| YAP1 | 11q22.1 | 1 | 0.75 | 1 | 0.88 | 0.711 | 0.744 | (B) BRAF Class 3 Lung |
| FLCN | 17p11.2 | 1 | 0.56 | 1 | 0.66 | 0.711 | 0.744 | (B) BRAF Class 3 Lung |
| CBFB | 16q22.1 | 1 | 0.67 | 1 | 0.77 | 0.714 | 0.744 | (B) BRAF Class 3 Lung |
| CTCF | 16q22.1 | 1 | 0.67 | 1 | 0.77 | 0.714 | 0.744 | (B) BRAF Class 3 Lung |
| CDKN1A | 6p21.2 | 1 | 0.57 | 1 | 0.66 | 0.714 | 0.744 | (B) BRAF Class 3 Lung |
| GREM1 | 15q13.3 | 1 | 0.72 | 1 | 0.83 | 0.715 | 0.744 | (B) BRAF Class 3 Lung |
| ID3 | 1p36.12 | 1 | 0.61 | 1 | 0.70 | 0.715 | 0.744 | (B) BRAF Class 3 Lung |
| PDCD1LG2 | 9p24.1 | 1 | 0.67 | 1 | 0.77 | 0.716 | 0.744 | (B) BRAF Class 3 Lung |
| TMPRSS2 | 21q22.3 | 1 | 0.68 | 1 | 0.77 | 0.719 | 0.744 | (B) BRAF Class 3 Lung |
| RHOA | 3p21.31 | 1 | 0.74 | 1 | 0.83 | 0.721 | 0.744 | (B) BRAF Class 3 Lung |
| CALR | 19p13.13 | 1 | 0.73 | 1 | 0.82 | 0.721 | 0.744 | (B) BRAF Class 3 Lung |
| PLCG2 | 16q23.3 | 1 | 1.18 | 1 | 1.30 | 0.726 | 0.744 | (B) BRAF Class 3 Lung |
| SYK | 9q22.2 | 1 | 0.79 | 1 | 0.87 | 0.728 | 0.744 | (B) BRAF Class 3 Lung |
| SLIT2 | 4p15.31 | 1 | 3.23 | 1 | 3.57 | 0.728 | 0.744 | (B) BRAF Class 3 Lung |
| TOP2A | 17q21.2 | 1 | 3.23 | 1 | 3.57 | 0.728 | 0.744 | (B) BRAF Class 3 Lung |
| HIST1H3F | 6p22.2 | 1 | 1.35 | 1 | 1.47 | 0.73 | 0.744 | (B) BRAF Class 3 Lung |
| HIST1H3I | 6p22.1 | 1 | 1.35 | 1 | 1.47 | 0.73 | 0.744 | (B) BRAF Class 3 Lung |
| HIST1H3J | 6p22.1 | 1 | 1.35 | 1 | 1.47 | 0.73 | 0.744 | (B) BRAF Class 3 Lung |
| HIST2H3C | 1q21.2 | 1 | 1.35 | 1 | 1.47 | 0.73 | 0.744 | (B) BRAF Class 3 Lung |
| CARM1 | 19p13.2 | 1 | 3.03 | 1 | 2.78 | 0.731 | 0.744 | (A) BRAF Class 1 Lung |
| UPF1 | 19p13.11 | 1 | 3.03 | 1 | 2.78 | 0.731 | 0.744 | (A) BRAF Class 1 Lung |
| ZNF703 | 8p11.23 | 1 | 2.63 | 1 | 2.86 | 0.732 | 0.744 | (B) BRAF Class 3 Lung |
| CD70 | 19p13.3 | 0 | 0.00 | 1 | 9.09 | 0.733 | 0.744 | (B) BRAF Class 3 Lung |
| HIST2H3D | 1q21.2 | 1 | 1.37 | 1 | 1.47 | 0.734 | 0.744 | (B) BRAF Class 3 Lung |
| BBC3 | 19q13.32 | 1 | 1.32 | 1 | 1.39 | 0.738 | 0.744 | (B) BRAF Class 3 Lung |
| MDC1 | 6p21.33 | 1 | 1.32 | 1 | 1.39 | 0.738 | 0.744 | (B) BRAF Class 3 Lung |
| PIK3R3 | 1p34.1 | 1 | 1.32 | 1 | 1.39 | 0.738 | 0.744 | (B) BRAF Class 3 Lung |
| IKBKE | 1q32.1 | 1 | 1.02 | 1 | 1.08 | 0.738 | 0.744 | (B) BRAF Class 3 Lung |
| BIRC3 | 11q22.2 | 1 | 1.28 | 1 | 1.35 | 0.738 | 0.744 | (B) BRAF Class 3 Lung |
| PIK3CB | 3q22.3 | 1 | 1.09 | 1 | 1.14 | 0.74 | 0.744 | (B) BRAF Class 3 Lung |
| FGF19 | 11q13.3 | 1 | 1.02 | 1 | 1.06 | 0.741 | 0.744 | (B) BRAF Class 3 Lung |
| FGF3 | 11q13.3 | 1 | 1.02 | 1 | 1.06 | 0.741 | 0.744 | (B) BRAF Class 3 Lung |
| ASXL2 | 2p23.3 | 1 | 1.25 | 1 | 1.30 | 0.742 | 0.744 | (B) BRAF Class 3 Lung |

| | | | | | | | | |
|---|---|---|---|---|---|---|---|---|
| HIST1H1C | 6p22.2 | 1 | 1.30 | 1 | 1.30 | 0.752 | 0.753 | (B) BRAF Class 3 Lung |
| SLITRK6 | 13q31.1 | 1 | 5.26 | 1 | 5.26 | 0.757 | 0.757 | (B) BRAF Class 3 Lung |

#### Supplementary Table 18

| Gene | Cytoband | Class 1 Occurrence | Class 1 % | Class 3 Occurrence | Class 3 % | Sum Class | pval | qval | Most Enriched In |
|---|---|---|---|---|---|---|---|---|---|
| NF1 | 17q11.2 | 5 | 2.76 | 19 | 12.18 | 24 | 0.0007 | 0.0496 | (B) BRAF Class 3 Lung |
| RBM10 | Xp11.3 | 5 | 3.42 | 20 | 15.50 | 25 | 0.0004 | 0.0342 | (B) BRAF Class 3 Lung |
| PTPRD | 9p24.1-p23 | 5 | 4.72 | 22 | 21.57 | 27 | 0.0002 | 0.0211 | (B) BRAF Class 3 Lung |
| ERBB4 | 2q34 | 2 | 0.87 | 17 | 8.85 | 19 | 6E-05 | 0.0066 | (B) BRAF Class 3 Lung |
| TP53 | 17p13.1 | 74 | 31.22 | 101 | 50.75 | 175 | 3E-05 | 0.0032 | (B) BRAF Class 3 Lung |
| KRAS | 12p12.1 | 9 | 3.80 | 41 | 20.40 | 50 | 3E-08 | 5E-06 | (B) BRAF Class 3 Lung |
| SETD2 | 3p21.31 | 71 | 39.44 | 15 | 9.80 | 86 | 2E-10 | 4E-08 | (A) BRAF Class 1 Lung |
| KEAP1 | 19p13.2 | 3 | 1.79 | 40 | 27.40 | 43 | 5E-12 | 2E-09 | (B) BRAF Class 3 Lung |
| STK11 | 19p13.3 | 6 | 2.58 | 57 | 29.53 | 63 | 6E-16 | 4E-13 | (B) BRAF Class 3 Lung |

#### Supplementary Table 19

| Order Top-Bottom | Gene Name | baseMean | Log2fold | Padj |
|---|---|---|---|---|
| 1 | RPL9 | 1666.244384 | 2.317000847 | 0.000160543 |
| 2 | ULBP2 | 140.8712516 | 2.434393434 | 0.000601389 |
| 3 | TNNT1 | 296.6811702 | 4.137277 | 8.10E-07 |
| 4 | CAMK2N2 | 59.06447634 | 3.058896495 | 2.53E-05 |
| 5 | CTXN1 | 136.0597098 | 2.026784946 | 0.000301396 |
| 6 | PRDM8 | 174.9976023 | 2.416698115 | 3.73E-05 |
| 7 | TNFSF9 | 503.3596949 | 2.610316536 | 0.000246127 |
| 8 | KREMEN2 | 69.41735792 | 2.560230883 | 0.001135519 |
| 9 | FBXL16 | 367.5164531 | 2.238197522 | 1.88E-05 |
| 10 | NXPH4 | 470.8964822 | 3.924858407 | 0.000823545 |
| 11 | PCDHGB2 | 55.31821948 | 2.403870457 | 0.002552256 |
| 12 | LCK | 877.4494665 | 2.145765071 | 0.006741252 |
| 13 | AGAP2 | 215.3257302 | 2.229595312 | 0.001135519 |
| 14 | GABRP | 276.9324926 | 2.759307656 | 0.001815614 |
| 15 | RARRES1 | 1612.298008 | 2.069852147 | 0.007935528 |
| 16 | XKR9 | 123.6605397 | 2.470784901 | 0.000497559 |
| 17 | PNMA2 | 291.0881523 | 2.81568931 | 0.000605296 |
| 18 | DNAH2 | 287.8749658 | 2.364566392 | 0.000740865 |
| 19 | SERPINB5 | 3362.768977 | 2.240567517 | 0.00624963 |
| 20 | MMP25 | 132.3700973 | 2.160646858 | 0.00545536 |
| 21 | MUC16 | 89.23768704 | 7.575294146 | 5.20E-05 |
| 22 | HLA-DRB1 | 6537.501887 | 2.546361022 | 0.000235543 |
| 23 | HLA-DRB5 | 2449.000521 | 3.267712814 | 6.50E-05 |
| 24 | HLA-DQB1 | 2787.898324 | 2.718441899 | 0.000765523 |
| 25 | CIITA | 681.4537759 | 2.169068129 | 0.006022734 |
| 26 | KLRD1 | 68.87185643 | 2.1593233 | 0.004885937 |
| 27 | CXCL13 | 328.7230905 | 2.847377299 | 0.000531015 |
| 28 | HLA-DQA1 | 2141.51232 | 2.364247804 | 0.009128088 |
| 29 | NCF1 | 133.0346488 | 2.274023703 | 0.000783913 |
| 30 | GZMK | 74.69666023 | 2.499083107 | 0.006650741 |
| 31 | CTSW | 111.4441409 | 2.335442178 | 0.001641521 |
| 32 | IDO1 | 1428.629624 | 3.532389789 | 0.003306982 |
| 33 | ZNF683 | 60.22438755 | 3.044384385 | 0.000448434 |
| 34 | GZMH | 85.39310696 | 2.54265187 | 0.001802937 |
| 35 | PRF1 | 344.0719738 | 2.198614 | 0.000262613 |
| 36 | NKG7 | 213.9101048 | 2.397787893 | 0.000588099 |
| 37 | CCL5 | 947.2631221 | 2.284541569 | 0.000795865 |
| 38 | CD7 | 247.0465395 | 2.072018506 | 0.000504502 |
| 39 | PDCD1 | 70.46131539 | 2.040652636 | 0.002443752 |
| 40 | LAG3 | 210.2316171 | 2.521552637 | 0.000716886 |
| 41 | GZMA | 254.4557691 | 2.308218794 | 0.000537623 |
| 42 | GNLY | 281.2343533 | 3.5089871 | 3.62E-07 |

| | | | | |
|---|---|---|---|---|
| 43 | STMN3 | 915.5251243 | 2.323901837 | 0.000641501 |
| 44 | ROBO3 | 101.7617046 | 2.002490864 | 4.47E-05 |
| 45 | ATP1A3 | 138.7985365 | 2.957509733 | 0.00343791 |
| 46 | RPS28 | 279.4668445 | 3.820190984 | 2.45E-05 |
| 47 | CP | 180.62172 | 4.55846136 | 2.77E-05 |
| 48 | KYNU | 171.4223602 | 2.145716645 | 0.006603027 |
| 49 | BATF3 | 53.95598629 | 2.509781605 | 0.00061613 |
| 50 | SECTM1 | 1270.375664 | 2.185981349 | 0.000226823 |
| 51 | MT2A | 2264.701411 | 2.255495294 | 1.31E-06 |
| 52 | APOE | 3318.450048 | 2.249295247 | 0.004665639 |
| 53 | CCL18 | 841.5907051 | 2.704890488 | 0.004182623 |
| 54 | CCL3 | 277.1697307 | 2.451395883 | 0.000138359 |
| 55 | CCL4 | 241.0015629 | 2.383854702 | 3.07E-06 |
| 56 | CSF3R | 251.6658136 | 2.205187956 | 0.004665639 |
| 57 | CD274 | 68.39458902 | 2.253187107 | 0.004883191 |
| 58 | HAPLN3 | 689.6216771 | 2.325201394 | 9.37E-05 |
| 59 | TYMP | 2373.395181 | 2.366928612 | 0.000278962 |
| 60 | MARCO | 154.2416033 | 2.970295074 | 0.005383272 |
| 61 | TREM1 | 87.20363937 | 2.60395415 | 3.56E-05 |
| 62 | SLC11A1 | 260.7953025 | 2.209385325 | 8.15E-05 |
| 63 | GAS1 | 188.4449411 | 2.394107441 | 0.004182623 |
| 64 | COL10A1 | 591.8519452 | 2.48302786 | 0.004340292 |
| 65 | HOPX | 288.6391829 | 2.135738679 | 0.001157765 |
| 66 | ALOX5 | 697.2159983 | 2.161687386 | 1.64E-05 |
| 67 | CRIP1 | 3216.64978 | 2.350519394 | 4.74E-05 |
| 68 | GABRB3 | 69.75254653 | 4.618578602 | 0.002395863 |
| 69 | MIA | 172.5418903 | 4.723143882 | 0.000255679 |
| 70 | AQP5 | 627.5156506 | 6.914840016 | 1.34E-06 |
| 71 | PIWIL1 | 475.1123947 | 3.325683724 | 0.000387292 |
| 72 | TRPV6 | 99.5545305 | 5.086205526 | 1.11E-06 |
| 73 | VNN1 | 1093.301657 | 4.761737925 | 1.24E-08 |
| 74 | VNN2 | 151.1778909 | 4.444761586 | 3.18E-13 |
| 75 | SAA2 | 183.9166514 | 3.467657557 | 0.009023404 |
| 76 | CHST4 | 100.7762171 | 4.169942232 | 0.003884916 |
| 77 | TM4SF4 | 441.2532165 | 4.182705495 | 0.002176162 |
| 78 | PIPOX | 56.83800815 | -2.970536369 | 0.003306982 |
| 79 | CALB1 | 142.0222531 | 4.532119103 | 0.008503518 |
| 80 | AQP3 | 1462.794255 | 2.391753712 | 0.005745024 |
| 81 | GFI1 | 171.5066916 | 2.045220702 | 0.000401899 |
| 82 | PDE10A | 217.6135103 | 2.402062886 | 0.002434538 |
| 83 | DMBT1 | 9795.1678 | 3.510905688 | 0.00973388 |
| 84 | REG1A | 24504.00461 | 4.40815288 | 0.006278288 |
| 85 | ACHE | 365.7597377 | 3.548993146 | 2.35E-06 |

| | | | | |
|---|---|---|---|---|
| 86 | HSD17B2 | 637.2398258 | 2.523891582 | 0.000628696 |
| 87 | EDN3 | 143.713204 | 2.590322882 | 0.000870361 |
| 88 | PLAC8 | 1243.02475 | 2.356645674 | 0.000210276 |
| 89 | REG4 | 24343.29654 | 3.033547759 | 0.000472826 |
| 90 | B3GNT7 | 850.5476039 | 2.401467938 | 0.002110922 |
| 91 | ARHGEF4 | 135.3377332 | 3.489159449 | 3.24E-06 |
| 92 | TFF2 | 512.3835697 | 4.16534548 | 0.000995831 |
| 93 | TFF1 | 4841.887521 | 3.356928525 | 0.001147083 |
| 94 | MUC6 | 1707.034517 | 4.496372083 | 0.006735322 |
| 95 | FAM83A | 56.39858807 | 3.09677702 | 0.004818907 |
| 96 | ALDH1L1 | 466.563087 | 3.068830103 | 0.00842534 |
| 97 | NXF3 | 119.6926351 | 6.928835815 | 9.55E-08 |
| 98 | CIDEC | 57.55638055 | 2.227100706 | 0.00973388 |
| 99 | ALPI | 66.09498239 | 3.178667523 | 0.002706086 |
| 100 | NTSR1 | 257.1255111 | 4.210268872 | 0.001008956 |
| 101 | MSLN | 1936.718121 | 2.659228424 | 0.009784406 |
| 102 | HSPA6 | 404.8857839 | 2.954246656 | 0.000472826 |
| 103 | CRABP2 | 164.5726701 | 2.658728373 | 0.005575094 |
| 104 | IGFL2 | 71.56988784 | 3.310266084 | 0.001313561 |
| 105 | NLRP2 | 139.0425547 | 5.290544614 | 3.24E-06 |
| 106 | SCEL | 68.11033312 | 4.560600748 | 0.000705058 |
| 107 | AHNAK2 | 1681.107388 | 2.35257367 | 0.005837202 |
| 108 | ANO1 | 1939.251122 | 2.414140425 | 5.39E-05 |
| 109 | SLC4A11 | 274.8410881 | 2.04591285 | 0.006707152 |
| 110 | CD177 | 299.573456 | 3.728965027 | 0.002072968 |
| 111 | KLK10 | 2481.038558 | 2.840485893 | 0.002239817 |
| 112 | RHOF | 1461.31859 | 3.125083421 | 5.62E-10 |
| 113 | KRT7 | 696.7843498 | 3.849726041 | 0.00018541 |
| 114 | ANGPTL4 | 254.6982032 | 2.157940977 | 0.000221967 |
| 115 | PTPRR | 321.5783122 | 2.438262025 | 2.01E-05 |
| 116 | IL1RN | 750.6987004 | 2.060763434 | 0.000573929 |
| 117 | GJB5 | 203.60905 | 4.523432724 | 1.52E-07 |
| 118 | GJB4 | 174.3410084 | 2.233863845 | 0.004617675 |
| 119 | SPON1 | 1808.591645 | 2.173115385 | 0.009023404 |
| 120 | PI3 | 2613.366712 | 3.08395157 | 0.008414744 |
| 121 | CXCL5 | 1516.786874 | 4.154669449 | 0.000655156 |
| 122 | CCL3L1 | 78.75532037 | 2.69490044 | 0.00080629 |
| 123 | OSM | 195.2754942 | 2.824465328 | 0.000823545 |
| 124 | G0S2 | 252.0349432 | 2.013956745 | 0.001053394 |
| 125 | MMP1 | 3323.883833 | 2.525102036 | 0.004340292 |
| 126 | SERPINB2 | 69.06622818 | 5.683571467 | 1.33E-06 |
| 127 | SPRR1B | 55.0250889 | 5.188563763 | 0.005356656 |
| 128 | MFAP3L | 75.12091666 | -2.293458485 | 0.000233049 |

| | | | | |
|---|---|---|---|---|
| 129 | SEMA5A | 482.5491239 | -2.012723165 | 0.000571123 |
| 130 | VANGL2 | 132.3179558 | -2.358489377 | 0.000531015 |
| 131 | ABAT | 256.2696002 | -2.010689654 | 0.000763336 |
| 132 | SHROOM2 | 146.9227438 | -2.553988375 | 0.001600887 |
| 133 | EPHB1 | 84.02821971 | -2.756137808 | 0.003527698 |
| 134 | SLC19A3 | 136.4303834 | -4.491010135 | 2.89E-07 |
| 135 | CLIC6 | 97.5807973 | -2.210364681 | 0.00320402 |
| 136 | SLC4A8 | 60.85596199 | -2.102120318 | 0.003514747 |
| 137 | TNFRSF19 | 71.45778046 | -2.825666828 | 0.001147083 |
| 138 | NOTUM | 358.6035169 | -4.339997623 | 7.17E-05 |
| 139 | PLCB4 | 1017.473229 | -2.248506464 | 0.001608817 |
| 140 | SULT1B1 | 111.287813 | -2.530629432 | 0.007397894 |
| 141 | MME | 252.2658296 | -2.603890639 | 0.000234762 |
| 142 | SELENBP1 | 2819.59889 | -2.471830435 | 0.000401899 |
| 143 | CFTR | 1825.432778 | -2.257672554 | 0.004569572 |
| 144 | GRM8 | 91.71818182 | -4.696027472 | 1.89E-08 |
| 145 | ENPP3 | 154.5669501 | -5.385580855 | 1.03E-07 |
| 146 | C10orf99 | 538.6001834 | -4.036179592 | 0.001830386 |
| 147 | VAV3 | 513.0521852 | -2.659271744 | 0.002579106 |
| 148 | PTPRD | 193.5175017 | -2.978649042 | 0.001943803 |
| 149 | CTTNBP2 | 93.46395161 | -3.514645396 | 1.71E-05 |
| 150 | PTPRO | 442.5006881 | -4.530897521 | 1.54E-12 |
| 151 | ZNF470 | 52.09637545 | -2.006604426 | 0.008815117 |
| 152 | GSPT2 | 119.2927724 | -2.21416819 | 0.005116392 |
| 153 | ALPK3 | 243.5539963 | -3.31110622 | 2.82E-09 |
| 154 | ABCB1 | 325.1383508 | -3.088253018 | 0.000401899 |
| 155 | ZNF606 | 52.16701965 | -2.252358553 | 0.000295664 |
| 156 | APCDD1 | 897.0181026 | -4.295795827 | 5.76E-17 |
| 157 | MYEF2 | 91.58608013 | -3.506740389 | 0.000507274 |
| 158 | FHOD3 | 84.93409626 | -2.767221995 | 4.58E-06 |
| 159 | CAB39L | 214.0829324 | -2.900318951 | 3.74E-09 |
| 160 | NKD1 | 350.445154 | -4.159158119 | 1.90E-11 |
| 161 | SNCAIP | 75.93273294 | -3.840024552 | 2.51E-10 |
| 162 | TMEM150C | 87.86194877 | -3.956994709 | 4.21E-10 |
| 163 | APOLD1 | 635.3283597 | -3.464821713 | 5.63E-14 |
| 164 | KIT | 111.3994731 | -2.266251267 | 0.001453214 |
| 165 | LINC00667 | 399.8289122 | -2.388419595 | 0.000295664 |
| 166 | ASCL2 | 644.9903024 | -2.195572197 | 0.003514747 |
| 167 | ACSL6 | 69.9054484 | -3.533772134 | 0.009096023 |
| 168 | RNF43 | 2468.001111 | -2.37416574 | 2.22E-08 |
| 169 | HUNK | 303.8910921 | -3.031186962 | 0.000160543 |
| 170 | ZNRF3 | 715.8040866 | -2.059534502 | 4.44E-12 |
| 171 | AXIN2 | 1515.014469 | -3.281895308 | 5.17E-10 |

| | | | | |
|---|---|---|---|---|
| 172 | LRP4 | 1237.961971 | -2.866807612 | 0.000711729 |
| 173 | TDGF1 | 191.9388241 | -2.752464053 | 0.003458094 |
| 174 | QPRT | 449.0898142 | -2.89589446 | 0.001053394 |
| 175 | CEL | 446.2501037 | -6.702197927 | 5.63E-14 |
| 176 | MEX3A | 184.8249804 | -2.198609261 | 1.88E-05 |
| 177 | VWA2 | 359.3933097 | -2.068886156 | 0.003089136 |
| 178 | CKB | 3269.235156 | -2.010149504 | 0.003306982 |
| 179 | PRSS1 | 72.37716233 | 4.656835585 | 0.004680583 |
| 180 | NPC1L1 | 85.12532322 | 3.401412241 | 0.009023404 |
| 181 | DKK4 | 62.78370224 | -4.287802692 | 0.000178355 |
| 182 | PPBP | 59.50151876 | -4.092333515 | 0.000280288 |

#### Supplementary Table 20

| Order Top-Bottom | Gene Name | BaseMean | Log2Fold | Padj |
|---|---|---|---|---|
| 1 | BASP1 | 1991.682287 | -3.04617294 | 2.56E-05 |
| 2 | BMP6 | 216.2069608 | -2.3992458 | 0.004089324 |
| 3 | KCNE4 | 321.8142862 | -2.50543228 | 0.001364075 |
| 4 | FGL1 | 461.4412359 | -4.37488469 | 0.004868716 |
| 5 | MSMB | 267.6393493 | -3.58282777 | 0.004960652 |
| 6 | S100P | 4287.019019 | -4.02917303 | 0.000113036 |
| 7 | FXYD4 | 77.76775369 | -4.75232928 | 0.009130309 |
| 8 | PKP1 | 1087.093752 | -4.30633323 | 5.07E-05 |
| 9 | PGC | 13310.94901 | -5.36807606 | 0.001887339 |
| 10 | SLC14A2 | 60.4817057 | -5.86622041 | 7.09E-05 |
| 11 | DMRTA2 | 59.72440587 | -5.11826765 | 0.003316384 |
| 12 | H19 | 1696.997807 | -2.86525207 | 0.00269027 |
| 13 | HIF3A | 491.5341559 | -2.75048239 | 0.008689278 |
| 14 | KIF1A | 208.3848923 | -5.30872103 | 0.000568906 |
| 15 | BRDT | 223.4073313 | -6.05597208 | 4.25E-05 |
| 16 | LYPD3 | 310.0660511 | -2.33322247 | 0.003560997 |
| 17 | CLDN6 | 288.4014359 | -7.8477286 | 3.10E-08 |
| 18 | CCBP2 | 175.600186 | -3.11744806 | 0.002055062 |
| 19 | GLDC | 69.33817315 | -3.1906583 | 0.001037161 |
| 20 | FAM133A | 54.54823118 | -5.53507773 | 6.66E-06 |
| 21 | MAGEA6 | 85.72081032 | -9.04387762 | 1.19E-05 |
| 22 | MAGEA3 | 79.95012208 | -9.00014753 | 5.35E-05 |
| 23 | PRAME | 305.6517174 | -4.97247076 | 0.000198842 |
| 24 | ALG1L | 110.0759779 | -2.24994974 | 0.00477379 |
| 25 | QPRT | 765.9969822 | -3.5132685 | 8.10E-10 |
| 26 | IGF2BP3 | 619.3176094 | -5.44621921 | 6.56E-14 |
| 27 | VTCN1 | 129.9859196 | -2.52998939 | 0.004249702 |
| 28 | HIST3H2A | 63.73819124 | -2.96136592 | 0.001263655 |
| 29 | JSRP1 | 111.2631545 | -2.87682689 | 0.000788202 |
| 30 | CNTD2 | 105.0546407 | -2.59866618 | 0.005389397 |
| 31 | TM4SF19 | 53.75447508 | -3.42055157 | 0.000109901 |
| 32 | SRCRB4D | 61.26249412 | -2.01569731 | 0.008503333 |
| 33 | NRIP3 | 184.0862603 | -2.27329516 | 0.002393827 |
| 34 | BHMT2 | 374.6664489 | -3.64148025 | 0.002838472 |
| 35 | HORMAD1 | 123.837467 | -7.03689628 | 7.49E-08 |
| 36 | C7orf68 | 726.3802824 | -2.37248497 | 0.001039469 |
| 37 | CHODL | 55.50329682 | -2.82047543 | 0.004481393 |
| 38 | SRXN1 | 2862.415791 | -2.27232161 | 0.003534435 |
| 39 | AKR1C2 | 10822.97555 | -7.3552602 | 1.09E-08 |
| 40 | AKR1C1 | 17033.14031 | -5.49882997 | 5.36E-05 |
| 41 | CABYR | 277.97816 | -5.46285451 | 1.53E-11 |
| 42 | NR0B1 | 107.5019223 | -8.88182145 | 0.003960808 |
| 43 | ABCC2 | 92.60806207 | -3.57942188 | 0.00472124 |
| 44 | TSKU | 1500.089863 | -2.00059117 | 1.55E-06 |

| | | | | |
|---|---|---|---|---|
| 45 | NOTUM | 116.1487508 | -3.57060887 | 0.000120748 |
| 46 | SRPK3 | 145.051429 | -2.95111659 | 0.000127505 |
| 47 | GNG4 | 93.42462765 | -3.23908721 | 0.00202274 |
| 48 | C13orf29 | 119.8410901 | -2.48904398 | 1.74E-05 |
| 49 | SYNGR3 | 64.17341038 | -2.23739477 | 0.001352524 |
| 50 | IGFBPL1 | 62.64348196 | -2.50914303 | 0.005878152 |
| 51 | RND2 | 168.943963 | -2.9240821 | 0.000537837 |
| 52 | UCHL1 | 1320.402395 | -4.91707815 | 5.03E-09 |
| 53 | LY6K | 427.6173186 | -3.18047063 | 0.007306968 |
| 54 | MGC87042 | 62.61306691 | -2.38479596 | 0.000641494 |
| 55 | SMC1B | 58.38637887 | -4.47930963 | 0.000197948 |
| 56 | NEFH | 138.293667 | -3.93721031 | 0.000159396 |
| 57 | STAG3 | 83.6009206 | -2.39569713 | 0.00839336 |
| 58 | PEG10 | 1868.990902 | -2.91797288 | 0.003100555 |
| 59 | PI15 | 132.0175945 | -2.59559763 | 0.002487537 |
| 60 | FST | 55.60173085 | -2.81198029 | 0.001585498 |
| 61 | RGS4 | 100.9632317 | -2.02841479 | 0.006936498 |
| 62 | KCND2 | 68.9329061 | -2.27616828 | 0.005189949 |
| 63 | SLC2A5 | 180.297309 | -2.14801792 | 0.006651391 |
| 64 | MSI1 | 96.00029951 | -4.04436629 | 0.000378573 |
| 65 | ASCL1 | 131.7065432 | -6.59377408 | 2.16E-05 |
| 66 | LGI2 | 316.9219186 | -2.40310035 | 0.002019914 |
| 67 | HOXB9 | 100.4262474 | -5.07228649 | 0.001896977 |
| 68 | HOXB6 | 169.9107943 | -2.74480803 | 0.000239023 |
| 69 | LOC145837 | 149.6394489 | -4.5843476 | 5.17E-05 |
| 70 | CTNND2 | 101.1493037 | -5.52127308 | 2.06E-05 |
| 71 | NPTX1 | 486.3886195 | -4.5910722 | 3.31E-06 |
| 72 | KCNMB4 | 217.4074754 | -2.35518284 | 0.001004463 |
| 73 | RNF183 | 55.17376383 | -3.19863518 | 0.001439672 |
| 74 | PDIA2 | 98.74254281 | -4.65918096 | 0.000307759 |
| 75 | NEB | 67.22330626 | -2.97350159 | 0.000239023 |
| 76 | IGF2BP1 | 151.0534532 | -7.32889766 | 1.39E-07 |
| 77 | GRIN2D | 140.026104 | -3.40999295 | 3.73E-06 |
| 78 | CARD14 | 205.4443586 | -2.46181205 | 0.001034449 |
| 79 | YBX2 | 108.7550503 | -2.94102441 | 0.004868716 |
| 80 | GRIK5 | 121.7565898 | -3.3281945 | 0.007925001 |
| 81 | MLLT11 | 222.5404515 | -2.63747783 | 0.000341528 |
| 82 | SV2A | 210.0322176 | -2.56882099 | 0.001547112 |
| 83 | TAS1R3 | 114.8691954 | -3.45912698 | 2.40E-05 |
| 84 | PAX9 | 366.4557826 | -2.49205176 | 0.005659923 |
| 85 | SP8 | 57.40562554 | -7.50115719 | 0.000197948 |
| 86 | FKBP10 | 3367.016537 | -2.0224493 | 0.001004463 |
| 87 | HOXC6 | 56.09175771 | -2.54374461 | 0.004666252 |
| 88 | EFCAB4B | 95.90507269 | -2.78881696 | 0.000692032 |
| 89 | SYT13 | 137.5069667 | -3.76591875 | 0.00744518 |

| | | | | |
|---|---|---|---|---|
| 90 | DYNC1I1 | 52.26310596 | -2.41774877 | 0.002521515 |
| 91 | CPS1 | 1313.855835 | -6.60262713 | 2.51E-06 |
| 92 | SLC16A14 | 602.0271595 | -2.55769766 | 0.003287405 |
| 93 | STK31 | 78.97026157 | -3.18999777 | 1.21E-06 |
| 94 | LGSN | 460.4330994 | -7.1024969 | 5.46E-19 |
| 95 | APOD | 1186.902867 | -2.33047082 | 0.005457509 |
| 96 | FLNC | 378.5212364 | -2.21860654 | 0.003960808 |
| 97 | FSCN1 | 3144.791688 | -2.11595072 | 0.00342456 |
| 98 | SERPINE1 | 2148.618493 | -2.32741076 | 0.000950254 |
| 99 | POPDC3 | 52.06568637 | -5.9112724 | 3.55E-06 |
| 100 | PFN2 | 2146.940405 | -2.57108355 | 5.57E-06 |
| 101 | TMEFF1 | 70.19697517 | -2.17930753 | 0.00065547 |
| 102 | TNNT2 | 70.6690026 | 2.749695154 | 0.007366076 |
| 103 | COL17A1 | 795.3817922 | 3.231584946 | 4.82E-05 |
| 104 | SDR16C5 | 1448.800032 | 2.427868644 | 2.06E-05 |
| 105 | MFSD2A | 1028.888013 | 2.235840102 | 1.58E-07 |
| 106 | NRGN | 842.0539927 | 2.660630347 | 0.000587268 |
| 107 | SFTPA2 | 122419.0089 | 3.429175914 | 0.002475744 |
| 108 | SFTPA1 | 66286.41552 | 3.692359912 | 0.000947493 |
| 109 | LRRK2 | 3216.995799 | 3.550884517 | 6.35E-07 |
| 110 | IGFBP6 | 704.1101371 | 3.3578567 | 8.59E-05 |
| 111 | ANKRD22 | 742.6668663 | 2.1541354 | 0.00488791 |
| 112 | CDKL2 | 579.7823817 | 2.222261687 | 0.000981746 |
| 113 | SCN1A | 57.55320631 | 3.0403623 | 1.99E-05 |
| 114 | PTPN13 | 2678.363686 | 3.122816602 | 2.51E-05 |
| 115 | MS4A2 | 109.7154661 | 2.032291553 | 0.009279951 |
| 116 | HLF | 753.2660685 | 3.282167739 | 1.42E-05 |
| 117 | BAMBI | 519.5095407 | 2.441916976 | 0.001518386 |
| 118 | ANKRD24 | 60.0761099 | 3.135677556 | 1.09E-08 |
| 119 | DOC2A | 94.98393895 | 3.543154724 | 0.000649237 |
| 120 | TMEM27 | 223.9158587 | 2.658028709 | 0.003960808 |
| 121 | NOVA1 | 97.81666585 | 2.797401621 | 0.000981746 |
| 122 | MAPT | 81.32032097 | 2.381357889 | 6.66E-06 |
| 123 | HSD17B6 | 383.612422 | 2.323645288 | 0.001518386 |
| 124 | CYS1 | 134.3161906 | 2.350897326 | 0.003969204 |
| 125 | HHIP | 696.2489313 | 5.549862095 | 5.03E-09 |
| 126 | HP | 3370.755906 | 3.421157475 | 0.004996818 |
| 127 | SHISA3 | 1520.592127 | 5.985128088 | 9.17E-10 |
| 128 | ITGA10 | 80.17158777 | 2.170007577 | 0.001137218 |
| 129 | SERPIND1 | 453.0855351 | 4.5570019 | 2.20E-05 |
| 130 | PLCXD3 | 97.48848046 | 3.78215359 | 1.63E-06 |
| 131 | CYP21A2 | 50.03718651 | 2.583307684 | 0.000171447 |
| 132 | MEG3 | 141.3443947 | 3.082577835 | 0.000120748 |
| 133 | TPRG1 | 73.71630814 | 2.807295772 | 3.73E-05 |
| 134 | VSIG1 | 1911.464931 | 4.006587697 | 0.000444411 |

| | | | | |
|---|---|---|---|---|
| 135 | MMP1 | 2743.972315 | 2.933674965 | 0.008771783 |
| 136 | DEPDC7 | 115.4848135 | 2.989516149 | 0.000135809 |
| 137 | FABP4 | 91.35319531 | 2.67358205 | 0.004960652 |
| 138 | ANXA10 | 74.41296454 | 5.092755624 | 0.000422051 |
| 139 | DUOX2 | 209.1227546 | 3.041134589 | 5.20E-05 |
| 140 | FGFBP1 | 210.4287912 | 2.562019561 | 0.007759896 |
| 141 | HABP2 | 901.1698355 | 3.152667873 | 0.002838472 |
| 142 | TM4SF4 | 158.9858977 | 4.124309799 | 0.000207418 |
| 143 | MICALCL | 86.21706742 | 2.015971647 | 0.00342456 |
| 144 | SLC26A9 | 709.527527 | 2.0225684 | 0.004257242 |
| 145 | PRG4 | 168.2448738 | 3.017083685 | 0.000618528 |
| 146 | ERBB4 | 97.21431506 | 3.9781141 | 4.29E-06 |
| 147 | CD96 | 347.2624304 | 2.294900485 | 0.00080468 |
| 148 | PTGFR | 158.5877139 | 3.830272147 | 2.51E-05 |
| 149 | GRIK3 | 54.32481385 | 4.341200018 | 0.000104531 |
| 150 | ARNT2 | 1133.302883 | 3.173994526 | 6.85E-05 |
| 151 | MACROD2 | 997.5061907 | 3.113765744 | 0.004868716 |
| 152 | HSD17B2 | 173.5618837 | 4.136421072 | 0.000176119 |
| 153 | RASGRF1 | 361.5427839 | 2.617755751 | 0.004100272 |
| 154 | TP63 | 338.1328729 | 3.274306769 | 0.000568906 |
| 155 | ARHGAP20 | 207.4498347 | 2.067684235 | 0.000484089 |
| 156 | ITGB3 | 360.3350515 | 2.471155253 | 0.002055062 |
| 157 | HPGD | 6109.763406 | 3.619790977 | 1.01E-05 |
| 158 | IFIT1 | 1033.521986 | 2.122407208 | 0.000712058 |
| 159 | IFI44L | 998.4376368 | 2.527780592 | 0.000432791 |
| 160 | MX1 | 3202.825354 | 2.005611334 | 9.63E-05 |

#### Supplementary Table 21

| Order Top-Bottom | Gene Name | BaseMean | Log2Fold | Padj |
|---|---|---|---|---|
| 1 | COCH | 69.38317703 | 2.13835419 | 0.00073914 |
| 2 | PLN | 71.30797485 | 2.96027122 | 5.89E-08 |
| 3 | THBS2 | 9237.742981 | 2.13822069 | 7.82E-05 |
| 4 | PRIMA1 | 202.3381646 | 2.66918221 | 0.00019916 |
| 5 | GRIK3 | 763.5946 | 2.39695399 | 0.00340502 |
| 6 | MSI1 | 59.12081906 | 2.02494585 | 0.00202115 |
| 7 | MDFI | 345.9511031 | 2.94552187 | 2.66E-09 |
| 8 | DLX1 | 251.9935032 | 2.57304768 | 1.30E-06 |
| 9 | HOXB9 | 341.1460616 | 2.82722505 | 8.61E-06 |
| 10 | KCNH2 | 62.02106336 | 2.57486687 | 5.35E-05 |
| 11 | TFAP2B | 118.8683425 | 4.10037515 | 0.00021092 |
| 12 | RELN | 155.7643055 | 2.41557683 | 0.00138101 |
| 13 | TSPAN11 | 484.9223737 | 2.41126536 | 7.66E-06 |
| 14 | REM1 | 80.51163096 | 2.0226429 | 0.00028455 |
| 15 | IGFBP3 | 15146.54122 | 2.66971881 | 2.81E-07 |
| 16 | ITIH5 | 5656.775172 | 4.56320391 | 3.59E-11 |
| 17 | BAALC | 2073.274252 | 2.28547294 | 0.00021092 |
| 18 | ABCA6 | 396.788253 | 2.47547686 | 2.05E-05 |
| 19 | OLIG2 | 122.3952933 | 2.82748352 | 0.00026266 |
| 20 | PRSS35 | 104.1886871 | 2.78732132 | 0.00012285 |
| 21 | SEMA3E | 191.8860138 | 3.37847891 | 1.55E-06 |
| 22 | FMOD | 2540.68223 | 2.54015402 | 1.46E-06 |
| 23 | SOD3 | 941.7961052 | 2.25604195 | 2.05E-05 |
| 24 | CT45A1 | 57.42103053 | 6.57625135 | 0.00083848 |
| 25 | RYR2 | 139.1883561 | 2.2998174 | 0.00035465 |
| 26 | PRG4 | 72.6310566 | 2.39905852 | 0.00027905 |
| 27 | PLA2G2A | 221.5575043 | 3.10894852 | 2.18E-06 |
| 28 | PLIN4 | 82.55889472 | 3.29323709 | 8.20E-08 |
| 29 | RGS11 | 54.7930414 | 2.42825618 | 2.67E-05 |
| 30 | NPTX2 | 1527.172897 | 2.28104897 | 0.00866664 |
| 31 | TGFA | 255.4754795 | 2.40782369 | 5.61E-05 |
| 32 | VAT1L | 257.7262567 | 3.70058011 | 1.06E-06 |
| 33 | SYTL5 | 303.6103696 | 2.57721184 | 0.00139464 |
| 34 | CHGB | 71.56219106 | 2.26045755 | 0.00065122 |
| 35 | NGEF | 68.39898177 | 3.01661864 | 7.97E-05 |
| 36 | NTSR1 | 94.2712983 | 3.51314928 | 1.72E-06 |
| 37 | PCSK1 | 479.53272 | 2.76202382 | 7.78E-05 |
| 38 | ACPP | 117.1456126 | 2.59247139 | 5.73E-05 |
| 39 | GALNT5 | 166.8141691 | 2.36031803 | 7.52E-05 |
| 40 | SEC14L4 | 57.52928297 | 4.39251247 | 2.81E-07 |
| 41 | KLK6 | 100.8151699 | 4.36894021 | 1.65E-08 |
| 42 | CAMK2B | 50.95899656 | 2.32572001 | 4.34E-05 |
| 43 | TESC | 365.4323434 | 3.27040371 | 1.83E-09 |
| 44 | HAPLN1 | 516.8272226 | 5.55671605 | 3.59E-11 |

| | | | | |
|---|---|---|---|---|
| 45 | SERPINA3 | 12985.02344 | 2.1786553 | 0.00227004 |
| 46 | PLA2G5 | 82.53102714 | 2.93852003 | 4.70E-08 |
| 47 | PODNL1 | 238.5849253 | 3.10736101 | 1.22E-09 |
| 48 | MCHR1 | 324.9093496 | 2.17610732 | 0.00442403 |
| 49 | MT1L | 101.5207003 | 2.11507207 | 0.00065122 |
| 50 | HKDC1 | 323.9559787 | 2.59969696 | 2.17E-05 |
| 51 | FRAS1 | 379.0896873 | 2.16060881 | 5.28E-05 |
| 52 | ATP1B2 | 863.8725203 | 2.71212292 | 1.06E-06 |
| 53 | FXYD1 | 192.3758103 | 2.92174456 | 7.53E-07 |
| 54 | LGI4 | 1677.852316 | 2.60934944 | 6.78E-07 |
| 55 | TF | 2880.090475 | 3.41643305 | 7.53E-07 |
| 56 | LOC286002 | 66.69278122 | 4.38243378 | 5.71E-13 |
| 57 | SLC26A4 | 230.2837411 | 2.79019035 | 7.77E-10 |
| 58 | OLFM3 | 58.77164091 | 5.8054934 | 3.18E-07 |
| 59 | BHLHE22 | 97.97304488 | 2.73227136 | 4.16E-07 |
| 60 | RAPGEF4 | 248.5918724 | 2.28128525 | 1.06E-06 |
| 61 | CLEC1A | 144.9997418 | 2.46315529 | 1.26E-06 |
| 62 | C1QTNF3 | 2264.131905 | 3.96423079 | 1.80E-10 |
| 63 | CNTN6 | 64.16167112 | 4.23740507 | 1.06E-06 |
| 64 | F2RL2 | 336.9019413 | 2.14697653 | 0.00158603 |
| 65 | RNF175 | 180.9612808 | 2.91053928 | 1.30E-06 |
| 66 | F5 | 573.263136 | 3.34586413 | 1.04E-07 |
| 67 | FABP7 | 964.4124344 | 2.04827989 | 0.00995119 |
| 68 | C16orf89 | 77.04373692 | 2.48081849 | 2.61E-05 |
| 69 | HHATL | 1088.218969 | 7.04991668 | 1.21E-13 |
| 70 | TSIX | 106.8446863 | 2.69828285 | 0.00756967 |
| 71 | DLK1 | 51.00012897 | 7.02249105 | 7.37E-07 |
| 72 | PDE11A | 65.15412363 | 2.07170393 | 0.00026954 |
| 73 | LRRTM1 | 54.23743543 | 4.74921152 | 2.32E-07 |
| 74 | GJB6 | 120.0570362 | 3.60343648 | 5.40E-05 |
| 75 | GJB2 | 918.0515185 | 2.07712959 | 0.00131079 |
| 76 | NELL1 | 992.0113456 | 5.6465414 | 3.59E-11 |
| 77 | TFCP2L1 | 130.8043466 | 3.19079816 | 5.32E-09 |
| 78 | HPGD | 1523.989622 | 2.56031411 | 6.31E-05 |
| 79 | AGXT2L1 | 79.01094536 | 2.53266262 | 0.00089167 |
| 80 | KIF17 | 229.3583753 | 2.37035196 | 5.41E-05 |
| 81 | BAAT | 554.0960453 | 4.78239331 | 6.75E-10 |
| 82 | RLBP1 | 462.1957905 | 2.08839345 | 0.00073914 |
| 83 | BMX | 121.3188478 | 3.56285794 | 5.75E-08 |
| 84 | FAM19A5 | 398.1139322 | 2.12710274 | 0.00035465 |
| 85 | APOC2 | 1795.520985 | 2.37374233 | 0.0004612 |
| 86 | MMP1 | 593.3608557 | 2.63817339 | 0.00011826 |
| 87 | MMP3 | 59.19677439 | 2.86237505 | 0.0001758 |
| 88 | C6orf114 | 76.05669324 | -2.2249151 | 0.00022746 |
| 89 | RORC | 75.5671336 | 2.33232386 | 0.00016453 |

| | | | | |
|---|---|---|---|---|
| 90 | APCDD1 | 684.5775582 | 2.00561623 | 0.00017699 |
| 91 | TMPRSS13 | 101.5446282 | 2.99425497 | 1.55E-06 |
| 92 | CD24 | 144.8812473 | -2.2503963 | 1.43E-05 |
| 93 | RPH3A | 62.33260144 | -3.8550904 | 3.52E-10 |
